# Alzheimer’s disease-related dysregulation of protein synthesis causes key pathological features with ageing

**DOI:** 10.1101/2019.12.20.884783

**Authors:** Anshua Ghosh, Keiko Mizuno, Sachin S. Tiwari, Petroula Proitsi, Beatriz Gomez Perez-Nievas, Elizabeth Glennon, Rocio T. Martinez-Nunez, K. Peter Giese

**Author notes:** corresponding author, Maurice Wohl Clinical Neuroscience Institute, 5 Cutcombe Road, King’s College London, London, SE5 9RX, UK.

## Abstract

Alzheimer’s disease (AD) is characterised by Aβ and tau pathology as well as synaptic degeneration. Recently, it was suggested that the development of these key disease features may at least in part be due to increased protein synthesis that is regulated by fragile X mental retardation protein (FMRP) and its binding partner CYFIP2. Using an unbiased screen, we show that exposure of primary neurons to Aβ increases FMRP-regulated protein synthesis and involves a reduction of CYFIP2 levels. Modelling this CYFIP2 reduction in mice, we find Aβ accumulation, development of pre-tangle tau pathology, gliosis, loss of synapses, and deficits in memory formation with ageing. We conclude that reducing endogenous CYFIP2 expression is sufficient to cause some key features of AD in mice. We therefore propose a central role for CYFIP2 in AD as a modulator of protein synthesis, highlighting a novel direction for therapeutic targeting.

## Introduction

Amyloid-β (Aβ) plaques^1^ and phosphorylated tau tangles^2^ are classical histological features found in brains of patients suffering from Alzheimer’s disease (AD), a neurodegenerative condition that causes loss of memory and other cognitive impairments in older age^3^. The best correlate of memory loss is the progressive degeneration of synapses, which occurs prior to neuronal death^4^. Early changes in Aβ metabolism may result in synaptic degeneration via the production of soluble Aβ_1-42_ oligomers^5, 6^, a consequence of excessive amyloidogenic processing of the amyloid precursor protein (APP)^7^. In cultured neurons, Aβ_1-42_ oligomers have been suggested to locally stimulate the synthesis of APP at synapses, via an mRNA translation-dependent mechanism that involves the RNA-binding protein fragile X mental retardation protein (FMRP)^8, 9^. In this manner, Aβ_1-42_ may drive its own production, resulting in a feed-forward loop of toxicity at a particularly vulnerable location. FMRP binds to hundreds of mRNAs in the brain, including *App* mRNA^10–12^. Its primary cellular function is thought to be the inhibition of mRNA translation, which requires FMRP-mRNA interaction^10, 13, 14^, although in some instances FMRP may be involved in both nucleo-cytoplasmic and dendritic mRNA transport^15, 16^.

FMRP binds to the highly conserved cytoplasmic FMRP-interacting proteins 1 and 2 (CYFIP1 and 2, also known as Sra-1 and Pir121, respectively)^17^. Both proteins are expressed in various tissues including the brain^18^, where they localise to both excitatory^19^ and inhibitory^20^ synapses of hippocampal neurons. Although little is known about their precise function due to their relatively recent discovery, they have both overlapping and unique functions^21, 22^. CYFIP1 is thought to repress cap-dependent translation of specific mRNAs by interacting with the eukaryotic initiation factor 4E (eIF4E), and CYFIP2 has an identical eIF4E-binding motif to CYFIP1^21, 23^. In addition to regulating translation of mRNA, CYFIP1 and CYFIP2 are part of the Wiskott-Aldrich syndrome protein (WASP)-family verprolin-homologous protein (WAVE) complex that regulates actin polymerisation at synapses^24, 25^. CYFIP2 protein expression is reduced in post-mortem AD brain and in an AD mouse model^22^. Modelling the AD-related reduced CYFIP2 expression in heterozygous *Cyfip2* null mutant mice (*Cyfip2*^+/-^), which have no overt phenotypes^18^, led to increased APP protein expression, enhanced tau phosphorylation at CaMKII phospho-sites in synapses, elevated soluble Aβ_1-42_ levels, and changes in synapse morphology in the young adult hippocampus^22^.

Taken together, these findings suggested that amyloid accumulation, tau pathology, and synaptic degeneration are due to, at least in part, an Aβ-induced elevation of protein synthesis that involves CYFIP2. Here, we tested this hypothesis in primary neuronal cultures and found that nanomolar amounts of Aβ_1-42_ preparations increase net protein synthesis and modulate the association of specific putative FMRP targets with ribosomes. Mechanistically, Aβ_1-42_ causes MAPK-interacting kinase Mnk1-dependent phosphorylation of eIF4E and dissociation between eIF4E and CYFIP2, ultimately resulting in ubiquitination and reduction of CYFIP2 levels. Since early changes in Aβ metabolism reduce CYFIP2 expression, we studied the phenotype of heterozygous *Cyfip2* null mutants in older age, as ageing is the biggest risk factor for AD, and homozygous knockouts are not viable^18^. We establish that aged *Cyfip2* heterozygotes develop amyloid and tau pathology, gliosis, spine loss and more severe memory impairment than in young age, indicating that reduction of a single endogenous mouse gene can cause key aspects of AD-type pathology in the mouse brain. Therefore, our work proposes a central role for CYFIP2 in AD pathogenesis, as a potential modulator of Aβ-dependent mRNA translation.

## Results

### Aβ_1-42_ regulates differential translation of FMRP target mRNAs in neurons

Previous studies have provided circumstantial evidence that sub-micromolar doses of Aβ_1-42_ may enhance protein synthesis^26^; however, the alternative may be a difference in protein turnover. For the purpose of this study, we selected synthetic rat peptides rather than human peptides. Although it has been suggested that the rodent Aβ sequence is unable to aggregate^27, 28^, overexpression of murine Aβ in the mouse brain can result in Aβ accumulation^29^. Therefore, no *a priori* assumptions were made regarding the amyloidogenic properties of the peptide, to keep the model system as physiological as possible. To directly monitor whether Aβ_1-42_ impacts on protein synthesis, we used the SUnSET assay^30^ which utilises incorporation of puromycin, a structural analogue of aminoacyl tRNAs, into the nascent polypeptide chain, to directly reflect the rate of mRNA translation *in vitro*. Primary rat cortical neurons were cultured for 27 days *in vitro* (DIV), before being treated with a preparation of 100 nM synthetic rat Aβ_1-42_ peptides including oligomers (Supplementary Fig. 1) for 24 hours, or vehicle as control. Puromycin was added during the last 10 minutes of treatment to provide a snapshot of newly synthesised proteins and determine whether Aβ_1-42_ exposure modulates this rate. Cells were lysed and samples subjected to western blot were analysed using an anti-puromycin antibody. When normalised to signal for total protein using Revert™, we found that Aβ_1-42_ treatment led to a significant increase in protein synthesis (Fig. 1a, b; q=4.88, p<0.05). Prior treatment with translational elongation inhibitor cycloheximide (CHX) blocked the increase in puromycin signal by Aβ_1-42_, confirming it is translation dependent (Fig. 1a, b; q=0.14, p=0.99). To determine which mRNAs might be regulated at the level of translation by Aβ_1-42_ treatment, we performed paired RNA-sequencing in total cytosolic extracts and ribosomal-enriched fractions using sucrose cushions (Fig. 1c) from rat cortical neurons treated for 24 hours with Aβ_1-42_ or vehicle. A minimum of 42M reads were generated per sample (Supplementary Table 1). Correlation analysis showed that samples cluster differently depending on the subcellular fraction (Supplementary Fig. 2), suggesting that mRNAs pelleted in the sucrose cushion were distinct to total mRNAs. Differential expression between total and ribosomal-bound mRNAs showed 356 unique genes in vehicle-treated cells and 1136 unique genes in Aβ_1-42_-treated cells, with an overlap of 2624 genes (Fig. 1d, Supplementary Table 2). These data suggest that Aβ_1-42_ exposure modulates the binding of specific mRNAs to ribosomes. We interrogated the presence of previously reported FMRP targets^12^ in these differentially expressed genes and detected FMRP targets in both datasets (Fig 1e, Supplementary Fig. 3). About 11% of the differentially ribosome-bound mRNAs were FMRP targets in both vehicle and Aβ_1-42_-treated cells. However, in Aβ_1-42_-treated cells there were more unique differentially ribosome-bound FMRP targets than in control (Fig. 1e, Supplementary Table 2). This suggests that Aβ_1-42_ exposure may modulate the translation of specific FMRP-bound mRNAs. Unsupervised hierarchical clustering of the identified unique FMRP targets that showed differential binding to ribosomes upon Aβ_1-42_ exposure (not present in vehicle-treated cells) revealed clear groups of genes with differences between vehicle and Aβ_1-42_ treatment and subcellular fraction (Fig. 1f). Biological pathway analysis of these genes with Reactome^31^ were suggestive of pathways enriched in neuronal and synaptic function, such as ‘protein-protein interaction at synapses’ and ‘neurexins and neuroligins’ (Supplementary Table 3). Altogether, our data strongly suggest a role for Aβ in the ribosomal association and translation of specific genes modulated by FMRP.

**Figure 1.**
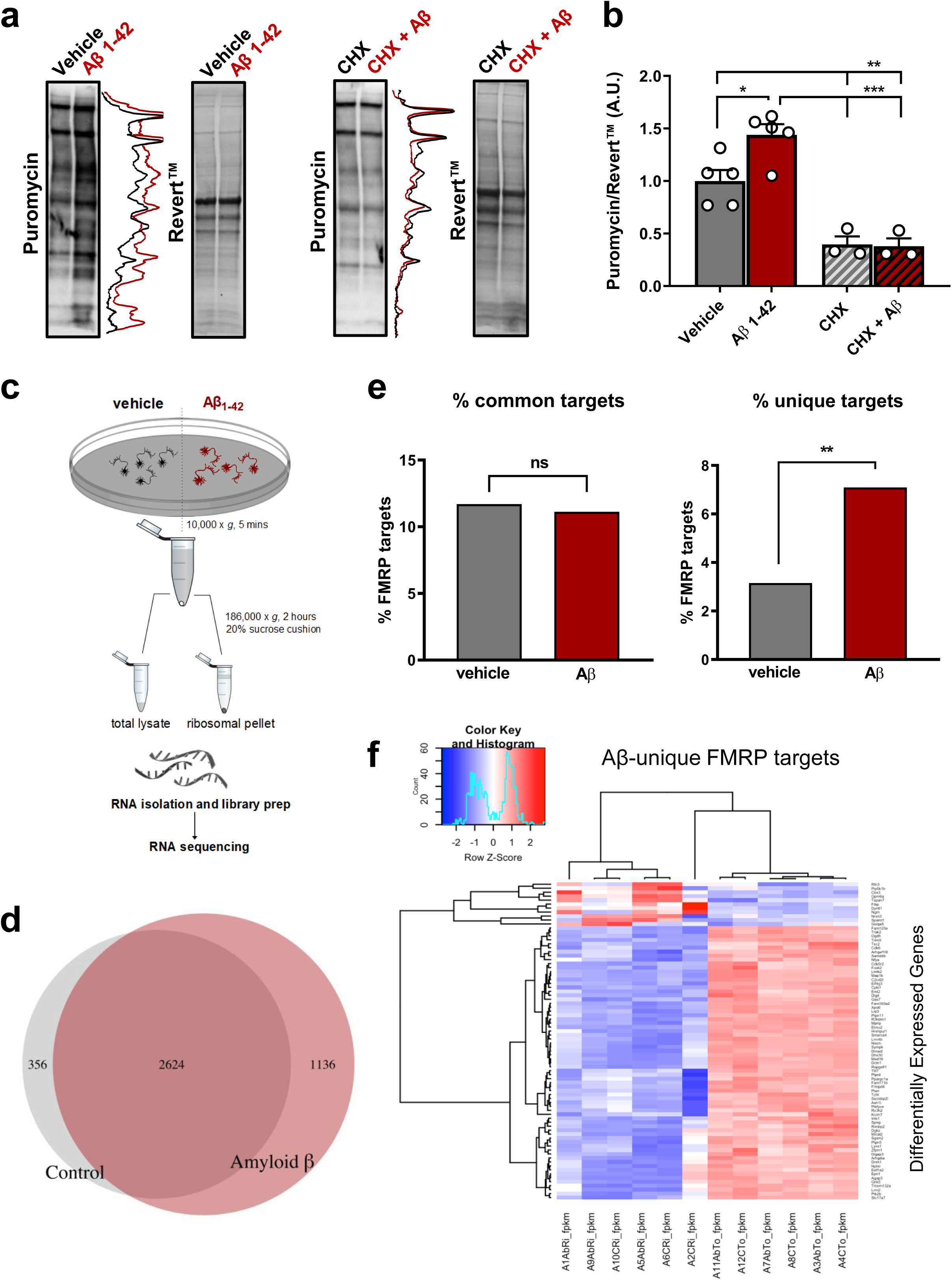
Aβ_1-42_ increases protein synthesis in primary cortical neurons by promoting ribosomal association of FMRP-bound mRNAs. **a** and **b** Protein synthesis is increased in primary neurons following 24-hour treatment with 100 nM Aβ_1-42_ preparations including oligomers, as determined by puromycin levels normalised to Revert™, a total protein stain. This effect is blocked in the presence of 10 µM cycloheximide (CHX). **c** Schematic describing the protocol used to prepare ribosome-enriched pellets from primary neurons for RNA sequencing. **d** Overlap between mRNA transcripts that are differentially bound to ribosome (versus total) within vehicle- and Aβ_1-42_-treated groups. **e** Enrichment of FMRP targets in vehicle vs Aβ cells. When considering all putative FMRP-binding targets (common and different between vehicle and Aβ-treated cells) no enrichment is observed (left). Considering unique targets that are only present either in vehicle or Aβ-treated cells (right) shows an enrichment for FMRP targets in mRNAs differentially bound to polyribosomes in the presence of Aβ. **f** Heatmap showing expression changes of FMRP targets between total and ribosomal RNAs unique to Aβ, using unsupervised hierarchical clustering. Data represent mean ± SEM. n=3-5 biological replicates (individual experiments shown as white dots). \**p* < 0.05, \*\**p* < 0.01, \*\*\**p* < 0.001.

### Aβ_1-42_ impacts on protein synthesis via the Mnk/eIF4E/CYFIP axis and reduces CYFIP2 levels

Since treatment of cortical neurons with Aβ_1-42_ preparations increased protein synthesis and appeared to modulate the translation of specific FMRP-regulated mRNAs (Fig. 1), we further explored the mechanism of this control. CYFIP1 and 2 exist as part of the FMRP translational repressor complex^17^, while interacting with the translation initiation factor eIF4E which is essential for initiation of cap-dependent mRNA translation in neurons^21^. We hypothesised that the increase in protein synthesis by Aβ_1-42_ may depend on phosphorylation of eIF4E at Ser209, as this phosphorylation is elevated in post-mortem AD brain^61^ and because it increases translation of specific transcripts^32^. To test this idea, we pharmacologically blocked eIF4E phosphorylation at Ser209 and assessed its impact on Aβ_1-42_-induced protein synthesis in primary neurons (Fig. 2a-d). We used CGP 57380, a competitive inhibitor of MAPK-interacting protein kinase 1 (Mnk1)^33^, a kinase whose only validated substrate is eIF4E^34, 35^. We found that the Aβ_1-42_ treatment significantly increased eIF4E phosphorylation at Ser209 (q=4.89, p<0.05) and the Mnk1 inhibitor blocked this increase (q=0.06, p>0.99) (Fig. 2a, b). Blocking eIF4E phosphorylation prevented an Aβ_1-42_-induced increase in protein synthesis (Fig. 2c, d; t=0.014, p=0.98), suggesting the increased translation by Aβ_1-42_ is dependent on eIF4E phosphorylation at Ser209. Blocking eIF4E phosphorylation did not alter basal protein synthesis (t=0.21, p=0.99, data not shown), in accordance with previous reports^36^.

**Figure 2.**
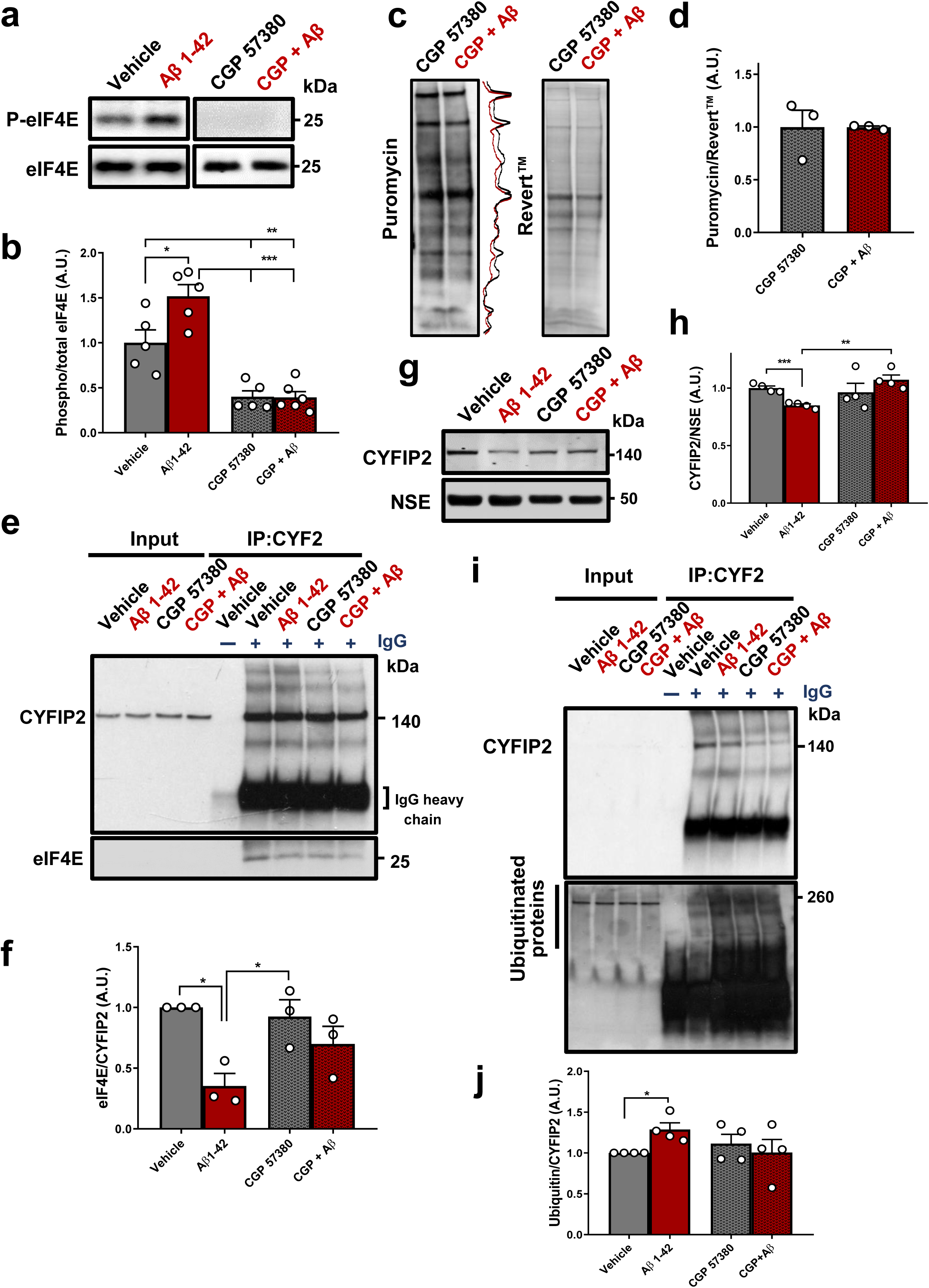
Aβ_1-42_ regulates protein synthesis in neurons via an eIF4E-CYFIP2 axis, resulting in CYFIP2 reduction. **a** and **b** An Mnk1 inhibitor (10 µM CGP 57380) reduces eIF4E phosphorylation at Ser209 and prevents Aβ_1-42_-induced increase in eIF4E phosphorylation. **c** and **d** CPG 57380 treatments prevent Aβ_1-42_-induced protein synthesis in primary neurons, as determined by SUnSET. **e** and **f** Aβ_1-42_ treatment reduces eIF4E binding to CYFIP2, as determined by co-immunoprecipitation. This effect is prevented by CGP 57380 treatment. **g** and **h** Aβ_1-42_ treatment reduces CYFIP2 levels when normalised to levels of neuronal marker NSE, which is prevented by CGP 57380 treatment. **i** and **j** Aβ_1-42_ treatment leads to ubiquitination of CYFIP2, as determined by co-IP, which is prevented by CGP 57380 treatment. Data represent mean ± SEM. n=3-6 biological replicates (individual experiments shown as white dots). \**p* < 0.05, \*\**p* < 0.01, \*\*\**p* < 0.001.

CYFIP1 represses cap-dependent translation of specific mRNAs by interacting with eIF4E and specifically blocking the eIF4E-eIF4G interaction^37^. As CYFIP2 has an identical eIF4E-binding motif to CYFIP1^21, 23^, we hypothesised it could also interact with eIF4E. We found that CYFIP2 co-immunoprecipitated with eIF4E, in rat primary cortical neurons at 28 DIV, and in the mouse hippocampus, even in synaptosome-enriched fractions (Supplementary Fig. 4). These data suggest that CYFIP2 interacts with eIF4E. Next, we tested whether Aβ_1-42_-induced phosphorylation of eIF4E would disrupt the interaction between CYFIP2 and eIF4E, using co-immunoprecipitation experiments in the presence or absence of Aβ_1-42_ (Fig. 2e, f). We found that treatment with Aβ_1-42_ alone significantly reduced the amount of eIF4E bound to CYFIP2 compared to vehicle-treated cells (Fig. 2e, f; q=5.73, p<0.05). Further, blocking Mnk1 prevented Aβ_1-42_-induced dissociation of eIF4E and CYFIP2 (Fig. 2e, f; q=2.01, p=0.52), and the inhibitor itself did not have any effect (Fig. 2e, f; q=0.66, p=0.96). We also found that Aβ_1-42_ treatment significantly reduced binding of eIF4E to CYFIP1 (Supplementary Fig. 5a, b; t=3.24, p<0.05).

We studied whether the dissociation of eIF4E and CYFIP2 could ultimately lead to reduction of CYFIP2 expression, using immunoblotting to detect CYFIP2 levels in cell lysates which were then normalised to levels of a neuronal marker, neuron specific enolase (NSE). We found that Aβ_1-42_ treatment led to significant CYFIP2 reduction when normalised to NSE (Fig. 2g, h; t=6.92, p<0.001), which was prevented by Mnk1 inhibition (Fig. 2g, h; t=1.23, p=0.27). CYFIP1 levels were not changed when normalised to NSE (Supplementary Fig. 5c, d; t=0.46, p=0.67). Additionally, the reduction of CYFIP2 expression could not be explained by synapse loss as levels of pre-synaptic marker α-synaptotagmin were not altered by Aβ_1-42_ treatment when normalised to NSE levels (Supplementary Fig. 5c, d; t=0.4, p=0.71). Levels of α-synaptotagmin were, however, reduced after five days of treatment with 100 nM Aβ_1-42_ (Supplementary Fig. 5e, f; t=3.8, p<0.05), indicating CYFIP2 expression is likely reduced prior to synapse loss.

We hypothesised that the reduction of CYFIP2 may result from its degradation by the proteasome. We therefore stripped the immunoblots for CYFIP2 and CYFIP1 co-IPs and re-probed with an antibody against ubiquitin (Ub) that recognises ubiquitinated proteins. We detected a significant increase in levels of Ub bound to CYFIP2 in Aβ_1-42_-treated cells compared to vehicle-treated cells (Fig. 2i, j; t=3.55, p<0.05), although the possibility that ubiquitination of proteins interacting with CYFIP2 was also detected cannot be excluded. Inhibition of Mnk1 prevented the effect of Aβ_1-42_ on CYFIP2 ubiquitination (Fig. 2i, j; t=0.56, p=0.60), and the inhibitor itself did not have any effect (Fig. 2i, j; t=1.03, p=0.38). In contrast, no difference was found in levels of Ub bound to CYFIP1 in Aβ_1-42_-treated cells compared to vehicle-treated cells (Supplementary Fig. 5g, h; t=0.23, p=0.84).

To test whether this pathway is genetically associated with AD, we screened variants in five genes, *MKNK1*, *MKNK2*, *CYFIP2*, *CYFIP1*, and *EIF4E*, for associations with AD (Supplementary Fig. 6). A total of 1727 variants were identified for the five genes from published summary data^38^. Following the creation of tag SNPs in order to exclude SNPs in high linkage disequilibrium (LD), there were 763 selected SNPs within 10 kb of the coding regions of each gene (Supplementary Table 4). After correcting for multiple testing there were no SNPs associated with AD. The strongest association was observed with SNP rs1258047 ∼6.3 kb downstream of *MKNK1* (OR=1.474, 95% CI: 1.21 - 1.79, p=0.0001, Q=0.075); this SNP is an *MKNK1-AS1* intronic variant and is also upstream of *KNCN*:2 kb. Mining data from the GTEx project^39^ showed that rs1258047 (or any other variants in LD (r^2^>0.8)) is not an expression quantitative trait locus (eQTL) for any gene. Taken together, this suggests that no mutations in the chosen pathway associate with AD, although more extensive studies will be more conclusive.

### CYFIP2 reduction leads to age-dependent accumulation of pre-tangle-like structures in the mouse brain

Given that CYFIP2 is downregulated in post-mortem AD brain^22^, in a mouse model of AD^22^, and in primary cortical neurons following treatment with Aβ_1-42_ oligomers (Fig. 2i, j), we speculated that its reduction may be important in the pathogenesis of AD. Previously, we reported that 3-4 month old heterozygous *Cyfip2* null mutants (*Cyfip2*^+/-^), having about 50% reduced CYFIP2 expression, have increased levels of tau phosphorylation at Ser214, possibly due to a post-transcriptional increase in αCaMKII levels^22^. Here, we also found increased tau phosphorylation at Ser416, a αCaMKII-specific phospho-site^40, 41^, when normalised to levels of total tau (Supplementary Fig. 7a, b; t=2.38, p<0.05). Further, in the young, adult *Cyfip2*^+/-^ mice no differences were found at pathology-associated sites of tau phosphorylation AT-8 (Ser202/Thr205) (Supplementary Fig. 7c, d; t=0.42, p=0.69) or PHF-1 (Ser396/Ser404) (Supplementary Fig. 7c, d; t=0.8, p=0.44).

However, we hypothesised that priming of tau phosphorylation by αCaMKII in a CYFIP2-dependent manner combined with age-related factors could result in pathological tau phosphorylation at the AT-8 and PHF-1 phosphorylation sites^42^. This idea was tested with immunohistochemistry in 12-month-old *Cyfip2*^+/-^ mice and wild-type littermates, including a group of 3-month-old wild-type mice to control for the effects of ‘healthy’ ageing. In the hippocampus, we observed an increase in neuritic layers across most sub-regions of both AT-8 and PHF-1 immunoreactivity between the 3-month-old and 12-month-old wild-type mice (Fig. 3a, b; AT-8: CA1 q=6.57, p<0.01, CA3 q=7.26, p<0.01, DG q=5.61, p<0.05; PHF-1: CA1 q=5.94, p<0.01, CA3 q=13.86, p<0.0001, DG q=2.19, p=0.32). In 12-month-old *Cyfip2*^+/-^ mice there was a further increase for AT-8 immunoreactivity, specifically in the *stratum oriens* (*so*) and *stratum radiatum* (*sr*) of area CA1 (Fig. 3a, b; q=7.44, p<0.01), and in the molecular layer (ml) of the dentate gyrus (DG) (Fig. 3a, b; q=4.4, p<0.05). No changes were observed in area CA3 (q=0.34, p=0.97, data not shown). Similar results were found for PHF-1 immunoreactivity, with a significant increase in area CA1 (Fig. 3a, b; q=4.77, p<0.05), and in DG (Fig. 3a, b; q=5.4, p<0.05), but no difference in area CA3 (q=2.14, p=0.34, data not shown).

**Figure 3.**
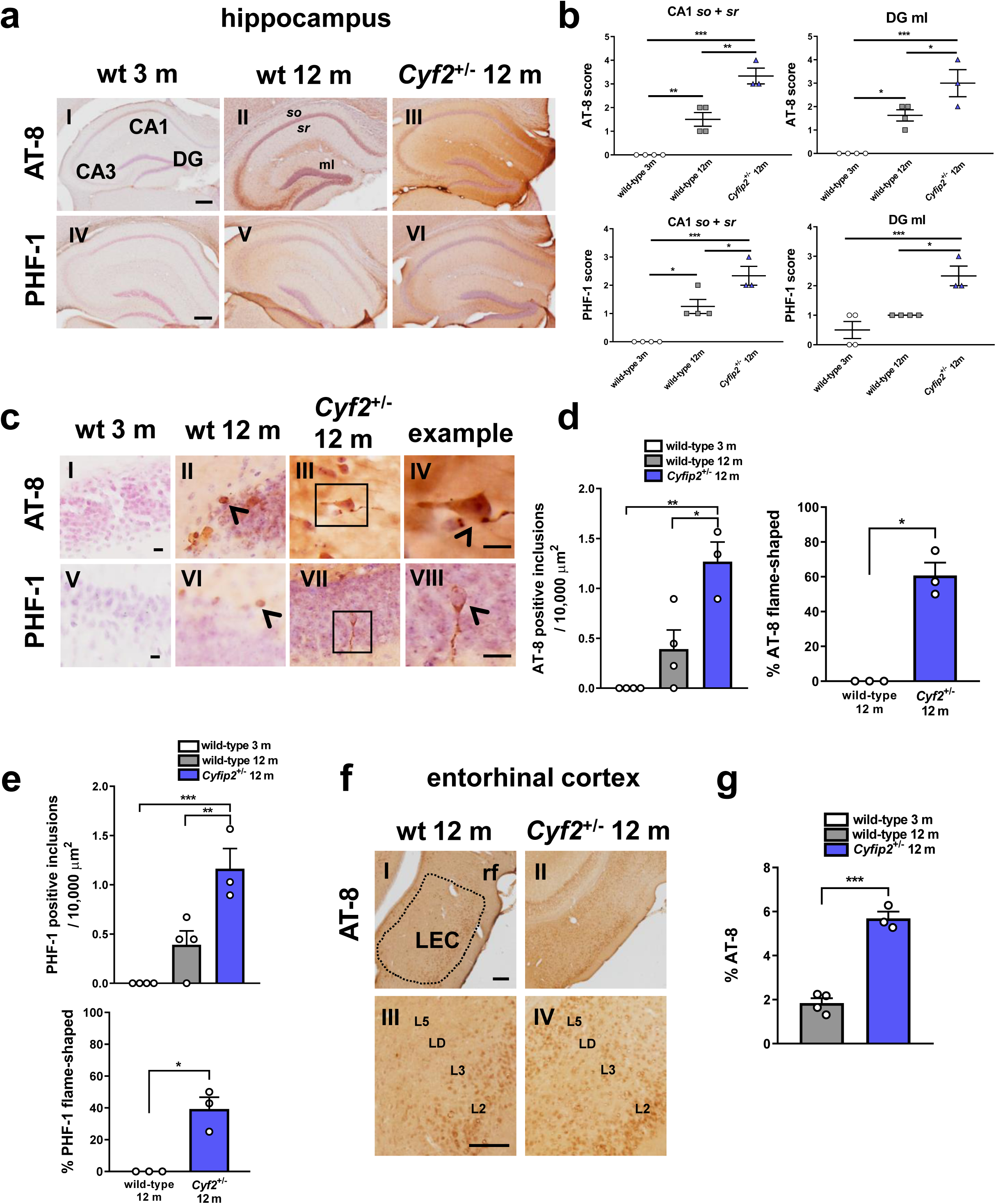
Reduced CYFIP2 expression results in phospho-tau accumulation in hippocampal formation and entorhinal cortex of 12-month-old mice. **a** Representative images of dorsal hippocampal sections from wild-type mice at 3 months and 12 months, and *Cyfip2*^+/-^ mice at 12 months, immunohistochemically stained for AT-8 (**I-III**) and PHF-1 (**IV-VI**) phospho-tau and haemotoxylin to label nuclei. Scale bars are 200 µm. **b** Analysis of DAB staining in the *so* and *sr* of area CA1 and the **ml** of the dentate gyrus showing increased phospho-tau in these brain areas in *Cyfip2*^+/-^ mice. **c** Higher magnification images of hippocampal sections labelled with AT-8 (**I-IV**) and PHF-1 (**V-VIII**). **d** and **e** Quantification of number of phospho-tau inclusions showing increase in *Cyfip2*^+/-^ mice. Scale bars are 10 µm**. f** Representative images of AT-8 staining in **LEC** of 12-month-old wild-type and *Cyfip2*^+/-^ mice (**I-II**). Higher magnification images showing laminar distribution of AT-8 in the LEC (**III-IV**). **g** Quantification of AT-8 positive staining in LEC across all layers showing increase in *Cyfip2*^+/-^ mice. Scale bars are 100 µm. Data represent mean ± SEM. n=3-4 animals per condition (individual animals shown as white dots). \**p* < 0.05, \*\**p* < 0.01, \*\*\**p* < 0.001. **L2-5** layers 2-5 LD *lamina dissecans* LEC lateral entorhinal cortex **ml** molecular layer **rf** rhinal fissure so *stratum oriens* sr *stratum radiatum*.

Higher magnification images revealed the presence of more AT8- and PHF-1-positive inclusions in 12-month-old *Cyfip2*^+/-^ mice in comparison to wild-type littermates (Fig. 3c-e; AT-8: q=5.6, p<0.05; PHF-1: q=5.7, p<0.01). In contrast, 12-month-old wild-type mice did not have significantly more inclusions than 3-month-old wild-type mice (Fig. 3c-e; AT-8: q=2.7, p=0.20; PHF-1: q=3.16, p=0.12). Importantly, among the inclusions in the *Cyfip2*^+/-^ hippocampus, several had a flame shape characteristic of pre-tangles and NFTs, which were never found in the wild-type littermates (Fig. 3c-e; AT-8: t=8.17, p<0.01; PHF-1: t=5.28, p<0.05). These are likely to be pre-tangles due to the lack of positive staining in the hippocampus with Amytracker™, which, like Thioflavin S, detects cross-β-sheet structures including NFTs^43^ (see next section; Supplementary Fig. 11a IV-VI).

AT-8 and PHF-1 immunoreactivity was increased in hippocampal area CA1 and DG, but not in area CA3, of 12-month-old *Cyfip2*^+/-^ mice (Fig. 3a, b). These two sub-regions receive direct inputs from the entorhinal cortex where tau pathology is believed to initiate^44, 45^. Therefore, we examined AT-8 and PHF-1 immunoreactivity in the lateral entorhinal cortex (LEC) of 12-month-old wild-type and *Cyfip2*^+/-^ mice. AT-8 immunoreactivity was significantly increased in LEC of 12-month-old *Cyfip2*^+/-^ mice compared to wild-type littermates (Fig. 3f, g; t=10.23, p<0.001). A qualitative comparison of the laminar distribution of AT-8 revealed that in wild-type mice, the staining was restricted to layers 2 and 3, as has been described previously in ‘healthy ageing’ studies^46^, whereas in *Cyfip2*^+/-^ mice, AT-8 immunoreactivity had progressed into deeper layers, including layer 5 (Fig. 3f, III-IV). No differences were found for PHF-1 immunohistochemistry in LEC of 12-month-old wild-type or *Cyfip2*^+/-^ mice (Supplementary Fig. 8a, b; t=0.67, p=0.55).

To test whether increased AT-8 and PHF-1 immunoreactivity in the hippocampus could be due to augmented levels of tau protein, which itself is subject to local synthesis^47, 48^, we assessed tau immunostaining in mouse brain sections. No differences were found in dendritic layers for CA1 or DG between 3-month-old and 12-month-old wild-type mice (Supplementary Fig. 9a-c; CA1 q=1.39, p=0.60; DG q=2.41, p=0.26). A significant increase was found in *so* and *sr* of area CA1 of 12-month-old *Cyfip2*^+/-^ mice compared to age-matched wild-types (Supplementary Fig. 9a, b; q=1.39, p=0.048), but no difference was found in the ml of the DG (Supplementary Fig. 9a, c; q=0.56, p=0.92). Since the change in tau expression is much smaller, or even not detected, in comparison to the observed upregulation in tau phosphorylation at the AT8 and PHF-1 sites (Fig. 3a, b), it is unlikely that the changes in tau phosphorylation are simply due to altered levels of tau protein expression. The differences observed were not due to background staining or age-related build-up of proteins (Supplementary Fig. 10).

### CYFIP2 reduction results in Aβ-positive accumulations in the mouse thalamus

Previously, we reported that 3-4-month-old *Cyfip2*^+/-^ mice had higher levels of soluble oligomeric Aβ_1-42_ in the hippocampus compared to age-matched wild-type mice^22^. We therefore hypothesised that increased levels of oligomeric Aβ_1-42_ could aggregate with age to result in Aβ_1-42_ accumulation in older mice, as it has been reported that mouse Aβ_1-42_ can aggregate, although less efficiently than human Aβ_1-42_ ^29^. We used immunohistochemical methods to search for amyloid plaques in 12-month-old wild-type and *Cyfip2*^+/-^ mice, using 3-month-old wild-type mice as control for effects of healthy ageing.

We used an anti-Aβ 4G8 antibody (aa 17-24) to detect endogenous rodent Aβ_1-42_ in mouse brain sections. In the hippocampus no positive immunohistochemical staining was observed for 4G8 across the three groups (Supplementary Fig. 11a I-III). However, in the thalamus, although no differences were observed for 4G8-positive staining between 3-month-old and 12-month-old wild-type mice (Fig. 4a I and II, b; q=0.11, p=0.99), a significant increase was found in 12-month-old *Cyfip2*^+/-^ mice when compared to age-matched controls (Fig. 4a II and III, b; q=6.07, p<0.01). This was reflected by the observation of several plaque-like inclusions localised predominantly to the somatosensory nuclei of the thalamus in these mice (Fig.4a, IV).

**Figure 4.**
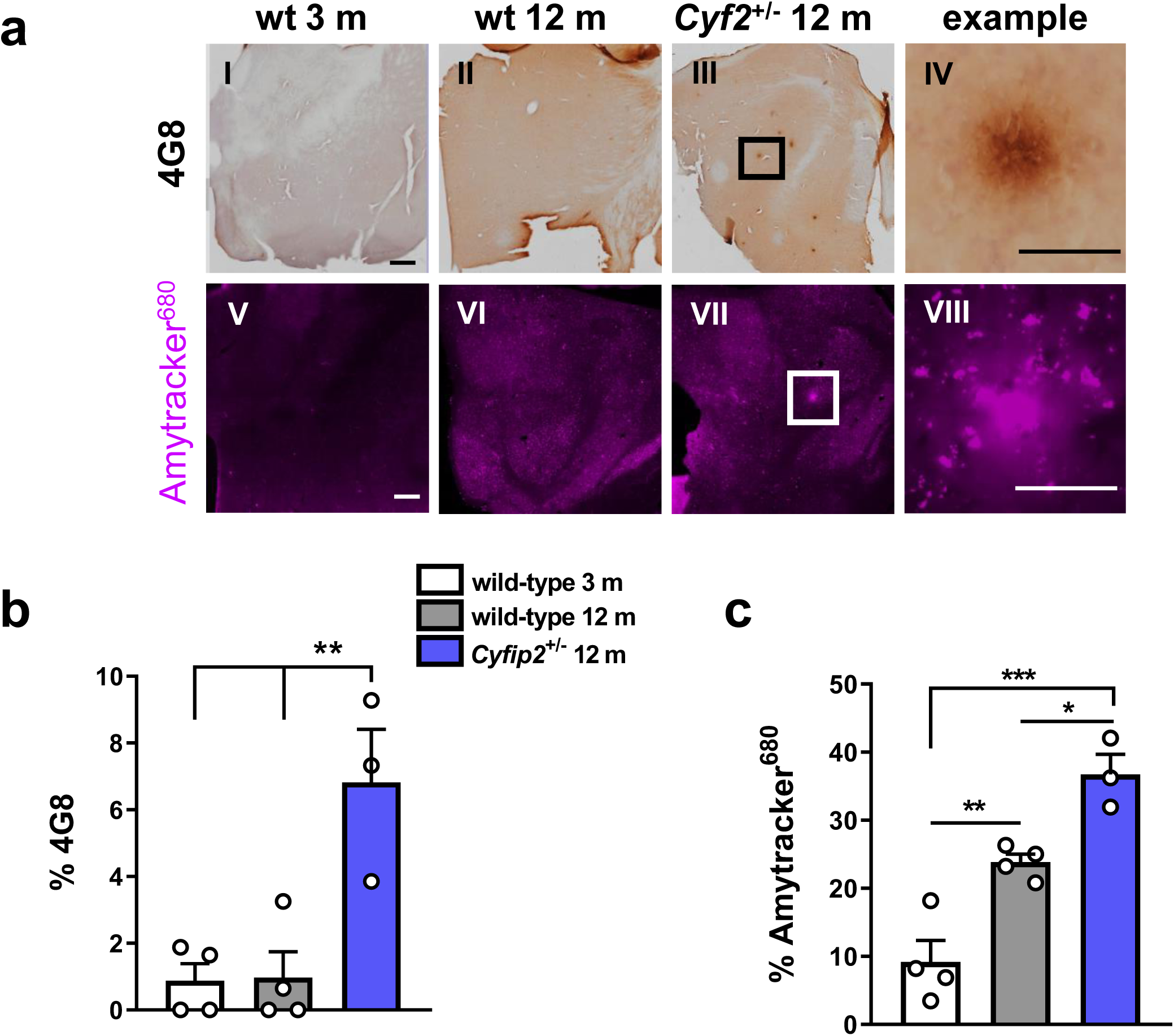
Reduced CYFIP2 expression results in Aβ accumulation in the thalamus of 12-month-old mice. **a** Representative images of the thalamus from dorsal brain sections of wild-type mice at 3 months and 12 months, and *Cyfip2*^+/-^ mice at 12 months, labelled with an anti-4G8 antibody to detect Aβ (**I-IV**) and Amytracker^680^ (**V-VIII**). Scale bars are 50 µm in ‘example’ panels and 500 µm in all other panels. **b** Quantification of 4G8 staining, expressed as % of area sampled, detects Aβ accumulation only in *Cyfip2*^+/-^ mice, but not in young adult or aged wild-type mice. **c** Quantification of Amytracker^680^ staining, expressed as % of area sampled, reveals a significant increase in *Cyfip2*^+/-^ mice. Data represent mean ± SEM. n=3-4 animals per condition (individual animals shown as white dots). \**p* < 0.05, \*\**p* < 0.01, \*\*\**p* < 0.001.

To further investigate the identity of these inclusions we also used the Amytracker™ range of fluorescent tracer molecules on brain sections. These dyes function in a manner similar to Thioflavin S, a cell-permeable benzothiazole dye that fluoresces upon binding to cross-β-sheet structures^43^, a defining feature of amyloid fibrils, and is therefore typically used for detection of amyloid plaques. Amytracker™ is designed for improved amyloid detection compared to Thioflavin compounds, detecting levels of both fibrillar and pre-fibrillar proteins and peptides. The dye revealed a trend for an increase in staining between 3-month-old and 12-month-old wild-type hippocampus (Supplementary Fig. 11a IV and V, b; q=3.6, p=0.08), but no further change was observed in 12-month-old *Cyfip2*^+/-^ mice (Supplementary Fig. 11a VI, b; q=0.05, p=0.99). However, in the thalamus, Amytracker^680^ showed significantly increased staining in 12-month-old wild-type mice compared to 3-month-old wild-types (Fig. 4a V and VI, c; q=6.04, p<0.01). This suggests that the dye most likely recognises other amyloid-like proteins besides Aβ, which are increased with ageing^49^. A further significant increase was found in 12-month-old *Cyfip2*^+/-^ mice (Fig. 4a VII, c; q=4.92, p<0.05). The dye also detected thalamic inclusions that resembled amyloid plaques, which were not present in age-matched wild-type mice (Fig.4a, VIII). These data strongly suggest that Aβ_1-42_ can accumulate in an age-dependent manner upon CYFIP2 reduction, although further characterisation of these inclusions is required.

### CYFIP2 reduction leads to astrocytic and microglial responses in the mouse hippocampus

Although plaques and tangles are two of the characteristic hallmarks of AD, there are several other pathological features such as gliosis, which is thought to be reactive to amyloid and tau pathology. To test whether 12-month-old *Cyfip2*^+/-^ mice have an astroglial response, mouse brain sections were analysed for glial fibrillary acidic protein (GFAP) expression, using immunofluorescence. Increased numbers of GFAP-positive astrocytes were found across all hippocampal sub-regions except the hilus in 12-month-old wild-type mice compared to 3-month-old wild-types (Fig. 5a, b; CA1 q=5.42, p<0.05, CA3 q=7.25, p<0.01, DG q=14.49, p<0.0001, hilus q=0.67, p=0.88). A further significant increase was observed in 12-month-old *Cyfip2*^+/-^ mice compared to age-matched wild-types, specifically in area CA1 and DG, but not area CA3 or hilus (Fig. 5a, b; CA1 q=5.59, p<0.05, CA3 q=0.16, p=0.99, DG q=5.54, p<0.05, hilus q=1.56, p=0.54).

**Figure 5.**
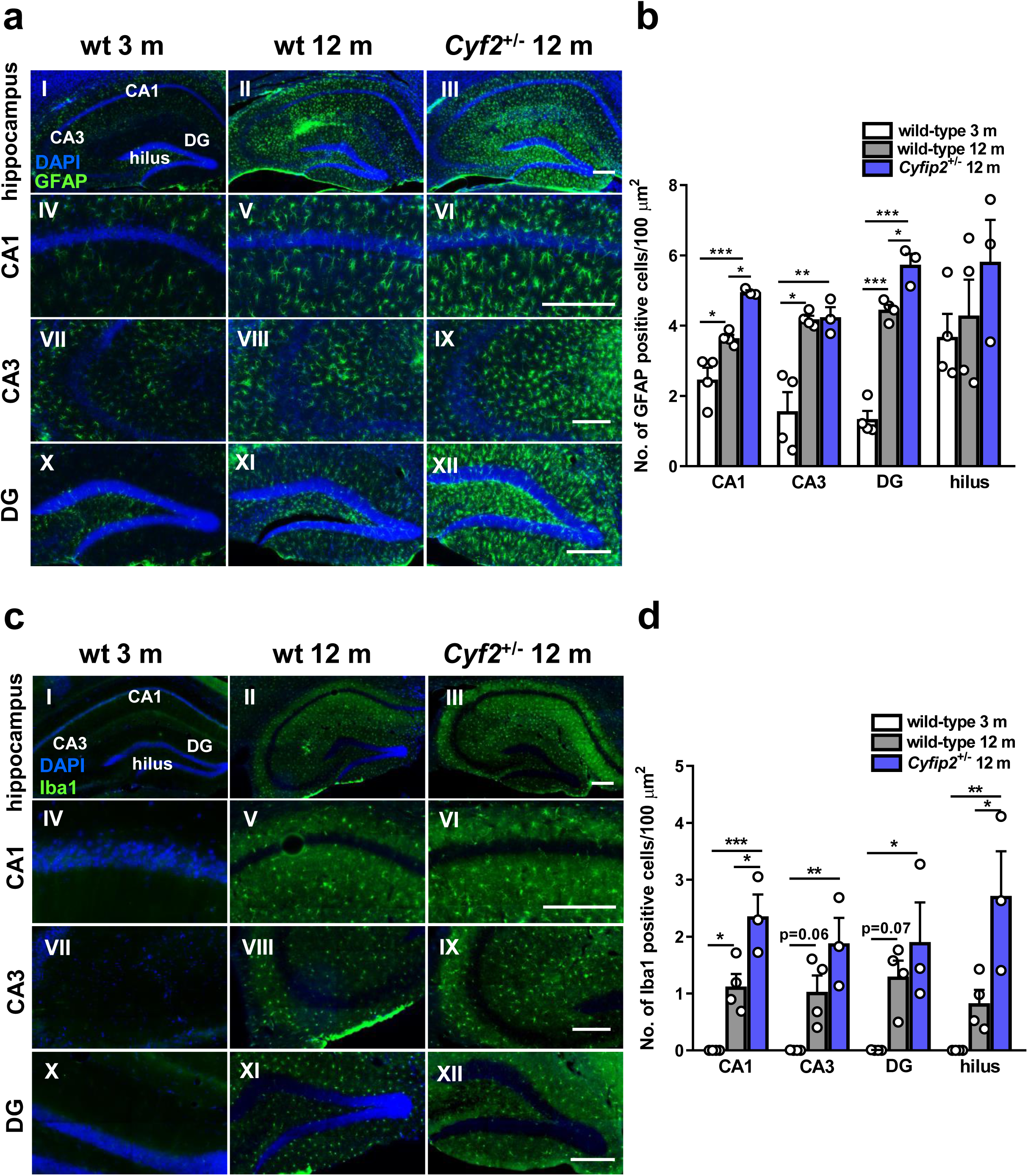
Reduced CYFIP2 expression results in astrocytic and microglial responses in specific hippocampal sub-regions of 12-month-old mice. **a** Representative images of dorsal hippocampal sections (**I-III**) from wild-type mice at 3 months and 12 months, and *Cyfip2*^+/-^ mice at 12 months, stained for GFAP. Higher magnification images of CA1 (**IV-VI**), CA3 (**VII-IX**) and DG (**X-XII**). Scale bars are 200 µm. **b** Quantification of GFAP-positive astrocytes in hippocampal sub-regions shown indicates an elevated astrocytic response in area CA1 and DG of *Cyfip2*^+/-^ mice in comparison with wild-type littermates. **c** Representative images of dorsal hippocampal sections (**I-III**) from wild-type mice at 3 months and 12 months, and *Cyfip2*^+/-^ mice at 12 months, stained for Iba1. Higher magnification images of CA1 (**IV-VI**), CA3 (**VII-IX**) and DG (**X-XII**). Scale bars are 200 µm. **b** Quantification of Iba1-positive microglia in hippocampal sub-regions shown reveals an increased microglial response in area CA1 and hilus of *Cyfip2*^+/-^ mice in comparison with wild-type littermates. Data represent mean ± SEM. n=3-4 animals per condition (individual animals shown as white dots). \**p* < 0.05, \*\**p* < 0.01, \*\*\**p* < 0.001.

To test for the presence of a microglial response in 12-month-old *Cyfip2*^+/-^ mice, we used an antibody against ionised calcium-binding adapter molecule 1 (Iba1), a microglia/macrophage-specific calcium-binding protein, to ubiquitously detect microglia in mouse brain sections by immunofluorescence. We found a significant increase in the number of Iba1-positive microglia in the CA1 region of 12-month-old wild-type mice compared to 3-month-old wild-type mice, and a trend for an increase in area CA3 and DG (Fig. 5c, d; CA1 q=5.22, p<0.05, CA3 q=3.9, p=0.06, DG q=3.74, p=0.07, hilus q=2.25, p=0.30). In 12-month-old *Cyfip2*^+/-^ mice we found a further increase in area CA1 and hilus compared to age-matched controls (Fig. 5c, d; CA1 q=5.31, p<0.05, CA3 q=3.01, p=0.15, DG q=1.62, p=0.51, hilus q=4.74, p<0.05). Taken together, our data show that *Cyfip2*^+/-^ mice display hippocampal gliosis.

Given that Aβ accumulations were found in the thalamus of aged *Cyfip2*^+/-^ mice and glial responses have been reported near the vicinity of plaques^50, 51^, astrocytes and microglia were also quantified in the thalamus of mouse brain sections. Increased numbers of GFAP-positive astrocytes (Supplementary Fig. 12a I-III, b; q=6.07, p<0.01) and Iba1-positive microglia (Supplementary Fig. 12a IV-VI, c; q=6.54, p<0.01) were found in 12-month-old wild-type mice compared to 3-month-old wild-type mice. No further changes in astrocyte and microglial numbers (Supplementary Fig. 12a-c; GFAP q=1.01, p=0.76; Iba1 q=0.31, p=0.97) were observed in the thalamus of 12-month-old *Cyfip2*^+/-^ mice. These data may suggest that the glial response in *Cyfip2*^+/-^ mice is reactive to phospho-tau or soluble Aβ_1-42_^22^, rather than Aβ accumulation.

### CYFIP2 reduction leads to dendritic spine loss in hippocampal CA1 neurons and impacts on contextual fear memory formation

Previously, we reported a change in dendritic spine morphology on apical dendrites of hippocampal CA1 neurons in 3-4-month-old *Cyfip2*^+/-^ mice compared to age-matched wild-type mice^22^. Specifically, young, adult *Cyfip2*^+/-^ mice have more immature long/thin-type spines and fewer mature stubby/mushroom-type spines, but the overall spine number is not changed^22^. Since with ageing CYFIP2 reduction causes tau pathology and gliosis in the hippocampus (Figs. 3 and 5), we tested whether this may result in spine loss in area CA1. We used Golgi-Cox impregnation to label neurons in brains from 12-month-old wild-type and *Cyfip2*^+/-^ mice, and quantified dendritic spines in the *stratum radiatum* of hippocampal CA1. We found that 12-month-old *Cyfip2*^+/-^ mice had an overall reduction in the number of dendritic spines compared to age-matched wild-types (Fig. 6a, b; t=2.84, p<0.05). Upon stratification, there was a trend for reduction in the number of stubby spines (Fig. 6c; t=3.04, p=0.07) and a significant reduction in the number of mushroom spines (Fig. 6c; t=3.76, p<0.05), but no difference in thin spines (Fig. 6c; t=0.09, p=0.93) or filopodia (Fig. 6c; t=0.98, p=0.42).

**Figure 6.**
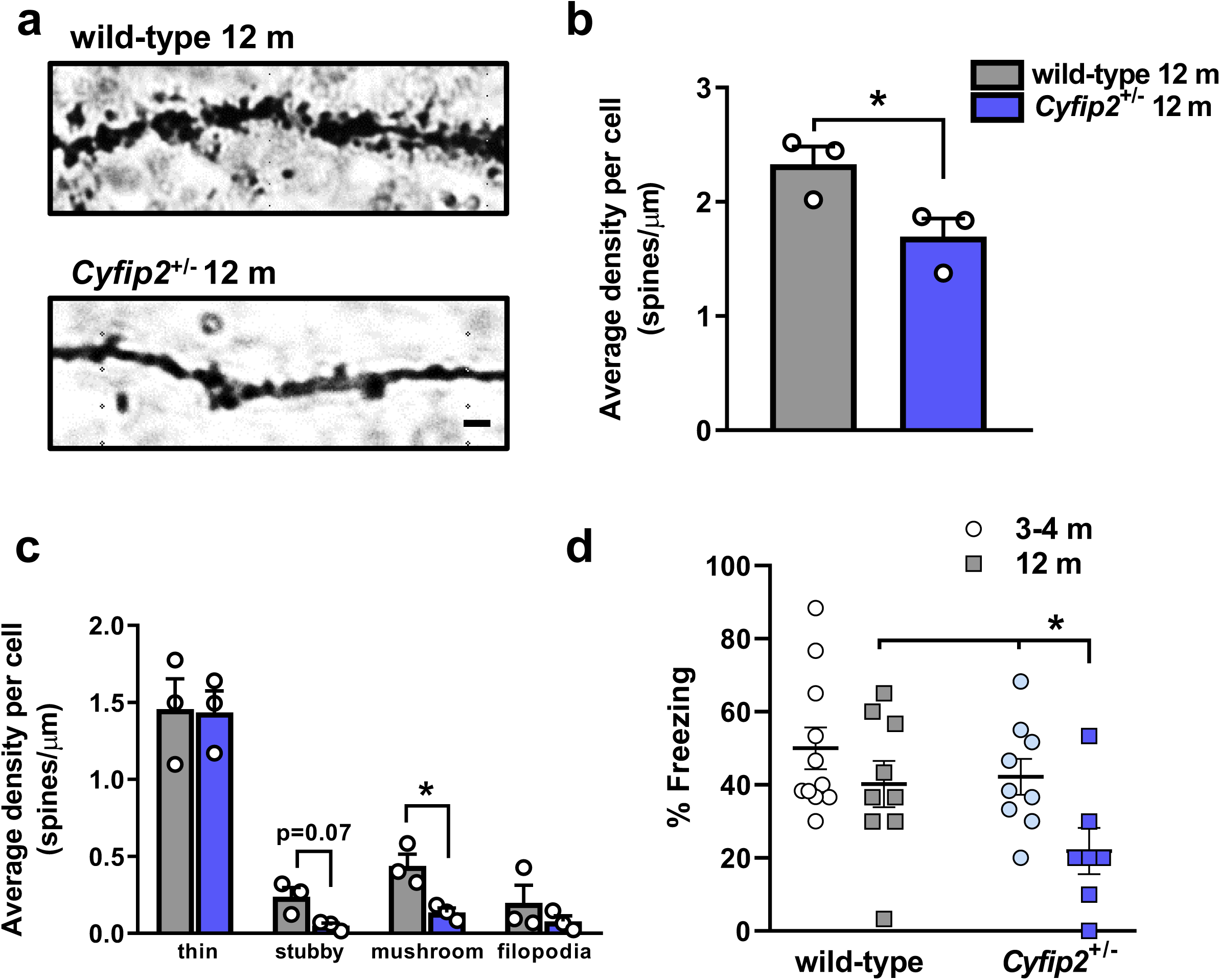
Reduced CYFIP2 expression causes loss of dendritic spines and age-dependent impairment in contextual memory formation. **a** Representative images of apical dendrites from hippocampal CA1 neurons impregnated with Golgi-Cox stain, from 12-month-old wild-type and *Cyfip2*^+/-^ mice. Scale bar is 1 µm. **b** Quantification of total number of dendritic spines per µm length of dendrite reveals spine loss in the *Cyfip2*^+/-^ mice. **c** Quantification of numbers of dendritic spines stratified according to morphological parameters, indicating loss of mushroom and stubby spines in *Cyfip2*^+/-^ mice. **d** Quantification of percentage freezing in 3-4-month-old and 12-month-old wild-type and *Cyfip2*^+/-^ mice in the contextual fear-conditioning paradigm. Aged *Cyfip2*^+/-^ mice have significantly impaired contextual fear memory. Data represent mean ± SEM. n=3 animals per condition for spine analysis, n=7-11 animals per condition for behaviour (individual animals shown as dots or squares). \**p* < 0.05.

The age-related spine loss in *Cyfip2*^+/-^ mice is expected to exacerbate learning and memory deficits. We tested this in hippocampus-dependent contextual fear conditioning, using *Cyfip2*^+/-^ mice and wild-type littermates at 3-4 months or 12 months of age. We found significant differences in genotype and age for freezing (Fig. 6d; effect of genotype F_1,32_=4.84, p<0.05; effect of age F_1,32_=6.47, p<0.05; genotype and age interaction F_1,32_=0.78, p=0.38). Post-hoc analysis showed that 12-month-old *Cyfip2*^+/-^ mice froze significantly less than 3-4-month-old *Cyfip2*^+/-^ mice (q=3.25, p<0.05) and 12-month-old wild-type mice (q=2.92, p<0.05). These results strongly suggest that ageing exacerbates memory impairment in *Cyfip2*^+/-^ mice.

## Discussion

Loss of synapses in AD is the best correlate of memory impairment and may result from build-up of amyloid-beta (Aβ_1-42_) peptides during disease progression. However, the precise roles of Aβ_1-42_ in synapse modification are poorly understood. Here, we show that in neurons Aβ_1-42_ regulates mRNA translation and modulates ribosomal association of several FMRP-bound mRNAs, primarily affecting synthesis of proteins related to neuronal and synaptic function. We demonstrate that this is likely to be via a pathway involving Mnk1 and eIF4E, which results in reduction of the 4E-binding protein CYFIP2, most likely due to ubiquitination. Finally, we find that reduced expression of CYFIP2 in 12-month-old mice causes several phenotypes of the disease. These include development of phospho-tau accumulation in the hippocampus and entorhinal cortex, thalamic Aβ accumulation, hippocampal gliosis, dendritic spine loss, and impaired hippocampus-dependent memory formation.

We propose a model where CYFIP2 reduction functions as both a cause and consequence of AD pathogenesis (Fig. 7). Reduction of CYFIP2 levels may result from activation of the Mnk1-eIF4E-CYFIP axis by Aβ_1-42_ (Fig. 2). Once lost, a state of prolonged protein synthesis is proposed to further contribute to the toxic loop of Aβ_1-42_ production and tau hyperphosphorylation via increased synthesis of APP and αCaMKII protein, respectively^22^. Ageing processes may then interact to drive accumulation of Aβ_1-42_ and hyperphosphorylated tau (Figs. 3 and 4) in the form of plaques and tangles. It is possible that an age-related increase in GSK3β activity dependent on its phosphorylation at Tyr216 is a factor in driving pathological tau hyperphosphorylation^52^ (Supplementary Fig. 13). With age, CYFIP2 reduction also promotes a glial response (Fig. 5), which is likely to be secondary to neuronal injury due to the absence of CYFIP2 expression in glial cells^22^. Functionally, CYFIP2 reduction may cause a failure of immature spines to stabilise into mushroom spines via reduced WAVE function^53^, resulting in age-dependent loss of dendritic spines (Fig. 6). This correlates with an initial spatial memory retention deficit^22^, which is then further exacerbated to an additional impairment in contextual memory formation with age (Fig. 6).

**Figure 7.**
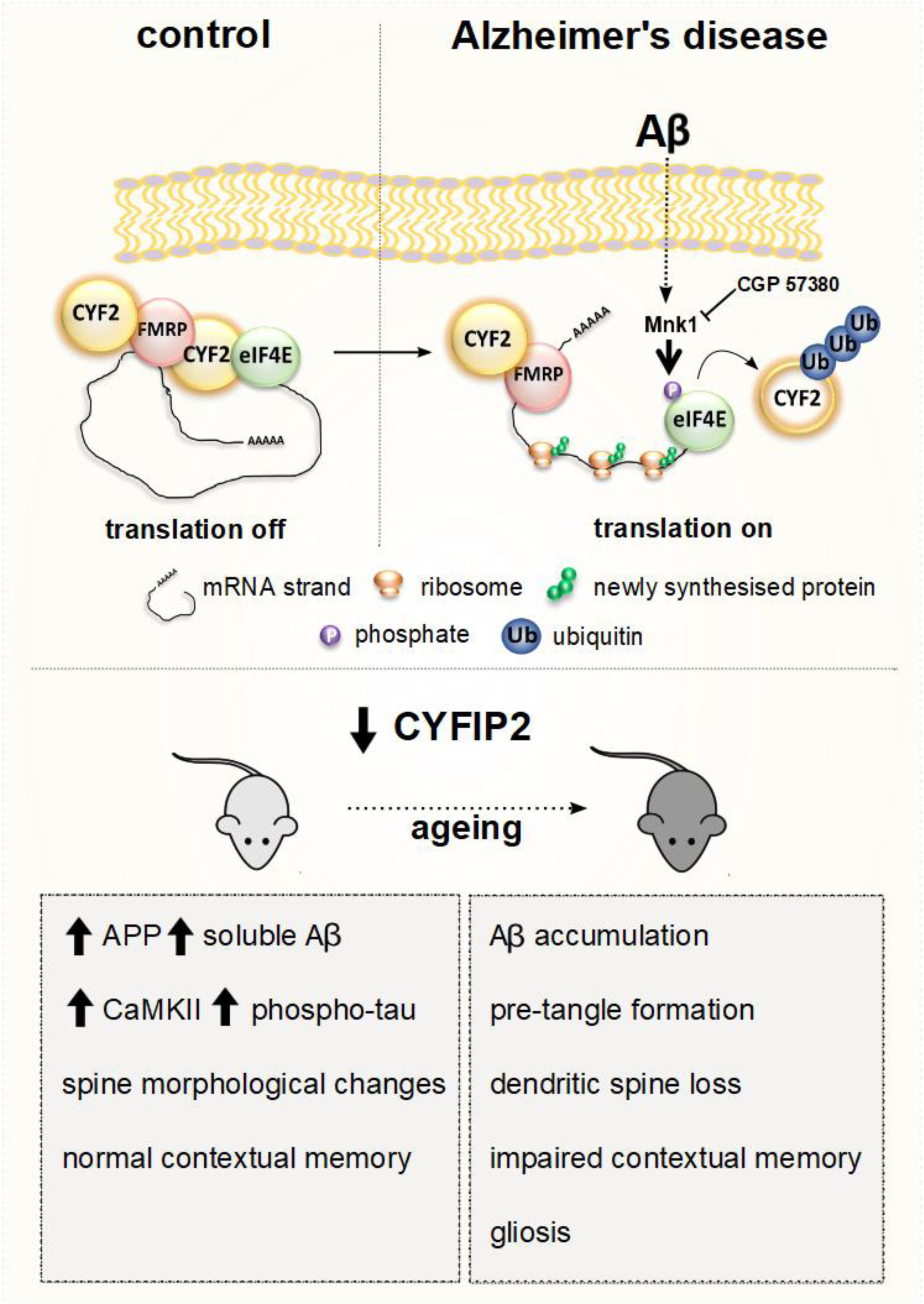
Proposed model of CYFIP2 dysregulation as a cause and consequence in Alzheimer’s disease pathogenesis. The schematic in the upper panel shows that in neurons of the healthy brain, mRNAs bound by FMRP are translationally repressed via the eIF4E-binding protein CYFIP2. In Alzheimer’s disease, increased production and/or reduced clearance of Aβ results in dissociation of this complex and prolonged translation of a subset of these mRNAs due to reduction of CYFIP2, which can be blocked by inhibition of Mnk1. The lower panel shows that in the young-adult mouse brain, loss of CYFIP2 elevates Aβ production and tau phosphorylation (at CaMKII sites), causes changes in dendritic spine morphology, and impairs spatial, but not contextual, memory. These effects are exacerbated by ageing, leading to accumulation of Aβ and of tau hyperphosphorylated at AT-8 and PHF-1 sites, dendritic spine loss, contextual memory impairment, and gliosis.

The presence of key AD-like characteristics in 12-month-old *Cyfip2*^+/-^ mice suggests it is useful as a model of AD. Amyloid pathology has previously been detected upon overexpression of mutant *human* APP, or humanisation of the murine Aβ_1-42_ sequence in a knock-in model^54^. Rodent Aβ_1-42_ differs from the human sequence by three amino acids (Arg^5^, Tyr^10^, and His^13^ are replaced by Gly, Phe and Arg respectively); as His^13^ is required for Aβ_1-42_ fibrillogenesis, it makes rodent Aβ_1-42_ less prone to forming amyloid aggregates^27^. A proteolytic processing difference between species may result in protection of the mouse sequence against β-secretase processing^28^; however, overexpression of murine Aβ_1-42_ can lead to formation of morphologically distinct plaques in an age-dependent manner, suggesting rodent Aβ_1-42_ can aggregate, but less efficiently than the human peptide^29^. At 12 months, *Cyfip2*^+/-^ mice develop Aβ_1-42_ accumulations in the thalamus, a vulnerable brain region^55^, which is understudied in AD research. Further characterisation will therefore be required of these Aβ_1-42_-positive inclusions in the *Cyfip2*^+/-^ mouse model.

AD-related tau pathology is particularly difficult to model, generally only occurring independently when non-physiological amounts of tau protein are overexpressed or when a point mutant version of tau is expressed, with the limitation that *MAPT* mutations do not cause AD^54^. Similarly, pre-tangle-like tau inclusions are only found following co-expression of familial AD-causing mutant human APP^54^, and not in double knock-in mice having both humanised APP and tau^56^. Instead, the *Cyfip2*^+/-^ mouse model, which lacks one copy of an endogenous gene, rendering lower expression of CYFIP2 similar as that present in post-mortem AD brain^22^, shows stereotypical progression of phospho-tau accumulation^44, 57^, and may therefore be a more physiological model of sporadic AD-related tau pathology. This has the promise to allow mechanistic understanding of the emergence of both amyloid and tau abnormalities in AD.

Our study shows direct evidence that nanomolar concentrations of Aβ_1-42_ preparations that include oligomers increase mRNA translation, particularly of specific transcripts that are predicted to be regulated by FMRP. The presence of unique candidates within both vehicle- and Aβ_1-42_-treated groups suggests the regulation of translation by Aβ_1-42_ is complex, as some mRNAs are preferentially translated or untranslated, while the majority of mRNA transcripts are unaffected. Further analysis will be required to identify common elements of these transcripts, as well as association or localisation to distinct polyribosomal populations. Our data suggests evidence for a biphasic regulation of protein synthesis in AD. In earlier stages there could be an increase in translation of specific mRNA transcripts, eventually leading to deposition of particular proteins. In later stages of the disease global protein synthesis may be shunted as a compensatory response^58^, ultimately leading to neurodegeneration^59^. Several lines of evidence suggest that prior to inducing neurotoxicity Aβ_1-42_ can function as a beneficial agent^60^, with its functions ranging from anti-microbial to enhancement of long-term potentiation and memory, akin to a neurotrophic factor. Our finding that nanomolar doses of Aβ_1-42_ enhance translation overall, via a pathway also modulated by the neurotrophin BDNF^61^, lends support to this idea. As Aβ_1-42_-induced signalling results in dissociation between eIF4E and both CYFIP1 and CYFIP2, it might regulate the translation of both CYFIP1 and 2-regulated mRNAs. However, the reduction of CYFIP2, but not CYFIP1, may act as a switch between reversible and irreversible processes, driving pathology.

Given that our data suggest that CYFIP2 reduction by Aβ_1-42_ can be blocked by Mnk1 inhibition, and CYFIP2 reduction results in age-dependent AD-like pathology in the mouse brain, we propose Mnk1 as a therapeutic target in AD. Our study finds tentative evidence that Mnk1 dysregulation may be associated with AD, and a previous report indicates excessive eIF4E phosphorylation in the AD brain^62^. The usefulness of Mnk inhibitors for the treatment of AD has already been noted, in part due to the role of these kinases in tau phosphorylation (patent no. WO2009065596A2)^63^. Our study sheds further light on the involvement of Mnk1 in AD, highlighting novel therapeutic avenues. The compound used in this study is unstable *in vivo*; however, a different Mnk inhibitor such as eFT508 which is already in clinical trials for treatment of solid tumours and lymphomas^33^ may be suitable.

Altogether, our study suggests that CYFIP2 reduction plays a pivotal role in AD pathogenesis and modelling its dysregulation will provide more physiological models of the disease. Our data also identify an upstream pathway which may be implicated in disease pathogenesis and can therefore be targeted for therapeutic intervention.

## Materials & Methods

### *Cyfip2*^+/-^ mice

As detailed previously^22^, *Cyfip2* heterozygous null mutant mice on the C57BL/6N genetic background were obtained from the Wellcome Trust Sanger Institute (Wellcome Trust Genome Campus, Cambridge, UK). Mice designated *Cyfip2*^+/-^ have a *Cyfip2*^tm1a(EUCOMM)Wtsi^ (ID:33461) allele generated by the European Conditional Mouse Mutagenesis Program (EUCOMM) which uses a ‘knockout-first’ design^18, 64^. Mice were maintained on the C57BL/6N genetic background. Animals were housed on a 12-hour light/dark cycle with food and water available *ad libitum* and genotyped by PCR as described previously^22^. All procedures were undertaken in accordance with the UK Animals (Scientific Procedures) Act 1986.

### Fear conditioning memory

All animals used for experiments were handled 2 min per day either for 3 days (3-4-month-old) or 5 days (12-month-old) before conditioning. All experiments were performed during the light cycle. During training, each mouse was placed into the chamber (Med Associates Inc., St. Albans, VT, USA) in a soundproof box. After a 120-second introductory period, a tone (75 dB, 10 kHz) was presented for 30 seconds, and the mouse received a 2-second foot shock (0.7 mA) which co-terminated with the tone. After an additional 30 seconds, the mouse was returned to the home cage. 24 hours after training, the mice were brought back to the conditioning chamber for 5 minutes to test for contextual fear memory. Freezing behaviour during each 2-second shock was scored every 5 seconds, blind to genotype.

### Immunohistochemistry/immunofluorescence (IHC/IF)

For histological analyses of aged mice, 12-month-old female *Cyfip2*^+/-^ mutants and wild-type littermates were used. To control for the effects of normal ageing, a group of 3-month-old female wild-type mice also on the C57BL/6N background were added to the analysis. Mice were anaesthetised with an intraperitoneal injection of Euthatal® (Merial, Toulouse, France), and intra-cardially perfused with a 4% paraformaldehyde solution after an initial PBS wash. Whole brains were removed and stored overnight in in 30% sucrose in PBS, then cut sagittally to separate left and right hemispheres, snap-frozen in isopentane, and stored at −80°C until ready for cutting and processing. Brains were mounted onto stages using cryo-embedding compound and coronally cut into 40 µm thick sections. For every animal (biological replicate), two sections from the same brain region (technical replicate) were stained per target. Prior to processing, floating sections were washed in TBS, then blocked in 2% normal goat serum in TBS with 0.1% TritonX-100 and incubated overnight with primary antibodies diluted in blocking solution. Antibodies used were AT-8 (Thermo Fisher MN1020, 1:100), PHF-1 (Peter Davies, 1:100), tau (Dako A0024, 1:1000), Iba1 (Wako Chemicals 019-19741, 1:200), GFAP (Dako Z0334, 1:1000), pGSK3β (Tyr216) (Abcam ab75745, 1:300), 4G8 (Calbiochem NE1002, 1:100). Sections were washed in TBS before incubating in a species-appropriate HRP-tagged (IHC) or fluorophore conjugated (IF) secondary antibody, diluted directly in blocking solution.

For IHC, sections were incubated in 2.5% v/v H_2_O_2_ in methanol for 30 minutes. After secondary antibody incubation sections were washed in TBS and incubated in 3,3’Diaminobenzidine (DAB) (Sigma) for 2-10 minutes depending on the antibody used, until the tissue appeared visibly brown, in small batches and blind to group and genotype. Sections were then washed twice in TBS for 10 minutes, mounted onto glass slides and allowed to dry. Slides were processed for counterstaining of nuclei using haematoxylin. Briefly, mounted sections were gently rinsed with water and then placed in Gill’s haematoxylin stain (Vector Laboratories) before being differentiated in 0.5% acid alcohol, washed, and dehydrated through a series of ethanol solutions (70%, 95%, 95%, 100%, 100%). Finally, sections were cleared in two changes of xylene before being coverslipped with xylene-based mounting media (Thermo Scientific) using an automated coverslipping system (Thermo Scientific). For IF, slides were coverslipped manually using a fluorescence-compatible mounting medium (Invitrogen ProLong™ Diamond Antifade mountant with DAPI).

Slides were imaged using the Olympus Slidescanner VS120 with Brightfield or Fluorescence mode. Images were taken with the 40x objective (NA 0.95, 0.17μm/pixel). Images were taken as automated maximal projections of z-stacks using the maximal density of focal points. Exposure times for fluorescent images were determined manually prior to the scan, and automatically maintained across sections for the same experiment.

### Amytracker™ staining

Manufacturer’s instructions were used to visualise protein aggregates in mouse brain sections with Amytracker™680 (Ebba Biotech). Briefly, sections were rinsed in PBS and fixed in ice-cold (−20°C) ethanol at RT for 10 minutes. Tissue sections were rehydrated in a 1:1 mixture of ethanol and water for 5 minutes and in PBS for 5 minutes. Sections were incubated with Amytracker™680 dye, diluted 1:1000 in PBS, for 30 minutes room temperature. Sections were washed in PBS, mounted onto glass slides with ProLong™ Diamond (Invitrogen), and coverslipped. Once dry, fluorescence was visualised using the Cy5 filter set of the Olympus Slidescanner.

### Dendritic spine analysis

Brains from 12-month-old female *Cyfip2*^+/-^ mutants and wild-type littermates were used to analyse spine density and morphology and processed for modified Golgi-Cox impregnation according to manufacturer’s instructions (FD Rapid GolgiStain™ kit, FD NeuroTechnologies, USA). Briefly, brains were isolated as quickly as possible, rinsed in water and immersed in impregnation solutions A and B at room temperature for 2 weeks in the dark. Tissue was transferred into Golgi solution C for 72 hours in the dark at room temperature. Brains were rapidly frozen by dipping into iso-pentane pre-cooled with dry ice and stored at −80°C until ready for sectioning. Coronal sections of 80 µm thickness were cut and mounted on double gelatine-coated slides. Sections were rinsed, stained with solutions D and E, dehydrated through an ethanol series, and cleared with xylene, before coverslipping with Permount®. Slides were allowed to dry overnight before imaging and analysis. Pyramidal neurons in the CA1 region of dorsal hippocampal sections were identified by their triangular soma shape and numerous dendritic spines. A 100x Plan Apo oil-immersion objective (N.A. 1.40, 0.07 μm/pixel) on the Eclipse Ti2 inverted microscope (Nikon) was used to image z-stacks of secondary and tertiary dendrites longer than 10 µm in the *stratum radiatum* (apical dendrites). Fifty dendritic segments were imaged and analysed for each group, and 10-12 cells were imaged per animal. Dendrites were reconstructed in 3D using the Neurolucida 360 system (MBF Bioscience, USA) and dendritic spines were identified and classified as thin, stubby, mushroom or filopodia using parameters based on three-dimensional structures of dendritic spines^65^.

### Primary neuronal culture

Primary cortical neuronal cultures were prepared from Sprague-Dawley rat E18 embryos, as described. Cells were seeded into culture plates coated with 0.2 mg/mL poly-D-lysine (Sigma) at a density of approximately 937 cells/mm^2^. Cells were cultured in Neurobasal medium supplemented with 2% B27, 0.5 mM L-glutamine, and 1% penicillin/streptomycin (Life Technologies, UK). After 4 days *in vitro* (DIV), 200 μM of D,L-amino-phosphonovalerate (D,L-APV, Abcam) was added to the media to maintain neuronal health over long-term culture and to reduce cell death due to excitotoxicity. Fifty percent media changes were performed weekly until desired time in culture was reached (27-28 DIV).

### Synthetic Aβ_1-42_ oligomer preparation

Oligomers from synthetic rat Aβ_1-42_ peptide (Calbiochem) were prepared as detailed^66^. Briefly, a stock solution was prepared at 100 μM in 200 mM HEPES (pH 8.5). The solution was gently agitated for 30 min at room temperature, aliquoted, and stored at −80°C. Once defrosted for treatments, aliquots were not subjected to any further freeze-thaw. This method of Aβ_1-42_ preparation is thought to form oligomers and not higher molecular weight or fibrillar aggregate forms^66^. To verify the composition of the Aβ_1-42_ generated, the NativePAGE™ (Invitrogen) system was used, and manufacturer’s instructions were followed. To confirm the presence of oligomers, Aβ_1-42_ peptides were treated in a 1:1 ratio with guanidine hydrochloride (GnHCl) which breaks down oligomers. An antibody raised against the juxtamembrane extracellular domain of Aβ_1-42_, spanning amino acids 17-24 (clone 4G8, Millipore MAB1561, diluted 1:1000), was used for detection of rodent Aβ_1-42_ oligomers. As a control, commonly used human Aβ_1-42_ peptides solubilised either in hexafluoroisopropanol (HFIP) or trifluoroacetic acid (TFA) were also prepared by removal of solvent and oligomerisation in PBS at 37°C for 3 hours (Supplementary Fig. 1).

### Pharmacological treatments

All treatments were performed directly in cell culture media. Stock solutions of cycloheximide (Sigma) and Mnk1 inhibitor compound CGP 57380 (Tocris) in DMSO were diluted to final concentrations in the cell culture medium, using equal volumes of DMSO as control. For oligomeric Aβ_1-42_ treatments, aliquots were defrosted just prior to treatment, and diluted to a final concentration of 100 nM in cell culture medium, using equal volumes of HEPES buffer as control. The surface sensing of translation (SUnSET) assay was used to monitor protein synthesis in cell cultures, following a published protocol^56^. Briefly, puromycin (Sigma) was diluted to a final concentration of 10 μg/ml in culture medium for the last 10 minutes of the treatment. Samples were lysed and run as a western blot, using an anti-puromycin antibody (Kerafast EQ0001, 1:1000). Prior to blocking and primary antibody incubation, total protein levels on the membrane were detected using the Revert™ Total Protein Stain Kit (LI-COR) as per manufacturer’s instructions.

### Cell lysis

At the end of the treatment culture medium was removed and plates were placed on an ice-block. Cells were rinsed briefly with ice-cold PBS, then lysed in equal volumes of lysis buffer (20 mM Tris pH 7.4, 150 mM NaCl, 1% Triton X-100, 5 mM EDTA pH 8) supplemented with protease and phosphatase inhibitor cocktails (Sigma), using an ice-cold plastic cell scraper. Lysates were frozen at −20°C. Protein amounts were quantified using a BCA kit (Pierce) and proteins were detected by immunoblotting.

### Immunoprecipitation

For immunoprecipitation (IP) in cultured neurons, cells were briefly rinsed in ice-cold PBS, then lysed on ice in cold IP buffer (50 mM Tris pH 7.4, 150 mM NaCl, 1% Triton X-100) supplemented with protease inhibitor cocktail (Sigma). For IP in the total mouse hippocampus, frozen tissue was homogenised in the same IP buffer with 20 strokes in a Dounce homogeniser (Smith Scientific). The cell lysate/brain homogenate was centrifuged at 16,089 x *g*, 4°C for 1 minute and the supernatant used as input for IP. For IP in synaptosomes in mouse hippocampus, crude synaptosomal fractions were freshly prepared, as detailed previously ^22^. Protein samples were incubated with pre-cleared magnetic Dynabeads™ Protein A (Invitrogen) and primary antibodies against CYFIP1 (Millipore) or CYFIP2 (GeneTex) were added in a 1:50 ratio to the lysate-bead mixture. Tubes were rotated at 4°C overnight, spun down briefly, and the supernatant removed using a magnetic rack. Beads were washed 3x in cold IP buffer, before adding Laemmli sample buffer and boiling at 95°C for 10 minutes. This IP fraction was used fresh for protein detection by immunoblotting. For certain analyses, values were normalised to those of the vehicle control to minimise error between biological replicates. For quantification of protein ubiquitination, cells treated with the proteasomal inhibitor MG132 were used as a positive control to determine the correct molecular weights of ubiquitinated proteins.

### Western blotting

Protein samples were diluted and boiled at 95°C in Laemmli sample buffer. 10-20 µg of proteins were separated on 4–15% Criterion™ TGX™ precast gels (Bio-Rad) and then transferred onto a methanol-activated 0.2 μm polyvinylidene fluoride (PVDF) membrane (Bio-Rad). Membranes were blocked at room temperature for 1 h (5% milk or BSA in TBST pH 7.5) and then incubated in primary antibody diluted in blocking buffer overnight at 4°C. Primary antibodies were detected using horseradish peroxidase (HRP)-conjugated secondary antibodies (Dako) and chemiluminescent reagent (Thermo Scientific), and signals in the linear range obtained by exposing membranes to X-ray films (Amersham). Prior to probing with other primary antibodies, the membranes were washed in western blot stripping buffer (Santa Cruz Biotechnology). Primary antibodies used were against CYFIP1 (Millipore 07-531, 1:10,000), CYFIP2 (GeneTex GTX124387, 1:10,000), phospho-eIF4E Ser209 (Cell Signaling 9741 1:1000), eIF4E (Cell Signaling 2067, 1:1000), NSE (Millipore AB951, 1:60,000), α-synaptotagmin (Sigma S2177, 1:30,000), AT-8 phospho-tau (Invitrogen MN1020, 1:1000), PHF-1 phospho-tau (Peter Davies, 1:1000), total tau (Dako, A0024, 1:10,000) and ubiquitin (Millipore 05-1307, 1:1000). Signals were analysed with ImageJ software (NIH) and normalised to appropriate loading controls.

### Ribosomal RNA extraction

For analysis of ribosome-bound mRNA, primary rat cortical neurons were seeded onto 20 mm PDL-coated culture dishes at a density of 636 cells/mm^2^ and treated with the Aβ_1-42_ preparation at 27 DIV for 24 hours. During the last 10 minutes of treatment on 28 DIV 100 µg/mL cycloheximide was added to the culture media to inhibit ribosome drop-off. Once media was removed, cells were placed on a cold block and washed with ice-cold PBS plus 100 µg/mL cycloheximide. Cells were then scraped into ice-cold polysome lysis buffer (10 mM HEPES-KOH (pH 7.4), 5 mM MgCl_2_, 150 mM KCl, 1% NP-40) freshly supplemented with 0.5 mM DTT, 100 U/mL RNasin RNase inhibitor (Promega), 100 µg/mL cycloheximide, and EDTA-free protease inhibitors (Cell Signaling). The lysate was syringed three times through a 23G needle on ice, then centrifuged at 10,000 x *g* for 5 min at 4 °C. The supernatant was layered onto a 20% sucrose cushion prepared in the same lysis buffer and then centrifuged at 186,000 x *g* for 2 h at 4 °C in a TLA-55 rotor ultracentrifuge (Beckman Coulter). RNA was isolated from the ribosome-enriched pellet using TRIzol LS reagent according to manufacturer’s instructions (Invitrogen).

### RNA sequencing and analysis

Library preparation and RNA-sequencing was done by Novogene (Hong Kong). Libraries were made using polyadenylated mRNA isolation and employing NEBNext® Ultra™ RNA Library Prep Kit for Illumina® (NEB, USA). Three samples had lower RNA amounts and only a higher efficiency bead capture was employed for those, maintaining the same library preparation kit otherwise. Base Quality and Phred score were calculated employing the Illumina CASAVA v1.8 software with more than 92% of all reads for all samples having a Q30. Bowtie v2.2.3 was used to index the reference genome and trimmed paired-end reads were aligned using TopHat v2.0.12. HTSeq v0.6.1 using the union mode was employed to count mapped reads and generate FPKM (Fragments per Kilobase of transcript sequence)^67^. Differential gene expression analysis was performed employing DESeq^68^ with Benjamini and Hochberg’s FDR correction. Identification of putative FMRP fragments was done by cross-referencing differentially expressed genes between total and ribosome-enriched fractions with previously published datasets^12^ employing R. Heatmap (unsupervised clustering using Pearson), Venn diagram and correlation plot were made using R. All packages are available in CRAN.

### Statistical analysis

All data was analysed using GraphPad Prism version 8. Data was tested for normality with a Shapiro-Wilk test. Data points falling beyond 2*σ (SD) from the mean were considered statistical outliers and removed from every analysis. Statistical significance was determined by Student’s t-test or one-way or two-way analysis of variance (ANOVA) with post-hoc Tukey’s correction or Student-Newman-Keuls method. For non-parametric data, Kruskal-Wallis one-way ANOVA with Dunn’s testing for multiple comparisons was used. For FMRP analysis a Kolmogorov-Smirnov test was employed to compare cumulative distributions between targets and non-targets; a Fisher’s exact 2-tailed test was done to determine enrichment of FMRP targets in genes differentially present in ribosomal-enriched fractions. Results having *p*<0.05 were considered significant in every analysis.

### Genome-wide association studies (GWAS)

Single Nucleotide Polymorphisms (SNPs) within 10 kb upstream and downstream of the coding region of each gene were screened for associations with AD using the latest Genome Wide Association (GWA) and meta-analysis of diagnosed Alzheimer’s disease (AD)^38^. To account for linkage disequilibrium (LD), tag SNPs were created with the Priority Pruner software (version 0.1.4, http://prioritypruner.sourceforge.net/index.html), which prunes SNPs by removing all those that are in LD with other SNPs in the dataset. The 1000 Genomes (Phase 3) was used as a reference panel for pruning using the Kunkle et al., dataset p-values for prioritisation. SNPs with an r^2^ > 0.8 and within 250KB were determined to be in LD. For the selected SNPs, an FDR threshold of 0.05 was applied to correct for multiple testing when looking for associations with AD (“FDRtools” Rstudio, Version 1.2.1335).

To gain insight into potential regulatory effects of the genetic variants associated with AD in the five genes, expression quantitative trait loci (eQTL) data from the Genotype-Tissue Expression (GTEx)(Version 6) Project Consortium^39^ and BRAINEAC (www.braineac.org/)^69^ were used to determine whether variants that showed associations with AD (after correction for multiple testing) affect gene expression as eQTLs. GTEx and BRAINEAC are high-quality databases of matched genotype and gene expression measurements, which enable the quantification of effects of SNPs on gene expression in various tissues, including various brain tissues.

The BRAINEAC eQTLs dataset includes 10 brain regions from 134 neuropathologically normal individuals of European descent. The 10 brain regions are cerebellar cortex, frontal cortex, hippocampus, medulla, occipital cortex, putamen, substantia nigra, temporal cortex, thalamus, and intralobular white matter.

The GTEx (version 6) eQTLs dataset includes 53 tissues, 544 donors and 8555 samples. 10 human brain tissues were selected including anterior cingulate cortex, caudate basal ganglia, cerebellar hemisphere, cerebellum, cortex, frontal cortex BA9, hippocampus, hypothalamus, nucleus accumbens basal ganglia, and putamen basal ganglia, which include at least 70 samples.

## Acknowledgements

We thank Kei Cho, Wendy Noble, and Alessio Delogu for helpful discussions. We also thank Peter Davies for kindly providing us with PHF-1 antibody, Wendy Noble for Iba1 antibody and Amytracker™ dye, and Claire Troakes for 4G8 antibody. This work was supported by an MRC IoPPN excellence PhD studentship and an Alzheimer’s Research UK (ARUK) King’s College London Network Centre pump priming grant to AG, and an ARUK pilot grant (ARUK-PPG2018A-002) to KPG. The use of the Nikon Eclipse Ti2 inverted microscope was possible due to funding by ARUK-EG2013B-1 and the Nikon Wohl Cellular Imaging Centre (WCIC).

## Author contributions

A.G. and K.P.G. designed the project and wrote the manuscript. A.G. performed most of the experiments and analysed the data. R.T.M.N. advised on the polyribosome experiment, analysed the RNA sequencing data and wrote the relevant sections. K.M. and S.S.T. performed behavioural experiments and K.M. wrote the relevant sections. P.P. performed the GWAS analysis and wrote the relevant sections. B.G.P.N. performed the Native PAGE. E.G. provided all the primary neurons used in this study. All authors discussed the results and commented on the manuscript.

## Materials & Correspondence

All requests for materials and any correspondence should be addressed to.

