## Supplementary figures and tables for "Alzheimer’s disease-related dysregulation of protein synthesis causes key pathological features with ageing"

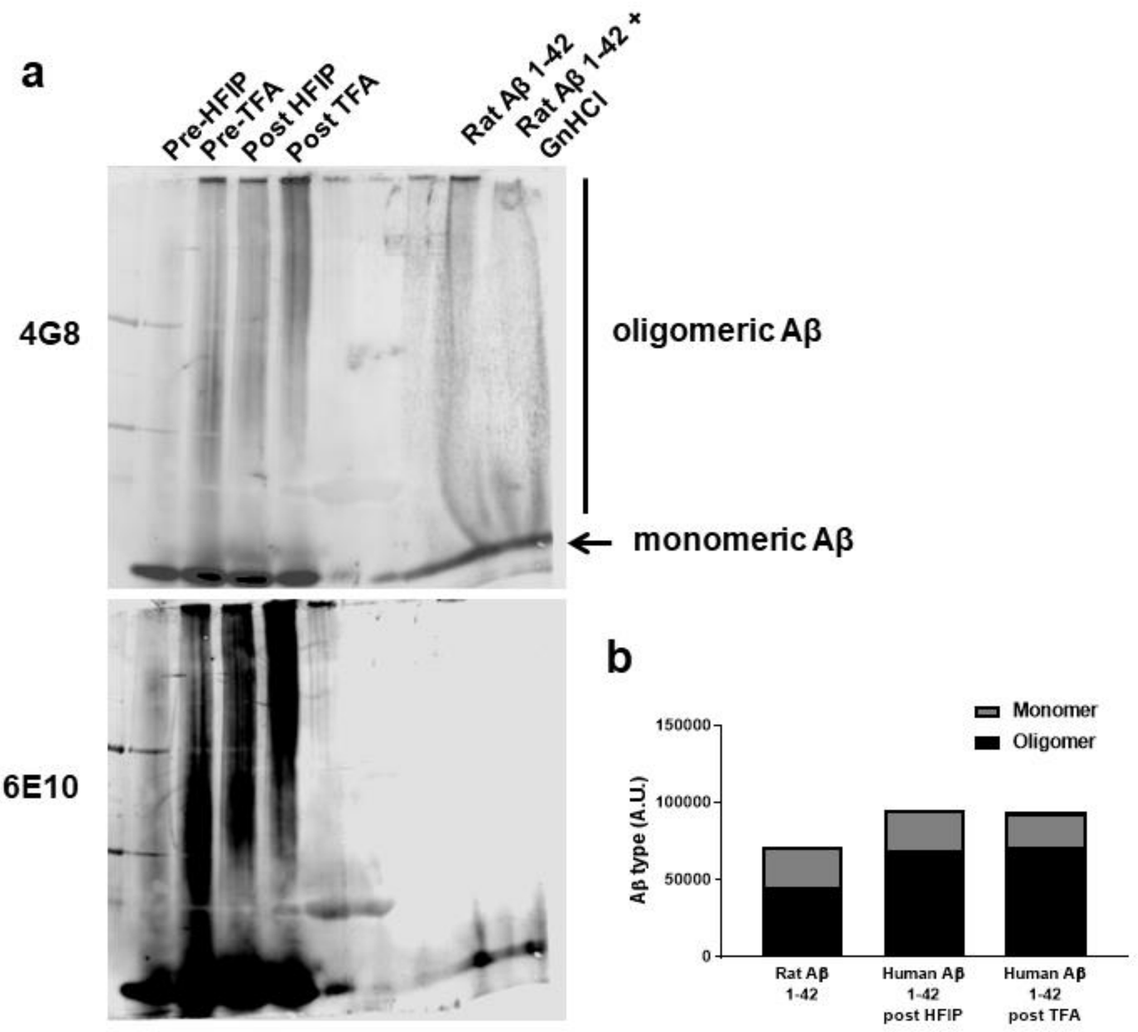


**Supplementary Figure 1. Composition of $A\beta_{1-42}$ peptides used in this study, as determined by Native-PAGE. a** Western blot to profile $A\beta_{1-42}$ generated. The first two lanes ("pre") show monomeric human $A\beta_{1-42}$ peptides solubilised in hexafluoroisopropanol (HFIP) or trifluoroacetic acid (TFA). The second two lanes ("post") show these peptides following evaporation of solvent and oligomerisation in PBS at 37°C for 3 hours. The last two lanes show synthetic rat $A\beta_{1-42}$ peptides prepared as detailed, before and after treatment with guanidine hydrochloride (GnHCl) which breaks down oligomers. The anti-4G8 (Aβ, aa 17-24) antibody is able to detect both the human and rat peptides, whereas the anti-6E10 (Aβ, aa 1-16) antibody can only detect the human peptide due to the sequence differences between species in this region of the peptide. **b** Quantification of the ratio of monomers to oligomers produced by the three different methods as detected by the anti-4G8 antibody, showing that the method used in this study with rat $A\beta_{1-42}$ peptides generates fewer oligomers compared to monomers than standard methods using the human $A\beta_{1-42}$ peptide. Only rat peptides were used in this study.

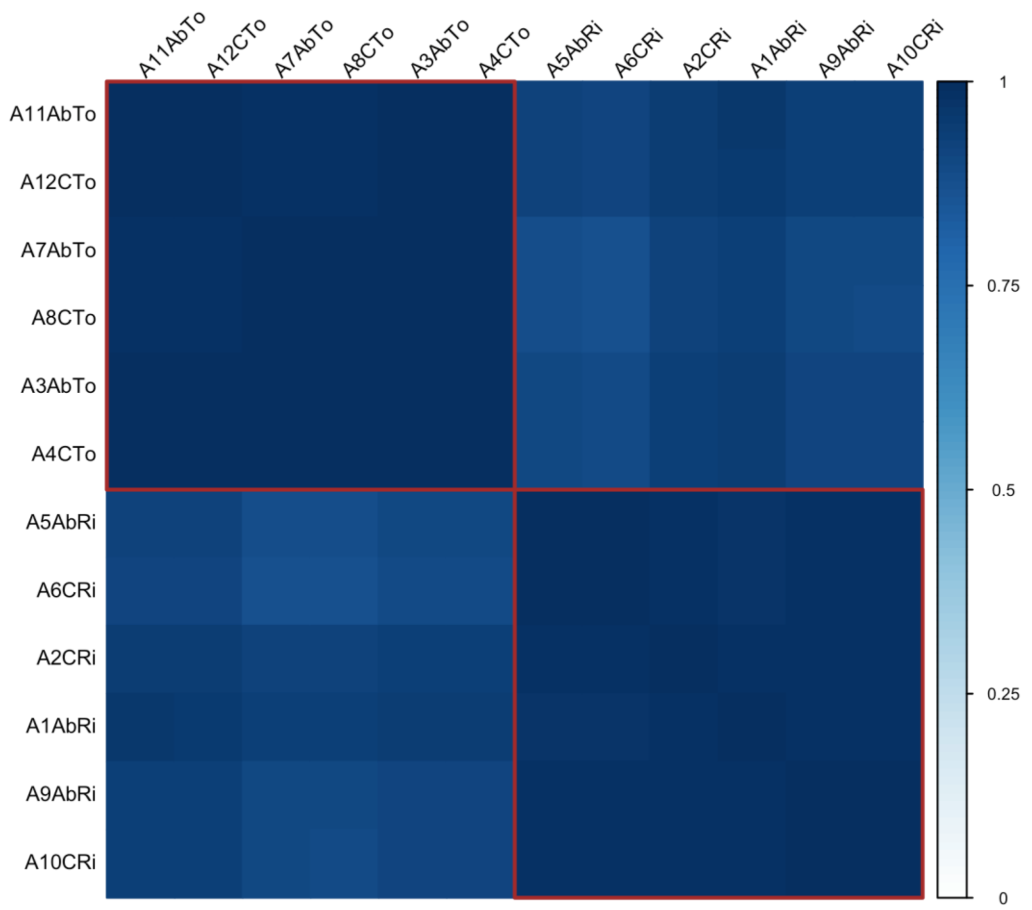


**Supplementary Figure 2.** Correlation plot showing a closer correlation within total mRNA samples and ribosome-enriched samples, in a subcellular fraction-dependent manner. This suggests that the ribosomal enrichment protocol results in distinct mRNA populations which are similar across different biological replicates. All mRNAs detected (FPKM) were employed in generating these plots.

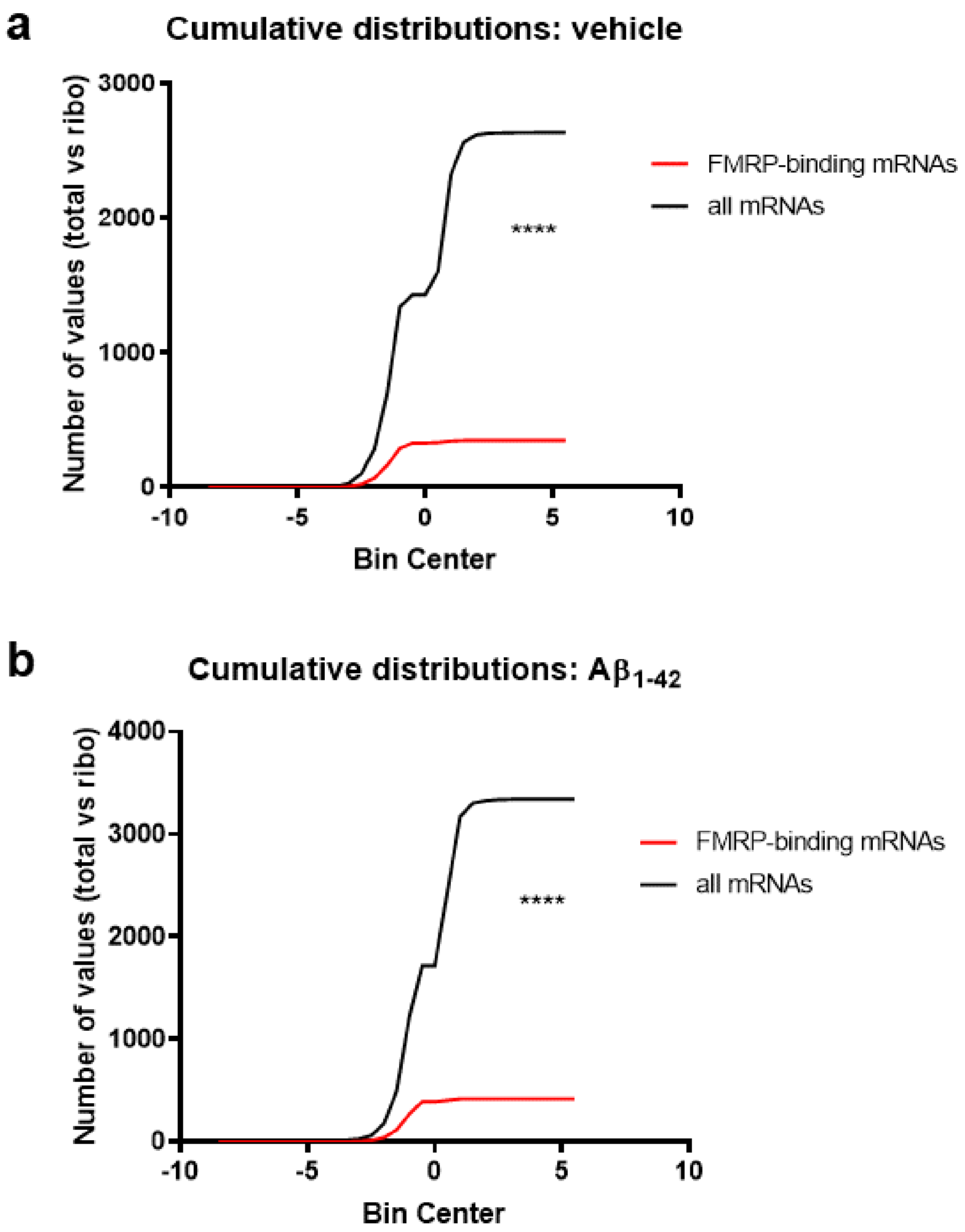


**Supplementary Figure 3. FMRP targets are present in both vehicle and Aβ$_{1\text{-}42}$-treated cells. a** and **b** Cumulative distributions of mRNA changes of 'Total' versus 'Ribo' in vehicle (**a**) and Aβ$_{1\text{-}42}$ (**b**) for FMRP versus non-FMRP targets. These graphs show that the prediction of FMRP binding causes a change in the distribution of mRNA changes between total and ribosome-enriched fractions in both vehicle- and Aβ-treated cells. **** $p < 0.0001$.

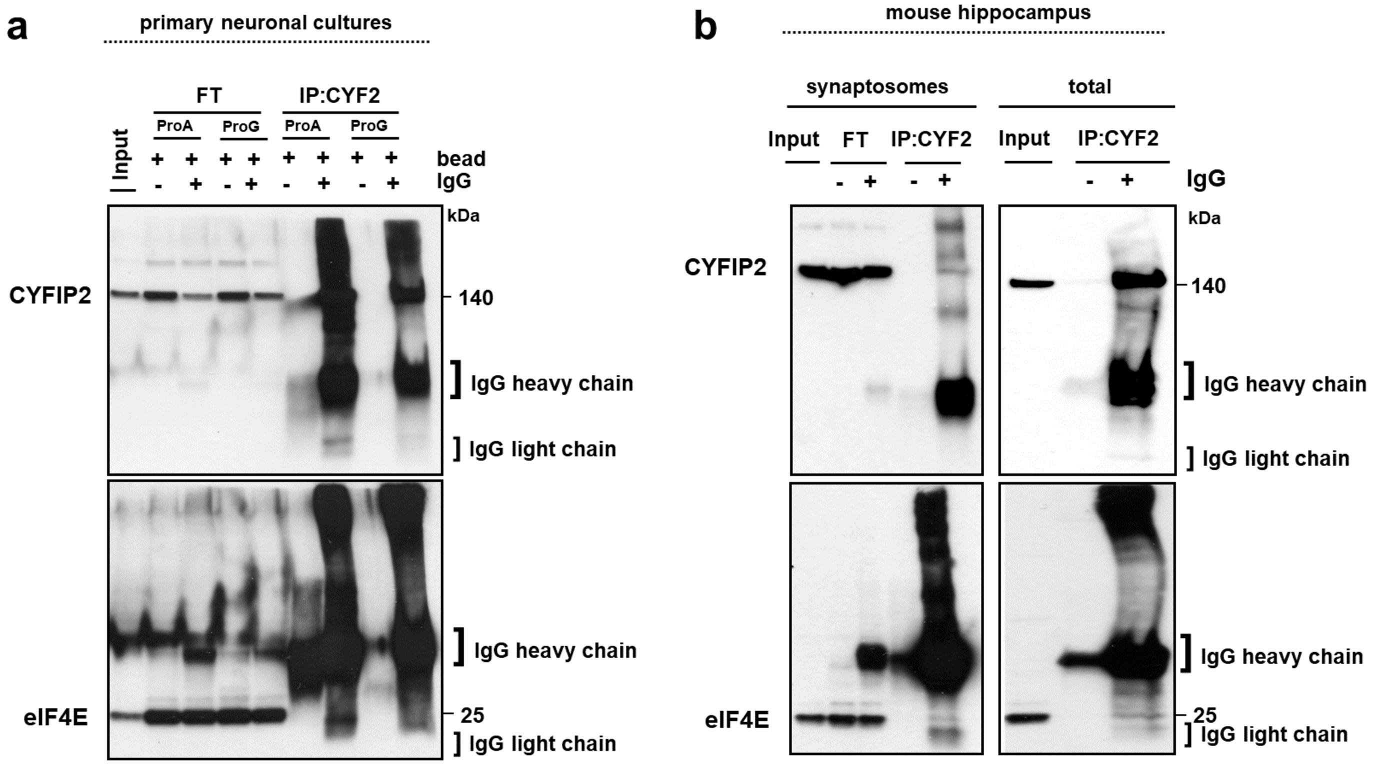


**Supplementary Figure 4. CYFIP2 co-immunoprecipitates with eIF4E in rat primary cortical neurons and in the mouse hippocampus. a** Blot showing immunoprecipitation of CYFIP2 using a primary antibody against rabbit CYFIP2 (IgG) in lysates prepared from rat primary cortical neurons at 28 DIV. Beads covalently coupled with either Protein A (**ProA**) or Protein G (**ProG**) were tested to determine efficacy. The input, FT, and IP fractions were visualised by immunoblotting for CYFIP2 and eIF4E. **b** Blots showing immunoprecipitation of CYFIP2 using the same primary antibody against rabbit CYFIP2 (IgG) in either the crude synaptosomal fraction (left) or in the total homogenate (right) prepared fresh from the mouse hippocampus. Protein A-coupled beads were used, and the input, FT, and IP were visualised by immunoblotting for CYFIP2 and eIF4E. **FT** flow-through, **IP** immunoprecipitate.

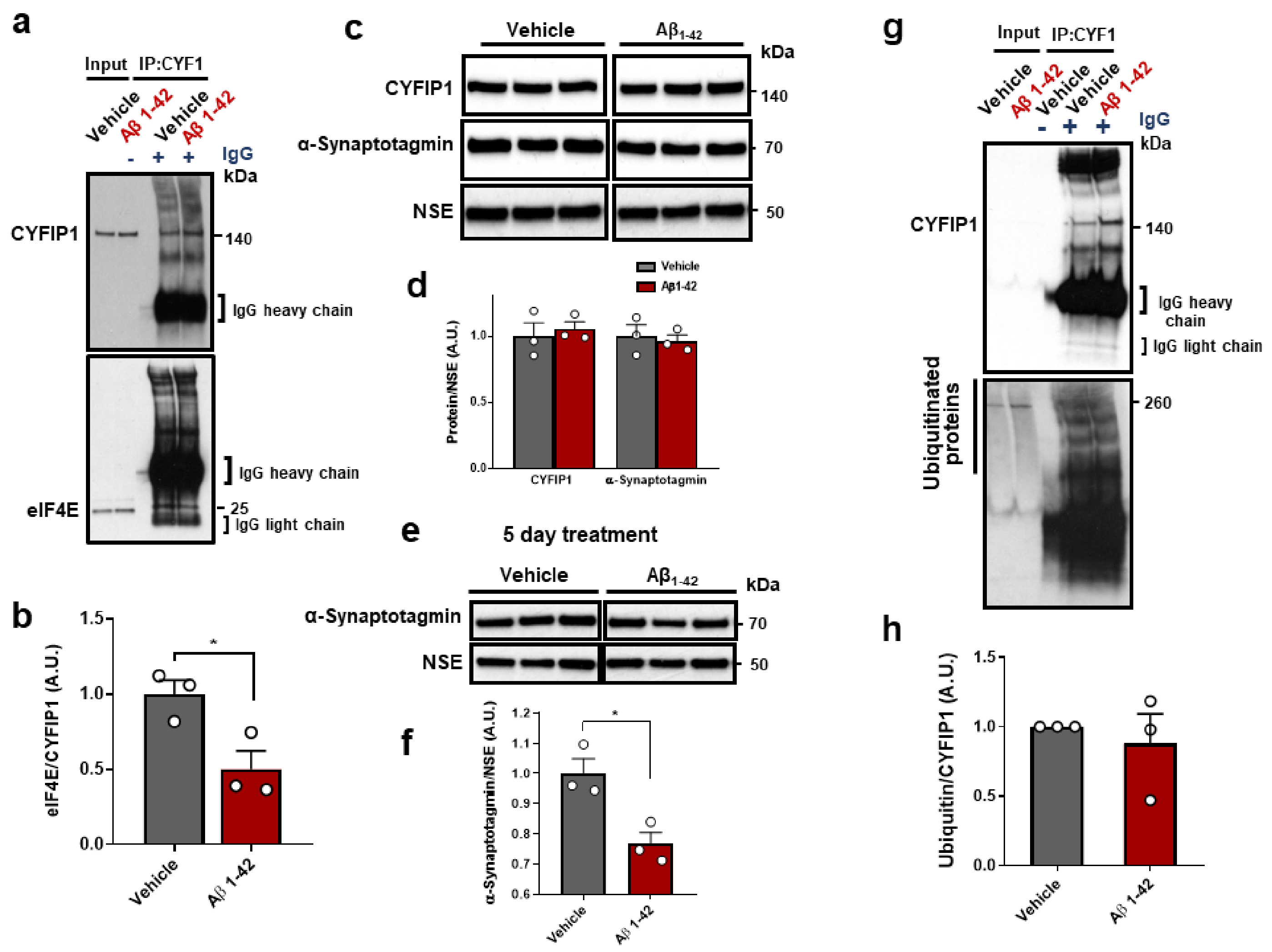


**Supplementary Figure 5. $A\beta_{1-42}$ treatment results in dissociation between CYFIP1 and eIF4E but does not change CYFIP1 levels.** **a** Blots showing immunoprecipitation of CYFIP1 using a primary antibody against rabbit CYFIP1 (IgG) in lysates prepared from rat primary cortical neurons at 28 DIV, treated for 24 hours with vehicle or 100 nM Aβ1-42. The input, FT, and IP fractions were visualised by immunoblotting for CYFIP1 and eIF4E. **b** Quantification of eIF4E bound to CYFIP1 pulled down in the IP fraction in vehicle- and $A\beta_{1-42}$-treated cells. **c** Blots showing levels of CYFIP1, α-synaptotagmin and NSE in 28 DIV primary cortical neurons after 24-hour treatment with vehicle or $A\beta_{1-42}$ at 100 nM. **d** Quantification of CYFIP1 and α-synaptotagmin normalised to levels of NSE. **e** Blots showing levels of α-synaptotagmin and NSE in 28 DIV primary cortical neurons after 5-day treatment with vehicle or $A\beta_{1-42}$ at 100 nM. **f** Quantification of α-synaptotagmin normalised to levels of NSE. **g** Blots showing immunoprecipitation of CYFIP1 in primary cortical neurons treated for 24 hours with vehicle or 100 nM $A\beta_{1-42}$. The input, FT, and IP fractions were visualised by immunoblotting for CYFIP1 and ubiquitin. **h** Quantification of ubiquitinated proteins normalised to CYFIP1 pulled down in the IP fraction in vehicle- and A$\beta_{1-42}$-treated cells. Data represent mean ± SEM. n=3 biological replicates (individual experiments shown as white dots). * $p < 0.05$. **FT** flow-through, **IP** immunoprecipitate.

**a**

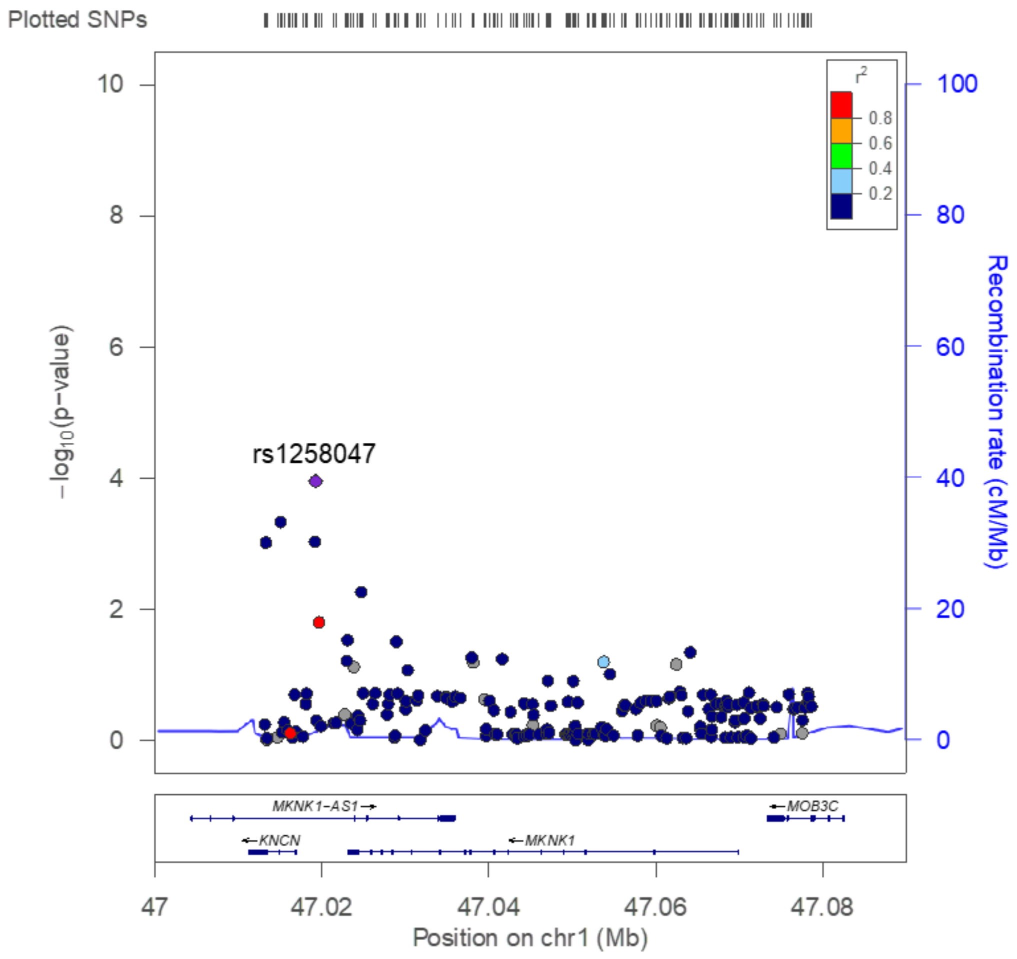
Plotted SNPs
rs1258047
−log10(p−value)
Recombination rate (cM/Mb)
r2
MKNK1−AS1
KNCN
MKNK1
MOB3C
Position on chr1 (Mb)

**b**

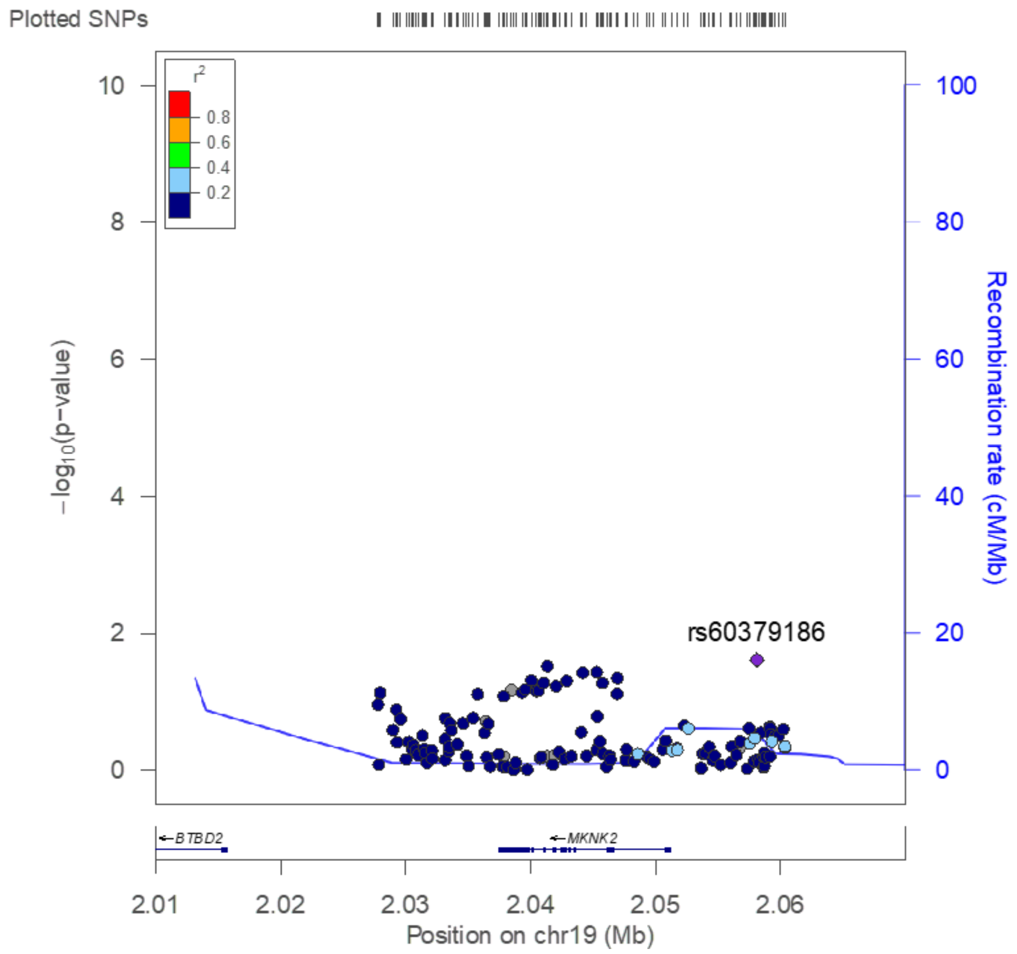
Plotted SNPs
rs60379186
−log10(p−value)
Recombination rate (cM/Mb)
r2
BTBD2
MKNK2
Position on chr19 (Mb)

**c**

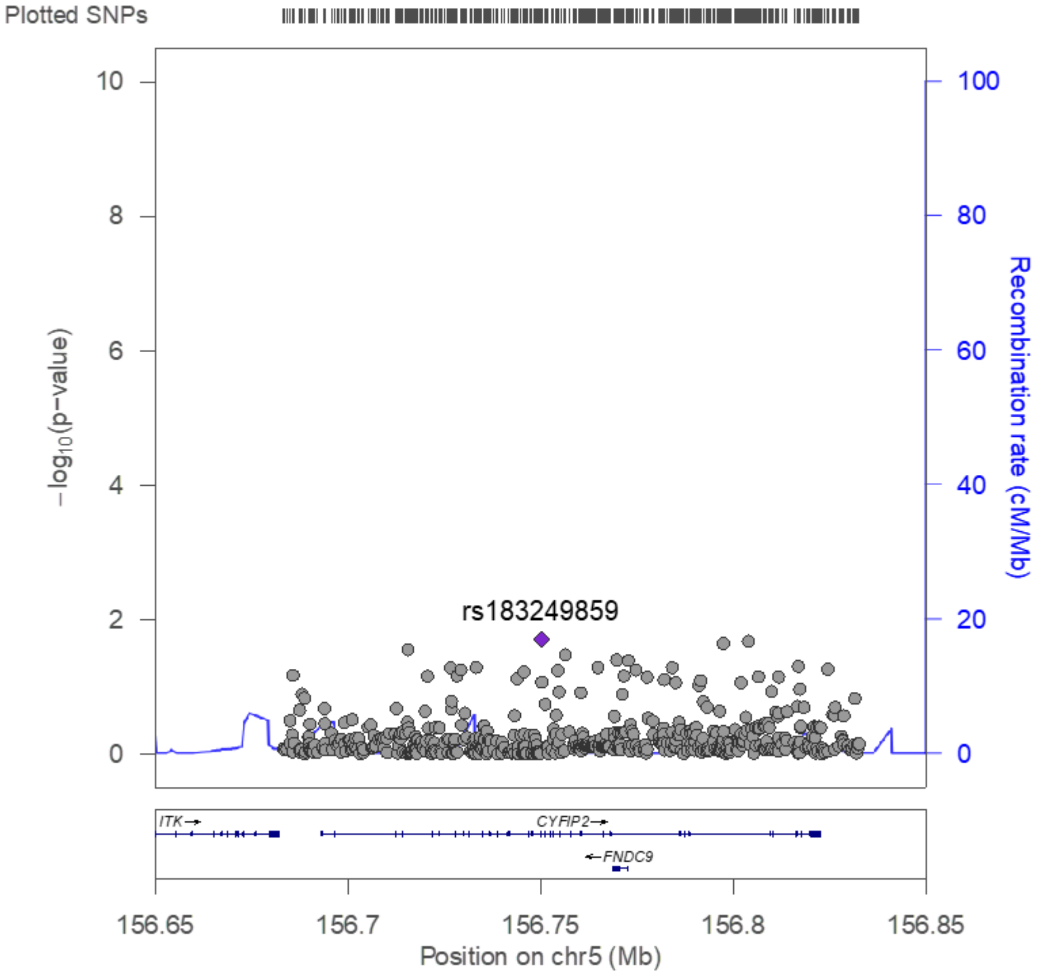
Plotted SNPs
10
8
6
4
2
0
−log10(p−value)
100
80
60
40
20
0
Recombination rate (cM/Mb)
rs183249859
ITK
CYFIP2
FNDC9
156.65
156.7
156.75
156.8
156.85
Position on chr5 (Mb)

**d**

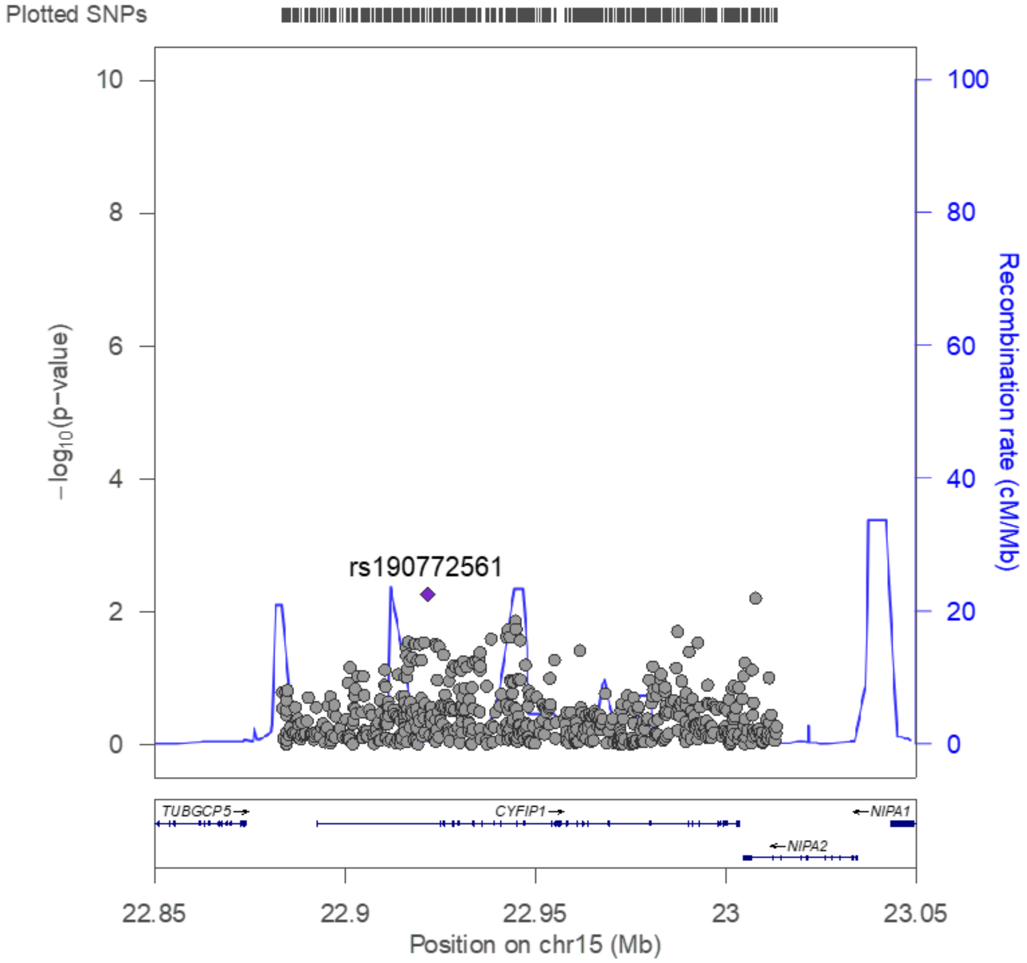
Plotted SNPs
10
8
6
4
2
0
−log10(p−value)
100
80
60
40
20
0
Recombination rate (cM/Mb)
rs190772561
TUBGCP5
CYFIP1
NIPA1
NIPA2
22.85
22.9
22.95
23
23.05
Position on chr15 (Mb)

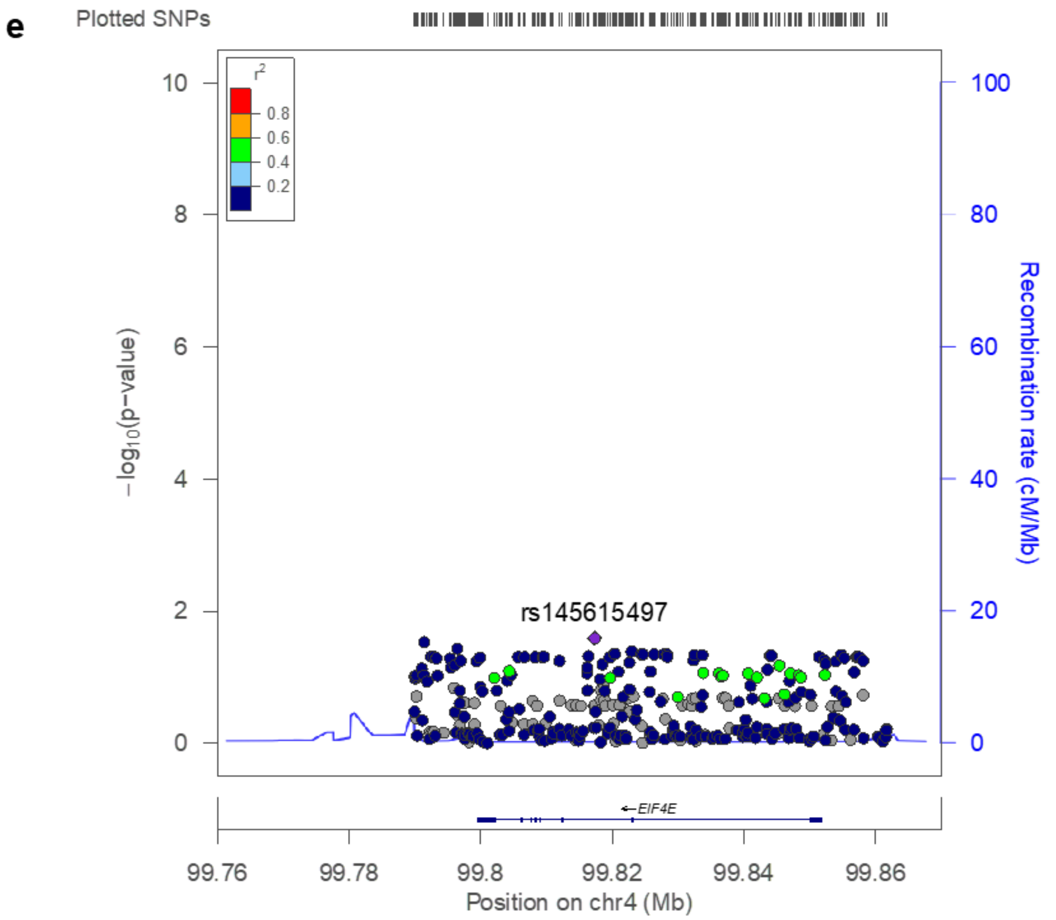


**Supplementary Figure 6. Plots showing single nucleotide polymorphisms (SNPs) within five genes, *MKNK1*, *MKNK2*, *CYFIP2*, *CYFIP1*, and *EIF4E*, tested for association with Alzheimer's disease (AD).** No SNPs were associated with AD following correction for multiple testing. No usable LD information was available for the top SNP for *CYFIP1* and *CYFIP2* SNPs (shown as grey dots).

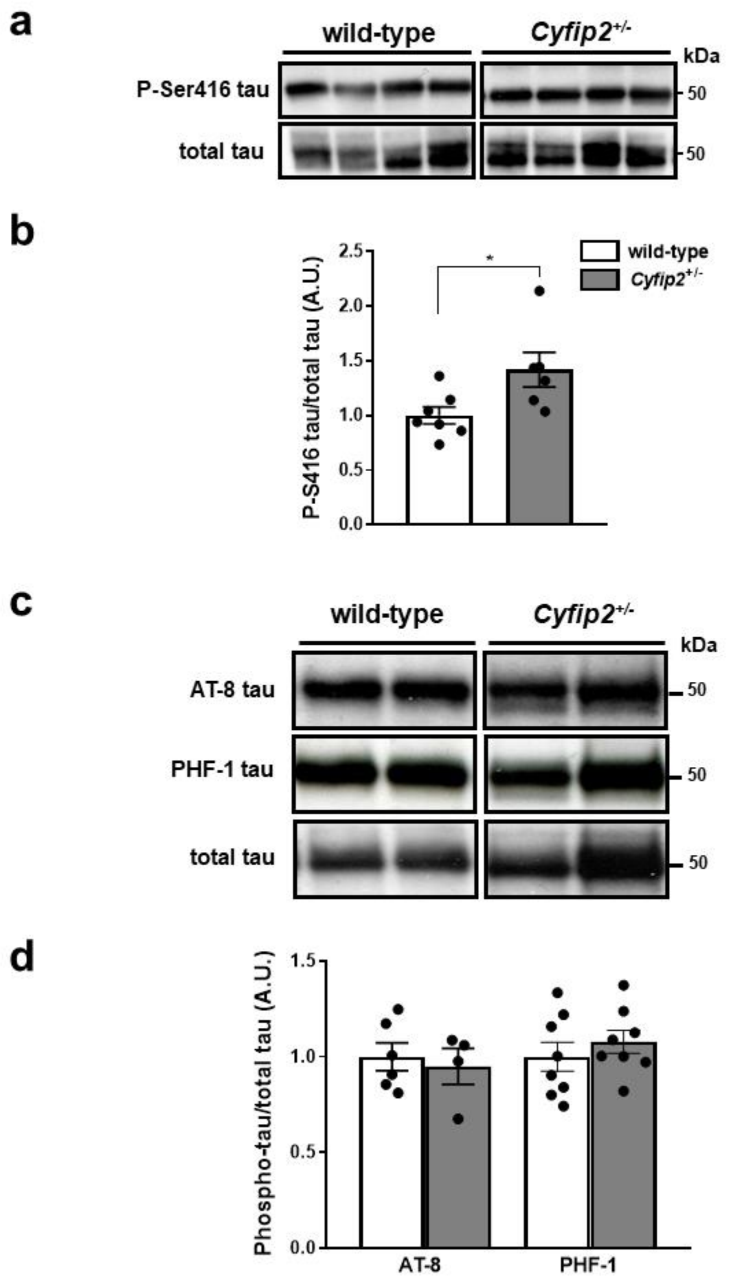


**Supplementary Figure 7. Tau phosphorylation at Ser416, but not at AT-8 or PHF-1 sites, is increased in hippocampal synaptosomes of 3-4-month-old *Cyfip2*$^{+/-}$ mice. a** Representative blots of phospho-Ser416 tau and total tau in synaptosomes prepared from hippocampal tissue from 3-4-month-old *Cyfip2*$^{+/-}$ mice and wild-type littermates. **b** Quantification of phospho-Ser416 levels normalised to total tau reveals a significant increase in young, adult *Cyfip2*$^{+/-}$ mice. **c** Representative blots of AT-8 and PHF-1 phospho-tau and total tau in synaptosomes prepared from hippocampal tissue from *Cyfip2*$^{+/-}$ and wild-type mice. **d** Quantification of AT-8 and PHF-1 levels normalised to total tau in the synaptosomal fraction does not detect a difference between genotypes. Data represent mean ± SEM. n=4-8 animals per condition (individual animals shown as black dots). * $p < 0.05$.

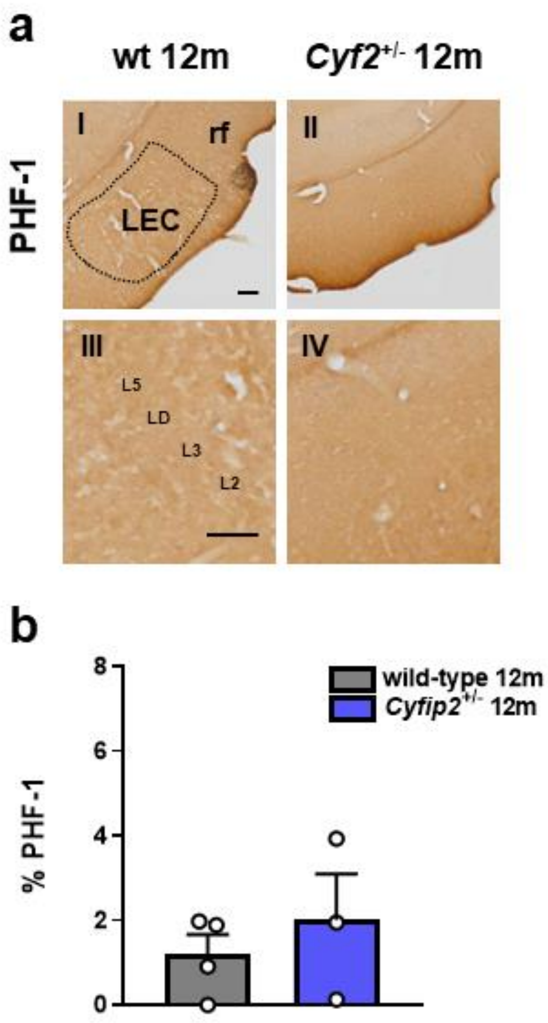


**Supplementary Figure 8. Negligible PHF-1 positive immunostaining in the entorhinal cortex of 12-month-old *Cyfip2*$^{+/-}$ and wild-type mice. a** Representative images of PHF-1 staining in **LEC** of 12-month-old *Cyfip2*$^{+/-}$ and wild-type mice (**I-II**). Higher magnification images of PHF-1 in the LEC (**III-IV**). **b** Quantification of PHF-1 positive staining in LEC does not detect a difference between genotypes. Scale bars are 100 µm. Data represent mean ± SEM. n=3-4 animals per condition (individual animals shown as white dots). **L2-5** layers 2-5 **LD** *lamina dissecans* **LEC** lateral entorhinal cortex **rf** rhinal fissure.

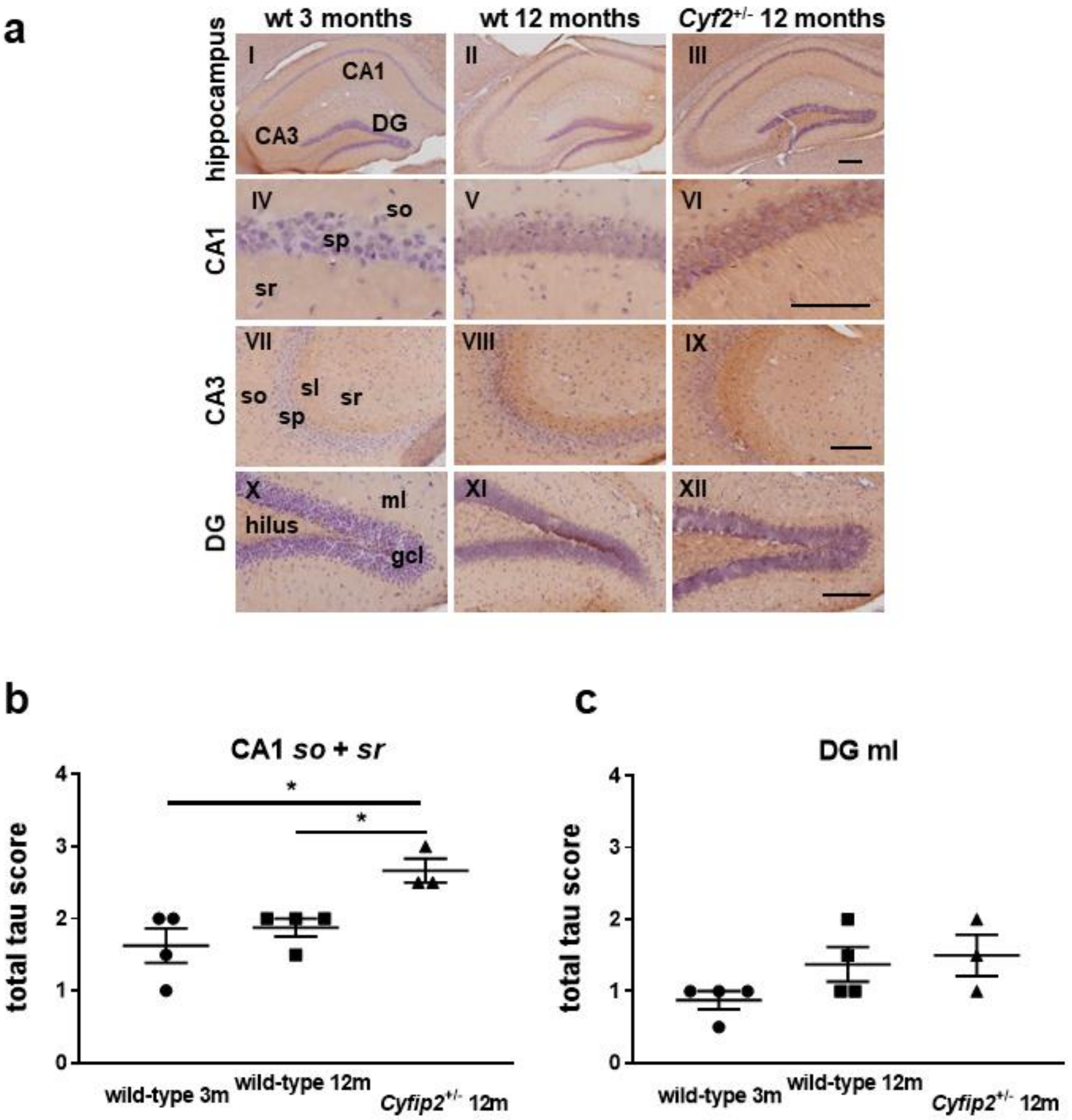


**Supplementary Figure 9. Restricted change in tau levels in 12-month-old *Cyfip2*$^{+/-}$ mice. a** Representative images of dorsal hippocampal sections from wild-type mice at 3 months and 12 months, and *Cyfip2*$^{+/-}$ mice at 12 months, immunohistochemically stained for total tau (**I-III**) and haemotoxylin to label nuclei. Higher magnification images of area CA1 (**IV-VI**), CA3 (**VII-IX**) and DG (**X-XII**). Scale bars are 200 µm. **b** Analysis of DAB staining in ***so*** and ***sr*** of area CA1 and the **ml** of the dentate gyrus indicates that tau levels are increased in hippocampal area CA1, but not DG in *Cyfip2*$^{+/-}$ mice. Importantly, this change is much more restricted than the elevation of tau phosphorylation at AT8 and PHF-1 sites in these mutants (see, Fig. 3a, b). Data represent mean ± SEM. n=3-4 animals per condition (individual animals shown as data points). * $p < 0.05$, ** $p < 0.01$. **gcl** granular cell layer **ml** molecular layer **sl** *stratum lucidum* **so** *stratum oriens* **sp** *stratum pyramidale* **sr** *stratum radiatum*.

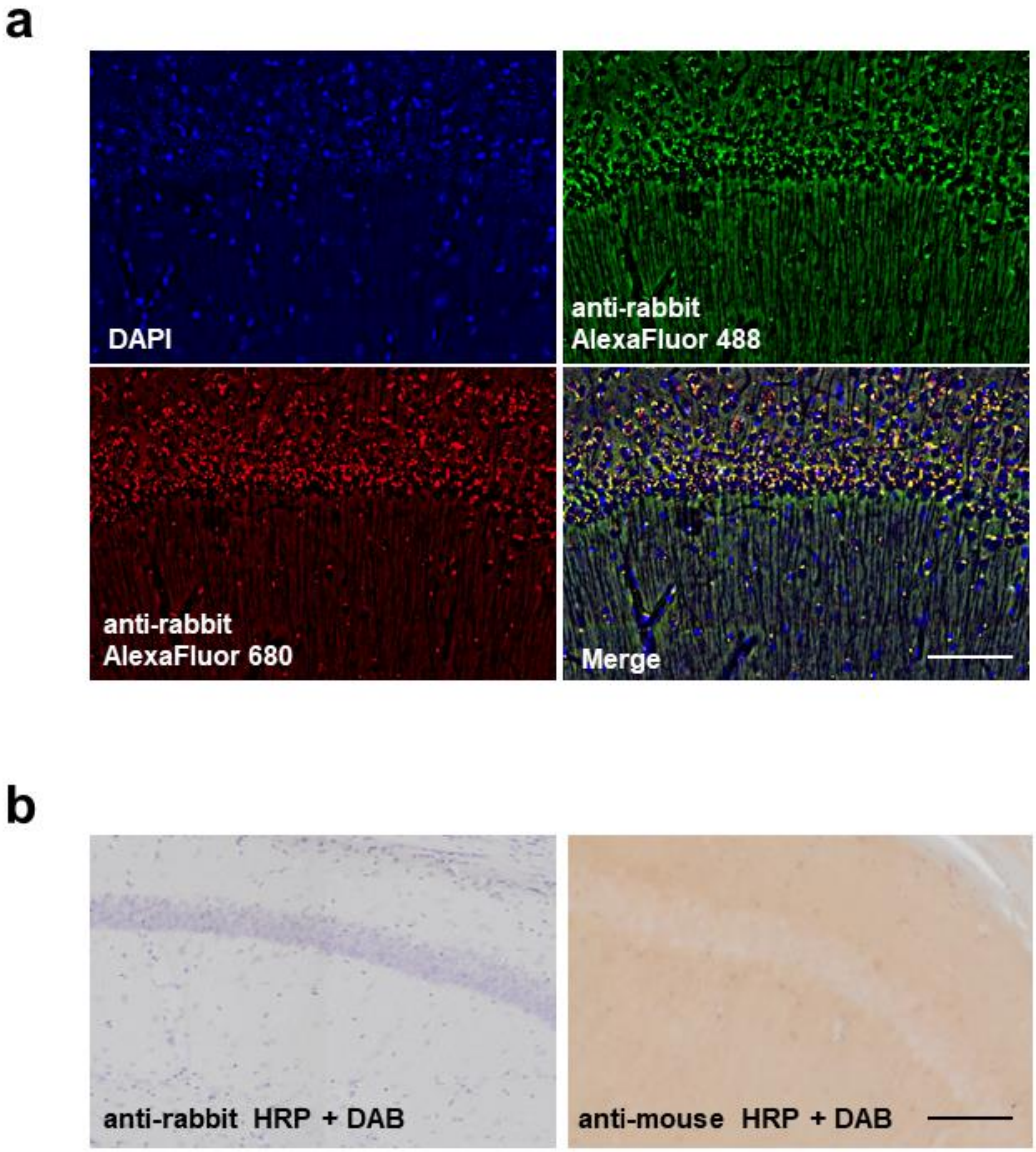


**Supplementary Figure 10. Staining detected in aged tissues is specific and not due to background or age-related protein build-up. a** Images of area CA1 of dorsal hippocampus from 12-month-old wild-type mice treated only with relevant fluorescent secondary antibodies and DAPI, and no primary antibodies. **b** Images of area CA1 of dorsal hippocampus from 12-month-old wild-type mice treated only with relevant HRP-tagged secondary antibodies and DAB, and counter-stained with haematoxylin, and no primary antibodies. Scale bars are 100 µm.

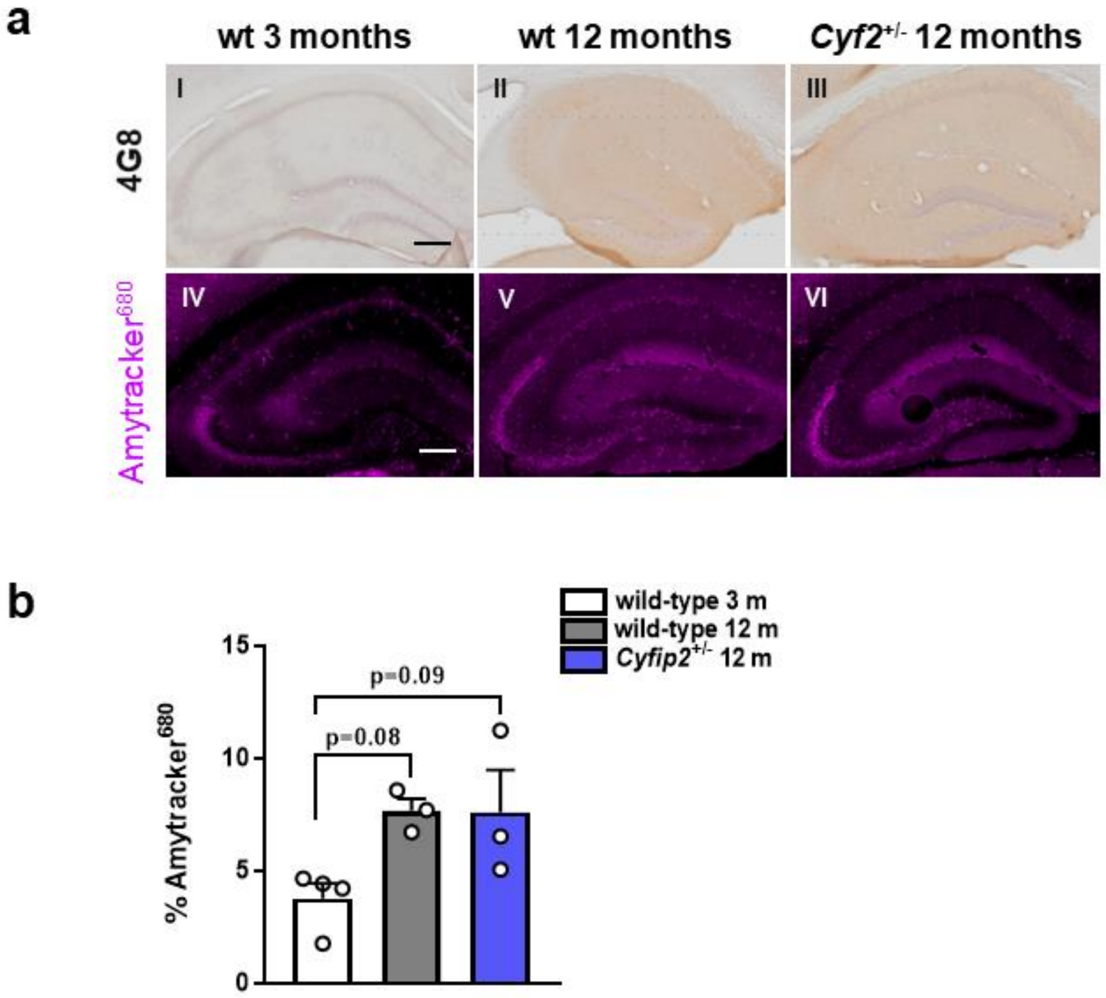


**Supplementary Figure 11. No changes in 4G8 or Amytracker$^{680}$ staining in the hippocampus of 12-month-old wild-type and *Cyfip2*$^{+/-}$ mice. a** Representative images of dorsal hippocampal sections from wild-type mice at 3 months and 12 months, and *Cyfip2*$^{+/-}$ mice at 12 months, immunohistochemically stained for anti-4G8 antibody to detect Aβ (**I-III**) and Amytracker$^{680}$ (**IV-VI**). Nuclei were labelled with haemotoxylin or DAPI, respectively. Scale bars are 200 µm. **b** Quantification of Amytracker$^{680}$ staining, expressed as % of area sampled does not reveal a difference between aged genotypes. No 4G8-positive staining was detected across groups in the hippocampus, although aged tissues have higher background staining. Data represent mean ± SEM. n=3-4 animals per condition (individual animals shown as white dots).

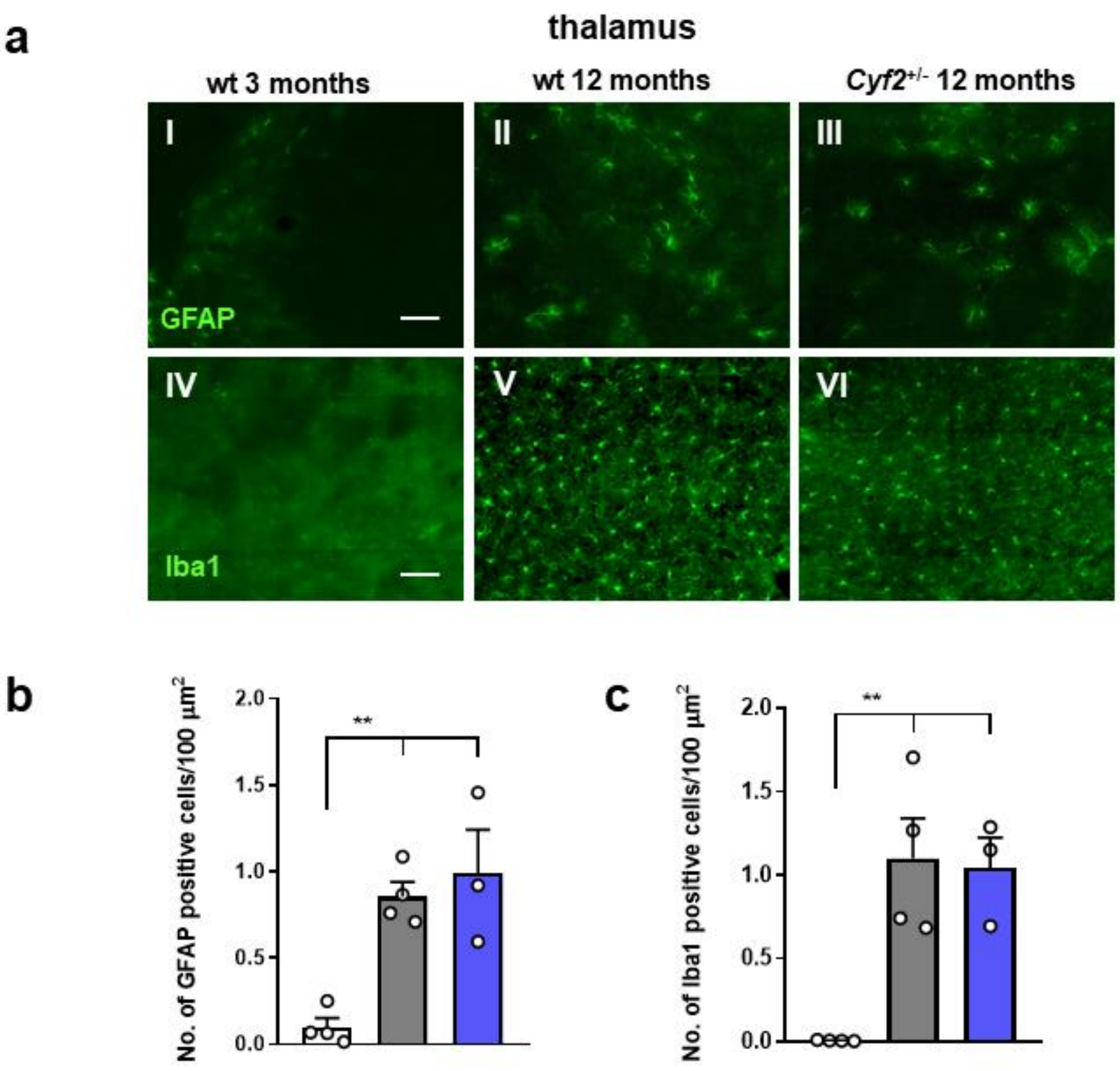


**Supplementary Figure 12. No evidence for thalamic gliosis in *Cyfip2*$^{+/-}$ mice. a** Representative images of thalamic regions from wild-type mice at 3 months and 12 months, and *Cyfip2*$^{+/-}$ mice at 12 months, immunohistochemically stained for GFAP (**I-III**) or Iba1 (**IV-VI**). Scale bars are 100 µm. **b** and **c** Quantification of number of GFAP-positive cells (**b**) and number of Iba1-positive microglia (**c**) in the mouse thalamus does not detect any difference between the genotypes at 12 months of age. Data represent mean ± SEM. n=3-4 animals per condition (individual animals shown as white dots). * $p < 0.05$, ** $p < 0.01$, *** $p < 0.001$.

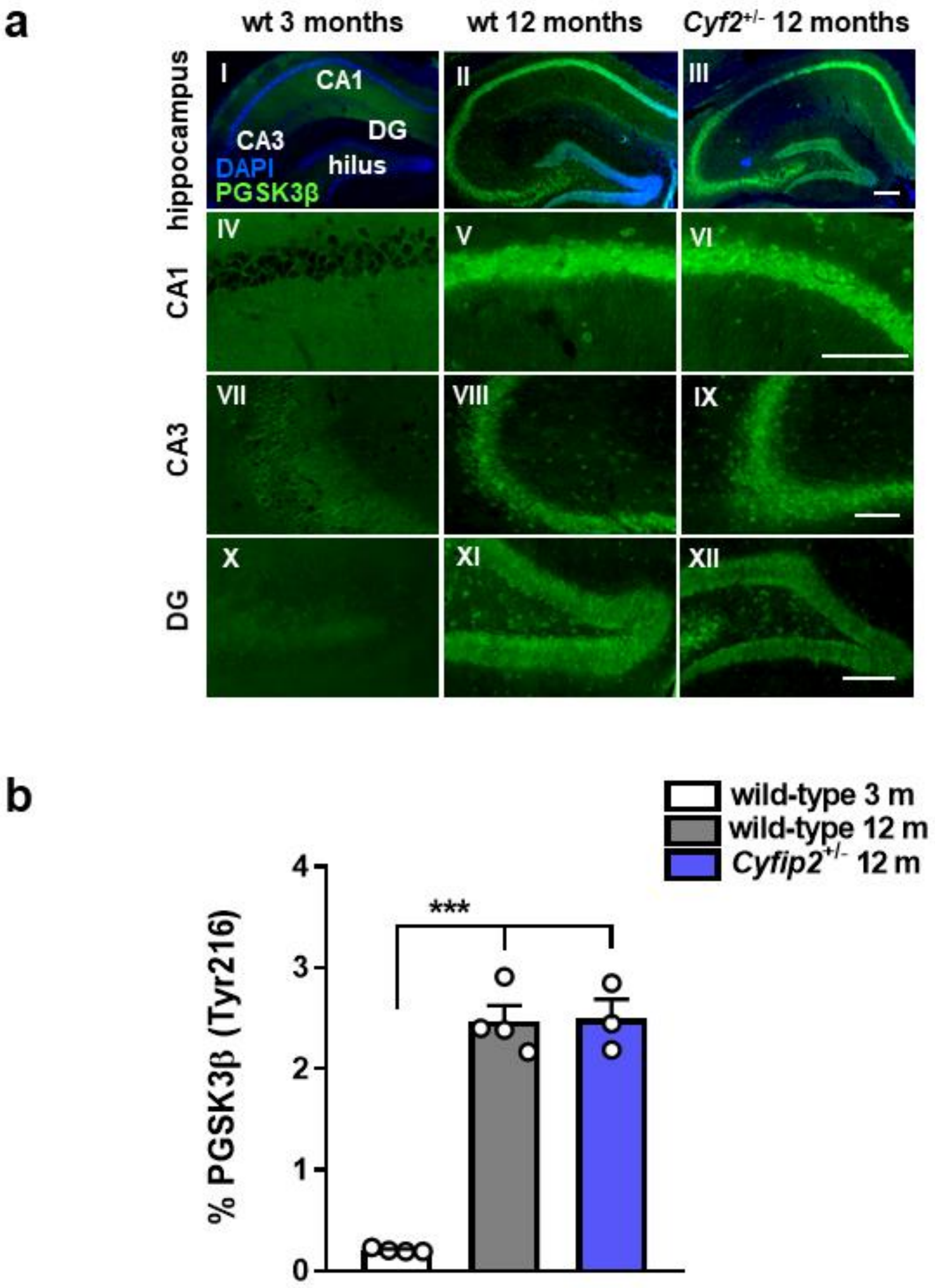


**Supplementary Figure 13. Phosphorylation of GSK3β at Tyr216 is increased in the mouse hippocampus with ageing. a** Representative images of dorsal hippocampal sections (**I-III**) from wild-type mice at 3 months and 12 months, and *Cyfip2*$^{+/-}$ mice at 12 months, stained for GSK3β phosphorylated at Tyr216. Higher magnification images of CA1 (**IV-VI**), CA3 (**VII-IX**) and DG (**X-XII**). Scale bars are 200 µm. **b** Quantification of phosphoGSK3β (Tyr216) across the entire dorsal hippocampus demonstrates that this phosphorylation event is increased with ageing. Data represent mean ± SEM. n=3-4 animals per condition (individual animals shown as white dots). *** $p < 0.001$.

**Supplementary Tables 1-3 provided as Excel files.**

**Supplementary Table 1.** RNA sequencing information of samples used for analysis of total and ribosomal RNAs following A$\beta_{1-42}$ treatment.

**Supplementary Table 2.** Log2 fold changes and corresponding p-values for transcripts showing statistically significant changes in expression between total and ribosomal fractions, under either control conditions or following A$\beta_{1-42}$ exposure. The first two tabs display non-FMRP-binding targets, and the second two tabs display FMRP-binding targets.

**Supplementary Table 3.** Pathway analysis of genes differentially regulated in ribosomes under control conditions and following A$\beta_{1-42}$ exposure.

**Supplementary Table 4.** Table of SNPs in five genes of interest following exclusion of SNPs in high linkage disequilibrium (LD).

| **Gene** | **Chr** | **Start*** | **End*** | **N SNPs before pruning** | **N SNPs after pruning** | **Best SNP** | **Best SNP position** | **Best SNP type** | **Best SNP P-Value** | **Best SNP Q-Value**** |
|---|---|---|---|---|---|---|---|---|---|---|
| *MKNK1* | 1 | *4023090* | *47082515* | 205 | 85 | rs1258047 | 47019254 T>C | MKNK1-AS1: Intronic KNCN:2KB Upstream Variant | 0.0001 | 0.075 |
| *MKNK2* | 19 | *2037470* | *2051243* | 144 | 89 | rs60379186 | 2058122A>G | Upstream MNK2 | 0.024 | 0.705 |
| *CYFIP1* | 15 | 22892667 | 23003603 | 675 | 308 | rs190772561 | 22921657 C>T | Intronic | 0.006 | 0.705 |
| *CYFIP2* | 5 | *156693091* | *156822606* | 593 | 201 | rs183249859 | 156750164G>C | Intronic | 0.020 | 0.704 |
| *EIF4E* | 4 | *99799607* | *99851786* | 253 | 80 | rs145615497 | 99817301 C>G | Intronic | 0.025 | 0.707 |

*Gene coding region (GRCh37/hg19).

**Q-value was obtained using the FDR tools package in R.

| Sample name | Raw reads | Clean reads | raw bases | clean bases | Error rate(%) | Q20(%) | Q30(%) |
|---|---|---|---|---|---|---|---|
| A1AbRi | 43188656 | 42404378 | 6.5G | 6.4G | 0.03 | 97.63 | 93.48 |
| A5AbRi | 56594866 | 55553840 | 8.5G | 8.3G | 0.03 | 97.93 | 94.15 |
| A9AbRi | 46311014 | 45196054 | 6.9G | 6.8G | 0.02 | 98.17 | 94.75 |
| A2CRi | 44472562 | 42476518 | 6.7G | 6.4G | 0.03 | 97.51 | 93.25 |
| A6CRi | 51063824 | 49814428 | 7.7G | 7.5G | 0.03 | 97.79 | 93.83 |
| A10CRi | 45833744 | 44052390 | 6.9G | 6.6G | 0.03 | 97.56 | 93.33 |
| A3AbTo | 54859076 | 53346420 | 8.2G | 8G | 0.03 | 97.59 | 93.54 |
| A7AbTo | 51356448 | 49390724 | 7.7G | 7.4G | 0.03 | 97.35 | 92.83 |
| A11AbTo | 51347978 | 50539652 | 7.7G | 7.6G | 0.02 | 98.15 | 94.8 |
| A4CTo | 49619414 | 48626636 | 7.4G | 7.3G | 0.02 | 97.96 | 94.34 |
| A8CTo | 51821142 | 50861920 | 7.8G | 7.6G | 0.02 | 98.03 | 94.5 |
| A12CTo | 46009708 | 44722012 | 6.9G | 6.7G | 0.03 | 97.86 | 94.15 |

| GC content(%) |
| --- |
| 48.79 |
| 47.12 |
| 48.09 |
| 49.58 |
| 46.91 |
| 48.01 |
| 50.24 |
| 50.46 |
| 50.79 |
| 50.59 |
| 50.44 |
| 50.74 |

| Associated_Gene_Name | log2FoldChange | padj |
|---|---|---|
| Aatk | -0.67017 | 0.0021168 |
| Abca2 | -1.0418 | 3.70E-07 |
| Abca3 | -2.2234 | 4.17E-11 |
| Acly | -0.98624 | 0.0043427 |
| Adarb1 | -0.5686 | 0.011143 |
| Adcy2 | 0.78638 | 3.03E-04 |
| Adcy5 | -0.92034 | 1.32E-05 |
| Aff1 | -0.93577 | 0.0063764 |
| Aff3 | -1.3219 | 1.85E-09 |
| Agap1 | -1.4526 | 1.66E-12 |
| Agap2 | -0.95686 | 1.89E-06 |
| Agap3 | -0.51668 | 0.022784 |
| Agrn | -0.89731 | 1.67E-05 |
| Ahdc1 | -0.87689 | 6.94E-05 |
| Akap6 | -1.3355 | 6.40E-11 |
| Akt3 | -0.96431 | 7.74E-06 |
| Ank1 | -1.1466 | 1.59E-06 |
| Ank2 | -1.9127 | 2.40E-22 |
| Ankrd11 | -1.3335 | 4.31E-11 |
| Ankrd52 | -2.3632 | 7.06E-28 |
| Ap2a1 | -0.87764 | 2.06E-05 |
| Apba1 | -1.8335 | 2.58E-20 |
| Apc | -0.93323 | 3.42E-05 |
| Apc2 | -1.3077 | 8.64E-11 |
| Arhgap21 | -0.62592 | 0.0045746 |
| Arhgap23 | -1.1459 | 2.63E-08 |
| Arhgap32 | -1.3427 | 7.71E-11 |
| Arhgap33 | -1.4064 | 1.31E-12 |
| Arhgdia | -0.47794 | 0.035101 |
| Arhgef10l | -0.53925 | 0.022818 |
| Arhgef11 | -0.68301 | 0.0026589 |
| Arhgef12 | -1.3262 | 2.14E-10 |
| Arhgef7 | -0.65137 | 0.0037125 |
| Arid1a | -0.77176 | 4.30E-04 |
| Arid2 | -0.95049 | 4.34E-05 |
| Arl3 | 0.90946 | 3.24E-05 |
| Arnt2 | -0.71747 | 7.20E-04 |
| Arrb1 | -1.8673 | 1.87E-18 |
| Ash1l | -0.69289 | 0.0018339 |
| Atf4 | 0.61611 | 0.0044573 |

| | | |
|---|---|---|
| Atf7ip | -1.2121 | 9.82E-08 |
| Atg2b | -0.58531 | 0.0099462 |
| Atp13a2 | -0.79279 | 1.36E-04 |
| Atp2b4 | -1.4491 | 6.57E-05 |
| Atp6v1f | 0.85592 | 3.61E-04 |
| Atxn1 | -1.3158 | 6.57E-09 |
| Auts2 | -1.0926 | 7.42E-04 |
| B3gat1 | -0.76948 | 7.26E-04 |
| Bcl9l | -1.8031 | 6.71E-19 |
| Bcr | -1.1364 | 4.40E-07 |
| Birc6 | -1.394 | 1.70E-11 |
| Bmpr2 | -1.6396 | 3.44E-15 |
| Brd4 | -0.94049 | 1.05E-05 |
| Brsk2 | -1.2788 | 1.12E-09 |
| Bsn | -1.5025 | 8.14E-12 |
| Bzrap1 | -1.1299 | 4.90E-08 |
| C2cd2l | -0.74708 | 5.42E-04 |
| Cabin1 | -0.94661 | 1.19E-05 |
| Cacna1b | -0.87865 | 3.41E-05 |
| Cacna1d | -0.81378 | 3.25E-04 |
| Cacna1e | -2.7534 | 2.87E-40 |
| Cacnb1 | -0.99997 | 2.22E-06 |
| Cadps | -0.82759 | 9.20E-05 |
| Calr | 0.64405 | 0.0037636 |
| Camk2a | -1.0634 | 1.41E-07 |
| Camkk2 | -0.67534 | 0.007147 |
| Camta2 | -0.90579 | 1.26E-05 |
| Caskin1 | -1.0502 | 2.25E-07 |
| Cbx3 | 0.57006 | 0.010058 |
| Cbx6 | -0.78452 | 1.89E-04 |
| Cdc42bpa | -1.041 | 5.03E-07 |
| Cdc42bpb | -0.97923 | 1.37E-06 |
| Cdk5r1 | -0.92705 | 1.61E-05 |
| Cdk5r2 | -0.50847 | 0.028972 |
| Cdk6 | -2.5638 | 0.017883 |
| Cdkl5 | -1.5869 | 2.26E-08 |
| Celf5 | -0.82627 | 6.24E-05 |
| Celsr2 | -2.239 | 3.15E-29 |
| Chd3 | -0.82969 | 4.13E-05 |
| Chd4 | -0.99368 | 1.55E-06 |
| Chd5 | -1.729 | 6.35E-18 |

| | | |
|---|---|---|
| Chn1 | 0.7653 | 0.0054018 |
| Chst2 | -0.69153 | 0.0014902 |
| Cit | -1.5848 | 1.41E-13 |
| Clip3 | -1.3544 | 6.09E-11 |
| Cplx1 | -0.51337 | 0.030112 |
| Cplx2 | -1.2351 | 2.53E-08 |
| Crebbp | -0.63776 | 0.012732 |
| Crtc1 | -1.3766 | 3.81E-10 |
| Ctnnd2 | -1.0625 | 1.61E-07 |
| Cul9 | -0.67765 | 0.0043783 |
| Dag1 | -0.87375 | 0.0046204 |
| Dagla | -0.95534 | 6.22E-06 |
| Dctn1 | -0.54838 | 0.012127 |
| Dgkz | -0.68309 | 0.0014578 |
| Dhx30 | -0.49465 | 0.032604 |
| Dicer1 | -0.70786 | 0.0026742 |
| Dido1 | -0.73211 | 0.0017893 |
| Dip2a | -1.125 | 2.17E-07 |
| Dip2c | -0.9305 | 1.34E-05 |
| Diras2 | -1.2774 | 1.60E-04 |
| Dlg2 | -0.9661 | 4.88E-06 |
| Dlg4 | -0.60477 | 0.0053271 |
| Dlg5 | -1.2692 | 4.15E-09 |
| Dlgap1 | -0.76346 | 3.27E-04 |
| Dlgap2 | -1.3924 | 3.85E-06 |
| Dlgap3 | -0.76711 | 2.54E-04 |
| Dlgap4 | -0.84152 | 6.66E-05 |
| Dmwd | -0.58385 | 0.010331 |
| Dnm1 | -0.59035 | 0.0052438 |
| Dock3 | -1.1321 | 5.01E-08 |
| Dock4 | -1.4618 | 1.77E-12 |
| Dopey2 | -0.61327 | 0.0097814 |
| Dot1l | -1.2533 | 1.97E-08 |
| Dscam | -1.0115 | 5.82E-06 |
| Dscaml1 | -1.1482 | 1.82E-07 |
| Dst | -1.7258 | 2.10E-18 |
| Dtx1 | -0.63225 | 0.0064604 |
| Dusp8 | -1.0761 | 3.84E-07 |
| Dync1h1 | -1.2072 | 2.06E-09 |
| Dynll1 | 0.65333 | 0.0024668 |
| Eef1a1 | 0.69538 | 0.0042707 |

| | | |
|---|---|---|
| Eef1a2 | -0.61213 | 0.004164 |
| Ehmt1 | -0.67732 | 0.0031201 |
| Ehmt2 | -0.67733 | 0.0017677 |
| Eif4g1 | -1.0726 | 1.74E-06 |
| Eif4g3 | -0.57169 | 0.010472 |
| Elfn2 | -1.7479 | 6.37E-17 |
| Elmo2 | -0.56716 | 0.014629 |
| Eml2 | -0.52897 | 0.020154 |
| Ep300 | -1.9351 | 2.88E-10 |
| Ep400 | -0.70755 | 0.0015818 |
| Epn1 | -0.51582 | 0.020769 |
| Extl3 | -0.97665 | 6.97E-06 |
| Fam120a | -0.5381 | 0.016734 |
| Fam160a2 | -0.66033 | 0.0041654 |
| Fam171b | -0.50224 | 0.033221 |
| Fam65a | -0.65296 | 0.0033813 |
| Fasn | -0.89668 | 0.001937 |
| Fat1 | -0.99515 | 8.60E-04 |
| Fat3 | -1.2594 | 2.74E-05 |
| Fbxl16 | -1.0776 | 1.15E-07 |
| Fbxl19 | -0.63988 | 0.0041881 |
| Fbxo41 | -1.5141 | 6.06E-13 |
| Fcho1 | -1.0128 | 6.01E-04 |
| Fibp | 0.50219 | 0.046282 |
| Fkbp3 | 0.87771 | 3.27E-04 |
| Foxk2 | -0.48122 | 0.043828 |
| Foxo3 | -1.1771 | 1.19E-06 |
| Frmpd4 | -0.56597 | 0.020176 |
| Fry | -0.51072 | 0.027807 |
| Gabbr2 | -0.64468 | 0.0030903 |
| Gabrd | 0.96976 | 0.0069565 |
| Gas7 | -0.46181 | 0.047086 |
| Gcn1l1 | -0.75609 | 4.78E-04 |
| Gnaz | -0.75453 | 6.43E-04 |
| Golga5 | 0.57087 | 0.032054 |
| Gpam | -1.3099 | 2.81E-04 |
| Gpm6a | 0.98656 | 9.58E-06 |
| Gria1 | -0.76179 | 2.39E-04 |
| Grik3 | -2.0214 | 1.32E-20 |
| Grik5 | -0.82249 | 6.46E-05 |
| Grin2a | -2.5045 | 9.32E-28 |

| | | |
|---|---|---|
| Grin2b | -3.0017 | 1.39E-43 |
| Grm5 | -1.3362 | 8.66E-11 |
| Gsk3b | -1.6193 | 1.88E-14 |
| Gtf3c1 | -0.63485 | 0.0045385 |
| Hcn2 | -0.73194 | 0.001376 |
| Hdac4 | -1.8046 | 1.64E-16 |
| Hdlbp | -0.64631 | 0.0028549 |
| Herc2 | -0.70904 | 9.05E-04 |
| Hint1 | 0.80327 | 1.09E-04 |
| Hipk2 | -2.9621 | 4.23E-34 |
| Hipk3 | -1.1293 | 1.38E-06 |
| Hivep3 | -1.9023 | 2.15E-17 |
| Hnrnpul1 | -0.53609 | 0.01456 |
| Htt | -0.94068 | 1.23E-05 |
| Igsf9b | -3.621 | 2.53E-45 |
| Inpp4a | -1.316 | 1.15E-10 |
| Ints1 | -0.54664 | 0.025304 |
| Ipo4 | -0.61932 | 0.0064156 |
| Iqsec3 | -0.77741 | 2.20E-04 |
| Itsn1 | -1.1609 | 5.69E-04 |
| Jph4 | -1.0209 | 3.52E-07 |
| Kalrn | -0.57744 | 0.012087 |
| Kcna2 | -2.4873 | 2.30E-24 |
| Kcnb1 | -1.151 | 1.71E-07 |
| Kcnc3 | -1.2343 | 1.86E-09 |
| Kcnh1 | -0.61112 | 0.014424 |
| Kcnh7 | -0.90566 | 0.041808 |
| Kcnma1 | -1.9241 | 3.86E-16 |
| Kcnq2 | -1.8065 | 5.06E-19 |
| Kcnq3 | -1.9761 | 1.30E-04 |
| Kcnt1 | -1.3299 | 9.90E-08 |
| Kdm4b | -0.60003 | 0.0077174 |
| Kdm6b | -1.3377 | 8.72E-09 |
| Kif1a | -1.7308 | 8.06E-19 |
| Kif21b | -2.9013 | 2.70E-42 |
| Kif3c | -0.64802 | 0.0026585 |
| Kif5c | -1.0369 | 2.83E-07 |
| Kndc1 | -1.2549 | 9.22E-09 |
| Lhfpl4 | -1.4694 | 3.85E-11 |
| Lmtk2 | -0.57839 | 0.010574 |
| Lrp1 | -1.0987 | 3.75E-08 |

| | | |
|---|---|---|
| Lrp3 | -0.56259 | 0.013156 |
| Lrp8 | -1.4882 | 1.30E-11 |
| Lrrc4b | -0.54347 | 0.011958 |
| Lrrc7 | -1.7407 | 5.93E-16 |
| Lrrc8b | -1.1852 | 1.15E-05 |
| Lrrn2 | -0.48723 | 0.032178 |
| Lynx1 | -0.5892 | 0.010956 |
| Macf1 | -1.3672 | 8.11E-12 |
| Magi2 | -0.87944 | 4.42E-05 |
| Man2a2 | -0.85787 | 1.18E-04 |
| Map1a | -0.95415 | 8.62E-06 |
| Map1b | -0.58494 | 0.0075749 |
| Map2 | -1.0283 | 2.81E-07 |
| Map7d1 | -0.80495 | 1.38E-04 |
| Mapk8ip3 | -0.81988 | 8.19E-05 |
| Mapkbp1 | -1.1068 | 6.60E-07 |
| Med13l | -1.3201 | 6.57E-10 |
| Med16 | -0.57274 | 0.013072 |
| Mef2d | -1.364 | 1.23E-10 |
| Mfhas1 | -1.3419 | 4.65E-09 |
| Mib1 | -1.3634 | 6.08E-11 |
| Mical2 | -0.55125 | 0.012995 |
| Mink1 | -0.77681 | 1.92E-04 |
| Mprip | -0.52045 | 0.021316 |
| Mtss1l | -1.4121 | 4.69E-11 |
| Myo16 | -0.77577 | 3.80E-04 |
| Myo18a | -0.81636 | 1.24E-04 |
| Myt1l | -1.0231 | 7.13E-07 |
| Nat8l | -0.71318 | 9.40E-04 |
| Nav1 | -2.1793 | 1.56E-26 |
| Nav2 | -1.1196 | 1.71E-07 |
| Ncam1 | -0.82865 | 4.48E-05 |
| Ncan | -2.4589 | 7.79E-10 |
| Ncdn | -0.81 | 9.01E-05 |
| Ncoa2 | -1.4271 | 2.81E-11 |
| Ncor2 | -1.1818 | 7.95E-09 |
| Ncs1 | -1.0721 | 3.13E-07 |
| Ndst1 | -1.8841 | 9.06E-19 |
| Neurl4 | -0.6836 | 0.0015911 |
| Nf1 | -0.6316 | 0.0052869 |
| Nfic | -1.7213 | 2.34E-10 |

| | | |
|---|---|---|
| Nfix | -1.8945 | 6.33E-20 |
| Nfya | -0.58019 | 0.042418 |
| Nfyb | 0.73987 | 0.01142 |
| Ngrn | 0.58691 | 0.014757 |
| Nisch | -0.49169 | 0.024746 |
| Nlgn2 | -1.423 | 1.43E-11 |
| Nlgn3 | -0.96335 | 3.26E-06 |
| Nos1 | -1.995 | 2.30E-18 |
| Nptxr | -0.74287 | 0.0019001 |
| Nr3c1 | 0.85731 | 1.15E-04 |
| Nr3c1 | -1.0045 | 8.07E-04 |
| Nrxn2 | -0.95733 | 1.89E-06 |
| Nrxn3 | 0.74231 | 0.014777 |
| Nsd1 | -1.2199 | 5.14E-09 |
| Ntrk2 | -1.0636 | 4.99E-06 |
| Ntrk3 | -1.2243 | 1.97E-08 |
| Ogdh | -0.5316 | 0.021759 |
| Pacs1 | -0.63416 | 0.0042079 |
| Pacs2 | -1.311 | 3.09E-10 |
| Pak6 | -0.99128 | 0.0027883 |
| Pcdh10 | -0.63431 | 0.0046259 |
| Pcdh7 | -1.1577 | 9.59E-08 |
| Pcdh9 | -0.75465 | 0.0012088 |
| Pclo | -0.77651 | 3.71E-04 |
| Pcnx | -0.98624 | 1.50E-06 |
| Pde4dip | -0.74236 | 4.42E-04 |
| Pdzd2 | -1.8205 | 1.81E-04 |
| Peg3 | -1.4539 | 1.80E-09 |
| Phactr1 | -0.99763 | 5.06E-06 |
| Pik3r2 | -0.7021 | 0.0011084 |
| Pikfyve | -0.80907 | 3.33E-04 |
| Pip5k1b | 0.73522 | 0.036945 |
| Pip5k1c | -0.78529 | 1.49E-04 |
| Pitpnm2 | -0.62802 | 0.0045822 |
| Pkd1 | -1.2851 | 4.90E-10 |
| Plcb1 | -0.76171 | 5.42E-04 |
| Plch2 | -1.0786 | 2.24E-05 |
| Plec | -1.3642 | 9.74E-04 |
| Plxna1 | -2.0591 | 9.16E-24 |
| Plxna2 | -0.73115 | 7.78E-04 |
| Plxna4 | -2.2788 | 4.95E-27 |

| | | |
|---|---|---|
| Polr2a | -1.563 | 1.81E-14 |
| Ppargc1a | -0.62039 | 0.03263 |
| Ppfia3 | -0.81791 | 1.03E-04 |
| Ppm1e | -1.5374 | 4.47E-08 |
| Prex1 | -1.1198 | 8.21E-08 |
| Prkce | -1.3704 | 8.83E-12 |
| Ptch1 | -1.1299 | 2.50E-05 |
| Pten | -0.69504 | 0.0029352 |
| Ptk2b | -0.5423 | 0.015307 |
| Ptma | 0.57562 | 0.0085591 |
| Ptpn11 | -0.49068 | 0.029297 |
| Ptpn5 | -0.57782 | 0.010845 |
| Ptprd | -0.51062 | 0.041183 |
| Ptprf | -0.98126 | 1.52E-06 |
| Ptprg | -1.0562 | 5.23E-07 |
| Ptprj | -0.7749 | 6.47E-04 |
| Ptprs | -0.75298 | 4.01E-04 |
| Ptprt | -2.0294 | 1.83E-16 |
| R3hdm1 | -0.55515 | 0.013634 |
| Ralgapb | -0.95047 | 1.18E-05 |
| Ralgds | -0.63193 | 0.0047132 |
| Rap1gap | -0.84115 | 6.31E-05 |
| Rapgef1 | -1.2366 | 3.69E-09 |
| Rapgefl1 | -0.57396 | 0.010455 |
| Rc3h1 | -1.6894 | 2.16E-12 |
| Rc3h2 | -0.9278 | 3.25E-04 |
| Rere | -0.80968 | 1.61E-04 |
| Rgs7bp | -1.166 | 1.30E-06 |
| Rhob | -0.72012 | 0.009353 |
| Rhobtb2 | -0.54103 | 0.026418 |
| Rimbp2 | -0.56281 | 0.012819 |
| Rpl13a | 1.028 | 3.84E-07 |
| Rpsa | 0.96974 | 4.50E-04 |
| Rtn3 | 0.46792 | 0.034313 |
| Rusc2 | -0.60713 | 0.0058384 |
| Ryr2 | -0.91848 | 2.36E-05 |
| Samd4b | -0.57586 | 0.014709 |
| Sap130 | -1.354 | 2.85E-05 |
| Sash1 | -0.96115 | 0.010114 |
| Sbf1 | -1.0975 | 3.89E-08 |
| Scn8a | -1.7922 | 1.02E-18 |

| | | |
|---|---|---|
| Sec16a | -1.0208 | 9.58E-06 |
| Secisbp2l | -0.48912 | 0.038438 |
| Setd5 | -0.57099 | 0.010754 |
| Setx | -0.64704 | 0.0060683 |
| Sgip1 | -1.3941 | 9.43E-08 |
| Sgsm2 | -0.55143 | 0.015107 |
| Shank1 | -2.0452 | 1.28E-25 |
| Shank2 | -2.1579 | 1.28E-14 |
| Sipa1l1 | -0.72779 | 9.14E-04 |
| Sipa1l3 | -0.98295 | 4.41E-05 |
| Slc12a5 | -1.0798 | 2.06E-07 |
| Slc17a7 | -0.57677 | 0.0091917 |
| Slc1a2 | -1.4044 | 0.0072787 |
| Slc24a2 | -1.8025 | 2.78E-10 |
| Slc4a3 | -0.94032 | 1.67E-05 |
| Slc4a8 | -2.6355 | 2.21E-29 |
| Slc8a1 | -1.9382 | 2.15E-20 |
| Slc8a2 | -0.72467 | 0.0010711 |
| Smarca4 | -0.45845 | 0.043496 |
| Smarcc2 | -0.72819 | 6.09E-04 |
| Smg1 | -1.005 | 2.95E-06 |
| Snap91 | -0.80345 | 1.10E-04 |
| Snph | -1.0041 | 8.16E-07 |
| Snrk | -0.81342 | 2.89E-04 |
| Sobp | -1.2284 | 3.07E-09 |
| Sod1 | 0.98921 | 7.26E-05 |
| Sort1 | -1.3176 | 1.11E-09 |
| Sparcl1 | 0.65037 | 0.038384 |
| Speg | -0.602 | 0.0077954 |
| Spen | -2.3178 | 1.90E-28 |
| Spred1 | -0.96556 | 1.03E-04 |
| Sptan1 | -0.61689 | 0.0044105 |
| Sptb | -2.3539 | 8.23E-28 |
| Sptbn1 | -1.5336 | 6.12E-15 |
| Srcin1 | -1.5768 | 2.03E-15 |
| Srgap3 | -1.8117 | 8.38E-19 |
| Srpk1 | -0.78594 | 8.93E-04 |
| Stox2 | -1.6454 | 7.60E-16 |
| Stxbp1 | -0.62837 | 0.0031446 |
| Stxbp5 | -0.84245 | 1.14E-04 |
| Sympk | -0.62801 | 0.0046243 |

| | | |
|---|---|---|
| Syn1 | -0.82349 | 7.39E-05 |
| Syngap1 | -1.1855 | 2.35E-09 |
| Syngr1 | -0.90408 | 1.67E-05 |
| Synj1 | -0.80878 | 1.36E-04 |
| Synpo | -0.8391 | 2.43E-04 |
| Syt7 | -1.5324 | 2.71E-13 |
| Taok1 | -1.6471 | 8.86E-15 |
| Taok2 | -0.65966 | 0.0021703 |
| Tcf4 | -0.56378 | 0.017833 |
| Tln2 | -1.7211 | 9.04E-13 |
| Tmem132a | -0.51599 | 0.027158 |
| Tmem63b | -0.62971 | 0.0041659 |
| Tmem8b | -0.67523 | 0.0046913 |
| Tmod2 | -0.99374 | 3.08E-06 |
| Tnik | -1.1773 | 8.60E-09 |
| Tnk2 | -1.138 | 4.98E-08 |
| Tnks | -1.4474 | 1.54E-12 |
| Tnrc18 | -2.0028 | 3.40E-24 |
| Tnrc6b | -0.56541 | 0.013414 |
| Tppp | -1.0097 | 2.30E-06 |
| Trak2 | -0.56984 | 0.011523 |
| Trim2 | -1.3537 | 2.91E-11 |
| Trim9 | -0.5015 | 0.036842 |
| Trio | -1.2184 | 2.83E-09 |
| Trrap | -1.0195 | 9.29E-07 |
| Tsc2 | -0.51866 | 0.028985 |
| Tspan7 | 0.5287 | 0.015388 |
| Ttbk1 | -1.9297 | 7.58E-16 |
| Ttbk2 | -1.5599 | 1.18E-10 |
| Ttll7 | -0.59847 | 0.01327 |
| Tulp4 | -1.6169 | 2.07E-15 |
| Ubap2l | -0.54151 | 0.014626 |
| Ube2a | 0.65146 | 0.0042042 |
| Ube2o | -0.80162 | 1.83E-04 |
| Ube3b | -1.1423 | 4.55E-07 |
| Ulk1 | -0.48813 | 0.041893 |
| Ulk2 | -0.68354 | 0.0031965 |
| Unc13a | -1.3935 | 4.74E-08 |
| Unc13b | -1.0828 | 9.83E-04 |
| Usp22 | -1.2057 | 2.24E-09 |
| Wdr13 | -0.84046 | 9.55E-05 |

| | | |
|---|---|---|
| Wdr6 | -0.62426 | 0.006442 |
| Wnk2 | -1.3199 | 7.67E-11 |
| Xpo6 | -0.53751 | 0.018653 |
| Zc3h4 | -1.5004 | 3.22E-12 |
| Zcchc14 | -1.5949 | 9.26E-13 |
| Zfhx2 | -1.6254 | 8.84E-05 |
| Zfpm1 | -1.0715 | 0.0014492 |
| Zhx3 | -1.1039 | 3.63E-04 |
| Zmiz1 | -0.90259 | 1.17E-05 |
| Zyg11b | -1.3211 | 8.69E-10 |

| Associated_Gene_Name | log2FoldChange | padj |
|---|---|---|
| AABR07000156.1 | -3.4929 | 0.013201 |
| AABR07001099.1 | 0.67153 | 0.034874 |
| AABR07006458.1 | 0.84655 | 6.52E-04 |
| AABR07006627.1 | 0.683 | 0.011237 |
| AABR07007385.1 | 0.68578 | 0.007639 |
| AABR07008016.1 | 1.2105 | 0.0026971 |
| AABR07008066.2 | 1.1549 | 0.013013 |
| AABR07012054.1 | 1.4889 | 0.0025683 |
| AABR07016640.1 | 1.7664 | 0.017061 |
| AABR07016841.1 | 1.4878 | 0.0078401 |
| AABR07018124.1 | -3.7841 | 0.034538 |
| AABR07018226.1 | 1.119 | 2.79E-05 |
| AABR07019083.1 | -1.7011 | 2.70E-12 |
| AABR07019412.1 | -1.992 | 5.17E-11 |
| AABR07020786.1 | -0.97778 | 0.0020886 |
| AABR07020860.1 | -1.0674 | 0.027203 |
| AABR07026311.1 | 0.57403 | 0.011751 |
| AABR07029573.1 | 0.69488 | 0.032262 |
| AABR07030603.1 | -1.8261 | 2.27E-06 |
| AABR07030781.1 | 0.91388 | 1.02E-05 |
| AABR07031533.1 | -0.48613 | 0.034619 |
| AABR07031734.2 | -1.0629 | 0.0038083 |
| AABR07036087.1 | 1.0021 | 0.018666 |
| AABR07043823.2 | 1.2431 | 0.021018 |
| AABR07044825.1 | -0.72708 | 0.01491 |
| AABR07048211.1 | -2.2993 | 1.29E-08 |
| AABR07049948.1 | -2.3124 | 1.47E-04 |
| AABR07051947.1 | 0.99688 | 0.0061563 |
| AABR07053185.1 | 0.72341 | 0.012947 |
| AABR07053188.1 | -4.9777 | 0.0043775 |
| AABR07061755.1 | -1.1846 | 7.27E-05 |
| AABR07064349.1 | -1.6782 | 1.91E-07 |
| AABR07065406.1 | 1.0328 | 7.54E-05 |
| AABR07065923.1 | 1.9162 | 0.019748 |
| AABR07066510.1 | 1.1978 | 0.010173 |
| AABR07067499.1 | 1.282 | 0.0084738 |
| AABR07067736.1 | 1.5883 | 8.99E-04 |
| AABR07071445.1 | -2.2109 | 1.32E-09 |
| AABR07071876.1 | 0.81183 | 0.010085 |
| AABR07072537.1 | -1.0738 | 4.47E-04 |

| | | |
|---|---|---|
| Aamdc | 1.4696 | 7.63E-07 |
| Abat | -1.1575 | 5.16E-05 |
| Abca5 | -0.61597 | 0.014303 |
| Abca7 | -1.9329 | 6.12E-11 |
| Abcd3 | 0.5107 | 0.029904 |
| Abcg4 | -0.59768 | 0.011619 |
| Abhd14b | 1.0286 | 0.013464 |
| Abhd2 | -1.3424 | 1.86E-05 |
| Abhd3 | 1.1448 | 0.0051234 |
| Abl2 | -1.6975 | 1.67E-08 |
| Ablim2 | -0.56973 | 0.022387 |
| Ablim3 | -1.1962 | 2.97E-05 |
| Abt1 | 0.53893 | 0.04225 |
| AC095947.2 | 3.4048 | 0.046102 |
| AC096601.1 | -1.1409 | 0.022864 |
| AC109942.1 | 0.89569 | 0.030174 |
| AC115420.2 | -1.6853 | 0.036493 |
| AC117330.1 | -1.1035 | 0.018382 |
| AC126148.1 | -1.0647 | 0.0070875 |
| AC134160.1 | -1.8745 | 0.028247 |
| AC135400.1 | 1.4068 | 7.67E-06 |
| Acaa2 | 1.1793 | 1.54E-07 |
| Acaca | -1.0739 | 9.65E-06 |
| Acacb | -1.3975 | 0.046276 |
| Acadl | 0.89801 | 1.65E-04 |
| Acadm | 0.71286 | 0.009505 |
| Acap2 | -0.84598 | 1.89E-04 |
| Acap3 | -0.91887 | 1.04E-05 |
| Acd | -0.55334 | 0.035529 |
| Ackr3 | 0.96975 | 5.51E-04 |
| Acot13 | 1.0522 | 0.0020232 |
| Acot4 | 1.0685 | 0.019351 |
| Acot7 | -0.46542 | 0.048483 |
| Acot9 | 0.83444 | 2.05E-04 |
| Acox3 | -0.60034 | 0.022625 |
| Acp1 | 0.81404 | 1.66E-04 |
| Actn2 | -0.7263 | 0.010917 |
| Actn4 | -0.85293 | 5.72E-05 |
| Actr10 | 0.55679 | 0.013806 |
| Actr1b | -0.4812 | 0.038384 |
| Actr6 | 1.0516 | 3.76E-06 |

| | | |
|---|---|---|
| Acvr1b | -0.526 | 0.03103 |
| Acvr2b | -1.5677 | 1.10E-04 |
| Acyp2 | 0.77229 | 0.029033 |
| Adam11 | -0.9997 | 3.03E-06 |
| Adam12 | -2.6147 | 7.11E-08 |
| Adam15 | -0.6336 | 0.0051405 |
| Adam19 | -0.85582 | 1.34E-04 |
| Adam22 | -1.3322 | 1.25E-09 |
| Adam23 | -1.526 | 1.78E-13 |
| Adamts10 | -1.1544 | 4.57E-04 |
| Adamts17 | -1.4926 | 2.98E-06 |
| Adamts20 | -0.93738 | 0.035681 |
| Adamts3 | -0.93259 | 4.80E-04 |
| Adamts7 | -1.2666 | 0.0029608 |
| Adamtsl1 | -0.99513 | 0.0322 |
| Adamtsl5 | -1.9923 | 0.006045 |
| Adck3 | 0.70161 | 0.0053359 |
| Adcy9 | -1.6343 | 3.64E-12 |
| Adcyap1r1 | -1.2023 | 2.97E-05 |
| Add2 | -1.7448 | 9.73E-18 |
| Adgra2 | -1.0131 | 0.0063854 |
| Adgrb1 | -0.86515 | 2.55E-05 |
| Adgrb2 | -1.0263 | 2.47E-07 |
| Adgre1 | -2.1078 | 0.041603 |
| Adgrl1 | -1.4559 | 1.72E-13 |
| Adhfe1 | 1.2382 | 1.24E-06 |
| Adi1 | 0.81222 | 0.0023494 |
| Adipor2 | -0.67276 | 0.0038537 |
| Adora1 | -1.0003 | 3.76E-06 |
| Aen | -0.93809 | 7.56E-05 |
| Agbl3 | 0.97507 | 0.010482 |
| Agfg2 | -0.83545 | 0.024196 |
| Ago2 | -1.9533 | 2.61E-12 |
| Ago3 | -2.2383 | 1.23E-06 |
| Ago4 | -0.85482 | 0.014704 |
| Agpat1 | -0.50423 | 0.02845 |
| Agpat4 | 0.48313 | 0.048395 |
| Agps | -1.082 | 5.32E-05 |
| Ahcyl2 | -0.52222 | 0.035339 |
| Aif1 | 5.1204 | 0.026823 |
| Aimp1 | 1.0374 | 2.80E-05 |

| | | |
|---|---|---|
| Ak4 | -0.48687 | 0.041986 |
| Ak6 | 0.85214 | 0.002255 |
| Akap2 | -1.0845 | 1.56E-05 |
| Akap5 | -0.68266 | 0.014368 |
| Akap7 | 1.0124 | 1.30E-04 |
| Akip1 | 0.97191 | 2.42E-04 |
| Akna | -0.98687 | 0.021756 |
| Akt2 | -0.63632 | 0.0074263 |
| Aktip | 0.5205 | 0.025694 |
| Aldh1l2 | -1.565 | 0.043097 |
| Aldh5a1 | -0.6366 | 0.015347 |
| Aldh6a1 | 0.92543 | 2.04E-05 |
| Alg10 | -0.79672 | 0.0027141 |
| Alg12 | 0.67561 | 0.013086 |
| Alg8 | 0.73185 | 0.030617 |
| Alkbh1 | 0.61916 | 0.019493 |
| Alkbh2 | 0.75814 | 0.041105 |
| Alkbh5 | -0.82958 | 0.0025308 |
| Alkbh7 | 0.8387 | 0.041282 |
| Alkbh8 | -0.87694 | 0.0026479 |
| Alms1 | 0.5431 | 0.033093 |
| Alpk1 | -1.0433 | 0.0040805 |
| Amd1 | 0.63189 | 0.0045789 |
| Amer1 | -1.676 | 1.68E-07 |
| Amer2 | -1.2339 | 4.68E-07 |
| Amer3 | -0.57478 | 0.03108 |
| Amn1 | 0.99967 | 1.12E-05 |
| Amot | -0.85097 | 0.041183 |
| Amotl1 | -1.6065 | 1.24E-05 |
| Amz1 | -0.78023 | 0.0063852 |
| Anapc10 | 0.94415 | 2.53E-04 |
| Anapc11 | 0.89787 | 0.013121 |
| Anapc13 | 1.206 | 1.80E-07 |
| Anapc15 | 0.85473 | 4.58E-04 |
| Angel1 | -0.62759 | 0.020553 |
| Ankfy1 | -0.96093 | 0.0011759 |
| Ankle1 | -3.1151 | 5.04E-09 |
| Ankrd13a | -0.56256 | 0.024908 |
| Ankrd13b | -1.0662 | 2.66E-07 |
| Ankrd24 | 0.85209 | 0.006529 |
| Ankrd28 | -0.50534 | 0.03569 |

| | | |
|---|---|---|
| Ankrd33b | -1.1128 | 0.0033751 |
| Ankrd34a | -0.51853 | 0.031772 |
| Ankrd35 | -1.4183 | 0.023455 |
| Ankrd37 | 1.2042 | 0.022046 |
| Ankrd49 | 0.71037 | 0.0040381 |
| Ankrd50 | -0.68454 | 0.015224 |
| Ankrd53 | -0.75731 | 0.010058 |
| Anks1a | -0.57452 | 0.019318 |
| Anks1b | -0.7372 | 6.69E-04 |
| Anks6 | -0.66725 | 0.026462 |
| Anln | -1.1371 | 0.017809 |
| Ano6 | -0.74431 | 0.011938 |
| Antxr1 | -0.96903 | 2.83E-04 |
| Anxa5 | 0.70851 | 0.0040789 |
| Anxa7 | 0.49429 | 0.047213 |
| Aox1 | 0.89311 | 0.0083912 |
| Ap1m1 | -0.63906 | 0.0040477 |
| Ap1s3 | 1.0925 | 0.0028004 |
| Ap3s1 | 0.86908 | 2.39E-04 |
| Ap4s1 | 0.78113 | 0.001734 |
| Ap5z1 | -1.3013 | 1.06E-04 |
| Apaf1 | -1.1564 | 1.10E-06 |
| Apba2 | -0.77645 | 1.54E-04 |
| Aplp2 | 0.51617 | 0.019011 |
| Apoa1bp | 0.60525 | 0.018382 |
| Apoo | 0.62423 | 0.010246 |
| Apopt1 | 0.90077 | 0.0026334 |
| Ar | -2.2826 | 0.04823 |
| Araf | 0.73562 | 7.20E-04 |
| Arap1 | -1.0813 | 7.57E-06 |
| Arfgef2 | -1.0217 | 5.45E-06 |
| Arfgef3 | -1.937 | 3.42E-16 |
| Arhgap19 | -1.5942 | 0.0044562 |
| Arhgap26 | -1.4333 | 1.29E-09 |
| Arhgap31 | -1.7109 | 0.0024457 |
| Arhgap35 | -1.7364 | 1.01E-17 |
| Arhgap39 | -0.55364 | 0.014468 |
| Arhgap42 | -1.1682 | 0.0046342 |
| Arhgap44 | -0.55922 | 0.015154 |
| Arhgef1 | -1.3218 | 8.40E-05 |
| Arhgef10 | -0.77419 | 0.0038269 |

| | | |
|---|---|---|
| Arhgef3 | 0.85781 | 0.0026435 |
| Arhgef39 | -0.7601 | 0.0097905 |
| Arid3a | -0.8194 | 0.039874 |
| Arid5b | -0.84006 | 9.76E-04 |
| Arl1 | 0.65282 | 0.0032009 |
| Arl13b | 0.76406 | 0.0033123 |
| Arl2bp | 0.4957 | 0.025566 |
| Arl5b | -0.68515 | 0.012619 |
| Arl6ip1 | 0.6464 | 0.0029722 |
| Arl6ip5 | 0.62373 | 0.0042707 |
| Armcx1 | 0.89311 | 0.0010498 |
| Armcx6 | 0.78186 | 0.004524 |
| Armt1 | 0.719 | 0.0044033 |
| Arpc3 | 0.55337 | 0.022012 |
| Arpc5l | 0.51876 | 0.022821 |
| Arrdc4 | -0.70801 | 0.048942 |
| Arse | -0.99048 | 2.42E-04 |
| Arv1 | 0.64362 | 0.015832 |
| Asb1 | -1.2267 | 1.04E-05 |
| Asb7 | -0.71594 | 0.031626 |
| Ascc2 | -0.69881 | 0.011943 |
| Ascc3 | 0.77067 | 0.0047544 |
| Asmt | -1.0816 | 0.021339 |
| Asmtl | 0.71423 | 0.0016471 |
| Asnsd1 | 0.90209 | 4.76E-05 |
| Aspdh | -0.86742 | 0.03015 |
| Asphd1 | 0.75325 | 0.0019039 |
| Aspm | -1.0268 | 0.029297 |
| Aspscr1 | -0.65472 | 0.0099462 |
| Asxl2 | -0.84495 | 0.016352 |
| Asxl3 | -1.1415 | 4.02E-04 |
| Atad1 | 0.83057 | 1.01E-04 |
| Atat1 | -0.9369 | 2.51E-05 |
| Atf6 | -0.87686 | 3.80E-04 |
| Atf6b | -0.5087 | 0.024101 |
| Atf7 | -0.83188 | 0.01792 |
| Atg12 | 0.7187 | 0.006444 |
| Atg4c | 0.59797 | 0.019744 |
| Atg5 | 0.56924 | 0.037843 |
| Atl1 | 0.61712 | 0.0049785 |
| Atl2 | 0.50288 | 0.037029 |

| | | |
|---|---|---|
| Atp11a | -0.97691 | 1.79E-04 |
| Atp13a3 | -1.7727 | 3.52E-06 |
| Atp1b3 | 0.72633 | 0.001153 |
| Atp5e | 1.1028 | 0.012962 |
| Atp5g2 | 0.65445 | 0.0094236 |
| Atp5g3 | 0.45953 | 0.045807 |
| Atp5h | 0.77798 | 3.05E-04 |
| Atp5i | 1.077 | 0.017816 |
| Atp5j | 0.94356 | 0.0042538 |
| Atp5j2 | 1.21 | 0.0040789 |
| Atp5l | 1.0746 | 0.0034644 |
| Atp5o | 0.75219 | 0.002684 |
| Atp5sl | 0.49583 | 0.046911 |
| Atp6ap2 | 0.48298 | 0.036194 |
| Atp6v0b | 0.58556 | 0.031218 |
| Atp6v0e1 | 1.0745 | 0.021853 |
| Atp6v1d | 0.65516 | 0.00189 |
| Atp6v1e1 | 0.59587 | 0.0053271 |
| Atp6v1g1 | 0.86742 | 2.43E-05 |
| Atp6v1h | 0.62818 | 0.0040638 |
| Atp8a1 | -0.9692 | 1.18E-04 |
| Atp8a2 | -1.8066 | 1.11E-13 |
| Atp8b2 | -0.94331 | 3.70E-05 |
| Atpif1 | 0.92121 | 0.0010089 |
| Atraid | 1.1696 | 7.18E-08 |
| Atrip | 0.8742 | 0.0074516 |
| Atrnl1 | -1.1409 | 1.03E-07 |
| Atxn1l | -1.2803 | 2.87E-06 |
| Atxn2 | -0.49519 | 0.042573 |
| Atxn2l | -1.1628 | 1.22E-08 |
| Atxn7 | -0.75058 | 0.04855 |
| Atxn7l3 | -0.60879 | 0.0096266 |
| Aurka | -0.93017 | 0.010291 |
| Aurkb | -1.497 | 0.0094708 |
| Avl9 | -1.0842 | 4.65E-06 |
| Axl | -0.80209 | 0.022499 |
| AY172581.2 | 1.3812 | 0.0037083 |
| AY172581.24 | 1.824 | 0.02818 |
| AY172581.9 | 1.5267 | 0.046439 |
| B2m | 1.1528 | 0.0016114 |
| B3galnt1 | 0.7753 | 3.76E-04 |

| | | |
|---|---|---|
| B3galt6 | -0.71449 | 0.047644 |
| B3gat3 | 0.59747 | 0.0062178 |
| B4galnt1 | -0.82786 | 2.88E-04 |
| B4galt4 | 0.80218 | 0.038803 |
| B9d1 | 0.81635 | 0.04844 |
| Baat | 1.4817 | 0.032327 |
| Bace1 | -0.61143 | 0.0077861 |
| Bace2 | -0.63638 | 0.013059 |
| Bach1 | -0.73406 | 0.013536 |
| Bach2 | -1.583 | 5.93E-07 |
| Bag1 | 0.74485 | 0.0015307 |
| Bahcc1 | -2.0202 | 1.26E-18 |
| Bahd1 | -1.1048 | 4.24E-06 |
| Bambi | 1.2014 | 0.013588 |
| Bbs10 | 0.79913 | 0.0054479 |
| Bbs5 | 0.97107 | 2.59E-04 |
| Bbx | -1.5695 | 2.61E-05 |
| Bcap29 | 0.68425 | 0.0037649 |
| Bcar1 | -0.60657 | 0.0096629 |
| Bcas2 | 0.85503 | 1.06E-04 |
| Bccip | 0.74459 | 0.0021833 |
| Bcl10 | 0.69207 | 0.0056945 |
| Bcl11b | -0.88911 | 9.46E-04 |
| Bcl2 | -1.6608 | 4.26E-05 |
| Bcl2l1 | -0.51832 | 0.027143 |
| Bcl2l13 | -0.77792 | 0.0039735 |
| Bcl9 | -0.68199 | 0.0023144 |
| Bdh1 | -0.58227 | 0.016193 |
| Bdh2 | 0.89318 | 0.012352 |
| Bean1 | -1.3449 | 0.0019543 |
| Best3 | -3.9404 | 0.01232 |
| Bet1 | 1.0076 | 7.81E-06 |
| Bex2 | 0.4663 | 0.042721 |
| Bhlhb9 | 0.51163 | 0.038419 |
| Bicd1 | -0.85978 | 2.32E-04 |
| Bloc1s1 | 0.65357 | 0.020111 |
| Bloc1s2 | 0.99689 | 3.67E-06 |
| Bloc1s5 | 0.74155 | 0.0028667 |
| Blzf1 | 0.60755 | 0.021339 |
| Bmp3 | -1.2079 | 1.04E-04 |
| Bnip3 | 0.62089 | 0.0036798 |

| | | |
|---|---|---|
| Boc | 0.67207 | 0.026622 |
| Bola1 | 0.74954 | 0.011626 |
| Bop1 | -0.51482 | 0.03114 |
| Borcs5 | -0.76982 | 0.0082385 |
| Borcs8 | 0.70991 | 0.011465 |
| Bpgm | 0.66483 | 0.0029517 |
| Bphl | 0.86345 | 7.98E-04 |
| Braf | -1.0421 | 2.44E-05 |
| Brap | -1.0693 | 7.67E-06 |
| Brd3 | -0.94085 | 2.43E-05 |
| Bre | 0.63933 | 0.0065685 |
| Brix1 | 0.62452 | 0.0090507 |
| Brk1 | 0.84392 | 1.95E-04 |
| Brpf3 | -1.1715 | 3.22E-06 |
| Brwd3 | -1.0563 | 1.87E-04 |
| Bspry | -0.89259 | 0.0035772 |
| Btbd1 | 0.51731 | 0.02481 |
| Btbd2 | -0.53355 | 0.022854 |
| Btbd7 | -0.60049 | 0.03263 |
| Btbd9 | -0.63515 | 0.013814 |
| Btf3 | 0.51472 | 0.023728 |
| Btf3l4 | 0.59987 | 0.0092487 |
| Btg1 | 0.66719 | 0.024013 |
| Btg2 | 0.71767 | 0.041844 |
| Btg3 | 0.57604 | 0.038803 |
| Btrc | -0.63059 | 0.0067666 |
| Bub1b | -1.66 | 2.72E-04 |
| Bud31 | 0.74157 | 0.0013762 |
| C1d | 0.86626 | 7.27E-05 |
| C1galt1c1 | 0.91868 | 8.24E-05 |
| C2cd2 | -0.86933 | 0.0010474 |
| C2cd4b | -1.1094 | 0.014023 |
| Cab39l | 0.71361 | 0.0077152 |
| Cabp1 | -0.59333 | 0.021903 |
| Cabp4 | 0.94762 | 0.011414 |
| Cachd1 | -0.89235 | 0.023464 |
| Cacna1c | -1.7453 | 3.39E-16 |
| Cacna1h | -1.5832 | 4.36E-14 |
| Cacna2d1 | -0.78432 | 3.41E-04 |
| Cacna2d2 | -1.319 | 1.54E-07 |
| Cacnb4 | -0.85919 | 1.74E-04 |

| | | |
|---|---|---|
| Cacng2 | -0.84347 | 7.20E-04 |
| Cacng4 | -1.374 | 1.22E-08 |
| Cacybp | 0.57983 | 0.01003 |
| Cad | -0.65035 | 0.008479 |
| Calm2 | 0.72329 | 6.37E-04 |
| Caln1 | -2.4626 | 2.13E-13 |
| Camk1d | -0.87037 | 2.97E-04 |
| Camk2g | -0.84094 | 1.13E-04 |
| Camk4 | -0.57037 | 0.037057 |
| Camlg | 0.58286 | 0.013855 |
| Canx | 0.54236 | 0.015491 |
| Capn15 | -0.94157 | 3.42E-04 |
| Car12 | -1.3845 | 0.034303 |
| Carm1 | -1.0958 | 5.07E-07 |
| Casc3 | -0.72649 | 0.0013326 |
| Casp3 | 0.50631 | 0.049537 |
| Cbarp | -0.59643 | 0.030112 |
| Cbfa2t2 | -1.2042 | 3.58E-07 |
| Cbfa2t3 | -1.1549 | 1.60E-05 |
| Cbfb | 0.57904 | 0.027764 |
| Cbl | -2.5318 | 4.99E-25 |
| Cblb | -0.7672 | 0.012899 |
| Cbr4 | 0.57806 | 0.03885 |
| Cbx4 | -0.63528 | 0.0087182 |
| Cby1 | 0.65948 | 0.0042455 |
| Cc2d1a | -0.87853 | 4.76E-05 |
| Ccar1 | 0.83505 | 2.33E-04 |
| Ccbe1 | -1.8152 | 8.89E-04 |
| Ccdc112 | 0.59655 | 0.021174 |
| Ccdc12 | 0.54496 | 0.044117 |
| Ccdc126 | 0.89952 | 4.64E-04 |
| Ccdc138 | 1.0074 | 0.011239 |
| Ccdc142 | -0.95978 | 3.50E-04 |
| Ccdc146 | 1.6677 | 0.0056118 |
| Ccdc15 | -1.772 | 0.0016548 |
| Ccdc152 | 2.5941 | 0.027916 |
| Ccdc157 | -1.4475 | 5.05E-07 |
| Ccdc175 | 1.3506 | 2.22E-05 |
| Ccdc177 | -0.7776 | 0.023269 |
| Ccdc181 | 0.50213 | 0.049537 |
| Ccdc188 | -1.9259 | 5.85E-04 |

| | | |
|---|---|---|
| Ccdc43 | 0.61288 | 0.020992 |
| Ccdc50 | -0.91779 | 1.18E-04 |
| Ccdc53 | 0.92985 | 2.86E-04 |
| Ccdc58 | 0.91635 | 0.0010636 |
| Ccdc59 | 0.68799 | 0.0089551 |
| Ccdc6 | -1.5319 | 2.24E-09 |
| Ccdc62 | 0.70345 | 0.046339 |
| Ccdc85a | -0.66952 | 0.018496 |
| Ccdc88a | -0.71568 | 0.001513 |
| Ccdc88c | -1.109 | 0.0059708 |
| Ccdc90b | 0.77563 | 0.0056191 |
| Ccdc92 | -0.52478 | 0.028209 |
| Ccdc93 | -0.8597 | 0.0027859 |
| Ccdc97 | -0.59032 | 0.014046 |
| Ccl2 | 3.1292 | 0.012962 |
| Ccna2 | -1.3953 | 7.98E-04 |
| Ccnd1 | -0.71983 | 0.039774 |
| Ccndbp1 | 0.51742 | 0.044346 |
| Ccnf | -1.3705 | 0.049033 |
| Ccng2 | 0.62379 | 0.0075407 |
| Ccnh | 0.54423 | 0.025115 |
| Ccni | 0.91654 | 1.00E-05 |
| Ccnl1 | 0.6394 | 0.015155 |
| Ccnt2 | 0.6552 | 0.0039575 |
| Ccser2 | -0.8944 | 6.92E-05 |
| Cct2 | 0.69428 | 0.0015184 |
| Cct3 | 0.6091 | 0.0068632 |
| Cct4 | 0.56875 | 0.014376 |
| Cct5 | 0.58833 | 0.0090942 |
| Cct6a | 0.49352 | 0.033966 |
| Cct8 | 0.61355 | 0.0061303 |
| Cd164 | 0.80636 | 2.51E-04 |
| Cd302 | 1.6496 | 0.0010759 |
| Cd47 | -0.68043 | 0.0050472 |
| Cd59 | 1.3718 | 3.35E-08 |
| Cd63 | 1.1251 | 0.012214 |
| Cd81 | 0.65994 | 0.0022182 |
| Cd83 | 0.85249 | 0.015409 |
| Cdadc1 | 0.88624 | 5.75E-05 |
| Cdan1 | -0.72543 | 0.0087803 |
| Cdc123 | 0.67852 | 0.0021519 |

| | | |
|---|---|---|
| Cdc20 | -1.5476 | 0.0049085 |
| Cdc26 | 0.90838 | 3.74E-05 |
| Cdc45 | -1.3253 | 0.0042481 |
| Cdc73 | -0.65965 | 0.021187 |
| Cdca5 | -2.0542 | 0.017097 |
| Cdh22 | -1.1832 | 1.17E-04 |
| Cdh23 | -2.4801 | 1.24E-05 |
| Cdh24 | -0.93199 | 0.0021104 |
| Cdhr1 | -1.637 | 0.0031287 |
| Cdk12 | -0.55904 | 0.029342 |
| Cdk18 | -0.87855 | 0.031091 |
| Cdk2 | -0.99154 | 0.0042057 |
| Cdon | -0.76323 | 0.009396 |
| Cdr2 | -0.82166 | 0.01908 |
| Cdr2l | -1.3915 | 2.56E-07 |
| Cds2 | -1.7382 | 6.21E-16 |
| Cdt1 | -1.3744 | 1.25E-04 |
| Cdv3 | -1.102 | 1.01E-06 |
| Cdyl2 | -1.1163 | 9.24E-04 |
| Cecr2 | -1.2261 | 9.38E-04 |
| Cecr6 | -1.182 | 7.42E-06 |
| Celf2 | -0.85302 | 6.55E-05 |
| Celf3 | -0.61851 | 0.00522 |
| Cenpt | -1.8842 | 0.0093658 |
| Cep164 | -0.67793 | 0.015292 |
| Cep170b | -1.2405 | 8.93E-10 |
| Cep19 | 0.63354 | 0.0057535 |
| Cep350 | -1.1718 | 2.98E-07 |
| Cep85l | -0.69185 | 0.040128 |
| Cept1 | 0.95105 | 1.68E-05 |
| Cers6 | -1.0826 | 9.25E-05 |
| Cetn4 | 0.95488 | 0.04728 |
| Cfap36 | 0.72657 | 7.38E-04 |
| Cfb | -2.012 | 0.03545 |
| Cfdp1 | 0.60011 | 0.0085147 |
| Cfl2 | 0.55254 | 0.014961 |
| Cgrrf1 | 0.96178 | 7.98E-05 |
| Chad | #NAME? | 0.0059454 |
| Chchd1 | 1.0204 | 0.018061 |
| Chchd3 | 0.7518 | 8.94E-04 |
| Chd1 | -0.70302 | 0.0065695 |

| | | |
|---|---|---|
| Chd7 | -1.2577 | 0.0043544 |
| Cherp | -0.83425 | 1.42E-04 |
| Chl1 | -0.51633 | 0.024037 |
| Chmp2a | 0.5248 | 0.023783 |
| Chmp3 | 0.56045 | 0.015384 |
| Chmp5 | 0.67467 | 0.0024523 |
| Chmp7 | -0.56055 | 0.017019 |
| Chpf2 | -0.58905 | 0.012236 |
| Chrac1 | 1.0525 | 6.86E-04 |
| Chrd | -1.2984 | 1.58E-04 |
| Chrna4 | -1.133 | 6.52E-04 |
| Chrna7 | -1.0219 | 0.0030765 |
| Chrnb2 | -1.2521 | 1.19E-08 |
| Chst1 | -0.63125 | 0.0039145 |
| Chst11 | -1.3561 | 9.49E-09 |
| Chst3 | -1.5545 | 3.01E-04 |
| Chst7 | -1.3395 | 1.88E-08 |
| Chtf18 | -1.2645 | 4.63E-04 |
| Churc1 | 1.1535 | 0.015568 |
| Cinp | 0.69077 | 0.0023124 |
| Cir1 | 0.8555 | 0.037185 |
| Cisd1 | 0.51118 | 0.026574 |
| Cisd2 | 0.72214 | 9.69E-04 |
| Ckap2l | -1.5638 | 0.022054 |
| Ckmt1b | 1.2148 | 5.30E-04 |
| Cks1b | 0.80056 | 0.0086443 |
| Clasrp | -1.019 | 0.017563 |
| Clcc1 | 0.90869 | 5.83E-05 |
| Clcn5 | -1.9028 | 4.22E-04 |
| Clcn6 | -1.0296 | 4.68E-06 |
| Clcn7 | -0.83175 | 2.23E-04 |
| Cldn10 | 2.2037 | 3.40E-05 |
| Cldn12 | 0.64825 | 0.0058732 |
| Cldnd1 | 0.94869 | 8.62E-06 |
| Clint1 | -0.7393 | 0.027398 |
| Clip2 | -0.50149 | 0.030178 |
| Clip4 | -0.64734 | 0.02658 |
| Clmn | -2.2647 | 8.01E-15 |
| Clns1a | 0.50335 | 0.033195 |
| Clock | -1.2944 | 5.05E-07 |
| Clpb | -0.71909 | 0.0023022 |

| | | |
|---|---|---|
| Clstn2 | -1.6742 | 6.56E-14 |
| Clu | 1.4287 | 4.94E-06 |
| Cluap1 | 0.97025 | 1.29E-05 |
| Cluh | -1.1082 | 9.50E-08 |
| Clybl | 0.90247 | 0.044485 |
| Cmbl | 0.86625 | 0.018712 |
| Cmc2 | 1.2193 | 1.09E-04 |
| Cmip | -1.3132 | 4.31E-11 |
| Cml5 | 2.0098 | 0.045477 |
| Cmtm4 | -1.5897 | 4.47E-13 |
| Cnbp | 0.58943 | 0.007094 |
| Cnep1r1 | 0.64983 | 0.0071355 |
| Cnih1 | 0.68235 | 0.0023472 |
| Cnih4 | 0.60653 | 0.024385 |
| Cnksr2 | -0.6211 | 0.0067182 |
| Cnnm1 | -0.88038 | 1.48E-04 |
| Cnnm2 | -0.6234 | 0.026722 |
| Cnnm4 | -0.85961 | 0.0072813 |
| Cnot7 | 0.73406 | 6.77E-04 |
| Cnot8 | 0.58887 | 0.0098942 |
| Cnpy2 | 0.49833 | 0.044723 |
| Cntn1 | -0.51182 | 0.019164 |
| Cntn2 | -1.094 | 7.31E-04 |
| Cntn4 | 1.3015 | 0.0024102 |
| Cntnap1 | -1.1415 | 2.68E-08 |
| Col11a2 | -2.5036 | 0.015279 |
| Col16a1 | -2.3252 | 3.80E-04 |
| Col19a1 | -1.725 | 8.40E-08 |
| Col23a1 | -1.4226 | 0.028715 |
| Col25a1 | -0.94774 | 0.0051824 |
| Col27a1 | -3.0218 | 2.99E-04 |
| Col4a1 | -1.114 | 0.0017319 |
| Col4a2 | -1.1746 | 7.60E-08 |
| Col5a1 | -2.2645 | 0.012632 |
| Col5a3 | -2.8455 | 1.36E-09 |
| Col6a3 | -3.286 | 3.10E-07 |
| Col6a4 | -2.1974 | 7.01E-09 |
| Commd1 | 0.80483 | 3.33E-04 |
| Commd2 | 0.64516 | 0.011137 |
| Commd3 | 0.70633 | 0.0011 |
| Commd5 | 0.77045 | 0.01253 |

| | | |
|---|---|---|
| Commd6 | 1.034 | 6.37E-06 |
| Commd7 | 0.78002 | 0.0010385 |
| Commd8 | 0.85644 | 9.57E-05 |
| Commd9 | 0.75461 | 0.013122 |
| Coprs | 0.7284 | 0.0013634 |
| Cops2 | 0.46904 | 0.048043 |
| Cops4 | 0.703 | 0.0013456 |
| Coq3 | 0.70136 | 0.0034996 |
| Coro2a | -1.1352 | 4.25E-05 |
| Coro7 | -0.7599 | 0.0011219 |
| Coro7 | -1.7102 | 0.0013702 |
| Cotl1 | -0.8977 | 1.66E-04 |
| Cox10 | -0.90975 | 0.0013776 |
| Cox16 | 0.87426 | 0.006502 |
| Cox17 | 1.4926 | 6.70E-04 |
| Cox20 | 0.7892 | 6.37E-04 |
| Cox5b | 0.68209 | 0.023031 |
| Cox6a1 | 0.7347 | 0.034138 |
| Cox6b1 | 1.0901 | 0.0037682 |
| Cox6b1 | 0.96821 | 7.47E-05 |
| Cox6c | 1.2082 | 0.0059708 |
| Cox7a2l | 0.71502 | 8.23E-04 |
| Cox7a2l2 | 1.2401 | 0.0060089 |
| Cox8a | 0.86232 | 0.045129 |
| Cpd | -0.68294 | 0.0026136 |
| Cpeb2 | -1.0227 | 1.40E-05 |
| Cpeb3 | -1.4915 | 2.82E-08 |
| Cpeb4 | -0.69268 | 0.0022947 |
| Cpne6 | -0.61444 | 0.0061406 |
| Cpne7 | -1.0491 | 0.0044949 |
| Cpne8 | 0.95153 | 0.031126 |
| Cpq | 1.3852 | 0.031972 |
| Cpsf1 | -0.65606 | 0.0041019 |
| Cpt2 | 0.97614 | 5.86E-04 |
| Cramp1 | -1.1269 | 1.30E-04 |
| Crb2 | -1.3886 | 0.0022319 |
| Crcp | 1.0625 | 2.04E-06 |
| Creb5 | -1.1445 | 0.01345 |
| Creg1 | 1.2345 | 1.37E-08 |
| Creg2 | -1.4495 | 5.63E-08 |
| Crhr1 | -0.7114 | 0.035978 |

| | | |
|---|---|---|
| Cript | 1.0631 | 4.64E-06 |
| Crkl | -0.60599 | 0.010084 |
| Crocc | -1.2852 | 1.77E-07 |
| Crtc3 | -1.5923 | 1.05E-04 |
| Cryzl1 | 0.73278 | 0.012359 |
| Csdc2 | -0.46355 | 0.044034 |
| Csgalnact2 | 0.71009 | 0.0076772 |
| Csmd1 | -0.73628 | 0.0011219 |
| Csmd2 | -1.8093 | 1.17E-07 |
| Csmd3 | -0.72654 | 0.0034781 |
| Csnk1g1 | -1.4802 | 1.32E-04 |
| Cspg4 | -1.3894 | 7.94E-04 |
| Csrnp3 | -1.5668 | 1.66E-08 |
| Cst3 | 1.5182 | 0.0085736 |
| Cstb | 0.93607 | 0.047793 |
| Cstf2t | 0.52371 | 0.021537 |
| Ctdsp2 | -0.64531 | 0.029468 |
| Ctif | -1.0358 | 5.59E-05 |
| Ctnnd1 | -0.50615 | 0.028882 |
| Ctsa | 0.52372 | 0.027468 |
| Ctsh | 1.1074 | 2.63E-06 |
| Ctsl | 0.83265 | 1.49E-04 |
| Ctxn2 | 1.4404 | 0.0051824 |
| Cuedc1 | -0.83394 | 4.01E-04 |
| Cwc15 | 0.80283 | 2.78E-04 |
| Cwc22 | 0.67121 | 0.011207 |
| Cwc25 | 0.55984 | 0.044247 |
| Cxcl12 | 0.84684 | 0.0037636 |
| Cxcr4 | 1.2378 | 1.90E-04 |
| Cyb5a | 1.1182 | 1.64E-07 |
| Cyb5b | -0.51189 | 0.029468 |
| Cyba | 1.5929 | 1.70E-04 |
| Cycs | 0.47964 | 0.046911 |
| Cyp26b1 | 1.4385 | 3.25E-04 |
| Cyp4f39 | -3.0069 | 0.030046 |
| Cyp4f5 | -0.91692 | 0.0091367 |
| Cyp4v3 | 1.2485 | 0.033311 |
| Cyp7b1 | 1.1615 | 0.0068237 |
| Dab1 | -0.81851 | 4.56E-04 |
| Dad1 | 0.99489 | 5.56E-04 |
| Dap | 0.72967 | 0.0076322 |

| | | |
|---|---|---|
| Dars | 0.49509 | 0.038247 |
| Dazap2 | 0.64901 | 0.0029856 |
| Dbn1 | -0.92755 | 8.47E-06 |
| Dbnl | -1.1461 | 5.63E-08 |
| Dcaf15 | -0.97498 | 3.27E-05 |
| Dcaf17 | -1.359 | 4.48E-06 |
| Dcakd | -0.70315 | 0.0010634 |
| Dcbld1 | -0.78942 | 0.0033453 |
| Dcc | -1.9882 | 7.04E-17 |
| Dchs1 | -1.228 | 0.0049334 |
| Dclk2 | -0.8238 | 1.61E-04 |
| Dcp1a | -0.73307 | 0.034853 |
| Dcp2 | -1.2725 | 2.59E-04 |
| Dctn5 | 0.60337 | 0.0081319 |
| Dctn6 | 0.74893 | 0.0012965 |
| Dcun1d1 | 0.54645 | 0.048233 |
| Ddit3 | 0.62233 | 0.036686 |
| Ddt | 0.92733 | 9.83E-06 |
| Ddx17 | -0.6372 | 0.0033127 |
| Ddx21 | -0.62884 | 0.0057102 |
| Ddx50 | 0.54685 | 0.026131 |
| Ddx54 | -1.0949 | 1.08E-06 |
| Decr1 | 1.0433 | 5.61E-04 |
| Degs1 | 0.59703 | 0.01025 |
| Dek | 0.54255 | 0.021003 |
| Dennd2a | -0.74611 | 0.0015596 |
| Dennd3 | -0.8317 | 0.0072149 |
| Dennd4b | -0.65019 | 0.0070824 |
| Dennd6b | -0.59983 | 0.036134 |
| Denr | 0.60352 | 0.0068557 |
| Deptor | -1.3754 | 0.0026322 |
| Derl1 | 0.45624 | 0.04935 |
| Desi1 | -0.50939 | 0.033662 |
| Desi2 | -0.79325 | 0.010148 |
| Dfnb31 | -0.70852 | 0.0068637 |
| Dgkd | -0.66418 | 0.0023472 |
| Dgke | -0.78616 | 9.02E-04 |
| Dgkg | -1.0416 | 1.13E-04 |
| Dgkh | -2.4978 | 1.78E-14 |
| Dgki | -1.7245 | 5.62E-14 |
| Dgkq | -0.99274 | 1.38E-05 |

| | | |
|---|---|---|
| Dhcr24 | -1.5437 | 0.020698 |
| Dhrs7 | 1.3283 | 0.0023706 |
| Dhx16 | -0.62201 | 0.043247 |
| Dhx33 | -0.62233 | 0.037545 |
| Dhx37 | -0.93213 | 4.39E-04 |
| Dhx9 | -0.4771 | 0.039929 |
| Diaph1 | -0.999 | 6.36E-05 |
| Diras1 | -0.523 | 0.026134 |
| Dlat | -0.62961 | 0.0053382 |
| Dlgap5 | -1.3034 | 0.010124 |
| Dlx6as | 1.3071 | 0.010349 |
| Dmpk | -0.80595 | 0.0083849 |
| Dmrt3 | -1.7752 | 1.14E-04 |
| Dmrta2 | -0.7902 | 0.024293 |
| Dnai1 | 1.4335 | 0.0082356 |
| Dnajb13 | 0.78446 | 0.024616 |
| Dnajb9 | 0.82828 | 2.71E-04 |
| Dnajc16 | -0.96258 | 0.0010365 |
| Dnajc18 | -0.60595 | 0.016383 |
| Dnajc19 | 0.88326 | 1.13E-04 |
| Dnajc24 | 0.83445 | 0.002153 |
| Dnajc5 | -1.1005 | 1.14E-07 |
| Dnajc8 | 0.93474 | 1.24E-05 |
| Dner | 0.65655 | 0.0044499 |
| Dnlz | 0.63364 | 0.026364 |
| Dnm3 | -0.7718 | 6.99E-04 |
| Dnmbp | -0.98215 | 0.0010513 |
| Dnmt1 | -0.48796 | 0.045288 |
| Dnmt3a | -1.7832 | 8.84E-18 |
| Doc2a | -0.72875 | 0.018681 |
| Dok6 | -1.5513 | 2.50E-06 |
| Dpagt1 | 0.55383 | 0.025913 |
| Dpcd | 0.83246 | 0.0075435 |
| Dph1 | -0.72981 | 0.012988 |
| Dph3 | 0.85325 | 0.013351 |
| Dpm1 | 0.52243 | 0.029492 |
| Dpm2 | 0.75112 | 6.67E-04 |
| Dpp9 | -0.68935 | 0.0038555 |
| Dpy19l3 | -1.1139 | 1.40E-05 |
| Dpy30 | 1.1126 | 5.82E-07 |
| Dpysl3 | -1.2918 | 3.31E-09 |

| | | |
|---|---|---|
| Dpysl5 | -1.8325 | 4.14E-15 |
| Dram2 | 0.71109 | 0.022129 |
| Draxin | -1.4289 | 0.022602 |
| Drg1 | 0.56614 | 0.018293 |
| Drp2 | -0.62429 | 0.020992 |
| Dsel | 0.60576 | 0.028604 |
| Dstn | 0.81798 | 9.96E-05 |
| Dtd2 | 0.74494 | 0.0049334 |
| Dtl | -1.3694 | 0.0050777 |
| Dtwd1 | 0.89855 | 0.04219 |
| Dtx4 | -0.84315 | 9.13E-04 |
| Dtymk | 0.63696 | 0.0098986 |
| Dus3l | -0.5776 | 0.036282 |
| Dus4l | 0.83813 | 0.014748 |
| Dusp1 | 0.79242 | 0.031921 |
| Dusp12 | 0.67149 | 0.014348 |
| Dusp15 | -0.64558 | 0.0080727 |
| Dusp18 | -1.0105 | 1.11E-05 |
| Dvl1 | -0.77643 | 4.22E-04 |
| Dvl3 | -1.9396 | 1.27E-20 |
| Dynlrb1 | 1.1159 | 1.15E-05 |
| Dynlrb2 | 2.0694 | 0.017238 |
| Dynlt1 | 0.68961 | 0.040319 |
| Dynlt3 | 0.76116 | 3.46E-04 |
| Dyrk1a | -0.549 | 0.019354 |
| Dyrk1b | -1.3164 | 5.25E-08 |
| Dzank1 | -0.96285 | 3.48E-05 |
| E2f3 | -1.0864 | 3.95E-04 |
| E2f7 | -2.8678 | 1.24E-05 |
| E2f8 | -2.4076 | 1.60E-04 |
| Ebag9 | 0.94815 | 2.88E-05 |
| Ech1 | 0.61954 | 0.0087853 |
| Echdc1 | 0.81484 | 3.09E-04 |
| Eci1 | 0.82246 | 0.0017878 |
| Ect2 | -1.418 | 0.022308 |
| Eda2r | -1.6891 | 0.011439 |
| Edem3 | -1.0588 | 2.21E-06 |
| Eef1b2 | 1.0059 | 1.41E-06 |
| Eef1d | 0.67123 | 0.0039145 |
| Eef1e1 | 0.81587 | 2.55E-04 |
| Eef1g | 0.71839 | 0.0055885 |

| | | |
|---|---|---|
| Eef2k | -0.55448 | 0.017809 |
| Efcab7 | 1.0879 | 0.042265 |
| Efr3b | -1.6042 | 1.51E-14 |
| Efs | -0.96515 | 0.027729 |
| Egflam | 2.0468 | 0.0072926 |
| Egr2 | 1.1686 | 0.0020432 |
| Ehd3 | -0.63065 | 0.0047301 |
| Eid1 | 0.50136 | 0.048167 |
| Eif1a | 0.52729 | 0.030011 |
| Eif1ad | 0.56186 | 0.024314 |
| Eif2a | 0.54167 | 0.032966 |
| Eif2b3 | 0.55384 | 0.044712 |
| Eif3e | 0.78508 | 3.48E-04 |
| Eif3h | 0.63054 | 0.034248 |
| Eif3i | 0.67128 | 0.0030294 |
| Eif3k | 0.98049 | 5.86E-04 |
| Eif3l | 0.5713 | 0.017469 |
| Eif3m | 0.95896 | 9.06E-06 |
| Eif4a2 | 0.63538 | 0.0029687 |
| Eif5a2 | -0.89629 | 0.0099399 |
| Eif5b | 0.69006 | 0.0022504 |
| Elavl2 | 0.66946 | 0.01318 |
| Elavl3 | -1.0269 | 1.18E-06 |
| Ell3 | 0.99357 | 0.028498 |
| Elmsan1 | -1.2101 | 2.26E-06 |
| Elof1 | 0.63772 | 0.0069802 |
| Elovl1 | 0.79079 | 0.0053646 |
| Elp5 | 0.52405 | 0.040353 |
| Emc2 | 0.63133 | 0.0055744 |
| Emc3 | 0.58865 | 0.0091316 |
| Emc4 | 0.5396 | 0.023462 |
| Emc6 | 0.67069 | 0.0043215 |
| Emc7 | 0.89188 | 2.44E-05 |
| Eme2 | -0.94192 | 1.10E-04 |
| Eml1 | -0.64272 | 0.006787 |
| Eml3 | -0.80391 | 0.021339 |
| Eml6 | -1.6499 | 1.01E-08 |
| Enah | -1.0739 | 1.15E-06 |
| Endod1 | -0.85074 | 0.0022215 |
| Enho | 0.60596 | 0.012259 |
| Enkur | 1.0096 | 3.04E-05 |

| | | |
|---|---|---|
| Enpp2 | 1.4049 | 0.017527 |
| Enpp4 | 0.5658 | 0.038983 |
| Enpp5 | 0.49255 | 0.030174 |
| Entpd5 | -0.64835 | 0.032394 |
| Entpd7 | -1.0868 | 0.036671 |
| Eny2 | 0.59971 | 0.0077194 |
| Epb4.1 | -1.4611 | 9.66E-09 |
| Epc2 | 0.52772 | 0.031529 |
| Epha10 | -1.1741 | 0.01486 |
| Epha6 | -1.6967 | 4.78E-10 |
| Epha8 | -1.3381 | 3.53E-04 |
| Ephb2 | -0.87665 | 2.84E-04 |
| Epm2a | -1.415 | 5.80E-08 |
| Epm2aip1 | -1.0807 | 0.0020876 |
| Eps8 | -0.53608 | 0.032638 |
| Erbb4 | -1.3611 | 0.0010468 |
| Erc1 | -1.7222 | 3.15E-08 |
| Erc2 | -1.0047 | 6.37E-04 |
| Ercc2 | -0.95525 | 5.83E-04 |
| Ergic1 | -0.76418 | 0.0022543 |
| Ergic3 | 0.53018 | 0.020102 |
| Erh | 0.73468 | 5.73E-04 |
| Erich3 | -1.5895 | 6.35E-05 |
| Erlec1 | 0.89412 | 2.85E-05 |
| Ern1 | -1.2609 | 0.002155 |
| Erp29 | 0.62966 | 0.0055179 |
| Erp44 | 0.50276 | 0.041312 |
| Esco1 | 0.84906 | 3.48E-04 |
| Esd | 0.8537 | 9.63E-05 |
| Espl1 | -1.5215 | 1.57E-05 |
| Esyt2 | -0.60184 | 0.021299 |
| Etfa | 0.73509 | 0.001145 |
| Etfb | 0.53907 | 0.025186 |
| Etl4 | -1.1881 | 1.94E-05 |
| Etv3 | -0.72948 | 0.024975 |
| Etv4 | -1.2809 | 0.029778 |
| Etv5 | -0.48373 | 0.034312 |
| Etv6 | -1.4199 | 0.0034753 |
| Evi5 | -0.90912 | 3.48E-04 |
| Evi5l | -0.48354 | 0.040681 |
| Exoc8 | -0.78254 | 9.14E-04 |

| | | |
|---|---|---|
| Exosc1 | 0.63988 | 0.014528 |
| F8 | -0.56015 | 0.032172 |
| Faap100 | -0.65676 | 0.019748 |
| Fabp3 | 0.9043 | 0.038336 |
| Fam101b | -1.4684 | 0.0050204 |
| Fam102a | -1.4013 | 5.27E-07 |
| Fam102b | -1.0211 | 6.79E-04 |
| Fam103a1 | 0.77996 | 7.15E-04 |
| Fam104a | 0.54942 | 0.036105 |
| Fam126a | -1.4843 | 4.68E-07 |
| Fam129b | -0.9191 | 9.32E-04 |
| Fam133b | 0.56894 | 0.016858 |
| Fam135b | -1.79 | 2.02E-13 |
| Fam136a | 0.86351 | 0.03939 |
| Fam149b1 | 0.51367 | 0.041312 |
| Fam151b | 0.77702 | 0.011626 |
| Fam155b | -0.72363 | 0.0051628 |
| Fam162a | 1.0609 | 8.68E-07 |
| Fam163b | -1.2091 | 2.53E-08 |
| Fam168a | -0.90897 | 1.31E-05 |
| Fam168b | -1.3316 | 1.76E-07 |
| Fam171a1 | -0.53171 | 0.030743 |
| Fam172a | 0.52908 | 0.026462 |
| Fam173b | 0.60348 | 0.029014 |
| Fam174a | 0.8478 | 1.93E-04 |
| Fam175b | 0.64567 | 0.0069099 |
| Fam181b | -0.6397 | 0.023626 |
| Fam184b | -0.84812 | 0.0011124 |
| FAM187A | -1.9556 | 0.0094336 |
| Fam189a1 | -1.2848 | 1.17E-06 |
| Fam189b | -0.81019 | 3.59E-04 |
| Fam199x | -1.0998 | 0.0039477 |
| Fam213a | 0.68711 | 0.0046255 |
| Fam219a | -0.85929 | 5.98E-05 |
| Fam222b | -0.69957 | 0.0035205 |
| Fam229b | 1.1724 | 0.016275 |
| Fam45a | 1.2884 | 3.87E-09 |
| Fam53c | -0.90875 | 9.64E-05 |
| Fam64a | -1.8495 | 0.0082943 |
| Fam73b | -0.57669 | 0.014439 |
| Fam83d | -1.2257 | 0.036088 |

| | | |
|---|---|---|
| Fam89a | -1.2252 | 0.014439 |
| Fam8a1 | -0.63293 | 0.0082883 |
| Fam92a1 | 0.86791 | 1.28E-04 |
| Fam96a | 1.2115 | 7.50E-07 |
| Fam96b | 0.93716 | 0.0013628 |
| Fan1 | -0.69614 | 0.025529 |
| Fanca | -1.4794 | 0.0010057 |
| Fanci | -1.0109 | 0.011568 |
| Fancl | 0.74776 | 0.024862 |
| Farp1 | -0.71073 | 0.0011535 |
| Fars2 | 0.53514 | 0.042214 |
| Farsb | 0.53077 | 0.028676 |
| Fastk | -0.51221 | 0.037158 |
| Faxc | -1.351 | 2.33E-07 |
| Fbrs | -1.0491 | 3.03E-05 |
| Fbxl17 | -0.59482 | 0.020412 |
| Fbxl18 | -1.9132 | 2.07E-16 |
| Fbxl2 | -0.7121 | 0.0070843 |
| Fbxl20 | -1.2193 | 3.99E-05 |
| Fbxl22 | 1.2184 | 0.0050472 |
| Fbxl5 | 0.91674 | 2.83E-05 |
| Fbxl7 | -1.7123 | 0.0039554 |
| Fbxo10 | -0.62658 | 0.0085368 |
| Fbxo11 | 0.64995 | 0.0068044 |
| Fbxo42 | -1.1307 | 1.46E-06 |
| Fbxo8 | 0.77174 | 0.001286 |
| Fbxw2 | 0.68444 | 0.0022815 |
| Fbxw5 | -0.50593 | 0.040834 |
| Fbxw7 | 0.64985 | 0.006502 |
| Fbxw8 | -0.60426 | 0.017641 |
| Fcf1 | 0.96991 | 9.99E-04 |
| Fcgrt | 1.0647 | 3.50E-05 |
| Fchsd1 | -0.8612 | 0.0045154 |
| Fdx1 | 0.68808 | 0.01908 |
| Fdx1l | 0.5373 | 0.027729 |
| Fem1a | -0.97088 | 2.23E-05 |
| Fem1b | -0.89353 | 5.13E-05 |
| Fer | -0.61958 | 0.045988 |
| Fgd1 | -1.2036 | 3.22E-06 |
| Fgf22 | -0.69735 | 0.02324 |
| Fgf9 | 0.7118 | 0.014044 |

| | | |
|---|---|---|
| Fggy | 0.71186 | 0.046881 |
| Fh | 0.53728 | 0.02236 |
| Fis1 | 0.92606 | 1.84E-05 |
| Fjx1 | -0.46708 | 0.046882 |
| Fkbp2 | 0.89643 | 4.13E-05 |
| Fkbp5 | -0.7151 | 0.0032938 |
| Flrt1 | -1.57 | 9.66E-09 |
| Flvcr1 | -1.1197 | 9.20E-04 |
| Flywch1 | -0.82795 | 1.22E-04 |
| Flywch2 | 0.75064 | 0.038581 |
| Fmc1 | 0.82343 | 0.007147 |
| Fmn1 | -2.0715 | 0.045266 |
| Fmnl1 | -1.0115 | 7.98E-06 |
| Fnbp1 | -0.92071 | 0.00575 |
| Fnbp1l | -0.50442 | 0.032507 |
| Fndc3b | -1.6953 | 1.38E-10 |
| Fnip2 | -1.1365 | 7.64E-06 |
| Fnta | 0.52225 | 0.036304 |
| Foxk1 | -1.7708 | 1.19E-09 |
| Foxm1 | -1.7217 | 1.41E-05 |
| Foxn2 | -1.0533 | 0.001067 |
| Foxn3 | -1.4761 | 4.41E-08 |
| Foxo1 | -1.0189 | 5.33E-04 |
| Foxp4 | -0.7844 | 6.30E-04 |
| Foxred2 | -0.94107 | 0.0072493 |
| Fra10ac1 | 0.57758 | 0.029703 |
| Fras1 | -2.3581 | 2.84E-18 |
| Frem2 | -1.6704 | 0.0011962 |
| Frg1 | 0.9757 | 0.03629 |
| Frmd4a | -1.1321 | 1.57E-06 |
| Frmd4b | -1.0289 | 0.034138 |
| Frmd8 | -0.68046 | 0.018401 |
| Frmpd1 | -1.044 | 8.09E-05 |
| Frrs1l | -1.4564 | 6.82E-07 |
| Fryl | -0.94221 | 4.46E-05 |
| Fscn2 | -1.2633 | 0.028032 |
| Fuca1 | 0.56629 | 0.024964 |
| Fuca2 | 0.89933 | 4.39E-04 |
| Fundc1 | 0.91077 | 7.74E-05 |
| Furin | -0.68371 | 0.0041813 |
| Fut9 | -1.738 | 9.37E-08 |

| | | |
|---|---|---|
| Fxn | 0.78255 | 0.0018824 |
| Fxr2 | -0.51371 | 0.02996 |
| Fxyd1 | 1.5659 | 1.69E-05 |
| Fzd3 | -1.503 | 5.06E-06 |
| Fzd7 | -1.3325 | 0.043686 |
| Fzd9 | -0.71024 | 0.018054 |
| Fzr1 | -0.85216 | 1.29E-04 |
| Gab2 | -1.0703 | 4.94E-04 |
| Gabarap | 0.71573 | 7.83E-04 |
| Gabarapl2 | 0.76037 | 3.11E-04 |
| Gabrb1 | -2.1842 | 1.55E-16 |
| Gabrb2 | -0.96691 | 0.0018025 |
| Gadd45g | 0.76873 | 0.0029503 |
| Gadd45gip1 | 0.6536 | 0.0051921 |
| Gal3st4 | 0.98291 | 0.01949 |
| Galc | 0.63234 | 0.041905 |
| Galnt10 | -1.3444 | 0.0272 |
| Galnt14 | -0.97602 | 0.0064449 |
| Galnt16 | -0.62141 | 0.048188 |
| Galnt7 | -0.76611 | 0.0074263 |
| Galnt9 | -0.75825 | 0.0033631 |
| Gan | -2.4346 | 3.28E-15 |
| Gareml | -0.87289 | 0.033014 |
| Gas2 | 0.95452 | 0.014116 |
| Gas2l3 | -2.2871 | 6.81E-05 |
| Gatad2a | -0.75532 | 8.59E-04 |
| Gatad2b | -1.0258 | 1.88E-04 |
| Gatm | 0.58103 | 0.011331 |
| Gatsl2 | -0.90537 | 3.02E-05 |
| Gcsh | 0.81532 | 2.44E-04 |
| Gda | -0.87216 | 5.15E-05 |
| Gdf11 | -0.9491 | 0.0033412 |
| Gdi2 | 0.49938 | 0.036335 |
| Gemin2 | 0.90978 | 0.016408 |
| Gemin5 | -0.56982 | 0.029297 |
| Gfm2 | -0.69633 | 0.0053299 |
| Gfod1 | -1.3995 | 9.81E-09 |
| Gfpt1 | -0.79185 | 4.59E-04 |
| Gfra1 | -0.67667 | 0.027928 |
| Gga1 | -0.72632 | 0.0017331 |
| Ggct | 0.7909 | 0.0092172 |

| | | |
|---|---|---|
| Ggh | 0.69381 | 0.0048989 |
| Ggnbp2 | 0.54539 | 0.018966 |
| Ggt7 | -0.50255 | 0.043826 |
| Ghrl | -1.4441 | 6.36E-05 |
| Gid4 | 0.50718 | 0.048655 |
| Gigyf1 | -1.3527 | 1.24E-05 |
| Ginm1 | 0.89572 | 4.50E-04 |
| Gipr | -1.678 | 0.0063408 |
| Gja1 | 1.1434 | 0.0012795 |
| Gjc1 | 0.78348 | 0.041671 |
| Glg1 | -0.87828 | 0.00112 |
| Glipr1 | 1.1346 | 0.0081688 |
| Glo1 | 0.69605 | 0.0014955 |
| Glod4 | 0.55542 | 0.018701 |
| Glrb | 0.52142 | 0.019118 |
| Glrx | 0.82204 | 0.010294 |
| Glrx2 | 0.60523 | 0.0070407 |
| Glrx3 | 0.59758 | 0.0068632 |
| Gltscr1 | -1.0086 | 3.78E-05 |
| Gm1043 | -1.397 | 0.004568 |
| Gm884 | 0.78413 | 0.0016864 |
| Gmfg | 1.5404 | 0.0030921 |
| Gmps | -0.55295 | 0.034303 |
| Gna11 | -1.0134 | 2.40E-06 |
| Gna13 | -1.112 | 1.15E-04 |
| Gnaq | -0.9518 | 1.42E-05 |
| Gnb3 | -1.9786 | 1.38E-04 |
| Gnb4 | -0.65686 | 0.0030114 |
| Gng10 | 1.127 | 2.50E-08 |
| Gng13 | 1.1806 | 1.72E-04 |
| Gng5 | 1.1647 | 0.0024909 |
| Gng7 | -0.67703 | 0.012389 |
| Gnpda2 | 0.66854 | 0.026663 |
| Gnpnat1 | 0.91388 | 0.0038979 |
| Gnptab | -0.82081 | 4.40E-04 |
| Golga7 | 0.56485 | 0.010987 |
| Golga7b | -0.63928 | 0.01951 |
| Gon4l | -0.95095 | 2.51E-05 |
| Gorab | 0.65917 | 0.017468 |
| Gpatch2 | -1.1745 | 7.68E-05 |
| Gpatch4 | 0.66088 | 0.023702 |

| | | |
|---|---|---|
| Gpbp1l1 | 0.60029 | 0.038156 |
| Gpc1 | -0.50577 | 0.026612 |
| Gpc2 | -0.83119 | 0.02324 |
| Gpc6 | -0.89925 | 0.036892 |
| Gpd2 | -0.86093 | 4.90E-04 |
| Gpm6b | 0.53215 | 0.019881 |
| Gpr161 | -1.3492 | 9.05E-06 |
| Gpr19 | 0.71737 | 0.0069349 |
| Gpr26 | -1.8328 | 2.55E-12 |
| Gprasp1 | 0.68102 | 0.0018354 |
| Gprasp2 | 0.47882 | 0.044525 |
| Gprin3 | -1.2074 | 0.004568 |
| Gpsm1 | -0.60217 | 0.0084002 |
| Gpx4 | 0.93364 | 4.90E-05 |
| Gpx8 | 1.3769 | 3.17E-09 |
| Gramd4 | -0.85328 | 4.91E-04 |
| Grasp | -0.85995 | 0.0038269 |
| Grcc10 | 0.97279 | 0.00203 |
| Greb1l | -1.4037 | 0.0016127 |
| Grik4 | -0.67089 | 0.0064325 |
| Grin2d | -1.1299 | 0.0039145 |
| Grin3b | -1.247 | 2.77E-04 |
| Grip2 | -0.75545 | 0.0013424 |
| Grk6 | -0.84987 | 1.39E-04 |
| Grm3 | -1.0981 | 0.019118 |
| Grpel1 | 0.64276 | 0.0041145 |
| Grwd1 | -0.62648 | 0.035042 |
| Gse1 | -1.691 | 1.39E-13 |
| Gsk3a | -0.47027 | 0.034887 |
| Gskip | 0.73943 | 0.0067697 |
| Gsta4 | 1.6064 | 2.07E-12 |
| Gstk1 | 0.98022 | 2.05E-04 |
| Gstm1 | 1.6176 | 1.00E-10 |
| Gstm2 | 0.7268 | 0.0016402 |
| Gstt1 | 1.2924 | 0.016593 |
| Gtdc1 | 0.76243 | 0.016935 |
| Gtf2a1 | 0.88646 | 4.61E-05 |
| Gtf2h5 | 1.1388 | 2.64E-08 |
| Gtf3c6 | 0.82338 | 4.25E-04 |
| Gtpbp1 | -0.79143 | 6.79E-04 |
| Gtpbp10 | 0.73851 | 0.0079743 |

| | | |
|---|---|---|
| Gtpbp8 | 0.63951 | 0.019593 |
| Gtse1 | -1.1748 | 9.83E-04 |
| Gucy1a2 | -1.7791 | 1.49E-08 |
| Gucy1b3 | 0.50234 | 0.03733 |
| Guk1 | 0.81805 | 0.024188 |
| Gulp1 | 0.75836 | 0.024644 |
| Gzf1 | 0.56696 | 0.026978 |
| H1fx | -0.69162 | 0.0075022 |
| H2afj | 0.70945 | 0.028709 |
| H2afz | 0.52916 | 0.026352 |
| H3f3a | 0.58842 | 0.0075233 |
| H3f3b | 0.68587 | 0.001153 |
| H3f3c | 2.9634 | 0.030758 |
| H6pd | -1.0053 | 0.0023722 |
| Hadhb | 0.55925 | 0.015751 |
| Hat1 | 0.60493 | 0.018113 |
| Hbp1 | 0.61281 | 0.014826 |
| Hcn3 | -0.68967 | 0.0059335 |
| Hcn4 | -1.4181 | 3.24E-08 |
| Hdac9 | -0.55114 | 0.028934 |
| Hddc3 | 0.94142 | 0.0018709 |
| Hdhd2 | 0.70992 | 0.0012088 |
| Heatr4 | 0.95344 | 0.020907 |
| Hebp1 | 0.77666 | 6.01E-04 |
| Hectd4 | -1.6376 | 2.81E-16 |
| Hecw1 | -0.83364 | 2.88E-04 |
| Hecw2 | -1.009 | 1.15E-05 |
| Heg1 | -1.8169 | 2.73E-10 |
| Helz | -1.5298 | 2.41E-10 |
| Heph | 1.1829 | 0.013837 |
| Hes5 | -1.5265 | 1.61E-06 |
| Hexb | 0.50525 | 0.037859 |
| Hgs | -0.4847 | 0.043745 |
| Hhip | -3.2965 | 2.22E-07 |
| Hhipl1 | -0.86676 | 0.0019506 |
| Hiat1 | 0.65395 | 0.0046051 |
| Hibch | 0.64826 | 0.012995 |
| Hid1 | -0.65993 | 0.0025 |
| Hif1an | -1.3482 | 1.27E-07 |
| Hint3 | 0.75577 | 0.0029687 |
| Hip1 | -0.88551 | 7.35E-04 |

| | | |
|---|---|---|
| Hip1r | -0.55688 | 0.019832 |
| Hjurp | -1.6537 | 0.015602 |
| Hk2 | -1.3424 | 0.024014 |
| Hlcs | -1.4172 | 8.48E-05 |
| Hltf | 0.68304 | 0.0077987 |
| Hmbox1 | -1.9978 | 4.49E-15 |
| Hmcn1 | -2.3893 | 0.029466 |
| Hmg20b | -0.76074 | 0.024775 |
| Hmgcll1 | 0.67781 | 0.014929 |
| Hmgn3 | 0.72047 | 0.0013093 |
| Hnrnpa0 | -0.8973 | 2.11E-04 |
| Hnrnpul2 | -0.71586 | 8.87E-04 |
| Homer3 | -0.87183 | 0.011471 |
| Hr | -0.71221 | 0.047954 |
| Hrsp12 | 1.7418 | 6.64E-16 |
| Hs6st1 | -0.50053 | 0.03744 |
| Hs6st3 | -2.1749 | 4.21E-10 |
| Hsbp1 | 0.76507 | 1.93E-04 |
| Hscb | 0.99319 | 1.51E-04 |
| Hsd17b10 | 0.74442 | 0.0015942 |
| Hsd17b11 | 1.2256 | 4.61E-06 |
| Hsd17b12 | 0.72202 | 0.0013733 |
| Hsdl2 | 0.84595 | 3.12E-04 |
| Hsp90b1 | 0.85229 | 6.62E-05 |
| Hspa12a | -0.4687 | 0.038736 |
| Hspa2 | 0.70043 | 0.0044198 |
| Hspa5 | 0.54032 | 0.020242 |
| Hspa8 | 1.9122 | 4.99E-08 |
| Hspa8 | 1.0573 | 2.27E-07 |
| Hspb11 | 0.7354 | 0.020714 |
| Hspd1 | 0.6181 | 0.0063765 |
| Hspg2 | -1.5331 | 0.0037636 |
| Hunk | -1.4209 | 0.0015124 |
| Hyal1 | -1.8013 | 1.04E-04 |
| Hypk | 1.2217 | 5.51E-07 |
| Ica1 | 0.69983 | 0.011472 |
| Icam5 | -0.48888 | 0.037511 |
| Ice1 | -0.6753 | 0.0028481 |
| Icmt | -0.92766 | 2.71E-04 |
| Ict1 | 0.62304 | 0.013631 |
| Id2 | 1.3104 | 0.0011193 |

| | | |
|---|---|---|
| Ifi44 | 1.0657 | 0.013308 |
| Ifngr1 | 0.5428 | 0.044009 |
| Ift122 | -0.5538 | 0.038374 |
| Ift140 | -0.5514 | 0.023874 |
| Ift20 | 1.2558 | 2.37E-09 |
| Ift46 | 0.74947 | 0.016235 |
| Igbp1 | 0.81518 | 9.91E-04 |
| Igdcc3 | -1.1299 | 0.012236 |
| Igdcc4 | -0.97084 | 0.031857 |
| Igf1r | -1.1533 | 0.001264 |
| Igf2bp3 | -1.181 | 0.036652 |
| Igf2r | -0.49419 | 0.034874 |
| Igip | 2.3846 | 6.77E-06 |
| Iglon5 | -0.72819 | 0.0074357 |
| Igsf1 | 2.1572 | 0.0030722 |
| Igsf10 | 0.64415 | 0.039861 |
| Ikbip | 0.62875 | 0.022047 |
| Ikbkap | -0.59319 | 0.0085286 |
| Ikzf4 | -0.98768 | 0.0026435 |
| Il16 | -1.1893 | 0.040225 |
| Il1r1 | -2.1936 | 0.026967 |
| Ildr2 | -1.6162 | 1.12E-12 |
| Immp1l | 1.0207 | 8.21E-06 |
| Impact | 0.82272 | 6.40E-05 |
| Impad1 | 0.5396 | 0.016683 |
| Impdh1 | -0.51608 | 0.031453 |
| Incenp | -0.97534 | 0.001906 |
| Ing3 | 0.58688 | 0.030911 |
| Ing4 | 0.5911 | 0.010943 |
| Ino80 | -0.64441 | 0.017453 |
| Ino80d | -2.6168 | 8.14E-15 |
| Inpp5j | -0.81895 | 8.59E-04 |
| Insig2 | 0.50813 | 0.042813 |
| Insr | -1.2275 | 8.63E-08 |
| Ints4 | 0.60184 | 0.016409 |
| Ints6 | -0.84757 | 0.024163 |
| Ipo9 | -0.51718 | 0.027279 |
| Iqcb1 | 0.73254 | 0.0026435 |
| Iqcc | 0.84057 | 0.01168 |
| Iqce | -0.84534 | 0.0010423 |
| Iqgap1 | -0.91176 | 7.02E-04 |

| | | |
|---|---|---|
| Iqgap3 | -1.9671 | 1.24E-04 |
| Iqub | 0.92724 | 0.048669 |
| Ireb2 | -0.57658 | 0.015506 |
| Irs1 | -1.2708 | 0.0018082 |
| Isca1 | 0.53783 | 0.014424 |
| Iscu | 0.85615 | 4.83E-05 |
| Islr2 | -1.2384 | 1.66E-07 |
| Itfg1 | 0.59616 | 0.0058487 |
| Itga3 | -0.93728 | 1.60E-04 |
| Itga6 | -0.99684 | 0.0035351 |
| Itga9 | -1.1599 | 0.015171 |
| Itgae | 1.1412 | 2.25E-04 |
| Itgav | -1.1841 | 0.0094994 |
| Itgb3bp | 1.0432 | 0.001915 |
| Itgb8 | -1.2288 | 4.23E-05 |
| Itm2b | 0.8012 | 1.16E-04 |
| Itpk1 | -0.66729 | 0.0046259 |
| Itpkb | -1.0069 | 0.0091181 |
| Jag2 | -0.86798 | 2.02E-04 |
| Jagn1 | 0.63678 | 0.0055692 |
| Jakmip1 | -0.58144 | 0.013818 |
| Jam2 | 0.50365 | 0.033911 |
| Jam3 | 0.7904 | 0.025765 |
| Jarid2 | -0.63313 | 0.0096844 |
| Jmjd7 | -1.8612 | 0.0013709 |
| Jtb | 0.62929 | 0.0065685 |
| Jup | -0.64871 | 0.0053271 |
| Kat2a | -0.81596 | 2.10E-04 |
| Kat6a | -1.3432 | 7.83E-09 |
| Katnb1 | -0.60084 | 0.011862 |
| Kazn | -1.2198 | 4.01E-08 |
| Kbtbd11 | -1.6175 | 8.28E-05 |
| Kbtbd2 | 0.64448 | 0.0082943 |
| Kcna1 | -2.2339 | 1.02E-17 |
| Kcnab2 | -1.2456 | 4.21E-07 |
| Kcnd3 | -1.6133 | 2.46E-12 |
| Kcnf1 | -0.6226 | 0.01003 |
| Kcng3 | -2.031 | 1.92E-07 |
| Kcnh2 | -0.99321 | 8.92E-05 |
| Kcnip3 | -1.1159 | 2.43E-05 |
| Kcnj10 | -1.4276 | 3.34E-04 |

| | | |
|---|---|---|
| Kcnj4 | -0.91337 | 3.57E-05 |
| Kcnj6 | -1.6349 | 1.93E-09 |
| Kcnk10 | -0.87058 | 0.0014348 |
| Kcnk12 | -1.2376 | 9.03E-07 |
| Kcnk9 | -1.3616 | 6.30E-09 |
| Kcnn2 | 0.8639 | 5.77E-04 |
| Kcnn3 | -1.444 | 1.82E-07 |
| Kcnq5 | -0.67439 | 0.0050472 |
| Kcns3 | 1.2572 | 0.0022815 |
| Kctd10 | -0.49426 | 0.037202 |
| Kctd17 | -0.50055 | 0.040326 |
| Kctd20 | -0.84562 | 0.0011481 |
| Kctd3 | -0.57336 | 0.011829 |
| Kctd7 | -0.77733 | 0.01577 |
| Kdelr1 | 0.483 | 0.038827 |
| Kdelr2 | 0.65354 | 0.0057004 |
| Kdm4c | 0.63854 | 0.012735 |
| Khsrp | -0.95853 | 1.70E-05 |
| Kif13a | -0.96852 | 2.74E-05 |
| Kif13b | -1.0177 | 3.31E-04 |
| Kif14 | -2.1474 | 2.34E-05 |
| Kif18b | -1.968 | 4.93E-05 |
| Kif1c | -0.52368 | 0.020992 |
| Kif23 | -1.0554 | 0.014015 |
| Kif26a | -1.5207 | 0.0012915 |
| Kif26b | -1.8785 | 4.03E-07 |
| Kif28p | 1.4713 | 0.017776 |
| Kif2c | -1.5045 | 0.017362 |
| Kif3a | -0.5486 | 0.015488 |
| Kifap3 | 0.46195 | 0.040177 |
| Kifc1 | -1.6026 | 0.0017437 |
| Kifc3 | -0.4793 | 0.042813 |
| Kin | 0.80157 | 0.009099 |
| Kirrel | -1.4883 | 8.48E-06 |
| Kiz | 0.77558 | 0.0072015 |
| Klc2 | -0.80337 | 1.28E-04 |
| Klf12 | -1.9483 | 4.37E-10 |
| Klf13 | -1.6641 | 4.39E-07 |
| Klf7 | -1.3201 | 8.30E-05 |
| Klf7 | -1.3709 | 5.03E-05 |
| Klf8 | -1.547 | 0.0051395 |

| | | |
|---|---|---|
| Klf9 | -0.49637 | 0.035109 |
| Klhdc10 | -1.2673 | 7.50E-09 |
| Klhl11 | -1.1473 | 1.35E-06 |
| Klhl15 | -0.80324 | 0.033941 |
| Klhl17 | -1.1465 | 1.52E-04 |
| Klhl18 | -0.74621 | 0.0046317 |
| Klhl23 | 0.56874 | 0.014681 |
| Klhl3 | -1.711 | 1.43E-08 |
| Klhl7 | 0.74047 | 5.85E-04 |
| Kmt2a | -2.1055 | 1.97E-23 |
| Kmt2c | -0.78921 | 3.57E-04 |
| Kmt2e | -0.48826 | 0.040599 |
| Kntc1 | -1.0942 | 2.59E-04 |
| Kpna5 | -0.67404 | 0.04689 |
| Kpna6 | -0.90446 | 2.18E-04 |
| Kpnb1 | -0.92284 | 9.16E-06 |
| Krtcap2 | 0.88505 | 2.05E-04 |
| Ksr1 | -2.0445 | 3.94E-18 |
| Ksr2 | -2.8102 | 5.13E-18 |
| L2hgdh | 0.55391 | 0.049581 |
| L3mbtl3 | -0.60194 | 0.035978 |
| l7Rn6 | 0.58725 | 0.011366 |
| Lace1 | 0.84193 | 0.037966 |
| Lactb2 | 0.52312 | 0.042526 |
| Lamp2 | 0.97373 | 0.0014102 |
| Lamtor2 | 1.0515 | 6.77E-06 |
| Lamtor3 | 0.75802 | 8.79E-04 |
| Lamtor4 | 1.0841 | 2.96E-06 |
| Lamtor5 | 1.0548 | 6.08E-07 |
| Lancl3 | -1.2924 | 3.34E-05 |
| Laptm4a | 0.9824 | 1.91E-06 |
| Larp1 | -1.387 | 2.13E-11 |
| Lbhd1 | 1.1512 | 2.86E-05 |
| Lcmt1 | 0.72951 | 0.0011952 |
| Lcor | -1.4682 | 2.54E-08 |
| Ldhb | 0.77676 | 1.81E-04 |
| Ldoc1l | -1.4067 | 8.28E-09 |
| Lef1 | -2.5766 | 0.014263 |
| Leprot | 0.68915 | 0.0040811 |
| Letm1 | -0.89652 | 4.80E-05 |
| Lgals1 | 1.6328 | 0.0043458 |

| | | |
|---|---|---|
| Lgi2 | -1.0013 | 0.042079 |
| Lgmn | 0.88956 | 6.45E-05 |
| Lhfp | 0.86388 | 0.029461 |
| Lhx4 | -1.9858 | 0.0037072 |
| Lifr | -1.823 | 5.47E-13 |
| Limch1 | -1.1205 | 1.92E-07 |
| Limd1 | -0.96268 | 0.012389 |
| Limk1 | -1.089 | 1.11E-06 |
| Lin7c | 0.59877 | 0.0075233 |
| Lingo3 | -0.91065 | 0.0018878 |
| Lipt2 | 0.72779 | 0.029813 |
| Lix1 | 0.54357 | 0.026595 |
| Lman2l | -0.73319 | 0.012903 |
| Lmbrd1 | 0.94602 | 6.97E-06 |
| Lmln | -0.64336 | 0.027607 |
| Lmnb1 | -0.99493 | 1.19E-04 |
| Lnx2 | -0.78006 | 0.012383 |
| LOC100174910 | 0.84579 | 0.0012495 |
| LOC100233176 | 0.90092 | 1.80E-04 |
| LOC100294508 | -0.49249 | 0.032435 |
| LOC100359687 | 1.3615 | 9.29E-04 |
| LOC100359922 | 0.76835 | 2.66E-04 |
| LOC100360056 | 1.0712 | 0.047042 |
| LOC100360057 | 0.83617 | 0.010323 |
| LOC100360087 | 0.98008 | 0.0059764 |
| LOC100360120 | 0.48721 | 0.034944 |
| LOC100360218 | 0.99664 | 5.08E-04 |
| LOC100360522 | 1.2323 | 0.0010468 |
| LOC100360573 | 1.8671 | 9.65E-06 |
| LOC100360647 | 1.1653 | 1.29E-07 |
| LOC100360726 | -1.6134 | 0.007639 |
| LOC100360791 | 2.5063 | 0.007222 |
| LOC100361087 | 1.0291 | 0.045498 |
| LOC100361993 | 1.2538 | 0.0058459 |
| LOC100362027 | 1.4558 | 0.016631 |
| LOC100362814 | -1.731 | 2.26E-16 |
| LOC100362980 | 0.93862 | 3.37E-04 |
| LOC100363675 | -0.85266 | 1.06E-04 |
| LOC100364435 | 0.9201 | 2.14E-04 |
| LOC100365839 | 1.002 | 0.047532 |
| LOC100909548 | 0.48738 | 0.04674 |

| | | |
|---|---|---|
| LOC100909840 | 0.61283 | 0.0043293 |
| LOC100909954 | -1.491 | 1.62E-09 |
| LOC100910137 | 0.77583 | 0.0010218 |
| LOC100910370 | 0.71814 | 0.0012188 |
| LOC100910438 | 0.53921 | 0.046518 |
| LOC100910508 | -0.94569 | 2.68E-04 |
| LOC100910577 | 0.81684 | 0.0049597 |
| LOC100910732 | -0.64652 | 0.041341 |
| LOC100910779 | 0.86806 | 0.003955 |
| LOC100911293 | 0.86828 | 0.027166 |
| LOC100911372 | 1.4053 | 0.042587 |
| LOC100911440 | -0.51236 | 0.026768 |
| LOC100911483 | 1.1982 | 2.79E-07 |
| LOC100911485 | -0.93092 | 0.0048262 |
| LOC100911585 | 0.91836 | 0.044985 |
| LOC100911664 | 0.55996 | 0.015386 |
| LOC100911685 | -1.0371 | 3.25E-04 |
| LOC100911840 | 1.3066 | 0.034329 |
| LOC100911956 | 0.91733 | 0.013588 |
| LOC100912042 | 0.89192 | 0.0022729 |
| LOC100912163 | 0.95363 | 2.44E-04 |
| LOC100912182 | 1.3378 | 1.67E-04 |
| LOC100912195 | 0.88261 | 0.023507 |
| LOC100912369 | 0.53185 | 0.022355 |
| LOC100912429 | 0.76639 | 0.0052411 |
| LOC100912445 | -3.0561 | 5.54E-08 |
| LOC100912481 | -0.78988 | 0.0044136 |
| LOC100912924 | 0.85148 | 0.0032476 |
| LOC102547897 | -1.5327 | 0.010698 |
| LOC102549817 | 1.2266 | 0.027729 |
| LOC102549842 | 0.74329 | 0.0083778 |
| LOC102550385 | 0.81346 | 0.0050761 |
| LOC102550391 | 0.6983 | 0.032129 |
| LOC102552166 | -1.9916 | 0.0018016 |
| LOC102552640 | -0.86441 | 0.023788 |
| LOC102552920 | 0.58112 | 0.029877 |
| LOC102554232 | 0.78381 | 5.42E-04 |
| LOC102555167 | -0.51796 | 0.038285 |
| LOC102555377 | 1.1624 | 2.86E-05 |
| LOC102555672 | 1.0889 | 2.28E-06 |
| LOC103689975 | -1.9627 | 3.48E-05 |

| | | |
|---|---|---|
| LOC103689983 | 0.94554 | 8.05E-06 |
| LOC103690024 | -0.81808 | 0.0061313 |
| LOC103690050 | -3.4228 | 3.80E-06 |
| LOC103690054 | 1.2181 | 7.19E-04 |
| LOC103690139 | 1.1096 | 0.025324 |
| LOC103690171 | 1.1287 | 0.0053749 |
| LOC103693210 | 0.83648 | 2.80E-04 |
| LOC103693430 | 1.0823 | 0.0062756 |
| LOC103693776 | -0.79859 | 1.36E-04 |
| LOC103694877 | 0.80131 | 9.24E-04 |
| LOC257650 | 1.0918 | 0.0028375 |
| LOC288913 | 0.63724 | 0.004568 |
| LOC301124 | 0.87227 | 0.0027371 |
| LOC361985 | 0.97673 | 2.94E-06 |
| LOC498555 | 0.85609 | 0.030528 |
| LOC499781 | -1.0511 | 0.0036809 |
| LOC501110 | 1.1378 | 6.41E-05 |
| LOC678813 | 0.92406 | 0.014435 |
| LOC679894 | -0.71643 | 0.020957 |
| LOC680227 | -7.1172 | 7.33E-04 |
| LOC681282 | 0.84158 | 0.0091431 |
| LOC681367 | 0.84914 | 0.01632 |
| LOC683593 | -1.9 | 1.84E-13 |
| LOC683897 | 0.81822 | 0.0018881 |
| LOC684797 | 0.4948 | 0.037095 |
| LOC687679 | 1.0616 | 0.0043738 |
| LOC689499 | -1.7548 | 5.30E-14 |
| LOC689991 | -1.0989 | 6.07E-07 |
| LOC690422 | -1.0106 | 0.0020739 |
| LOC690871 | 1.1144 | 0.0022287 |
| LOC691056 | -0.62531 | 0.0038872 |
| LOC691153 | -0.78193 | 0.0064978 |
| LOC691427 | 1.922 | 0.026364 |
| LOC691807 | 1.1581 | 1.97E-08 |
| Lonp1 | -0.57665 | 0.0098942 |
| Lonrf2 | -2.0594 | 9.10E-23 |
| Lonrf3 | -0.68057 | 0.04923 |
| Lpar4 | 1.4966 | 1.03E-04 |
| Lpgat1 | -0.66795 | 0.0032316 |
| Lpin3 | -1.8902 | 0.0043766 |
| Lpp | -4.0358 | 8.22E-09 |

| | | |
|---|---|---|
| Lppr2 | -1.0494 | 2.64E-07 |
| Lrba | -1.1539 | 8.37E-05 |
| Lrch3 | -1.1109 | 4.00E-04 |
| Lrch4 | -1.2149 | 1.06E-04 |
| Lrfn1 | -0.56425 | 0.014348 |
| Lrif1 | 0.85966 | 0.0014307 |
| Lrig1 | -0.65066 | 0.024746 |
| Lrp1b | -0.85796 | 0.027298 |
| Lrp1b | 2.6045 | 0.003699 |
| Lrp5 | -1.0355 | 9.74E-05 |
| Lrp6 | -1.1609 | 5.07E-07 |
| Lrrc16b | -1.4664 | 1.54E-12 |
| Lrrc26 | -1.4174 | 0.018796 |
| Lrrc3b | 1.2893 | 0.013346 |
| Lrrc40 | 0.86266 | 1.64E-04 |
| Lrrc49 | 0.52301 | 0.024974 |
| Lrrc4c | 0.55703 | 0.035485 |
| Lrrc57 | 1.0018 | 8.48E-05 |
| Lrrc58 | -1.3448 | 0.0039814 |
| Lrrc75b | -0.84035 | 0.017164 |
| Lrrc8c | -1.046 | 5.38E-04 |
| Lrrcc1 | 0.74 | 0.0086026 |
| Lrrn3 | 0.68892 | 0.018314 |
| Lsamp | -0.89764 | 3.05E-05 |
| Lsg1 | 0.58872 | 0.022914 |
| Lsm1 | 0.87618 | 2.58E-04 |
| Lsm11 | -1.3074 | 8.52E-06 |
| Lsm3 | 1.2594 | 6.76E-06 |
| Lsm5 | 1.2193 | 6.25E-04 |
| Lsm6 | 0.83144 | 0.003449 |
| Lsm7 | 0.96772 | 0.0069339 |
| Ltbp4 | -1.2045 | 2.60E-06 |
| Lurap1 | -0.74591 | 0.040442 |
| Lxn | 0.81039 | 0.038576 |
| Lypd5 | -2.3124 | 0.0028108 |
| Lypd6 | -0.95897 | 0.014013 |
| Lyrm1 | 0.75547 | 0.0074607 |
| Lyrm2 | 0.73897 | 0.0023298 |
| Lyrm5 | 1.0137 | 5.68E-04 |
| Lysmd2 | 0.53832 | 0.031529 |
| Lyz2 | 1.2087 | 0.014303 |

| | | |
|---|---|---|
| Lztfl1 | 1.0733 | 6.88E-06 |
| Lztr1 | -0.51716 | 0.030718 |
| Lzts3 | -0.70752 | 0.002051 |
| Maats1 | 0.90689 | 0.034138 |
| Mafg | -0.83825 | 2.38E-04 |
| Magi1 | -1.0396 | 3.52E-06 |
| Magoh | 0.87682 | 3.63E-04 |
| Mal2 | 0.64834 | 0.0064996 |
| Malsu1 | 0.60386 | 0.026454 |
| Mamdc4 | -0.91861 | 0.045327 |
| Maml2 | -1.1863 | 0.0142 |
| Man1a2 | -0.81227 | 5.20E-04 |
| Man1b1 | -0.50156 | 0.032161 |
| Man2c1 | -0.56788 | 0.01832 |
| Manbal | 0.7373 | 0.0011481 |
| Manf | 0.68126 | 0.0023917 |
| Maob | 1.2662 | 0.013266 |
| Map1s | -0.50697 | 0.031627 |
| Map2k2 | -0.6511 | 0.0046342 |
| Map2k7 | -0.71743 | 0.0050032 |
| Map3k13 | -0.94807 | 0.0012584 |
| Map3k2 | -1.4378 | 3.02E-05 |
| Map3k9 | -1.9075 | 1.11E-18 |
| Map4k2 | -1.3261 | 3.88E-06 |
| Map6d1 | -1.0694 | 5.16E-05 |
| Map9 | -0.68912 | 0.0023235 |
| Mapk1ip1 | 1.1977 | 0.011613 |
| Mapk1ip1l | -0.53098 | 0.034525 |
| Mapk8ip2 | -1.5435 | 1.94E-14 |
| Mapt | -1.2913 | 1.59E-10 |
| Marcks | 0.59437 | 0.0074147 |
| Mark1 | -0.63319 | 0.005012 |
| Mark2 | -0.98797 | 6.91E-06 |
| Mark4 | -0.82269 | 1.24E-04 |
| Masp2 | 0.61246 | 0.02996 |
| MAST1 | -0.62601 | 0.0046776 |
| Mast3 | -0.66309 | 0.0051922 |
| Matk | -0.50866 | 0.041282 |
| Matr3 | 0.79314 | 1.38E-04 |
| Mau2 | -0.80592 | 4.05E-04 |
| Mbd1 | -0.58363 | 0.01304 |

| | | |
|---|---|---|
| Mbd6 | -1.4221 | 1.58E-05 |
| Mbip | 0.87597 | 0.0025221 |
| Mbnl2 | -0.66351 | 0.0051964 |
| Mcart1 | -1.4443 | 6.07E-04 |
| Mcc | -0.90242 | 0.011099 |
| Mccc1 | 0.57535 | 0.040216 |
| Mcee | 0.61831 | 0.018754 |
| Mcf2l | -0.77132 | 4.27E-04 |
| Mcm10 | -1.7495 | 0.012457 |
| Mcm2 | -0.67679 | 0.030251 |
| Mcm3 | -1.4517 | 1.71E-05 |
| Mcm3ap | -0.5475 | 0.018293 |
| Mcm5 | -1.5006 | 0.0021092 |
| Mcts1 | 0.80699 | 3.97E-04 |
| Mcu | -0.61476 | 0.014657 |
| Mcur1 | -1.1042 | 3.82E-04 |
| Mdc1 | -1.7006 | 1.85E-09 |
| Mdga1 | -1.3721 | 1.81E-08 |
| Mdga2 | -0.82481 | 0.0017331 |
| Mdk | 0.97697 | 1.55E-05 |
| Mdn1 | -0.64549 | 0.0064949 |
| Mdp1 | 0.55957 | 0.042691 |
| Med10 | 0.49919 | 0.040504 |
| Med11 | 1.0144 | 4.40E-04 |
| Med12 | -1.0441 | 0.0033116 |
| Med12l | -2.1045 | 4.51E-14 |
| Med15 | -0.64429 | 0.0054042 |
| Med19 | 0.84529 | 0.0091238 |
| Med21 | 0.72681 | 0.007556 |
| Med25 | -0.8082 | 1.70E-04 |
| Med28 | 1.084 | 9.99E-07 |
| Med30 | 0.88314 | 2.54E-04 |
| Med31 | 0.98205 | 0.0046611 |
| Med4 | 0.91288 | 1.12E-04 |
| Med6 | 0.88218 | 4.21E-04 |
| Med7 | 0.59259 | 0.041593 |
| Megf11 | -0.74985 | 0.018562 |
| Megf9 | -0.72018 | 0.0013382 |
| Melk | -1.3965 | 0.028034 |
| Mesdc2 | 0.67817 | 0.002227 |
| Mest | -0.63026 | 0.011536 |

| | | |
|---|---|---|
| Mettl25 | 0.74148 | 0.036815 |
| Mettl5 | 0.81486 | 0.0010242 |
| Mettl9 | 0.96977 | 8.20E-06 |
| Mex3d | -0.73362 | 0.0027485 |
| Mff | 0.78505 | 2.10E-04 |
| Mfn2 | 1.0105 | 0.029676 |
| Mfsd12 | -1.1739 | 4.07E-07 |
| Mga | -0.65729 | 0.0064978 |
| Mgam | -1.105 | 0.00203 |
| Mgat3 | -0.85168 | 7.57E-05 |
| Mgat4b | -0.59557 | 0.0094708 |
| Mgat5 | -1.514 | 0.0068448 |
| MGC116121 | 0.80427 | 0.0061406 |
| MGC94335 | 0.92687 | 1.95E-04 |
| Mgrn1 | -0.64319 | 0.0029496 |
| Mgst3 | 0.59963 | 0.0079879 |
| Micall1 | -1.7402 | 5.76E-11 |
| Mid1 | -1.0584 | 0.0067956 |
| Midn | -1.2059 | 4.89E-06 |
| Mief1 | -0.71579 | 0.0057342 |
| Mien1 | 0.66154 | 0.0029359 |
| Mier1 | 0.48498 | 0.042228 |
| Mier2 | -0.8033 | 0.001056 |
| Minos1 | 0.74218 | 0.043893 |
| Mir3557 | 3.5788 | 0.022054 |
| Mir3597-1 | -2.2377 | 0.0039018 |
| Mir6333 | -1.8392 | 9.06E-04 |
| Mki67 | -1.6666 | 3.61E-04 |
| Mkks | 0.77467 | 3.82E-04 |
| Mkl1 | -0.89634 | 9.11E-05 |
| Mknk1 | -0.68905 | 0.024301 |
| Mknk2 | -0.87915 | 7.68E-05 |
| Mkrn2 | 0.55951 | 0.020992 |
| Mlc1 | 0.87145 | 6.43E-04 |
| Mllt1 | -0.5413 | 0.018931 |
| Mllt4 | -0.8677 | 6.33E-05 |
| Mlst8 | -0.83616 | 5.49E-04 |
| Mlxip | -1.8069 | 9.92E-12 |
| Mmaa | 0.94053 | 0.0017319 |
| Mmab | -1.2731 | 0.0042538 |
| Mmachc | 0.5873 | 0.045054 |

| | | |
|---|---|---|
| Mmp15 | -1.0182 | 9.72E-05 |
| Mmp16 | -1.0558 | 1.74E-06 |
| Mn1 | -0.99421 | 8.83E-06 |
| Mnat1 | 0.61772 | 0.016593 |
| Mnt | -0.59913 | 0.0093211 |
| Mob1b | -0.72875 | 0.004524 |
| Mob4 | 0.6657 | 0.0020447 |
| Mon1b | -0.78702 | 0.0018045 |
| Morf4l2 | 0.47809 | 0.035161 |
| Morn2 | 1.4377 | 0.0010343 |
| Mpc1 | 0.94264 | 1.06E-04 |
| Mpc2 | 0.81552 | 0.011213 |
| Mpdu1 | 0.60072 | 0.01526 |
| Mphosph10 | 0.56275 | 0.04157 |
| Mphosph6 | 0.98652 | 1.41E-05 |
| Mpp3 | -0.74777 | 9.89E-04 |
| Mras | -0.77383 | 2.71E-04 |
| Mre11a | -0.97101 | 4.36E-04 |
| Mroh1 | -1.8016 | 5.01E-08 |
| Mroh6 | -3.2302 | 0.040442 |
| Mrp63 | 0.80823 | 4.69E-04 |
| Mrpl13 | 1.1791 | 2.84E-06 |
| Mrpl14 | 0.87342 | 0.0051964 |
| Mrpl18 | 1.1003 | 1.47E-06 |
| Mrpl20 | 0.76567 | 0.0072954 |
| Mrpl21 | 0.88709 | 2.19E-04 |
| Mrpl22 | 0.80056 | 0.0021816 |
| Mrpl23 | 0.79761 | 6.70E-04 |
| Mrpl27 | 0.90724 | 4.11E-05 |
| Mrpl28 | 0.52878 | 0.030134 |
| Mrpl32 | 0.77065 | 0.0036534 |
| Mrpl33 | 0.89544 | 0.0060462 |
| Mrpl34 | 0.88331 | 0.0072551 |
| Mrpl35 | 0.86605 | 1.68E-04 |
| Mrpl39 | 0.76083 | 0.003748 |
| Mrpl41 | 0.63581 | 0.0060653 |
| Mrpl42 | 1.1025 | 5.37E-04 |
| Mrpl44 | 0.56852 | 0.024395 |
| Mrpl46 | 0.72999 | 0.0029122 |
| Mrpl48 | 1.0552 | 1.35E-06 |
| Mrpl49 | 0.48012 | 0.040357 |

| | | |
|---|---|---|
| Mrpl51 | 0.49451 | 0.046018 |
| Mrpl52 | 1.0153 | 1.30E-05 |
| Mrpl54 | 0.80858 | 3.89E-04 |
| Mrps14 | 0.86922 | 7.27E-05 |
| Mrps15 | 0.88094 | 1.10E-04 |
| Mrps16 | 0.59215 | 0.018231 |
| Mrps17 | 0.88341 | 7.46E-04 |
| Mrps18a | 0.66137 | 0.0040065 |
| Mrps18c | 1.0093 | 4.48E-04 |
| Mrps21 | 1.2026 | 1.14E-04 |
| Mrps22 | 0.59167 | 0.017688 |
| Mrps24 | 0.71257 | 0.0025833 |
| Mrps25 | 0.6602 | 0.0032361 |
| Mrps26 | 0.54019 | 0.02475 |
| Mrps28 | 0.9002 | 6.44E-04 |
| Mrps31 | 0.84596 | 7.66E-04 |
| Mrps33 | 0.95806 | 1.23E-05 |
| Mrps35 | 0.82992 | 4.04E-04 |
| Mrps7 | 0.63415 | 0.011799 |
| Mrps9 | 0.73688 | 0.0019917 |
| Msantd2 | 1.0501 | 0.012332 |
| Msantd4 | 0.5849 | 0.012404 |
| Msi1 | -0.77548 | 0.0024909 |
| Msi2 | -0.91298 | 0.016215 |
| Msmp | -1.1143 | 3.01E-04 |
| Mt-atp6 | 2.449 | 4.24E-06 |
| Mt-atp8 | 2.7582 | 4.16E-10 |
| Mt-co1 | 1.693 | 2.63E-09 |
| Mt-co2 | 2.2007 | 3.75E-06 |
| Mt-cox3 | 2.0212 | 3.70E-05 |
| Mt-cytb | 1.767 | 2.99E-08 |
| Mt-nd1 | 2.2337 | 4.25E-07 |
| Mt-nd2 | 2.3464 | 5.61E-05 |
| Mt-nd4 | 2.305 | 3.02E-14 |
| Mt-nd4l | 1.8312 | 1.27E-05 |
| Mt-nd5 | 1.4759 | 5.23E-10 |
| Mt-nd6 | 2.1683 | 2.40E-28 |
| Mt3 | 1.5462 | 7.11E-08 |
| Mtcl1 | -0.84239 | 0.0026805 |
| Mterf1 | 1.0948 | 2.28E-04 |
| Mterf3 | 0.64678 | 0.018791 |

| | | |
|---|---|---|
| Mterf4 | 0.624 | 0.038803 |
| Mtf1 | -0.5883 | 0.04935 |
| Mtg1 | 0.6345 | 0.01104 |
| Mthfs | 0.70521 | 0.041844 |
| Mtif2 | 0.59087 | 0.035134 |
| Mtmr9 | -1.0058 | 4.31E-06 |
| Mtr | -0.73103 | 0.021028 |
| Mtrf1 | 0.98101 | 2.06E-04 |
| Mtx3 | -1.4174 | 5.38E-05 |
| Mvb12b | -0.66357 | 0.0015957 |
| Mxi1 | 1.3297 | 7.95E-09 |
| Mybbp1a | -0.69352 | 0.0016382 |
| Mybl2 | -1.7203 | 2.85E-04 |
| Myeov2 | 1.1947 | 5.87E-04 |
| Myh6 | -2.6699 | 8.92E-15 |
| Myl12b | -0.51563 | 0.035978 |
| Myo1c | -0.68083 | 0.016312 |
| Myo5b | -0.83556 | 2.94E-04 |
| Myo9a | -1.1741 | 1.39E-07 |
| Myo9b | -1.2689 | 3.37E-09 |
| Mypop | -0.56192 | 0.047477 |
| Myrip | -0.73167 | 0.002157 |
| N4bp1 | -0.92958 | 3.11E-04 |
| N6amt1 | 0.85638 | 0.0016778 |
| N6amt2 | 0.92756 | 4.57E-04 |
| Naa20 | 0.67802 | 0.0027857 |
| Naa25 | -0.57422 | 0.023875 |
| Naa38 | 1.1016 | 4.52E-05 |
| Naaladl1 | -0.98831 | 0.021921 |
| Nabp1 | 0.86995 | 0.019387 |
| Naca | 0.61412 | 0.0044862 |
| Nacc1 | -1.2473 | 3.82E-09 |
| Nanos3 | 1.4368 | 0.021329 |
| Nanp | 0.58343 | 0.029568 |
| Nap1l3 | 0.4995 | 0.034345 |
| Napb | -0.70422 | 0.0014018 |
| Natd1 | -1.1539 | 0.0033127 |
| Nbeal2 | -1.9264 | 1.77E-07 |
| Ncald | 0.48131 | 0.041105 |
| Ncam2 | -0.70965 | 0.0010958 |
| Ncaph | -1.4967 | 3.25E-05 |

| | | |
|---|---|---|
| Ncbp2 | 0.50692 | 0.034078 |
| Nck1 | 0.63523 | 0.038074 |
| Nckipsd | -0.5105 | 0.044626 |
| Ncln | -0.85176 | 7.27E-05 |
| Ncoa3 | -0.74531 | 0.0065666 |
| Ncoa4 | 0.52715 | 0.021024 |
| Ncoa7 | -0.61179 | 0.0087853 |
| Ndnf | 1.651 | 1.07E-06 |
| Ndor1 | -0.57258 | 0.036942 |
| Ndp | 1.0134 | 6.06E-05 |
| Ndrg3 | -0.55656 | 0.031626 |
| Ndst4 | -2.5773 | 0.0014008 |
| Ndufa1 | 1.0632 | 0.0065871 |
| Ndufa11 | 1.0239 | 2.18E-04 |
| Ndufa12 | 0.59337 | 0.0075233 |
| Ndufa2 | 1.1498 | 0.0063852 |
| Ndufa4 | 1.0788 | 0.0019543 |
| Ndufa5 | 0.81384 | 3.77E-04 |
| Ndufa6 | 1.2842 | 1.72E-04 |
| Ndufa7 | 0.91444 | 5.67E-04 |
| Ndufa8 | 0.94282 | 0.00334 |
| Ndufab1 | 0.63492 | 0.024701 |
| Ndufaf5 | 0.96197 | 2.49E-05 |
| Ndufaf7 | 0.52121 | 0.044342 |
| Ndufb10 | 0.99094 | 2.34E-06 |
| Ndufb11 | 0.74035 | 0.0082943 |
| Ndufb2 | 0.90859 | 0.03569 |
| Ndufb3 | 0.98904 | 0.0012831 |
| Ndufb4 | 0.83101 | 0.027349 |
| Ndufb5 | 1.1174 | 8.28E-05 |
| Ndufb6 | 0.76401 | 0.003427 |
| Ndufb7 | 0.74496 | 0.010791 |
| Ndufb8 | 0.75366 | 0.0057068 |
| Ndufb9 | 0.87529 | 0.0018748 |
| Ndufc1 | 1.2069 | 6.60E-07 |
| Ndufc2 | 1.0271 | 3.00E-04 |
| Ndufs3 | 0.52836 | 0.024341 |
| Ndufs4 | 0.75802 | 0.0022251 |
| Ndufs5 | 0.91205 | 0.010688 |
| Ndufs8 | 0.607 | 0.0070413 |
| Ndufv2 | 0.75851 | 3.00E-04 |

| | | |
|---|---|---|
| Ndufv3-ps1 | 0.60339 | 0.01407 |
| Neb | -0.98728 | 0.041504 |
| Nebl | -0.63328 | 0.014843 |
| Necab2 | -0.49977 | 0.040132 |
| Nedd4l | -0.63764 | 0.0046289 |
| Nedd8 | 0.88441 | 0.0066714 |
| Negr1 | -0.6524 | 0.0067229 |
| Nek2l1 | -1.3368 | 0.032178 |
| Nell1 | 0.61303 | 0.018701 |
| Nenf | 1.3254 | 6.91E-04 |
| Neto1 | -0.90605 | 4.14E-05 |
| Neurl1 | -0.54173 | 0.033138 |
| Neurl1b | -1.7461 | 4.43E-12 |
| NEWGENE_1310139 | -2.6262 | 4.30E-05 |
| NEWGENE_1582994 | 0.58036 | 0.046623 |
| Nf2 | -0.54928 | 0.013999 |
| Nfam1 | -0.81963 | 0.011811 |
| Nfasc | -0.89431 | 3.33E-05 |
| Nfe2l1 | -0.65431 | 0.032444 |
| Nfia | -1.1571 | 1.09E-05 |
| Nfib | -1.6129 | 1.03E-07 |
| Nfkbia | 0.63517 | 0.029835 |
| Nfu1 | 0.58509 | 0.016734 |
| Nfyc | 0.76955 | 0.0017035 |
| Ngdn | 0.59821 | 0.015286 |
| Ngfrap1 | 0.61352 | 0.0043113 |
| Nhlrc2 | -0.79658 | 0.0070707 |
| Nhp2 | 0.66201 | 0.014971 |
| Nhs | -0.86338 | 0.045701 |
| Nhsl2 | -1.4813 | 1.20E-07 |
| Nid1 | -1.4542 | 0.0021589 |
| Nif3l1 | 0.63705 | 0.024921 |
| Nin | -0.74632 | 0.019755 |
| Nip7 | 0.60379 | 0.023732 |
| Nipa1 | -0.52652 | 0.036926 |
| Nipal2 | 1.4296 | 0.0068193 |
| Nipal3 | -1.2984 | 6.06E-08 |
| Nipsnap3b | 0.59841 | 0.048447 |
| Nkain1 | -0.68773 | 0.011449 |
| Nkain3 | -0.82842 | 0.047633 |
| Nkain4 | 1.1697 | 3.12E-06 |

| | | |
|---|---|---|
| Nkd1 | -1.0868 | 0.0010683 |
| Nkd2 | 0.65042 | 0.034119 |
| Nlrc5 | -1.8321 | 0.013818 |
| Nmbr | 2.2172 | 0.014645 |
| Nme3 | 1.1396 | 1.38E-05 |
| Nmnat2 | -0.92966 | 2.67E-05 |
| Nmral1 | 0.67783 | 0.037202 |
| Nmrk1 | 0.87195 | 0.0084228 |
| Nob1 | 0.57783 | 0.015844 |
| Nol3 | 0.69291 | 0.043496 |
| Nol4l | -1.5258 | 9.45E-09 |
| Nol6 | -0.51642 | 0.027236 |
| Nol7 | 0.62992 | 0.0051967 |
| Nol9 | -0.54277 | 0.026524 |
| Nom1 | -0.54645 | 0.038237 |
| Nop10 | 1.1737 | 0.0086784 |
| Nop56 | 0.54627 | 0.017481 |
| Notch1 | -1.4154 | 5.74E-06 |
| Notch2 | -1.3797 | 0.0018087 |
| Notch3 | -2.2019 | 2.98E-11 |
| Nova1 | -0.82194 | 0.0021147 |
| Nova2 | -1.3526 | 5.23E-10 |
| Npap60 | -0.69249 | 0.0036065 |
| Npas1 | -0.63445 | 0.049799 |
| Npc2 | 0.59654 | 0.012336 |
| Nphp4 | -0.92337 | 6.34E-04 |
| Npm1 | 0.5635 | 0.011565 |
| Nptx1 | -2.0729 | 1.82E-17 |
| Npy1r | 1.0406 | 0.010148 |
| Nr2c2 | -1.2656 | 1.07E-08 |
| Nr3c2 | -0.82003 | 0.0074873 |
| Nrarp | -1.6368 | 0.0052939 |
| Nrbp2 | -1.0905 | 0.0010152 |
| Nrg1 | -0.96149 | 0.010568 |
| Nrn1 | 0.8277 | 1.38E-04 |
| Nrp2 | -1.5986 | 2.24E-05 |
| Nrsn1 | 1.9195 | 4.93E-04 |
| Nsa2 | 0.69521 | 0.0022039 |
| Nsl1 | -0.97148 | 0.037545 |
| Nsmce1 | 0.6778 | 0.0058241 |
| Nsrp1 | 0.81874 | 0.0014485 |

| | | |
|---|---|---|
| Nt5c1a | -1.3381 | 0.046656 |
| Nt5c3a | 0.88448 | 1.99E-04 |
| Nt5dc1 | 0.7964 | 0.0048632 |
| Nt5dc3 | -1.512 | 1.70E-12 |
| Ntn1 | -1.6142 | 0.0044112 |
| Ntn3 | -1.3227 | 9.96E-05 |
| Ntng2 | -1.1737 | 0.0021064 |
| Ntpcr | 0.69282 | 0.0042707 |
| Ntsr2 | 1.4001 | 0.029877 |
| Nuak1 | -0.96307 | 1.38E-04 |
| Nubpl | 0.65119 | 0.02928 |
| Nucb2 | 0.80709 | 0.0012743 |
| Nucks1 | -0.55398 | 0.017669 |
| Nudt11 | -1.1826 | 0.042342 |
| Nudt2 | 0.99143 | 1.03E-05 |
| Nudt6 | 1.0635 | 2.33E-04 |
| Nudt7 | 1.3061 | 0.010965 |
| Numa1 | -0.59667 | 0.0072129 |
| Numbl | -0.52513 | 0.025916 |
| Nup188 | -0.59521 | 0.021441 |
| Nup210 | -0.75069 | 0.0013301 |
| Nup43 | 0.75591 | 0.011481 |
| Nupr1l | 1.0689 | 0.0043866 |
| Nutf2 | 0.61411 | 0.0065903 |
| Nxph4 | -1.6481 | 0.0054144 |
| Nxt1 | 0.88078 | 0.0070875 |
| Nxt2 | 1.234 | 4.79E-05 |
| Nyap2 | -1.1175 | 9.64E-05 |
| Nynrin | -0.79219 | 0.0057794 |
| Oat | 0.62237 | 0.0044309 |
| Ociad1 | 0.68923 | 0.0011568 |
| Ocln | 1.7708 | 0.0011026 |
| Odf3 | -1.4994 | 0.045612 |
| Ogfod1 | -0.83085 | 8.03E-04 |
| Olig1 | -1.0056 | 1.59E-06 |
| Onecut1 | -1.5277 | 0.041145 |
| Opa1 | -0.51274 | 0.027903 |
| Opcml | -1.292 | 1.38E-10 |
| Orai2 | -0.71052 | 0.022083 |
| Orc5 | 0.5684 | 0.020809 |
| Orc6 | 0.61133 | 0.045153 |

| | | |
|---|---|---|
| Ormdl2 | 0.82095 | 0.001126 |
| Osbp2 | -0.57839 | 0.012764 |
| Osbpl5 | -0.69393 | 0.0048138 |
| Osbpl6 | -0.69 | 0.0039145 |
| Osbpl9 | 0.72255 | 0.0023182 |
| Oser1 | 0.73273 | 0.0043541 |
| Ost4 | 1.2658 | 0.0016009 |
| Ostc | 0.83494 | 1.61E-04 |
| Ostm1 | -0.57348 | 0.014974 |
| Otof | -2.3111 | 3.05E-20 |
| Otud3 | -0.98284 | 0.004524 |
| Otud4 | -0.87515 | 5.16E-04 |
| Otud7a | -0.55322 | 0.024902 |
| Ovol1 | 2.3353 | 0.021075 |
| Ovol2 | 1.2186 | 0.0074423 |
| Oxld1 | 0.60321 | 0.041341 |
| P3h3 | -1.1034 | 0.027209 |
| Pabpc1 | 0.56106 | 0.013855 |
| Paf1 | 0.59015 | 0.012611 |
| Pafah2 | -1.0944 | 0.036194 |
| Paics | 0.65106 | 0.0053271 |
| Paip2 | 0.88283 | 2.91E-05 |
| Pak3 | -1.3122 | 3.25E-05 |
| Pak7 | 0.5992 | 0.022436 |
| Palm | -0.5517 | 0.038094 |
| Palm2 | -1.3991 | 6.11E-07 |
| Pam16 | 0.92076 | 0.0036113 |
| Pan2 | -0.75996 | 0.0058277 |
| Pank3 | -1.0347 | 6.61E-04 |
| Panx1 | -0.69803 | 0.0035556 |
| Papd4 | 0.62918 | 0.016237 |
| Pappa | -1.8301 | 0.035978 |
| Paqr8 | -0.96237 | 0.0089218 |
| Paqr9 | -1.3306 | 2.93E-05 |
| Pard3 | -0.69683 | 0.0087901 |
| Pard3b | -1.3342 | 0.0079973 |
| Parm1 | -0.76186 | 0.0046792 |
| Parva | -0.72221 | 0.02497 |
| Pbrm1 | -0.93992 | 3.84E-05 |
| Pbx1 | -0.89805 | 3.91E-05 |
| Pc | -0.50866 | 0.042342 |

| | | |
|---|---|---|
| Pcbd2 | 1.2875 | 9.27E-06 |
| Pcbp3 | -0.6204 | 0.011967 |
| Pcdh11x | -0.99058 | 0.037095 |
| Pcdh15 | -1.9746 | 2.47E-04 |
| Pcdh17 | -1.151 | 2.74E-08 |
| Pcdh19 | -0.99851 | 5.14E-06 |
| Pcdha13 | -1.1871 | 8.04E-08 |
| Pcdha2 | -1.0475 | 4.96E-04 |
| Pcdha3 | -0.87026 | 0.0029171 |
| Pcdhb10 | 0.79471 | 0.020878 |
| Pcdhb2 | -2.0688 | 5.32E-05 |
| Pcdhga3 | -1.0034 | 0.0010152 |
| Pcdhga4 | -0.93249 | 0.0046913 |
| Pcdhga5 | -1.028 | 0.0023144 |
| Pcdhga7 | -1.0318 | 4.22E-05 |
| Pcdhga8 | -1.0731 | 0.0052198 |
| Pcdhgb5 | -0.76563 | 0.0094533 |
| Pcdhgb6 | -0.86307 | 0.044745 |
| Pcdhgb8 | -0.98469 | 0.0011314 |
| Pcgf3 | -0.61397 | 0.013925 |
| Pcgf6 | 0.67184 | 0.019114 |
| Pcmtd2 | 0.83534 | 0.0016554 |
| Pcnp | 0.67477 | 0.0023606 |
| Pcnxl2 | -0.8529 | 1.45E-04 |
| Pcnxl3 | -1.0487 | 8.89E-07 |
| Pcsk1 | 0.9142 | 0.0017081 |
| Pcyt1b | -1.1251 | 1.40E-06 |
| Pcyt2 | -0.51705 | 0.038031 |
| Pdcd10 | 0.54206 | 0.020147 |
| Pdcd11 | -0.6764 | 0.002922 |
| Pdcd2 | 0.86382 | 2.43E-04 |
| Pdcd5 | 1.1041 | 4.29E-07 |
| Pdcd6 | 0.69728 | 0.0021802 |
| Pdcl | 0.54392 | 0.026595 |
| Pde10a | -1.1628 | 3.30E-07 |
| Pde3a | -0.9796 | 0.0044156 |
| Pde4a | -0.93612 | 7.91E-05 |
| Pde4d | -1.01 | 2.74E-05 |
| Pdgfb | -1.1008 | 3.34E-05 |
| Pdia4 | 0.96791 | 6.74E-05 |
| Pdia6 | 0.61431 | 0.0062536 |

| | | |
|---|---|---|
| Pdlim4 | 0.59676 | 0.01793 |
| Pdlim5 | -0.84823 | 0.045701 |
| Pdp1 | 0.76217 | 9.46E-04 |
| Pdp2 | -0.77125 | 0.031521 |
| Pdpk1 | -1.3282 | 2.29E-09 |
| Pdpr | -1.4969 | 1.90E-09 |
| Pdrg1 | 0.66412 | 0.01879 |
| Pds5a | -0.72599 | 0.0021699 |
| Pdxdc1 | -0.76259 | 0.0054958 |
| Pdxk | -1.1607 | 9.54E-05 |
| Pdzd3 | -0.79626 | 0.010505 |
| Peak1 | -1.6286 | 1.66E-12 |
| Peli2 | -0.81035 | 0.013393 |
| Per3 | -0.70484 | 0.016187 |
| Pet100 | 0.84227 | 2.41E-04 |
| Pex12 | 0.65517 | 0.038826 |
| Pex2 | 0.71373 | 0.0047299 |
| Pex5l | -1.4595 | 2.22E-11 |
| Pex6 | -0.54019 | 0.023455 |
| Pfas | -0.74055 | 0.0011366 |
| Pfdn1 | 1.3445 | 1.89E-06 |
| Pfdn2 | 1.1159 | 1.04E-05 |
| Pfdn4 | 1.0576 | 0.0016585 |
| Pfdn5 | 1.1439 | 3.69E-08 |
| Pfkfb2 | -0.88912 | 1.31E-04 |
| Pfkfb3 | -1.0967 | 0.025825 |
| Pfkl | -0.68777 | 0.0047444 |
| Pgam2 | -1.0574 | 1.67E-04 |
| Pgap1 | -0.90577 | 0.0048009 |
| Pgpep1 | -0.72187 | 0.011438 |
| Pgrmc1 | 0.79266 | 1.94E-04 |
| Phactr2 | -1.2921 | 0.0012407 |
| Phactr4 | -1.2066 | 6.69E-04 |
| Phax | 0.5371 | 0.029752 |
| Phb | 0.47496 | 0.041752 |
| Phc2 | -0.59504 | 0.0083778 |
| Phc3 | -1.6674 | 4.15E-10 |
| Phf1 | -0.59503 | 0.030897 |
| Phf2 | -0.73309 | 0.0014255 |
| Phf21b | -0.98487 | 0.026797 |
| Phf5a | 0.94345 | 3.22E-05 |

| | | |
|---|---|---|
| Phip | -0.74964 | 0.0029896 |
| Phka2 | -1.9747 | 7.30E-06 |
| Phlda1 | -0.89339 | 0.017086 |
| Phlda3 | -0.68826 | 0.0063765 |
| Phlpp2 | -0.696 | 0.0044484 |
| Phospho1 | -0.93805 | 0.0017605 |
| Phospho2 | 0.65764 | 0.0058384 |
| Phpt1 | 0.8695 | 3.80E-04 |
| Pi4k2a | -0.83752 | 5.68E-04 |
| Pigc | 0.54688 | 0.033249 |
| Pigf | 0.67512 | 0.030718 |
| Pigk | 0.5203 | 0.038419 |
| Pign | -0.99418 | 0.0011759 |
| Pigp | 0.96815 | 7.39E-05 |
| Pigu | 0.82005 | 7.30E-04 |
| Pigw | 1.1322 | 3.14E-05 |
| Pik3c2b | -0.97234 | 0.0020204 |
| Pik3r1 | -0.86924 | 9.60E-05 |
| Pik3r3 | -1.0501 | 0.016816 |
| Pik3r5 | -3.8137 | 0.030346 |
| Pin4 | 1.3516 | 2.69E-08 |
| Pip4k2b | -1.0959 | 2.22E-07 |
| Pir | 1.0016 | 0.031537 |
| Pitpna | -0.48254 | 0.040499 |
| Pitpnm3 | -1.3161 | 1.04E-04 |
| Pkig | 0.95029 | 4.76E-05 |
| Pkn1 | -0.66366 | 0.0047279 |
| Pknox1 | -1.3263 | 9.01E-05 |
| Pla2g16 | 1.7246 | 6.09E-05 |
| Pla2g4b | -1.1691 | 0.0095923 |
| Pla2g4e | -0.92682 | 0.042204 |
| Pla2g7 | 2.2631 | 2.90E-05 |
| Plagl1 | -1.7 | 9.31E-05 |
| Plcb3 | -0.68991 | 0.03817 |
| Plcxd3 | -1.3888 | 2.22E-07 |
| Pld5 | 1.7424 | 2.44E-05 |
| Plekha1 | 0.57201 | 0.012792 |
| Plekhf2 | 0.64994 | 0.017661 |
| Plekhg1 | -0.73333 | 0.023452 |
| Plekhg2 | -1.1883 | 0.0065532 |
| Plekhg4 | -2.4213 | 0.0016697 |

| | | |
|---|---|---|
| Plekhg5 | -0.95426 | 3.83E-05 |
| Plekhm1 | -0.69626 | 0.0043456 |
| Plekhm2 | -0.51788 | 0.027545 |
| Plekhm3 | -1.2916 | 6.27E-05 |
| Plekho2 | -1.5865 | 2.88E-05 |
| Plgrkt | 0.73614 | 0.0082385 |
| Plk1 | -1.1348 | 0.0016471 |
| Plk3 | -0.81481 | 0.023507 |
| Plk4 | -0.86487 | 0.049953 |
| Plrg1 | 0.80245 | 4.57E-04 |
| Pls3 | 0.90248 | 3.61E-05 |
| Pltp | 0.83845 | 0.024431 |
| Plxnc1 | -1.0058 | 6.45E-04 |
| Pm20d2 | -1.1972 | 6.99E-04 |
| Pmel | -1.1874 | 0.023578 |
| Pml | -1.1994 | 0.048548 |
| Pmpcb | 0.54791 | 0.022192 |
| Pnkd | -0.48528 | 0.041862 |
| Pnpla6 | -0.76758 | 6.24E-04 |
| Pnpla8 | 0.83227 | 7.33E-05 |
| Pnrc2 | 0.61165 | 0.0064081 |
| Pofut1 | -0.71505 | 0.022838 |
| Poglut1 | 0.54848 | 0.02922 |
| Pold1 | -1.0739 | 9.42E-04 |
| Pole4 | 0.95383 | 1.62E-05 |
| Polg | -1.0106 | 1.18E-04 |
| Polr2d | 0.93945 | 9.77E-05 |
| Polr2j | 0.81564 | 0.0069524 |
| Polr2k | 1.1454 | 9.24E-04 |
| Polr3k | 0.7915 | 2.97E-04 |
| Polrmt | -0.82714 | 8.60E-04 |
| Pom121 | -0.85596 | 1.21E-04 |
| Pomgnt1 | -0.55899 | 0.020992 |
| Pomt2 | -0.59899 | 0.022144 |
| Pon2 | 1.0339 | 1.58E-06 |
| Pop7 | 0.7856 | 0.0018777 |
| Pou6f1 | -0.65842 | 0.017135 |
| Pp2d1 | 0.97009 | 0.0072309 |
| Ppa2 | 0.73737 | 0.0022259 |
| Ppard | -0.97158 | 3.25E-04 |
| Ppargc1b | -1.8705 | 8.60E-09 |

| | | |
|---|---|---|
| Ppfia4 | -0.5382 | 0.02363 |
| Ppib | 0.83626 | 8.24E-05 |
| Ppid | 0.56451 | 0.014229 |
| Ppie | 0.65338 | 0.017882 |
| Ppil1 | 0.73334 | 0.0073275 |
| Ppm1f | -0.73184 | 0.0029814 |
| Ppm1h | -1.1264 | 3.20E-07 |
| Ppm1k | 0.51458 | 0.037617 |
| Ppm1l | -1.9882 | 1.86E-11 |
| Ppp1cb | 0.55854 | 0.011201 |
| Ppp1r16b | -1.1016 | 9.78E-07 |
| Ppp1r36 | 1.7014 | 0.014424 |
| Ppp1r37 | -0.57105 | 0.012561 |
| Ppp1r3c | 0.60757 | 0.022722 |
| Ppp1r3e | -1.3816 | 0.0015346 |
| Ppp1r9a | -2.1184 | 1.66E-22 |
| Ppp2cb | 0.53014 | 0.018653 |
| Ppp2r1b | 0.67989 | 0.0090858 |
| Ppp2r3c | 0.74063 | 0.0040381 |
| Ppp2r5d | -0.50905 | 0.031219 |
| Ppp3r1 | 0.53802 | 0.014376 |
| Ppp4r2 | 0.49638 | 0.034488 |
| Ppp4r3b | 0.56389 | 0.014787 |
| Ppp6r1 | -0.76372 | 3.09E-04 |
| Pprc1 | -0.8923 | 3.10E-04 |
| Ppt1 | 0.85384 | 5.75E-05 |
| Pragmin | -0.72647 | 0.0030203 |
| Prdm10 | -0.78283 | 0.023265 |
| Prdm11 | -1.5796 | 6.79E-04 |
| Prdm15 | -1.1158 | 1.85E-04 |
| Prdx1 | 0.8425 | 5.00E-05 |
| Prdx3 | 0.51326 | 0.026551 |
| Prdx4 | 1.5063 | 1.43E-10 |
| Prdx6 | 1.2746 | 3.93E-06 |
| Prelid3b | 0.82571 | 0.0019205 |
| Prep | -0.51689 | 0.033222 |
| Prkaa2 | -1.2826 | 1.41E-07 |
| Prkab2 | -0.85286 | 0.0035751 |
| Prkar2a | -1.2064 | 8.96E-08 |
| Prkca | -1.6068 | 3.30E-07 |
| Prmt10 | 0.9528 | 7.14E-05 |

| | | |
|---|---|---|
| Prmt6 | 0.66008 | 0.02818 |
| Prorsd1 | 1.2806 | 0.030586 |
| Proser1 | -0.66521 | 0.011472 |
| Prpf3 | 0.74956 | 0.0022387 |
| Prpf38a | 0.69186 | 0.0039145 |
| Prr11 | -1.601 | 0.012819 |
| Prr14l | -1.1405 | 7.80E-07 |
| Prr16 | 1.5771 | 1.21E-05 |
| Prr36 | -1.1787 | 4.85E-08 |
| Prr5 | -0.69477 | 0.0089969 |
| Prrc1 | -0.6645 | 0.028209 |
| Prrc2a | -1.2953 | 4.21E-11 |
| Prrc2b | -2.0085 | 1.09E-20 |
| Prrt1 | -0.90128 | 0.0054932 |
| Prrt3 | -0.57095 | 0.022043 |
| Prrt4 | -0.54628 | 0.044747 |
| Prtfdc1 | 0.71554 | 0.0025558 |
| Prune2 | -0.68724 | 0.00334 |
| Psd3 | -1.6721 | 1.16E-15 |
| Psenen | 0.96351 | 6.99E-05 |
| Psma1 | 0.64971 | 0.0034528 |
| Psma2 | 0.73787 | 6.01E-04 |
| Psma3 | 0.62632 | 0.0045705 |
| Psma4 | 0.70651 | 0.0013244 |
| Psma5 | 0.66316 | 0.00283 |
| Psma6 | 0.69832 | 0.0012407 |
| Psmb1 | 0.73268 | 5.56E-04 |
| Psmb2 | 0.59074 | 0.0078525 |
| Psmb4 | 0.79555 | 1.55E-04 |
| Psmb5 | 0.56116 | 0.017658 |
| Psmb6 | 0.71437 | 9.51E-04 |
| Psmc2 | 0.64937 | 0.0034519 |
| Psmc5 | 0.48845 | 0.034964 |
| Psmc6 | 0.65459 | 0.0033544 |
| Psmd3 | -0.48921 | 0.03545 |
| Psme1 | 0.75732 | 0.0020382 |
| Psme2 | 0.88188 | 4.51E-04 |
| Psmg1 | 0.67769 | 0.0035738 |
| Psmg2 | 0.62541 | 0.021255 |
| Psmg3 | 1.045 | 3.13E-05 |
| Psmg4 | 1.1968 | 1.24E-05 |

| | | |
|---|---|---|
| Pstpip2 | -1.8203 | 2.03E-04 |
| Ptar1 | -1.308 | 2.02E-04 |
| Ptchd2 | -0.83205 | 0.0018489 |
| Ptger1 | -0.98536 | 2.70E-04 |
| Ptk7 | -0.68253 | 0.010917 |
| Ptpmt1 | 0.70635 | 0.0045376 |
| Ptpn1 | -0.59273 | 0.010388 |
| Ptpn14 | -2.7621 | 3.43E-12 |
| Ptpn23 | -1.0824 | 7.44E-07 |
| Ptpn4 | -0.87323 | 0.010987 |
| Ptpn9 | -0.64403 | 0.0095955 |
| Ptpre | -0.63856 | 0.031671 |
| Ptprn | -0.58437 | 0.013588 |
| Ptprz1 | -0.79174 | 0.0010712 |
| Pts | 0.85031 | 0.0015922 |
| Pttg1ip | 0.58847 | 0.013837 |
| Pura | -0.85505 | 6.99E-04 |
| Purg | -1.3749 | 1.16E-06 |
| Pvrl1 | -1.4058 | 3.91E-09 |
| Pxmp2 | 1.0913 | 1.01E-04 |
| Pxn | -0.84561 | 0.0011177 |
| Pycr1 | -0.79483 | 0.021464 |
| Pyurf | 0.63726 | 0.04242 |
| Qser1 | -0.6028 | 0.042596 |
| Rab11fip2 | -0.74082 | 0.0051964 |
| Rab11fip4 | -1.0905 | 1.94E-07 |
| Rab13 | 0.75507 | 0.015838 |
| Rab18 | 0.54062 | 0.017989 |
| Rab23 | -0.64313 | 0.010868 |
| Rab28 | 0.8695 | 5.14E-05 |
| Rab29 | 0.72733 | 0.0075157 |
| Rab36 | -0.52305 | 0.045921 |
| Rab3c | -1.4111 | 1.22E-10 |
| Rab3d | -0.78627 | 0.0027211 |
| Rab9a | 0.68923 | 0.0059067 |
| Rabac1 | 0.78647 | 0.0021981 |
| Rabgap1l | -1.2393 | 1.20E-07 |
| Rabggtb | 0.6422 | 0.0031107 |
| Rad17 | 0.57855 | 0.035646 |
| Rad21 | 0.48851 | 0.03944 |
| Rad54l2 | -0.98919 | 1.47E-04 |

| | | |
|---|---|---|
| Ralgapa2 | -1.1465 | 9.45E-06 |
| Ralgps1 | -1.475 | 2.83E-09 |
| Ralgps2 | -0.89859 | 0.0054042 |
| Ralyl | 0.73247 | 0.041903 |
| Rangrf | 0.96065 | 8.89E-04 |
| Rap1a | 0.64234 | 0.004346 |
| Rap2b | -0.9168 | 3.03E-04 |
| Rapgef6 | -0.73812 | 0.0044485 |
| Raph1 | -1.3415 | 8.63E-08 |
| Rars | 0.74894 | 0.0011326 |
| Rasa1 | -0.58575 | 0.011374 |
| Rasa3 | -0.95386 | 8.87E-04 |
| Rasal1 | -0.77046 | 3.65E-04 |
| Rasal2 | -0.95778 | 2.64E-04 |
| Rasd2 | -1.0621 | 2.02E-06 |
| Rassf8 | -1.0007 | 0.016674 |
| Raver1 | -0.82312 | 2.81E-04 |
| Raver2 | -0.64085 | 0.0062645 |
| Rb1cc1 | 0.52819 | 0.021785 |
| Rbak | 0.64384 | 0.046882 |
| Rbbp9 | 0.87386 | 4.99E-04 |
| Rbfox2 | -1.616 | 1.19E-10 |
| Rbfox3 | -0.88978 | 0.0012584 |
| Rbks | 0.76696 | 0.025304 |
| Rbm12b | 0.79666 | 0.0015725 |
| Rbm15 | -1.2028 | 0.0011256 |
| Rbm15b | -0.68431 | 0.0028549 |
| Rbm19 | -1.3233 | 6.78E-06 |
| Rbm25 | 0.72832 | 8.85E-04 |
| Rbm25l1 | -0.67243 | 0.012156 |
| Rbm27 | -0.67618 | 0.0066425 |
| Rbm33 | -0.79011 | 0.0041916 |
| Rbm48 | 0.99138 | 3.66E-04 |
| Rbm7 | 0.88374 | 0.0014311 |
| Rbm8a | 0.79 | 3.28E-04 |
| Rbpjl | -2.2618 | 0.039422 |
| Rcc1 | -1.0819 | 1.32E-04 |
| Rcc2 | -1.3102 | 2.04E-06 |
| Rchy1 | 0.74631 | 0.0014578 |
| Rcn2 | 0.78481 | 0.00022135 |
| Rcor1 | -1.3313 | 9.78E-04 |

| | | |
|---|---|---|
| Rell2 | -1.0528 | 0.0042455 |
| Rer1 | 0.56443 | 0.016275 |
| Rexo1 | -0.89738 | 3.24E-05 |
| Rfwd2 | 0.55173 | 0.023099 |
| Rfx1 | -0.98834 | 9.66E-05 |
| Rfx3 | -1.3152 | 1.32E-08 |
| RGD1304567 | 0.88819 | 2.05E-04 |
| RGD1304694 | -1.0206 | 4.47E-06 |
| RGD1304704 | 0.62329 | 0.0061554 |
| RGD1304884 | -1.158 | 1.21E-06 |
| RGD1305089 | 0.64703 | 0.033614 |
| RGD1305455 | -0.62312 | 0.0087625 |
| RGD1305645 | 2.4856 | 9.21E-11 |
| RGD1306063 | 1.0567 | 8.48E-06 |
| RGD1306215 | 0.58697 | 0.024287 |
| RGD1306233 | 1.0325 | 0.045701 |
| RGD1306271 | -2.0345 | 3.28E-20 |
| RGD1306941 | 0.56469 | 0.02363 |
| RGD1307929 | -0.48996 | 0.032372 |
| RGD1308147 | 0.84295 | 4.56E-04 |
| RGD1308430 | 0.64893 | 0.0059801 |
| RGD1308706 | 0.76742 | 0.0039145 |
| RGD1309534 | 0.64813 | 0.022046 |
| RGD1309651 | 1.9739 | 0.043805 |
| RGD1309730 | 1.2651 | 6.75E-07 |
| RGD1309779 | 0.85878 | 0.045298 |
| RGD1310081 | -0.63753 | 0.0083778 |
| RGD1310429 | -0.6038 | 0.013971 |
| RGD1310553 | 0.60229 | 0.01233 |
| RGD1310769 | 0.74794 | 0.018974 |
| RGD1310819 | -0.60049 | 0.0071179 |
| RGD1311745 | 0.86226 | 0.0014495 |
| RGD1311747 | 0.83129 | 2.21E-04 |
| RGD1359158 | 0.94107 | 3.48E-04 |
| RGD1560065 | 0.71086 | 0.0013939 |
| RGD1560108 | -0.76309 | 5.02E-04 |
| RGD1560175 | -0.8194 | 0.002043 |
| RGD1560394 | 1.3284 | 0.0091181 |
| RGD1560398 | 0.56203 | 0.021524 |
| RGD1561034 | -0.92005 | 0.0013702 |
| RGD1561149 | -1.3419 | 6.22E-05 |

| | | |
|---|---|---|
| RGD1561444 | 0.83687 | 0.0047588 |
| RGD1561590 | 0.51661 | 0.025827 |
| RGD1561897 | -1.3712 | 1.28E-09 |
| RGD1561931 | -1.9515 | 1.89E-18 |
| RGD1562136 | 0.53479 | 0.026663 |
| RGD1562310 | -0.92217 | 1.31E-04 |
| RGD1562339 | -1.2539 | 4.68E-07 |
| RGD1562747 | 0.64766 | 0.017882 |
| RGD1562987 | 0.80547 | 1.22E-04 |
| RGD1563441 | 0.93667 | 0.020916 |
| RGD1563986 | -1.3782 | 6.30E-09 |
| RGD1564053 | -0.55015 | 0.031772 |
| RGD1564093 | 0.54525 | 0.019615 |
| RGD1565033 | 0.67098 | 0.022852 |
| RGD1565059 | 1.1582 | 0.0023285 |
| RGD1565363 | 1.1574 | 1.21E-06 |
| RGD1565616 | -0.6467 | 0.0037373 |
| RGD1565641 | 1.2056 | 5.03E-07 |
| RGD1565685 | 0.67761 | 0.014534 |
| RGD1565784 | 0.59155 | 0.013911 |
| RGD1566099 | 0.75232 | 0.010638 |
| RGD1566239 | 0.75291 | 0.020051 |
| RGD1566320 | 0.93081 | 6.76E-05 |
| RGD1566386 | 0.79032 | 0.0043866 |
| RGD735065 | 0.49228 | 0.042691 |
| Rgl2 | -0.9542 | 0.0040814 |
| Rgma | -0.71919 | 0.008343 |
| Rgmb | -0.59259 | 0.014911 |
| Rgp1 | -1.1828 | 1.78E-06 |
| Rgs10 | 1.3225 | 0.0062536 |
| Rgs14 | -0.84749 | 5.27E-04 |
| Rgs2 | 0.89184 | 6.02E-05 |
| Rgs6 | -0.66776 | 0.01276 |
| Rgs7 | 0.48888 | 0.035938 |
| Rhbdf1 | -1.1885 | 1.10E-06 |
| Rheb | 0.54226 | 0.014435 |
| Rhot2 | -0.52233 | 0.030172 |
| Rhpn2 | 0.92383 | 0.026663 |
| Ric1 | -0.8002 | 0.001247 |
| Rictor | -0.86134 | 5.10E-04 |
| Rimkla | -1.0348 | 1.49E-04 |

| | | |
|---|---|---|
| Rims1 | -0.77148 | 3.92E-04 |
| Rims4 | -1.6481 | 3.38E-13 |
| Rlim | -1.2229 | 1.57E-05 |
| Rmdn1 | 1.0396 | 2.51E-05 |
| Rmi1 | 0.79308 | 0.0016852 |
| Rmi2 | -2.3922 | 1.75E-04 |
| Rmnd1 | 0.58233 | 0.019989 |
| Rmnd5a | -0.86934 | 3.05E-04 |
| Rn50_1_1359.1 | 1.3284 | 2.37E-07 |
| Rn50_10_0635.1 | #NAME? | 5.15E-05 |
| Rn50_10_0702.4 | 1.2946 | 0.0029309 |
| Rn50_13_0822.1 | -2.1829 | 2.69E-05 |
| Rn50_13_0829.4 | -0.64962 | 0.010107 |
| Rn50_13_0839.5 | 0.90842 | 0.002925 |
| Rn50_5_0825.2 | 1.6677 | 0.020187 |
| Rn50_X_0635.3 | 1.0651 | 5.94E-04 |
| Rn50_X_0667.2 | 1.6823 | 0.0043255 |
| Rn50_X_0746.3 | 1.3967 | 1.18E-05 |
| Rn50_X_0747.1 | -1.0237 | 7.67E-06 |
| Rn50_X_0752.3 | -7.1215 | 6.07E-04 |
| Rn60_14_0788.1 | 1.5268 | 3.73E-09 |
| Rnaseh1 | 0.74232 | 0.011741 |
| Rnasek | 0.96963 | 0.0018981 |
| Rnaset2 | 1.2513 | 6.01E-04 |
| Rnd2 | 1.3319 | 7.52E-05 |
| Rnd3 | 0.9593 | 4.15E-04 |
| Rnf103 | 0.55064 | 0.017728 |
| Rnf112 | -0.48705 | 0.041844 |
| Rnf123 | -0.75907 | 6.69E-04 |
| Rnf126 | -0.55665 | 0.018796 |
| Rnf13 | 0.90538 | 2.72E-05 |
| Rnf138 | 0.90187 | 1.25E-04 |
| Rnf139 | 0.71278 | 0.0027371 |
| Rnf144a | -1.2348 | 2.50E-07 |
| Rnf146 | 0.74327 | 8.03E-04 |
| Rnf150 | -1.2416 | 2.73E-06 |
| Rnf157 | -1.4427 | 8.40E-08 |
| Rnf165 | -0.9295 | 1.14E-05 |
| Rnf169 | -0.81059 | 0.023556 |
| Rnf181 | 0.64826 | 0.010266 |
| Rnf182 | -0.79596 | 0.015543 |

| | | |
|---|---|---|
| Rnf187 | -0.50944 | 0.022844 |
| Rnf2 | 0.53165 | 0.026364 |
| Rnf213 | -1.2732 | 0.0052022 |
| Rnf214 | -0.84516 | 4.52E-04 |
| Rnf38 | -0.93134 | 4.27E-04 |
| Rnf40 | -0.51705 | 0.031772 |
| Rnf44 | -1.2761 | 5.48E-09 |
| Rnf7 | 1.0096 | 2.89E-06 |
| Rnpepl1 | -0.6698 | 0.019572 |
| Robo1 | -1.1935 | 3.18E-05 |
| Rock1 | 0.71847 | 0.014725 |
| Rock2 | -0.61319 | 0.0078233 |
| Rpain | 0.71083 | 0.017882 |
| Rpap1 | -0.77455 | 0.0023144 |
| Rpe | 0.64322 | 0.0081319 |
| Rpf2 | 0.71374 | 0.0033544 |
| Rpgrip1 | 0.75459 | 0.037126 |
| Rpl10a | 0.80392 | 9.78E-04 |
| Rpl11 | 1.236 | 9.45E-10 |
| Rpl12 | 1.067 | 2.11E-07 |
| Rpl14 | 1.029 | 1.11E-05 |
| Rpl15 | 0.87527 | 2.15E-05 |
| Rpl17 | 1.0805 | 7.78E-06 |
| Rpl18 | 0.98047 | 1.36E-05 |
| Rpl18a | 1.1444 | 1.01E-06 |
| Rpl19 | 1.155 | 7.19E-04 |
| Rpl21 | 1.351 | 1.70E-11 |
| Rpl21 | 1.1384 | 0.0081883 |
| Rpl22l1 | 1.1132 | 0.00524 |
| Rpl23 | 1.2897 | 1.39E-06 |
| Rpl23a | 1.3311 | 4.98E-07 |
| Rpl24 | 1.0821 | 2.80E-04 |
| Rpl26 | 1.4479 | 4.02E-06 |
| Rpl26-ps2 | 1.4106 | 0.029461 |
| Rpl27 | 1.1202 | 0.002634 |
| Rpl27a | 1.2445 | 2.25E-04 |
| Rpl28 | 1.1481 | 5.74E-05 |
| Rpl29 | 1.06 | 2.21E-07 |
| Rpl3 | 0.79065 | 2.32E-04 |
| Rpl31 | 1.2823 | 4.97E-05 |
| Rpl32 | 1.4834 | 5.95E-08 |

| | | |
|---|---|---|
| Rpl34 | 1.2475 | 8.87E-04 |
| Rpl35 | 1.4055 | 7.67E-06 |
| Rpl35a | 1.4349 | 8.60E-04 |
| Rpl36 | 1.1101 | 0.016535 |
| Rpl36al | 1.0154 | 6.69E-04 |
| Rpl37a | 1.3265 | 0.015095 |
| Rpl39 | 1.2687 | 3.61E-04 |
| Rpl4 | 0.92447 | 8.13E-06 |
| Rpl41 | 1.4127 | 0.0019419 |
| Rpl5 | 1.0591 | 2.39E-07 |
| Rpl6 | 0.70139 | 0.0010532 |
| Rpl7 | 1.2832 | 1.52E-10 |
| Rpl9 | 1.2377 | 6.74E-09 |
| Rplp0 | 0.71992 | 0.018163 |
| Rpn1 | 0.51055 | 0.025115 |
| Rpn2 | 0.63303 | 0.0058649 |
| Rpo1-3 | 0.63649 | 0.0046913 |
| Rpp21 | 0.91057 | 7.88E-05 |
| Rpp30 | 0.79601 | 0.0012503 |
| Rps10 | 1.0347 | 6.24E-05 |
| Rps11 | 1.2204 | 2.67E-05 |
| Rps12 | 1.2966 | 2.24E-04 |
| Rps14 | 1.3679 | 9.65E-06 |
| Rps15 | 1.2439 | 4.12E-04 |
| Rps15a | 1.3007 | 9.20E-06 |
| Rps16 | 1.231 | 1.58E-05 |
| Rps17 | 1.2986 | 4.79E-08 |
| Rps18 | 0.96126 | 2.29E-04 |
| Rps2 | 0.86452 | 0.0056263 |
| Rps20 | 1.1204 | 0.0497 |
| Rps23 | 1.2429 | 4.56E-10 |
| Rps24 | 1.2948 | 7.94E-05 |
| Rps25 | 1.418 | 1.11E-04 |
| Rps27 | 1.4518 | 0.0090114 |
| Rps27a | 1.0961 | 0.02922 |
| Rps28 | 1.6211 | 0.0021658 |
| Rps29 | 1.4324 | 0.020553 |
| Rps3 | 1.0271 | 3.63E-07 |
| Rps3a | 1.231 | 2.21E-06 |
| Rps4x | 1.378 | 2.39E-09 |
| Rps5 | 1.3105 | 8.78E-11 |

| | | |
|---|---|---|
| Rps6ka1 | -0.76719 | 0.027044 |
| Rps6ka2 | -0.69933 | 0.0020186 |
| Rps6ka3 | -0.58853 | 0.025339 |
| Rps6ka4 | -0.78249 | 9.33E-04 |
| Rps7 | 1.2275 | 1.08E-06 |
| Rrm2b | -0.70296 | 0.03017 |
| Rrp12 | -0.54391 | 0.043404 |
| Rrp15 | 0.59736 | 0.021663 |
| Rrp1b | -1.0269 | 8.87E-04 |
| Rsrc2 | 0.58903 | 0.013351 |
| RT1-CE7 | -4.312 | 0.036079 |
| RT1-M3-1 | 0.68959 | 0.022043 |
| RT1-N2 | 0.80276 | 0.038379 |
| RT1-N3 | 0.68867 | 0.01102 |
| RT1-T24-1 | -2.7157 | 0.013728 |
| RT1-T24-3 | -4.3917 | 0.042885 |
| Rtel1 | -0.82145 | 0.040574 |
| Rtn4rl2 | -0.65695 | 0.035784 |
| Rtp2 | #NAME? | 0.0059062 |
| Runx1t1 | -0.84165 | 0.013056 |
| Rwdd2a | 0.75132 | 0.038111 |
| Rxra | -0.75502 | 0.0036351 |
| Rxrg | -0.78137 | 0.014303 |
| S100a1 | 1.5524 | 1.39E-10 |
| S100a10 | 0.8161 | 2.64E-04 |
| S100a16 | 1.0795 | 0.012891 |
| S100b | 1.2687 | 3.70E-07 |
| Sall3 | -1.1578 | 4.76E-05 |
| Samd12 | -1.1511 | 2.59E-06 |
| Samd14 | -0.85686 | 2.09E-04 |
| Samd5 | -1.2671 | 1.14E-05 |
| Samd8 | -1.1184 | 2.22E-07 |
| Sap25 | -2.6184 | 0.013916 |
| Sapcd2 | -1.5818 | 3.71E-04 |
| Sar1b | 0.78877 | 2.77E-04 |
| Sarm1 | -0.98756 | 6.65E-06 |
| Sarnp | 1.0667 | 1.75E-06 |
| Sat1 | 1.4586 | 4.43E-12 |
| Sat2 | 0.64567 | 0.03885 |
| Satb2 | -0.60512 | 0.012707 |
| Sbk1 | -0.79544 | 1.65E-04 |

| | | |
|---|---|---|
| Sbno2 | -0.82958 | 0.0022287 |
| Scai | -1.2582 | 6.13E-07 |
| Scfd1 | 0.54386 | 0.028764 |
| Scg2 | 0.73758 | 0.019331 |
| Scg3 | 0.83162 | 9.79E-04 |
| Scg5 | 1.3298 | 0.0063105 |
| Scly | -0.71928 | 0.012218 |
| Scn2a | -1.0322 | 5.27E-07 |
| Scn2b | -0.99005 | 8.08E-06 |
| Scnm1 | 0.68917 | 0.014944 |
| Scoc | 0.48289 | 0.029752 |
| Scpep1 | 0.96342 | 4.33E-05 |
| Scrib | -0.84398 | 8.94E-04 |
| Scrn1 | -0.8806 | 2.11E-05 |
| Scrt1 | -1.3354 | 1.07E-08 |
| Scrt2 | -1.8556 | 7.13E-07 |
| Scube1 | -1.2922 | 7.67E-09 |
| Scube3 | -4.1927 | 5.68E-08 |
| Sdc2 | 0.82519 | 0.041324 |
| Sdc3 | -1.1527 | 1.22E-08 |
| Sdc4 | 0.91074 | 8.29E-04 |
| Sdcbp | 0.71765 | 6.09E-04 |
| Sdccag3 | 0.60073 | 0.0093462 |
| Sdccag8 | 0.84193 | 0.02392 |
| Sdf2 | 0.95339 | 7.50E-06 |
| Sdf4 | 0.50537 | 0.037754 |
| Sdhaf2 | 0.59184 | 0.010173 |
| Sdhaf3 | 0.91382 | 0.0041299 |
| Sdhaf4 | 0.95919 | 7.37E-04 |
| Sdhb | 0.60909 | 0.0058241 |
| Sdhc | 0.6236 | 0.0081388 |
| Sdhd | 0.48622 | 0.032766 |
| Sdk1 | -1.2924 | 6.68E-06 |
| Sdk2 | -1.7532 | 1.58E-11 |
| Sec11a | 0.7669 | 6.92E-04 |
| Sec11c | 0.69608 | 0.0023096 |
| Sec14l2 | -1.0023 | 0.003909 |
| Sec22a | 0.59647 | 0.029324 |
| Sec22c | -1.1936 | 8.84E-06 |
| Sec31a | -0.7461 | 9.33E-04 |
| Sec61b | 0.69899 | 0.0033813 |

| | | |
|---|---|---|
| Sec62 | 0.96493 | 3.90E-06 |
| Sel1l | -0.88383 | 3.32E-05 |
| Selk | 0.9532 | 8.48E-05 |
| Sema4b | -0.94962 | 0.0014955 |
| Sema4g | -0.537 | 0.032515 |
| Sema6a | -1.0758 | 1.21E-04 |
| Sema6b | -0.59701 | 0.0068205 |
| Sepn1 | -0.64013 | 0.010172 |
| Serf1 | 0.56911 | 0.021796 |
| Serf2 | 1.0451 | 0.010963 |
| Serinc1 | 1.1953 | 1.39E-09 |
| Serinc3 | 0.72481 | 5.39E-04 |
| Serinc4 | 0.75462 | 0.0016294 |
| Serp2 | 0.84649 | 0.031134 |
| Serpinb6 | 0.88449 | 0.0014502 |
| Serpinb9 | 0.94615 | 0.017076 |
| Serpine2 | 0.46438 | 0.048447 |
| Sertad2 | -0.98315 | 0.013572 |
| Sesn2 | -1.0056 | 0.020794 |
| Sesn3 | -1.0824 | 1.99E-04 |
| Set | -0.64111 | 0.0055182 |
| Setbp1 | -1.9342 | 3.68E-14 |
| Setd1b | -1.9346 | 7.95E-18 |
| Setd7 | -1.4843 | 2.62E-07 |
| Sez6 | -0.78821 | 4.86E-04 |
| Sez6l | -1.1824 | 3.82E-09 |
| Sf3a2 | -0.987 | 4.05E-04 |
| Sf3b5 | 0.76965 | 5.30E-04 |
| Sf3b6 | 0.91734 | 9.82E-05 |
| Sfmbt2 | -1.7319 | 7.61E-04 |
| Sfr1 | 0.93942 | 1.22E-05 |
| Sft2d1 | 0.88217 | 8.45E-05 |
| Sgce | 0.71355 | 0.0024197 |
| Sgol2 | -2.3567 | 0.0097876 |
| Sgpp2 | -1.5462 | 0.0084738 |
| Sh2d1a | 1.7431 | 8.07E-04 |
| Sh3bp1 | -1.3397 | 0.011519 |
| Sh3d19 | -1.0274 | 1.71E-05 |
| Sh3gl3 | 0.63304 | 0.0073172 |
| Sh3pxd2a | -2.2358 | 5.98E-14 |
| Sh3pxd2b | -2.0884 | 3.94E-15 |

| | | |
|---|---|---|
| Sh3rf1 | -0.89843 | 1.03E-04 |
| Sh3rf3 | -1.2596 | 5.51E-06 |
| Shc2 | -0.60461 | 0.010066 |
| Shc3 | -2.1528 | 3.58E-13 |
| Shc4 | -1.0876 | 0.035855 |
| Shfm1 | 1.0452 | 2.02E-04 |
| Shisa5 | 0.61322 | 0.014012 |
| Shisa6 | -2.1887 | 1.78E-14 |
| Shkbp1 | -0.89345 | 0.012865 |
| Shroom2 | 0.5032 | 0.024924 |
| Shroom3 | -0.6657 | 0.0095239 |
| Shtn1 | -0.61127 | 0.015224 |
| Siah3 | -2.14 | 1.75E-07 |
| Sidt2 | -0.56366 | 0.013059 |
| Sik1 | -1.3987 | 0.0010295 |
| Sik3 | -0.62944 | 0.0051141 |
| Sin3b | -0.74613 | 7.19E-04 |
| Sipa1 | -0.99268 | 0.035042 |
| Sirt3 | 0.47077 | 0.045701 |
| Siva1 | 0.68719 | 0.04074 |
| Ska2 | 1.0057 | 2.56E-04 |
| Skil | -1.0028 | 3.32E-05 |
| Skp1 | 0.77919 | 1.29E-04 |
| Slain1 | 0.76391 | 0.003622 |
| Slc10a1 | 1.77 | 0.0067721 |
| Slc16a14 | -1.5106 | 0.0044323 |
| Slc16a2 | -1.2656 | 8.45E-06 |
| Slc16a7 | -1.035 | 1.35E-04 |
| Slc17a6 | 0.82077 | 0.0023472 |
| Slc18a2 | 1.4018 | 0.011923 |
| Slc20a1 | 0.62434 | 0.0062645 |
| Slc22a12 | -1.211 | 8.32E-04 |
| Slc25a14 | 0.80951 | 0.0054142 |
| Slc25a17 | 0.67349 | 0.0044507 |
| Slc25a2 | 1.2656 | 0.015286 |
| Slc25a20 | 0.55975 | 0.025125 |
| Slc25a28 | 0.767 | 0.0015126 |
| Slc25a29 | -0.71679 | 0.0074607 |
| Slc25a33 | 0.51834 | 0.038422 |
| Slc25a5 | 0.58807 | 0.034917 |
| Slc25a51 | 0.78544 | 0.0033029 |

| | | |
|---|---|---|
| Slc27a4 | -0.60188 | 0.0091053 |
| Slc29a4 | -0.85561 | 1.05E-04 |
| Slc2a12 | 1.4063 | 3.40E-05 |
| Slc2a13 | -0.73704 | 0.0031827 |
| Slc30a6 | 0.57986 | 0.034593 |
| Slc30a9 | 0.54924 | 0.013513 |
| Slc35a1 | 0.78327 | 0.0010109 |
| Slc35d2 | 2.5439 | 0.0044146 |
| Slc35e2b | -0.75576 | 0.0037569 |
| Slc35f4 | 0.92378 | 0.0019755 |
| Slc35f6 | 0.60431 | 0.014381 |
| Slc35g2 | 1.3909 | 0.0016337 |
| Slc35g3 | -1.9054 | 0.0030436 |
| Slc36a1 | -1.5329 | 1.72E-09 |
| Slc36a4 | -0.67073 | 0.0095878 |
| Slc38a1 | -1.0932 | 6.28E-07 |
| Slc38a7 | -0.64384 | 0.014136 |
| Slc39a11 | -0.70069 | 0.012666 |
| Slc39a9 | -0.80143 | 8.63E-04 |
| Slc3a1 | 0.80733 | 2.66E-04 |
| Slc41a1 | -0.6487 | 0.0037682 |
| Slc44a5 | -1.4357 | 0.0011326 |
| Slc45a4 | -1.0712 | 9.28E-06 |
| Slc4a7 | -1.1273 | 1.10E-06 |
| Slc6a11 | -1.0089 | 0.025246 |
| Slc6a15 | 0.67452 | 0.016002 |
| Slc6a5 | -2.1375 | 0.024287 |
| Slc6a6 | -0.78598 | 0.0011568 |
| Slc7a14 | -1.5998 | 1.77E-10 |
| Slc7a6os | 0.57996 | 0.017256 |
| Slc9a1 | -0.85673 | 1.06E-04 |
| Slc9a3r1 | -1.0477 | 4.76E-06 |
| Slc9a3r2 | -0.62883 | 0.01793 |
| Slc9a7 | -1.5569 | 4.45E-12 |
| Slc9a8 | -0.57384 | 0.041834 |
| Slc9b2 | 1.072 | 0.010311 |
| Slco1c1 | 0.69387 | 0.027345 |
| Slco3a1 | -1.4204 | 7.05E-10 |
| Slco5a1 | -1.8832 | 0.0022359 |
| Slf2 | 1.0499 | 1.17E-05 |
| Slit1 | -1.322 | 7.75E-08 |

| | | |
|---|---|---|
| Slit3 | -1.0986 | 5.23E-07 |
| Slitrk2 | -0.88261 | 0.0011366 |
| Slk | -0.77425 | 0.011698 |
| Slmap | -0.6005 | 0.0095891 |
| Slx4ip | -1.1848 | 0.0097944 |
| Smad5 | -0.96156 | 7.53E-05 |
| Smap2 | -0.48062 | 0.038756 |
| Smarcc1 | -0.68483 | 0.0027077 |
| Smcr8 | -0.92569 | 2.77E-04 |
| Smg5 | -0.88692 | 6.85E-05 |
| Smim11 | 1.0866 | 7.87E-04 |
| Smim13 | -0.69206 | 0.011065 |
| Smim14 | 0.46005 | 0.048725 |
| Smim15 | 0.95031 | 4.14E-05 |
| Smim18 | 0.90125 | 0.0031054 |
| Smim19 | 0.55162 | 0.048415 |
| Smim3 | 1.1754 | 0.021329 |
| Smim7 | 0.49092 | 0.034853 |
| Smim8 | 1.0682 | 4.45E-05 |
| Smoc1 | -0.88932 | 4.10E-04 |
| Smu1 | 0.50361 | 0.041312 |
| Smurf1 | -1.1957 | 4.41E-07 |
| Snapc5 | 0.5661 | 0.024621 |
| Snapin | 0.99 | 3.02E-06 |
| Sncb | -0.61459 | 0.0064325 |
| Snf8 | 0.66838 | 0.0031116 |
| Snrnp25 | 1.0949 | 2.84E-06 |
| Snrnp35 | 0.58833 | 0.022565 |
| Snrpa1 | 0.56134 | 0.027601 |
| Snrpb2 | 0.76685 | 0.0017233 |
| Snrpd1 | 0.7467 | 8.07E-04 |
| Snrpd2 | 0.94039 | 5.68E-06 |
| Snrpd3 | 0.4806 | 0.047788 |
| Snrpe | 1.123 | 2.41E-07 |
| Snrpf | 0.8073 | 4.92E-04 |
| Sntb2 | -2.0905 | 6.23E-04 |
| Snu13 | 0.57536 | 0.010374 |
| Snx19 | -0.54941 | 0.018683 |
| Snx29 | -0.97351 | 0.0023699 |
| Snx3 | 0.62362 | 0.0038177 |
| Snx30 | -1.0867 | 5.13E-06 |

| | | |
|---|---|---|
| Snx32 | -0.5431 | 0.028108 |
| Snx5 | 0.68727 | 0.002372 |
| Soga3 | -0.97406 | 4.37E-06 |
| Sorbs1 | -1.0746 | 1.10E-04 |
| Sorcs1 | -0.52043 | 0.046018 |
| Sorcs2 | -0.67683 | 0.012236 |
| Sorcs3 | -0.93685 | 4.20E-04 |
| Sox11 | -0.9394 | 5.81E-05 |
| Sox12 | -0.77794 | 8.77E-04 |
| Sox5 | -0.8731 | 0.0050795 |
| Sox6 | -1.6075 | 5.18E-05 |
| Sox9 | -1.1437 | 1.59E-05 |
| Sp2 | -0.7412 | 0.016571 |
| Spag7 | 0.62908 | 0.0084757 |
| Sparc | 0.98642 | 2.07E-06 |
| Spata13 | -1.9809 | 5.20E-05 |
| Spata2 | -0.58464 | 0.01303 |
| Spata7 | 0.82461 | 0.0018814 |
| Spata9 | 0.89514 | 0.0056462 |
| Spcs1 | 1.0437 | 9.75E-07 |
| Spcs2 | 0.66082 | 0.0023324 |
| Specc1 | -0.84495 | 0.046926 |
| Spock3 | 1.5509 | 0.031682 |
| Spop | 0.8732 | 1.13E-04 |
| Spred2 | -1.0715 | 2.16E-06 |
| Sprtn | -0.65677 | 0.028604 |
| Spryd3 | -0.55017 | 0.015386 |
| Spryd7 | 0.91371 | 2.63E-05 |
| Spta1 | 1.5063 | 2.21E-07 |
| Sptlc2 | -0.73369 | 0.0046473 |
| Sptssa | 1.2208 | 1.31E-08 |
| Spty2d1 | -0.69891 | 0.0272 |
| Src | -0.62704 | 0.0067835 |
| Srgap1 | -1.7637 | 4.21E-11 |
| Srgap2 | -0.87633 | 6.41E-05 |
| Srm | -0.64036 | 0.049723 |
| Srp14 | 0.92172 | 5.97E-05 |
| Srp19 | 0.81379 | 2.20E-04 |
| Srp54a | 0.5603 | 0.016023 |
| Srp9 | 0.88634 | 4.14E-05 |
| Srrm3 | -0.88282 | 2.22E-04 |

| | | |
|---|---|---|
| Srrm4 | -1.7345 | 6.74E-11 |
| Srrt | -0.69488 | 0.0025683 |
| Srsf5 | 0.54355 | 0.016122 |
| Ss18l2 | 1.0227 | 8.42E-05 |
| Ssbp1 | 0.56287 | 0.049229 |
| Ssh1 | -1.8093 | 1.01E-08 |
| Ssh2 | -1.1226 | 8.02E-07 |
| Ssna1 | 0.6684 | 0.0063073 |
| Ssr2 | 0.64536 | 0.0036219 |
| Ssr3 | 0.86899 | 4.10E-05 |
| Ssu72 | 0.81597 | 0.0014544 |
| St18 | -1.4122 | 0.039127 |
| St3gal1 | -0.95517 | 5.99E-04 |
| St6gal2 | -0.74477 | 0.033524 |
| St7l | 0.74691 | 0.001456 |
| St8sia3 | -1.331 | 2.46E-09 |
| Stag2 | 0.65528 | 0.0037636 |
| Stambp | 0.80558 | 7.00E-04 |
| Stard10 | 0.75541 | 0.0035 |
| Stat5b | -0.64204 | 0.018653 |
| Stbd1 | 0.9119 | 0.0045154 |
| Stc2 | -1.2114 | 0.011465 |
| Stk11ip | -0.6062 | 0.011663 |
| Stk35 | -0.60332 | 0.037912 |
| Stoml2 | 0.51917 | 0.033564 |
| Ston2 | -1.578 | 8.88E-05 |
| Strn | -1.1105 | 1.26E-06 |
| Stt3a | 0.48313 | 0.045317 |
| Stx16 | 0.52858 | 0.026892 |
| Stx1b | -0.48391 | 0.037899 |
| Stx4 | 0.60127 | 0.014859 |
| Stx8 | 0.93373 | 0.0016461 |
| Stxbp3 | 1.2014 | 3.40E-07 |
| Stxbp4 | -1.2447 | 9.58E-04 |
| Stxbp5l | -1.3488 | 7.61E-04 |
| Sub1 | 0.8709 | 1.94E-05 |
| Suclg2 | 0.58225 | 0.017044 |
| Sufu | -1.3808 | 7.95E-06 |
| Sugt1 | 0.93323 | 2.78E-05 |
| Sulf2 | -0.94608 | 0.013346 |
| Sult1d1 | 1.6292 | 0.029395 |

| | | |
|---|---|---|
| Sult2b1 | -1.0285 | 6.36E-05 |
| Sumf2 | 0.71289 | 0.018031 |
| Sumo1 | 1.0743 | 8.79E-07 |
| Sumo3 | 0.48166 | 0.043003 |
| Sumo4 | 1.1918 | 2.58E-06 |
| Supt4h1 | 1.1894 | 4.31E-04 |
| Supt5h | -0.77132 | 2.52E-04 |
| Supt6h | -0.6899 | 0.0012862 |
| Surf1 | 1.5728 | 0.001004 |
| Susd6 | -1.0471 | 2.92E-04 |
| Svbp | 0.87931 | 6.95E-04 |
| Svop | -0.57501 | 0.0099952 |
| Swi5 | 0.47887 | 0.034488 |
| Syap1 | 0.62645 | 0.030232 |
| Syn2 | -0.65793 | 0.0023057 |
| Syn3 | -1.2774 | 5.23E-07 |
| Synj2 | -0.84672 | 0.015488 |
| Synpr | 1.1615 | 0.04454 |
| Syt11 | -0.50547 | 0.021115 |
| Syt12 | -0.82621 | 0.0074516 |
| Syt14 | -0.9281 | 0.0063407 |
| Syt6 | -1.5591 | 2.84E-05 |
| Sytl5 | -1.0145 | 0.047678 |
| Szt2 | -1.3535 | 3.82E-09 |
| Tacc3 | -1.3833 | 0.00068847 |
| Taf13 | 1.118 | 1.35E-07 |
| Taf3 | 0.61134 | 0.0051269 |
| Taf9 | 1.0158 | 0.026164 |
| Tanc1 | -1.6101 | 5.93E-12 |
| Tank | 0.98519 | 8.95E-04 |
| Taok3 | -0.96632 | 2.13E-05 |
| Tarsl2 | -0.79101 | 0.001781 |
| Tas1r1 | 0.66203 | 0.03263 |
| Tatdn1 | 0.75733 | 0.005527 |
| Tatdn2 | -0.70048 | 0.0024668 |
| Tbc1d10a | -0.65381 | 0.014007 |
| Tbc1d13 | -0.99898 | 7.27E-05 |
| Tbc1d16 | -1.7612 | 1.83E-12 |
| Tbc1d5 | -0.59424 | 0.016686 |
| Tbca | 1.0602 | 0.00213 |
| Tbce | 0.52269 | 0.034619 |

| | | |
|---|---|---|
| Tbcel | -1.1543 | 1.16E-04 |
| Tbl2 | -0.499 | 0.048233 |
| Tbr1 | -0.81486 | 7.39E-04 |
| Tceal1 | 0.9488 | 1.94E-05 |
| Tceal7 | 1.7811 | 4.14E-05 |
| Tceal8 | 1.0786 | 3.84E-07 |
| Tceb1 | 0.76907 | 4.07E-04 |
| Tceb2 | 0.98409 | 0.0085368 |
| Tcf7l1 | -0.69175 | 0.041312 |
| Tcof1 | -0.70195 | 0.0089144 |
| Tcp1 | 0.57576 | 0.014369 |
| Tcp11l1 | -1.6176 | 6.02E-08 |
| Tcp11l2 | 1.4057 | 0.026462 |
| Tcte1 | 1.2371 | 9.21E-05 |
| Tctex1d2 | 1.0715 | 2.39E-06 |
| Tead1 | -1.9987 | 1.89E-08 |
| Tec | 0.95968 | 0.030077 |
| Tecpr1 | -0.53606 | 0.016099 |
| Tecpr2 | -0.99629 | 1.46E-05 |
| Tecr | 0.56975 | 0.01232 |
| Tefm | 0.68209 | 0.035339 |
| Tekt2 | 1.9689 | 0.048853 |
| Tenm2 | -0.78381 | 2.53E-04 |
| Tenm4 | -2.1081 | 3.94E-25 |
| Tesc | -0.71262 | 0.0061568 |
| Tet1 | -1.6639 | 3.36E-08 |
| Tet2 | -0.75211 | 0.0033177 |
| Tet3 | -2.098 | 8.14E-21 |
| Tfb2m | 0.58149 | 0.020992 |
| Tfe3 | -0.81531 | 0.0016272 |
| Tfpi | 0.89674 | 0.0049041 |
| Tfr2 | -1.0639 | 0.02995 |
| Tgfa | -1.0737 | 0.0070582 |
| Tgfbr3l | -1.8431 | 0.012819 |
| Tgs1 | 0.58802 | 0.020906 |
| Thoc7 | 0.76147 | 7.85E-04 |
| Thyn1 | 0.59698 | 0.012423 |
| Tial1 | -0.67245 | 0.0046307 |
| Ticrr | -1.9797 | 1.94E-05 |
| Timeless | -0.77766 | 0.019787 |
| Timm10b | 1.0504 | 0.003698 |

| | | |
|---|---|---|
| Timm13 | 0.7417 | 4.27E-04 |
| Timm17a | 0.72411 | 9.37E-04 |
| Timm21 | 0.85296 | 4.66E-04 |
| Timm23 | 0.53728 | 0.019787 |
| Timm8a1 | 0.8509 | 0.015267 |
| Timm8b | 1.1283 | 8.73E-04 |
| Timm9 | 0.60863 | 0.014303 |
| Timp2 | -1.0452 | 1.89E-07 |
| Tipin | 1.1827 | 5.64E-04 |
| Tipinl1 | 1.2653 | 1.82E-05 |
| Tiprl | 0.5978 | 0.010203 |
| Tjap1 | -0.82895 | 0.012808 |
| Tk1 | -0.75517 | 0.047135 |
| Tle2 | -1.3699 | 4.45E-07 |
| Tm2d1 | 0.93219 | 2.50E-05 |
| Tm2d2 | 0.59065 | 0.0098558 |
| Tm9sf2 | 0.71071 | 9.64E-04 |
| Tmbim4 | 1.2169 | 6.34E-08 |
| Tmco1 | 0.72193 | 0.0034575 |
| Tmed2 | 0.49648 | 0.036066 |
| Tmed4 | 0.84709 | 0.010708 |
| Tmed7 | 0.78558 | 2.94E-04 |
| Tmed8 | -1.372 | 2.86E-06 |
| Tmem104 | -0.89468 | 0.0036658 |
| Tmem107 | 0.80988 | 0.035832 |
| Tmem108 | -0.89203 | 5.97E-04 |
| Tmem114 | -3.3227 | 0.017164 |
| Tmem126a | 0.75046 | 0.0019032 |
| Tmem126b | 0.83668 | 3.59E-04 |
| Tmem128 | 0.85394 | 1.47E-04 |
| Tmem131 | -0.47418 | 0.049444 |
| Tmem132b | -1.9503 | 6.21E-16 |
| Tmem132c | -0.83633 | 0.029512 |
| Tmem132d | -1.21 | 1.08E-06 |
| Tmem136 | 0.72645 | 0.0045077 |
| Tmem138 | 0.69344 | 0.028914 |
| Tmem141 | 1.1839 | 0.001949 |
| Tmem145 | -0.59064 | 0.012154 |
| Tmem147 | 0.58038 | 0.013204 |
| Tmem14c | 0.49882 | 0.037202 |
| Tmem161b | 0.59195 | 0.024414 |

| | | |
|---|---|---|
| Tmem165 | 0.723 | 0.015876 |
| Tmem170b | -1.5018 | 5.25E-08 |
| Tmem179b | 1.2003 | 3.12E-04 |
| Tmem181 | -0.5739 | 0.015224 |
| Tmem183a | 0.48034 | 0.04689 |
| Tmem184c | 0.54603 | 0.023233 |
| Tmem185b | 0.55853 | 0.029752 |
| Tmem186 | 0.95109 | 4.33E-05 |
| Tmem19 | 0.79741 | 8.94E-04 |
| Tmem191c | -0.9708 | 0.0097895 |
| Tmem192 | 0.6016 | 0.034571 |
| Tmem199 | 0.52369 | 0.031626 |
| Tmem201 | -1.2227 | 1.73E-07 |
| Tmem205 | 1.134 | 9.33E-07 |
| Tmem208 | 0.68379 | 0.0026589 |
| Tmem216 | 0.82435 | 0.015514 |
| Tmem218 | 0.96765 | 0.022893 |
| Tmem219 | 0.63555 | 0.0301 |
| Tmem223 | 0.71461 | 0.0099043 |
| Tmem245 | -1.7928 | 4.00E-13 |
| Tmem251 | 0.95523 | 3.46E-04 |
| Tmem256 | 0.93224 | 0.022076 |
| Tmem258b | 1.0188 | 0.04074 |
| Tmem259 | -0.64386 | 0.0036113 |
| Tmem261 | 0.83949 | 0.0053131 |
| Tmem38b | 0.93458 | 5.75E-04 |
| Tmem41b | 0.80552 | 2.23E-04 |
| Tmem47 | 0.95419 | 3.75E-06 |
| Tmem50a | 0.78544 | 2.33E-04 |
| Tmem59 | 0.83509 | 9.64E-05 |
| Tmem60 | 0.84878 | 4.89E-04 |
| Tmem63c | -0.58711 | 0.024326 |
| Tmem69 | 0.68933 | 0.0050044 |
| Tmem70 | 0.51554 | 0.035241 |
| Tmem71 | -1.6878 | 0.020394 |
| Tmem80 | 0.7214 | 0.0019755 |
| Tmem82 | -3.1158 | 6.69E-04 |
| Tmem94 | -1.1626 | 2.22E-07 |
| Tmem97 | 0.77701 | 0.046882 |
| Tmem9b | 0.52881 | 0.02501 |
| Tmod3 | 0.63189 | 0.021726 |

| | | |
|---|---|---|
| Tmsb4x | 1.2016 | 3.92E-06 |
| Tmsbl1 | 0.78174 | 5.42E-04 |
| Tmx2 | 0.64295 | 0.0036949 |
| Tmx4 | 0.60789 | 0.0057818 |
| Tnfaip8l1 | -1.779 | 0.047236 |
| Tnks1bp1 | -0.68134 | 0.0020749 |
| Tnnc1 | 1.0696 | 0.0075233 |
| Tnnc2 | 2.787 | 0.0053924 |
| Tnpo1 | -1.098 | 5.11E-06 |
| Tnr | -1.3752 | 0.0064978 |
| Tnrc6a | -0.69561 | 0.0033412 |
| Tnrc6c | -1.3569 | 7.55E-11 |
| Tns1 | -2.0191 | 1.25E-04 |
| Tob1 | 0.60578 | 0.01398 |
| Tom1l1 | 1.0125 | 2.48E-04 |
| Tom1l2 | -1.3343 | 2.38E-10 |
| Tomm20 | 0.80808 | 1.09E-04 |
| Tomm22 | 0.61081 | 0.0060537 |
| Tomm6 | 0.77508 | 0.0020608 |
| Tomm7 | 1.2159 | 0.0029058 |
| Tomm70a | -0.68905 | 0.0019995 |
| Tonsl | -1.1876 | 0.002331 |
| Top1 | -1.463 | 6.18E-05 |
| Top3a | -1.2394 | 0.0050044 |
| Topors | 0.69059 | 0.0030274 |
| Tox2 | -0.8443 | 0.015224 |
| Tox3 | 0.76025 | 0.023783 |
| Tp53 | -0.86015 | 2.75E-04 |
| Tp53bp1 | -0.52115 | 0.021024 |
| Tp53i11 | -0.56267 | 0.047203 |
| Tp53inp2 | -1.9223 | 7.11E-05 |
| Tp53rk | 0.68368 | 0.042371 |
| Tpp2 | 0.68161 | 0.0023216 |
| Tpt1 | 1.1787 | 0.0035256 |
| Tpx2 | -1.243 | 0.027327 |
| Tra2a | 0.53771 | 0.024213 |
| Trabd2b | -2.4873 | 0.0080343 |
| Traf3 | -1.1975 | 6.39E-09 |
| Traf4 | -0.93402 | 0.014769 |
| Traf6 | -1.0986 | 0.0017813 |
| Traf7 | -0.64463 | 0.0065666 |

| | | |
|---|---|---|
| Tram1l1 | 0.60045 | 0.011286 |
| Trappc1 | 0.70952 | 0.01632 |
| Trappc13 | 0.46858 | 0.048116 |
| Trappc2 | 0.53973 | 0.017547 |
| Trappc2b | 1.1022 | 2.76E-05 |
| Trappc2l | 0.92021 | 1.72E-04 |
| Trappc3l | -3.4151 | 0.010565 |
| Trappc6b | 0.85931 | 0.042872 |
| Trerf1 | -0.71925 | 0.010911 |
| Trhde | -0.57691 | 0.046241 |
| Trim14 | 1.46 | 0.048271 |
| Trim23 | 0.6643 | 0.0051922 |
| Trim44 | -1.2117 | 1.40E-08 |
| Trim62 | -1.0342 | 0.01193 |
| Trim66 | -2.1871 | 8.94E-08 |
| Trim67 | -2.2141 | 1.24E-09 |
| Trim8 | -0.46317 | 0.045883 |
| Triqk | 0.78197 | 0.03569 |
| Trit1 | 0.61472 | 0.017882 |
| Trmt1 | -0.556 | 0.022332 |
| Trmt10b | 0.73137 | 0.0085302 |
| Trmt10c | 0.79536 | 9.26E-04 |
| Trmt112 | 0.6372 | 0.0064923 |
| Trmt12 | 0.62357 | 0.037126 |
| Trmt5 | 0.59462 | 0.042949 |
| Trnt1 | 0.74586 | 0.0016294 |
| Trove2 | -0.82506 | 6.37E-04 |
| Trpc3 | -0.7273 | 0.03263 |
| Trpc5 | -1.2501 | 1.53E-07 |
| Trpm2 | -0.72877 | 0.035114 |
| Trps1 | -1.1336 | 0.012995 |
| Tsc1 | -0.88879 | 1.38E-04 |
| Tsen15 | 0.92175 | 9.21E-05 |
| Tsg101 | 0.52802 | 0.023288 |
| Tspan3 | 0.53034 | 0.017708 |
| Tspan31 | 0.75836 | 7.95E-04 |
| Tspan6 | 1.2369 | 5.31E-09 |
| Tspyl5 | -0.58153 | 0.032435 |
| Tsr3 | 0.55216 | 0.030622 |
| Tst | 0.584 | 0.020995 |
| Tstd3 | 0.63133 | 0.011002 |

| | | |
|---|---|---|
| Ttc28 | -1.2759 | 2.25E-06 |
| Ttc39b | -1.3194 | 5.73E-05 |
| Ttl | -0.81918 | 3.51E-04 |
| Ttll12 | -0.64156 | 0.010291 |
| Ttll4 | -1.2897 | 4.81E-06 |
| Ttll8 | -2.5922 | 0.022792 |
| Tub | -1.4123 | 3.64E-11 |
| Tuft1 | -1.0458 | 0.042129 |
| Tulp3 | -1.1078 | 1.72E-04 |
| Tusc2 | 0.45528 | 0.047675 |
| Tusc3 | 0.62728 | 0.010482 |
| Tvp23b | 0.74275 | 0.0012531 |
| Txn1 | 0.75104 | 0.043786 |
| Txn2 | 0.52018 | 0.025512 |
| Txndc12 | 0.99923 | 4.38E-06 |
| Txndc15 | 0.67189 | 0.0030326 |
| Txndc17 | 0.7581 | 0.0018077 |
| Txndc9 | 0.65638 | 0.004164 |
| Tyk2 | -1.0886 | 2.50E-04 |
| Tysnd1 | -0.6547 | 0.012704 |
| Tyw5 | 0.72728 | 0.043893 |
| U2af1 | 0.55552 | 0.020302 |
| U2af1l4 | 0.67024 | 0.0038898 |
| Uba3 | 0.50106 | 0.031342 |
| Uba52 | 1.1569 | 1.13E-04 |
| Uba6 | 0.55514 | 0.028676 |
| Ubac2 | 0.53632 | 0.030174 |
| Ubash3b | -0.98829 | 1.93E-04 |
| Ube2b | 0.91038 | 0.0055004 |
| Ube2e1 | 0.6076 | 0.0096028 |
| Ube2l3 | 0.4645 | 0.048591 |
| Ube2v2 | 0.86912 | 3.72E-05 |
| Ubfd1 | -0.64158 | 0.010951 |
| Ubn2 | -1.0941 | 1.57E-05 |
| Ubr1 | -0.61473 | 0.0095942 |
| Ubr2 | -0.91171 | 4.39E-05 |
| Ubr4 | -0.93055 | 7.60E-06 |
| Ubxn11 | 0.64268 | 0.024832 |
| Ubxn2b | -0.65814 | 0.01137 |
| Ubxn7 | -1.3457 | 0.0024021 |
| Ubxn8 | 0.73029 | 0.0043292 |

| | | |
|---|---|---|
| Uchl5 | 0.58504 | 0.028079 |
| Uck2 | -1.4118 | 6.58E-08 |
| Ufc1 | 0.71956 | 9.94E-04 |
| Ufl1 | 0.52153 | 0.047829 |
| Ufm1 | 0.6821 | 0.0028566 |
| Ufsp2 | 0.8105 | 3.92E-04 |
| Uggt1 | -0.86315 | 7.62E-05 |
| Ugp2 | 0.59022 | 0.012225 |
| Ugt1a8 | 0.90891 | 0.0063205 |
| Uhrf1 | -1.0524 | 0.0031735 |
| Uhrf1bp1 | -1.6794 | 6.92E-11 |
| Umps | 0.62844 | 0.014657 |
| Unc119b | -0.96652 | 3.65E-04 |
| Unc50 | 0.70025 | 0.0024341 |
| Unc5b | -1.5136 | 5.44E-05 |
| Unc5c | -1.6932 | 2.52E-11 |
| Unc5d | -1.575 | 4.28E-12 |
| Unc79 | -0.55765 | 0.021254 |
| Unk | -0.94895 | 3.50E-04 |
| Upf1 | -1.2285 | 6.70E-09 |
| Uprt | -1.7653 | 2.22E-07 |
| Uqcc2 | 0.83038 | 0.010424 |
| Uqcc3 | 1.2673 | 1.93E-04 |
| Uqcrb | 1.2535 | 0.002043 |
| Uqcrb | 1.1634 | 7.31E-05 |
| Uqcrh | 1.0351 | 0.0011366 |
| Uqcrq | 0.87445 | 0.026665 |
| Uri1 | 0.65961 | 0.0039654 |
| Urod | 1.0825 | 3.20E-06 |
| Use1 | 0.71739 | 0.0022686 |
| Usp19 | -0.61375 | 0.0063105 |
| Usp24 | -1.1291 | 6.37E-08 |
| Usp31 | -1.3522 | 1.06E-09 |
| Usp37 | -0.66881 | 0.024188 |
| Usp53 | 0.8297 | 0.001205 |
| Usp54 | -0.54009 | 0.020845 |
| Usp7 | -0.50075 | 0.032121 |
| Ust | -1.1237 | 3.95E-04 |
| Utp11l | 0.53298 | 0.029282 |
| Utrn | -0.71543 | 0.0054142 |
| Uvrag | -0.63574 | 0.01991 |

| | | |
|---|---|---|
| Vac14 | -0.79105 | 7.89E-04 |
| Vamp3 | 0.79928 | 5.43E-04 |
| Vamp4 | 0.54263 | 0.017351 |
| Vamp8 | 2.0222 | 0.0016383 |
| Vangl2 | -1.1728 | 4.31E-06 |
| Vars | -0.56571 | 0.015386 |
| Vars2 | -0.91411 | 3.10E-04 |
| Vash1 | -1.3745 | 2.81E-04 |
| Vav2 | -0.79921 | 0.0010817 |
| Vcam1 | 0.75945 | 0.025498 |
| Vdac3 | 0.61705 | 0.0043402 |
| Vimp | 0.68547 | 0.0026018 |
| Vipr1 | -1.0636 | 0.028427 |
| Vkorc1 | 0.7786 | 0.0026255 |
| Vma21 | 0.74868 | 8.58E-04 |
| Vmp1 | 0.65053 | 0.0031359 |
| Vps13a | -0.97196 | 4.93E-05 |
| Vps13c | -0.89431 | 5.34E-05 |
| Vps28 | 0.7575 | 5.38E-04 |
| Vps29 | 0.58415 | 0.0076169 |
| Vps45 | 0.54074 | 0.026987 |
| Vps52 | -0.72774 | 8.68E-04 |
| Vsig10l | -1.0205 | 0.041341 |
| Vstm2a | 0.48652 | 0.045122 |
| Vta1 | 0.75154 | 5.44E-04 |
| Vti1a | -0.68039 | 0.0133 |
| Vwa5b1 | -1.4359 | 0.013393 |
| Vwa5b2 | -1.0416 | 0.0077954 |
| Wasf2 | -1.6925 | 1.14E-05 |
| Wasf3 | -0.9551 | 1.04E-05 |
| Wbp2 | -0.98152 | 2.31E-06 |
| Wbp5 | 0.90917 | 2.17E-05 |
| Wbscr17 | -1.5539 | 3.69E-08 |
| Wdfy1 | -0.67947 | 0.0046913 |
| Wdr59 | -0.81332 | 0.0017505 |
| Wdr61 | 0.57248 | 0.016215 |
| Wdr62 | -4.2099 | 0.0057325 |
| Wdr7 | -1.022 | 1.10E-06 |
| Wdr82 | -0.66463 | 0.0029927 |
| Wdr83os | 0.71989 | 0.001141 |
| Wdr89 | 1.1531 | 0.0014307 |

| | | |
|---|---|---|
| Wdr90 | -1.7544 | 2.41E-06 |
| Wdtc1 | -1.1106 | 2.12E-06 |
| Whsc1 | -0.66927 | 0.0029849 |
| Whsc1l1 | -0.55095 | 0.022721 |
| Wipf2 | -1.7266 | 1.22E-13 |
| Wiz | -0.74336 | 7.40E-04 |
| Wls | 0.6808 | 0.042949 |
| Wnk3 | -1.2433 | 3.90E-06 |
| Wnt4 | -1.1409 | 0.0012795 |
| Wnt7b | -1.2193 | 3.39E-04 |
| Wrb | 0.59541 | 0.0095346 |
| Wwc3 | -0.86626 | 0.0016278 |
| Wwtr1 | -1.0769 | 0.040353 |
| Xbp1 | 0.54182 | 0.018562 |
| Xkr4 | -1.8325 | 1.40E-16 |
| Xkr6 | -2.2297 | 1.66E-12 |
| Xkr7 | -1.2207 | 2.31E-04 |
| Xpa | 0.87892 | 0.015506 |
| Xpnpep3 | -0.76677 | 0.01154 |
| Xpot | -0.50609 | 0.026691 |
| Xpr1 | -1.1574 | 1.07E-07 |
| Xrcc1 | -0.74986 | 0.001088 |
| Xrcc4 | 0.67618 | 0.013252 |
| Xrcc5 | 0.69624 | 0.0072149 |
| Xrn1 | -0.85563 | 0.0014328 |
| Yae1d1 | 0.66661 | 0.0053754 |
| Yaf2 | 0.57606 | 0.010566 |
| Yap1 | -0.95337 | 0.002955 |
| Ybx1-ps3 | 0.55174 | 0.018078 |
| Yeats2 | -0.57189 | 0.021749 |
| Yeats4 | 0.59744 | 0.012991 |
| Yipf4 | 0.60921 | 0.0074336 |
| Yipf5 | 0.55549 | 0.016512 |
| Yipf6 | 0.4865 | 0.044745 |
| Ypel2 | -0.56974 | 0.015997 |
| Ypel5 | 0.76265 | 3.75E-04 |
| Ythdf3 | 0.51381 | 0.027599 |
| Zadh2 | 0.5928 | 0.03112 |
| Zak | -1.3508 | 1.59E-06 |
| Zbtb16 | -1.2627 | 8.63E-08 |
| Zbtb37 | -2.4316 | 2.15E-08 |

| | | |
|---|---|---|
| Zbtb4 | -0.71438 | 0.0016019 |
| Zbtb40 | -1.3216 | 2.97E-05 |
| Zbtb41 | -0.67515 | 0.01761 |
| Zbtb45 | -0.61872 | 0.018555 |
| Zbtb46 | -0.57626 | 0.04648 |
| Zbtb47 | -0.82446 | 1.58E-04 |
| Zbtb7a | -0.78247 | 6.75E-04 |
| Zbtb8os | 0.93797 | 0.0038459 |
| Zc3h12b | -0.77474 | 0.0264 |
| Zc3h12c | -1.1889 | 3.04E-04 |
| Zc3h18 | -0.59072 | 0.013133 |
| Zcchc10 | 1.0305 | 0.0035384 |
| Zcchc12 | 0.75489 | 4.07E-04 |
| Zcchc17 | 0.58219 | 0.011473 |
| Zcchc6 | -0.54498 | 0.02799 |
| Zcrb1 | 0.58603 | 0.012652 |
| Zdbf2 | -1.5721 | 2.51E-10 |
| Zdhhc2 | 1.2072 | 0.0013221 |
| Zdhhc21 | -1.2695 | 2.58E-05 |
| Zdhhc8 | -0.95582 | 1.16E-05 |
| Zfand1 | 0.68799 | 0.011698 |
| Zfand6 | 0.76204 | 9.16E-04 |
| Zfhx3 | -2.8409 | 3.91E-09 |
| Zfp105 | 0.60748 | 0.042576 |
| Zfp110 | 0.66144 | 0.008088 |
| Zfp133 | 0.70841 | 0.014818 |
| Zfp142 | -0.53827 | 0.030506 |
| Zfp18 | 0.64397 | 0.023934 |
| Zfp180 | 0.77183 | 4.69E-04 |
| Zfp236 | -1.3051 | 1.20E-06 |
| Zfp260 | 0.92587 | 3.84E-05 |
| Zfp277 | 0.63658 | 0.023077 |
| Zfp286a | 0.96836 | 5.66E-04 |
| Zfp287 | 0.62187 | 0.01699 |
| Zfp317 | 0.65449 | 0.0083391 |
| Zfp319 | -1.0628 | 8.20E-05 |
| Zfp329 | 0.60382 | 0.013729 |
| Zfp335 | -0.65863 | 0.0062062 |
| Zfp341 | -0.80798 | 0.046881 |
| Zfp347 | 1.1439 | 8.48E-05 |
| Zfp354a | 0.88155 | 0.0030284 |

| | | |
|---|---|---|
| Zfp354b | 0.90676 | 0.030081 |
| Zfp367 | 0.73354 | 0.0065778 |
| Zfp36l2 | 0.55756 | 0.028439 |
| Zfp385a | -0.62547 | 0.010373 |
| Zfp386 | 0.72223 | 0.0050504 |
| Zfp407 | -0.90728 | 0.0034858 |
| Zfp418 | 0.75661 | 0.0070875 |
| Zfp422 | 0.56984 | 0.017688 |
| Zfp426 | 0.91117 | 0.0070009 |
| Zfp455 | 0.88763 | 0.049401 |
| Zfp462 | -1.0703 | 1.26E-06 |
| Zfp472 | 1.0755 | 8.01E-04 |
| Zfp483 | -0.52895 | 0.038031 |
| Zfp496 | -0.60081 | 0.035602 |
| Zfp503 | -0.89914 | 0.024737 |
| Zfp507 | -0.69461 | 0.020242 |
| Zfp512b | -0.76057 | 4.83E-04 |
| Zfp516 | -1.0296 | 3.80E-04 |
| Zfp518b | -0.85361 | 0.0037032 |
| Zfp526 | -1.4051 | 1.05E-04 |
| Zfp560 | 1.1375 | 2.64E-06 |
| Zfp597 | -0.67073 | 0.022319 |
| Zfp598 | -0.81143 | 3.94E-04 |
| Zfp609 | -0.79992 | 2.02E-04 |
| Zfp618 | -1.4459 | 0.003673 |
| Zfp637 | 1.1641 | 1.06E-04 |
| Zfp638 | 0.4751 | 0.049428 |
| Zfp652 | -1.3794 | 2.38E-07 |
| Zfp68 | 0.70716 | 0.0022824 |
| Zfp691 | 0.99381 | 0.0038313 |
| Zfp709 | 0.82788 | 0.0042926 |
| Zfp780b-ps1 | 0.64016 | 0.013582 |
| Zfp827 | -1.3827 | 1.54E-08 |
| Zfp831 | -3.2681 | 5.48E-06 |
| Zfp84 | 0.64643 | 0.0091917 |
| Zfp869 | 0.83575 | 0.0051713 |
| Zfp871 | -1.7103 | 2.38E-10 |
| Zfp879 | 0.91329 | 0.016594 |
| Zfp90 | 0.69606 | 0.043711 |
| Zfp952 | 0.57843 | 0.049454 |
| Zfp954 | 0.85447 | 0.0033634 |

| | | |
|---|---|---|
| Zfp955a | 0.71272 | 0.0018339 |
| Zfp958 | 0.93184 | 0.036672 |
| Zfr2 | -0.6499 | 0.0044323 |
| Zfx | 0.65739 | 0.0081204 |
| Zgpat | -0.69083 | 0.0058899 |
| Zhx1 | 0.51652 | 0.026343 |
| Zkscan1 | -0.5574 | 0.02925 |
| Zmat2 | 0.63256 | 0.0040811 |
| Zmat3 | -1.1231 | 5.86E-08 |
| Zmym1 | 0.72457 | 0.048167 |
| Zmym5 | 0.74356 | 0.00198 |
| Zmym6 | 0.71905 | 0.0061119 |
| Zmynd8 | -0.60223 | 0.0099043 |
| Znf48 | -0.68043 | 0.012561 |
| Znhit3 | 0.65323 | 0.010187 |
| Znrf1 | -0.6298 | 0.0094291 |
| Zswim5 | -1.3581 | 1.96E-07 |
| Zswim6 | -0.6802 | 0.017075 |
| Zswim8 | -0.87775 | 4.41E-05 |
| Zufsp | 0.75178 | 0.0075233 |
| Zzef1 | -1.0009 | 8.06E-06 |

| Associated_Gene_Name | log2FoldChange | padj |
|---|---|---|
| Aatk | -0.76177 | 0.011912 |
| Abca2 | -1.1431 | 4.69E-05 |
| Abca3 | -2.8933 | 2.08E-17 |
| Acly | -0.8715 | 0.0046575 |
| Adarb1 | -0.66311 | 0.034728 |
| Adcy2 | 0.70792 | 0.028723 |
| Adcy5 | -1.0555 | 3.12E-04 |
| Aff1 | -1.6834 | 6.82E-05 |
| Aff3 | -1.5652 | 2.39E-07 |
| Agap1 | -1.5698 | 3.17E-08 |
| Agap2 | -0.94716 | 6.92E-04 |
| Agrn | -1.1694 | 3.28E-05 |
| Ahdc1 | -1.0067 | 7.06E-04 |
| Akap6 | -1.5192 | 2.05E-05 |
| Akt3 | -1.0402 | 7.91E-04 |
| Ank1 | -1.5898 | 4.37E-07 |
| Ank2 | -2.3035 | 2.69E-17 |
| Ankrd11 | -1.577 | 1.51E-08 |
| Ankrd52 | -2.6171 | 2.10E-19 |
| Ap2a1 | -0.85471 | 0.0035143 |
| Apba1 | -2.2377 | 1.12E-13 |
| Apc | -0.9236 | 0.032203 |
| Apc2 | -1.5168 | 3.87E-08 |
| Arhgap21 | -0.68499 | 0.02963 |
| Arhgap23 | -1.4066 | 4.48E-07 |
| Arhgap32 | -1.5083 | 1.21E-07 |
| Arhgap33 | -1.5423 | 1.44E-08 |
| Arhgef11 | -0.88672 | 0.0041041 |
| Arhgef12 | -1.614 | 2.67E-08 |
| Arhgef7 | -0.73046 | 0.022259 |
| Arid1a | -0.85709 | 0.0047244 |
| Arid2 | -1.1582 | 2.30E-04 |
| Arl3 | 1.3003 | 0.042644 |
| Arnt2 | -0.81569 | 0.0061276 |
| Arrb1 | -2.2607 | 1.25E-14 |
| Atf4 | 0.90955 | 0.0018426 |
| Atf7ip | -1.1648 | 2.54E-04 |
| Atg2b | -0.91742 | 0.0031143 |
| Atp13a2 | -0.91386 | 0.0014477 |
| Atp2b4 | -1.8065 | 1.22E-07 |

| | | |
|---|---|---|
| Atp6v1f | 1.1571 | 0.0070723 |
| Atxn1 | -1.7071 | 2.53E-08 |
| Auts2 | -1.4483 | 5.27E-06 |
| B3gat1 | -1.0358 | 2.74E-04 |
| Bcl9l | -2.1071 | 5.34E-14 |
| Bcr | -1.1729 | 1.20E-04 |
| Birc6 | -1.6676 | 1.68E-06 |
| Bmpr2 | -1.8366 | 1.80E-10 |
| Brd4 | -0.96699 | 9.43E-04 |
| Brsk2 | -1.6028 | 1.84E-08 |
| Bsn | -1.4316 | 1.16E-07 |
| Bzrap1 | -1.4422 | 2.91E-07 |
| Cabin1 | -0.9853 | 8.95E-04 |
| Cacna1b | -1.0438 | 3.51E-04 |
| Cacna1d | -1.1573 | 7.50E-05 |
| Cacna1e | -3.2389 | 1.78E-29 |
| Cacnb1 | -0.88146 | 0.0022096 |
| Cadps | -1.1028 | 1.86E-04 |
| Calr | 0.77981 | 0.011416 |
| Camk2a | -1.3203 | 2.34E-06 |
| Camk2n1 | 1.2048 | 0.0047251 |
| Camkk2 | -0.8035 | 0.015828 |
| Camta2 | -0.86452 | 0.0029422 |
| Caskin1 | -1.2264 | 1.02E-05 |
| Cbx6 | -0.89286 | 0.0020713 |
| Cdc42bpa | -1.3816 | 1.56E-06 |
| Cdc42bpb | -1.068 | 1.74E-04 |
| Cdk5r1 | -1.13 | 1.21E-04 |
| Cdkl5 | -1.7429 | 2.41E-04 |
| Celf5 | -0.98589 | 5.47E-04 |
| Celsr2 | -2.6543 | 4.83E-22 |
| Chd3 | -1.0257 | 3.32E-04 |
| Chd4 | -1.0192 | 3.91E-04 |
| Chd5 | -1.9837 | 6.89E-13 |
| Chn1 | 1.0202 | 0.013432 |
| Chst2 | -0.81569 | 0.009029 |
| Cib1 | 0.85986 | 0.049137 |
| Cit | -1.7943 | 8.58E-10 |
| Clip3 | -1.8263 | 4.67E-11 |
| Cplx2 | -1.5087 | 1.07E-07 |
| Crebbp | -0.7759 | 0.023025 |

| | | |
|---|---|---|
| Crtc1 | -1.8185 | 7.36E-10 |
| Ctnnd2 | -1.3508 | 2.84E-06 |
| Cul9 | -0.81975 | 0.0071248 |
| Dag1 | -1.0394 | 0.0038422 |
| Dagla | -1.1917 | 3.02E-05 |
| Dicer1 | -0.90745 | 0.006052 |
| Dido1 | -0.71844 | 0.035347 |
| Dip2a | -1.3908 | 2.59E-06 |
| Dip2c | -1.1535 | 1.21E-04 |
| Diras2 | -1.642 | 9.61E-09 |
| Dlc1 | -1.4523 | 2.16E-04 |
| Dlg2 | -1.1379 | 0.029054 |
| Dlg5 | -1.6443 | 2.63E-08 |
| Dlgap1 | -0.91628 | 0.0027632 |
| Dlgap2 | -1.6514 | 2.38E-06 |
| Dlgap4 | -0.81301 | 0.0063561 |
| Dock3 | -1.3345 | 3.77E-06 |
| Dock4 | -1.6994 | 2.97E-09 |
| Dock9 | -0.71917 | 0.022231 |
| Dopey2 | -0.77225 | 0.013564 |
| Dot1l | -1.4948 | 4.60E-07 |
| Dscam | -1.2182 | 6.64E-05 |
| Dscaml1 | -1.579 | 6.28E-08 |
| Dst | -1.9159 | 1.89E-12 |
| Dtx1 | -0.89105 | 0.0033984 |
| Dusp8 | -1.2557 | 2.15E-05 |
| Dync1h1 | -1.293 | 1.87E-06 |
| Eef1a1 | 0.7361 | 0.015168 |
| Egr1 | 1.3514 | 0.041678 |
| Ehmt1 | -0.87492 | 0.0076169 |
| Ehmt2 | -0.69712 | 0.022259 |
| Eif4g1 | -1.0693 | 2.62E-04 |
| Elfn2 | -2.0025 | 2.47E-12 |
| Ep300 | -1.7418 | 3.57E-06 |
| Ep400 | -0.74364 | 0.018028 |
| Extl3 | -1.0429 | 8.09E-04 |
| Fam65a | -0.77782 | 0.0084696 |
| Fasn | -0.96646 | 9.71E-04 |
| Fat1 | -1.1292 | 1.12E-04 |
| Fat3 | -1.6101 | 2.80E-04 |
| Fat4 | -0.91802 | 0.0052075 |

| | | |
|---|---|---|
| Fbxl16 | -1.1851 | 3.11E-05 |
| Fbxl19 | -0.68252 | 0.027995 |
| Fbxo41 | -1.9457 | 9.14E-12 |
| Fcho1 | -1.3158 | 2.40E-06 |
| Fkbp3 | 1.0751 | 0.0023182 |
| Foxo3 | -1.6457 | 3.19E-07 |
| Fry | -0.74604 | 0.017487 |
| Gabbr2 | -0.6813 | 0.03242 |
| Gabrd | 1.254 | 5.88E-04 |
| Gcn1l1 | -0.84164 | 0.0051651 |
| Gnaz | -0.71206 | 0.025657 |
| Gpam | -1.5058 | 9.62E-05 |
| Gria1 | -1.0951 | 1.47E-04 |
| Grik3 | -2.7472 | 2.16E-20 |
| Grin2a | -2.4382 | 2.44E-15 |
| Grin2b | -2.8143 | 3.20E-23 |
| Grm5 | -1.6019 | 3.26E-06 |
| Gsk3b | -1.8308 | 3.59E-10 |
| Gtf3c1 | -0.76588 | 0.014121 |
| Hcn2 | -0.66068 | 0.044438 |
| Hdac4 | -2.2403 | 4.34E-14 |
| Hdlbp | -0.63688 | 0.045345 |
| Herc2 | -0.7433 | 0.02011 |
| Hint1 | 1.1502 | 4.66E-05 |
| Hipk2 | -2.8751 | 5.14E-22 |
| Hipk3 | -1.1623 | 3.62E-04 |
| Hivep3 | -2.3426 | 2.73E-15 |
| Htt | -1.2065 | 3.95E-05 |
| Igsf9b | -3.4207 | 2.71E-25 |
| Inpp4a | -1.6447 | 5.06E-09 |
| Ipo4 | -0.72149 | 0.022382 |
| Iqsec3 | -0.73531 | 0.015848 |
| Itsn1 | -1.363 | 2.85E-06 |
| Jph4 | -1.2203 | 1.03E-05 |
| Kalrn | -0.70557 | 0.03008 |
| Kcna2 | -2.4256 | 2.93E-14 |
| Kcnb1 | -1.1419 | 1.38E-04 |
| Kcnc3 | -1.4284 | 2.39E-07 |
| Kcnh1 | -0.70283 | 0.03664 |
| Kcnma1 | -1.8994 | 9.31E-11 |
| Kcnq2 | -2.011 | 6.23E-13 |

| | | |
|---|---|---|
| Kcnq3 | -2.2262 | 9.09E-06 |
| Kcnt1 | -1.5518 | 9.50E-07 |
| Kdm4b | -0.72916 | 0.018071 |
| Kdm6b | -1.635 | 1.05E-07 |
| Kif1a | -2.0308 | 8.85E-14 |
| Kif21b | -3.1536 | 2.95E-29 |
| Kif3c | -0.66282 | 0.034734 |
| Kif5c | -1.16 | 9.47E-04 |
| Kndc1 | -1.6188 | 3.22E-08 |
| Lhfpl4 | -1.657 | 2.63E-08 |
| Llgl1 | -1.0024 | 0.0069596 |
| Lrp1 | -1.3835 | 2.75E-07 |
| Lrp8 | -1.6524 | 2.89E-08 |
| Lrrc7 | -1.9173 | 2.58E-11 |
| Lrrc8b | -1.3136 | 0.0033947 |
| Macf1 | -1.6949 | 6.51E-10 |
| Magi2 | -1.1682 | 8.26E-05 |
| Man2a2 | -1.2286 | 4.89E-05 |
| Map1a | -0.92329 | 0.0015691 |
| Map2 | -1.3737 | 3.02E-05 |
| Map4k4 | -0.64308 | 0.046703 |
| Map7d1 | -0.74809 | 0.010356 |
| Mapk8ip3 | -1.0475 | 2.69E-04 |
| Mapkbp1 | -1.5382 | 2.88E-07 |
| Med13l | -1.6871 | 6.26E-09 |
| Mef2d | -1.6479 | 9.98E-09 |
| Mfhas1 | -1.4454 | 2.93E-04 |
| Mib1 | -1.7991 | 1.11E-06 |
| Mink1 | -0.83722 | 0.003784 |
| Mtss1l | -1.8821 | 6.87E-12 |
| Myo16 | -0.81211 | 0.0092924 |
| Myo18a | -0.93185 | 0.0017598 |
| Myt1l | -1.4557 | 3.75E-07 |
| Nat8l | -0.69467 | 0.028388 |
| Nav1 | -2.5017 | 2.39E-19 |
| Nav2 | -1.2763 | 1.08E-05 |
| Ncam1 | -1.139 | 8.33E-05 |
| Ncan | -2.0886 | 3.18E-07 |
| Ncdn | -0.80596 | 0.0068777 |
| Ncoa2 | -1.9094 | 9.66E-11 |
| Ncor2 | -1.3076 | 1.65E-06 |

| | | |
|---|---|---|
| Ncs1 | -1.1363 | 1.04E-04 |
| Ndst1 | -2.0591 | 1.37E-12 |
| Neurl4 | -0.68484 | 0.027722 |
| Nf1 | -0.81232 | 0.013179 |
| Nfic | -1.8969 | 8.50E-08 |
| Nfix | -2.1257 | 6.82E-14 |
| Nfyb | 0.76901 | 0.032939 |
| Nlgn2 | -1.7617 | 2.06E-10 |
| Nlgn3 | -1.0477 | 3.30E-04 |
| Nos1 | -2.6048 | 4.79E-19 |
| Nr3c1 | 0.69322 | 0.029412 |
| Nrxn2 | -1.0059 | 2.95E-04 |
| Nsd1 | -1.4533 | 1.14E-06 |
| Ntrk2 | -1.2658 | 9.38E-06 |
| Ntrk3 | -1.5594 | 1.07E-07 |
| Pacs1 | -0.63096 | 0.045845 |
| Pacs2 | -1.5306 | 2.26E-07 |
| Pak6 | -0.87999 | 0.005048 |
| Pcdh10 | -0.73769 | 0.021765 |
| Pcdh7 | -1.5327 | 2.39E-07 |
| Pcdh9 | -0.94987 | 0.0098835 |
| Pclo | -0.94056 | 0.021478 |
| Pcnx | -1.1259 | 1.04E-04 |
| Pde4dip | -0.86646 | 0.003836 |
| Pdzd2 | -2.1957 | 1.68E-12 |
| Peg3 | -1.6863 | 6.55E-06 |
| Phactr1 | -1.2008 | 5.90E-05 |
| Pik3r2 | -0.67125 | 0.025844 |
| Pip5k1c | -0.81111 | 0.0062155 |
| Pitpnm2 | -0.81397 | 0.0059531 |
| Pkd1 | -1.4844 | 6.30E-08 |
| Plcb1 | -1.0091 | 0.012243 |
| Plch2 | -1.3135 | 2.33E-06 |
| Plec | -1.6992 | 3.15E-09 |
| Plxna1 | -2.4832 | 9.03E-19 |
| Plxna2 | -0.90869 | 0.0026747 |
| Plxna4 | -2.5597 | 8.76E-19 |
| Plxnb1 | -1.5718 | 0.0084313 |
| Polr2a | -1.7288 | 6.87E-10 |
| Ppfia3 | -0.76923 | 0.011741 |
| Ppm1e | -1.7687 | 4.80E-07 |

| | | |
|---|---|---|
| Prex1 | -1.2994 | 6.15E-06 |
| Prkce | -1.5933 | 1.49E-08 |
| Ptch1 | -1.2085 | 2.61E-04 |
| Ptma | 0.64182 | 0.041498 |
| Ptprf | -1.2055 | 2.03E-05 |
| Ptprg | -1.339 | 3.97E-06 |
| Ptprj | -0.94709 | 0.034016 |
| Ptprs | -0.84991 | 0.0025239 |
| Ptprt | -2.1728 | 1.18E-13 |
| Ralgapb | -1.1885 | 9.82E-05 |
| Ralgds | -0.62838 | 0.044748 |
| Rap1gap | -0.72686 | 0.014375 |
| Rapgef1 | -1.4066 | 1.12E-06 |
| Rc3h1 | -1.696 | 5.07E-05 |
| Rere | -1.1013 | 1.17E-04 |
| Rgs7bp | -1.3397 | 4.50E-05 |
| Rhob | -1.1091 | 9.81E-05 |
| Rhobtb2 | -0.78908 | 0.016064 |
| Rph3a | -1.025 | 0.016763 |
| Rpl13a | 1.2846 | 0.02828 |
| Rpsa | 1.3008 | 0.01084 |
| Rusc2 | -0.81143 | 0.0060804 |
| Ryr2 | -1.277 | 8.59E-06 |
| Sap130 | -1.9127 | 1.07E-07 |
| Sash1 | -1.2728 | 0.0028018 |
| Sbf1 | -1.249 | 5.81E-06 |
| Scn8a | -2.0776 | 1.37E-13 |
| Sec16a | -1.4046 | 2.34E-05 |
| Setd5 | -0.65707 | 0.042007 |
| Setx | -0.73826 | 0.036087 |
| Sgip1 | -1.5589 | 1.52E-06 |
| Shank1 | -2.3312 | 5.32E-18 |
| Shank2 | -2.3656 | 2.49E-12 |
| Sipa1l1 | -0.94573 | 0.0019158 |
| Sipa1l3 | -1.1012 | 5.37E-04 |
| Slc12a5 | -1.3615 | 1.47E-06 |
| Slc1a2 | -1.8138 | 6.33E-04 |
| Slc24a2 | -2.1914 | 1.01E-08 |
| Slc4a3 | -1.041 | 3.72E-04 |
| Slc4a8 | -3.1136 | 8.30E-24 |
| Slc8a1 | -2.1011 | 1.97E-13 |

| | | |
|---|---|---|
| Slc8a2 | -0.84259 | 0.0057985 |
| Smarcc2 | -0.72144 | 0.014884 |
| Smg1 | -1.4285 | 7.92E-06 |
| Snap91 | -0.99339 | 7.45E-04 |
| Snph | -1.0936 | 1.13E-04 |
| Snrk | -0.91424 | 0.0044137 |
| Sobp | -1.3723 | 2.00E-06 |
| Sod1 | 1.1155 | 5.09E-04 |
| Sort1 | -1.3916 | 1.15E-06 |
| Spen | -2.8082 | 1.43E-22 |
| Spred1 | -1.1664 | 8.06E-04 |
| Sptan1 | -0.74492 | 0.010955 |
| Sptb | -2.5322 | 1.24E-18 |
| Sptbn1 | -1.9387 | 9.57E-13 |
| Srcin1 | -1.8719 | 1.35E-11 |
| Srgap3 | -2.2373 | 1.25E-15 |
| Srpk1 | -0.82186 | 0.012568 |
| Stox2 | -1.7705 | 4.23E-10 |
| Stxbp1 | -0.66569 | 0.033036 |
| Stxbp5 | -1.2249 | 3.95E-05 |
| Syn1 | -0.87836 | 0.0025865 |
| Syngap1 | -1.3485 | 1.48E-06 |
| Syngr1 | -1.1015 | 1.96E-04 |
| Synj1 | -0.91398 | 0.0029029 |
| Synpo | -0.97487 | 0.0011438 |
| Syt7 | -1.7253 | 9.16E-10 |
| Taok1 | -1.557 | 9.54E-05 |
| Taok2 | -0.68421 | 0.023493 |
| Tln2 | -2.0372 | 1.70E-12 |
| Tmem63b | -0.6344 | 0.044847 |
| Tmem8b | -1.0623 | 9.06E-04 |
| Tmod2 | -1.1094 | 2.10E-04 |
| Tnik | -1.4905 | 1.67E-07 |
| Tnk2 | -1.1923 | 1.75E-05 |
| Tnks | -1.74 | 1.14E-08 |
| Tnrc18 | -2.5252 | 2.72E-20 |
| Tnrc6b | -0.69551 | 0.03138 |
| Tppp | -1.1744 | 2.63E-04 |
| Trim2 | -1.6289 | 5.32E-06 |
| Trio | -1.2775 | 8.36E-06 |
| Trrap | -1.066 | 1.47E-04 |

| | | |
|---|---|---|
| Ttbk1 | -2.2519 | 8.58E-14 |
| Ttbk2 | -1.7357 | 1.83E-07 |
| Tulp4 | -1.9328 | 1.11E-11 |
| Ubap2l | -0.66165 | 0.044721 |
| Ube2a | 0.74463 | 0.017234 |
| Ube2o | -1.0198 | 5.71E-04 |
| Ube3b | -1.3705 | 5.53E-06 |
| Ulk1 | -0.74141 | 0.017887 |
| Ulk2 | -0.94126 | 0.0032289 |
| Unc13a | -1.7334 | 5.99E-10 |
| Unc13b | -1.2554 | 5.09E-04 |
| Usp22 | -1.4608 | 2.77E-07 |
| Wdr13 | -1.1992 | 4.52E-05 |
| Wdr6 | -0.85825 | 0.0042495 |
| Wnk2 | -1.517 | 2.11E-08 |
| Wwc1 | -0.89706 | 0.0023697 |
| Zc3h4 | -1.6975 | 9.28E-09 |
| Zcchc14 | -1.7513 | 8.32E-09 |
| Zfhx2 | -2.0067 | 1.71E-05 |
| Zhx3 | -0.95424 | 0.0045413 |
| Zmiz1 | -1.0482 | 2.11E-04 |
| Zyg11b | -1.5487 | 2.06E-05 |

| Associated_Gene_Name | log2FoldChange | padj |
|---|---|---|
| AABR07006627.1 | 0.80811 | 0.030306 |
| AABR07010030.1 | 1.1593 | 0.0082537 |
| AABR07011857.1 | -1.6556 | 0.014521 |
| AABR07012054.1 | 1.7583 | 0.0081555 |
| AABR07015079.1 | Inf | 0.01557 |
| AABR07016640.1 | 1.7923 | 0.046343 |
| AABR07016841.1 | 1.9033 | 0.02804 |
| AABR07018124.1 | #NAME? | 6.42E-04 |
| AABR07019083.1 | -1.8538 | 7.24E-09 |
| AABR07019093.1 | -1.2335 | 0.0096057 |
| AABR07019412.1 | -2.5795 | 1.23E-09 |
| AABR07020786.1 | -1.1735 | 0.012027 |
| AABR07020860.1 | -1.3602 | 0.02204 |
| AABR07026311.1 | 0.83371 | 0.017605 |
| AABR07026997.1 | 2.4556 | 0.0016753 |
| AABR07030603.1 | -1.7015 | 3.30E-04 |
| AABR07030781.1 | 0.90691 | 0.0064628 |
| AABR07031533.1 | -0.78683 | 0.015349 |
| AABR07031734.2 | -1.0331 | 0.029051 |
| AABR07043601.1 | -1.5406 | 0.019628 |
| AABR07043823.2 | 1.4306 | 0.010317 |
| AABR07045621.1 | 1.1598 | 0.017827 |
| AABR07048211.1 | -2.9885 | 2.49E-12 |
| AABR07049948.1 | -2.7308 | 6.39E-04 |
| AABR07051947.1 | 1.0351 | 0.0036638 |
| AABR07053185.1 | 0.76075 | 0.045634 |
| AABR07055280.1 | -1.4127 | 0.019101 |
| AABR07057196.1 | 1.1981 | 0.036483 |
| AABR07061755.1 | -1.4151 | 1.16E-04 |
| AABR07064349.1 | -2.0629 | 5.28E-07 |
| AABR07065406.1 | 1.0776 | 0.0016637 |
| AABR07065531.1 | -1.47 | 0.0089666 |
| AABR07067499.1 | 1.4129 | 1.65E-04 |
| AABR07067736.1 | 1.8156 | 0.034822 |
| AABR07068242.1 | -1.7814 | 0.0013504 |
| AABR07071445.1 | -2.2111 | 5.11E-07 |
| AABR07072537.1 | -1.181 | 1.76E-05 |
| AABR07072748.1 | -2.0468 | 0.013061 |
| AABR07073039.1 | 1.8382 | 0.020499 |
| Aadat | 1.5931 | 0.0017177 |
| Aamdc | 1.4224 | 0.0016391 |
| Abat | -1.3515 | 2.40E-06 |
| Abca5 | -0.89195 | 0.012404 |

| | | |
|---|---|---|
| Abca7 | -1.6807 | 3.16E-07 |
| Abcg4 | -0.74446 | 0.018636 |
| Abhd14b | 0.92986 | 0.02963 |
| Abhd15 | -2.5791 | 0.021929 |
| Abhd2 | -1.5063 | 1.02E-05 |
| Abhd3 | 0.92878 | 0.0024345 |
| Abl2 | -1.7699 | 8.22E-08 |
| Ablim2 | -0.7831 | 0.017983 |
| Ablim3 | -1.2557 | 5.01E-04 |
| AC096601.1 | -1.0613 | 0.040681 |
| AC115420.2 | -1.6542 | 9.93E-04 |
| AC126148.1 | -1.2518 | 0.013002 |
| AC128848.1 | 1.3781 | 1.11E-04 |
| AC134160.1 | -2.8813 | 0.0049438 |
| AC135310.2 | -3.0505 | 0.0010069 |
| AC135400.1 | 1.5206 | 8.18E-06 |
| Acaa2 | 1.1735 | 1.64E-04 |
| Acaca | -1.258 | 8.89E-05 |
| Acacb | -1.3873 | 0.03256 |
| Acadl | 0.88366 | 0.0065307 |
| Acadm | 0.77569 | 0.038086 |
| Acap2 | -1.1332 | 0.020327 |
| Acap3 | -1.0443 | 2.85E-04 |
| Acot13 | 0.99675 | 0.0030684 |
| Acot9 | 0.78579 | 0.0096283 |
| Acox3 | -0.98276 | 0.003807 |
| Acp1 | 0.88395 | 0.0031108 |
| Actn4 | -0.84013 | 0.0047889 |
| Actr6 | 0.92988 | 0.0025736 |
| Acvr1b | -0.85496 | 0.014373 |
| Acvr2b | -2.2646 | 3.18E-06 |
| Acyp1 | 0.83161 | 0.015122 |
| Adam11 | -1.3875 | 1.31E-06 |
| Adam12 | -2.4733 | 2.09E-07 |
| Adam19 | -1.0425 | 5.26E-04 |
| Adam22 | -1.6575 | 3.17E-08 |
| Adam23 | -1.5619 | 1.37E-07 |
| Adamts10 | -1.5338 | 1.04E-04 |
| Adamts17 | -1.5898 | 6.66E-05 |
| Adamts18 | -1.3671 | 0.037335 |
| Adamts20 | -1.0667 | 0.049768 |
| Adamts3 | -1.3261 | 4.52E-04 |
| Adamts7 | -1.5095 | 3.02E-04 |
| Adamtsl1 | -1.8501 | 0.0014262 |

| | | |
|---|---|---|
| Adcy9 | -1.8383 | 2.75E-09 |
| Adcyap1r1 | -1.5623 | 8.70E-08 |
| Add2 | -2.0021 | 9.53E-13 |
| Adgra2 | -1.2778 | 0.0072791 |
| Adgrb1 | -0.83623 | 0.0036799 |
| Adgrb2 | -0.93477 | 7.08E-04 |
| Adgrl1 | -1.6711 | 9.11E-10 |
| Adhfe1 | 1.0364 | 0.002779 |
| Adi1 | 0.75314 | 0.045538 |
| Adipor2 | -0.77773 | 0.025911 |
| Adora1 | -0.90026 | 0.00335 |
| Aen | -0.81014 | 0.01351 |
| Ager | -1.6823 | 0.022831 |
| Agfg2 | -0.69304 | 0.040218 |
| Ago2 | -1.8041 | 3.24E-06 |
| Ago3 | -2.5631 | 1.82E-06 |
| Agps | -1.3909 | 0.010804 |
| Ahcyl2 | -0.66443 | 0.04612 |
| Ahi1 | -0.70132 | 0.033346 |
| Aimp1 | 1.2184 | 0.015046 |
| Ak6 | 1.0939 | 0.0015563 |
| Akap13 | -0.89098 | 0.03181 |
| Akap2 | -1.189 | 0.0016877 |
| Akap7 | 0.87211 | 0.0036 |
| Akip1 | 1.1933 | 0.0026008 |
| Akna | -1.2118 | 0.001058 |
| Akt2 | -0.71874 | 0.034009 |
| Aktip | 0.65235 | 0.042585 |
| Alb | 4.0764 | 0.026652 |
| Aldh5a1 | -0.87656 | 0.040516 |
| Aldh6a1 | 0.75072 | 0.016246 |
| Alkbh5 | -0.93901 | 0.011807 |
| Alkbh7 | 1.2547 | 3.53E-04 |
| Alox12 | 1.7121 | 0.044484 |
| Alpk1 | -1.3794 | 0.0019925 |
| Amer1 | -1.8281 | 4.67E-05 |
| Amer2 | -1.3435 | 2.84E-05 |
| Amn1 | 0.95689 | 0.0020347 |
| Amot | -1.2837 | 5.26E-04 |
| Amotl1 | -1.3324 | 0.0099856 |
| Amz1 | -0.99151 | 0.0065942 |
| Anapc10 | 1.0173 | 0.0022794 |
| Anapc11 | 1.095 | 0.016876 |
| Anapc13 | 1.6637 | 1.74E-08 |

| | | |
|---|---|---|
| Anapc15 | 1.1053 | 0.012574 |
| Ankfy1 | -1.271 | 0.0042495 |
| Ankle1 | -3.0085 | 1.01E-06 |
| Ankrd13b | -1.0232 | 3.75E-04 |
| Ankrd24 | 1.0177 | 0.02273 |
| Ankrd28 | -0.85062 | 0.0097159 |
| Ankrd33b | -1.1366 | 0.0096283 |
| Ankrd45 | -0.92271 | 0.037761 |
| Ankrd49 | 0.81414 | 0.012306 |
| Ankrd63 | -2.5033 | 0.0075374 |
| Anks1a | -0.82174 | 0.014002 |
| Anks1b | -0.80583 | 0.011721 |
| Anks6 | -0.87219 | 0.014267 |
| Anln | -1.3816 | 0.0037739 |
| Antxr1 | -1.4357 | 3.75E-05 |
| Anxa5 | 0.84028 | 0.0084903 |
| Anxa7 | 0.7529 | 0.029056 |
| Ap1g2 | -1.7336 | 4.68E-04 |
| Ap3s1 | 1.0743 | 1.76E-04 |
| Ap4s1 | 1.0481 | 0.0010839 |
| Ap5z1 | -1.518 | 4.79E-04 |
| Apaf1 | -1.3601 | 3.35E-05 |
| Apba2 | -0.85012 | 0.0037205 |
| Apoo | 0.82863 | 0.010286 |
| Apopt1 | 1.1909 | 1.31E-04 |
| Aprt | 0.68874 | 0.046343 |
| Araf | 0.65899 | 0.034647 |
| Arap1 | -1.1588 | 2.48E-04 |
| Arfgef2 | -1.1949 | 1.02E-04 |
| Arfgef3 | -2.2641 | 5.08E-15 |
| Arhgap26 | -1.5875 | 1.89E-06 |
| Arhgap35 | -1.9455 | 6.28E-12 |
| Arhgap6 | -1.1834 | 0.0032606 |
| Arhgef1 | -1.4138 | 2.58E-06 |
| Arhgef10 | -1.3994 | 2.26E-04 |
| Arhgef3 | 0.84147 | 0.0086457 |
| Arhgef33 | 1.0643 | 0.040999 |
| Arid3a | -1.3329 | 0.0025064 |
| Arid5b | -1.1354 | 0.0064906 |
| Arl1 | 0.66795 | 0.036684 |
| Arl2bp | 0.65559 | 0.037001 |
| Arl4a | 0.745 | 0.022383 |
| Arl6ip1 | 0.62201 | 0.044692 |
| Arl6ip5 | 0.78187 | 0.0097251 |

| | | |
|---|---|---|
| Armcx1 | 0.80675 | 0.028613 |
| Arpc3 | 0.86409 | 0.0041229 |
| Arpc5l | 0.84497 | 0.0056262 |
| Arse | -1.169 | 3.03E-04 |
| Asb1 | -1.2539 | 5.41E-05 |
| Asb7 | -0.92772 | 0.030529 |
| Ascc3 | 0.81738 | 0.019538 |
| Asnsd1 | 0.97764 | 0.0013162 |
| Aspdh | -1.0247 | 0.037966 |
| Asxl3 | -1.4409 | 3.40E-04 |
| Atad1 | 0.79177 | 0.0067248 |
| Atat1 | -1.0009 | 9.11E-04 |
| Atf7 | -1.2258 | 0.00746 |
| Atg12 | 0.75935 | 0.013376 |
| Atox1 | 1.1614 | 8.33E-05 |
| Atp11a | -1.1222 | 0.0021357 |
| Atp13a3 | -2.0716 | 2.48E-05 |
| Atp1b3 | 0.8292 | 0.0065177 |
| Atp5e | 1.5902 | 7.30E-09 |
| Atp5g2 | 0.98834 | 0.0028744 |
| Atp5g3 | 0.73739 | 0.032928 |
| Atp5h | 1.0335 | 3.00E-04 |
| Atp5i | 1.7103 | 9.62E-09 |
| Atp5j | 1.1018 | 0.0046034 |
| Atp5j2 | 1.4652 | 8.01E-07 |
| Atp5l | 1.2901 | 2.72E-06 |
| Atp5o | 0.90851 | 0.0019931 |
| Atp6v0b | 0.83139 | 0.029051 |
| Atp6v0e1 | 1.359 | 1.03E-05 |
| Atp6v1d | 0.68639 | 0.027095 |
| Atp6v1e1 | 0.71845 | 0.034564 |
| Atp6v1g1 | 1.0655 | 1.76E-04 |
| Atp6v1h | 0.78267 | 0.0098561 |
| Atp8a1 | -1.2593 | 0.0039142 |
| Atp8a2 | -2.1133 | 5.66E-11 |
| Atp8b2 | -1.1894 | 8.26E-05 |
| Atpif1 | 1.3162 | 4.71E-04 |
| Atraid | 1.2546 | 2.26E-05 |
| Atrnl1 | -1.3074 | 3.50E-04 |
| Atxn1l | -1.7225 | 9.47E-05 |
| Atxn2l | -1.2264 | 8.24E-06 |
| Aurkb | -1.8424 | 4.95E-05 |
| Avl9 | -1.4816 | 4.35E-06 |
| AY172581.24 | 2.0527 | 0.0099976 |

| | | |
|---|---|---|
| AY172581.9 | 1.7252 | 0.01457 |
| B2m | 1.1425 | 1.83E-04 |
| B3galnt1 | 0.68845 | 0.02859 |
| B3galt6 | -1.1889 | 0.0034304 |
| B3gat3 | 0.94828 | 0.0044939 |
| B3gntl1 | -1.2795 | 0.02188 |
| B4galnt1 | -0.95225 | 0.0021389 |
| B4galt4 | 0.78116 | 0.040434 |
| B9d1 | 1.1875 | 0.0079089 |
| Bace1 | -0.81399 | 0.010182 |
| Bace2 | -0.75113 | 0.03223 |
| Bach2 | -2.0603 | 2.63E-07 |
| Bag1 | 0.82629 | 0.011411 |
| Bahcc1 | -2.5151 | 2.90E-16 |
| Bahd1 | -1.3895 | 1.12E-05 |
| Bambi | 1.3041 | 0.0040171 |
| Bard1 | -1.3552 | 0.0109 |
| Bcap29 | 0.72129 | 0.023617 |
| Bcas2 | 0.89976 | 0.0097617 |
| Bccip | 0.76663 | 0.020494 |
| Bcl11b | -1.2134 | 4.58E-04 |
| Bcl2 | -1.4332 | 0.018758 |
| Bcl2l13 | -0.90803 | 0.012285 |
| Bcl9 | -1.0467 | 4.99E-04 |
| Bdh2 | 0.82897 | 0.047222 |
| Bean1 | -1.4917 | 0.0047315 |
| Bend3 | -1.4903 | 0.012285 |
| Bet1 | 0.86446 | 0.0048759 |
| Bicd1 | -1.1351 | 0.0062541 |
| Bloc1s1 | 1.0207 | 9.37E-04 |
| Bloc1s2 | 1.1843 | 4.32E-05 |
| Bloc1s5 | 0.85515 | 0.011269 |
| Bmp2k | -0.79906 | 0.039015 |
| Bmp3 | -1.4131 | 2.19E-04 |
| Bmp6 | 1.7941 | 0.020745 |
| Bnip3 | 0.64681 | 0.034677 |
| Bola3 | 0.93076 | 0.0023697 |
| Borcs5 | -0.75329 | 0.040331 |
| Borcs8 | 0.95454 | 0.0032762 |
| Bpgm | 0.94151 | 0.0014792 |
| Bphl | 0.85889 | 0.020745 |
| Braf | -1.2268 | 2.00E-04 |
| Brap | -1.2256 | 1.68E-04 |
| Brd3 | -1.1435 | 1.68E-04 |

| | | |
|---|---|---|
| Bre | 0.88169 | 0.0049013 |
| Brk1 | 1.1271 | 7.02E-05 |
| Brpf3 | -1.1737 | 4.95E-04 |
| Brwd3 | -1.1048 | 0.027722 |
| Btbd7 | -1.1447 | 0.035047 |
| Btbd9 | -0.78046 | 0.027821 |
| Btf3 | 0.75629 | 0.016454 |
| Btrc | -0.76228 | 0.020618 |
| Bub1b | -1.271 | 0.0033795 |
| Bud31 | 0.93954 | 0.0026979 |
| C1d | 0.92452 | 0.0019097 |
| C1galt1c1 | 1.0199 | 0.0010733 |
| C1ql3 | -0.95775 | 0.0013222 |
| C2cd2 | -1.0982 | 0.0019515 |
| C2cd3 | -0.71931 | 0.04824 |
| Cab39l | 1.1478 | 4.83E-04 |
| Cachd1 | -1.3935 | 2.48E-06 |
| Cacna1c | -1.902 | 5.38E-11 |
| Cacna1h | -1.7378 | 3.03E-10 |
| Cacna2d1 | -1.0781 | 0.023758 |
| Cacna2d2 | -1.5242 | 3.57E-06 |
| Cacnb4 | -1.0741 | 0.019992 |
| Cacng2 | -0.75073 | 0.029477 |
| Cacng4 | -1.6292 | 3.17E-08 |
| Cacybp | 0.67498 | 0.027687 |
| Cad | -0.99913 | 0.0020574 |
| Cadm1 | -0.84996 | 0.0053974 |
| Calm2 | 0.72093 | 0.017556 |
| Caln1 | -2.7541 | 4.93E-20 |
| Calr3 | 1.2135 | 0.014578 |
| Camk1d | -1.1087 | 3.44E-04 |
| Camk2g | -0.97096 | 0.0012911 |
| Capn15 | -0.9235 | 0.0040954 |
| Carm1 | -1.1486 | 9.21E-05 |
| Casc3 | -0.92061 | 0.0031695 |
| Caskin2 | -1.0397 | 0.034011 |
| Cbfa2t2 | -1.2635 | 9.13E-05 |
| Cbfa2t3 | -1.4977 | 5.82E-06 |
| Cbl | -2.4297 | 3.19E-14 |
| Cblb | -0.98215 | 0.010286 |
| Cbr4 | 0.93119 | 0.0081161 |
| Cbx4 | -0.64998 | 0.048291 |
| Cby1 | 0.78496 | 0.012599 |
| Cc2d1a | -0.69482 | 0.019541 |

| | | |
|---|---|---|
| Ccar1 | 0.77351 | 0.016401 |
| Ccbe1 | -2.433 | 3.40E-04 |
| Ccdc103 | -2.2074 | 0.041448 |
| Ccdc12 | 0.9875 | 0.0072583 |
| Ccdc13 | -1.0795 | 0.024767 |
| Ccdc142 | -1.3147 | 4.28E-05 |
| Ccdc149 | -0.76455 | 0.042547 |
| Ccdc15 | -1.5087 | 0.021395 |
| Ccdc157 | -1.4276 | 4.92E-05 |
| Ccdc175 | 1.5492 | 0.0052253 |
| Ccdc177 | -1.0383 | 0.0060541 |
| Ccdc188 | -1.6148 | 3.84E-04 |
| Ccdc190 | 2.0197 | 0.035047 |
| Ccdc34 | 0.79103 | 0.020768 |
| Ccdc43 | 0.77735 | 0.015884 |
| Ccdc50 | -0.89927 | 0.038861 |
| Ccdc53 | 1.1059 | 7.84E-04 |
| Ccdc58 | 1.2312 | 4.48E-04 |
| Ccdc59 | 1.0025 | 0.0028792 |
| Ccdc6 | -1.6165 | 1.26E-06 |
| Ccdc62 | 0.98586 | 0.036872 |
| Ccdc85a | -1.006 | 0.0017398 |
| Ccdc88a | -0.96296 | 0.04655 |
| Ccdc88c | -1.138 | 0.025832 |
| Ccdc90b | 0.80657 | 0.019337 |
| Ccnf | -1.1892 | 0.011793 |
| Ccni | 1.018 | 3.26E-04 |
| Cct2 | 0.86782 | 0.0033124 |
| Cct3 | 0.77306 | 0.0486 |
| Cct4 | 0.71484 | 0.045461 |
| Cct5 | 0.76309 | 0.013077 |
| Cct8 | 0.72898 | 0.016971 |
| Cd24 | 0.65838 | 0.039992 |
| Cd302 | 1.4098 | 0.035442 |
| Cd47 | -0.76462 | 0.03008 |
| Cd59 | 1.2501 | 9.92E-06 |
| Cd63 | 1.0616 | 2.03E-04 |
| Cd81 | 0.70468 | 0.023557 |
| Cdadc1 | 1.1119 | 2.11E-04 |
| Cdan1 | -0.75575 | 0.03618 |
| Cdc123 | 0.69795 | 0.031622 |
| Cdc26 | 0.90044 | 0.0032898 |
| Cdh22 | -0.96616 | 0.0048676 |
| Cdh23 | -2.0049 | 0.0015603 |

| | | |
|---|---|---|
| Cdh24 | -1.2889 | 7.20E-05 |
| Cdhr1 | -1.9895 | 2.77E-04 |
| Cdk12 | -0.75575 | 0.037554 |
| Cdk2 | -1.0833 | 0.015884 |
| Cdon | -1.0632 | 0.0035273 |
| Cdr2l | -1.5367 | 1.03E-05 |
| Cds2 | -1.942 | 3.00E-11 |
| Cdv3 | -1.4928 | 1.40E-06 |
| Cdyl2 | -1.6438 | 9.07E-05 |
| Cecr2 | -1.505 | 0.022607 |
| Cecr6 | -1.4139 | 1.32E-05 |
| Celf2 | -1.0922 | 1.71E-04 |
| Celf3 | -0.64828 | 0.032446 |
| Celsr1 | -1.8878 | 0.0028294 |
| Cep164 | -0.97726 | 0.0068052 |
| Cep170b | -1.4876 | 9.46E-08 |
| Cep19 | 0.64977 | 0.046052 |
| Cep350 | -1.3232 | 4.73E-04 |
| Cep85l | -1.0429 | 0.016454 |
| Cept1 | 0.99577 | 8.23E-04 |
| Cers6 | -1.1803 | 0.026934 |
| Cfap36 | 0.91997 | 0.0018434 |
| Cfb | -4.0817 | 0.0019033 |
| Cfdp1 | 0.74376 | 0.018929 |
| Cfl2 | 0.64915 | 0.033113 |
| Cgrrf1 | 1.1632 | 2.53E-04 |
| Chchd1 | 1.2098 | 4.87E-05 |
| Chchd3 | 0.92857 | 0.0022148 |
| Chchd5 | 1.1118 | 8.98E-04 |
| Chd7 | -1.5078 | 0.0062608 |
| Cherp | -0.7956 | 0.0077685 |
| Chmp2a | 0.80392 | 0.010844 |
| Chmp3 | 0.81388 | 0.0081281 |
| Chmp5 | 0.79485 | 0.0095834 |
| Chpf2 | -0.65137 | 0.04655 |
| Chrac1 | 0.98618 | 0.005804 |
| Chrd | -1.4553 | 1.75E-05 |
| Chrna4 | -1.1607 | 2.18E-04 |
| Chrna5 | -1.6562 | 3.60E-05 |
| Chrna7 | -1.3357 | 7.83E-04 |
| Chrnb2 | -1.4072 | 2.73E-06 |
| Chst1 | -0.6832 | 0.024133 |
| Chst11 | -1.4637 | 2.29E-05 |
| Chst3 | -2.0449 | 2.64E-04 |

| | | |
|---|---|---|
| Chst7 | -1.5018 | 3.58E-06 |
| Chtf18 | -1.2239 | 0.0044329 |
| Churc1 | 1.0732 | 0.0019202 |
| Cinp | 0.81858 | 0.0081122 |
| Cisd1 | 0.77268 | 0.012337 |
| Cisd2 | 0.77538 | 0.0090533 |
| Cisd3 | 0.88021 | 0.025089 |
| Ckmt1b | 1.4941 | 0.0033042 |
| Cks1b | 0.88402 | 0.020281 |
| Clasrp | -0.98199 | 0.0014388 |
| Clcc1 | 0.64664 | 0.049966 |
| Clcn1 | -1.5834 | 0.024107 |
| Clcn5 | -1.6472 | 0.0085176 |
| Clcn6 | -1.2228 | 1.76E-04 |
| Clcn7 | -0.91324 | 0.004024 |
| Cldn1 | 4.404 | 5.14E-05 |
| Cldn10 | 2.1821 | 9.52E-05 |
| Cldnd1 | 0.90716 | 0.0020929 |
| Clic1 | 1.2426 | 0.044988 |
| Clmn | -2.7964 | 3.17E-14 |
| Clock | -1.5316 | 1.65E-04 |
| Clpb | -0.67831 | 0.046571 |
| Clspn | -1.2961 | 0.0064434 |
| Clstn2 | -1.9992 | 2.97E-11 |
| Clu | 1.379 | 0.004859 |
| Cluap1 | 1.0148 | 0.010603 |
| Cluh | -1.3273 | 4.05E-06 |
| Cmbl | 1.0296 | 0.023062 |
| Cmc2 | 1.7414 | 2.14E-06 |
| Cmip | -1.598 | 5.77E-09 |
| Cml1 | 1.101 | 0.028666 |
| Cml5 | 2.6718 | 0.0033348 |
| Cmtm4 | -1.8101 | 9.31E-06 |
| Cnbp | 0.65301 | 0.041337 |
| Cnih1 | 0.87733 | 0.00303 |
| Cnih4 | 0.71542 | 0.045098 |
| Cnnm1 | -0.94837 | 0.0032749 |
| Cnnm4 | -0.96791 | 0.01316 |
| Cnot7 | 0.74273 | 0.015848 |
| Cntn1 | -0.74995 | 0.021151 |
| Cntn2 | -1.5702 | 3.14E-06 |
| Cntnap1 | -1.1133 | 1.20E-04 |
| Coa3 | 0.72054 | 0.029775 |
| Col11a2 | -2.3651 | 6.53E-05 |

| | | |
|---|---|---|
| Col16a1 | -2.3064 | 0.0064756 |
| Col19a1 | -1.8074 | 7.16E-05 |
| Col20a1 | -1.6541 | 0.0044137 |
| Col23a1 | -2.2984 | 1.31E-05 |
| Col25a1 | -1.2899 | 4.27E-04 |
| Col27a1 | -2.9573 | 3.64E-05 |
| Col4a1 | -1.2842 | 0.01224 |
| Col4a2 | -1.8569 | 1.89E-10 |
| Col5a1 | -2.4122 | 0.0027834 |
| Col5a3 | -2.9446 | 4.28E-05 |
| Col6a3 | -2.0158 | 5.37E-04 |
| Col6a4 | -2.2936 | 2.53E-08 |
| Col6a6 | -2.1724 | 0.034931 |
| Commd1 | 1.1482 | 0.010285 |
| Commd3 | 0.91193 | 0.0021389 |
| Commd6 | 1.4229 | 1.98E-06 |
| Commd7 | 0.84218 | 0.0094277 |
| Commd8 | 0.92527 | 0.0022096 |
| Commd9 | 0.88997 | 0.004804 |
| Coprs | 1.2957 | 3.51E-04 |
| Cops4 | 0.74958 | 0.029051 |
| Coq10b | 0.94397 | 0.016085 |
| Coq3 | 0.8957 | 0.0046311 |
| Coro2a | -1.3499 | 1.15E-04 |
| Coro7 | -0.83087 | 0.0058671 |
| Coro7 | -1.7088 | 3.47E-05 |
| Cotl1 | -0.79037 | 0.015415 |
| Cox10 | -1.0606 | 0.026221 |
| Cox16 | 0.94973 | 0.0035524 |
| Cox17 | 1.8648 | 8.49E-08 |
| Cox5a | 0.93586 | 0.001347 |
| Cox5b | 0.97287 | 0.0012184 |
| Cox6a1 | 1.1254 | 1.65E-04 |
| Cox6b1 | 1.4945 | 7.34E-06 |
| Cox6b1 | 1.9899 | 0.0022134 |
| Cox6c | 1.4585 | 6.65E-08 |
| Cox7a2l | 0.9833 | 7.92E-04 |
| Cox7a2l2 | 1.6094 | 1.39E-08 |
| Cox7c | 1.1331 | 0.0025204 |
| Cox8a | 1.2073 | 1.31E-05 |
| Cpd | -0.89726 | 0.0042495 |
| Cpeb2 | -1.4121 | 6.36E-06 |
| Cpeb3 | -1.9876 | 6.26E-09 |
| Cpeb4 | -0.92115 | 0.036872 |

| | | |
|---|---|---|
| Cpne2 | -1.0784 | 8.36E-04 |
| Cpsf1 | -0.76387 | 0.012537 |
| Cramp1 | -1.4133 | 1.66E-04 |
| Crb2 | -1.7296 | 0.0033795 |
| Crcp | 1.1257 | 2.23E-04 |
| Creb5 | -1.6562 | 0.0060733 |
| Creg1 | 1.0691 | 3.40E-04 |
| Creg2 | -1.745 | 3.27E-07 |
| Crem | 1.008 | 0.0027325 |
| Crhbp | 1.5826 | 5.44E-06 |
| Crim1 | -1.0497 | 6.46E-04 |
| Cript | 1.0035 | 6.14E-04 |
| Crkl | -0.82553 | 0.015554 |
| Crocc | -1.4082 | 2.31E-05 |
| Crtc3 | -1.319 | 0.0060901 |
| Cryga | 1.1451 | 0.025211 |
| Cryzl1 | 0.71275 | 0.029498 |
| Csmd1 | -1.0787 | 3.56E-04 |
| Csmd2 | -2.8209 | 9.66E-11 |
| Csmd3 | -1.2126 | 2.82E-04 |
| Csnk1g1 | -1.1401 | 0.022831 |
| Cspg4 | -1.216 | 8.68E-05 |
| Csrnp3 | -1.6809 | 3.23E-05 |
| Csrp1 | 0.6581 | 0.049653 |
| Cst3 | 1.3844 | 1.05E-04 |
| Cstb | 1.1379 | 8.82E-05 |
| Ctdsp2 | -1.0632 | 0.0038169 |
| Ctif | -1.1473 | 8.38E-04 |
| Ctnnd1 | -0.67667 | 0.040311 |
| Ctsh | 0.94487 | 0.0036611 |
| Ctsl | 0.88758 | 0.0033795 |
| Cttn | -1.1748 | 0.0010981 |
| Ctxn2 | 1.5188 | 0.027205 |
| Cuedc1 | -0.99404 | 0.0019296 |
| Cwc15 | 0.9964 | 0.0012559 |
| Cxcl12 | 1.1238 | 0.0046986 |
| Cyb5a | 1.2399 | 2.10E-05 |
| Cyba | 1.1132 | 0.046642 |
| Cycs | 0.69755 | 0.024656 |
| Cyp26b1 | 1.3151 | 0.042666 |
| Cyp2j16 | 3.2516 | 0.014088 |
| Cyp4f4 | -1.1813 | 0.033346 |
| Cyp4v3 | 1.2959 | 0.0081194 |
| Cyp7b1 | 0.95582 | 0.024002 |

| | | |
|---|---|---|
| Cystm1 | 0.85258 | 0.0061763 |
| Dab1 | -0.79412 | 0.019638 |
| Dad1 | 1.2028 | 2.34E-05 |
| Dap | 1.0464 | 0.0021509 |
| Dbi | 1.3218 | 0.0010355 |
| Dbn1 | -0.92045 | 0.0010425 |
| Dbnl | -1.0113 | 4.99E-04 |
| Dcaf15 | -0.88783 | 0.0051907 |
| Dcaf17 | -1.9072 | 5.22E-07 |
| Dcakd | -0.72492 | 0.017518 |
| Dcbld1 | -0.93149 | 0.014061 |
| Dcc | -2.1327 | 1.40E-11 |
| Dchs1 | -1.5727 | 1.06E-05 |
| Dclk2 | -1.0315 | 4.84E-04 |
| Dcp2 | -1.0343 | 0.032699 |
| Dctn3 | 0.92601 | 0.019674 |
| Dctn5 | 0.68367 | 0.031131 |
| Dctn6 | 0.77759 | 0.011659 |
| Dctpp1 | 1.094 | 0.023128 |
| Ddit3 | 0.90791 | 0.0038824 |
| Ddit4l | 1.0819 | 0.029843 |
| Ddt | 1.1229 | 4.14E-04 |
| Ddx17 | -0.96413 | 0.0011808 |
| Ddx54 | -0.9545 | 0.0020885 |
| Decr1 | 1.0577 | 0.0016486 |
| Degs1 | 0.6923 | 0.03359 |
| Dennd2a | -1.055 | 7.88E-04 |
| Dennd3 | -1.0722 | 0.0061175 |
| Dennd4b | -0.67092 | 0.042094 |
| Dennd6b | -0.76869 | 0.029421 |
| Dera | 0.80504 | 0.047709 |
| Derl1 | 0.65434 | 0.034405 |
| Dfnb31 | -0.92333 | 0.0045413 |
| Dgkd | -0.74556 | 0.015801 |
| Dgke | -0.84129 | 0.014054 |
| Dgkg | -1.3435 | 5.93E-06 |
| Dgkh | -2.7315 | 9.09E-12 |
| Dgki | -1.611 | 9.00E-08 |
| Dgkq | -1.0632 | 4.52E-04 |
| Dhcr24 | -1.3484 | 0.034726 |
| Dhrs7 | 1.3573 | 0.021012 |
| Dhrs7b | 0.9467 | 0.0070655 |
| Dhx33 | -1.0237 | 0.0056456 |
| Dhx37 | -0.92487 | 0.007296 |

| | | |
|---|---|---|
| Diaph1 | -1.2018 | 2.00E-04 |
| Diaph3 | -2.5962 | 0.0039755 |
| Disc1 | -1.5204 | 0.038729 |
| Dlat | -0.79547 | 0.014402 |
| Dll3 | -1.3913 | 0.026287 |
| Dlx1 | 0.69035 | 0.032361 |
| Dlx6as | 1.2257 | 0.020822 |
| Dmpk | -0.92949 | 0.029533 |
| Dmrt3 | -1.9552 | 8.00E-05 |
| Dnaaf3 | -1.1965 | 0.037436 |
| Dnah11 | -1.0882 | 0.036179 |
| Dnah5 | -1.5713 | 0.034011 |
| Dnajb9 | 0.77046 | 0.016254 |
| Dnajc15 | 0.73918 | 0.02179 |
| Dnajc16 | -1.2163 | 1.21E-04 |
| Dnajc19 | 1.0929 | 3.43E-04 |
| Dnajc24 | 0.93946 | 0.0089752 |
| Dnajc4 | 0.82335 | 0.020662 |
| Dnajc5 | -1.2118 | 3.70E-05 |
| Dnajc8 | 1.0213 | 6.46E-04 |
| Dnlz | 0.92476 | 0.0039662 |
| Dnm3 | -1.1497 | 1.89E-04 |
| Dnmbp | -0.96646 | 0.021151 |
| Dnmt3a | -2.0628 | 6.11E-13 |
| Dock5 | -1.9382 | 0.016149 |
| Dok6 | -1.6123 | 9.65E-06 |
| Dpcd | 1.1175 | 0.021361 |
| Dpf3 | -1.4588 | 0.015536 |
| Dph3 | 0.97544 | 0.020219 |
| Dpm2 | 0.81925 | 0.0089727 |
| Dpm3 | 1.2877 | 2.80E-04 |
| Dpy19l3 | -1.2155 | 0.002075 |
| Dpy30 | 0.9938 | 0.0012743 |
| Dpysl3 | -1.7004 | 1.03E-09 |
| Dpysl5 | -2.6053 | 2.30E-12 |
| Draxin | -1.9341 | 0.0030186 |
| Drd2 | -0.95628 | 0.032924 |
| Drp2 | -0.91324 | 0.011348 |
| Dstn | 0.9168 | 0.0014325 |
| Dtd2 | 0.75497 | 0.036125 |
| Dtx4 | -0.98864 | 0.0037912 |
| Dtymk | 0.85182 | 0.0098496 |
| Dusp18 | -1.2048 | 1.05E-04 |
| Dusp4 | -1.0202 | 0.0085271 |

| | | |
|---|---|---|
| Dvl1 | -0.64402 | 0.042578 |
| Dvl3 | -2.2672 | 1.90E-15 |
| Dynlrb1 | 1.2841 | 2.57E-04 |
| Dynlt1 | 0.82461 | 0.0099538 |
| Dynlt3 | 0.99359 | 4.57E-04 |
| Dyrk1b | -0.92049 | 0.0035513 |
| Dzank1 | -1.0503 | 0.020984 |
| E2f3 | -1.0567 | 0.010177 |
| E2f7 | -2.0008 | 0.025757 |
| E2f8 | -1.9029 | 0.013803 |
| Ebag9 | 0.88672 | 0.0040161 |
| Ebf4 | -0.93658 | 0.01543 |
| Ece1 | -0.68686 | 0.04504 |
| Echdc1 | 0.80172 | 0.0084998 |
| Eci1 | 0.78023 | 0.03242 |
| Eda | -1.0571 | 0.016278 |
| Eda2r | -2.1393 | 0.0018024 |
| Edc3 | -0.75258 | 0.045203 |
| Edem3 | -1.219 | 1.10E-04 |
| Eef1b2 | 1.3674 | 1.07E-06 |
| Eef1d | 0.87412 | 0.034026 |
| Eef1e1 | 0.99297 | 7.93E-04 |
| Eef1g | 1.0484 | 9.06E-04 |
| Eef2k | -0.89967 | 0.0029239 |
| Efna1 | 2.2399 | 0.023704 |
| Efr3b | -2.0421 | 1.20E-12 |
| Ei24 | 0.66507 | 0.035377 |
| Eif3e | 0.92051 | 0.0017482 |
| Eif3h | 0.98684 | 0.0010474 |
| Eif3i | 0.86361 | 0.0063819 |
| Eif3l | 0.76596 | 0.014884 |
| Eif3m | 0.94997 | 0.0014759 |
| Elavl3 | -1.1889 | 5.05E-05 |
| Elmsan1 | -1.6514 | 5.98E-07 |
| Elof1 | 1.0553 | 0.013472 |
| Emc2 | 0.72723 | 0.022414 |
| Emc6 | 0.91518 | 0.0037852 |
| Emc7 | 1.0288 | 4.17E-04 |
| Eme2 | -0.83944 | 0.010113 |
| Eml1 | -1.0548 | 0.0010297 |
| Eml3 | -1.0519 | 0.0067562 |
| Eml5 | -0.73028 | 0.030818 |
| Eml6 | -1.8665 | 2.39E-07 |
| Enah | -1.2764 | 3.08E-05 |

| | | |
|---|---|---|
| Enkur | 1.1192 | 4.76E-04 |
| Enpp2 | 1.288 | 0.0083499 |
| Epb4.1 | -1.3271 | 0.011983 |
| Epha10 | -1.4526 | 0.0021763 |
| Epha6 | -2.2271 | 1.39E-10 |
| Epha8 | -1.5204 | 3.84E-04 |
| Ephb2 | -0.73566 | 0.025905 |
| Epm2a | -1.4597 | 2.48E-05 |
| Epm2aip1 | -1.3501 | 0.0011712 |
| Eps8 | -0.82938 | 0.0109 |
| Erbb4 | -1.6283 | 0.0048879 |
| Erc1 | -1.784 | 1.49E-04 |
| Ercc2 | -1.1285 | 0.001361 |
| Ergic1 | -0.97802 | 0.0034868 |
| Erh | 0.82994 | 0.0059618 |
| Erich3 | -1.4021 | 8.99E-04 |
| Erlec1 | 0.95236 | 0.0011983 |
| Ern1 | -1.506 | 0.0031958 |
| Esco1 | 0.69978 | 0.036518 |
| Esd | 0.95947 | 0.0013112 |
| Espl1 | -1.5466 | 2.47E-04 |
| Esyt2 | -0.89231 | 0.0096963 |
| Etfa | 0.74032 | 0.01872 |
| Etl4 | -1.2899 | 1.27E-04 |
| Etv4 | -1.0204 | 0.0074465 |
| Etv6 | -1.1518 | 0.0099109 |
| Exoc8 | -0.96931 | 0.0030343 |
| F8 | -1.027 | 0.012001 |
| Faap100 | -1.0388 | 0.002888 |
| Fabp3 | 1.354 | 1.56E-04 |
| Fabp5 | 0.93974 | 0.0010816 |
| Fam102a | -1.5961 | 1.16E-05 |
| Fam102b | -0.89856 | 0.027556 |
| Fam103a1 | 0.80249 | 0.012453 |
| Fam104a | 0.85212 | 0.011319 |
| Fam124a | -1.0132 | 0.0041729 |
| Fam126a | -1.4951 | 6.69E-04 |
| Fam129b | -0.73274 | 0.032198 |
| Fam133b | 0.65372 | 0.045293 |
| Fam135b | -2.1956 | 5.22E-12 |
| Fam136a | 0.85583 | 0.036684 |
| Fam151b | 0.73866 | 0.039522 |
| Fam160b1 | -0.79834 | 0.040424 |
| Fam162a | 1.1065 | 1.50E-04 |

| | | |
|---|---|---|
| Fam163b | -1.1993 | 4.57E-05 |
| Fam168a | -1.1878 | 5.11E-05 |
| Fam168b | -1.6129 | 8.97E-07 |
| Fam171a1 | -0.75543 | 0.028723 |
| Fam174a | 0.93581 | 0.0019143 |
| Fam175b | 0.6776 | 0.042027 |
| Fam184b | -1.1158 | 0.0015463 |
| Fam185a | -0.91517 | 0.026181 |
| Fam189a1 | -1.3652 | 1.67E-05 |
| Fam189b | -0.79074 | 0.0097433 |
| Fam199x | -1.8721 | 1.07E-04 |
| Fam213a | 0.89965 | 0.0035913 |
| Fam219a | -1.0129 | 6.39E-04 |
| Fam222a | -1.3465 | 0.0043475 |
| Fam229b | 1.179 | 0.0037622 |
| Fam45a | 1.3111 | 5.91E-06 |
| Fam53c | -1.2794 | 3.56E-04 |
| Fam89a | -1.1942 | 0.0062608 |
| Fam8a1 | -0.7597 | 0.027091 |
| Fam92a1 | 0.81815 | 0.010047 |
| Fam96a | 1.3387 | 2.01E-05 |
| Fam96b | 1.218 | 0.0010995 |
| Fan1 | -1.061 | 0.0083677 |
| Fanca | -1.6623 | 0.0012596 |
| Fanci | -1.187 | 0.01025 |
| Farp1 | -0.83871 | 0.0040986 |
| Fars2 | 0.75049 | 0.025639 |
| Faxc | -1.5357 | 6.34E-06 |
| Fbn2 | -1.5665 | 0.046995 |
| Fbrs | -1.2162 | 1.42E-04 |
| Fbrsl1 | -0.87752 | 0.0041064 |
| Fbxl18 | -2.2045 | 1.20E-12 |
| Fbxl20 | -1.7045 | 4.23E-06 |
| Fbxl22 | 1.3953 | 0.034348 |
| Fbxl5 | 0.84549 | 0.0060714 |
| Fbxl7 | -1.9783 | 0.040931 |
| Fbxo10 | -0.81228 | 0.013363 |
| Fbxo16 | 1.3786 | 1.15E-04 |
| Fbxo42 | -1.3953 | 1.31E-05 |
| Fbxw2 | 0.79892 | 0.009382 |
| Fbxw7 | 0.76939 | 0.045942 |
| Fbxw8 | -0.69868 | 0.046286 |
| Fcf1 | 1.1707 | 1.68E-04 |
| Fcgrt | 1.3737 | 0.033673 |

| | | |
|---|---|---|
| Fchsd1 | -1.1921 | 8.06E-04 |
| Fem1a | -0.8447 | 0.0091361 |
| Fem1b | -1.0161 | 0.0010772 |
| Fgd1 | -1.4586 | 1.45E-05 |
| Fgd3 | -0.97552 | 0.019123 |
| Fgd5 | -1.8831 | 7.30E-05 |
| Fgfbp3 | 0.74003 | 0.040172 |
| Fgfr1 | -0.82759 | 0.0075864 |
| Fh | 0.72317 | 0.02813 |
| Fis1 | 1.2682 | 2.40E-04 |
| Fkbp2 | 1.1748 | 0.034007 |
| Flnb | -0.80368 | 0.013118 |
| Flrt1 | -1.5808 | 6.08E-06 |
| Flvcr1 | -1.1941 | 0.0054396 |
| Flywch1 | -0.8005 | 0.0081998 |
| Flywch2 | 1.0905 | 0.01816 |
| Fmc1 | 1.2395 | 1.93E-04 |
| Fmn1 | -2.9543 | 0.0050179 |
| Fmnl1 | -0.95873 | 0.0017794 |
| Fnbp1 | -1.1406 | 2.10E-04 |
| Fnbp1l | -0.87135 | 0.0093836 |
| Fndc3b | -2.0642 | 2.35E-09 |
| Fnip2 | -1.4283 | 5.98E-05 |
| Fosb | 1.5597 | 4.95E-04 |
| Foxk1 | -2.1115 | 8.90E-09 |
| Foxn3 | -1.4039 | 0.0019507 |
| Foxo1 | -1.6954 | 3.49E-06 |
| Foxp4 | -0.87577 | 0.0099284 |
| Foxred2 | -1.2316 | 0.0028844 |
| Fra10ac1 | 0.83215 | 0.01506 |
| Fras1 | -2.8623 | 1.41E-16 |
| Frem2 | -1.7482 | 0.0034889 |
| Frmd4a | -1.3941 | 1.12E-05 |
| Frmpd1 | -1.0575 | 0.0011699 |
| Frrs1l | -1.5305 | 1.65E-05 |
| Fryl | -1.2778 | 7.50E-05 |
| Fth1 | 1.2697 | 0.025538 |
| Ftl1 | 2.2437 | 0.023709 |
| Fuca2 | 0.93051 | 0.029092 |
| Fundc1 | 0.96308 | 0.0016016 |
| Furin | -0.6724 | 0.047338 |
| Fut9 | -1.3572 | 0.0013618 |
| Fxn | 0.79806 | 0.021556 |
| Fxyd1 | 1.6703 | 8.18E-08 |

| | | |
|---|---|---|
| Fzd1 | -0.74306 | 0.025103 |
| Fzd3 | -1.6579 | 2.53E-05 |
| Fzr1 | -0.73907 | 0.015485 |
| G6pc3 | 0.68389 | 0.045218 |
| Gab2 | -1.5531 | 5.78E-05 |
| Gabarap | 0.92309 | 0.0020246 |
| Gabarapl2 | 0.82747 | 0.0062253 |
| Gabrb1 | -2.1627 | 1.97E-12 |
| Gabrb2 | -1.183 | 0.032046 |
| Gadd45gip1 | 0.96589 | 0.0050705 |
| Galnt16 | -0.64074 | 0.038207 |
| Galnt7 | -0.83031 | 0.03223 |
| Gan | -3.1093 | 2.57E-12 |
| Gareml | -1.1984 | 5.26E-04 |
| Gatad2a | -0.73156 | 0.027032 |
| Gatad2b | -1.3178 | 1.93E-04 |
| Gatsl2 | -1.3929 | 2.22E-06 |
| Gcsh | 0.94776 | 0.0016112 |
| Gda | -1.0413 | 0.010163 |
| Gdf11 | -1.2136 | 0.010887 |
| Gemin6 | 0.87302 | 0.033656 |
| Gemin7 | 0.87281 | 0.020703 |
| Gfm2 | -0.86657 | 0.0085674 |
| Gfod1 | -1.6625 | 1.75E-05 |
| Gga1 | -0.71377 | 0.023241 |
| Ggct | 1.018 | 0.002563 |
| Ghrl | -1.8721 | 2.25E-05 |
| Gigyf1 | -1.8201 | 2.67E-07 |
| Ginm1 | 1.0184 | 0.0021043 |
| Gja1 | 0.95143 | 0.001217 |
| Glb1l | -1.0137 | 0.007776 |
| Glg1 | -0.96727 | 0.0015914 |
| Gli3 | -1.2136 | 0.015226 |
| Glipr1 | 1.4456 | 0.029275 |
| Glo1 | 0.94039 | 0.0015668 |
| Glod4 | 0.78828 | 0.013796 |
| Glrx | 0.82517 | 0.018458 |
| Glrx2 | 0.80189 | 0.0084998 |
| Glrx3 | 0.67202 | 0.031341 |
| Gltscr1 | -1.2265 | 1.15E-04 |
| Gm1043 | -2.0218 | 6.21E-04 |
| Gmfg | 1.4013 | 0.02848 |
| Gmppb | -0.79262 | 0.044067 |
| Gna11 | -1.179 | 8.33E-05 |

| | | |
|---|---|---|
| Gna13 | -1.692 | 0.0014792 |
| Gnaq | -1.1613 | 0.0050933 |
| Gnb3 | -2.2923 | 9.82E-05 |
| Gnb4 | -1.1515 | 0.0017493 |
| Gng10 | 1.2542 | 5.36E-06 |
| Gng3 | 1.008 | 0.0015116 |
| Gng5 | 1.3432 | 2.97E-06 |
| Gnl3l | -0.70893 | 0.04386 |
| Gnpnat1 | 0.81163 | 0.020325 |
| Gnptab | -0.98614 | 0.002022 |
| Golga7 | 0.66706 | 0.030737 |
| Gon4l | -1.0702 | 4.36E-04 |
| Gpatch2 | -1.4475 | 1.76E-04 |
| Gpd2 | -0.97896 | 0.023185 |
| Gpr153 | -0.91485 | 0.037216 |
| Gpr161 | -1.7204 | 5.21E-06 |
| Gpr26 | -1.8487 | 2.53E-08 |
| Gpr3 | 1.1889 | 0.041906 |
| Gprasp1 | 0.60946 | 0.043833 |
| Gprin3 | -1.799 | 2.52E-04 |
| Gpsm1 | -0.63123 | 0.046207 |
| Gpt | -1.1961 | 5.33E-04 |
| Gpx1 | 1.374 | 0.0026909 |
| Gpx4 | 1.1682 | 3.63E-05 |
| Gpx8 | 1.3568 | 9.45E-06 |
| Gramd4 | -1.0859 | 7.52E-04 |
| Grcc10 | 1.2625 | 0.01264 |
| Grin2d | -1.1305 | 0.0060491 |
| Grin3a | -1.4301 | 3.63E-06 |
| Grip2 | -1.1016 | 3.90E-04 |
| Grk6 | -0.88966 | 0.0034773 |
| Grm3 | -0.92374 | 0.012609 |
| Grpel1 | 0.83356 | 0.031561 |
| Gse1 | -2.2791 | 3.75E-14 |
| Gskip | 0.9368 | 0.0066492 |
| Gsta4 | 1.7329 | 0.0076166 |
| Gstk1 | 1.0332 | 0.0029831 |
| Gstm1 | 1.5639 | 1.75E-05 |
| Gstt1 | 1.7742 | 0.0032779 |
| Gtdc1 | 0.84546 | 0.012982 |
| Gtf2a1 | 0.83363 | 0.0060497 |
| Gtf2h5 | 1.1848 | 2.81E-05 |
| Gtf3c6 | 0.69685 | 0.032594 |
| Gtpbp1 | -0.8565 | 0.0076906 |

| | | |
|---|---|---|
| Gtse1 | -1.1663 | 0.0026833 |
| Gucy1a2 | -1.6533 | 2.81E-05 |
| Guk1 | 1.1211 | 0.0078545 |
| H2afz | 0.70548 | 0.034302 |
| H6pd | -0.89799 | 0.044595 |
| Hadh | 0.91491 | 0.034156 |
| Hcfc1r1 | 0.65415 | 0.049016 |
| Hcn4 | -1.7394 | 3.82E-07 |
| Hcrtr1 | -2.3045 | 0.010406 |
| Hddc3 | 1.122 | 2.31E-04 |
| Hdhd2 | 0.71463 | 0.022585 |
| Heatr4 | 1.2602 | 0.044615 |
| Hebp1 | 0.76297 | 0.018036 |
| Hectd4 | -2.0322 | 1.72E-13 |
| Hecw1 | -0.93115 | 0.047277 |
| Hecw2 | -1.2194 | 3.21E-04 |
| Heg1 | -1.8687 | 7.50E-07 |
| Helz | -1.9051 | 2.75E-09 |
| Hes5 | -1.4236 | 4.79E-05 |
| Hhip | -3.3933 | 8.26E-05 |
| Hhipl1 | -1.0695 | 0.0025974 |
| Hiat1 | 0.77254 | 0.023033 |
| Hid1 | -0.68328 | 0.031962 |
| Hif1an | -1.647 | 5.82E-07 |
| Hif3a | -2.8177 | 0.046834 |
| Higd1a | 0.66196 | 0.040016 |
| Higd2a | 0.78958 | 0.046658 |
| Hint2 | 0.71945 | 0.03986 |
| Hint3 | 0.81651 | 0.01702 |
| Hip1 | -0.8567 | 0.0068777 |
| Hjurp | -1.5059 | 0.0023946 |
| Hlcs | -1.3263 | 0.0056067 |
| Hmbox1 | -2.1675 | 7.77E-11 |
| Hmgn3 | 0.91404 | 0.0026909 |
| Hnrnpa0 | -1.1344 | 3.74E-05 |
| Hnrnpul2 | -0.81686 | 0.0076494 |
| Hrsp12 | 1.7393 | 9.16E-10 |
| Hs6st3 | -2.2719 | 4.38E-08 |
| Hsbp1 | 0.89931 | 0.0016691 |
| Hscb | 0.97252 | 0.0046353 |
| Hsd17b10 | 0.94028 | 0.0031211 |
| Hsd17b11 | 1.2534 | 3.07E-04 |
| Hsd17b12 | 0.65134 | 0.037497 |
| Hsdl2 | 0.79789 | 0.015356 |

| | | |
|---|---|---|
| Hsp90b1 | 0.91407 | 0.0017728 |
| Hspa8 | 1.9178 | 0.042255 |
| Hspa8 | 1.3607 | 4.12E-04 |
| Hspb11 | 0.82767 | 0.040287 |
| Hspd1 | 0.69635 | 0.021222 |
| Hspg2 | -1.8426 | 0.0071931 |
| Htr1a | -1.5139 | 0.023904 |
| Hunk | -1.9219 | 4.37E-04 |
| Hyal1 | -1.6656 | 0.01102 |
| Hypk | 1.5933 | 2.67E-07 |
| Iars2 | -0.99162 | 0.0062044 |
| Ica1 | 0.86468 | 0.008679 |
| Ice1 | -0.86211 | 0.0064628 |
| Icmt | -1.0542 | 0.0060816 |
| Ict1 | 0.87501 | 0.0095133 |
| Id2 | 1.2549 | 0.0011636 |
| Idua | -0.91655 | 0.013984 |
| Ift122 | -0.79433 | 0.016418 |
| Ift140 | -0.7544 | 0.018772 |
| Ift20 | 1.4317 | 4.72E-07 |
| Ift46 | 0.78738 | 0.028656 |
| Igbp1 | 0.97301 | 0.0031953 |
| Igdcc3 | -1.0856 | 0.032854 |
| Igdcc4 | -1.5164 | 0.0025403 |
| Igf1r | -1.5096 | 1.67E-05 |
| Igf2bp3 | -1.7768 | 0.03066 |
| Igf2r | -0.65325 | 0.049526 |
| Igfbp2 | -0.81165 | 0.012434 |
| Igip | 2.5503 | 4.28E-05 |
| Iglon5 | -0.76548 | 0.030273 |
| Igsf1 | 1.845 | 0.0068251 |
| Ikbkap | -0.81012 | 0.010873 |
| Ikbke | -1.3936 | 0.047282 |
| Ikzf4 | -1.288 | 0.0013064 |
| Il17rd | -3.1904 | 5.42E-08 |
| Ildr2 | -1.7866 | 4.37E-07 |
| Immp1l | 0.9812 | 0.0014984 |
| Impact | 0.69427 | 0.01981 |
| Ing2 | 0.70448 | 0.043538 |
| Ino80d | -2.8356 | 2.32E-11 |
| Inpp5j | -0.81714 | 0.01084 |
| Insr | -1.4333 | 3.65E-06 |
| Ints6 | -1.0073 | 0.032349 |
| Iqcb1 | 0.82055 | 0.011938 |

| | | |
|---|---|---|
| Iqcc | 1.0105 | 0.014865 |
| Iqce | -1.0879 | 9.98E-04 |
| Iqgap1 | -1.0631 | 0.0026372 |
| Iqgap3 | -1.8146 | 6.98E-04 |
| Iqub | 0.82934 | 0.031635 |
| Irf2bpl | -0.85026 | 0.0051328 |
| Irs1 | -1.4679 | 0.011403 |
| Isca1 | 0.6576 | 0.03359 |
| Isca2 | 0.67965 | 0.036695 |
| Iscu | 1.1409 | 0.012665 |
| Islr2 | -1.8231 | 3.02E-09 |
| Itga3 | -1.1216 | 5.02E-04 |
| Itga9 | -2.2523 | 1.20E-04 |
| Itgae | 1.0523 | 0.0099538 |
| Itgb3bp | 1.0436 | 0.012833 |
| Itgb8 | -1.4975 | 0.0087569 |
| Itgbl1 | -0.73048 | 0.049212 |
| Itm2b | 0.70683 | 0.021597 |
| Itpkb | -1.1052 | 0.0035018 |
| Jade2 | -0.98815 | 0.025211 |
| Jag2 | -1.1292 | 2.86E-04 |
| Jagn1 | 0.80413 | 0.010961 |
| Jak3 | -1.0439 | 0.01183 |
| Jarid2 | -0.87598 | 0.0087286 |
| Jmjd7 | -1.8398 | 0.0049941 |
| Josd2 | 0.80211 | 0.038684 |
| Jph1 | -1.1688 | 0.012434 |
| Jtb | 0.67812 | 0.036135 |
| Junb | 1.0861 | 2.69E-04 |
| Jup | -0.73931 | 0.018036 |
| Kat2a | -0.68021 | 0.03078 |
| Kat6a | -1.6038 | 5.50E-08 |
| Kazn | -1.6199 | 9.00E-08 |
| Kbtbd11 | -2.1003 | 3.87E-08 |
| Kbtbd7 | -1.8682 | 8.84E-05 |
| Kcna1 | -2.2613 | 7.20E-11 |
| Kcnab2 | -1.4327 | 9.58E-07 |
| Kcnd3 | -1.9119 | 9.23E-10 |
| Kcnf1 | -0.93764 | 0.0046551 |
| Kcng3 | -2.8494 | 7.40E-09 |
| Kcnh2 | -0.76965 | 0.026049 |
| Kcnj10 | -1.7979 | 8.18E-06 |
| Kcnj4 | -0.74594 | 0.017704 |
| Kcnj6 | -1.8547 | 1.78E-07 |

| | | |
|---|---|---|
| Kcnk10 | -1.1061 | 0.0026645 |
| Kcnk12 | -1.2359 | 1.73E-04 |
| Kcnk9 | -1.9696 | 3.81E-10 |
| Kcnn2 | 0.92473 | 0.0058396 |
| Kcnn3 | -1.3195 | 1.27E-04 |
| Kcnq5 | -0.84603 | 0.01952 |
| Kcns3 | 1.259 | 9.30E-04 |
| Kcnv1 | 0.89015 | 0.009352 |
| Kctd20 | -1.3241 | 7.48E-05 |
| Kctd3 | -0.704 | 0.027604 |
| Kdelr1 | 0.6499 | 0.047438 |
| Kdelr2 | 0.7199 | 0.02405 |
| Khnyn | -1.2012 | 0.0062038 |
| Khsrp | -0.9861 | 0.0014798 |
| Kif13a | -1.2478 | 2.10E-04 |
| Kif13b | -1.3094 | 3.08E-04 |
| Kif14 | -1.439 | 0.049719 |
| Kif18b | -1.5946 | 7.83E-04 |
| Kif26b | -1.7867 | 3.31E-05 |
| Kif28p | 1.8798 | 0.016717 |
| Kif2c | -1.4344 | 0.022043 |
| Kif3a | -0.68656 | 0.038887 |
| Kifc1 | -1.2888 | 6.98E-04 |
| Kirrel | -1.6011 | 1.43E-04 |
| Kit | -1.0143 | 9.65E-04 |
| Klf12 | -1.9839 | 1.11E-06 |
| Klf13 | -2.2185 | 1.98E-10 |
| Klf7 | -1.8477 | 3.33E-04 |
| Klf7 | -1.548 | 4.92E-04 |
| Klf8 | -2.174 | 0.0017104 |
| Klhdc10 | -1.7538 | 2.48E-07 |
| Klhl11 | -1.349 | 0.0019725 |
| Klhl17 | -1.2757 | 2.60E-04 |
| Klhl18 | -0.94845 | 0.0079927 |
| Klhl3 | -1.6242 | 2.43E-05 |
| Kmt2a | -2.5629 | 2.50E-19 |
| Kmt2c | -1.0794 | 3.99E-04 |
| Kmt2e | -0.67973 | 0.034302 |
| Kntc1 | -1.4179 | 3.13E-04 |
| Kpna5 | -0.8765 | 0.048291 |
| Kpna6 | -0.84706 | 0.012307 |
| Kpnb1 | -0.93563 | 0.0019902 |
| Krt10 | 0.84366 | 0.042578 |
| Krtcap2 | 1.0064 | 8.50E-04 |

| | | |
|---|---|---|
| Ksr1 | -2.619 | 6.39E-14 |
| Ksr2 | -2.7465 | 1.05E-11 |
| L3mbtl1 | -2.473 | 0.00283 |
| L3mbtl3 | -0.82276 | 0.033998 |
| l7Rn6 | 0.75268 | 0.015944 |
| Lamp2 | 0.75446 | 0.04121 |
| Lamp5 | 1.5326 | 3.72E-04 |
| Lamtor2 | 1.2593 | 9.24E-06 |
| Lamtor3 | 0.77985 | 0.014346 |
| Lamtor4 | 1.2945 | 4.23E-04 |
| Lamtor5 | 1.0688 | 3.19E-04 |
| Lancl3 | -1.9254 | 1.76E-07 |
| Laptm4a | 0.81687 | 0.005445 |
| Larp1 | -1.3457 | 3.36E-06 |
| Lats1 | -0.96032 | 0.047385 |
| Lbhd1 | 1.127 | 0.0013935 |
| Lcmt1 | 0.88165 | 0.0042354 |
| Lcor | -1.6222 | 2.24E-04 |
| Ldhb | 0.82981 | 0.033441 |
| Ldoc1l | -1.4867 | 3.93E-06 |
| Leng8 | -1.4113 | 0.020703 |
| Leprot | 0.72043 | 0.022259 |
| Letm1 | -0.94056 | 0.002386 |
| Lgals1 | 1.6494 | 1.74E-04 |
| Lhfp | 0.80954 | 0.020808 |
| Lhfpl5 | 1.4451 | 0.047954 |
| Lhx4 | -1.7169 | 0.021835 |
| Lifr | -2.0282 | 1.30E-09 |
| Limch1 | -1.395 | 9.21E-05 |
| Limd1 | -1.1368 | 0.0059413 |
| Lime1 | -0.9676 | 0.010259 |
| Limk1 | -1.121 | 1.76E-04 |
| Lingo3 | -1.1277 | 0.0021113 |
| Lipg | -2.4846 | 0.0329 |
| Lipt2 | 0.87966 | 0.031963 |
| Lman2l | -0.90409 | 0.018186 |
| Lmbrd1 | 0.85032 | 0.003904 |
| Lmnb1 | -1.0719 | 0.0014649 |
| LOC100174910 | 1.0436 | 0.0024798 |
| LOC100233176 | 1.1825 | 1.64E-04 |
| LOC100359687 | 1.8199 | 4.52E-05 |
| LOC100359922 | 1.1768 | 0.0046034 |
| LOC100360056 | 1.3135 | 2.84E-06 |
| LOC100360057 | 1.3016 | 2.72E-05 |

| | | |
|---|---|---|
| LOC100360087 | 1.5062 | 0.0011003 |
| LOC100360218 | 0.92529 | 0.013307 |
| LOC100360522 | 1.8208 | 1.35E-04 |
| LOC100360573 | 1.6299 | 6.98E-04 |
| LOC100360647 | 1.6476 | 0.0028411 |
| LOC100360726 | -1.6192 | 0.028654 |
| LOC100360791 | 2.855 | 0.025146 |
| LOC100361240 | 1.9858 | 0.038282 |
| LOC100361543 | 0.75227 | 0.016876 |
| LOC100361993 | 2.2457 | 1.49E-04 |
| LOC100362027 | 1.5954 | 3.08E-09 |
| LOC100362384 | 2.2649 | 0.039781 |
| LOC100362814 | -2.0765 | 6.11E-13 |
| LOC100362980 | 1.2934 | 6.84E-05 |
| LOC100363675 | -1.059 | 0.0099142 |
| LOC100364191 | 1.731 | 0.0080769 |
| LOC100909840 | 0.76427 | 0.0098835 |
| LOC100909954 | -1.8109 | 1.11E-08 |
| LOC100910046 | -1.3479 | 0.020646 |
| LOC100910137 | 0.6909 | 0.044621 |
| LOC100910370 | 0.97157 | 0.0012637 |
| LOC100910427 | 0.76297 | 0.03986 |
| LOC100910508 | -0.77722 | 0.042296 |
| LOC100910577 | 0.8075 | 0.036483 |
| LOC100910732 | -0.9959 | 0.03401 |
| LOC100911293 | 1.1346 | 0.0025113 |
| LOC100911372 | 1.3019 | 2.87E-04 |
| LOC100911483 | 1.4938 | 1.87E-05 |
| LOC100911664 | 0.68851 | 0.031864 |
| LOC100911685 | -1.2454 | 7.31E-04 |
| LOC100911840 | 0.8464 | 0.0065302 |
| LOC100911956 | 1.1639 | 0.0068375 |
| LOC100912042 | 1.0725 | 0.026837 |
| LOC100912163 | 0.81845 | 0.020748 |
| LOC100912182 | 1.8163 | 3.60E-11 |
| LOC100912228 | 1.9537 | 9.32E-05 |
| LOC100912445 | -2.5515 | 8.10E-05 |
| LOC100912458 | 0.80155 | 0.041242 |
| LOC100912481 | -1.1285 | 0.0020994 |
| LOC102550385 | 0.85665 | 0.022832 |
| LOC102550391 | 0.87497 | 0.019214 |
| LOC102552166 | -2.1719 | 0.0053969 |
| LOC102552640 | -0.76688 | 0.023904 |
| LOC102554232 | 0.7767 | 0.016113 |

| | | |
|---|---|---|
| LOC102555377 | 1.0347 | 0.0034764 |
| LOC102555672 | 0.81529 | 0.010764 |
| LOC103689975 | -3.1954 | 4.14E-08 |
| LOC103689983 | 0.91929 | 0.013118 |
| LOC103690024 | -1.1315 | 0.0031267 |
| LOC103690050 | -1.719 | 0.039923 |
| LOC103690171 | 1.0874 | 0.0098688 |
| LOC103693210 | 0.93306 | 0.0030793 |
| LOC103693430 | 1.4788 | 3.93E-08 |
| LOC103693776 | -0.85085 | 0.0033311 |
| LOC103694877 | 1.1962 | 0.0017627 |
| LOC257650 | 1.3925 | 0.0028294 |
| LOC288913 | 0.88391 | 0.0039894 |
| LOC301124 | 1.1338 | 7.02E-05 |
| LOC361985 | 1.1021 | 1.37E-04 |
| LOC498555 | 1.4115 | 0.037396 |
| LOC499781 | -1.0173 | 0.023724 |
| LOC501110 | 1.0329 | 0.0065838 |
| LOC679894 | -1.0223 | 0.025958 |
| LOC680227 | -7.5364 | 1.87E-04 |
| LOC683593 | -1.8837 | 1.70E-08 |
| LOC683897 | 0.86327 | 0.01509 |
| LOC687679 | 1.4597 | 5.42E-07 |
| LOC689499 | -2.1491 | 2.11E-12 |
| LOC689574 | 0.77091 | 0.02319 |
| LOC689991 | -1.5931 | 6.06E-08 |
| LOC690871 | 1.4986 | 1.59E-05 |
| LOC691427 | 1.814 | 0.031341 |
| LOC691807 | 1.0754 | 1.87E-04 |
| LOC691921 | 1.0569 | 0.016173 |
| Lonrf2 | -2.3935 | 1.73E-16 |
| Lpar4 | 1.11 | 0.005919 |
| Lpp | -3.0875 | 3.56E-05 |
| Lppr2 | -1.1703 | 3.09E-05 |
| Lrba | -1.286 | 0.0025342 |
| Lrch3 | -1.2981 | 0.0079927 |
| Lrch4 | -1.0394 | 0.0061068 |
| Lrif1 | 0.75115 | 0.030349 |
| Lrp1b | -1.0651 | 0.049895 |
| Lrp5 | -1.1984 | 2.81E-04 |
| Lrp6 | -1.3706 | 1.62E-05 |
| Lrrc16b | -1.4617 | 7.95E-08 |
| Lrrc26 | -2.8707 | 7.39E-05 |
| Lrrc3b | 1.3927 | 0.02963 |

| | | |
|---|---|---|
| Lrrc40 | 0.81938 | 0.0094812 |
| Lrrc57 | 0.91616 | 0.008377 |
| Lrrc58 | -1.4949 | 0.0076047 |
| Lrrc75b | -1.4275 | 0.0011641 |
| Lrrc8c | -1.1871 | 0.0025064 |
| Lsamp | -0.99289 | 0.0011558 |
| Lsm1 | 0.89902 | 0.0070108 |
| Lsm11 | -1.4912 | 5.97E-07 |
| Lsm12 | 0.6651 | 0.038766 |
| Lsm3 | 1.2156 | 5.53E-05 |
| Lsm5 | 1.5854 | 2.01E-07 |
| Lsm6 | 1.0849 | 4.35E-04 |
| Lsm7 | 1.347 | 8.97E-05 |
| Ltbp3 | -0.78945 | 0.015647 |
| Ltbp4 | -1.606 | 7.35E-08 |
| Lxn | 0.79668 | 0.010781 |
| Lypd5 | -2.0698 | 0.010624 |
| Lyrm1 | 0.81733 | 0.029179 |
| Lyrm2 | 1.0015 | 0.0014477 |
| Lyrm5 | 1.3384 | 1.33E-04 |
| Lyrm9 | 0.78585 | 0.023762 |
| Lysmd2 | 0.70982 | 0.036526 |
| Lyz2 | 1.0522 | 0.0049884 |
| Lztfl1 | 1.0776 | 7.22E-04 |
| Lzts3 | -0.91273 | 0.0029359 |
| Mafg | -0.959 | 0.0031108 |
| Magi1 | -1.1272 | 3.64E-04 |
| Magoh | 0.79586 | 0.020623 |
| Magohb | 1.0744 | 0.0084114 |
| Mal2 | 0.75168 | 0.048516 |
| Malsu1 | 0.84497 | 0.014969 |
| Mamdc4 | -1.0868 | 0.030056 |
| Man1a2 | -1.0134 | 0.0184 |
| Man2c1 | -0.76913 | 0.013096 |
| Manbal | 1.0193 | 0.043391 |
| Manf | 0.87891 | 0.0040925 |
| Map2k7 | -1.0345 | 0.0019515 |
| Map3k13 | -1.0772 | 0.0052837 |
| Map3k2 | -1.8065 | 4.32E-05 |
| Map3k9 | -2.1712 | 4.05E-14 |
| Map4k2 | -1.2085 | 1.96E-04 |
| Map6d1 | -0.77955 | 0.044357 |
| Map7 | -1.0213 | 0.0013756 |
| Mapk1ip1 | 1.1412 | 0.020352 |

| | | |
|---|---|---|
| Mapk8ip2 | -1.7872 | 4.03E-11 |
| Mapt | -1.6598 | 2.35E-09 |
| Mark2 | -1.0652 | 3.86E-04 |
| Mark4 | -0.89929 | 0.0018269 |
| MAST1 | -0.61763 | 0.044357 |
| Matr3 | 0.82546 | 0.0040162 |
| Mau2 | -0.68385 | 0.034284 |
| Mbd1 | -0.81913 | 0.0095111 |
| Mbd6 | -1.9406 | 4.50E-06 |
| Mbip | 0.82947 | 0.032733 |
| Mbnl2 | -1.0355 | 0.0010349 |
| Mcee | 0.90428 | 0.0076973 |
| Mcf2l | -1.0481 | 3.29E-04 |
| Mcm10 | -1.5873 | 0.001792 |
| Mcm3 | -1.0014 | 0.022191 |
| Mcm3ap | -0.6472 | 0.049506 |
| Mcm5 | -1.4464 | 1.97E-04 |
| Mcts1 | 0.93366 | 0.0026286 |
| Mcur1 | -1.3102 | 0.0011292 |
| Mdc1 | -1.8684 | 2.89E-07 |
| Mdga1 | -2.1966 | 1.33E-07 |
| Mdga2 | -0.94881 | 0.0073455 |
| Mdk | 1.131 | 0.0069908 |
| Mdn1 | -0.82213 | 0.010112 |
| Mea1 | 0.92393 | 0.034467 |
| Med11 | 1.002 | 0.0077285 |
| Med12 | -1.2835 | 0.0011131 |
| Med12l | -2.3011 | 8.27E-11 |
| Med19 | 1.0631 | 0.003556 |
| Med21 | 0.71759 | 0.049402 |
| Med25 | -0.82293 | 0.0047648 |
| Med28 | 1.419 | 0.0016361 |
| Med30 | 0.94917 | 0.0038613 |
| Med31 | 1.0495 | 0.0016115 |
| Med4 | 0.95119 | 0.0028215 |
| Med6 | 0.99613 | 0.0034542 |
| Med7 | 0.77998 | 0.036097 |
| Med8 | 0.77249 | 0.048029 |
| Megf11 | -0.86429 | 0.0061995 |
| Megf9 | -0.95738 | 0.038214 |
| Mettl23 | 0.80286 | 0.046606 |
| Mettl5 | 0.90058 | 0.0072201 |
| Mettl9 | 1.0902 | 2.16E-04 |
| Mex3d | -0.95127 | 0.0035959 |

| | | |
|---|---|---|
| Mff | 0.8239 | 0.0065295 |
| Mfsd12 | -1.3447 | 1.71E-05 |
| Mga | -0.88625 | 0.03596 |
| Mgam | -1.7127 | 3.94E-04 |
| Mgat3 | -0.89645 | 0.0039662 |
| Mgat5 | -1.8777 | 6.55E-04 |
| MGC94335 | 1.032 | 0.0016675 |
| Mgrn1 | -0.77359 | 0.012453 |
| Mgst3 | 0.74936 | 0.019166 |
| Micall1 | -2.0103 | 7.54E-09 |
| Mid1 | -1.659 | 7.84E-04 |
| Midn | -1.4278 | 8.01E-07 |
| Mief1 | -1.0047 | 0.005048 |
| Mien1 | 0.95849 | 0.0015102 |
| Mier2 | -0.66004 | 0.048317 |
| Minos1 | 0.95482 | 0.0010581 |
| Mir132 | 2.0169 | 0.039378 |
| Mir186 | -1.372 | 0.01363 |
| Mir3597-1 | -1.8274 | 0.014895 |
| Mir6333 | -2.6008 | 6.33E-04 |
| Mki67 | -1.3313 | 0.019312 |
| Mkks | 0.7979 | 0.010239 |
| Mkl1 | -1.1348 | 2.27E-04 |
| Mknk1 | -0.75673 | 0.043626 |
| Mknk2 | -0.86617 | 0.0050752 |
| Mlc1 | 0.77085 | 0.010975 |
| Mllt4 | -1.4074 | 1.60E-06 |
| Mlst8 | -0.69047 | 0.048222 |
| Mlxip | -2.5336 | 1.87E-13 |
| Mmab | -1.2664 | 0.013891 |
| Mmp15 | -1.0919 | 2.23E-04 |
| Mmp16 | -1.0832 | 5.20E-04 |
| Mn1 | -1.3843 | 3.99E-06 |
| Mnt | -0.69146 | 0.02828 |
| Mob3b | -1.3649 | 0.0061708 |
| Mogat1 | 1.2901 | 0.038609 |
| Mon1b | -0.8543 | 0.015647 |
| Mov10 | -0.91093 | 0.039108 |
| Mpc1 | 0.91397 | 0.0014939 |
| Mpc2 | 1.0537 | 5.87E-04 |
| Mpdu1 | 0.91155 | 0.0052075 |
| Mphosph6 | 1.0862 | 4.26E-04 |
| Mpp3 | -0.76073 | 0.014631 |
| Mras | -0.66711 | 0.03237 |

| | | |
|---|---|---|
| Mroh1 | -2.3213 | 1.14E-08 |
| Mroh2a | -1.9489 | 0.038317 |
| Mroh6 | -3.1591 | 0.034929 |
| Mrp63 | 0.91159 | 0.0042354 |
| Mrpl12 | 0.84124 | 0.028141 |
| Mrpl13 | 1.412 | 3.01E-06 |
| Mrpl15 | 0.68989 | 0.041153 |
| Mrpl16 | 0.79433 | 0.020615 |
| Mrpl18 | 1.3649 | 4.00E-06 |
| Mrpl20 | 0.90526 | 0.0026025 |
| Mrpl21 | 1.0796 | 0.0011683 |
| Mrpl22 | 1.1062 | 8.90E-04 |
| Mrpl23 | 0.9582 | 0.0088716 |
| Mrpl27 | 1.1846 | 1.17E-04 |
| Mrpl32 | 0.82157 | 0.017906 |
| Mrpl33 | 1.057 | 2.93E-04 |
| Mrpl34 | 1.3016 | 0.026218 |
| Mrpl35 | 0.94101 | 0.00283 |
| Mrpl36 | 0.8089 | 0.01557 |
| Mrpl39 | 0.8854 | 0.0096549 |
| Mrpl42 | 1.4142 | 1.38E-06 |
| Mrpl43 | 0.71358 | 0.028515 |
| Mrpl46 | 0.83541 | 0.011169 |
| Mrpl48 | 1.1774 | 7.97E-05 |
| Mrpl49 | 0.6739 | 0.035047 |
| Mrpl52 | 1.4104 | 5.00E-05 |
| Mrpl54 | 1.2429 | 7.09E-05 |
| Mrps14 | 0.86758 | 0.004788 |
| Mrps15 | 1.0537 | 6.64E-04 |
| Mrps16 | 0.72735 | 0.032837 |
| Mrps17 | 1.0467 | 5.61E-04 |
| Mrps18a | 0.93673 | 0.042369 |
| Mrps18c | 1.3206 | 1.85E-05 |
| Mrps21 | 1.3985 | 1.10E-06 |
| Mrps24 | 1.0039 | 0.0093868 |
| Mrps25 | 0.97096 | 0.017568 |
| Mrps33 | 1.0943 | 2.93E-04 |
| Mrps35 | 1.062 | 0.0045425 |
| Mrps7 | 1.1177 | 0.0015494 |
| Mrps9 | 0.94888 | 0.0029046 |
| Msantd2 | 0.95542 | 0.0035469 |
| Msi1 | -1.0244 | 0.0033109 |
| Msi2 | -1.1313 | 0.010742 |
| Msmp | -1.3259 | 4.92E-04 |

| | | |
|---|---|---|
| Mt-atp6 | 2.7032 | 1.20E-06 |
| Mt-atp8 | 2.9889 | 6.46E-25 |
| Mt-co1 | 1.812 | 3.91E-09 |
| Mt-co2 | 2.5584 | 3.06E-22 |
| Mt-cox3 | 2.2962 | 3.77E-09 |
| Mt-cytb | 1.9562 | 5.09E-09 |
| Mt-nd1 | 2.4357 | 8.21E-11 |
| Mt-nd2 | 2.5734 | 8.11E-07 |
| Mt-nd3 | 1.9769 | 0.022231 |
| Mt-nd4 | 2.4553 | 4.27E-13 |
| Mt-nd4l | 2.2313 | 0.021098 |
| Mt-nd5 | 1.5848 | 0.0017104 |
| Mt-nd6 | 2.5952 | 2.41E-22 |
| Mt2A | 1.7723 | 0.001361 |
| Mt3 | 1.8449 | 9.47E-12 |
| Mtcl1 | -1.0533 | 0.0029667 |
| Mterf1 | 1.1464 | 0.0022148 |
| Mterf4 | 0.81894 | 0.028459 |
| Mtg1 | 0.75734 | 0.030851 |
| Mthfr | -1.1273 | 0.023662 |
| Mtif2 | 0.74167 | 0.043673 |
| Mtmr9 | -1.3826 | 3.86E-05 |
| Mtr | -1.0908 | 0.036605 |
| Mtrf1 | 0.98579 | 0.012377 |
| Mtus2 | -0.90725 | 0.03107 |
| Mtx3 | -1.5796 | 1.91E-04 |
| Mustn1 | 1.6951 | 0.023762 |
| Mvb12b | -0.62321 | 0.04504 |
| Mxi1 | 1.2391 | 6.84E-05 |
| Mybbp1a | -0.73477 | 0.020499 |
| Mybl2 | -1.4022 | 0.0090257 |
| Mycbp | 0.86682 | 0.037891 |
| Mydgf | 0.70439 | 0.029843 |
| Myeov2 | 1.4979 | 5.83E-08 |
| Myh6 | -1.7656 | 6.54E-04 |
| Myo5b | -0.76242 | 0.01981 |
| Myo9a | -1.2689 | 8.36E-04 |
| Myo9b | -1.5809 | 6.83E-05 |
| Myrip | -0.81842 | 0.013988 |
| N4bp1 | -1.1657 | 6.82E-04 |
| N4bp2 | -1.2134 | 0.035377 |
| N6amt1 | 0.82957 | 0.0198 |
| N6amt2 | 1.1128 | 0.0041449 |
| Naa20 | 0.77578 | 0.012947 |

| | | |
|---|---|---|
| Naa38 | 1.6472 | 2.66E-04 |
| Naca | 0.89064 | 0.0028787 |
| Nacc1 | -1.3013 | 1.23E-05 |
| Nbeal2 | -2.0797 | 1.22E-04 |
| Ncam2 | -1.0221 | 0.0076973 |
| Ncaph | -1.1703 | 0.009643 |
| Ncln | -0.78236 | 0.0082798 |
| Ncoa7 | -0.85727 | 0.0091482 |
| Ndp | 1.0411 | 0.0015528 |
| Ndrg3 | -0.85026 | 0.022047 |
| Ndst4 | -2.939 | 1.24E-04 |
| Ndufa1 | 1.261 | 5.81E-06 |
| Ndufa11 | 1.2446 | 1.20E-05 |
| Ndufa12 | 0.96678 | 0.0065191 |
| Ndufa2 | 1.4269 | 7.54E-06 |
| Ndufa4 | 1.4029 | 8.97E-07 |
| Ndufa5 | 0.8942 | 0.0021604 |
| Ndufa6 | 1.4851 | 1.26E-06 |
| Ndufa7 | 1.3021 | 4.87E-06 |
| Ndufa8 | 1.1542 | 0.0025965 |
| Ndufab1 | 0.80489 | 0.0072411 |
| Ndufaf3 | 0.73524 | 0.027628 |
| Ndufaf5 | 1.1491 | 0.0058733 |
| Ndufb10 | 1.3714 | 8.64E-05 |
| Ndufb11 | 0.98589 | 8.32E-04 |
| Ndufb2 | 1.148 | 0.0014 |
| Ndufb3 | 1.1403 | 7.67E-05 |
| Ndufb4 | 1.1084 | 1.11E-04 |
| Ndufb5 | 1.6671 | 1.73E-06 |
| Ndufb6 | 0.93034 | 0.0015305 |
| Ndufb7 | 1.1682 | 0.022699 |
| Ndufb8 | 0.95281 | 0.0011227 |
| Ndufb9 | 1.2573 | 6.67E-06 |
| Ndufc1 | 1.417 | 5.44E-07 |
| Ndufc2 | 1.1319 | 5.97E-05 |
| Ndufs3 | 0.84234 | 0.024193 |
| Ndufs4 | 0.77993 | 0.0097964 |
| Ndufs5 | 1.2715 | 5.24E-04 |
| Ndufs8 | 0.79736 | 0.010943 |
| Ndufv2 | 0.9418 | 0.0011983 |
| Ndufv3-ps1 | 1.169 | 0.0084998 |
| Nebl | -0.79525 | 0.034903 |
| Nedd4l | -0.63908 | 0.049001 |
| Nedd8 | 1.1258 | 6.82E-05 |

| | | |
|---|---|---|
| Nek2l1 | -1.6335 | 0.01375 |
| Nenf | 1.8497 | 1.04E-07 |
| Neo1 | -0.93106 | 0.0019143 |
| Neto1 | -0.99277 | 0.0013386 |
| Neurl1b | -2.0702 | 5.42E-06 |
| NEWGENE_1310139 | -3.3323 | 2.39E-05 |
| Nfasc | -1.08 | 1.97E-04 |
| Nfatc1 | -1.2382 | 0.011878 |
| Nfia | -1.4234 | 1.30E-05 |
| Nfib | -1.9512 | 1.25E-05 |
| Nfu1 | 0.75705 | 0.023153 |
| Nfyc | 0.69421 | 0.038456 |
| Ngdn | 0.7839 | 0.02804 |
| Nhlrc2 | -1.1334 | 0.0020737 |
| Nhp2 | 0.89475 | 0.0085444 |
| Nhsl2 | -2.1378 | 1.87E-09 |
| Nid1 | -1.6954 | 0.0072583 |
| Nip7 | 0.68908 | 0.049966 |
| Nipal2 | 2.0299 | 4.65E-04 |
| Nipal3 | -1.8004 | 1.57E-08 |
| Nlrc5 | -1.61 | 0.037566 |
| Nme1 | 0.6846 | 0.03198 |
| Nme2 | 0.68581 | 0.031663 |
| Nme3 | 1.3042 | 0.016246 |
| Nmnat2 | -1.3083 | 1.58E-04 |
| Nmrk1 | 0.81984 | 0.047431 |
| Nol4l | -2.0869 | 1.63E-09 |
| Nol7 | 0.72177 | 0.025639 |
| Nop10 | 1.3317 | 0.001144 |
| Notch1 | -1.5057 | 0.0025974 |
| Notch2 | -1.8464 | 8.28E-06 |
| Notch3 | -2.3846 | 1.89E-12 |
| Nova2 | -1.4044 | 1.36E-06 |
| Npap60 | -0.73826 | 0.026527 |
| Npas4 | 2.958 | 1.87E-05 |
| Npc2 | 0.7239 | 0.023789 |
| Nphp4 | -1.1433 | 7.23E-04 |
| Npr3 | -1.0677 | 0.0090931 |
| Nptx1 | -2.6524 | 1.80E-10 |
| Nr2c2 | -1.4925 | 1.36E-05 |
| Nr3c2 | -0.94138 | 0.03986 |
| Nr4a1 | 1.0563 | 0.01557 |
| Nrarp | -1.4148 | 4.05E-05 |
| Nrbp2 | -1.4808 | 5.14E-08 |

| | | |
|---|---|---|
| Nrn1 | 1.1226 | 0.0044842 |
| Nrp1 | -1.1082 | 6.72E-04 |
| Nrp2 | -1.7641 | 2.01E-05 |
| Nrsn1 | 1.9537 | 0.037536 |
| Nsa2 | 0.87281 | 0.0034666 |
| Nsmce1 | 0.80328 | 0.029661 |
| Nsrp1 | 0.91237 | 0.0077467 |
| Nt5c1a | -1.4608 | 0.025676 |
| Nt5c3a | 0.82915 | 0.0072302 |
| Nt5dc3 | -1.8453 | 2.60E-10 |
| Ntn1 | -1.5181 | 8.69E-04 |
| Ntn3 | -1.4951 | 6.52E-04 |
| Ntpcr | 0.90559 | 0.0060587 |
| Ntsr2 | 1.191 | 0.040552 |
| Nuak1 | -1.1299 | 8.24E-04 |
| Nucb2 | 0.89792 | 0.007898 |
| Nudt2 | 1.0887 | 3.62E-04 |
| Nudt6 | 1.0585 | 0.0040783 |
| Numa1 | -0.81902 | 0.0067509 |
| Nup188 | -0.83433 | 0.012643 |
| Nup210 | -0.81708 | 0.012901 |
| Nupr1l | 1.3316 | 0.0070099 |
| Nutf2 | 0.83686 | 0.0054514 |
| Nxt2 | 0.94097 | 0.026527 |
| Nyap2 | -1.4662 | 0.0071207 |
| Nynrin | -1.3422 | 9.51E-05 |
| Ociad1 | 0.79288 | 0.0085762 |
| Odf3 | -2.6113 | 7.66E-04 |
| Ogfod1 | -1.1641 | 5.35E-04 |
| Olig1 | -0.97386 | 8.47E-04 |
| Onecut1 | -2.2039 | 0.011579 |
| Opcml | -1.4773 | 2.85E-06 |
| Ormdl2 | 0.97332 | 0.0028407 |
| Osbpl6 | -0.94329 | 0.0037212 |
| Ost4 | 1.4582 | 1.08E-07 |
| Ostc | 0.97284 | 9.61E-04 |
| Otof | -3.194 | 9.18E-22 |
| Otud4 | -1.0839 | 0.0037929 |
| P3h3 | -1.4399 | 4.65E-04 |
| Pabpc1 | 0.73596 | 0.01221 |
| Padi2 | -1.1424 | 0.026218 |
| Paics | 0.70238 | 0.030384 |
| Paip2 | 0.8421 | 0.0036752 |
| Pak3 | -1.6431 | 5.34E-05 |

| | | |
|---|---|---|
| Pak4 | -0.887 | 0.024083 |
| Palm2 | -1.3415 | 0.0028742 |
| Pam16 | 1.2829 | 1.86E-04 |
| Pan2 | -1.0829 | 0.0019143 |
| Panx1 | -0.72827 | 0.027687 |
| Paqr8 | -1.3636 | 3.75E-04 |
| Paqr9 | -1.6645 | 4.66E-05 |
| Pard3 | -0.83315 | 0.021009 |
| Parm1 | -1.1327 | 0.0036257 |
| Pbdc1 | 1.0224 | 0.0014696 |
| Pbrm1 | -1.0359 | 0.014935 |
| Pbx1 | -1.2012 | 5.41E-05 |
| Pcbd2 | 1.3818 | 4.67E-05 |
| Pcdh15 | -1.909 | 5.87E-04 |
| Pcdh17 | -1.5513 | 7.71E-08 |
| Pcdh19 | -1.4257 | 2.65E-04 |
| Pcdha13 | -1.3156 | 4.44E-04 |
| Pcdha2 | -1.576 | 4.57E-04 |
| Pcdha3 | -1.3254 | 5.54E-04 |
| Pcdhb2 | -2.1457 | 4.93E-04 |
| Pcdhga1 | -1.1739 | 0.045934 |
| Pcdhga10 | -1.3256 | 0.0062716 |
| Pcdhga3 | -1.2065 | 5.19E-04 |
| Pcdhga4 | -0.97189 | 0.037545 |
| Pcdhga5 | -1.5828 | 2.00E-04 |
| Pcdhga7 | -1.45 | 1.64E-05 |
| Pcdhga8 | -1.7682 | 2.35E-04 |
| Pcdhga9 | -1.6215 | 0.03198 |
| Pcdhgb5 | -0.82823 | 0.040723 |
| Pcdhgb7 | -1.4225 | 3.19E-04 |
| Pcdhgb8 | -1.3032 | 7.62E-04 |
| Pcgf3 | -0.86468 | 0.029615 |
| Pcmtd2 | 0.6935 | 0.022275 |
| Pcnxl2 | -0.75624 | 0.016737 |
| Pcnxl3 | -1.186 | 3.44E-05 |
| Pcsk1 | 1.0569 | 0.0043933 |
| Pcyt1b | -1.1128 | 0.0097904 |
| Pdcd11 | -0.87027 | 0.0044371 |
| Pdcd2 | 0.91299 | 0.0037578 |
| Pdcd5 | 1.2916 | 1.12E-05 |
| Pdcd6 | 0.75106 | 0.016814 |
| Pde10a | -1.2986 | 1.82E-04 |
| Pde4a | -1.0665 | 6.96E-04 |
| Pde4d | -1.0506 | 0.0078631 |

| | | |
|---|---|---|
| Pdgfb | -0.9004 | 0.037988 |
| Pdia4 | 0.7052 | 0.042503 |
| Pdia6 | 0.74203 | 0.041785 |
| Pdp1 | 0.71353 | 0.032096 |
| Pdp2 | -1.0023 | 0.024702 |
| Pdpk1 | -1.7854 | 6.89E-06 |
| Pdpr | -1.504 | 7.46E-06 |
| Pdrg1 | 0.98435 | 0.0012947 |
| Pds5a | -0.81367 | 0.016132 |
| Pdxdc1 | -0.84438 | 0.036388 |
| Pdxk | -0.79396 | 0.023696 |
| Pdzd7 | -1.2718 | 0.014073 |
| Peak1 | -1.6577 | 6.03E-08 |
| Peli2 | -0.97042 | 0.030056 |
| Per3 | -1.1871 | 0.0028126 |
| Pet100 | 1.1028 | 2.95E-04 |
| Pex5l | -1.6495 | 1.61E-08 |
| Pfas | -1.0361 | 7.09E-04 |
| Pfdn1 | 1.634 | 3.41E-09 |
| Pfdn2 | 1.2922 | 0.01119 |
| Pfdn4 | 1.3828 | 2.11E-06 |
| Pfdn5 | 1.4828 | 8.01E-07 |
| Pfkfb2 | -0.9628 | 0.0031544 |
| Pfkfb3 | -1.3716 | 0.0025105 |
| Pgam2 | -1.5307 | 1.96E-05 |
| Pgrmc1 | 0.82429 | 0.0051651 |
| Phactr2 | -1.3052 | 0.0073839 |
| Phactr4 | -1.1005 | 0.020266 |
| Phb | 0.77768 | 0.012449 |
| Phc3 | -1.8068 | 5.59E-06 |
| Phf2 | -0.88771 | 0.0046927 |
| Phf21b | -1.313 | 0.0091541 |
| Phf5a | 1.1085 | 2.21E-04 |
| Phka2 | -2.3553 | 1.07E-05 |
| Phlda1 | -0.92467 | 0.0051518 |
| Phlpp2 | -1.1531 | 0.010804 |
| Phospho2 | 0.85468 | 0.0071785 |
| Phpt1 | 0.97896 | 8.44E-04 |
| Pi4k2a | -0.84358 | 0.01314 |
| Pign | -1.1349 | 0.0055043 |
| Pigp | 1.0848 | 7.86E-04 |
| Pigu | 0.89231 | 0.0070053 |
| Pigw | 1.2105 | 4.36E-04 |
| Pik3c2b | -1.8981 | 1.13E-08 |

| | | |
|---|---|---|
| Pik3r1 | -0.92172 | 0.032064 |
| Pik3r3 | -1.2689 | 0.026002 |
| Pin4 | 1.7205 | 1.07E-08 |
| Pip4k2b | -1.2031 | 1.13E-04 |
| Pir | 1.0209 | 0.025113 |
| Pitpnm3 | -1.6527 | 8.55E-08 |
| Pkig | 1.3739 | 0.013912 |
| Pkn1 | -0.863 | 0.003607 |
| Pknox1 | -1.1101 | 0.010112 |
| Pla2g16 | 1.638 | 0.027797 |
| Pla2g4b | -1.4572 | 0.001496 |
| Pla2g7 | 1.9706 | 0.026285 |
| Plagl1 | -2.7337 | 9.07E-05 |
| Plcb3 | -0.84282 | 0.02623 |
| Plcxd3 | -1.4703 | 0.0032669 |
| Plekha7 | -2.3333 | 0.004092 |
| Plekhg1 | -1.107 | 0.0068977 |
| Plekhg5 | -0.83488 | 0.018604 |
| Plekhm1 | -0.99758 | 0.001537 |
| Plekhm3 | -1.9652 | 7.20E-07 |
| Plekhn1 | -1.1441 | 0.0061872 |
| Plgrkt | 0.91669 | 0.0093977 |
| Plrg1 | 0.74632 | 0.023218 |
| Pls3 | 0.77014 | 0.0096599 |
| Plxnc1 | -1.6129 | 7.77E-09 |
| Pm20d2 | -1.5825 | 4.35E-04 |
| Pmel | -1.0988 | 0.03986 |
| Pmpcb | 0.83716 | 0.0080045 |
| Pmvk | 0.77171 | 0.044692 |
| Pnoc | 1.5965 | 1.48E-05 |
| Pnpla6 | -0.90224 | 0.0035976 |
| Pnpla8 | 0.77355 | 0.0081111 |
| Pofut1 | -1.0167 | 0.0097964 |
| Pold1 | -1.1701 | 0.0042354 |
| Pole | -1.0637 | 0.0076924 |
| Pole4 | 0.97093 | 0.0015654 |
| Polg | -1.163 | 1.72E-04 |
| Polr2d | 1.0133 | 0.001353 |
| Polr2f | 0.88013 | 0.0035692 |
| Polr2j | 1.0719 | 4.52E-04 |
| Polr2k | 1.3966 | 1.62E-06 |
| Polr3k | 0.89114 | 0.0029738 |
| Polrmt | -0.84821 | 0.0082601 |
| Pom121 | -0.70495 | 0.028511 |

| | | |
|---|---|---|
| Pomc | 2.0736 | 0.030192 |
| Pon2 | 0.9676 | 0.0012637 |
| Postn | 1.5479 | 0.01702 |
| Pou2f1 | -1.3721 | 0.026274 |
| Ppa2 | 0.88589 | 0.0056325 |
| Ppargc1b | -2.4358 | 1.43E-09 |
| Ppdpf | 0.70539 | 0.031295 |
| Ppib | 1.0206 | 5.08E-04 |
| Ppil3 | 0.90633 | 0.018036 |
| Ppm1f | -1.0477 | 0.0016222 |
| Ppm1h | -1.0963 | 4.14E-04 |
| Ppm1l | -1.8413 | 1.19E-06 |
| Ppp1r16b | -1.334 | 6.45E-05 |
| Ppp1r1a | 0.98231 | 0.048208 |
| Ppp1r36 | 1.8852 | 0.0028124 |
| Ppp1r3e | -1.1125 | 0.047038 |
| Ppp1r9a | -2.2894 | 9.07E-14 |
| Ppp2cb | 0.76371 | 0.027554 |
| Ppp2r3c | 0.74796 | 0.033849 |
| Ppp3r1 | 0.61625 | 0.049852 |
| Ppp6r1 | -0.67178 | 0.02919 |
| Ppt1 | 0.81639 | 0.0070616 |
| Prdm10 | -1.0577 | 0.012787 |
| Prdm11 | -2.0766 | 4.56E-05 |
| Prdm15 | -1.4595 | 9.23E-05 |
| Prdx1 | 0.82483 | 0.0053449 |
| Prdx4 | 1.5105 | 1.05E-06 |
| Prdx6 | 1.2943 | 3.05E-04 |
| Prelid3b | 1.0443 | 0.0016585 |
| Prep | -0.70704 | 0.033849 |
| Prkaa2 | -1.3031 | 0.0010772 |
| Prkab2 | -1.2074 | 0.0012142 |
| Prkar2a | -1.4502 | 3.27E-06 |
| Prkca | -1.8081 | 1.81E-06 |
| Prorsd1 | 1.4112 | 0.040266 |
| Proser1 | -0.72224 | 0.043792 |
| Proser2 | -1.8007 | 0.0080988 |
| Prpf3 | 0.80125 | 0.020327 |
| Prpf38a | 0.69273 | 0.038329 |
| Prr11 | -1.8219 | 0.041105 |
| Prr14l | -1.4059 | 2.53E-04 |
| Prr36 | -1.5532 | 4.54E-08 |
| Prr5 | -0.97299 | 0.0042861 |
| Prrc1 | -0.96127 | 0.021167 |

| | | |
|---|---|---|
| Prrc2a | -1.314 | 1.50E-06 |
| Prrc2b | -2.3826 | 2.89E-18 |
| Prrt4 | -0.83007 | 0.016513 |
| Psd3 | -1.8621 | 2.71E-07 |
| Psenen | 1.0621 | 8.97E-04 |
| Psma1 | 0.68306 | 0.03027 |
| Psma2 | 0.90236 | 0.0023335 |
| Psma3 | 0.62732 | 0.041387 |
| Psma4 | 0.80482 | 0.008584 |
| Psma5 | 0.80913 | 0.0092365 |
| Psma6 | 0.6905 | 0.029002 |
| Psmb1 | 0.88689 | 0.0029359 |
| Psmb10 | 1.2924 | 0.0043827 |
| Psmb4 | 1.0573 | 2.49E-04 |
| Psmb5 | 1.1149 | 0.04504 |
| Psmb6 | 0.91572 | 0.037611 |
| Psmc1 | 0.694 | 0.04553 |
| Psmc2 | 0.91632 | 0.0039794 |
| Psmc4 | 0.83724 | 0.007807 |
| Psmc6 | 0.70921 | 0.020984 |
| Psme1 | 1.0602 | 0.04026 |
| Psme2 | 0.83454 | 0.018652 |
| Psmg2 | 0.9212 | 0.006819 |
| Psmg4 | 1.4413 | 2.01E-06 |
| Pstpip1 | 1.8016 | 7.14E-04 |
| Pstpip2 | -1.3872 | 0.023939 |
| Ptar1 | -1.7328 | 0.001788 |
| Ptchd2 | -1.5309 | 3.68E-06 |
| Ptger1 | -1.6356 | 6.32E-07 |
| Ptk7 | -0.9628 | 0.0059413 |
| Ptpmt1 | 0.99007 | 0.0021648 |
| Ptpn14 | -2.4692 | 1.01E-08 |
| Ptpn23 | -1.2964 | 7.43E-06 |
| Ptprv | -1.8872 | 0.010418 |
| Ptprz1 | -0.91271 | 0.0016789 |
| Ptrh2 | 0.70167 | 0.048576 |
| Pts | 1.0499 | 0.0019144 |
| Pura | -0.97328 | 0.003357 |
| Purg | -1.7161 | 2.57E-06 |
| Pvrl1 | -1.7876 | 1.19E-09 |
| Pxmp2 | 1.3472 | 1.96E-04 |
| Pxn | -0.84196 | 0.020865 |
| Pycr1 | -0.80067 | 0.03484 |
| Rab11fip4 | -1.0646 | 4.09E-04 |

| | | |
|---|---|---|
| Rab13 | 0.83987 | 0.014355 |
| Rab18 | 0.59945 | 0.049966 |
| Rab23 | -0.7921 | 0.024303 |
| Rab28 | 0.8677 | 0.0033795 |
| Rab3c | -1.5228 | 1.92E-05 |
| Rab3d | -1.1642 | 3.74E-04 |
| Rabgap1l | -1.5621 | 9.23E-05 |
| Rabggta | -0.74942 | 0.02409 |
| Rabggtb | 0.70859 | 0.025131 |
| Rad54l2 | -1.2687 | 2.57E-04 |
| Ralgapa2 | -1.3686 | 0.0053279 |
| Ralgps1 | -1.8283 | 2.30E-08 |
| Ralgps2 | -1.2356 | 0.0019143 |
| Rangrf | 0.81472 | 0.037853 |
| Rap1a | 0.69483 | 0.023969 |
| Rap2b | -0.97246 | 0.0056325 |
| Rapgef6 | -0.78018 | 0.035272 |
| Raph1 | -1.6412 | 6.97E-07 |
| Rarb | -1.5813 | 0.0090752 |
| Rars | 0.79509 | 0.012957 |
| Rasa1 | -0.76048 | 0.022058 |
| Rasa3 | -1.2331 | 1.99E-05 |
| Rasal1 | -0.85319 | 0.044045 |
| Rasal2 | -1.1444 | 0.0081111 |
| Rasd2 | -1.0857 | 3.19E-04 |
| Rassf8 | -1.3547 | 0.0080159 |
| Raver1 | -0.76956 | 0.013505 |
| Raver2 | -0.96872 | 0.0019553 |
| Rbfox2 | -1.8164 | 1.48E-10 |
| Rbfox3 | -1.0861 | 2.19E-04 |
| Rbks | 0.90183 | 0.036311 |
| Rbm19 | -1.2008 | 0.0021113 |
| Rbm24 | 1.2826 | 5.34E-04 |
| Rbm33 | -1.0576 | 0.012399 |
| Rbm48 | 1.0122 | 0.0061394 |
| Rbm8a | 0.87693 | 0.0043948 |
| Rcbtb1 | -0.72095 | 0.046343 |
| Rcc2 | -1.3733 | 9.68E-06 |
| Rchy1 | 0.76906 | 0.018277 |
| Rcn2 | 0.6497 | 0.038199 |
| Rcor1 | -1.8494 | 3.26E-04 |
| Reck | -0.88568 | 0.042743 |
| Recql4 | -1.4745 | 0.0061763 |
| Ren | -1.5026 | 0.028519 |

| | | |
|---|---|---|
| Rer1 | 0.67908 | 0.032131 |
| Rexo1 | -1.0422 | 4.03E-04 |
| Rfx1 | -0.96907 | 0.0028787 |
| Rfx3 | -1.5692 | 1.28E-04 |
| RGD1304567 | 0.83186 | 0.014061 |
| RGD1304694 | -1.0284 | 4.91E-04 |
| RGD1304704 | 0.78676 | 0.012337 |
| RGD1304884 | -1.1488 | 1.11E-04 |
| RGD1305464 | -0.86402 | 0.025844 |
| RGD1305537 | 0.94809 | 0.041748 |
| RGD1305587 | 0.93977 | 0.0085723 |
| RGD1305645 | 2.1007 | 3.81E-05 |
| RGD1306063 | 1.2884 | 1.71E-05 |
| RGD1306215 | 0.88309 | 0.012599 |
| RGD1306233 | 1.1794 | 0.044663 |
| RGD1306271 | -2.2868 | 6.62E-15 |
| RGD1306941 | 0.80295 | 0.010725 |
| RGD1308147 | 0.73398 | 0.027816 |
| RGD1308430 | 0.71999 | 0.029929 |
| RGD1309730 | 1.2837 | 0.0014854 |
| RGD1310081 | -0.82414 | 0.012611 |
| RGD1310769 | 0.91608 | 0.0018749 |
| RGD1311745 | 0.97703 | 0.005048 |
| RGD1311747 | 1.0299 | 0.0010614 |
| RGD1359158 | 0.76483 | 0.026401 |
| RGD1559786 | 0.67919 | 0.046038 |
| RGD1560065 | 0.81123 | 0.0092331 |
| RGD1560108 | -0.96937 | 8.23E-04 |
| RGD1560175 | -0.79424 | 0.041738 |
| RGD1560394 | 1.6416 | 0.020221 |
| RGD1560398 | 0.67818 | 0.03948 |
| RGD1560470 | -0.72218 | 0.021151 |
| RGD1560687 | 2.1767 | 0.02962 |
| RGD1561034 | -1.2105 | 0.0011278 |
| RGD1561113 | 0.99563 | 0.0016428 |
| RGD1561149 | -1.237 | 6.98E-04 |
| RGD1561590 | 0.74052 | 0.0198 |
| RGD1561897 | -1.4046 | 7.67E-04 |
| RGD1561931 | -2.0597 | 1.83E-09 |
| RGD1562310 | -1.1402 | 4.66E-04 |
| RGD1562339 | -1.5399 | 4.26E-06 |
| RGD1562747 | 0.76438 | 0.036051 |
| RGD1562987 | 0.8665 | 0.0036393 |
| RGD1563441 | 1.0107 | 0.013232 |

| | | |
|---|---|---|
| RGD1563941 | 1.3346 | 0.01366 |
| RGD1563986 | -1.6669 | 1.35E-07 |
| RGD1564093 | 0.818 | 0.036379 |
| RGD1565363 | 1.6445 | 5.04E-04 |
| RGD1565616 | -0.88357 | 0.0043222 |
| RGD1565641 | 1.37 | 1.03E-05 |
| RGD1565685 | 0.76715 | 0.037466 |
| RGD1565784 | 0.69133 | 0.042644 |
| RGD1566099 | 0.8666 | 0.020499 |
| RGD1566239 | 1.1413 | 2.79E-04 |
| RGD1566320 | 1.2397 | 3.94E-05 |
| RGD1566386 | 0.95168 | 0.0057864 |
| RGD735065 | 0.71411 | 0.029389 |
| Rgl2 | -1.1452 | 3.39E-04 |
| Rgma | -0.94018 | 0.0011212 |
| Rgmb | -0.99074 | 0.0025064 |
| Rgp1 | -1.1871 | 2.84E-04 |
| Rgs10 | 1.657 | 2.96E-08 |
| Rgs14 | -0.80529 | 0.020578 |
| Rgs2 | 1.1856 | 0.0021986 |
| Rgs6 | -0.88501 | 0.011859 |
| Rhbdf1 | -1.3447 | 3.47E-05 |
| Rheb | 0.64788 | 0.041752 |
| Ric1 | -1.0644 | 0.0021812 |
| Rictor | -1.0019 | 0.034067 |
| Rimkla | -1.1087 | 0.0020435 |
| Rims1 | -0.92156 | 0.0055272 |
| Rims3 | -1.0588 | 0.0090321 |
| Rims4 | -1.7533 | 6.49E-09 |
| Rin1 | -1.721 | 0.01264 |
| Ripply2 | 1.5552 | 0.012266 |
| Rlim | -1.782 | 1.80E-06 |
| Rmdn1 | 0.86561 | 0.010467 |
| Rmi2 | -2.2106 | 0.0033089 |
| Rmnd5a | -0.98183 | 0.0028744 |
| Rn50_1_1359.1 | 1.2769 | 1.87E-05 |
| Rn50_10_0635.1 | -4.7814 | 2.87E-04 |
| Rn50_10_0702.4 | 1.348 | 6.74E-04 |
| Rn50_13_0822.1 | -2.6675 | 1.03E-07 |
| Rn50_X_0635.3 | 1.3199 | 5.25E-04 |
| Rn50_X_0746.3 | 1.7562 | 0.049966 |
| Rn50_X_0747.1 | -1.0481 | 5.96E-04 |
| Rn50_X_0752.3 | -8.3883 | 4.14E-04 |
| Rn60_14_0788.1 | 2.0852 | 6.58E-13 |

| | | |
|---|---|---|
| Rnasek | 1.3469 | 1.43E-06 |
| Rnaset2 | 1.3501 | 1.31E-05 |
| Rnd2 | 1.5557 | 2.03E-08 |
| Rnd3 | 0.76537 | 0.035667 |
| Rnf113a1 | 0.77915 | 0.037694 |
| Rnf123 | -0.83827 | 0.0071785 |
| Rnf138 | 0.84053 | 0.0098263 |
| Rnf139 | 0.65675 | 0.043888 |
| Rnf144a | -1.8013 | 2.71E-08 |
| Rnf146 | 0.69894 | 0.028154 |
| Rnf150 | -1.3402 | 1.40E-04 |
| Rnf157 | -1.5035 | 2.94E-07 |
| Rnf165 | -1.0586 | 2.52E-04 |
| Rnf181 | 0.83905 | 0.0067385 |
| Rnf182 | -1.2959 | 0.0016932 |
| Rnf213 | -1.4537 | 0.0017131 |
| Rnf214 | -0.98697 | 0.0023635 |
| Rnf38 | -1.0131 | 0.005727 |
| Rnf44 | -1.5344 | 2.35E-07 |
| Rnf7 | 1.2183 | 3.16E-05 |
| Robo1 | -1.6433 | 2.40E-06 |
| Romo1 | 1.3092 | 0.0015276 |
| Rpa3 | 0.71006 | 0.048301 |
| Rpain | 0.80921 | 0.037436 |
| Rpap1 | -0.87106 | 0.010961 |
| Rpf2 | 0.96904 | 0.0020931 |
| Rpl10a | 1.172 | 3.10E-05 |
| Rpl11 | 1.6199 | 1.87E-05 |
| Rpl12 | 1.4105 | 6.60E-07 |
| Rpl14 | 1.3707 | 1.92E-06 |
| Rpl15 | 1.0915 | 1.42E-04 |
| Rpl17 | 1.3344 | 1.79E-06 |
| Rpl18 | 1.2584 | 0.0010297 |
| Rpl18a | 1.4879 | 2.62E-05 |
| Rpl19 | 2.0831 | 0.010191 |
| Rpl21 | 1.7488 | 1.24E-05 |
| Rpl21 | 1.6614 | 7.21E-09 |
| Rpl22l1 | 1.6071 | 6.66E-04 |
| Rpl23 | 1.5626 | 1.13E-08 |
| Rpl23a | 1.5824 | 6.53E-09 |
| Rpl24 | 1.3602 | 1.07E-06 |
| Rpl26 | 1.7386 | 1.69E-10 |
| Rpl26-ps2 | 1.9768 | 0.025425 |
| Rpl27 | 1.3236 | 1.55E-06 |

| | | |
|---|---|---|
| Rpl27a | 1.4909 | 5.01E-08 |
| Rpl28 | 1.4709 | 1.52E-07 |
| Rpl29 | 1.3847 | 4.89E-05 |
| Rpl3 | 0.9931 | 0.0078614 |
| Rpl31 | 1.537 | 1.71E-08 |
| Rpl32 | 1.8071 | 5.80E-08 |
| Rpl34 | 1.43 | 2.08E-07 |
| Rpl35 | 1.8033 | 7.49E-05 |
| Rpl35a | 1.2786 | 0.029843 |
| Rpl36 | 1.4237 | 3.40E-07 |
| Rpl36al | 1.2996 | 5.05E-06 |
| Rpl37a | 1.4662 | 6.30E-07 |
| Rpl38 | 2.2339 | 0.04386 |
| Rpl39 | 1.6948 | 9.20E-10 |
| Rpl4 | 1.1374 | 9.43E-04 |
| Rpl41 | 1.7687 | 6.13E-11 |
| Rpl5 | 1.2878 | 3.27E-05 |
| Rpl6 | 1.0728 | 0.025568 |
| Rpl7 | 1.4917 | 1.26E-07 |
| Rpl9 | 1.5702 | 3.93E-06 |
| Rplp0 | 1.0007 | 4.87E-04 |
| Rpn2 | 0.70241 | 0.029557 |
| Rpo1-3 | 0.80938 | 0.008778 |
| Rpp21 | 1.2117 | 0.0031392 |
| Rpp30 | 0.87811 | 0.0081111 |
| Rprd1a | -0.71789 | 0.046875 |
| Rps10 | 1.2796 | 6.12E-05 |
| Rps11 | 1.3784 | 5.69E-07 |
| Rps12 | 1.6058 | 4.47E-09 |
| Rps14 | 1.6436 | 1.52E-09 |
| Rps15 | 1.7194 | 7.08E-05 |
| Rps15a | 1.5769 | 7.74E-09 |
| Rps16 | 1.5197 | 5.31E-08 |
| Rps17 | 1.7253 | 4.79E-10 |
| Rps18 | 1.3507 | 1.14E-06 |
| Rps2 | 1.121 | 7.04E-05 |
| Rps20 | 1.4772 | 7.12E-08 |
| Rps23 | 1.5808 | 1.98E-05 |
| Rps24 | 1.6272 | 2.98E-09 |
| Rps25 | 1.7427 | 1.57E-10 |
| Rps27 | 1.7112 | 2.36E-10 |
| Rps27a | 1.4159 | 1.67E-07 |
| Rps27l | 0.94254 | 0.01854 |
| Rps28 | 2.1022 | 1.14E-11 |

| | | |
|---|---|---|
| Rps29 | 1.9371 | 3.12E-08 |
| Rps3 | 1.2499 | 2.30E-04 |
| Rps3a | 1.4807 | 6.40E-08 |
| Rps4x | 1.5327 | 2.12E-08 |
| Rps5 | 1.4598 | 3.64E-07 |
| Rps6ka1 | -0.8797 | 0.048794 |
| Rps6ka2 | -0.90956 | 0.0031714 |
| Rps7 | 1.5728 | 4.50E-06 |
| Rrp1b | -1.1122 | 0.036139 |
| Rtel1 | -0.96773 | 0.0021448 |
| Runx1t1 | -1.321 | 1.71E-05 |
| Rwdd3 | 0.98818 | 0.038916 |
| Rxra | -1.1738 | 0.0010657 |
| Ryr1 | -2.3668 | 0.0028744 |
| S100a1 | 1.7224 | 1.82E-08 |
| S100a10 | 0.89545 | 0.0030186 |
| S100a13 | 1.3762 | 1.85E-05 |
| S100a16 | 1.2325 | 7.21E-05 |
| S100b | 1.2186 | 1.03E-05 |
| Sall3 | -1.3657 | 5.60E-05 |
| Samd12 | -1.038 | 0.034719 |
| Samd14 | -0.87964 | 0.004799 |
| Samd5 | -1.5525 | 1.77E-04 |
| Samd8 | -1.2681 | 6.42E-04 |
| Sap25 | -2.7091 | 0.032486 |
| Sapcd2 | -1.3396 | 5.78E-05 |
| Sar1b | 0.77603 | 0.0109 |
| Sarm1 | -1.3543 | 3.29E-06 |
| Sarnp | 1.2198 | 2.18E-05 |
| Sat1 | 1.5804 | 2.22E-08 |
| Sat2 | 0.79015 | 0.048445 |
| Sbk1 | -0.98315 | 8.76E-04 |
| Sbno2 | -0.93464 | 0.0035976 |
| Scai | -1.318 | 0.012468 |
| Scg2 | 0.83729 | 0.028945 |
| Scg3 | 0.77556 | 0.012306 |
| Scg5 | 1.4116 | 0.004703 |
| Sclt1 | 0.9109 | 0.031688 |
| Scn2a | -1.2163 | 3.28E-05 |
| Scn2b | -1.0078 | 0.0014608 |
| Scn5a | -2.2863 | 0.020748 |
| Scpep1 | 0.73942 | 0.03008 |
| Scrib | -0.95879 | 0.0013521 |
| Scrn1 | -1.1171 | 1.04E-04 |

| | | |
|---|---|---|
| Scrt1 | -1.4777 | 3.18E-06 |
| Scrt2 | -2.1426 | 1.93E-06 |
| Scube1 | -1.5798 | 7.47E-08 |
| Scube3 | -3.7698 | 1.03E-09 |
| Sdc2 | 0.71615 | 0.033673 |
| Sdc3 | -1.4669 | 1.14E-07 |
| Sdc4 | 1.0712 | 3.61E-04 |
| Sdcbp | 0.62783 | 0.044995 |
| Sdf2 | 1.0497 | 4.01E-04 |
| Sdhaf3 | 0.81938 | 0.043478 |
| Sdhaf4 | 1.0337 | 7.13E-04 |
| Sdhb | 0.66583 | 0.039256 |
| Sdhc | 0.79378 | 0.0083305 |
| Sdhd | 0.65418 | 0.041814 |
| Sdk1 | -1.6631 | 2.77E-07 |
| Sdk2 | -2.3083 | 1.40E-11 |
| Sec11a | 0.85389 | 0.0062253 |
| Sec11c | 1.0735 | 0.0014939 |
| Sec14l2 | -0.82146 | 0.0094473 |
| Sec22c | -1.4073 | 2.66E-05 |
| Sec31a | -0.73887 | 0.019959 |
| Sec61b | 0.95495 | 0.0020141 |
| Sec61g | 1.2999 | 7.51E-04 |
| Sec62 | 0.94992 | 0.0043207 |
| Sel1l | -1.0572 | 4.97E-04 |
| Selk | 0.99604 | 6.38E-04 |
| Sema3e | -0.79187 | 0.044047 |
| Sema4b | -1.0424 | 0.0089625 |
| Sema4g | -0.77644 | 0.021928 |
| Sema5b | -1.3963 | 2.35E-05 |
| Sema6a | -1.1044 | 0.0033212 |
| Sepn1 | -1.0491 | 0.0015167 |
| Sepw1 | 1.2147 | 0.0026909 |
| Serf1 | 0.90507 | 0.0043046 |
| Serf2 | 1.625 | 7.39E-08 |
| Serinc1 | 1.0682 | 0.002779 |
| Serinc3 | 0.77805 | 0.0082813 |
| Serinc4 | 0.90557 | 0.0040723 |
| Serpina11 | 1.5411 | 0.0044661 |
| Serpinb6 | 0.88957 | 0.01825 |
| Sesn1 | 0.70705 | 0.042369 |
| Sesn2 | -1.1635 | 0.0024754 |
| Sesn3 | -1.3161 | 5.74E-04 |
| Set | -0.72884 | 0.028388 |

| | | |
|---|---|---|
| Setbp1 | -2.1175 | 1.66E-10 |
| Setd1b | -2.2968 | 2.22E-14 |
| Setd7 | -1.5676 | 1.02E-05 |
| Sez6 | -0.95695 | 4.58E-04 |
| Sez6l | -1.4555 | 1.56E-07 |
| Sf3a2 | -0.80795 | 0.015659 |
| Sf3b6 | 0.87146 | 0.0070596 |
| Sfmbt2 | -1.7894 | 0.0018196 |
| Sfr1 | 1.0044 | 7.81E-04 |
| Sft2d1 | 0.80856 | 0.010628 |
| Sgcd | -1.8951 | 0.031688 |
| Sgpp2 | -2.0983 | 0.0094488 |
| Sgsm1 | -0.64306 | 0.03663 |
| Sh2b2 | -1.1969 | 0.0109 |
| Sh2d1a | 1.6671 | 0.0031388 |
| Sh3bgrl3 | 0.8721 | 0.023413 |
| Sh3d19 | -1.2807 | 7.24E-04 |
| Sh3gl3 | 0.74184 | 0.020641 |
| Sh3pxd2a | -2.1772 | 1.32E-09 |
| Sh3pxd2b | -1.9141 | 2.18E-08 |
| Sh3rf1 | -1.1076 | 5.89E-04 |
| Sh3rf3 | -1.8816 | 7.53E-06 |
| Shc2 | -0.70967 | 0.025034 |
| Shc3 | -2.2597 | 2.07E-12 |
| Shc4 | -1.8236 | 0.0061763 |
| Shfm1 | 1.516 | 6.56E-08 |
| Shisa6 | -2.9192 | 6.39E-14 |
| Shroom3 | -0.87416 | 0.0087144 |
| Shroom4 | -1.9564 | 0.030804 |
| Shtn1 | -0.87249 | 0.0075801 |
| Siah3 | -2.6274 | 4.85E-08 |
| Sidt2 | -0.64521 | 0.040476 |
| Sik1 | -1.3955 | 0.039588 |
| Sik3 | -0.69121 | 0.030544 |
| Sin3b | -0.81773 | 0.0070099 |
| Sipa1 | -1.4013 | 0.0069372 |
| Sirt3 | 0.75767 | 0.019736 |
| Siva1 | 0.96218 | 0.014052 |
| Ska2 | 0.86671 | 0.014409 |
| Skil | -1.0725 | 0.0039616 |
| Skp1 | 0.90028 | 0.0018444 |
| Slc10a1 | 1.8935 | 0.024389 |
| Slc16a14 | -1.2605 | 0.0066219 |
| Slc16a2 | -1.7807 | 9.90E-10 |

| | | |
|---|---|---|
| Slc16a7 | -1.1041 | 0.0024678 |
| Slc18a2 | 1.2507 | 9.36E-04 |
| Slc20a1 | 0.68179 | 0.029512 |
| Slc22a12 | -1.2569 | 0.0041583 |
| Slc25a2 | 1.2984 | 0.022434 |
| Slc25a5 | 0.7114 | 0.019325 |
| Slc25a51 | 0.8867 | 0.0033813 |
| Slc2a12 | 1.8425 | 6.16E-06 |
| Slc30a10 | -1.0541 | 0.0097964 |
| Slc30a6 | 0.7097 | 0.049772 |
| Slc35e2b | -0.92782 | 0.041738 |
| Slc35f6 | 0.72961 | 0.028115 |
| Slc35g2 | 1.5704 | 0.037032 |
| Slc35g3 | -1.7627 | 0.034012 |
| Slc36a1 | -1.904 | 4.91E-06 |
| Slc36a4 | -0.9887 | 0.0043222 |
| Slc38a1 | -1.1595 | 1.37E-04 |
| Slc38a7 | -0.94882 | 0.005178 |
| Slc39a9 | -1.1007 | 0.0022083 |
| Slc41a1 | -0.7399 | 0.021066 |
| Slc44a5 | -1.699 | 0.0071636 |
| Slc45a4 | -1.267 | 3.39E-04 |
| Slc4a7 | -1.532 | 5.98E-05 |
| Slc6a11 | -1.2334 | 0.0021604 |
| Slc6a6 | -1.4029 | 6.76E-06 |
| Slc7a14 | -1.8829 | 3.43E-09 |
| Slc7a6os | 0.71125 | 0.036212 |
| Slc9a1 | -0.82794 | 0.0072121 |
| Slc9a3r1 | -0.78523 | 0.0158 |
| Slc9a5 | -0.87802 | 0.013077 |
| Slc9a7 | -1.9128 | 9.25E-08 |
| Slc9a8 | -0.7464 | 0.045842 |
| Slco3a1 | -1.4092 | 4.90E-06 |
| Slco5a1 | -2.3084 | 1.56E-04 |
| Slf2 | 0.84843 | 0.0044271 |
| Slirp | 1.0824 | 0.010349 |
| Slit1 | -1.5566 | 1.96E-08 |
| Slit3 | -1.1837 | 5.79E-05 |
| Slitrk2 | -0.82478 | 0.033673 |
| Slmap | -0.7348 | 0.030518 |
| Smad5 | -1.0853 | 0.0020703 |
| Smarcc1 | -0.80382 | 0.012334 |
| Smcr8 | -1.1301 | 0.014346 |
| Smdt1 | 0.76463 | 0.013054 |

| | | |
|---|---|---|
| Smg5 | -1.0158 | 8.29E-04 |
| Smim11 | 1.4405 | 1.92E-05 |
| Smim13 | -0.96426 | 0.0044164 |
| Smim18 | 0.81232 | 0.030562 |
| Smim3 | 1.2413 | 0.0058493 |
| Smim7 | 0.7201 | 0.022362 |
| Smim8 | 1.0379 | 0.003268 |
| Smoc1 | -0.99682 | 0.0017704 |
| Smurf1 | -1.6206 | 1.31E-07 |
| Snap47 | 0.71415 | 0.029477 |
| Snapin | 0.87387 | 0.003378 |
| Snf8 | 1.1221 | 0.0063598 |
| Snrnp25 | 1.2029 | 1.28E-04 |
| Snrpb2 | 0.8555 | 0.0090321 |
| Snrpc | 0.69951 | 0.048716 |
| Snrpd1 | 1.0567 | 3.26E-04 |
| Snrpd2 | 1.2933 | 6.28E-04 |
| Snrpd3 | 0.71982 | 0.026042 |
| Snrpe | 1.2564 | 1.48E-05 |
| Snrpf | 1.1775 | 1.03E-04 |
| Sntb2 | -1.8795 | 0.0047506 |
| Snx29 | -1.3504 | 1.05E-04 |
| Snx3 | 0.7634 | 0.012404 |
| Snx30 | -1.3593 | 0.002136 |
| Soga3 | -1.3728 | 1.18E-05 |
| Sorbs1 | -1.4989 | 6.89E-04 |
| Sorcs2 | -0.6812 | 0.030433 |
| Sorcs3 | -1.4289 | 1.66E-06 |
| Sox11 | -1.056 | 6.98E-04 |
| Sox12 | -0.96106 | 0.001508 |
| Sox5 | -0.99471 | 0.015485 |
| Sox6 | -1.8078 | 9.19E-04 |
| Sox9 | -1.3243 | 0.0011202 |
| Sp2 | -0.98739 | 0.011381 |
| Spag7 | 0.81578 | 0.022127 |
| Sparc | 0.91247 | 0.0024168 |
| Spata1 | 1.4364 | 0.040102 |
| Spata13 | -2.6056 | 9.92E-04 |
| Spata2 | -0.7529 | 0.023904 |
| Spcs1 | 1.1666 | 4.18E-05 |
| Spcs2 | 0.66434 | 0.036767 |
| Specc1 | -0.92463 | 0.0067385 |
| Spock1 | -0.68957 | 0.041738 |
| Spop | 0.92996 | 0.013185 |

| | | |
|---|---|---|
| Spred2 | -1.1659 | 1.66E-04 |
| Spryd7 | 0.75092 | 0.038174 |
| Spta1 | 1.2782 | 0.0012587 |
| Sptlc2 | -0.87234 | 0.013051 |
| Sptssa | 1.1198 | 1.17E-04 |
| Sqrdl | 1.4481 | 0.015119 |
| Src | -0.6798 | 0.040287 |
| Srgap1 | -1.9068 | 3.06E-08 |
| Srgap2 | -1.041 | 6.92E-04 |
| Srp14 | 1.0401 | 2.87E-04 |
| Srp19 | 1.0618 | 3.43E-04 |
| Srp9 | 0.98541 | 8.04E-04 |
| Srrm3 | -1.0868 | 3.00E-04 |
| Srrm4 | -2.0757 | 8.60E-10 |
| Srrt | -0.73537 | 0.015432 |
| Ss18l2 | 1.1053 | 1.37E-04 |
| Ssh1 | -2.0517 | 9.55E-08 |
| Ssh2 | -1.5172 | 6.59E-06 |
| Ssr2 | 0.83496 | 0.0072046 |
| Ssr3 | 0.88275 | 0.0022395 |
| Sst | 0.84499 | 0.0036411 |
| Ssu72 | 0.90659 | 0.0071892 |
| St18 | -2.3886 | 1.39E-07 |
| St3gal1 | -1.1352 | 0.0023635 |
| St7l | 0.70317 | 0.030509 |
| St8sia3 | -1.5296 | 2.23E-05 |
| Stambp | 0.9108 | 0.005048 |
| Stard10 | 1.2918 | 0.0083838 |
| Stard6 | 1.039 | 0.039133 |
| Stard9 | -1.5519 | 0.0066738 |
| Stat5b | -1.2091 | 4.47E-04 |
| Stbd1 | 0.95293 | 0.017605 |
| Stc2 | -1.3684 | 0.048006 |
| Stk11ip | -0.80152 | 0.014822 |
| Stk36 | -1.255 | 0.0023635 |
| Stoml2 | 0.7676 | 0.021231 |
| Ston2 | -1.7587 | 0.0014792 |
| Strn | -1.1423 | 0.015889 |
| Stt3a | 0.64368 | 0.042914 |
| Stx8 | 1.1944 | 1.51E-04 |
| Stxbp3 | 1.0197 | 0.0015218 |
| Stxbp4 | -1.0949 | 0.018823 |
| Stxbp5l | -1.3226 | 0.0068629 |
| Styx | -1.0339 | 0.049784 |

| | | |
|---|---|---|
| Sub1 | 0.90731 | 0.002535 |
| Sufu | -1.2119 | 0.0029455 |
| Sugt1 | 1.1664 | 0.0051442 |
| Sulf2 | -0.80354 | 0.010201 |
| Sult2b1 | -1.1705 | 2.53E-04 |
| Sumf2 | 0.84474 | 0.027857 |
| Sumo1 | 1.112 | 7.35E-05 |
| Sumo3 | 0.73394 | 0.0198 |
| Sumo4 | 1.409 | 8.75E-07 |
| Supt4h1 | 1.3559 | 3.24E-06 |
| Supt5h | -0.80552 | 0.0051948 |
| Supt6h | -0.67637 | 0.032135 |
| Susd6 | -1.2229 | 0.028723 |
| Svbp | 1.1642 | 0.0016691 |
| Swi5 | 0.73197 | 0.033673 |
| Syap1 | 0.85208 | 0.0077268 |
| Syn2 | -0.75234 | 0.014682 |
| Syn3 | -1.7158 | 2.01E-07 |
| Syt11 | -0.66539 | 0.042166 |
| Syt14 | -1.0844 | 0.017556 |
| Syt6 | -1.8999 | 1.46E-06 |
| Syt9 | -0.88892 | 0.034011 |
| Szt2 | -1.9158 | 1.41E-10 |
| Taf13 | 1.1865 | 2.48E-05 |
| Taf3 | 0.67747 | 0.029092 |
| Tanc1 | -1.9826 | 2.38E-10 |
| Taok3 | -0.91891 | 0.0038601 |
| Tarsl2 | -0.90474 | 0.0088622 |
| Tatdn1 | 0.76722 | 0.033925 |
| Tatdn2 | -0.66484 | 0.044984 |
| Tbc1d13 | -1.1916 | 3.56E-04 |
| Tbc1d16 | -2.0083 | 1.14E-07 |
| Tbc1d22a | -0.91383 | 0.017405 |
| Tbc1d32 | -0.97681 | 0.016965 |
| Tbc1d4 | -1.2142 | 0.02779 |
| Tbca | 2.1501 | 0.049895 |
| Tbca | 1.2745 | 6.81E-06 |
| Tbcel | -1.2835 | 5.25E-04 |
| Tbr1 | -1.3716 | 1.22E-05 |
| Tceal1 | 0.72659 | 0.027635 |
| Tceal7 | 1.4825 | 0.029398 |
| Tceal8 | 1.1078 | 1.17E-04 |
| Tceb1 | 0.81397 | 0.0090769 |
| Tceb2 | 1.2873 | 0.0045413 |

| | | |
|---|---|---|
| Tcf7l1 | -0.96478 | 0.013314 |
| Tcp1 | 0.67833 | 0.033456 |
| Tcp11l1 | -1.8606 | 1.57E-09 |
| Tcp11l2 | 0.97658 | 0.0049623 |
| Tcte1 | 1.5699 | 0.0079927 |
| Tctex1d2 | 1.1939 | 1.04E-04 |
| Tead1 | -2.366 | 1.26E-07 |
| Tecpr1 | -0.82246 | 0.009382 |
| Tecpr2 | -1.1543 | 2.24E-04 |
| Tecr | 0.68401 | 0.025708 |
| Tenm2 | -1.0783 | 2.10E-04 |
| Tenm4 | -2.4163 | 7.89E-16 |
| Tet1 | -1.9357 | 1.09E-06 |
| Tet2 | -1.0946 | 0.0016691 |
| Tet3 | -2.4968 | 2.20E-16 |
| Tfap2c | -2.3906 | 0.048794 |
| Tfe3 | -0.79177 | 0.033244 |
| Tfpi | 0.9283 | 0.019331 |
| Tfr2 | -1.5586 | 0.0030212 |
| Tgfa | -1.2768 | 0.0099424 |
| Thnsl2 | 1.0175 | 0.039528 |
| Thoc7 | 0.99175 | 0.0010839 |
| Tial1 | -0.85355 | 0.0068299 |
| Ticrr | -2.1554 | 1.58E-04 |
| Timm10 | 0.83715 | 0.0079089 |
| Timm10b | 1.1378 | 0.010335 |
| Timm13 | 1.1516 | 0.0042243 |
| Timm17a | 0.85726 | 0.004544 |
| Timm21 | 0.97203 | 0.0031108 |
| Timm23 | 0.72286 | 0.023001 |
| Timm8b | 1.5914 | 6.33E-09 |
| Timp2 | -1.1431 | 4.18E-05 |
| Tipin | 1.302 | 0.0016958 |
| Tipinl1 | 1.0079 | 0.0076393 |
| Tle2 | -1.4688 | 3.93E-06 |
| Tm2d1 | 0.9399 | 0.0018963 |
| Tm2d2 | 0.84886 | 0.0061618 |
| Tm2d3 | 0.83074 | 0.027819 |
| Tm9sf2 | 0.6769 | 0.026677 |
| Tmbim4 | 1.2217 | 3.83E-05 |
| Tmco5a | 1.0555 | 0.0093836 |
| Tmed4 | 0.72644 | 0.024002 |
| Tmed7 | 0.73699 | 0.020703 |
| Tmed8 | -2.2224 | 4.46E-09 |

| | | |
|---|---|---|
| Tmem104 | -1.3728 | 5.56E-04 |
| Tmem107 | 0.76137 | 0.048092 |
| Tmem108 | -0.91766 | 0.0063729 |
| Tmem114 | -2.7334 | 0.0034906 |
| Tmem126a | 0.97914 | 0.0018997 |
| Tmem126b | 0.84003 | 0.027276 |
| Tmem128 | 0.76016 | 0.016605 |
| Tmem132b | -2.1785 | 7.28E-12 |
| Tmem132d | -1.3586 | 4.31E-05 |
| Tmem134 | 0.69854 | 0.043203 |
| Tmem141 | 1.2672 | 0.0015249 |
| Tmem145 | -0.86759 | 0.0031544 |
| Tmem147 | 0.6676 | 0.044214 |
| Tmem165 | 0.86062 | 0.025165 |
| Tmem17 | 0.85223 | 0.017906 |
| Tmem170b | -1.6865 | 4.32E-04 |
| Tmem179b | 1.096 | 0.010418 |
| Tmem181 | -0.69464 | 0.04291 |
| Tmem186 | 0.95837 | 0.0024877 |
| Tmem19 | 0.84312 | 0.010717 |
| Tmem191c | -0.91388 | 0.049653 |
| Tmem201 | -1.3016 | 3.90E-05 |
| Tmem205 | 1.1855 | 1.54E-04 |
| Tmem208 | 0.74311 | 0.021388 |
| Tmem216 | 1.1091 | 0.0026821 |
| Tmem218 | 0.99602 | 0.0013118 |
| Tmem223 | 1.1201 | 5.67E-04 |
| Tmem243 | 0.87634 | 0.025832 |
| Tmem245 | -1.8025 | 3.28E-08 |
| Tmem251 | 1.2332 | 2.40E-04 |
| Tmem256 | 1.2214 | 3.35E-05 |
| Tmem258b | 1.1243 | 1.94E-04 |
| Tmem261 | 1.0351 | 0.001058 |
| Tmem38b | 0.9068 | 0.011177 |
| Tmem41b | 0.73804 | 0.015277 |
| Tmem47 | 0.87179 | 0.039324 |
| Tmem50a | 0.83554 | 0.014814 |
| Tmem59 | 0.74672 | 0.016669 |
| Tmem60 | 0.91259 | 0.0059688 |
| Tmem69 | 0.69895 | 0.037396 |
| Tmem74 | -1.307 | 2.24E-04 |
| Tmem80 | 0.78773 | 0.013854 |
| Tmem82 | -2.3489 | 0.0061068 |
| Tmem86b | -1.5864 | 0.014478 |

| | | |
|---|---|---|
| Tmem94 | -1.1957 | 6.85E-05 |
| Tmem97 | 0.69469 | 0.049774 |
| Tmem98 | 0.89556 | 0.012271 |
| Tmprss6 | -1.8724 | 0.040643 |
| Tmsb4x | 1.5047 | 2.69E-08 |
| Tmsbl1 | 0.87939 | 0.0040171 |
| Tmtc1 | -1.2897 | 0.033961 |
| Tmtc2 | -1.1515 | 0.0070148 |
| Tmx2 | 0.64958 | 0.033406 |
| Tnfrsf25 | -1.4083 | 0.03484 |
| Tnks1bp1 | -0.73983 | 0.014471 |
| Tnnc1 | 1.2109 | 0.0075896 |
| Tnpo1 | -1.2935 | 5.33E-04 |
| Tnr | -1.6304 | 0.0019543 |
| Tnrc6a | -0.91044 | 0.0269 |
| Tnrc6c | -1.6748 | 4.20E-09 |
| Tns1 | -2.1058 | 0.003569 |
| Tob2 | -1.0072 | 0.011786 |
| Tom1l1 | 0.94497 | 0.0093283 |
| Tom1l2 | -1.7671 | 9.16E-10 |
| Tomm20 | 1.0556 | 2.19E-04 |
| Tomm22 | 0.97032 | 8.98E-04 |
| Tomm6 | 1.2408 | 6.33E-04 |
| Tomm7 | 1.5192 | 5.31E-08 |
| Tomm70a | -0.7542 | 0.02338 |
| Tonsl | -1.099 | 0.022058 |
| Top1 | -1.4907 | 0.0011292 |
| Topors | 0.67063 | 0.043391 |
| Tox2 | -0.97145 | 0.029016 |
| Tp53 | -1.0202 | 0.0014792 |
| Tp53bp1 | -0.68094 | 0.029517 |
| Tp53inp2 | -1.9459 | 8.44E-07 |
| Tpcn2 | -1.4796 | 0.0051082 |
| Tpt1 | 1.4463 | 2.16E-07 |
| Trabd2b | -1.9541 | 0.039378 |
| Traf3 | -1.3469 | 3.06E-06 |
| Traf4 | -0.94639 | 0.0020994 |
| Traf6 | -1.331 | 0.011099 |
| Traf7 | -0.68522 | 0.036102 |
| Trappc1 | 0.73901 | 0.021388 |
| Trappc2b | 1.1146 | 0.0012984 |
| Trappc2l | 1.0975 | 1.97E-04 |
| Trappc4 | 0.70927 | 0.041506 |
| Trappc6b | 0.90886 | 0.020195 |

| | | |
|---|---|---|
| Trerf1 | -1.1948 | 3.82E-04 |
| Trhde | -0.82787 | 0.020768 |
| Trim25 | -2.4006 | 0.0043028 |
| Trim44 | -1.4095 | 4.19E-06 |
| Trim54 | 2.4144 | 0.0067248 |
| Trim62 | -1.9019 | 2.23E-06 |
| Trim66 | -2.4377 | 2.12E-06 |
| Trim67 | -2.7112 | 1.04E-08 |
| Trmt10c | 0.82748 | 0.011849 |
| Trmt112 | 1.0554 | 0.011632 |
| Trpc5 | -1.744 | 1.16E-07 |
| Tsc1 | -1.2217 | 9.23E-05 |
| Tsen15 | 0.9328 | 0.0041293 |
| Tspan31 | 0.94328 | 0.0016623 |
| Tspan6 | 1.0135 | 5.04E-04 |
| Tspyl5 | -0.77912 | 0.036135 |
| Tstd3 | 0.86433 | 0.0063767 |
| Ttc23 | 1.2399 | 0.016659 |
| Ttc28 | -1.7532 | 5.56E-08 |
| Ttc37 | -0.72839 | 0.037377 |
| Ttc39b | -1.416 | 0.0029239 |
| Ttl | -1.0156 | 0.0018681 |
| Ttll12 | -0.84247 | 0.010961 |
| Ttll4 | -1.4266 | 5.85E-05 |
| Ttll8 | -2.2498 | 0.025778 |
| Tub | -1.7077 | 4.03E-09 |
| Tulp3 | -1.2558 | 8.32E-04 |
| Tusc2 | 0.65587 | 0.043962 |
| Tusc3 | 0.69626 | 0.031694 |
| Tvp23b | 0.7504 | 0.015729 |
| Txlna | -0.81707 | 0.037554 |
| Txn1 | 0.77127 | 0.031875 |
| Txn2 | 0.80578 | 0.021569 |
| Txndc12 | 1.1456 | 2.24E-04 |
| Txndc17 | 0.84746 | 0.0091369 |
| Txndc9 | 0.67407 | 0.041448 |
| Tyk2 | -1.0434 | 0.0032898 |
| Tysnd1 | -0.71877 | 0.041387 |
| U2af1 | 0.87095 | 0.042027 |
| U2af1l4 | 0.8906 | 0.0045722 |
| Uba52 | 1.4593 | 6.41E-04 |
| Ubash3b | -1.2866 | 2.10E-04 |
| Ube2b | 0.76411 | 0.0124 |
| Ube2e1 | 0.70434 | 0.028501 |

| | | |
|---|---|---|
| Ube2v2 | 1.0552 | 1.88E-04 |
| Ubl5 | 0.95094 | 0.0040009 |
| Ubn2 | -1.1198 | 7.77E-04 |
| Ubr1 | -0.87449 | 0.031864 |
| Ubr2 | -1.1789 | 8.85E-05 |
| Ubr4 | -0.95572 | 9.08E-04 |
| Ubxn2b | -0.95455 | 0.0059745 |
| Ubxn7 | -1.5878 | 0.0038739 |
| Uck2 | -1.3651 | 9.44E-05 |
| Ufc1 | 0.97814 | 0.0010338 |
| Ufm1 | 0.78983 | 0.0099668 |
| Ufsp2 | 0.87664 | 0.0043562 |
| Uggt1 | -1.0447 | 6.72E-04 |
| Ugp2 | 0.66069 | 0.042644 |
| Uhrf1 | -0.93063 | 0.021265 |
| Uhrf1bp1 | -2.0127 | 9.87E-10 |
| Unc119b | -1.0856 | 0.0011372 |
| Unc50 | 0.66989 | 0.046995 |
| Unc5b | -1.8777 | 0.0028654 |
| Unc5c | -1.9224 | 3.99E-07 |
| Unc5d | -2.0456 | 1.94E-09 |
| Unc79 | -0.68577 | 0.038596 |
| Unk | -1.2644 | 1.98E-04 |
| Upf1 | -1.2101 | 2.55E-05 |
| Uprt | -1.8857 | 5.27E-06 |
| Uqcc2 | 1.2944 | 4.50E-05 |
| Uqcc3 | 1.6887 | 0.0015218 |
| Uqcrb | 1.5576 | 2.40E-06 |
| Uqcrb | 1.6615 | 1.96E-06 |
| Uqcrh | 1.3484 | 1.31E-06 |
| Uqcrq | 1.1 | 0.002095 |
| Uri1 | 0.65892 | 0.040819 |
| Urod | 1.1928 | 0.0031108 |
| Usp19 | -0.71376 | 0.020363 |
| Usp24 | -1.228 | 3.33E-05 |
| Usp31 | -1.325 | 1.31E-05 |
| Usp49 | -0.92591 | 0.014667 |
| Usp54 | -0.67278 | 0.040984 |
| Utp3 | 0.72291 | 0.030374 |
| Utrn | -1.0553 | 0.00153 |
| Uvrag | -0.91843 | 0.0094484 |
| Vamp3 | 0.84664 | 0.027995 |
| Vamp8 | 1.5184 | 0.0015407 |
| Vangl2 | -1.6368 | 3.12E-06 |

| | | |
|---|---|---|
| Vars2 | -1.0791 | 5.29E-04 |
| Vash1 | -1.6962 | 8.81E-07 |
| Vav2 | -1.2097 | 6.44E-05 |
| Vax2 | -2.3151 | 0.022414 |
| Vdac3 | 0.67817 | 0.029533 |
| Vimp | 0.84958 | 0.0055722 |
| Vipr1 | -1.2523 | 0.016513 |
| Vkorc1 | 0.98318 | 0.0038613 |
| Vmp1 | 0.6707 | 0.029389 |
| Vps13a | -1.2911 | 0.0043769 |
| Vps13c | -1.1919 | 0.001822 |
| Vps29 | 0.66577 | 0.02963 |
| Vps45 | 0.69174 | 0.038766 |
| Vps52 | -0.80966 | 0.0084114 |
| Vta1 | 0.76697 | 0.014291 |
| Vti1a | -0.75402 | 0.039482 |
| Vwa3a | -1.1784 | 0.018802 |
| Vwa5b1 | -1.6485 | 0.013324 |
| Vwa5b2 | -1.0747 | 0.0029239 |
| Wasf2 | -1.7803 | 0.0035976 |
| Wasf3 | -1.1181 | 0.003378 |
| Wbp2 | -1.1644 | 3.64E-05 |
| Wbp5 | 0.93519 | 0.0015494 |
| Wbscr17 | -1.766 | 1.37E-09 |
| Wdhd1 | -1.4004 | 0.04695 |
| Wdr53 | 0.93784 | 0.02405 |
| Wdr59 | -1.2479 | 2.09E-04 |
| Wdr61 | 0.67614 | 0.043355 |
| Wdr7 | -1.1668 | 9.54E-05 |
| Wdr82 | -0.91058 | 0.0038824 |
| Wdr83os | 0.86121 | 0.0047957 |
| Wdr90 | -1.694 | 3.22E-08 |
| Wdtc1 | -1.3995 | 1.41E-06 |
| Whsc1l1 | -0.88377 | 0.0083153 |
| Wipf2 | -2.3004 | 8.88E-14 |
| Wipf3 | -0.90463 | 0.0028787 |
| Wiz | -1.0016 | 8.21E-04 |
| Wnk3 | -1.5922 | 0.0054289 |
| Wnt7b | -1.4528 | 1.19E-06 |
| Wwc3 | -0.98805 | 0.0077118 |
| Xkr4 | -1.9123 | 2.08E-09 |
| Xkr6 | -1.9868 | 4.02E-07 |
| Xkr7 | -1.6508 | 6.30E-05 |
| Xpot | -0.7917 | 0.014827 |

| | | |
|---|---|---|
| Xpr1 | -1.4674 | 1.25E-06 |
| Xrcc1 | -0.74029 | 0.034096 |
| Xrn1 | -1.1826 | 0.0063848 |
| Yae1d1 | 0.7784 | 0.044273 |
| Yaf2 | 0.79331 | 0.0093987 |
| Yap1 | -1.3501 | 3.14E-04 |
| Ybx1-ps3 | 0.75625 | 0.014409 |
| Ybx2 | -1.2072 | 0.0019295 |
| Yipf4 | 0.69806 | 0.027923 |
| Yipf5 | 0.66255 | 0.041906 |
| Ypel2 | -0.80924 | 0.016221 |
| Ypel5 | 0.86457 | 0.0034193 |
| Zak | -1.3758 | 0.0021422 |
| Zbtb16 | -1.6765 | 9.09E-06 |
| Zbtb37 | -2.5965 | 6.96E-07 |
| Zbtb4 | -0.68244 | 0.038207 |
| Zbtb40 | -1.5165 | 0.0021267 |
| Zbtb46 | -0.81358 | 0.028574 |
| Zbtb47 | -0.96866 | 0.0014696 |
| Zbtb7a | -0.86087 | 0.0075864 |
| Zbtb8os | 1.0638 | 0.0013551 |
| Zc3h12b | -1.33 | 0.0029239 |
| Zc3h12c | -1.2941 | 0.01264 |
| Zcchc10 | 1.1365 | 0.0040528 |
| Zcchc17 | 0.76054 | 0.014346 |
| Zcrb1 | 0.79987 | 0.013232 |
| Zdbf2 | -1.4546 | 2.16E-05 |
| Zdhhc2 | 1.3115 | 0.01303 |
| Zdhhc21 | -1.786 | 1.75E-06 |
| Zdhhc22 | -0.78344 | 0.047349 |
| Zdhhc8 | -1.1323 | 1.38E-04 |
| Zfand6 | 0.85993 | 0.0064756 |
| Zfhx3 | -2.573 | 1.79E-06 |
| Zfp169 | -1.0419 | 0.023847 |
| Zfp180 | 0.93917 | 0.001822 |
| Zfp236 | -1.4794 | 0.0010531 |
| Zfp260 | 0.85129 | 0.0071157 |
| Zfp319 | -1.8027 | 1.42E-07 |
| Zfp335 | -0.89279 | 0.0040647 |
| Zfp341 | -1.0662 | 0.030804 |
| Zfp347 | 0.80713 | 0.049016 |
| Zfp367 | 0.78382 | 0.019614 |
| Zfp395 | -1.1775 | 0.018652 |
| Zfp407 | -0.98532 | 0.020583 |

| | | |
|---|---|---|
| Zfp426 | 1.01 | 0.0039499 |
| Zfp444 | -0.69478 | 0.042249 |
| Zfp462 | -1.3458 | 3.55E-05 |
| Zfp496 | -0.91282 | 0.016689 |
| Zfp512b | -1.032 | 3.85E-04 |
| Zfp516 | -1.2928 | 0.0096031 |
| Zfp526 | -1.4065 | 0.0011794 |
| Zfp560 | 0.91669 | 0.0050657 |
| Zfp593 | 0.90234 | 0.033628 |
| Zfp597 | -0.95945 | 0.030272 |
| Zfp598 | -0.94818 | 0.0019822 |
| Zfp609 | -0.91803 | 0.0025799 |
| Zfp618 | -1.2246 | 0.043306 |
| Zfp637 | 1.3091 | 0.0084447 |
| Zfp652 | -1.5886 | 4.80E-06 |
| Zfp68 | 0.64649 | 0.049197 |
| Zfp692 | -0.74708 | 0.023975 |
| Zfp710 | -0.97725 | 0.022494 |
| Zfp827 | -1.655 | 2.99E-07 |
| Zfp831 | -2.3541 | 9.43E-04 |
| Zfp871 | -1.5574 | 0.0056641 |
| Zfp879 | 1.1031 | 0.015154 |
| Zfp954 | 0.92076 | 0.013912 |
| Zfr2 | -0.72465 | 0.019719 |
| Zgpat | -0.94678 | 0.0021986 |
| Zim1 | -1.7246 | 0.038282 |
| Zkscan1 | -0.77721 | 0.038214 |
| Zmat2 | 0.82736 | 0.0070521 |
| Zmat3 | -1.0803 | 3.05E-04 |
| Zmynd8 | -0.8188 | 0.0092668 |
| Znrd1 | 0.81676 | 0.040655 |
| Znrf4 | -5.3302 | 0.0022214 |
| Zswim5 | -1.3834 | 6.06E-05 |
| Zswim6 | -0.91232 | 0.013133 |
| Zswim8 | -1.0335 | 3.49E-04 |
| Zzef1 | -1.2892 | 2.13E-05 |

**Pathway identifier**
R-HSA-6794362
R-HSA-6794361
R-HSA-451308
R-HSA-451306
R-HSA-8849932
R-HSA-388844
R-HSA-9630794
R-HSA-5339700
R-HSA-9632700
R-HSA-451326
R-HSA-5628897
R-HSA-8943723
R-HSA-9008059
R-HSA-447038
R-HSA-391160
R-HSA-209563
R-HSA-112316
R-HSA-5682910
R-HSA-196025
R-HSA-68616
R-HSA-8943724
R-HSA-210746
R-HSA-380320
R-HSA-190873
R-HSA-8951671
R-HSA-6791461
R-HSA-5659996
R-HSA-8854518
R-HSA-77595
R-HSA-169131
R-HSA-8941856
R-HSA-9630747
R-HSA-9632693
R-HSA-9630750
R-HSA-6791465
R-HSA-5250982
R-HSA-8939243
R-HSA-5250981
R-HSA-5250955
R-HSA-3371497
R-HSA-9617324
R-HSA-2644605
R-HSA-2644607
R-HSA-438066
R-HSA-193697
R-HSA-193634
R-HSA-5250958
R-HSA-380270
R-HSA-187706

R-HSA-428643
R-HSA-5674404
R-HSA-75067
R-HSA-112412
R-HSA-446205
R-HSA-2660826
R-HSA-2660825
R-HSA-8957275
R-HSA-8865999
R-HSA-9645135
R-HSA-72731
R-HSA-9032759
R-HSA-6785631
R-HSA-111457
R-HSA-6811436
R-HSA-165181
R-HSA-428540
R-HSA-8875555
R-HSA-5626978
R-HSA-210744
R-HSA-5250989
R-HSA-2691230
R-HSA-2691232
R-HSA-2132295
R-HSA-1296072
R-HSA-5626467
R-HSA-1963640
R-HSA-380259
R-HSA-380284
R-HSA-1368071
R-HSA-392851
R-HSA-2151209
R-HSA-549127
R-HSA-196843
R-HSA-194840
R-HSA-442982
R-HSA-8955332
R-HSA-2485179
R-HSA-210500
R-HSA-428930
R-HSA-111463
R-HSA-5620916
R-HSA-9634635
R-HSA-9022707
R-HSA-380287
R-HSA-8854521
R-HSA-111469
R-HSA-69560
R-HSA-69895
R-HSA-8934593

R-HSA-202040
R-HSA-9017802
R-HSA-5620912
R-HSA-6806942
R-HSA-400042
R-HSA-139910
R-HSA-179812
R-HSA-199418
R-HSA-110312
R-HSA-1169092
R-HSA-8941855
R-HSA-73772
R-HSA-392170
R-HSA-8875360
R-HSA-549132
R-HSA-113501
R-HSA-170984
R-HSA-1483255
R-HSA-111446
R-HSA-9034864
R-HSA-73856
R-HSA-9026519
R-HSA-9619229
R-HSA-9032845
R-HSA-2428933
R-HSA-392154
R-HSA-75892
R-HSA-114508
R-HSA-171007
R-HSA-168799
R-HSA-1253288
R-HSA-171319
R-HSA-389948
R-HSA-9619483
R-HSA-113510
R-HSA-446343
R-HSA-947581
R-HSA-9032500
R-HSA-71064
R-HSA-6787639
R-HSA-177504
R-HSA-416572
R-HSA-72695
R-HSA-5654704
R-HSA-167044
R-HSA-168276
R-HSA-5654719
R-HSA-429947
R-HSA-2565942
R-HSA-389960

R-HSA-380972
R-HSA-111459
R-HSA-190704
R-HSA-5654688
R-HSA-399955
R-HSA-210991
R-HSA-400685
R-HSA-5467340
R-HSA-8948747
R-HSA-4839735
R-HSA-5654699
R-HSA-389513
R-HSA-6807878
R-HSA-446388
R-HSA-1170546
R-HSA-3928663
R-HSA-5689901
R-HSA-68962
R-HSA-6807004
R-HSA-68867
R-HSA-1474151
R-HSA-2122948
R-HSA-72649
R-HSA-399956
R-HSA-2197563
R-HSA-380108
R-HSA-72187
R-HSA-8964539
R-HSA-112411
R-HSA-877312
R-HSA-8964616
R-HSA-8876493
R-HSA-110056
R-HSA-8983711
R-HSA-1257604
R-HSA-428542
R-HSA-5620922
R-HSA-170968
R-HSA-168253
R-HSA-71336
R-HSA-381426
R-HSA-6802952
R-HSA-169893
R-HSA-1660499
R-HSA-171286
R-HSA-180336
R-HSA-9027284
R-HSA-163359
R-HSA-1250347
R-HSA-72086

R-HSA-9013508
R-HSA-446652
R-HSA-418592
R-HSA-73817
R-HSA-6804116
R-HSA-6811438
R-HSA-71403
R-HSA-381183
R-HSA-5625900
R-HSA-165054
R-HSA-1632852
R-HSA-912694
R-HSA-380994
R-HSA-205043
R-HSA-381033
R-HSA-2219528
R-HSA-442742
R-HSA-434316
R-HSA-9620244
R-HSA-381676
R-HSA-8951936
R-HSA-389661
R-HSA-2559584
R-HSA-427389
R-HSA-456926
R-HSA-2151201
R-HSA-8851805
R-HSA-5205685
R-HSA-114452
R-HSA-9020558
R-HSA-8856828
R-HSA-68877
R-HSA-3214841
R-HSA-1362277
R-HSA-1980145
R-HSA-8956320
R-HSA-3134973
R-HSA-8876384
R-HSA-392518
R-HSA-1250196
R-HSA-69273
R-HSA-8856688
R-HSA-187687
R-HSA-190861
R-HSA-9013695
R-HSA-5467337
R-HSA-9609736
R-HSA-4839744
R-HSA-429914
R-HSA-432040

R-HSA-8854214
R-HSA-418886
R-HSA-9627069
R-HSA-156902
R-HSA-8874081
R-HSA-5663084
R-HSA-3928665
R-HSA-392517
R-HSA-111461
R-HSA-1482788
R-HSA-111458
R-HSA-936964
R-HSA-418597
R-HSA-2514859
R-HSA-8978934
R-HSA-6811558
R-HSA-210990
R-HSA-432142
R-HSA-351906
R-HSA-389958
R-HSA-5389840
R-HSA-9022699
R-HSA-5467348
R-HSA-5419276
R-HSA-201688
R-HSA-4839748
R-HSA-1433559
R-HSA-1660516
R-HSA-6802957
R-HSA-1483248
R-HSA-9615017
R-HSA-435368
R-HSA-192823
R-HSA-1660517
R-HSA-416482
R-HSA-180746
R-HSA-1855204
R-HSA-2514856
R-HSA-9006925
R-HSA-70263
R-HSA-6782315
R-HSA-6807070
R-HSA-69236
R-HSA-69231
R-HSA-72165
R-HSA-180786
R-HSA-445717
R-HSA-5620924
R-HSA-190828
R-HSA-5624958

R-HSA-5336415
R-HSA-75815
R-HSA-8851907
R-HSA-166208
R-HSA-1059683
R-HSA-399997
R-HSA-1989781
R-HSA-9027276
R-HSA-381042
R-HSA-400451
R-HSA-8863795
R-HSA-8849471
R-HSA-1296071
R-HSA-8951430
R-HSA-163560
R-HSA-6811442
R-HSA-389359
R-HSA-112409
R-HSA-6783310
R-HSA-111471
R-HSA-381771
R-HSA-159227
R-HSA-427359
R-HSA-110313
R-HSA-400206
R-HSA-350054
R-HSA-525793
R-HSA-69002
R-HSA-8849468
R-HSA-8866376
R-HSA-72706
R-HSA-175474
R-HSA-209543
R-HSA-445355
R-HSA-9022692
R-HSA-8985801
R-HSA-198203
R-HSA-8853659
R-HSA-159231
R-HSA-72662
R-HSA-2586552
R-HSA-2559582
R-HSA-187577
R-HSA-112310
R-HSA-8853334
R-HSA-156842
R-HSA-8875878
R-HSA-8856825
R-HSA-73893
R-HSA-5638303

R-HSA-5638302
R-HSA-177243
R-HSA-435354
R-HSA-400508
R-HSA-5693616
R-HSA-2559580
R-HSA-141444
R-HSA-141424
R-HSA-6807062
R-HSA-6807047
R-HSA-373753
R-HSA-5689603
R-HSA-4411364
R-HSA-437239
R-HSA-8876198
R-HSA-8866911
R-HSA-8979227
R-HSA-159230
R-HSA-6783589
R-HSA-1168372
R-HSA-8876725
R-HSA-5693579
R-HSA-5205647
R-HSA-1660514
R-HSA-446353
R-HSA-9013507
R-HSA-157858
R-HSA-5610783
R-HSA-8854050
R-HSA-198323
R-HSA-5610785
R-HSA-210993
R-HSA-159234
R-HSA-392451
R-HSA-71406
R-HSA-4791275
R-HSA-9614657
R-HSA-1834941
R-HSA-450513
R-HSA-2426168
R-HSA-5368287
R-HSA-210745
R-HSA-201722
R-HSA-9603381
R-HSA-425410
R-HSA-193639
R-HSA-156711
R-HSA-264642
R-HSA-181429
R-HSA-450408

R-HSA-6804754
R-HSA-2179392
R-HSA-180292
R-HSA-1266695
R-HSA-9617828
R-HSA-164944
R-HSA-69541
R-HSA-390466
R-HSA-2559585
R-HSA-109704
R-HSA-69618
R-HSA-181430
R-HSA-6804114
R-HSA-844456
R-HSA-5655291
R-HSA-199977
R-HSA-4641262
R-HSA-2173795
R-HSA-8878171
R-HSA-180534
R-HSA-1295596
R-HSA-203615
R-HSA-2467813
R-HSA-157579
R-HSA-1483249
R-HSA-212676
R-HSA-2299718
R-HSA-5621575
R-HSA-622312
R-HSA-9028731
R-HSA-8852276
R-HSA-9006335
R-HSA-8941858
R-HSA-9613354
R-HSA-5654710
R-HSA-5654720
R-HSA-975957
R-HSA-927802
R-HSA-9634638
R-HSA-391251
R-HSA-5663220
R-HSA-2500257
R-HSA-5654689
R-HSA-381038
R-HSA-3232142
R-HSA-1445148
R-HSA-5654732
R-HSA-983170
R-HSA-399719
R-HSA-5654695

R-HSA-450385
R-HSA-5654733
R-HSA-373755
R-HSA-1592230
R-HSA-5654706
R-HSA-180024
R-HSA-5654712
R-HSA-2219530
R-HSA-68882
R-HSA-73762
R-HSA-8854691
R-HSA-399721
R-HSA-159236
R-HSA-450604
R-HSA-6811434
R-HSA-5218920
R-HSA-70326
R-HSA-5654693
R-HSA-418885
R-HSA-5654727
R-HSA-5654726
R-HSA-112399
R-HSA-5655332
R-HSA-8853338
R-HSA-5654700
R-HSA-1655829
R-HSA-9615710
R-HSA-4615885
R-HSA-5625886
R-HSA-5668599
R-HSA-2028269
R-HSA-73886
R-HSA-4641265
R-HSA-888590
R-HSA-2428928
R-HSA-2730905
R-HSA-5250924
R-HSA-69306
R-HSA-8986944
R-HSA-2428924
R-HSA-4086398
R-HSA-2404192
R-HSA-381070
R-HSA-375165
R-HSA-446219
R-HSA-186712
R-HSA-204005
R-HSA-6807505
R-HSA-948021
R-HSA-1912408

R-HSA-5655253
R-HSA-5684996
R-HSA-381340
R-HSA-72737
R-HSA-69580
R-HSA-69563
R-HSA-72613
R-HSA-168273
R-HSA-9007101
R-HSA-445144
R-HSA-3928662
R-HSA-8940973
R-HSA-8939902
R-HSA-264876
R-HSA-2424491
R-HSA-9010553
R-HSA-3299685
R-HSA-1483257
R-HSA-114604
R-HSA-164952
R-HSA-69615
R-HSA-881907
R-HSA-418346
R-HSA-193648
R-HSA-5357956
R-HSA-202131
R-HSA-3371453
R-HSA-69656
R-HSA-9012852
R-HSA-397795
R-HSA-72306
R-HSA-193704
R-HSA-5637815
R-HSA-1236382
R-HSA-74751
R-HSA-1538133
R-HSA-432722
R-HSA-422356
R-HSA-2172127
R-HSA-425393
R-HSA-5625740
R-HSA-9013694
R-HSA-446193
R-HSA-1852241
R-HSA-977225
R-HSA-73854
R-HSA-6811440
R-HSA-2187338
R-HSA-5620920
R-HSA-202433

R-HSA-68875
R-HSA-182971
R-HSA-9034015
R-HSA-5655302
R-HSA-438064
R-HSA-109606
R-HSA-389356
R-HSA-936440
R-HSA-2559586
R-HSA-400253
R-HSA-9612973
R-HSA-5250941
R-HSA-2644603
R-HSA-2644606
R-HSA-2644602
R-HSA-2894858
R-HSA-2894862
R-HSA-8941326
R-HSA-1912422
R-HSA-983695
R-HSA-74752
R-HSA-3371556
R-HSA-5668914
R-HSA-1433557
R-HSA-2555396
R-HSA-1500931
R-HSA-5610787
R-HSA-3214858
R-HSA-73864
R-HSA-177929
R-HSA-9613829
R-HSA-5617833
R-HSA-112315
R-HSA-191273
R-HSA-5685942
R-HSA-69202
R-HSA-2122947
R-HSA-5250913
R-HSA-8857538
R-HSA-5674400
R-HSA-8983432
R-HSA-186763
R-HSA-512988
R-HSA-5339562
R-HSA-1226099
R-HSA-68886
R-HSA-6811555
R-HSA-112314
R-HSA-1169408
R-HSA-917937

R-HSA-983168
R-HSA-8936459
R-HSA-15869
R-HSA-4420097
R-HSA-72202
R-HSA-354192
R-HSA-9006921
R-HSA-381119
R-HSA-69275
R-HSA-5689896
R-HSA-453274
R-HSA-163685
R-HSA-9614085
R-HSA-204998
R-HSA-202424
R-HSA-9006115
R-HSA-442755
R-HSA-76009
R-HSA-194138
R-HSA-216083
R-HSA-3247509
R-HSA-4839726
R-HSA-8953750
R-HSA-447043
R-HSA-186797
R-HSA-5689880
R-HSA-165159
R-HSA-3108214
R-HSA-5663202
R-HSA-2871796
R-HSA-194315
R-HSA-3232118
R-HSA-1980143
R-HSA-6804758
R-HSA-446728
R-HSA-168643
R-HSA-8950505
R-HSA-5637812
R-HSA-5637810
R-HSA-9607240
R-HSA-196849
R-HSA-388841
R-HSA-416993
R-HSA-6804757
R-HSA-9616222
R-HSA-2029482
R-HSA-909733
R-HSA-168255
R-HSA-5654708
R-HSA-1169410

R-HSA-6791312
R-HSA-5654716
R-HSA-1834949
R-HSA-5673001
R-HSA-2173796
R-HSA-1643713
R-HSA-375276
R-HSA-397014
R-HSA-5654696
R-HSA-5654687
R-HSA-1227986
R-HSA-166016
R-HSA-1428517
R-HSA-451927
R-HSA-2559583
R-HSA-111465
R-HSA-9009391
R-HSA-8866910
R-HSA-877300
R-HSA-201681
R-HSA-3899300
R-HSA-1483206
R-HSA-166520
R-HSA-168254
R-HSA-162599
R-HSA-3769402
R-HSA-5687128
R-HSA-8948751
R-HSA-446203
R-HSA-69242
R-HSA-199991
R-HSA-983705
R-HSA-69206
R-HSA-6806834
R-HSA-162587
R-HSA-9020591
R-HSA-168898
R-HSA-187037
R-HSA-168928
R-HSA-6798695
R-HSA-75153
R-HSA-2029480
R-HSA-1236394
R-HSA-8878159
R-HSA-373752
R-HSA-5683057
R-HSA-72163
R-HSA-195253
R-HSA-937061
R-HSA-166166

R-HSA-3858494
R-HSA-111885
R-HSA-425366
R-HSA-4090294
R-HSA-72172
R-HSA-5654743
R-HSA-373760
R-HSA-202403
R-HSA-5654741
R-HSA-8953897
R-HSA-5688426
R-HSA-2262752
R-HSA-432720
R-HSA-6791226
R-HSA-5654736
R-HSA-5693567
R-HSA-4551638
R-HSA-2173793
R-HSA-5693538
R-HSA-2980736
R-HSA-6806003
R-HSA-450531
R-HSA-416476
R-HSA-69620
R-HSA-212165
R-HSA-3928664
R-HSA-5358351
R-HSA-983231
R-HSA-376176
R-HSA-5653656
R-HSA-76002
R-HSA-8868773
R-HSA-202733
R-HSA-5654738
R-HSA-453279
R-HSA-196854
R-HSA-5357905
R-HSA-190236
R-HSA-8878166
R-HSA-198933
R-HSA-449836
R-HSA-195258
R-HSA-2454202
R-HSA-9006934
R-HSA-913531
R-HSA-5621481
R-HSA-72203
R-HSA-72312
R-HSA-447115
R-HSA-71387

R-HSA-1280218
R-HSA-983169
R-HSA-195721
R-HSA-72766
R-HSA-199992
R-HSA-162909
R-HSA-170834
R-HSA-75893
R-HSA-5693532
R-HSA-418555
R-HSA-5619507
R-HSA-5617472
R-HSA-8848021
R-HSA-9006927
R-HSA-69278
R-HSA-2682334
R-HSA-9006936
R-HSA-73887
R-HSA-373076
R-HSA-1474244
R-HSA-8864260
R-HSA-8863678
R-HSA-8862803
R-HSA-425407
R-HSA-3700989
R-HSA-449147
R-HSA-157118
R-HSA-109581
R-HSA-73857
R-HSA-5357801
R-HSA-71291
R-HSA-9018519
R-HSA-162906
R-HSA-8957322
R-HSA-500792
R-HSA-212436
R-HSA-418594
R-HSA-74160
R-HSA-5633007
R-HSA-109582
R-HSA-73894
R-HSA-1640170
R-HSA-422475
R-HSA-8939211
R-HSA-1280215
R-HSA-556833
R-HSA-9006931
R-HSA-597592
R-HSA-5663205
R-HSA-3108232

R-HSA-2990846
R-HSA-388396
R-HSA-168249
R-HSA-8953854
R-HSA-168256
R-HSA-382551
R-HSA-1266738
R-HSA-392499
R-HSA-372790
R-HSA-1643685
R-HSA-162582
R-HSA-1430728

| **Pathway name** |
|---|
| Protein-protein interactions at synapses |
| Neurexins and neuroligins |
| Activation of Ca-permeable Kainate Receptor |
| Ionotropic activity of kainate receptors |
| Synaptic adhesion-like molecules |
| Receptor-type tyrosine-protein phosphatases |
| Evasion of Oncogene Induced Senescence Due to Defective p16INK4A binding to CDK4 and CDK6 |
| TCF7L2 mutants don't bind CTBP |
| Evasion of Oxidative Stress Induced Senescence Due to Defective p16INK4A binding to CDK4 and CDK6 |
| Activation of kainate receptors upon glutamate binding |
| TP53 Regulates Metabolic Genes |
| Regulation of PTEN mRNA translation |
| Interleukin-37 signaling |
| NrCAM interactions |
| Signal regulatory protein family interactions |
| Axonal growth stimulation |
| Neuronal System |
| LGI-ADAM interactions |
| Formation of annular gap junctions |
| Assembly of the ORC complex at the origin of replication |
| Regulation of PTEN gene transcription |
| Regulation of gene expression in endocrine-committed (NEUROG3+) progenitor cells |
| Recruitment of NuMA to mitotic centrosomes |
| Gap junction degradation |
| RUNX3 regulates YAP1-mediated transcription |
| RPIA deficiency: failed conversion of RU5P to R5P |
| RPIA deficiency: failed conversion of R5P to RU5P |
| AURKA Activation by TPX2 |
| Processing of Intronless Pre-mRNAs |
| Inhibition of PKR |
| RUNX3 regulates NOTCH signaling |
| Diseases of Cellular Senescence |
| Evasion of Oxidative Stress Induced Senescence Due to p16INK4A Defects |
| Evasion of Oncogene Induced Senescence Due to p16INK4A Defects |
| Pentose phosphate pathway disease |
| Toxicity of tetanus toxin (TeNT) |
| RUNX1 interacts with co-factors whose precise effect on RUNX1 targets is not known |
| Toxicity of botulinum toxin type F (BoNT/F) |
| Toxicity of botulinum toxin type D (BoNT/D) |
| HSP90 chaperone cycle for steroid hormone receptors (SHR) |
| Negative regulation of NMDA receptor-mediated neuronal transmission |
| FBXW7 Mutants and NOTCH1 in Cancer |
| Loss of Function of FBXW7 in Cancer and NOTCH1 Signaling |
| Unblocking of NMDA receptors, glutamate binding and activation |
| p75NTR regulates axonogenesis |
| Axonal growth inhibition (RHOA activation) |
| Toxicity of botulinum toxin type B (BoNT/B) |
| Recruitment of mitotic centrosome proteins and complexes |
| Signalling to p38 via RIT and RIN |

Organic anion transporters
PTEN Loss of Function in Cancer
Processing of Capped Intronless Pre-mRNA
SOS-mediated signalling
Synthesis of GDP-mannose
Constitutive Signaling by NOTCH1 t(7;9)(NOTCH1:M1580_K2555) Translocation Mutant
Signaling by NOTCH1 t(7;9)(NOTCH1:M1580_K2555) Translocation Mutant
Post-translational protein phosphorylation
MET activates PTPN11
STAT5 Activation
Recycling of eIF2:GDP
NTRK2 activates RAC1
ERBB2 Regulates Cell Motility
Release of apoptotic factors from the mitochondria
COPI-independent Golgi-to-ER retrograde traffic
Inhibition of TSC complex formation by PKB
Activation of RAC1
MET activates RAP1 and RAC1
TNFR1-mediated ceramide production
Regulation of gene expression in late stage (branching morphogenesis) pancreatic bud precursor cells
Toxicity of botulinum toxin type G (BoNT/G)
Signaling by NOTCH1 HD Domain Mutants in Cancer
Constitutive Signaling by NOTCH1 HD Domain Mutants
MHC class II antigen presentation
Voltage gated Potassium channels
RHO GTPases activate IQGAPs
GRB2 events in ERBB2 signaling
Loss of Nlp from mitotic centrosomes
Loss of proteins required for interphase microtubule organization from the centrosome
NR1D1 (REV-ERBA) represses gene expression
Prostacyclin signalling through prostacyclin receptor
Activation of PPARGC1A (PGC-1alpha) by phosphorylation
Organic cation transport
Vitamin B2 (riboflavin) metabolism
Rho GTPase cycle
Ras activation upon Ca2+ influx through NMDA receptor
Carboxyterminal post-translational modifications of tubulin
Activation of the phototransduction cascade
Glutamate Neurotransmitter Release Cycle
Thromboxane signalling through TP receptor
SMAC (DIABLO) binds to IAPs
VxPx cargo-targeting to cilium
Estrogen-stimulated signaling through PRKCZ
MECP2 regulates transcription factors
Centrosome maturation
Interaction between PHLDA1 and AURKA
SMAC, XIAP-regulated apoptotic response
Transcriptional activation of p53 responsive genes
Transcriptional activation of cell cycle inhibitor p21
Regulation of RUNX1 Expression and Activity

G-protein activation
Noncanonical activation of NOTCH3
Anchoring of the basal body to the plasma membrane
MET Receptor Activation
Adrenaline,noradrenaline inhibits insulin secretion
Activation of BMF and translocation to mitochondria
GRB2 events in EGFR signaling
Negative regulation of the PI3K/AKT network
Translesion synthesis by REV1
Activation of RAS in B cells
RUNX3 regulates CDKN1A transcription
RNA Polymerase I Promoter Escape
ADP signalling through P2Y purinoceptor 12
InlB-mediated entry of Listeria monocytogenes into host cell
Organic cation/anion/zwitterion transport
Inhibition of replication initiation of damaged DNA by RB1/E2F1
ARMS-mediated activation
PI Metabolism
Activation of BIM and translocation to mitochondria
Activated NTRK3 signals through RAS
RNA Polymerase II Transcription Termination
Activated NTRK2 signals through RAS
Activation of RAC1 downstream of NMDARs
Activated NTRK2 signals through CDK5
SHC-related events triggered by IGF1R
Nitric oxide stimulates guanylate cyclase
Platelet Adhesion to exposed collagen
Effects of PIP2 hydrolysis
p38MAPK events
Neurotoxicity of clostridium toxins
Downregulation of ERBB4 signaling
Telomere Extension By Telomerase
PD-1 signaling
Activation of AMPK downstream of NMDARs
E2F mediated regulation of DNA replication
Localization of the PINCH-ILK-PARVIN complex to focal adhesions
Molybdenum cofactor biosynthesis
Activated NTRK2 signals through FYN
Lysine catabolism
GDP-fucose biosynthesis
Retrograde neurotrophin signalling
Sema4D induced cell migration and growth-cone collapse
Formation of the ternary complex, and subsequently, the 43S complex
SHC-mediated cascade:FGFR3
Signalling to RAS
NS1 Mediated Effects on Host Pathways
SHC-mediated cascade:FGFR4
Deadenylation of mRNA
Regulation of PLK1 Activity at G2/M Transition
Formation of tubulin folding intermediates by CCT/TriC

Energy dependent regulation of mTOR by LKB1-AMPK
Activation of caspases through apoptosome-mediated cleavage
Oligomerization of connexins into connexons
SHC-mediated cascade:FGFR1
SEMA3A-Plexin repulsion signaling by inhibiting Integrin adhesion
Basigin interactions
Sema4D in semaphorin signaling
AXIN missense mutants destabilize the destruction complex
Regulation of PTEN localization
AXIN mutants destabilize the destruction complex, activating WNT signaling
SHC-mediated cascade:FGFR2
CTLA4 inhibitory signaling
COPI-mediated anterograde transport
Regulation of cytoskeletal remodeling and cell spreading by IPP complex components
Prolactin receptor signaling
EPHA-mediated growth cone collapse
Metalloprotease DUBs
Activation of the pre-replicative complex
Negative regulation of MET activity
Assembly of the pre-replicative complex
Tetrahydrobiopterin (BH4) synthesis, recycling, salvage and regulation
Activated NOTCH1 Transmits Signal to the Nucleus
Translation initiation complex formation
CRMPs in Sema3A signaling
NOTCH2 intracellular domain regulates transcription
Chemokine receptors bind chemokines
mRNA 3'-end processing
Glutamate and glutamine metabolism
MAPK1 (ERK2) activation
Regulation of IFNG signaling
G beta:gamma signalling through CDC42
InlA-mediated entry of Listeria monocytogenes into host cells
MAPK3 (ERK1) activation
OAS antiviral response
PIP3 activates AKT signaling
Regulation of commissural axon pathfinding by SLIT and ROBO
BBSome-mediated cargo-targeting to cilium
Frs2-mediated activation
Host Interactions with Influenza Factors
Pentose phosphate pathway
Regulation of Insulin-like Growth Factor (IGF) transport and uptake by Insulin-like Growth Factor Binding Proteins (IGF
Signaling by BRAF and RAF fusions
Prolonged ERK activation events
Synthesis of PIPs at the plasma membrane
Synthesis and processing of ENV and VPU
SHC1 events in EGFR signaling
Erythropoietin activates RAS
Glucagon signaling in metabolic regulation
SHC1 events in ERBB4 signaling
mRNA Capping

NOTCH3 Intracellular Domain Regulates Transcription
Interleukin-1 family signaling
ADP signalling through P2Y purinoceptor 1
Purine ribonucleoside monophosphate biosynthesis
TP53 Regulates Transcription of Genes Involved in G1 Cell Cycle Arrest
Intra-Golgi traffic
Citric acid cycle (TCA cycle)
ATF6 (ATF6-alpha) activates chaperone genes
RHO GTPases activate CIT
Rev-mediated nuclear export of HIV RNA
Macroautophagy
Regulation of IFNA signaling
ATF4 activates genes in response to endoplasmic reticulum stress
NRIF signals cell death from the nucleus
ATF6 (ATF6-alpha) activates chaperones
PI3K/AKT Signaling in Cancer
CREB1 phosphorylation through NMDA receptor-mediated activation of RAS signaling
Fatty Acids bound to GPR40 (FFAR1) regulate insulin secretion
Long-term potentiation
Glucagon-like Peptide-1 (GLP1) regulates insulin secretion
RUNX3 regulates p14-ARF
Glyoxylate metabolism and glycine degradation
Formation of Senescence-Associated Heterochromatin Foci (SAHF)
ERCC6 (CSB) and EHMT2 (G9a) positively regulate rRNA expression
Thrombin signalling through proteinase activated receptors (PARs)
Transcriptional activation of mitochondrial biogenesis
MET activates RAS signaling
Pink/Parkin Mediated Mitophagy
Activation of BH3-only proteins
Interleukin-2 signaling
Clathrin-mediated endocytosis
Mitotic Prometaphase
PKMTs methylate histone lysines
Transcription of E2F targets under negative control by DREAM complex
Signaling by NOTCH2
Nucleobase biosynthesis
LRR FLII-interacting protein 1 (LRRFIP1) activates type I IFN production
Listeria monocytogenes entry into host cells
Signal amplification
SHC1 events in ERBB2 signaling
Cyclin A/B1/B2 associated events during G2/M transition
Golgi-to-ER retrograde transport
Signalling to ERKs
Gap junction assembly
NOTCH4 Intracellular Domain Regulates Transcription
APC truncation mutants have impaired AXIN binding
Assembly and cell surface presentation of NMDA receptors
truncated APC mutants destabilize the destruction complex
Deadenylation-dependent mRNA decay
Vasopressin regulates renal water homeostasis via Aquaporins

TBC/RABGAPs
Netrin mediated repulsion signals
Regulation of the apoptosome activity
Peptide chain elongation
MET activates PTK2 signaling
Diseases of carbohydrate metabolism
EPH-ephrin mediated repulsion of cells
Rap1 signalling
Cytochrome c-mediated apoptotic response
Acyl chain remodelling of PC
Formation of apoptosome
Activation of IRF3/IRF7 mediated by TBK1/IKK epsilon
G alpha (z) signalling events
Inactivation, recovery and regulation of the phototransduction cascade
Metabolism of cofactors
PI5P, PP2A and IER3 Regulate PI3K/AKT Signaling
PECAM1 interactions
Platelet sensitization by LDL
Apoptotic cleavage of cell adhesion proteins
Cooperation of Prefoldin and TriC/CCT in actin and tubulin folding
Mitochondrial translation elongation
MECP2 regulates neuronal receptors and channels
Truncations of AMER1 destabilize the destruction complex
Mitochondrial translation termination
WNT mediated activation of DVL
AMER1 mutants destabilize the destruction complex
Regulation of KIT signaling
Synthesis of PIPs at the early endosome membrane
Oncogenic MAPK signaling
Synthesis of PIPs at the ER membrane
FOXO-mediated transcription of oxidative stress, metabolic and neuronal genes
Zinc efflux and compartmentalization by the SLC30 family
Viral mRNA Translation
Synthesis of PIPs at the late endosome membrane
G alpha (12/13) signalling events
Nuclear import of Rev protein
Synthesis of IP3 and IP4 in the cytosol
The phototransduction cascade
Intracellular signaling by second messengers
Gluconeogenesis
tRNA modification in the nucleus and cytosol
PTEN Regulation
G1 Phase
Cyclin D associated events in G1
mRNA Splicing - Minor Pathway
Extension of Telomeres
Aquaporin-mediated transport
Intraflagellar transport
Gap junction trafficking
ARL13B-mediated ciliary trafficking of INPP5E

Uptake and function of diphtheria toxin
Ubiquitin-dependent degradation of Cyclin D
MET activates PI3K/AKT signaling
mTORC1-mediated signalling
Interleukin-6 signaling
Acetylcholine regulates insulin secretion
PPARA activates gene expression
Erythropoietin activates Phosphoinositide-3-kinase (PI3K)
PERK regulates gene expression
Free fatty acids regulate insulin secretion
Downregulation of ERBB2 signaling
PTK6 Regulates RHO GTPases, RAS GTPase and MAP kinases
Potassium Channels
RUNX3 regulates WNT signaling
Triglyceride catabolism
Intra-Golgi and retrograde Golgi-to-ER traffic
CD28 dependent Vav1 pathway
RAF-independent MAPK1/3 activation
Fanconi Anemia Pathway
Apoptotic factor-mediated response
Synthesis, secretion, and inactivation of Glucagon-like Peptide-1 (GLP-1)
Transport of the SLBP independent Mature mRNA
SIRT1 negatively regulates rRNA expression
Translesion synthesis by Y family DNA polymerases bypasses lesions on DNA template
Regulation of lipid metabolism by Peroxisome proliferator-activated receptor alpha (PPARalpha)
Notch-HLH transcription pathway
Myogenesis
DNA Replication Pre-Initiation
PTK6 Regulates Proteins Involved in RNA Processing
Reelin signalling pathway
GTP hydrolysis and joining of the 60S ribosomal subunit
Assembly Of The HIV Virion
p75NTR recruits signalling complexes
Smooth Muscle Contraction
Regulation of MECP2 expression and activity
Regulation of cortical dendrite branching
PI3K/AKT activation
RET signaling
Transport of Mature mRNA Derived from an Intronless Transcript
Activation of the mRNA upon binding of the cap-binding complex and eIFs, and subsequent binding to 43S
Signaling by Leptin
Senescence-Associated Secretory Phenotype (SASP)
SCF(Skp2)-mediated degradation of p27/p21
Neurotransmitter release cycle
Signaling by FGFR3 fusions in cancer
Eukaryotic Translation Elongation
MET promotes cell motility
Cargo recognition for clathrin-mediated endocytosis
DNA Damage Bypass
Inhibition of Signaling by Overexpressed EGFR

Signaling by Overexpressed Wild-Type EGFR in Cancer
Interactions of Rev with host cellular proteins
Zinc transporters
Incretin synthesis, secretion, and inactivation
Presynaptic phase of homologous DNA pairing and strand exchange
Oxidative Stress Induced Senescence
Amplification of signal from unattached kinetochores via a MAD2 inhibitory signal
Amplification of signal from the kinetochores
Cholesterol biosynthesis via lathosterol
Cholesterol biosynthesis via desmosterol
Nephrin family interactions
UCH proteinases
Binding of TCF/LEF:CTNNB1 to target gene promoters
Recycling pathway of L1
RAB GEFs exchange GTP for GDP on RABs
TFAP2 (AP-2) family regulates transcription of cell cycle factors
Triglyceride metabolism
Transport of the SLBP Dependant Mature mRNA
Interleukin-6 family signaling
Downstream signaling events of B Cell Receptor (BCR)
Protein methylation
Homologous DNA Pairing and Strand Exchange
Mitophagy
Synthesis of PIPs at the Golgi membrane
Cell-extracellular matrix interactions
NOTCH3 Activation and Transmission of Signal to the Nucleus
Gap junction trafficking and regulation
Degradation of GLI2 by the proteasome
FBXL7 down-regulates AURKA during mitotic entry and in early mitosis
AKT phosphorylates targets in the cytosol
GLI3 is processed to GLI3R by the proteasome
Tie2 Signaling
Transport of Mature mRNAs Derived from Intronless Transcripts
G beta:gamma signalling through PI3Kgamma
Pyruvate metabolism and Citric Acid (TCA) cycle
Signaling by WNT in cancer
FOXO-mediated transcription of cell death genes
STING mediated induction of host immune responses
Tristetraprolin (TTP, ZFP36) binds and destabilizes mRNA
Activation of gene expression by SREBF (SREBP)
Mitochondrial translation
Regulation of gene expression in beta cells
Formation of the beta-catenin:TCF transactivating complex
Activated NTRK3 signals through PI3K
Metal ion SLC transporters
p75NTR signals via NF-kB
Polo-like kinase mediated events
Acetylcholine Neurotransmitter Release Cycle
Serotonin Neurotransmitter Release Cycle
AUF1 (hnRNP D0) binds and destabilizes mRNA

Regulation of TP53 Expression
EGFR Transactivation by Gastrin
GAB1 signalosome
Interleukin-7 signaling
FOXO-mediated transcription of cell cycle genes
Nef and signal transduction
Stabilization of p53
Chaperonin-mediated protein folding
Oncogene Induced Senescence
PI3K Cascade
Mitotic Spindle Checkpoint
Norepinephrine Neurotransmitter Release Cycle
TP53 Regulates Transcription of Genes Involved in G2 Cell Cycle Arrest
The NLRP3 inflammasome
Signaling by FGFR4 in disease
ER to Golgi Anterograde Transport
Disassembly of the destruction complex and recruitment of AXIN to the membrane
Downregulation of SMAD2/3:SMAD4 transcriptional activity
Transcriptional regulation by RUNX1
Vpu mediated degradation of CD4
Spry regulation of FGF signaling
eNOS activation
Separation of Sister Chromatids
Telomere Maintenance
Inositol phosphate metabolism
Dopamine Neurotransmitter Release Cycle
Condensation of Prophase Chromosomes
CD209 (DC-SIGN) signaling
Inflammasomes
Activated NTRK2 signals through FRS2 and FRS3
The role of GTSE1 in G2/M progression after G2 checkpoint
Signaling by Erythropoietin
Regulation of RUNX3 expression and activity
Lipophagy
PI-3K cascade:FGFR3
PI-3K cascade:FGFR4
Nonsense Mediated Decay (NMD) enhanced by the Exon Junction Complex (EJC)
Nonsense-Mediated Decay (NMD)
Estrogen-dependent nuclear events downstream of ESR-membrane signaling
Protein folding
RHO GTPases Activate Formins
Resolution of Sister Chromatid Cohesion
PI-3K cascade:FGFR1
XBP1(S) activates chaperone genes
SUMOylation of ubiquitinylation proteins
Translocation of SLC2A4 (GLUT4) to the plasma membrane
Negative regulation of FGFR3 signaling
Antigen Presentation: Folding, assembly and peptide loading of class I MHC
Trafficking of AMPA receptors
PI-3K cascade:FGFR2

Butyrate Response Factor 1 (BRF1) binds and destabilizes mRNA
Negative regulation of FGFR4 signaling
Semaphorin interactions
Mitochondrial biogenesis
FRS-mediated FGFR3 signaling
DARPP-32 events
FRS-mediated FGFR4 signaling
Constitutive Signaling by Aberrant PI3K in Cancer
Mitotic Anaphase
RNA Polymerase I Transcription Initiation
Interleukin-20 family signaling
Glutamate binding, activation of AMPA receptors and synaptic plasticity
Transport of Mature mRNA derived from an Intron-Containing Transcript
KSRP (KHSRP) binds and destabilizes mRNA
COPI-dependent Golgi-to-ER retrograde traffic
VEGFR2 mediated vascular permeability
Glucose metabolism
FRS-mediated FGFR1 signaling
DCC mediated attractive signaling
Negative regulation of FGFR2 signaling
Negative regulation of FGFR1 signaling
IRS-mediated signalling
Signaling by FGFR3 in disease
Signaling by FGFR3 point mutants in cancer
FRS-mediated FGFR2 signaling
Regulation of cholesterol biosynthesis by SREBP (SREBF)
Microautophagy
SUMOylation of DNA replication proteins
Activated PKN1 stimulates transcription of AR (androgen receptor) regulated genes KLK2 and KLK3
RHO GTPases Activate NADPH Oxidases
Signaling by Hippo
Chromosome Maintenance
Repression of WNT target genes
GABA synthesis, release, reuptake and degradation
IRS-related events triggered by IGF1R
Role of LAT2/NTAL/LAB on calcium mobilization
B-WICH complex positively regulates rRNA expression
DNA Replication
Transcriptional Regulation by MECP2
IGF1R signaling cascade
Ca2+ pathway
Signaling by Type 1 Insulin-like Growth Factor 1 Receptor (IGF1R)
IRE1alpha activates chaperones
NCAM signaling for neurite out-growth
Synthesis of substrates in N-glycan biosythesis
Regulation of beta-cell development
COPII-mediated vesicle transport
RNA polymerase II transcribes snRNA genes
Transport to the Golgi and subsequent modification
Pre-NOTCH Transcription and Translation

Signaling by FGFR2 in disease
MAPK1/MAPK3 signaling
Transcriptional regulation of white adipocyte differentiation
Cap-dependent Translation Initiation
p53-Dependent G1/S DNA damage checkpoint
p53-Dependent G1 DNA Damage Response
Eukaryotic Translation Initiation
Influenza Viral RNA Transcription and Replication
Rab regulation of trafficking
Signal transduction by L1
EPHB-mediated forward signaling
RUNX2 regulates osteoblast differentiation
Regulation of RUNX2 expression and activity
Insulin processing
DAP12 signaling
Regulation of expression of SLITs and ROBOs
Detoxification of Reactive Oxygen Species
Phospholipid metabolism
GPVI-mediated activation cascade
The role of Nef in HIV-1 replication and disease pathogenesis
G1/S DNA Damage Checkpoints
Gastrin-CREB signalling pathway via PKC and MAPK
Platelet homeostasis
NRAGE signals death through JNK
TNFR1-induced NFkappaB signaling pathway
Metabolism of nitric oxide: eNOS activation and regulation
Regulation of HSF1-mediated heat shock response
Cyclin A:Cdk2-associated events at S phase entry
Signaling by NOTCH3
G-protein beta:gamma signalling
tRNA processing
p75 NTR receptor-mediated signalling
Signaling by Ligand-Responsive EGFR Variants in Cancer
Constitutive Signaling by Ligand-Responsive EGFR Cancer Variants
Insulin receptor signalling cascade
G0 and Early G1
Golgi Associated Vesicle Biogenesis
Regulation of insulin secretion
DAP12 interactions
Transport of inorganic cations/anions and amino acids/oligopeptides
RHO GTPases activate PKNs
Signaling by NOTCH4
Biosynthesis of the N-glycan precursor (dolichol lipid-linked oligosaccharide, LLO) and transfer to a nascent protein
Organelle biogenesis and maintenance
Amyloid fiber formation
RNA Polymerase I Promoter Clearance
Retrograde transport at the Trans-Golgi-Network
Visual phototransduction
Cargo trafficking to the periciliary membrane
Generation of second messenger molecules

Mitotic Prophase
EGFR downregulation
Signaling by NTRK3 (TRKC)
Signaling by FGFR1 in disease
Post NMDA receptor activation events
Intrinsic Pathway for Apoptosis
CD28 co-stimulation
Negative regulators of DDX58/IFIH1 signaling
DNA Damage/Telomere Stress Induced Senescence
Circadian Clock
Autophagy
Negative epigenetic regulation of rRNA expression
Signaling by NOTCH1 in Cancer
Constitutive Signaling by NOTCH1 PEST Domain Mutants
Signaling by NOTCH1 PEST Domain Mutants in Cancer
Signaling by NOTCH1 HD+PEST Domain Mutants in Cancer
Constitutive Signaling by NOTCH1 HD+PEST Domain Mutants
RUNX2 regulates bone development
Pre-NOTCH Expression and Processing
Antigen activates B Cell Receptor (BCR) leading to generation of second messengers
Signaling by Insulin receptor
Cellular response to heat stress
Diseases of metabolism
Signaling by SCF-KIT
Mitotic Metaphase and Anaphase
Cell-Cell communication
Hedgehog 'off' state
RMTs methylate histone arginines
RNA Polymerase I Transcription
Signaling by EGFR
Chaperone Mediated Autophagy
Cilium Assembly
Transmission across Chemical Synapses
Cholesterol biosynthesis
HDR through Homologous Recombination (HRR)
Cyclin E associated events during G1/S transition
NOTCH1 Intracellular Domain Regulates Transcription
Positive epigenetic regulation of rRNA expression
PTK6 promotes HIF1A stabilization
Constitutive Signaling by AKT1 E17K in Cancer
Interleukin-15 signaling
Downstream signal transduction
Interleukin-3, Interleukin-5 and GM-CSF signaling
Uptake and actions of bacterial toxins
Signaling by FGFR in disease
M Phase
PI5P Regulates TP53 Acetylation
Neurotransmitter receptors and postsynaptic signal transmission
ISG15 antiviral mechanism
Iron uptake and transport

Antigen processing: Ubiquitination & Proteasome degradation
RUNX1 regulates genes involved in megakaryocyte differentiation and platelet function
Metabolism of nucleotides
VEGFA-VEGFR2 Pathway
Transport of Mature Transcript to Cytoplasm
Integrin alphaIIb beta3 signaling
Integrin signaling
Unfolded Protein Response (UPR)
G2/M Transition
Ovarian tumor domain proteases
Mitotic G2-G2/M phases
Integration of energy metabolism
FOXO-mediated transcription
Cell death signalling via NRAGE, NRIF and NADE
Downstream TCR signaling
Signaling by NTRK2 (TRKB)
Activation of NMDA receptors and postsynaptic events
Platelet Aggregation (Plug Formation)
Signaling by VEGF
Integrin cell surface interactions
Chromatin modifying enzymes
Chromatin organization
Transcriptional Regulation by E2F6
Neurofascin interactions
Signaling by PDGF
Ub-specific processing proteases
mTOR signalling
SUMOylation of DNA damage response and repair proteins
Diseases of signal transduction
FCERI mediated MAPK activation
Signaling by Rho GTPases
SUMOylation of transcription factors
Signaling by NOTCH1
Regulation of TP53 Activity through Acetylation
Cell junction organization
Nucleotide-binding domain, leucine rich repeat containing receptor (NLR) signaling pathways
Gene and protein expression by JAK-STAT signaling after Interleukin-12 stimulation
Signaling by EGFRvIII in Cancer
Constitutive Signaling by EGFRvIII
FLT3 Signaling
Metabolism of water-soluble vitamins and cofactors
Costimulation by the CD28 family
Trafficking of GluR2-containing AMPA receptors
Regulation of TP53 Degradation
Transcriptional regulation of granulopoiesis
Regulation of actin dynamics for phagocytic cup formation
Interferon alpha/beta signaling
Influenza Life Cycle
Downstream signaling of activated FGFR3
Antiviral mechanism by IFN-stimulated genes

TP53 Regulates Transcription of Cell Cycle Genes
Downstream signaling of activated FGFR4
Cytosolic sensors of pathogen-associated DNA
RAF/MAP kinase cascade
SMAD2/SMAD3:SMAD4 heterotrimer regulates transcription
Signaling by EGFR in Cancer
Peptide ligand-binding receptors
Muscle contraction
Downstream signaling of activated FGFR2
Downstream signaling of activated FGFR1
Signaling by ERBB2
Toll Like Receptor 4 (TLR4) Cascade
The citric acid (TCA) cycle and respiratory electron transport
Interleukin-2 family signaling
Cellular Senescence
Apoptotic cleavage of cellular proteins
Extra-nuclear estrogen signaling
TFAP2 (AP-2) family regulates transcription of growth factors and their receptors
Interferon gamma signaling
TCF dependent signaling in response to WNT
SUMOylation of transcription cofactors
Glycerophospholipid biosynthesis
Signaling by NTRKs
Influenza Infection
Late Phase of HIV Life Cycle
Deactivation of the beta-catenin transactivating complex
MAPK6/MAPK4 signaling
Regulation of PTEN stability and activity
Asparagine N-linked glycosylation
S Phase
Membrane Trafficking
Signaling by the B Cell Receptor (BCR)
G1/S Transition
Signaling by MET
HIV Life Cycle
Interleukin-12 signaling
Toll-like Receptor Cascades
Signaling by NTRK1 (TRKA)
DDX58/IFIH1-mediated induction of interferon-alpha/beta
Neutrophil degranulation
Apoptotic execution phase
Fcgamma receptor (FCGR) dependent phagocytosis
Signaling by ERBB4
Transcriptional regulation by RUNX3
Netrin-1 signaling
MAPK family signaling cascades
mRNA Splicing - Major Pathway
Degradation of beta-catenin by the destruction complex
TRIF(TICAM1)-mediated TLR4 signaling
MyD88-independent TLR4 cascade

Beta-catenin independent WNT signaling
Opioid Signalling
Transport of bile salts and organic acids, metal ions and amine compounds
SUMOylation of intracellular receptors
mRNA Splicing
Signaling by FGFR4
L1CAM interactions
TCR signaling
Signaling by FGFR3
Cellular responses to external stimuli
Deubiquitination
Cellular responses to stress
Lysosome Vesicle Biogenesis
Major pathway of rRNA processing in the nucleolus and cytosol
Signaling by FGFR1
HDR through Homologous Recombination (HRR) or Single Strand Annealing (SSA)
SUMOylation of chromatin organization proteins
Transcriptional activity of SMAD2/SMAD3:SMAD4 heterotrimer
Homology Directed Repair
Peptide hormone metabolism
Regulation of TP53 Expression and Degradation
Regulation of mRNA stability by proteins that bind AU-rich elements
G alpha (q) signalling events
Cell Cycle Checkpoints
Epigenetic regulation of gene expression
Ephrin signaling
Signaling by Hedgehog
Factors involved in megakaryocyte development and platelet production
Signaling by ROBO receptors
Vesicle-mediated transport
Platelet activation, signaling and aggregation
rRNA processing in the nucleus and cytosol
Cell surface interactions at the vascular wall
Signaling by FGFR2
Mitotic G1-G1/S phases
Metabolism of vitamins and cofactors
Regulation of TNFR1 signaling
Signaling by FGFR
Transcriptional regulation by RUNX2
Immunoregulatory interactions between a Lymphoid and a non-Lymphoid cell
Other interleukin signaling
RHO GTPase Effectors
Fc epsilon receptor (FCERI) signaling
Signaling by Receptor Tyrosine Kinases
Interferon Signaling
C-type lectin receptors (CLRs)
Processing of Capped Intron-Containing Pre-mRNA
rRNA processing
Interleukin-12 family signaling
Metabolism of carbohydrates

Adaptive Immune System
Class I MHC mediated antigen processing & presentation
Signaling by WNT
Translation
trans-Golgi Network Vesicle Budding
Host Interactions of HIV factors
Signaling by TGF-beta Receptor Complex
TNF signaling
DNA Double-Strand Break Repair
G alpha (s) signalling events
Activation of HOX genes during differentiation
Activation of anterior HOX genes in hindbrain development during early embryogenesis
Signaling by PTK6
Signaling by Non-Receptor Tyrosine Kinases
Cell Cycle, Mitotic
EPH-Ephrin signaling
Signaling by TGF-beta family members
Death Receptor Signalling
Class A/1 (Rhodopsin-like receptors)
Extracellular matrix organization
Transcriptional regulation by the AP-2 (TFAP2) family of transcription factors
Neurodegenerative Diseases
Deregulated CDK5 triggers multiple neurodegenerative pathways in Alzheimer's disease models
SLC-mediated transmembrane transport
Transcriptional Regulation by TP53
Signaling by Interleukins
Signaling by NOTCH
Apoptosis
RNA Polymerase II Transcription
Programmed Cell Death
Metabolism of amino acids and derivatives
Estrogen-dependent gene expression
HIV Infection
Metabolism of steroids
GPCR ligand binding
Generic Transcription Pathway
G alpha (i) signalling events
Gene expression (Transcription)
Regulation of TP53 Activity
Hemostasis
DNA Repair
Cell Cycle
Axon guidance
ESR-mediated signaling
Cytokine Signaling in Immune system
Metabolism of lipids
Signaling by Nuclear Receptors
Post-translational protein modification
Infectious disease
SUMO E3 ligases SUMOylate target proteins

SUMOylation
GPCR downstream signalling
Innate Immune System
Metabolism of RNA
Immune System
Transport of small molecules
Developmental Biology
Metabolism of proteins
Signaling by GPCR
Disease
Signal Transduction
Metabolism

| #Entities found | #Entities total | #Interactors found | #Interactors total | Entities ratio |
|---|---|---|---|---|
| 8 | 93 | 1 | 100 | 0.006562698 |
| 5 | 60 | 0 | 57 | 0.004233999 |
| 2 | 13 | 0 | 2 | 9.17E-04 |
| 2 | 14 | 0 | 2 | 9.88E-04 |
| 3 | 23 | 1 | 39 | 0.001623033 |
| 2 | 20 | 0 | 4 | 0.001411333 |
| 1 | 3 | 0 | 0 | 2.12E-04 |
| 1 | 3 | 0 | 0 | 2.12E-04 |
| 1 | 3 | 0 | 0 | 2.12E-04 |
| 2 | 34 | 0 | 2 | 0.002399266 |
| 4 | 124 | 2 | 92 | 0.008750265 |
| 2 | 29 | 1 | 23 | 0.002046433 |
| 4 | 36 | 0 | 202 | 0.002540399 |
| 1 | 7 | 0 | 2 | 4.94E-04 |
| 2 | 18 | 0 | 50 | 0.0012702 |
| 1 | 5 | 0 | 7 | 3.53E-04 |
| 13 | 498 | 8 | 959 | 0.035142192 |
| 1 | 14 | 0 | 1 | 9.88E-04 |
| 1 | 11 | 0 | 5 | 7.76E-04 |
| 0 | 6 | 1 | 18 | 4.23E-04 |
| 3 | 69 | 0 | 127 | 0.004869099 |
| 0 | 9 | 1 | 10 | 6.35E-04 |
| 2 | 97 | 0 | 0 | 0.006844965 |
| 1 | 12 | 0 | 10 | 8.47E-04 |
| 0 | 9 | 1 | 11 | 6.35E-04 |
| 0 | 2 | 1 | 20 | 1.41E-04 |
| 0 | 2 | 1 | 20 | 1.41E-04 |
| 2 | 74 | 0 | 32 | 0.005221932 |
| 1 | 19 | 0 | 4 | 0.001340766 |
| 0 | 2 | 1 | 24 | 1.41E-04 |
| 0 | 16 | 1 | 11 | 0.001129066 |
| 1 | 3 | 0 | 24 | 2.12E-04 |
| 1 | 3 | 0 | 24 | 2.12E-04 |
| 1 | 3 | 0 | 24 | 2.12E-04 |
| 0 | 8 | 1 | 20 | 5.65E-04 |
| 0 | 6 | 1 | 22 | 4.23E-04 |
| 2 | 39 | 0 | 86 | 0.002752099 |
| 0 | 9 | 1 | 20 | 6.35E-04 |
| 0 | 9 | 1 | 20 | 6.35E-04 |
| 2 | 70 | 0 | 56 | 0.004939666 |
| 1 | 27 | 0 | 3 | 0.0019053 |
| 0 | 6 | 1 | 25 | 4.23E-04 |
| 0 | 6 | 1 | 25 | 4.23E-04 |
| 1 | 28 | 0 | 4 | 0.001975866 |
| 1 | 12 | 0 | 21 | 8.47E-04 |
| 1 | 11 | 0 | 21 | 7.76E-04 |
| 0 | 11 | 1 | 22 | 7.76E-04 |
| 2 | 81 | 0 | 53 | 0.005715899 |
| 0 | 8 | 1 | 28 | 5.65E-04 |

| | | | | |
|---|---|---|---|---|
| 1 | 25 | 0 | 9 | 0.001764166 |
| 1 | 3 | 0 | 32 | 2.12E-04 |
| 1 | 28 | 0 | 8 | 0.001975866 |
| 0 | 9 | 1 | 28 | 6.35E-04 |
| 0 | 11 | 1 | 24 | 7.76E-04 |
| 0 | 8 | 1 | 29 | 5.65E-04 |
| 0 | 8 | 1 | 29 | 5.65E-04 |
| 2 | 109 | 0 | 27 | 0.007691765 |
| 1 | 5 | 0 | 33 | 3.53E-04 |
| 1 | 9 | 0 | 27 | 6.35E-04 |
| 0 | 10 | 1 | 28 | 7.06E-04 |
| 0 | 7 | 1 | 31 | 4.94E-04 |
| 0 | 19 | 2 | 19 | 0.001340766 |
| 0 | 5 | 1 | 34 | 3.53E-04 |
| 2 | 63 | 2 | 89 | 0.004445699 |
| 1 | 6 | 0 | 36 | 4.23E-04 |
| 0 | 15 | 1 | 28 | 0.0010585 |
| 0 | 13 | 1 | 33 | 9.17E-04 |
| 0 | 6 | 1 | 36 | 4.23E-04 |
| 0 | 20 | 1 | 22 | 0.001411333 |
| 0 | 7 | 1 | 36 | 4.94E-04 |
| 0 | 16 | 1 | 29 | 0.001129066 |
| 0 | 16 | 1 | 29 | 0.001129066 |
| 3 | 148 | 1 | 142 | 0.010443864 |
| 1 | 44 | 0 | 0 | 0.003104933 |
| 0 | 36 | 1 | 10 | 0.002540399 |
| 0 | 20 | 1 | 28 | 0.001411333 |
| 2 | 71 | 0 | 92 | 0.005010232 |
| 2 | 71 | 0 | 94 | 0.005010232 |
| 0 | 5 | 1 | 46 | 3.53E-04 |
| 0 | 23 | 1 | 28 | 0.001623033 |
| 1 | 13 | 0 | 37 | 9.17E-04 |
| 0 | 39 | 1 | 11 | 0.002752099 |
| 0 | 17 | 1 | 36 | 0.001199633 |
| 2 | 144 | 1 | 28 | 0.010161598 |
| 1 | 26 | 1 | 28 | 0.001834733 |
| 1 | 52 | 0 | 0 | 0.003669466 |
| 0 | 19 | 1 | 35 | 0.001340766 |
| 2 | 32 | 0 | 143 | 0.002258133 |
| 0 | 26 | 1 | 29 | 0.001834733 |
| 0 | 7 | 1 | 51 | 4.94E-04 |
| 0 | 25 | 1 | 30 | 0.001764166 |
| 0 | 11 | 1 | 42 | 7.76E-04 |
| 0 | 10 | 1 | 45 | 7.06E-04 |
| 2 | 83 | 0 | 94 | 0.005857032 |
| 0 | 4 | 1 | 50 | 2.82E-04 |
| 0 | 8 | 1 | 51 | 5.65E-04 |
| 0 | 6 | 1 | 50 | 4.23E-04 |
| 0 | 6 | 1 | 50 | 4.23E-04 |
| 2 | 32 | 2 | 152 | 0.002258133 |

| | | | | |
|---|---|---|---|---|
| 0 | 30 | 1 | 28 | 0.002117 |
| 0 | 8 | 2 | 50 | 5.65E-04 |
| 2 | 99 | 1 | 102 | 0.006986098 |
| 0 | 9 | 1 | 51 | 6.35E-04 |
| 0 | 32 | 1 | 28 | 0.002258133 |
| 0 | 5 | 1 | 56 | 3.53E-04 |
| 0 | 15 | 1 | 48 | 0.0010585 |
| 3 | 134 | 1 | 202 | 0.009455931 |
| 0 | 19 | 2 | 45 | 0.001340766 |
| 0 | 10 | 1 | 55 | 7.06E-04 |
| 0 | 8 | 2 | 57 | 5.65E-04 |
| 1 | 64 | 0 | 0 | 0.004516266 |
| 0 | 30 | 1 | 36 | 0.002117 |
| 0 | 19 | 1 | 48 | 0.001340766 |
| 0 | 52 | 1 | 12 | 0.003669466 |
| 0 | 13 | 1 | 55 | 9.17E-04 |
| 0 | 12 | 1 | 55 | 8.47E-04 |
| 3 | 112 | 2 | 240 | 0.007903465 |
| 1 | 5 | 0 | 62 | 3.53E-04 |
| 0 | 13 | 1 | 55 | 9.17E-04 |
| 1 | 67 | 0 | 0 | 0.004727966 |
| 0 | 14 | 1 | 55 | 9.88E-04 |
| 0 | 12 | 1 | 57 | 8.47E-04 |
| 0 | 10 | 1 | 59 | 7.06E-04 |
| 0 | 15 | 1 | 55 | 0.0010585 |
| 0 | 41 | 1 | 27 | 0.002893233 |
| 0 | 16 | 1 | 59 | 0.001129066 |
| 1 | 40 | 0 | 30 | 0.002822666 |
| 0 | 18 | 1 | 57 | 0.0012702 |
| 0 | 28 | 1 | 46 | 0.001975866 |
| 0 | 11 | 2 | 61 | 7.76E-04 |
| 0 | 12 | 1 | 66 | 8.47E-04 |
| 1 | 45 | 1 | 32 | 0.003175499 |
| 0 | 34 | 1 | 41 | 0.002399266 |
| 0 | 22 | 1 | 55 | 0.001552466 |
| 0 | 4 | 1 | 74 | 2.82E-04 |
| 0 | 27 | 1 | 49 | 0.0019053 |
| 0 | 11 | 3 | 68 | 7.76E-04 |
| 1 | 45 | 1 | 33 | 0.003175499 |
| 0 | 21 | 1 | 56 | 0.0014819 |
| 1 | 18 | 2 | 64 | 0.0012702 |
| 0 | 24 | 1 | 55 | 0.0016936 |
| 0 | 54 | 1 | 24 | 0.003810599 |
| 0 | 25 | 1 | 55 | 0.001764166 |
| 0 | 25 | 1 | 57 | 0.001764166 |
| 0 | 46 | 1 | 33 | 0.003246066 |
| 0 | 26 | 1 | 55 | 0.001834733 |
| 0 | 28 | 2 | 59 | 0.001975866 |
| 2 | 92 | 0 | 140 | 0.006492132 |
| 0 | 30 | 1 | 51 | 0.002117 |

| | | | | |
|---|---|---|---|---|
| 1 | 35 | 0 | 49 | 0.002469833 |
| 0 | 8 | 1 | 78 | 5.65E-04 |
| 0 | 3 | 1 | 80 | 2.12E-04 |
| 0 | 29 | 1 | 55 | 0.002046433 |
| 0 | 19 | 1 | 65 | 0.001340766 |
| 0 | 26 | 2 | 56 | 0.001834733 |
| 0 | 30 | 1 | 55 | 0.002117 |
| 0 | 14 | 1 | 74 | 9.88E-04 |
| 1 | 10 | 2 | 77 | 7.06E-04 |
| 0 | 15 | 1 | 74 | 0.0010585 |
| 0 | 32 | 1 | 55 | 0.002258133 |
| 1 | 25 | 0 | 60 | 0.001764166 |
| 2 | 107 | 0 | 136 | 0.007550632 |
| 0 | 8 | 1 | 81 | 5.65E-04 |
| 1 | 18 | 0 | 69 | 0.0012702 |
| 0 | 33 | 1 | 56 | 0.002328699 |
| 0 | 32 | 1 | 58 | 0.002258133 |
| 0 | 36 | 1 | 60 | 0.002540399 |
| 0 | 25 | 1 | 68 | 0.001764166 |
| 0 | 68 | 1 | 30 | 0.004798532 |
| 0 | 37 | 1 | 51 | 0.002610966 |
| 0 | 32 | 1 | 60 | 0.002258133 |
| 0 | 62 | 1 | 32 | 0.004375132 |
| 0 | 18 | 1 | 72 | 0.0012702 |
| 0 | 16 | 1 | 77 | 0.001129066 |
| 0 | 48 | 1 | 62 | 0.003387199 |
| 1 | 58 | 0 | 42 | 0.004092866 |
| 0 | 40 | 1 | 51 | 0.002822666 |
| 1 | 12 | 0 | 81 | 8.47E-04 |
| 1 | 16 | 0 | 77 | 0.001129066 |
| 0 | 21 | 1 | 74 | 0.0014819 |
| 0 | 14 | 1 | 81 | 9.88E-04 |
| 1 | 13 | 0 | 83 | 9.17E-04 |
| 0 | 15 | 1 | 80 | 0.0010585 |
| 7 | 315 | 3 | 805 | 0.022228495 |
| 0 | 12 | 1 | 84 | 8.47E-04 |
| 0 | 24 | 1 | 87 | 0.0016936 |
| 0 | 15 | 2 | 85 | 0.0010585 |
| 0 | 50 | 1 | 47 | 0.003528333 |
| 0 | 46 | 1 | 55 | 0.003246066 |
| 2 | 127 | 0 | 130 | 0.008961965 |
| 1 | 72 | 0 | 27 | 0.005080799 |
| 0 | 18 | 2 | 85 | 0.0012702 |
| 2 | 69 | 1 | 192 | 0.004869099 |
| 0 | 4 | 1 | 95 | 2.82E-04 |
| 0 | 18 | 1 | 85 | 0.0012702 |
| 0 | 18 | 1 | 85 | 0.0012702 |
| 0 | 40 | 1 | 66 | 0.002822666 |
| 0 | 20 | 1 | 85 | 0.001411333 |
| 0 | 35 | 1 | 73 | 0.002469833 |

| | | | | |
|---|---|---|---|---|
| 0 | 36 | 1 | 71 | 0.002540399 |
| 4 | 163 | 0 | 485 | 0.011502364 |
| 0 | 29 | 1 | 77 | 0.002046433 |
| 0 | 49 | 1 | 58 | 0.003457766 |
| 0 | 20 | 1 | 92 | 0.001411333 |
| 1 | 48 | 0 | 61 | 0.003387199 |
| 1 | 50 | 1 | 58 | 0.003528333 |
| 1 | 15 | 0 | 96 | 0.0010585 |
| 1 | 23 | 0 | 84 | 0.001623033 |
| 0 | 44 | 1 | 67 | 0.003104933 |
| 2 | 102 | 2 | 200 | 0.007197798 |
| 1 | 28 | 0 | 79 | 0.001975866 |
| 1 | 33 | 0 | 79 | 0.002328699 |
| 0 | 18 | 1 | 92 | 0.0012702 |
| 1 | 17 | 0 | 96 | 0.001199633 |
| 3 | 134 | 1 | 317 | 0.009455931 |
| 1 | 38 | 1 | 74 | 0.002681533 |
| 0 | 19 | 1 | 92 | 0.001340766 |
| 1 | 31 | 1 | 79 | 0.002187566 |
| 0 | 49 | 1 | 65 | 0.003457766 |
| 0 | 16 | 1 | 97 | 0.001129066 |
| 1 | 72 | 0 | 39 | 0.005080799 |
| 0 | 17 | 1 | 99 | 0.001199633 |
| 1 | 48 | 1 | 75 | 0.003387199 |
| 0 | 35 | 1 | 84 | 0.002469833 |
| 2 | 87 | 2 | 208 | 0.006139299 |
| 0 | 15 | 1 | 106 | 0.0010585 |
| 0 | 22 | 1 | 101 | 0.001552466 |
| 1 | 36 | 1 | 90 | 0.002540399 |
| 1 | 14 | 0 | 106 | 9.88E-04 |
| 2 | 161 | 0 | 133 | 0.011361231 |
| 2 | 207 | 0 | 86 | 0.014607297 |
| 1 | 49 | 0 | 74 | 0.003457766 |
| 0 | 25 | 1 | 95 | 0.001764166 |
| 0 | 38 | 1 | 82 | 0.002681533 |
| 0 | 62 | 1 | 58 | 0.004375132 |
| 0 | 7 | 2 | 112 | 4.94E-04 |
| 0 | 27 | 2 | 97 | 0.0019053 |
| 0 | 42 | 1 | 78 | 0.002963799 |
| 0 | 35 | 1 | 90 | 0.002469833 |
| 0 | 32 | 1 | 95 | 0.002258133 |
| 2 | 148 | 2 | 154 | 0.010443864 |
| 0 | 39 | 2 | 87 | 0.002752099 |
| 0 | 41 | 1 | 80 | 0.002893233 |
| 0 | 26 | 1 | 98 | 0.001834733 |
| 0 | 14 | 1 | 112 | 9.88E-04 |
| 1 | 48 | 2 | 78 | 0.003387199 |
| 0 | 14 | 1 | 112 | 9.88E-04 |
| 0 | 65 | 2 | 64 | 0.004586832 |
| 0 | 52 | 1 | 73 | 0.003669466 |

| | | | | |
|---|---|---|---|---|
| 1 | 49 | 0 | 80 | 0.003457766 |
| 1 | 10 | 1 | 115 | 7.06E-04 |
| 0 | 13 | 1 | 116 | 9.17E-04 |
| 0 | 97 | 1 | 28 | 0.006844965 |
| 0 | 32 | 1 | 95 | 0.002258133 |
| 0 | 81 | 1 | 46 | 0.005715899 |
| 1 | 55 | 1 | 73 | 0.003881166 |
| 0 | 24 | 1 | 108 | 0.0016936 |
| 0 | 15 | 1 | 118 | 0.0010585 |
| 0 | 40 | 1 | 86 | 0.002822666 |
| 0 | 13 | 1 | 118 | 9.17E-04 |
| 1 | 20 | 0 | 110 | 0.001411333 |
| 0 | 62 | 1 | 68 | 0.004375132 |
| 0 | 54 | 1 | 74 | 0.003810599 |
| 0 | 67 | 1 | 65 | 0.004727966 |
| 2 | 126 | 0 | 180 | 0.008891398 |
| 1 | 14 | 1 | 117 | 9.88E-04 |
| 1 | 23 | 1 | 107 | 0.001623033 |
| 0 | 11 | 2 | 118 | 7.76E-04 |
| 0 | 37 | 1 | 92 | 0.002610966 |
| 0 | 94 | 1 | 36 | 0.006633265 |
| 0 | 32 | 3 | 98 | 0.002258133 |
| 0 | 14 | 1 | 121 | 9.88E-04 |
| 0 | 94 | 1 | 37 | 0.006633265 |
| 1 | 12 | 0 | 121 | 8.47E-04 |
| 0 | 15 | 1 | 121 | 0.0010585 |
| 0 | 18 | 1 | 115 | 0.0012702 |
| 1 | 31 | 0 | 103 | 0.002187566 |
| 1 | 91 | 0 | 41 | 0.006421565 |
| 0 | 14 | 1 | 118 | 9.88E-04 |
| 1 | 49 | 2 | 88 | 0.003457766 |
| 0 | 8 | 1 | 127 | 5.65E-04 |
| 0 | 114 | 1 | 20 | 0.008044598 |
| 1 | 22 | 0 | 115 | 0.001552466 |
| 1 | 85 | 2 | 53 | 0.005998165 |
| 0 | 41 | 1 | 96 | 0.002893233 |
| 1 | 39 | 0 | 95 | 0.002752099 |
| 0 | 58 | 1 | 78 | 0.004092866 |
| 7 | 361 | 3 | 902 | 0.025474561 |
| 0 | 67 | 1 | 71 | 0.004727966 |
| 0 | 70 | 1 | 66 | 0.004939666 |
| 4 | 170 | 2 | 525 | 0.011996331 |
| 1 | 48 | 1 | 97 | 0.003387199 |
| 1 | 48 | 1 | 97 | 0.003387199 |
| 0 | 56 | 1 | 82 | 0.003951732 |
| 0 | 45 | 1 | 98 | 0.003175499 |
| 0 | 68 | 1 | 73 | 0.004798532 |
| 1 | 56 | 1 | 105 | 0.003951732 |
| 1 | 52 | 1 | 90 | 0.003669466 |
| 0 | 3 | 1 | 137 | 2.12E-04 |

| | | | | |
|---|---|---|---|---|
| 0 | 10 | 1 | 131 | 7.06E-04 |
| 0 | 54 | 2 | 86 | 0.003810599 |
| 0 | 10 | 1 | 130 | 7.06E-04 |
| 0 | 26 | 3 | 118 | 0.001834733 |
| 1 | 17 | 0 | 126 | 0.001199633 |
| 0 | 19 | 1 | 122 | 0.001340766 |
| 3 | 174 | 1 | 338 | 0.012278597 |
| 0 | 16 | 1 | 127 | 0.001129066 |
| 1 | 41 | 0 | 108 | 0.002893233 |
| 0 | 27 | 1 | 118 | 0.0019053 |
| 0 | 36 | 1 | 111 | 0.002540399 |
| 1 | 19 | 2 | 129 | 0.001340766 |
| 1 | 103 | 0 | 42 | 0.007268365 |
| 1 | 10 | 2 | 139 | 7.06E-04 |
| 0 | 42 | 2 | 105 | 0.002963799 |
| 3 | 218 | 3 | 318 | 0.01538353 |
| 0 | 17 | 2 | 134 | 0.001199633 |
| 1 | 28 | 0 | 126 | 0.001975866 |
| 0 | 42 | 1 | 121 | 0.002963799 |
| 0 | 18 | 1 | 141 | 0.0012702 |
| 1 | 43 | 0 | 110 | 0.003034366 |
| 0 | 39 | 2 | 120 | 0.002752099 |
| 0 | 45 | 1 | 110 | 0.003175499 |
| 0 | 48 | 2 | 113 | 0.003387199 |
| 3 | 176 | 1 | 366 | 0.01241973 |
| 0 | 28 | 1 | 132 | 0.001975866 |
| 1 | 32 | 0 | 124 | 0.002258133 |
| 0 | 88 | 2 | 79 | 0.006209865 |
| 0 | 7 | 1 | 152 | 4.94E-04 |
| 0 | 7 | 1 | 148 | 4.94E-04 |
| 0 | 120 | 1 | 46 | 0.008467998 |
| 0 | 27 | 1 | 132 | 0.0019053 |
| 0 | 16 | 1 | 146 | 0.001129066 |
| 0 | 55 | 1 | 104 | 0.003881166 |
| 0 | 39 | 1 | 118 | 0.002752099 |
| 0 | 4 | 1 | 153 | 2.82E-04 |
| 0 | 13 | 2 | 145 | 9.17E-04 |
| 1 | 43 | 1 | 125 | 0.003034366 |
| 1 | 46 | 2 | 120 | 0.003246066 |
| 0 | 66 | 3 | 103 | 0.004657399 |
| 1 | 13 | 0 | 147 | 9.17E-04 |
| 1 | 89 | 1 | 72 | 0.006280432 |
| 0 | 62 | 2 | 104 | 0.004375132 |
| 2 | 99 | 0 | 259 | 0.006986098 |
| 0 | 16 | 2 | 149 | 0.001129066 |
| 1 | 102 | 1 | 64 | 0.007197798 |
| 0 | 45 | 2 | 125 | 0.003175499 |
| 1 | 115 | 0 | 53 | 0.008115165 |
| 0 | 57 | 2 | 115 | 0.004022299 |
| 0 | 14 | 1 | 153 | 9.88E-04 |

| | | | | |
|---|---|---|---|---|
| 0 | 14 | 1 | 153 | 9.88E-04 |
| 0 | 46 | 1 | 124 | 0.003246066 |
| 0 | 18 | 1 | 151 | 0.0012702 |
| 1 | 47 | 0 | 120 | 0.003316633 |
| 0 | 41 | 1 | 137 | 0.002893233 |
| 2 | 114 | 2 | 270 | 0.008044598 |
| 1 | 94 | 0 | 78 | 0.006633265 |
| 1 | 94 | 0 | 78 | 0.006633265 |
| 0 | 14 | 1 | 154 | 9.88E-04 |
| 0 | 14 | 1 | 154 | 9.88E-04 |
| 0 | 25 | 1 | 150 | 0.001764166 |
| 1 | 98 | 1 | 119 | 0.006915532 |
| 1 | 10 | 0 | 160 | 7.06E-04 |
| 1 | 55 | 1 | 122 | 0.003881166 |
| 0 | 94 | 1 | 79 | 0.006633265 |
| 0 | 6 | 1 | 164 | 4.23E-04 |
| 0 | 66 | 2 | 106 | 0.004657399 |
| 0 | 40 | 2 | 140 | 0.002822666 |
| 1 | 30 | 0 | 149 | 0.002117 |
| 0 | 92 | 1 | 87 | 0.006492132 |
| 1 | 19 | 0 | 157 | 0.001340766 |
| 0 | 44 | 1 | 143 | 0.003104933 |
| 0 | 31 | 1 | 153 | 0.002187566 |
| 1 | 32 | 1 | 143 | 0.002258133 |
| 0 | 19 | 3 | 163 | 0.001340766 |
| 0 | 28 | 3 | 148 | 0.001975866 |
| 1 | 56 | 1 | 126 | 0.003951732 |
| 0 | 62 | 1 | 114 | 0.004375132 |
| 0 | 55 | 1 | 123 | 0.003881166 |
| 1 | 16 | 1 | 164 | 0.001129066 |
| 0 | 62 | 1 | 117 | 0.004375132 |
| 1 | 22 | 1 | 164 | 0.001552466 |
| 1 | 47 | 2 | 140 | 0.003316633 |
| 0 | 29 | 2 | 150 | 0.002046433 |
| 1 | 98 | 1 | 87 | 0.006915532 |
| 1 | 39 | 1 | 148 | 0.002752099 |
| 1 | 23 | 0 | 159 | 0.001623033 |
| 0 | 24 | 1 | 161 | 0.0016936 |
| 0 | 19 | 3 | 165 | 0.001340766 |
| 1 | 70 | 1 | 116 | 0.004939666 |
| 0 | 102 | 1 | 81 | 0.007197798 |
| 0 | 35 | 1 | 150 | 0.002469833 |
| 3 | 67 | 3 | 537 | 0.004727966 |
| 0 | 10 | 1 | 176 | 7.06E-04 |
| 0 | 33 | 1 | 157 | 0.002328699 |
| 0 | 19 | 1 | 176 | 0.001340766 |
| 0 | 23 | 1 | 172 | 0.001623033 |
| 1 | 26 | 0 | 165 | 0.001834733 |
| 1 | 23 | 0 | 170 | 0.001623033 |
| 0 | 56 | 1 | 141 | 0.003951732 |

| | | | | |
|---|---|---|---|---|
| 0 | 4 | 4 | 189 | 2.82E-04 |
| 0 | 12 | 2 | 184 | 8.47E-04 |
| 1 | 23 | 1 | 177 | 0.001623033 |
| 1 | 31 | 0 | 168 | 0.002187566 |
| 0 | 27 | 4 | 170 | 0.0019053 |
| 0 | 9 | 2 | 194 | 6.35E-04 |
| 0 | 59 | 1 | 144 | 0.004163432 |
| 0 | 100 | 1 | 98 | 0.007056665 |
| 1 | 42 | 5 | 169 | 0.002963799 |
| 1 | 58 | 0 | 140 | 0.004092866 |
| 1 | 110 | 0 | 107 | 0.007762332 |
| 1 | 30 | 0 | 170 | 0.002117 |
| 0 | 21 | 2 | 183 | 0.0014819 |
| 0 | 23 | 1 | 184 | 0.001623033 |
| 0 | 18 | 2 | 187 | 0.0012702 |
| 2 | 164 | 2 | 272 | 0.011572931 |
| 0 | 33 | 3 | 177 | 0.002328699 |
| 0 | 27 | 2 | 186 | 0.0019053 |
| 5 | 261 | 3 | 789 | 0.018417896 |
| 0 | 53 | 3 | 155 | 0.003740032 |
| 1 | 19 | 0 | 188 | 0.001340766 |
| 0 | 37 | 2 | 170 | 0.002610966 |
| 1 | 194 | 0 | 23 | 0.01368993 |
| 0 | 78 | 1 | 137 | 0.005504199 |
| 1 | 90 | 0 | 118 | 0.006350999 |
| 1 | 28 | 0 | 181 | 0.001975866 |
| 0 | 55 | 1 | 155 | 0.003881166 |
| 0 | 29 | 1 | 190 | 0.002046433 |
| 0 | 33 | 1 | 186 | 0.002328699 |
| 1 | 15 | 3 | 198 | 0.0010585 |
| 0 | 83 | 1 | 131 | 0.005857032 |
| 0 | 34 | 2 | 180 | 0.002399266 |
| 0 | 57 | 2 | 155 | 0.004022299 |
| 0 | 12 | 1 | 201 | 8.47E-04 |
| 1 | 24 | 0 | 190 | 0.0016936 |
| 1 | 25 | 0 | 190 | 0.001764166 |
| 0 | 124 | 1 | 103 | 0.008750265 |
| 0 | 124 | 1 | 103 | 0.008750265 |
| 0 | 29 | 1 | 190 | 0.002046433 |
| 0 | 106 | 1 | 108 | 0.007480065 |
| 1 | 149 | 0 | 72 | 0.010514431 |
| 1 | 134 | 0 | 86 | 0.009455931 |
| 1 | 28 | 0 | 190 | 0.001975866 |
| 2 | 95 | 1 | 337 | 0.006703832 |
| 0 | 43 | 1 | 178 | 0.003034366 |
| 0 | 79 | 3 | 140 | 0.005574765 |
| 1 | 34 | 0 | 188 | 0.002399266 |
| 0 | 102 | 1 | 129 | 0.007197798 |
| 2 | 37 | 2 | 400 | 0.002610966 |
| 1 | 31 | 0 | 190 | 0.002187566 |

| | | | | |
|---|---|---|---|---|
| 0 | 19 | 3 | 203 | 0.001340766 |
| 1 | 35 | 0 | 188 | 0.002469833 |
| 0 | 70 | 2 | 162 | 0.004939666 |
| 2 | 127 | 2 | 327 | 0.008961965 |
| 1 | 26 | 3 | 198 | 0.001834733 |
| 0 | 35 | 3 | 189 | 0.002469833 |
| 1 | 27 | 3 | 198 | 0.0019053 |
| 1 | 103 | 0 | 117 | 0.007268365 |
| 1 | 208 | 0 | 25 | 0.014677863 |
| 0 | 50 | 1 | 181 | 0.003528333 |
| 1 | 29 | 0 | 200 | 0.002046433 |
| 2 | 39 | 2 | 402 | 0.002752099 |
| 0 | 81 | 1 | 154 | 0.005715899 |
| 0 | 20 | 2 | 205 | 0.001411333 |
| 0 | 107 | 1 | 123 | 0.007550632 |
| 0 | 44 | 2 | 186 | 0.003104933 |
| 0 | 141 | 1 | 88 | 0.009949898 |
| 1 | 30 | 3 | 198 | 0.002117 |
| 0 | 19 | 1 | 212 | 0.001340766 |
| 1 | 41 | 0 | 188 | 0.002893233 |
| 1 | 39 | 0 | 190 | 0.002752099 |
| 1 | 64 | 1 | 164 | 0.004516266 |
| 0 | 41 | 2 | 189 | 0.002893233 |
| 0 | 41 | 2 | 189 | 0.002893233 |
| 1 | 33 | 3 | 198 | 0.002328699 |
| 1 | 86 | 1 | 147 | 0.006068732 |
| 0 | 32 | 1 | 201 | 0.002258133 |
| 0 | 50 | 1 | 189 | 0.003528333 |
| 0 | 49 | 1 | 181 | 0.003457766 |
| 0 | 38 | 1 | 191 | 0.002681533 |
| 0 | 22 | 1 | 219 | 0.001552466 |
| 0 | 105 | 1 | 137 | 0.007409498 |
| 1 | 16 | 1 | 217 | 0.001129066 |
| 1 | 35 | 0 | 202 | 0.002469833 |
| 1 | 68 | 1 | 171 | 0.004798532 |
| 0 | 107 | 1 | 130 | 0.007550632 |
| 0 | 62 | 1 | 182 | 0.004375132 |
| 0 | 141 | 2 | 122 | 0.009949898 |
| 2 | 100 | 7 | 372 | 0.007056665 |
| 1 | 71 | 1 | 171 | 0.005010232 |
| 1 | 80 | 1 | 162 | 0.005645332 |
| 1 | 72 | 1 | 171 | 0.005080799 |
| 2 | 101 | 1 | 368 | 0.007127232 |
| 0 | 69 | 2 | 178 | 0.004869099 |
| 0 | 131 | 1 | 115 | 0.009244231 |
| 0 | 67 | 1 | 179 | 0.004727966 |
| 0 | 77 | 2 | 182 | 0.005433632 |
| 1 | 94 | 2 | 158 | 0.006633265 |
| 2 | 219 | 2 | 280 | 0.015454096 |
| 0 | 89 | 3 | 163 | 0.006280432 |

| | | | | |
|---|---|---|---|---|
| 0 | 63 | 2 | 194 | 0.004445699 |
| 2 | 281 | 1 | 229 | 0.019829229 |
| 2 | 108 | 4 | 395 | 0.007621198 |
| 0 | 130 | 4 | 133 | 0.009173665 |
| 0 | 70 | 2 | 192 | 0.004939666 |
| 0 | 70 | 2 | 192 | 0.004939666 |
| 0 | 130 | 4 | 134 | 0.009173665 |
| 0 | 175 | 1 | 86 | 0.012349164 |
| 1 | 129 | 1 | 135 | 0.009103098 |
| 0 | 25 | 2 | 235 | 0.001764166 |
| 0 | 51 | 1 | 220 | 0.003598899 |
| 0 | 34 | 1 | 227 | 0.002399266 |
| 1 | 83 | 3 | 179 | 0.005857032 |
| 0 | 30 | 3 | 240 | 0.002117 |
| 0 | 35 | 1 | 232 | 0.002469833 |
| 0 | 183 | 1 | 84 | 0.012913697 |
| 0 | 65 | 1 | 197 | 0.004586832 |
| 3 | 325 | 3 | 398 | 0.022934161 |
| 1 | 43 | 2 | 227 | 0.003034366 |
| 0 | 29 | 2 | 243 | 0.002046433 |
| 0 | 72 | 2 | 206 | 0.005080799 |
| 0 | 23 | 2 | 247 | 0.001623033 |
| 1 | 123 | 2 | 149 | 0.008679698 |
| 1 | 63 | 1 | 210 | 0.004445699 |
| 0 | 30 | 2 | 244 | 0.002117 |
| 0 | 41 | 2 | 232 | 0.002893233 |
| 0 | 113 | 1 | 163 | 0.007974031 |
| 0 | 90 | 2 | 199 | 0.006350999 |
| 0 | 63 | 4 | 213 | 0.004445699 |
| 0 | 39 | 2 | 238 | 0.002752099 |
| 0 | 182 | 1 | 94 | 0.01284313 |
| 2 | 108 | 2 | 426 | 0.007621198 |
| 0 | 39 | 1 | 243 | 0.002752099 |
| 0 | 39 | 1 | 243 | 0.002752099 |
| 1 | 71 | 1 | 216 | 0.005010232 |
| 0 | 38 | 1 | 262 | 0.002681533 |
| 0 | 61 | 2 | 233 | 0.004304566 |
| 0 | 106 | 2 | 183 | 0.007480065 |
| 0 | 52 | 1 | 243 | 0.003669466 |
| 1 | 165 | 0 | 119 | 0.011643497 |
| 0 | 80 | 1 | 207 | 0.005645332 |
| 0 | 92 | 1 | 201 | 0.006492132 |
| 0 | 159 | 1 | 130 | 0.011220097 |
| 4 | 335 | 6 | 726 | 0.023639828 |
| 0 | 88 | 1 | 210 | 0.006209865 |
| 1 | 87 | 1 | 211 | 0.006139299 |
| 0 | 55 | 2 | 239 | 0.003881166 |
| 0 | 168 | 1 | 128 | 0.011855197 |
| 0 | 55 | 3 | 257 | 0.003881166 |
| 0 | 58 | 1 | 242 | 0.004092866 |

| | | | | |
|---|---|---|---|---|
| 0 | 135 | 1 | 160 | 0.009526498 |
| 1 | 37 | 0 | 260 | 0.002610966 |
| 0 | 26 | 2 | 271 | 0.001834733 |
| 0 | 57 | 2 | 242 | 0.004022299 |
| 1 | 95 | 3 | 203 | 0.006703832 |
| 1 | 59 | 1 | 264 | 0.004163432 |
| 0 | 39 | 2 | 260 | 0.002752099 |
| 0 | 36 | 1 | 267 | 0.002540399 |
| 0 | 71 | 2 | 240 | 0.005010232 |
| 2 | 104 | 3 | 456 | 0.007338932 |
| 2 | 122 | 4 | 457 | 0.008609131 |
| 0 | 89 | 1 | 220 | 0.006280432 |
| 0 | 68 | 1 | 239 | 0.004798532 |
| 0 | 68 | 1 | 239 | 0.004798532 |
| 0 | 68 | 1 | 239 | 0.004798532 |
| 0 | 68 | 1 | 239 | 0.004798532 |
| 0 | 68 | 1 | 239 | 0.004798532 |
| 0 | 43 | 1 | 263 | 0.003034366 |
| 0 | 113 | 3 | 192 | 0.007974031 |
| 0 | 103 | 1 | 203 | 0.007268365 |
| 1 | 96 | 1 | 217 | 0.006774398 |
| 0 | 135 | 1 | 181 | 0.009526498 |
| 0 | 220 | 1 | 87 | 0.015524663 |
| 1 | 50 | 3 | 271 | 0.003528333 |
| 1 | 211 | 0 | 109 | 0.014889563 |
| 2 | 133 | 4 | 457 | 0.009385364 |
| 0 | 124 | 2 | 196 | 0.008750265 |
| 1 | 53 | 1 | 270 | 0.003740032 |
| 1 | 89 | 1 | 231 | 0.006280432 |
| 2 | 59 | 2 | 522 | 0.004163432 |
| 0 | 20 | 2 | 295 | 0.001411333 |
| 2 | 208 | 4 | 421 | 0.014677863 |
| 5 | 352 | 7 | 897 | 0.024839461 |
| 0 | 72 | 2 | 243 | 0.005080799 |
| 0 | 72 | 1 | 259 | 0.005080799 |
| 0 | 88 | 3 | 248 | 0.006209865 |
| 0 | 56 | 1 | 272 | 0.003951732 |
| 1 | 80 | 2 | 254 | 0.005645332 |
| 0 | 9 | 4 | 312 | 6.35E-04 |
| 1 | 32 | 1 | 295 | 0.002258133 |
| 0 | 16 | 3 | 314 | 0.001129066 |
| 1 | 36 | 2 | 300 | 0.002540399 |
| 1 | 50 | 4 | 294 | 0.003528333 |
| 0 | 60 | 2 | 275 | 0.004233999 |
| 0 | 93 | 2 | 247 | 0.006562698 |
| 2 | 390 | 1 | 217 | 0.027520994 |
| 0 | 17 | 1 | 313 | 0.001199633 |
| 3 | 231 | 7 | 616 | 0.016300896 |
| 1 | 83 | 2 | 260 | 0.005857032 |
| 0 | 83 | 1 | 260 | 0.005857032 |

| | | | | |
|---|---|---|---|---|
| 1 | 315 | 0 | 26 | 0.022228495 |
| 1 | 78 | 1 | 270 | 0.005504199 |
| 0 | 254 | 1 | 91 | 0.017923929 |
| 2 | 125 | 4 | 511 | 0.008820831 |
| 1 | 90 | 3 | 282 | 0.006350999 |
| 0 | 39 | 1 | 316 | 0.002752099 |
| 0 | 39 | 1 | 316 | 0.002752099 |
| 3 | 154 | 1 | 700 | 0.010867264 |
| 2 | 212 | 3 | 426 | 0.01496013 |
| 1 | 43 | 3 | 312 | 0.003034366 |
| 2 | 214 | 3 | 426 | 0.015101263 |
| 0 | 144 | 2 | 210 | 0.010161598 |
| 2 | 110 | 5 | 519 | 0.007762332 |
| 1 | 80 | 2 | 282 | 0.005645332 |
| 1 | 124 | 1 | 250 | 0.008750265 |
| 1 | 32 | 5 | 332 | 0.002258133 |
| 1 | 112 | 5 | 259 | 0.007903465 |
| 0 | 53 | 1 | 316 | 0.003740032 |
| 2 | 134 | 4 | 531 | 0.009455931 |
| 0 | 86 | 2 | 290 | 0.006068732 |
| 2 | 256 | 1 | 438 | 0.018065062 |
| 2 | 256 | 1 | 438 | 0.018065062 |
| 1 | 46 | 3 | 334 | 0.003246066 |
| 0 | 9 | 2 | 355 | 6.35E-04 |
| 1 | 69 | 2 | 309 | 0.004869099 |
| 1 | 206 | 0 | 172 | 0.01453673 |
| 1 | 48 | 7 | 342 | 0.003387199 |
| 0 | 81 | 1 | 300 | 0.005715899 |
| 5 | 489 | 5 | 930 | 0.034507092 |
| 0 | 123 | 1 | 252 | 0.008679698 |
| 4 | 457 | 4 | 690 | 0.032248959 |
| 0 | 20 | 5 | 360 | 0.001411333 |
| 0 | 83 | 1 | 303 | 0.005857032 |
| 0 | 39 | 2 | 335 | 0.002752099 |
| 0 | 94 | 3 | 298 | 0.006633265 |
| 0 | 69 | 1 | 332 | 0.004869099 |
| 0 | 73 | 3 | 309 | 0.005151365 |
| 0 | 26 | 2 | 363 | 0.001834733 |
| 0 | 26 | 2 | 363 | 0.001834733 |
| 2 | 295 | 3 | 401 | 0.020817162 |
| 0 | 256 | 2 | 134 | 0.018065062 |
| 1 | 97 | 3 | 299 | 0.006844965 |
| 1 | 23 | 1 | 363 | 0.001623033 |
| 0 | 43 | 4 | 352 | 0.003034366 |
| 0 | 71 | 2 | 325 | 0.005010232 |
| 1 | 158 | 1 | 247 | 0.011149531 |
| 1 | 184 | 0 | 217 | 0.012984264 |
| 0 | 188 | 1 | 221 | 0.01326653 |
| 1 | 34 | 3 | 368 | 0.002399266 |
| 1 | 94 | 3 | 310 | 0.006633265 |

| | | | | |
|---|---|---|---|---|
| 0 | 65 | 3 | 350 | 0.004586832 |
| 1 | 35 | 3 | 368 | 0.002469833 |
| 0 | 77 | 3 | 338 | 0.005433632 |
| 1 | 274 | 1 | 142 | 0.019335262 |
| 0 | 39 | 2 | 366 | 0.002752099 |
| 0 | 46 | 2 | 363 | 0.003246066 |
| 0 | 202 | 2 | 218 | 0.014254463 |
| 0 | 256 | 1 | 151 | 0.018065062 |
| 1 | 41 | 3 | 368 | 0.002893233 |
| 1 | 41 | 3 | 368 | 0.002893233 |
| 0 | 67 | 4 | 346 | 0.004727966 |
| 2 | 144 | 0 | 593 | 0.010161598 |
| 1 | 229 | 1 | 202 | 0.016159763 |
| 1 | 47 | 3 | 384 | 0.003316633 |
| 2 | 198 | 6 | 545 | 0.013972197 |
| 0 | 38 | 2 | 393 | 0.002681533 |
| 0 | 109 | 4 | 326 | 0.007691765 |
| 0 | 21 | 3 | 411 | 0.0014819 |
| 1 | 250 | 0 | 186 | 0.017641663 |
| 4 | 216 | 9 | 1044 | 0.015242396 |
| 1 | 44 | 3 | 401 | 0.003104933 |
| 0 | 232 | 1 | 202 | 0.016371463 |
| 2 | 115 | 9 | 637 | 0.008115165 |
| 0 | 199 | 2 | 260 | 0.014042763 |
| 0 | 163 | 2 | 299 | 0.011502364 |
| 1 | 44 | 7 | 419 | 0.003104933 |
| 0 | 106 | 2 | 358 | 0.007480065 |
| 1 | 74 | 1 | 381 | 0.005221932 |
| 2 | 421 | 2 | 344 | 0.02970856 |
| 0 | 179 | 4 | 316 | 0.01263143 |
| 6 | 665 | 8 | 1161 | 0.046926822 |
| 0 | 190 | 1 | 294 | 0.013407664 |
| 0 | 150 | 4 | 359 | 0.010584998 |
| 1 | 87 | 3 | 402 | 0.006139299 |
| 0 | 177 | 2 | 320 | 0.012490297 |
| 0 | 84 | 3 | 401 | 0.005927599 |
| 2 | 184 | 0 | 645 | 0.012984264 |
| 1 | 90 | 5 | 411 | 0.006350999 |
| 0 | 96 | 2 | 400 | 0.006774398 |
| 1 | 480 | 0 | 0 | 0.033871992 |
| 0 | 54 | 2 | 445 | 0.003810599 |
| 1 | 193 | 1 | 315 | 0.013619363 |
| 1 | 81 | 3 | 414 | 0.005715899 |
| 1 | 118 | 4 | 397 | 0.008326865 |
| 1 | 59 | 3 | 446 | 0.004163432 |
| 2 | 332 | 3 | 518 | 0.023428128 |
| 2 | 185 | 4 | 707 | 0.01305483 |
| 1 | 90 | 3 | 425 | 0.006350999 |
| 1 | 107 | 0 | 443 | 0.007550632 |
| 1 | 107 | 0 | 443 | 0.007550632 |

| | | | | |
|---|---|---|---|---|
| 1 | 165 | 1 | 355 | 0.011643497 |
| 0 | 111 | 4 | 408 | 0.007832898 |
| 0 | 167 | 3 | 351 | 0.011784631 |
| 0 | 40 | 2 | 483 | 0.002822666 |
| 2 | 196 | 4 | 710 | 0.013831063 |
| 1 | 50 | 3 | 477 | 0.003528333 |
| 2 | 130 | 4 | 704 | 0.009173665 |
| 1 | 146 | 2 | 408 | 0.010302731 |
| 1 | 52 | 3 | 478 | 0.003669466 |
| 5 | 621 | 9 | 1082 | 0.04382189 |
| 2 | 288 | 5 | 622 | 0.020323195 |
| 4 | 515 | 7 | 915 | 0.036341825 |
| 0 | 43 | 2 | 487 | 0.003034366 |
| 0 | 189 | 3 | 368 | 0.013337097 |
| 1 | 61 | 3 | 478 | 0.004304566 |
| 0 | 124 | 1 | 442 | 0.008750265 |
| 0 | 62 | 3 | 479 | 0.004375132 |
| 0 | 55 | 4 | 495 | 0.003881166 |
| 0 | 130 | 1 | 445 | 0.009173665 |
| 1 | 147 | 3 | 401 | 0.010373298 |
| 0 | 46 | 8 | 500 | 0.003246066 |
| 0 | 93 | 4 | 459 | 0.006562698 |
| 1 | 282 | 2 | 267 | 0.019899795 |
| 1 | 279 | 2 | 320 | 0.019688095 |
| 1 | 139 | 2 | 440 | 0.009808764 |
| 0 | 23 | 4 | 529 | 0.001623033 |
| 0 | 168 | 2 | 394 | 0.011855197 |
| 1 | 194 | 6 | 374 | 0.01368993 |
| 0 | 235 | 4 | 347 | 0.016583163 |
| 6 | 824 | 8 | 1187 | 0.05814692 |
| 2 | 293 | 2 | 637 | 0.020676029 |
| 0 | 207 | 3 | 400 | 0.014607297 |
| 2 | 256 | 4 | 656 | 0.018065062 |
| 1 | 87 | 3 | 527 | 0.006139299 |
| 1 | 173 | 4 | 495 | 0.01220803 |
| 0 | 385 | 2 | 248 | 0.02716816 |
| 0 | 39 | 2 | 593 | 0.002752099 |
| 1 | 106 | 3 | 529 | 0.007480065 |
| 1 | 147 | 4 | 498 | 0.010373298 |
| 0 | 316 | 2 | 326 | 0.022299061 |
| 2 | 318 | 3 | 682 | 0.022440195 |
| 2 | 326 | 3 | 666 | 0.023004728 |
| 0 | 234 | 2 | 439 | 0.016512596 |
| 6 | 541 | 12 | 1609 | 0.038176558 |
| 2 | 392 | 3 | 605 | 0.027662127 |
| 0 | 202 | 1 | 489 | 0.014254463 |
| 2 | 256 | 5 | 833 | 0.018065062 |
| 0 | 245 | 3 | 443 | 0.017288829 |
| 0 | 96 | 3 | 577 | 0.006774398 |
| 0 | 457 | 2 | 218 | 0.032248959 |

| | | | | |
|---|---|---|---|---|
| 6 | 998 | 5 | 1197 | 0.070425517 |
| 1 | 465 | 1 | 228 | 0.032813492 |
| 4 | 330 | 10 | 1307 | 0.023286995 |
| 1 | 339 | 4 | 364 | 0.023922094 |
| 0 | 80 | 4 | 630 | 0.005645332 |
| 0 | 150 | 6 | 571 | 0.010584998 |
| 0 | 85 | 4 | 642 | 0.005998165 |
| 0 | 50 | 2 | 684 | 0.003528333 |
| 0 | 167 | 1 | 625 | 0.011784631 |
| 0 | 605 | 2 | 130 | 0.042692823 |
| 0 | 116 | 1 | 630 | 0.008185731 |
| 0 | 116 | 1 | 630 | 0.008185731 |
| 1 | 70 | 6 | 690 | 0.004939666 |
| 1 | 70 | 6 | 690 | 0.004939666 |
| 3 | 570 | 8 | 950 | 0.040222991 |
| 1 | 101 | 4 | 691 | 0.007127232 |
| 0 | 114 | 4 | 724 | 0.008044598 |
| 2 | 157 | 4 | 1072 | 0.011078964 |
| 0 | 452 | 2 | 400 | 0.031896126 |
| 0 | 329 | 2 | 535 | 0.023216428 |
| 0 | 52 | 4 | 795 | 0.003669466 |
| 0 | 30 | 4 | 851 | 0.002117 |
| 0 | 30 | 4 | 851 | 0.002117 |
| 1 | 421 | 3 | 483 | 0.02970856 |
| 4 | 485 | 11 | 1467 | 0.034224825 |
| 7 | 836 | 10 | 2014 | 0.05899372 |
| 0 | 257 | 6 | 679 | 0.018135629 |
| 1 | 187 | 3 | 796 | 0.013195964 |
| 16 | 1663 | 25 | 3504 | 0.117352339 |
| 1 | 195 | 3 | 807 | 0.013760497 |
| 1 | 662 | 1 | 288 | 0.046715122 |
| 0 | 154 | 4 | 818 | 0.010867264 |
| 0 | 262 | 6 | 733 | 0.018488462 |
| 1 | 321 | 3 | 676 | 0.022651895 |
| 0 | 643 | 2 | 419 | 0.045374356 |
| 14 | 1524 | 24 | 3367 | 0.107543575 |
| 0 | 567 | 5 | 518 | 0.040011291 |
| 16 | 1821 | 25 | 3684 | 0.12850187 |
| 0 | 178 | 8 | 1001 | 0.012560864 |
| 4 | 820 | 11 | 1501 | 0.057864653 |
| 0 | 367 | 3 | 955 | 0.025897961 |
| 3 | 682 | 10 | 1491 | 0.048126455 |
| 3 | 583 | 9 | 1469 | 0.041140357 |
| 0 | 255 | 6 | 1009 | 0.017994496 |
| 8 | 1245 | 13 | 2455 | 0.08785548 |
| 6 | 1444 | 7 | 1523 | 0.101898243 |
| 0 | 317 | 6 | 1015 | 0.022369628 |
| 9 | 1594 | 17 | 2533 | 0.11248324 |
| 0 | 540 | 11 | 1114 | 0.038105991 |
| 1 | 183 | 11 | 1533 | 0.012913697 |

| | | | | |
|---|---|---|---|---|
| 1 | 192 | 11 | 1591 | 0.013548797 |
| 2 | 1358 | 9 | 808 | 0.095829511 |
| 4 | 1324 | 6 | 1762 | 0.093430245 |
| 2 | 782 | 12 | 1628 | 0.05518312 |
| 14 | 2803 | 16 | 3476 | 0.197798321 |
| 1 | 967 | 5 | 942 | 0.068237951 |
| 6 | 1206 | 13 | 2698 | 0.08510338 |
| 13 | 2355 | 22 | 3612 | 0.166184461 |
| 2 | 1475 | 10 | 1162 | 0.104085809 |
| 6 | 1555 | 17 | 2691 | 0.109731141 |
| 19 | 3212 | 39 | 5222 | 0.22666008 |
| 7 | 3639 | 10 | 2191 | 0.25679204 |

| Entities pValue | Entities FDR | #Reactions found | #Reactions total | Reactions ratio |
|---|---|---|---|---|
| 1.42E-05 | 0.011870299 | 25 | 33 | 0.00270381 |
| 5.79E-04 | 0.242078661 | 15 | 19 | 0.001556739 |
| 0.003447982 | 0.677628804 | 2 | 2 | 1.64E-04 |
| 0.003908468 | 0.677628804 | 2 | 4 | 3.28E-04 |
| 0.004656389 | 0.677628804 | 5 | 8 | 6.55E-04 |
| 0.008536657 | 0.677628804 | 5 | 6 | 4.92E-04 |
| 0.017009743 | 0.677628804 | 1 | 1 | 8.19E-05 |
| 0.017009743 | 0.677628804 | 1 | 1 | 8.19E-05 |
| 0.017009743 | 0.677628804 | 1 | 1 | 8.19E-05 |
| 0.018374045 | 0.677628804 | 2 | 6 | 4.92E-04 |
| 0.0311029 | 0.677628804 | 9 | 34 | 0.002785744 |
| 0.034898381 | 0.677628804 | 13 | 24 | 0.001966407 |
| 0.045969942 | 0.677628804 | 1 | 14 | 0.001147071 |
| 0.050173452 | 0.677628804 | 1 | 4 | 3.28E-04 |
| 0.055332912 | 0.677628804 | 2 | 10 | 8.19E-04 |
| 0.060979632 | 0.677628804 | 2 | 2 | 1.64E-04 |
| 0.063672935 | 0.677628804 | 74 | 210 | 0.017206063 |
| 0.0822277 | 0.677628804 | 1 | 5 | 4.10E-04 |
| 0.087464812 | 0.677628804 | 1 | 2 | 1.64E-04 |
| 0.097850324 | 0.677628804 | 1 | 5 | 4.10E-04 |
| 0.098288706 | 0.677628804 | 13 | 15 | 0.001229005 |
| 0.102999055 | 0.677628804 | 4 | 4 | 3.28E-04 |
| 0.106868598 | 0.677628804 | 2 | 2 | 1.64E-04 |
| 0.108118656 | 0.677628804 | 3 | 4 | 3.28E-04 |
| 0.108118656 | 0.677628804 | 1 | 3 | 2.46E-04 |
| 0.113209289 | 0.677628804 | 1 | 1 | 8.19E-05 |
| 0.113209289 | 0.677628804 | 1 | 1 | 8.19E-05 |
| 0.117940006 | 0.677628804 | 2 | 2 | 1.64E-04 |
| 0.123304302 | 0.677628804 | 5 | 5 | 4.10E-04 |
| 0.133285384 | 0.677628804 | 1 | 2 | 1.64E-04 |
| 0.133285384 | 0.677628804 | 2 | 7 | 5.74E-04 |
| 0.1382336 | 0.677628804 | 1 | 2 | 1.64E-04 |
| 0.1382336 | 0.677628804 | 1 | 2 | 1.64E-04 |
| 0.1382336 | 0.677628804 | 1 | 2 | 1.64E-04 |
| 0.14315381 | 0.677628804 | 2 | 4 | 3.28E-04 |
| 0.148046171 | 0.677628804 | 1 | 5 | 4.10E-04 |
| 0.152673698 | 0.677628804 | 2 | 5 | 4.10E-04 |
| 0.152910839 | 0.677628804 | 1 | 5 | 4.10E-04 |
| 0.152910839 | 0.677628804 | 1 | 5 | 4.10E-04 |
| 0.154659448 | 0.677628804 | 1 | 12 | 9.83E-04 |
| 0.157747971 | 0.677628804 | 1 | 4 | 3.28E-04 |
| 0.162557719 | 0.677628804 | 1 | 1 | 8.19E-05 |
| 0.162557719 | 0.677628804 | 1 | 1 | 8.19E-05 |
| 0.167340239 | 0.677628804 | 5 | 5 | 4.10E-04 |
| 0.167340239 | 0.677628804 | 4 | 8 | 6.55E-04 |
| 0.167340239 | 0.677628804 | 2 | 6 | 4.92E-04 |
| 0.172095683 | 0.677628804 | 1 | 5 | 4.10E-04 |
| 0.174775585 | 0.677628804 | 3 | 3 | 2.46E-04 |
| 0.176824202 | 0.677628804 | 1 | 3 | 2.46E-04 |

| | | | | |
|---|---|---|---|---|
| 0.176824202 | 0.677628804 | 1 | 9 | 7.37E-04 |
| 0.181525948 | 0.677628804 | 1 | 1 | 8.19E-05 |
| 0.181525948 | 0.677628804 | 5 | 10 | 8.19E-04 |
| 0.181525948 | 0.677628804 | 1 | 2 | 1.64E-04 |
| 0.181525948 | 0.677628804 | 1 | 3 | 2.46E-04 |
| 0.181525948 | 0.677628804 | 1 | 5 | 4.10E-04 |
| 0.181525948 | 0.677628804 | 1 | 5 | 4.10E-04 |
| 0.18293767 | 0.677628804 | 1 | 1 | 8.19E-05 |
| 0.186201072 | 0.677628804 | 1 | 1 | 8.19E-05 |
| 0.186201072 | 0.677628804 | 3 | 3 | 2.46E-04 |
| 0.186201072 | 0.677628804 | 1 | 2 | 1.64E-04 |
| 0.186201072 | 0.677628804 | 1 | 2 | 1.64E-04 |
| 0.195472047 | 0.677628804 | 1 | 2 | 1.64E-04 |
| 0.200068195 | 0.677628804 | 2 | 4 | 3.28E-04 |
| 0.201495552 | 0.677628804 | 3 | 7 | 5.74E-04 |
| 0.209182547 | 0.677628804 | 2 | 2 | 1.64E-04 |
| 0.209182547 | 0.677628804 | 1 | 4 | 3.28E-04 |
| 0.213701043 | 0.677628804 | 2 | 5 | 4.10E-04 |
| 0.213701043 | 0.677628804 | 1 | 3 | 2.46E-04 |
| 0.213701043 | 0.677628804 | 1 | 4 | 3.28E-04 |
| 0.218193944 | 0.677628804 | 1 | 5 | 4.10E-04 |
| 0.218193944 | 0.677628804 | 1 | 6 | 4.92E-04 |
| 0.218193944 | 0.677628804 | 1 | 6 | 4.92E-04 |
| 0.222155678 | 0.677628804 | 5 | 26 | 0.002130274 |
| 0.222661395 | 0.677628804 | 1 | 1 | 8.19E-05 |
| 0.227103539 | 0.677628804 | 1 | 5 | 4.10E-04 |
| 0.231520517 | 0.677628804 | 2 | 4 | 3.28E-04 |
| 0.232855236 | 0.677628804 | 2 | 2 | 1.64E-04 |
| 0.23706378 | 0.677628804 | 3 | 3 | 2.46E-04 |
| 0.240279541 | 0.677628804 | 1 | 1 | 8.19E-05 |
| 0.244621868 | 0.677628804 | 1 | 4 | 3.28E-04 |
| 0.24893959 | 0.677628804 | 2 | 2 | 1.64E-04 |
| 0.24893959 | 0.677628804 | 1 | 9 | 7.37E-04 |
| 0.253232845 | 0.677628804 | 1 | 5 | 4.10E-04 |
| 0.253934498 | 0.677628804 | 3 | 5 | 4.10E-04 |
| 0.257501771 | 0.677628804 | 2 | 2 | 1.64E-04 |
| 0.257501771 | 0.677628804 | 1 | 6 | 4.92E-04 |
| 0.257501771 | 0.677628804 | 1 | 8 | 6.55E-04 |
| 0.260270608 | 0.677628804 | 3 | 8 | 6.55E-04 |
| 0.261746505 | 0.677628804 | 2 | 7 | 5.74E-04 |
| 0.261746505 | 0.677628804 | 1 | 4 | 3.28E-04 |
| 0.261746505 | 0.677628804 | 2 | 10 | 8.19E-04 |
| 0.261746505 | 0.677628804 | 1 | 6 | 4.92E-04 |
| 0.261746505 | 0.677628804 | 1 | 8 | 6.55E-04 |
| 0.262383217 | 0.677628804 | 6 | 6 | 4.92E-04 |
| 0.265967182 | 0.677628804 | 2 | 2 | 1.64E-04 |
| 0.265967182 | 0.677628804 | 4 | 10 | 8.19E-04 |
| 0.270163938 | 0.677628804 | 1 | 5 | 4.10E-04 |
| 0.270163938 | 0.677628804 | 1 | 5 | 4.10E-04 |
| 0.272947387 | 0.677628804 | 9 | 19 | 0.001556739 |

| | | | | |
|---|---|---|---|---|
| 0.274336907 | 0.677628804 | 2 | 6 | 4.92E-04 |
| 0.278486222 | 0.677628804 | 1 | 2 | 1.64E-04 |
| 0.279284339 | 0.677628804 | 8 | 9 | 7.37E-04 |
| 0.282612017 | 0.677628804 | 1 | 5 | 4.10E-04 |
| 0.282612017 | 0.677628804 | 1 | 6 | 4.92E-04 |
| 0.290793572 | 0.677628804 | 2 | 2 | 1.64E-04 |
| 0.290793572 | 0.677628804 | 1 | 3 | 2.46E-04 |
| 0.290964593 | 0.677628804 | 3 | 10 | 8.19E-04 |
| 0.294849596 | 0.677628804 | 1 | 5 | 4.10E-04 |
| 0.302892783 | 0.677628804 | 1 | 2 | 1.64E-04 |
| 0.306880204 | 0.677628804 | 5 | 6 | 4.92E-04 |
| 0.306880204 | 0.677628804 | 1 | 2 | 1.64E-04 |
| 0.306880204 | 0.677628804 | 1 | 4 | 3.28E-04 |
| 0.306880204 | 0.677628804 | 1 | 8 | 6.55E-04 |
| 0.306880204 | 0.677628804 | 1 | 12 | 9.83E-04 |
| 0.310845015 | 0.677628804 | 1 | 3 | 2.46E-04 |
| 0.310845015 | 0.677628804 | 1 | 6 | 4.92E-04 |
| 0.31433065 | 0.677628804 | 17 | 89 | 0.007292093 |
| 0.314787343 | 0.677628804 | 1 | 2 | 1.64E-04 |
| 0.314787343 | 0.677628804 | 1 | 4 | 3.28E-04 |
| 0.318707313 | 0.677628804 | 2 | 4 | 3.28E-04 |
| 0.318707313 | 0.677628804 | 1 | 4 | 3.28E-04 |
| 0.318707313 | 0.677628804 | 1 | 6 | 4.92E-04 |
| 0.318707313 | 0.677628804 | 1 | 6 | 4.92E-04 |
| 0.322605051 | 0.677628804 | 1 | 5 | 4.10E-04 |
| 0.322605051 | 0.677628804 | 1 | 7 | 5.74E-04 |
| 0.330334333 | 0.677628804 | 1 | 6 | 4.92E-04 |
| 0.330334333 | 0.677628804 | 1 | 9 | 7.37E-04 |
| 0.334166125 | 0.677628804 | 1 | 5 | 4.10E-04 |
| 0.334166125 | 0.677628804 | 5 | 40 | 0.003277345 |
| 0.33797618 | 0.677628804 | 2 | 5 | 4.10E-04 |
| 0.341764622 | 0.677628804 | 1 | 7 | 5.74E-04 |
| 0.345531572 | 0.677628804 | 3 | 4 | 3.28E-04 |
| 0.345531572 | 0.677628804 | 2 | 3 | 2.46E-04 |
| 0.345531572 | 0.677628804 | 1 | 4 | 3.28E-04 |
| 0.349277151 | 0.677628804 | 1 | 2 | 1.64E-04 |
| 0.349277151 | 0.677628804 | 1 | 6 | 4.92E-04 |
| 0.353001479 | 0.677628804 | 2 | 7 | 5.74E-04 |
| 0.353001479 | 0.677628804 | 2 | 13 | 0.001065137 |
| 0.353001479 | 0.677628804 | 1 | 7 | 5.74E-04 |
| 0.356704676 | 0.677628804 | 2 | 3 | 2.46E-04 |
| 0.356704676 | 0.677628804 | 1 | 6 | 4.92E-04 |
| 0.36038686 | 0.677628804 | 2 | 3 | 2.46E-04 |
| 0.36038686 | 0.677628804 | 1 | 4 | 3.28E-04 |
| 0.36038686 | 0.677628804 | 2 | 10 | 8.19E-04 |
| 0.36038686 | 0.677628804 | 1 | 6 | 4.92E-04 |
| 0.36404815 | 0.677628804 | 1 | 4 | 3.28E-04 |
| 0.367688662 | 0.677628804 | 2 | 4 | 3.28E-04 |
| 0.371040975 | 0.677628804 | 4 | 12 | 9.83E-04 |
| 0.371308515 | 0.677628804 | 1 | 2 | 1.64E-04 |

| | | | | |
|---|---|---|---|---|
| 0.371308515 | 0.677628804 | 2 | 7 | 5.74E-04 |
| 0.374907825 | 0.677628804 | 2 | 2 | 1.64E-04 |
| 0.374907825 | 0.677628804 | 2 | 3 | 2.46E-04 |
| 0.374907825 | 0.677628804 | 1 | 4 | 3.28E-04 |
| 0.374907825 | 0.677628804 | 1 | 8 | 6.55E-04 |
| 0.374907825 | 0.677628804 | 1 | 10 | 8.19E-04 |
| 0.378486705 | 0.677628804 | 1 | 12 | 9.83E-04 |
| 0.382045273 | 0.677628804 | 1 | 1 | 8.19E-05 |
| 0.385583642 | 0.677628804 | 4 | 4 | 3.28E-04 |
| 0.385583642 | 0.677628804 | 1 | 2 | 1.64E-04 |
| 0.385583642 | 0.677628804 | 1 | 4 | 3.28E-04 |
| 0.385583642 | 0.677628804 | 1 | 5 | 4.10E-04 |
| 0.387298268 | 0.677628804 | 2 | 12 | 9.83E-04 |
| 0.389101926 | 0.677628804 | 1 | 5 | 4.10E-04 |
| 0.389101926 | 0.677628804 | 2 | 14 | 0.001147071 |
| 0.392600237 | 0.677628804 | 1 | 4 | 3.28E-04 |
| 0.392600237 | 0.677628804 | 1 | 6 | 4.92E-04 |
| 0.392600237 | 0.677628804 | 1 | 9 | 7.37E-04 |
| 0.396078688 | 0.677628804 | 1 | 10 | 8.19E-04 |
| 0.396078688 | 0.677628804 | 1 | 12 | 9.83E-04 |
| 0.396078688 | 0.677628804 | 1 | 15 | 0.001229005 |
| 0.399537391 | 0.677628804 | 3 | 21 | 0.001720606 |
| 0.402976458 | 0.677628804 | 1 | 2 | 1.64E-04 |
| 0.402976458 | 0.677628804 | 1 | 5 | 4.10E-04 |
| 0.402976458 | 0.677628804 | 1 | 9 | 7.37E-04 |
| 0.402976458 | 0.677628804 | 1 | 18 | 0.001474805 |
| 0.406395997 | 0.677628804 | 3 | 3 | 2.46E-04 |
| 0.406395997 | 0.677628804 | 2 | 14 | 0.001147071 |
| 0.40979612 | 0.677628804 | 1 | 3 | 2.46E-04 |
| 0.40979612 | 0.677628804 | 1 | 4 | 3.28E-04 |
| 0.40979612 | 0.677628804 | 1 | 6 | 4.92E-04 |
| 0.413176936 | 0.677628804 | 2 | 5 | 4.10E-04 |
| 0.416538554 | 0.677628804 | 1 | 4 | 3.28E-04 |
| 0.416538554 | 0.677628804 | 1 | 15 | 0.001229005 |
| 0.417933749 | 0.677628804 | 47 | 86 | 0.007046293 |
| 0.41988108 | 0.677628804 | 1 | 5 | 4.10E-04 |
| 0.41988108 | 0.677628804 | 1 | 5 | 4.10E-04 |
| 0.423204624 | 0.677628804 | 2 | 7 | 5.74E-04 |
| 0.423204624 | 0.677628804 | 1 | 8 | 6.55E-04 |
| 0.423204624 | 0.677628804 | 2 | 20 | 0.001638673 |
| 0.427108476 | 0.677628804 | 1 | 14 | 0.001147071 |
| 0.429795189 | 0.677628804 | 5 | 5 | 4.10E-04 |
| 0.429795189 | 0.677628804 | 3 | 13 | 0.001065137 |
| 0.432966819 | 0.677628804 | 7 | 33 | 0.00270381 |
| 0.433062423 | 0.677628804 | 3 | 10 | 8.19E-04 |
| 0.433062423 | 0.677628804 | 1 | 4 | 3.28E-04 |
| 0.436311097 | 0.677628804 | 1 | 8 | 6.55E-04 |
| 0.439541316 | 0.677628804 | 1 | 6 | 4.92E-04 |
| 0.442753185 | 0.677628804 | 1 | 4 | 3.28E-04 |
| 0.442753185 | 0.677628804 | 1 | 11 | 9.01E-04 |

| | | | | |
|---|---|---|---|---|
| 0.445946807 | 0.677628804 | 1 | 22 | 0.00180254 |
| 0.447281751 | 0.677628804 | 1 | 79 | 0.006472757 |
| 0.449122283 | 0.677628804 | 1 | 7 | 5.74E-04 |
| 0.449122283 | 0.677628804 | 1 | 17 | 0.001392872 |
| 0.449122283 | 0.677628804 | 1 | 17 | 0.001392872 |
| 0.452279718 | 0.677628804 | 2 | 10 | 8.19E-04 |
| 0.452279718 | 0.677628804 | 2 | 17 | 0.001392872 |
| 0.455419211 | 0.677628804 | 3 | 5 | 4.10E-04 |
| 0.455419211 | 0.677628804 | 1 | 6 | 4.92E-04 |
| 0.455419211 | 0.677628804 | 1 | 10 | 8.19E-04 |
| 0.456078518 | 0.677628804 | 6 | 54 | 0.004424416 |
| 0.458540865 | 0.677628804 | 1 | 5 | 4.10E-04 |
| 0.458540865 | 0.677628804 | 1 | 7 | 5.74E-04 |
| 0.458540865 | 0.677628804 | 1 | 7 | 5.74E-04 |
| 0.46164478 | 0.677628804 | 3 | 10 | 8.19E-04 |
| 0.461751511 | 0.677628804 | 4 | 21 | 0.001720606 |
| 0.464731055 | 0.677628804 | 2 | 7 | 5.74E-04 |
| 0.464731055 | 0.677628804 | 1 | 5 | 4.10E-04 |
| 0.46779979 | 0.677628804 | 3 | 7 | 5.74E-04 |
| 0.470851084 | 0.677628804 | 2 | 11 | 9.01E-04 |
| 0.470851084 | 0.677628804 | 1 | 8 | 6.55E-04 |
| 0.470851084 | 0.677628804 | 1 | 23 | 0.001884474 |
| 0.473885035 | 0.677628804 | 1 | 2 | 1.64E-04 |
| 0.476901741 | 0.677628804 | 1 | 4 | 3.28E-04 |
| 0.476901741 | 0.677628804 | 2 | 14 | 0.001147071 |
| 0.482355472 | 0.677628804 | 21 | 32 | 0.002621876 |
| 0.482883806 | 0.677628804 | 3 | 10 | 8.19E-04 |
| 0.482883806 | 0.677628804 | 1 | 8 | 6.55E-04 |
| 0.485849357 | 0.677628804 | 3 | 19 | 0.001556739 |
| 0.485849357 | 0.677628804 | 2 | 18 | 0.001474805 |
| 0.487885629 | 0.677628804 | 18 | 34 | 0.002785744 |
| 0.487885629 | 0.677628804 | 7 | 15 | 0.001229005 |
| 0.488798048 | 0.677628804 | 2 | 22 | 0.00180254 |
| 0.488798048 | 0.677628804 | 1 | 12 | 9.83E-04 |
| 0.488798048 | 0.677628804 | 1 | 20 | 0.001638673 |
| 0.488798048 | 0.677628804 | 1 | 23 | 0.001884474 |
| 0.491729975 | 0.677628804 | 4 | 5 | 4.10E-04 |
| 0.491729975 | 0.677628804 | 3 | 13 | 0.001065137 |
| 0.491729975 | 0.677628804 | 4 | 18 | 0.001474805 |
| 0.491729975 | 0.677628804 | 1 | 6 | 4.92E-04 |
| 0.491729975 | 0.677628804 | 2 | 25 | 0.002048341 |
| 0.493379354 | 0.677628804 | 4 | 18 | 0.001474805 |
| 0.494645232 | 0.677628804 | 6 | 26 | 0.002130274 |
| 0.494645232 | 0.677628804 | 3 | 16 | 0.001310938 |
| 0.494645232 | 0.677628804 | 1 | 9 | 7.37E-04 |
| 0.497543912 | 0.677628804 | 1 | 1 | 8.19E-05 |
| 0.497543912 | 0.677628804 | 15 | 22 | 0.00180254 |
| 0.497543912 | 0.677628804 | 1 | 2 | 1.64E-04 |
| 0.500426109 | 0.677628804 | 2 | 10 | 8.19E-04 |
| 0.500426109 | 0.677628804 | 1 | 15 | 0.001229005 |

| | | | | |
|---|---|---|---|---|
| 0.503291916 | 0.677628804 | 1 | 14 | 0.001147071 |
| 0.506141426 | 0.677628804 | 2 | 4 | 3.28E-04 |
| 0.506141426 | 0.677628804 | 1 | 6 | 4.92E-04 |
| 0.50897473 | 0.677628804 | 1 | 5 | 4.10E-04 |
| 0.50897473 | 0.677628804 | 1 | 5 | 4.10E-04 |
| 0.50897473 | 0.677628804 | 2 | 44 | 0.00360508 |
| 0.51179192 | 0.677628804 | 3 | 9 | 7.37E-04 |
| 0.51179192 | 0.677628804 | 2 | 7 | 5.74E-04 |
| 0.51179192 | 0.677628804 | 3 | 12 | 9.83E-04 |
| 0.51179192 | 0.677628804 | 1 | 9 | 7.37E-04 |
| 0.51179192 | 0.677628804 | 1 | 10 | 8.19E-04 |
| 0.514593086 | 0.677628804 | 1 | 9 | 7.37E-04 |
| 0.514593086 | 0.677628804 | 1 | 13 | 0.001065137 |
| 0.514593086 | 0.677628804 | 1 | 19 | 0.001556739 |
| 0.514593086 | 0.677628804 | 1 | 27 | 0.002212208 |
| 0.516755959 | 0.677628804 | 2 | 7 | 5.74E-04 |
| 0.517378319 | 0.677628804 | 6 | 7 | 5.74E-04 |
| 0.517378319 | 0.677628804 | 4 | 9 | 7.37E-04 |
| 0.517378319 | 0.677628804 | 1 | 10 | 8.19E-04 |
| 0.520147709 | 0.677628804 | 1 | 6 | 4.92E-04 |
| 0.520147709 | 0.677628804 | 1 | 8 | 6.55E-04 |
| 0.520147709 | 0.677628804 | 3 | 26 | 0.002130274 |
| 0.522901345 | 0.677628804 | 1 | 1 | 8.19E-05 |
| 0.522901345 | 0.677628804 | 1 | 5 | 4.10E-04 |
| 0.525639316 | 0.677628804 | 2 | 4 | 3.28E-04 |
| 0.525639316 | 0.677628804 | 1 | 2 | 1.64E-04 |
| 0.525639316 | 0.677628804 | 1 | 8 | 6.55E-04 |
| 0.528361709 | 0.677628804 | 2 | 13 | 0.001065137 |
| 0.528361709 | 0.677628804 | 5 | 34 | 0.002785744 |
| 0.531068614 | 0.677628804 | 1 | 5 | 4.10E-04 |
| 0.531068614 | 0.677628804 | 6 | 34 | 0.002785744 |
| 0.531068614 | 0.677628804 | 1 | 7 | 5.74E-04 |
| 0.533760118 | 0.677628804 | 1 | 2 | 1.64E-04 |
| 0.533760118 | 0.677628804 | 3 | 7 | 5.74E-04 |
| 0.533760118 | 0.677628804 | 4 | 14 | 0.001147071 |
| 0.533760118 | 0.677628804 | 1 | 7 | 5.74E-04 |
| 0.533760118 | 0.677628804 | 1 | 12 | 9.83E-04 |
| 0.533760118 | 0.677628804 | 2 | 27 | 0.002212208 |
| 0.538915611 | 0.677628804 | 47 | 113 | 0.009258501 |
| 0.539097267 | 0.677628804 | 2 | 26 | 0.002130274 |
| 0.539097267 | 0.677628804 | 1 | 37 | 0.003031544 |
| 0.541644538 | 0.677628804 | 42 | 56 | 0.004588283 |
| 0.541743084 | 0.677628804 | 13 | 14 | 0.001147071 |
| 0.541743084 | 0.677628804 | 13 | 14 | 0.001147071 |
| 0.541743084 | 0.677628804 | 1 | 5 | 4.10E-04 |
| 0.544373845 | 0.677628804 | 1 | 21 | 0.001720606 |
| 0.544373845 | 0.677628804 | 1 | 25 | 0.002048341 |
| 0.546989634 | 0.677628804 | 7 | 12 | 9.83E-04 |
| 0.546989634 | 0.677628804 | 6 | 20 | 0.001638673 |
| 0.546989634 | 0.677628804 | 1 | 4 | 3.28E-04 |

| | | | | |
|---|---|---|---|---|
| 0.546989634 | 0.677628804 | 1 | 5 | 4.10E-04 |
| 0.549590535 | 0.677628804 | 3 | 5 | 4.10E-04 |
| 0.549590535 | 0.677628804 | 2 | 5 | 4.10E-04 |
| 0.549590535 | 0.677628804 | 2 | 7 | 5.74E-04 |
| 0.549590535 | 0.677628804 | 3 | 19 | 0.001556739 |
| 0.549590535 | 0.677628804 | 1 | 7 | 5.74E-04 |
| 0.54976679 | 0.677628804 | 6 | 41 | 0.003359279 |
| 0.55730475 | 0.677628804 | 1 | 7 | 5.74E-04 |
| 0.55730475 | 0.677628804 | 1 | 11 | 9.01E-04 |
| 0.559846936 | 0.677628804 | 1 | 7 | 5.74E-04 |
| 0.559846936 | 0.677628804 | 1 | 14 | 0.001147071 |
| 0.562374649 | 0.677628804 | 4 | 13 | 0.001065137 |
| 0.562374649 | 0.677628804 | 1 | 19 | 0.001556739 |
| 0.564887972 | 0.677628804 | 5 | 5 | 4.10E-04 |
| 0.564887972 | 0.677628804 | 1 | 17 | 0.001392872 |
| 0.567151752 | 0.677628804 | 8 | 48 | 0.003932814 |
| 0.56987177 | 0.677628804 | 3 | 6 | 4.92E-04 |
| 0.56987177 | 0.677628804 | 2 | 12 | 9.83E-04 |
| 0.577241553 | 0.677628804 | 1 | 13 | 0.001065137 |
| 0.579670222 | 0.677628804 | 9 | 26 | 0.002130274 |
| 0.579670222 | 0.677628804 | 1 | 7 | 5.74E-04 |
| 0.582085059 | 0.677628804 | 2 | 3 | 2.46E-04 |
| 0.582085059 | 0.677628804 | 1 | 5 | 4.10E-04 |
| 0.582085059 | 0.677628804 | 1 | 28 | 0.002294142 |
| 0.582836672 | 0.677628804 | 6 | 44 | 0.00360508 |
| 0.584486142 | 0.677628804 | 2 | 2 | 1.64E-04 |
| 0.584486142 | 0.677628804 | 2 | 14 | 0.001147071 |
| 0.584486142 | 0.677628804 | 2 | 21 | 0.001720606 |
| 0.58687355 | 0.677628804 | 4 | 7 | 5.74E-04 |
| 0.58687355 | 0.677628804 | 1 | 4 | 3.28E-04 |
| 0.591607645 | 0.677628804 | 1 | 3 | 2.46E-04 |
| 0.591607645 | 0.677628804 | 4 | 16 | 0.001310938 |
| 0.591607645 | 0.677628804 | 2 | 9 | 7.37E-04 |
| 0.591607645 | 0.677628804 | 1 | 11 | 9.01E-04 |
| 0.591607645 | 0.677628804 | 1 | 14 | 0.001147071 |
| 0.593954487 | 0.677628804 | 1 | 2 | 1.64E-04 |
| 0.596287959 | 0.677628804 | 1 | 5 | 4.10E-04 |
| 0.596287959 | 0.677628804 | 2 | 24 | 0.001966407 |
| 0.598608137 | 0.677628804 | 2 | 3 | 2.46E-04 |
| 0.598608137 | 0.677628804 | 3 | 6 | 4.92E-04 |
| 0.598608137 | 0.677628804 | 2 | 19 | 0.001556739 |
| 0.598608137 | 0.677628804 | 2 | 22 | 0.00180254 |
| 0.600915095 | 0.677628804 | 1 | 7 | 5.74E-04 |
| 0.601409341 | 0.677628804 | 13 | 36 | 0.002949611 |
| 0.60320891 | 0.677628804 | 2 | 9 | 7.37E-04 |
| 0.605489654 | 0.677628804 | 3 | 9 | 7.37E-04 |
| 0.605489654 | 0.677628804 | 3 | 12 | 9.83E-04 |
| 0.607757402 | 0.677628804 | 3 | 14 | 0.001147071 |
| 0.607757402 | 0.677628804 | 1 | 34 | 0.002785744 |
| 0.610012227 | 0.677628804 | 1 | 2 | 1.64E-04 |

| | | | | |
|---|---|---|---|---|
| 0.610012227 | 0.677628804 | 1 | 2 | 1.64E-04 |
| 0.610012227 | 0.677628804 | 1 | 16 | 0.001310938 |
| 0.612254202 | 0.677628804 | 1 | 13 | 0.001065137 |
| 0.612254202 | 0.677628804 | 1 | 14 | 0.001147071 |
| 0.6144834 | 0.677628804 | 1 | 5 | 4.10E-04 |
| 0.616655175 | 0.677628804 | 6 | 39 | 0.003195412 |
| 0.616699892 | 0.677628804 | 4 | 4 | 3.28E-04 |
| 0.616699892 | 0.677628804 | 4 | 4 | 3.28E-04 |
| 0.616699892 | 0.677628804 | 1 | 4 | 3.28E-04 |
| 0.616699892 | 0.677628804 | 1 | 4 | 3.28E-04 |
| 0.616699892 | 0.677628804 | 1 | 13 | 0.001065137 |
| 0.618903752 | 0.677628804 | 4 | 11 | 9.01E-04 |
| 0.621095049 | 0.677628804 | 4 | 4 | 3.28E-04 |
| 0.621095049 | 0.677628804 | 1 | 14 | 0.001147071 |
| 0.623273855 | 0.677628804 | 18 | 21 | 0.001720606 |
| 0.623273855 | 0.677628804 | 1 | 4 | 3.28E-04 |
| 0.623273855 | 0.677628804 | 1 | 24 | 0.001966407 |
| 0.625440241 | 0.677628804 | 2 | 3 | 2.46E-04 |
| 0.625440241 | 0.677628804 | 3 | 33 | 0.00270381 |
| 0.625440241 | 0.677628804 | 1 | 14 | 0.001147071 |
| 0.627594277 | 0.677628804 | 1 | 9 | 7.37E-04 |
| 0.629736032 | 0.677628804 | 1 | 8 | 6.55E-04 |
| 0.629736032 | 0.677628804 | 1 | 15 | 0.001229005 |
| 0.631865577 | 0.677628804 | 4 | 11 | 9.01E-04 |
| 0.631865577 | 0.677628804 | 3 | 10 | 8.19E-04 |
| 0.631865577 | 0.677628804 | 2 | 15 | 0.001229005 |
| 0.633982979 | 0.677628804 | 6 | 24 | 0.001966407 |
| 0.633982979 | 0.677628804 | 1 | 5 | 4.10E-04 |
| 0.633982979 | 0.677628804 | 1 | 6 | 4.92E-04 |
| 0.636088308 | 0.677628804 | 2 | 9 | 7.37E-04 |
| 0.636088308 | 0.677628804 | 1 | 5 | 4.10E-04 |
| 0.638181632 | 0.677628804 | 2 | 14 | 0.001147071 |
| 0.640263018 | 0.677628804 | 6 | 9 | 7.37E-04 |
| 0.640263018 | 0.677628804 | 1 | 4 | 3.28E-04 |
| 0.640263018 | 0.677628804 | 2 | 36 | 0.002949611 |
| 0.642332535 | 0.677628804 | 4 | 16 | 0.001310938 |
| 0.642332535 | 0.677628804 | 3 | 15 | 0.001229005 |
| 0.642332535 | 0.677628804 | 1 | 19 | 0.001556739 |
| 0.644390249 | 0.677628804 | 1 | 4 | 3.28E-04 |
| 0.646436227 | 0.677628804 | 31 | 42 | 0.003441213 |
| 0.646436227 | 0.677628804 | 2 | 17 | 0.001392872 |
| 0.648470535 | 0.677628804 | 1 | 12 | 9.83E-04 |
| 0.648893798 | 0.677628804 | 13 | 13 | 0.001065137 |
| 0.652504407 | 0.677628804 | 1 | 5 | 4.10E-04 |
| 0.652504407 | 0.677628804 | 1 | 20 | 0.001638673 |
| 0.654504101 | 0.677628804 | 2 | 13 | 0.001065137 |
| 0.66238944 | 0.677628804 | 7 | 15 | 0.001229005 |
| 0.66238944 | 0.677628804 | 2 | 6 | 4.92E-04 |
| 0.666264943 | 0.677628804 | 2 | 4 | 3.28E-04 |
| 0.666264943 | 0.677628804 | 1 | 4 | 3.28E-04 |

| | | | | |
|---|---|---|---|---|
| 0.668186124 | 0.677628804 | 2 | 5 | 4.10E-04 |
| 0.668186124 | 0.677628804 | 2 | 6 | 4.92E-04 |
| 0.67009634 | 0.677628804 | 6 | 11 | 9.01E-04 |
| 0.671995655 | 0.677628804 | 1 | 25 | 0.002048341 |
| 0.67388413 | 0.677628804 | 3 | 22 | 0.00180254 |
| 0.675761826 | 0.677628804 | 2 | 5 | 4.10E-04 |
| 0.675761826 | 0.677628804 | 2 | 15 | 0.001229005 |
| 0.675761826 | 0.677628804 | 1 | 19 | 0.001556739 |
| 0.677628804 | 0.677628804 | 5 | 18 | 0.001474805 |
| 0.677628804 | 0.677628804 | 1 | 6 | 4.92E-04 |
| 0.679485125 | 0.679485125 | 4 | 7 | 5.74E-04 |
| 0.679485125 | 0.679485125 | 2 | 6 | 4.92E-04 |
| 0.681330849 | 0.681330849 | 2 | 11 | 9.01E-04 |
| 0.681330849 | 0.681330849 | 1 | 18 | 0.001474805 |
| 0.683166036 | 0.683166036 | 2 | 15 | 0.001229005 |
| 0.684970277 | 0.684970277 | 5 | 39 | 0.003195412 |
| 0.684990746 | 0.684990746 | 2 | 7 | 5.74E-04 |
| 0.686805038 | 0.686805038 | 4 | 21 | 0.001720606 |
| 0.68782726 | 0.68782726 | 25 | 132 | 0.01081524 |
| 0.68860897 | 0.68860897 | 2 | 5 | 4.10E-04 |
| 0.68860897 | 0.68860897 | 1 | 11 | 9.01E-04 |
| 0.690402603 | 0.690402603 | 2 | 20 | 0.001638673 |
| 0.692185993 | 0.692185993 | 2 | 8 | 6.55E-04 |
| 0.692185993 | 0.692185993 | 1 | 23 | 0.001884474 |
| 0.692185993 | 0.692185993 | 1 | 71 | 0.005817288 |
| 0.693959199 | 0.693959199 | 2 | 5 | 4.10E-04 |
| 0.693959199 | 0.693959199 | 1 | 10 | 8.19E-04 |
| 0.693959199 | 0.693959199 | 1 | 11 | 9.01E-04 |
| 0.693959199 | 0.693959199 | 1 | 28 | 0.002294142 |
| 0.697475289 | 0.697475289 | 4 | 7 | 5.74E-04 |
| 0.697475289 | 0.697475289 | 1 | 10 | 8.19E-04 |
| 0.697475289 | 0.697475289 | 2 | 24 | 0.001966407 |
| 0.699218287 | 0.699218287 | 9 | 9 | 7.37E-04 |
| 0.699218287 | 0.699218287 | 2 | 5 | 4.10E-04 |
| 0.702674474 | 0.702674474 | 3 | 7 | 5.74E-04 |
| 0.704387774 | 0.704387774 | 3 | 7 | 5.74E-04 |
| 0.704387774 | 0.704387774 | 1 | 5 | 4.10E-04 |
| 0.704387774 | 0.704387774 | 1 | 6 | 4.92E-04 |
| 0.704387774 | 0.704387774 | 1 | 12 | 9.83E-04 |
| 0.704387774 | 0.704387774 | 1 | 28 | 0.002294142 |
| 0.706091288 | 0.706091288 | 3 | 27 | 0.002212208 |
| 0.707785069 | 0.707785069 | 4 | 8 | 6.55E-04 |
| 0.709469174 | 0.709469174 | 3 | 7 | 5.74E-04 |
| 0.709881005 | 0.709881005 | 2 | 47 | 0.003850881 |
| 0.711143656 | 0.711143656 | 2 | 3 | 2.46E-04 |
| 0.711143656 | 0.711143656 | 3 | 15 | 0.001229005 |
| 0.712808571 | 0.712808571 | 3 | 14 | 0.001147071 |
| 0.712808571 | 0.712808571 | 1 | 16 | 0.001310938 |
| 0.713473891 | 0.713473891 | 4 | 4 | 3.28E-04 |
| 0.714463973 | 0.714463973 | 3 | 7 | 5.74E-04 |

| | | | | |
|---|---|---|---|---|
| 0.714463973 | 0.714463973 | 2 | 6 | 4.92E-04 |
| 0.714463973 | 0.714463973 | 3 | 14 | 0.001147071 |
| 0.714463973 | 0.714463973 | 3 | 40 | 0.003277345 |
| 0.715848619 | 0.715848619 | 23 | 36 | 0.002949611 |
| 0.716109916 | 0.716109916 | 6 | 9 | 7.37E-04 |
| 0.716109916 | 0.716109916 | 2 | 12 | 9.83E-04 |
| 0.717746452 | 0.717746452 | 6 | 9 | 7.37E-04 |
| 0.717746452 | 0.717746452 | 1 | 2 | 1.64E-04 |
| 0.717746452 | 0.717746452 | 2 | 11 | 9.01E-04 |
| 0.717746452 | 0.717746452 | 1 | 6 | 4.92E-04 |
| 0.717746452 | 0.717746452 | 4 | 56 | 0.004588283 |
| 0.718206992 | 0.718206992 | 4 | 9 | 7.37E-04 |
| 0.719373636 | 0.719373636 | 1 | 4 | 3.28E-04 |
| 0.720991521 | 0.720991521 | 1 | 5 | 4.10E-04 |
| 0.720991521 | 0.720991521 | 1 | 11 | 9.01E-04 |
| 0.720991521 | 0.720991521 | 1 | 15 | 0.001229005 |
| 0.720991521 | 0.720991521 | 2 | 50 | 0.004096682 |
| 0.722600159 | 0.722600159 | 6 | 9 | 7.37E-04 |
| 0.722600159 | 0.722600159 | 3 | 12 | 9.83E-04 |
| 0.724199602 | 0.724199602 | 3 | 14 | 0.001147071 |
| 0.724199602 | 0.724199602 | 3 | 15 | 0.001229005 |
| 0.725789902 | 0.725789902 | 2 | 9 | 7.37E-04 |
| 0.725789902 | 0.725789902 | 4 | 28 | 0.002294142 |
| 0.725789902 | 0.725789902 | 2 | 19 | 0.001556739 |
| 0.727371113 | 0.727371113 | 6 | 9 | 7.37E-04 |
| 0.730506467 | 0.730506467 | 31 | 52 | 0.004260549 |
| 0.730506467 | 0.730506467 | 2 | 8 | 6.55E-04 |
| 0.730506467 | 0.730506467 | 1 | 8 | 6.55E-04 |
| 0.732060713 | 0.732060713 | 2 | 11 | 9.01E-04 |
| 0.732060713 | 0.732060713 | 1 | 14 | 0.001147071 |
| 0.732060713 | 0.732060713 | 2 | 30 | 0.002458009 |
| 0.736670335 | 0.736670335 | 1 | 27 | 0.002212208 |
| 0.738189337 | 0.738189337 | 6 | 7 | 5.74E-04 |
| 0.738189337 | 0.738189337 | 2 | 9 | 7.37E-04 |
| 0.741201331 | 0.741201331 | 2 | 12 | 9.83E-04 |
| 0.742694421 | 0.742694421 | 1 | 7 | 5.74E-04 |
| 0.744178972 | 0.744178972 | 1 | 3 | 2.46E-04 |
| 0.744178972 | 0.744178972 | 2 | 47 | 0.003850881 |
| 0.745254763 | 0.745254763 | 8 | 77 | 0.00630889 |
| 0.745655031 | 0.745655031 | 3 | 17 | 0.001392872 |
| 0.745655031 | 0.745655031 | 4 | 27 | 0.002212208 |
| 0.747122647 | 0.747122647 | 3 | 19 | 0.001556739 |
| 0.748476323 | 0.748476323 | 2 | 53 | 0.004342483 |
| 0.748581868 | 0.748581868 | 2 | 23 | 0.001884474 |
| 0.750032742 | 0.750032742 | 2 | 49 | 0.004014748 |
| 0.751475315 | 0.751475315 | 6 | 26 | 0.002130274 |
| 0.751475315 | 0.751475315 | 3 | 16 | 0.001310938 |
| 0.752909635 | 0.752909635 | 6 | 11 | 9.01E-04 |
| 0.756898316 | 0.756898316 | 5 | 60 | 0.004916018 |
| 0.758565315 | 0.758565315 | 3 | 28 | 0.002294142 |

| | | | | |
|---|---|---|---|---|
| 0.759959066 | 0.759959066 | 2 | 28 | 0.002294142 |
| 0.759993926 | 0.759993926 | 3 | 46 | 0.003768947 |
| 0.761018275 | 0.761018275 | 13 | 18 | 0.001474805 |
| 0.76134484 | 0.76134484 | 7 | 18 | 0.001474805 |
| 0.76134484 | 0.76134484 | 3 | 22 | 0.00180254 |
| 0.76134484 | 0.76134484 | 3 | 22 | 0.00180254 |
| 0.762722684 | 0.762722684 | 7 | 21 | 0.001720606 |
| 0.764092642 | 0.764092642 | 1 | 14 | 0.001147071 |
| 0.768155644 | 0.768155644 | 19 | 35 | 0.002867677 |
| 0.770825696 | 0.770825696 | 2 | 11 | 9.01E-04 |
| 0.773465263 | 0.773465263 | 5 | 26 | 0.002130274 |
| 0.774773722 | 0.774773722 | 1 | 24 | 0.001966407 |
| 0.776074689 | 0.776074689 | 6 | 32 | 0.002621876 |
| 0.776074689 | 0.776074689 | 2 | 16 | 0.001310938 |
| 0.776074689 | 0.776074689 | 2 | 17 | 0.001392872 |
| 0.776074689 | 0.776074689 | 1 | 20 | 0.001638673 |
| 0.776074689 | 0.776074689 | 1 | 34 | 0.002785744 |
| 0.777431663 | 0.777431663 | 18 | 218 | 0.017861532 |
| 0.778654316 | 0.778654316 | 7 | 25 | 0.002048341 |
| 0.778654316 | 0.778654316 | 2 | 20 | 0.001638673 |
| 0.781204481 | 0.781204481 | 3 | 26 | 0.002130274 |
| 0.782468619 | 0.782468619 | 3 | 9 | 7.37E-04 |
| 0.782468619 | 0.782468619 | 6 | 30 | 0.002458009 |
| 0.784975215 | 0.784975215 | 1 | 7 | 5.74E-04 |
| 0.786217755 | 0.786217755 | 2 | 5 | 4.10E-04 |
| 0.786217755 | 0.786217755 | 2 | 26 | 0.002130274 |
| 0.786217755 | 0.786217755 | 1 | 14 | 0.001147071 |
| 0.787453177 | 0.787453177 | 2 | 19 | 0.001556739 |
| 0.789902827 | 0.789902827 | 3 | 37 | 0.003031544 |
| 0.791117136 | 0.791117136 | 2 | 15 | 0.001229005 |
| 0.791117136 | 0.791117136 | 1 | 54 | 0.004424416 |
| 0.792774745 | 0.792774745 | 8 | 50 | 0.004096682 |
| 0.794718475 | 0.794718475 | 2 | 23 | 0.001884474 |
| 0.794718475 | 0.794718475 | 2 | 23 | 0.001884474 |
| 0.797085103 | 0.797085103 | 3 | 25 | 0.002048341 |
| 0.799424681 | 0.799424681 | 3 | 27 | 0.002212208 |
| 0.800584422 | 0.800584422 | 2 | 7 | 5.74E-04 |
| 0.804023911 | 0.804023911 | 6 | 34 | 0.002785744 |
| 0.804023911 | 0.804023911 | 2 | 33 | 0.00270381 |
| 0.804023911 | 0.804023911 | 1 | 75 | 0.006145023 |
| 0.805157287 | 0.805157287 | 2 | 20 | 0.001638673 |
| 0.805157287 | 0.805157287 | 1 | 20 | 0.001638673 |
| 0.805157287 | 0.805157287 | 2 | 63 | 0.005161819 |
| 0.807265675 | 0.807265675 | 42 | 86 | 0.007046293 |
| 0.807404583 | 0.807404583 | 5 | 33 | 0.00270381 |
| 0.808518576 | 0.808518576 | 2 | 10 | 8.19E-04 |
| 0.809626182 | 0.809626182 | 2 | 20 | 0.001638673 |
| 0.809626182 | 0.809626182 | 2 | 90 | 0.007374027 |
| 0.810727436 | 0.810727436 | 4 | 27 | 0.002212208 |
| 0.810727436 | 0.810727436 | 1 | 17 | 0.001392872 |

| | | | | |
|---|---|---|---|---|
| 0.810727436 | 0.810727436 | 1 | 34 | 0.002785744 |
| 0.811822374 | 0.811822374 | 6 | 22 | 0.00180254 |
| 0.812911033 | 0.812911033 | 2 | 19 | 0.001556739 |
| 0.812911033 | 0.812911033 | 2 | 35 | 0.002867677 |
| 0.815069656 | 0.815069656 | 8 | 39 | 0.003195412 |
| 0.815069656 | 0.815069656 | 12 | 60 | 0.004916018 |
| 0.815069656 | 0.815069656 | 3 | 19 | 0.001556739 |
| 0.815069656 | 0.815069656 | 2 | 13 | 0.001065137 |
| 0.815069656 | 0.815069656 | 2 | 18 | 0.001474805 |
| 0.815954447 | 0.815954447 | 13 | 59 | 0.004834084 |
| 0.816767183 | 0.816767183 | 9 | 80 | 0.006554691 |
| 0.817203586 | 0.817203586 | 1 | 12 | 9.83E-04 |
| 0.81826138 | 0.81826138 | 4 | 39 | 0.003195412 |
| 0.81826138 | 0.81826138 | 1 | 21 | 0.001720606 |
| 0.81826138 | 0.81826138 | 1 | 21 | 0.001720606 |
| 0.81826138 | 0.81826138 | 1 | 21 | 0.001720606 |
| 0.81826138 | 0.81826138 | 1 | 21 | 0.001720606 |
| 0.821398489 | 0.821398489 | 1 | 32 | 0.002621876 |
| 0.822432215 | 0.822432215 | 3 | 38 | 0.003113478 |
| 0.822432215 | 0.822432215 | 1 | 25 | 0.002048341 |
| 0.825497942 | 0.825497942 | 3 | 34 | 0.002785744 |
| 0.825497942 | 0.825497942 | 1 | 29 | 0.002376075 |
| 0.825497942 | 0.825497942 | 2 | 114 | 0.009340434 |
| 0.826508145 | 0.826508145 | 13 | 36 | 0.002949611 |
| 0.826508145 | 0.826508145 | 2 | 12 | 9.83E-04 |
| 0.827046943 | 0.827046943 | 6 | 60 | 0.004916018 |
| 0.828511191 | 0.828511191 | 3 | 32 | 0.002621876 |
| 0.83049131 | 0.83049131 | 4 | 22 | 0.00180254 |
| 0.83049131 | 0.83049131 | 2 | 14 | 0.001147071 |
| 0.830862045 | 0.830862045 | 15 | 48 | 0.003932814 |
| 0.831472853 | 0.831472853 | 3 | 24 | 0.001966407 |
| 0.832366955 | 0.832366955 | 19 | 50 | 0.004096682 |
| 0.833341594 | 0.833341594 | 48 | 157 | 0.01286358 |
| 0.833419067 | 0.833419067 | 3 | 32 | 0.002621876 |
| 0.833419067 | 0.833419067 | 1 | 19 | 0.001556739 |
| 0.835343 | 0.835343 | 3 | 20 | 0.001638673 |
| 0.837244902 | 0.837244902 | 4 | 16 | 0.001310938 |
| 0.839125025 | 0.839125025 | 2 | 7 | 5.74E-04 |
| 0.839125025 | 0.839125025 | 1 | 6 | 4.92E-04 |
| 0.839125025 | 0.839125025 | 2 | 18 | 0.001474805 |
| 0.841904914 | 0.841904914 | 2 | 17 | 0.001392872 |
| 0.843731664 | 0.843731664 | 3 | 16 | 0.001310938 |
| 0.844637177 | 0.844637177 | 6 | 38 | 0.003113478 |
| 0.845537488 | 0.845537488 | 6 | 65 | 0.005325686 |
| 0.849087305 | 0.849087305 | 10 | 106 | 0.008684965 |
| 0.849513948 | 0.849513948 | 10 | 66 | 0.00540762 |
| 0.849962049 | 0.849962049 | 1 | 8 | 6.55E-04 |
| 0.854494525 | 0.854494525 | 35 | 105 | 0.008603032 |
| 0.85510595 | 0.85510595 | 4 | 16 | 0.001310938 |
| 0.85510595 | 0.85510595 | 1 | 34 | 0.002785744 |

| | | | | |
|---|---|---|---|---|
| 0.858437698 | 0.858437698 | 4 | 9 | 7.37E-04 |
| 0.858437698 | 0.858437698 | 14 | 33 | 0.00270381 |
| 0.858437698 | 0.858437698 | 1 | 135 | 0.011061041 |
| 0.861320148 | 0.861320148 | 10 | 77 | 0.00630889 |
| 0.861693486 | 0.861693486 | 7 | 13 | 0.001065137 |
| 0.861693486 | 0.861693486 | 2 | 24 | 0.001966407 |
| 0.861693486 | 0.861693486 | 2 | 24 | 0.001966407 |
| 0.862295101 | 0.862295101 | 5 | 94 | 0.007701762 |
| 0.862578826 | 0.862578826 | 25 | 78 | 0.006390823 |
| 0.863293433 | 0.863293433 | 2 | 14 | 0.001147071 |
| 0.86382714 | 0.86382714 | 25 | 80 | 0.006554691 |
| 0.864086512 | 0.864086512 | 7 | 62 | 0.005079885 |
| 0.865680331 | 0.865680331 | 12 | 85 | 0.006964359 |
| 0.86798401 | 0.86798401 | 2 | 17 | 0.001392872 |
| 0.86798401 | 0.86798401 | 1 | 24 | 0.001966407 |
| 0.870269112 | 0.870269112 | 8 | 38 | 0.003113478 |
| 0.872514998 | 0.872514998 | 29 | 70 | 0.005735354 |
| 0.872514998 | 0.872514998 | 2 | 27 | 0.002212208 |
| 0.874029223 | 0.874029223 | 10 | 83 | 0.006800492 |
| 0.874722337 | 0.874722337 | 3 | 54 | 0.004424416 |
| 0.875182548 | 0.875182548 | 7 | 84 | 0.006882425 |
| 0.875182548 | 0.875182548 | 7 | 84 | 0.006882425 |
| 0.876172804 | 0.876172804 | 10 | 33 | 0.00270381 |
| 0.876891783 | 0.876891783 | 1 | 7 | 5.74E-04 |
| 0.876891783 | 0.876891783 | 3 | 28 | 0.002294142 |
| 0.876891783 | 0.876891783 | 1 | 40 | 0.003277345 |
| 0.877606623 | 0.877606623 | 8 | 28 | 0.002294142 |
| 0.877606623 | 0.877606623 | 2 | 24 | 0.001966407 |
| 0.877819325 | 0.877819325 | 33 | 289 | 0.02367882 |
| 0.878317348 | 0.878317348 | 2 | 20 | 0.001638673 |
| 0.878566193 | 0.878566193 | 11 | 117 | 0.009586235 |
| 0.881119567 | 0.881119567 | 2 | 11 | 9.01E-04 |
| 0.881810067 | 0.881810067 | 7 | 37 | 0.003031544 |
| 0.882496591 | 0.882496591 | 2 | 14 | 0.001147071 |
| 0.883179161 | 0.883179161 | 3 | 37 | 0.003031544 |
| 0.884532533 | 0.884532533 | 1 | 45 | 0.003687014 |
| 0.885203379 | 0.885203379 | 3 | 36 | 0.002949611 |
| 0.886533501 | 0.886533501 | 3 | 18 | 0.001474805 |
| 0.886533501 | 0.886533501 | 3 | 18 | 0.001474805 |
| 0.888807482 | 0.888807482 | 12 | 61 | 0.004997952 |
| 0.88979234 | 0.88979234 | 2 | 144 | 0.011798443 |
| 0.890432886 | 0.890432886 | 8 | 34 | 0.002785744 |
| 0.891069741 | 0.891069741 | 3 | 3 | 2.46E-04 |
| 0.891069741 | 0.891069741 | 14 | 30 | 0.002458009 |
| 0.891069741 | 0.891069741 | 2 | 27 | 0.002212208 |
| 0.893580674 | 0.893580674 | 4 | 24 | 0.001966407 |
| 0.894814539 | 0.894814539 | 2 | 20 | 0.001638673 |
| 0.89603422 | 0.89603422 | 1 | 51 | 0.004178615 |
| 0.896638793 | 0.896638793 | 10 | 23 | 0.001884474 |
| 0.896638793 | 0.896638793 | 5 | 31 | 0.002539943 |

| | | | | |
|---|---|---|---|---|
| 0.896638793 | 0.896638793 | 3 | 42 | 0.003441213 |
| 0.897239881 | 0.897239881 | 10 | 23 | 0.001884474 |
| 0.897239881 | 0.897239881 | 5 | 48 | 0.003932814 |
| 0.897239881 | 0.897239881 | 1 | 39 | 0.003195412 |
| 0.897837503 | 0.897837503 | 2 | 17 | 0.001392872 |
| 0.898431679 | 0.898431679 | 6 | 42 | 0.003441213 |
| 0.89902243 | 0.89902243 | 2 | 75 | 0.006145023 |
| 0.89902243 | 0.89902243 | 1 | 41 | 0.003359279 |
| 0.900193732 | 0.900193732 | 10 | 23 | 0.001884474 |
| 0.900774323 | 0.900774323 | 10 | 24 | 0.001966407 |
| 0.900774323 | 0.900774323 | 5 | 43 | 0.003523146 |
| 0.902894525 | 0.902894525 | 2 | 95 | 0.007783695 |
| 0.903063392 | 0.903063392 | 2 | 65 | 0.005325686 |
| 0.905300104 | 0.905300104 | 4 | 58 | 0.004752151 |
| 0.908657995 | 0.908657995 | 12 | 89 | 0.007292093 |
| 0.910147375 | 0.910147375 | 1 | 38 | 0.003113478 |
| 0.912221983 | 0.912221983 | 4 | 38 | 0.003113478 |
| 0.914748583 | 0.914748583 | 4 | 18 | 0.001474805 |
| 0.915245183 | 0.915245183 | 1 | 15 | 0.001229005 |
| 0.916022296 | 0.916022296 | 28 | 71 | 0.005817288 |
| 0.916229797 | 0.916229797 | 3 | 27 | 0.002212208 |
| 0.916717843 | 0.916717843 | 1 | 133 | 0.010897173 |
| 0.919245305 | 0.919245305 | 19 | 121 | 0.00991397 |
| 0.920522082 | 0.920522082 | 2 | 59 | 0.004834084 |
| 0.920985332 | 0.920985332 | 5 | 77 | 0.00630889 |
| 0.923261739 | 0.923261739 | 10 | 14 | 0.001147071 |
| 0.924153989 | 0.924153989 | 1 | 40 | 0.003277345 |
| 0.92459625 | 0.92459625 | 12 | 13 | 0.001065137 |
| 0.92624704 | 0.92624704 | 7 | 144 | 0.011798443 |
| 0.926339873 | 0.926339873 | 5 | 54 | 0.004424416 |
| 0.928160036 | 0.928160036 | 65 | 218 | 0.017861532 |
| 0.932529765 | 0.932529765 | 2 | 43 | 0.003523146 |
| 0.932923586 | 0.932923586 | 5 | 61 | 0.004997952 |
| 0.933704404 | 0.933704404 | 11 | 49 | 0.004014748 |
| 0.934091427 | 0.934091427 | 5 | 115 | 0.009422368 |
| 0.934858767 | 0.934858767 | 3 | 56 | 0.004588283 |
| 0.936157501 | 0.936157501 | 2 | 182 | 0.014911921 |
| 0.936366977 | 0.936366977 | 9 | 59 | 0.004834084 |
| 0.936738589 | 0.936738589 | 3 | 46 | 0.003768947 |
| 0.937840565 | 0.937840565 | 2 | 10 | 8.19E-04 |
| 0.938203644 | 0.938203644 | 1 | 57 | 0.004670217 |
| 0.939635059 | 0.939635059 | 4 | 42 | 0.003441213 |
| 0.939987747 | 0.939987747 | 4 | 52 | 0.004260549 |
| 0.9410336 | 0.9410336 | 23 | 47 | 0.003850881 |
| 0.9410336 | 0.9410336 | 6 | 37 | 0.003031544 |
| 0.944807392 | 0.944807392 | 4 | 86 | 0.007046293 |
| 0.945075771 | 0.945075771 | 9 | 9 | 7.37E-04 |
| 0.945360746 | 0.945360746 | 8 | 22 | 0.00180254 |
| 0.945997915 | 0.945997915 | 1 | 58 | 0.004752151 |
| 0.945997915 | 0.945997915 | 1 | 60 | 0.004916018 |

| | | | | |
|---|---|---|---|---|
| 0.946939883 | 0.946939883 | 4 | 51 | 0.004178615 |
| 0.947558794 | 0.947558794 | 4 | 58 | 0.004752151 |
| 0.947558794 | 0.947558794 | 3 | 67 | 0.005489553 |
| 0.948170548 | 0.948170548 | 4 | 25 | 0.002048341 |
| 0.948454571 | 0.948454571 | 10 | 14 | 0.001147071 |
| 0.948473767 | 0.948473767 | 13 | 41 | 0.003359279 |
| 0.948956636 | 0.948956636 | 5 | 54 | 0.004424416 |
| 0.949074939 | 0.949074939 | 2 | 52 | 0.004260549 |
| 0.949372912 | 0.949372912 | 13 | 43 | 0.003523146 |
| 0.949972631 | 0.949972631 | 21 | 269 | 0.022040147 |
| 0.950677722 | 0.950677722 | 8 | 77 | 0.00630889 |
| 0.950968749 | 0.950968749 | 15 | 184 | 0.015075789 |
| 0.951410881 | 0.951410881 | 2 | 8 | 6.55E-04 |
| 0.951978072 | 0.951978072 | 3 | 7 | 5.74E-04 |
| 0.951978072 | 0.951978072 | 13 | 48 | 0.003932814 |
| 0.952538699 | 0.952538699 | 1 | 42 | 0.003441213 |
| 0.953092838 | 0.953092838 | 3 | 15 | 0.001229005 |
| 0.953640562 | 0.953640562 | 6 | 38 | 0.003113478 |
| 0.953912042 | 0.953912042 | 1 | 50 | 0.004096682 |
| 0.954450282 | 0.954450282 | 3 | 63 | 0.005161819 |
| 0.954982291 | 0.954982291 | 16 | 35 | 0.002867677 |
| 0.954982291 | 0.954982291 | 5 | 26 | 0.002130274 |
| 0.955508139 | 0.955508139 | 5 | 35 | 0.002867677 |
| 0.955508139 | 0.955508139 | 7 | 56 | 0.004588283 |
| 0.956285516 | 0.956285516 | 3 | 34 | 0.002785744 |
| 0.957049426 | 0.957049426 | 3 | 11 | 9.01E-04 |
| 0.957049426 | 0.957049426 | 3 | 82 | 0.006718558 |
| 0.957800101 | 0.957800101 | 6 | 43 | 0.003523146 |
| 0.961362768 | 0.961362768 | 4 | 60 | 0.004916018 |
| 0.961943268 | 0.961943268 | 65 | 251 | 0.020565342 |
| 0.964059245 | 0.964059245 | 16 | 114 | 0.009340434 |
| 0.964208145 | 0.964208145 | 3 | 15 | 0.001229005 |
| 0.964944509 | 0.964944509 | 12 | 64 | 0.005243753 |
| 0.968372017 | 0.968372017 | 13 | 46 | 0.003768947 |
| 0.970705115 | 0.970705115 | 21 | 98 | 0.008029496 |
| 0.971388052 | 0.971388052 | 3 | 205 | 0.016796395 |
| 0.971723578 | 0.971723578 | 3 | 14 | 0.001147071 |
| 0.972055203 | 0.972055203 | 49 | 142 | 0.011634576 |
| 0.972867494 | 0.972867494 | 9 | 84 | 0.006882425 |
| 0.972867494 | 0.972867494 | 2 | 43 | 0.003523146 |
| 0.973136192 | 0.973136192 | 12 | 78 | 0.006390823 |
| 0.974331404 | 0.974331404 | 8 | 112 | 0.009176567 |
| 0.975019479 | 0.975019479 | 3 | 63 | 0.005161819 |
| 0.975386431 | 0.975386431 | 133 | 648 | 0.053092995 |
| 0.975723174 | 0.975723174 | 8 | 66 | 0.00540762 |
| 0.976172294 | 0.976172294 | 1 | 68 | 0.005571487 |
| 0.977158963 | 0.977158963 | 21 | 32 | 0.002621876 |
| 0.97767176 | 0.97767176 | 3 | 21 | 0.001720606 |
| 0.977934161 | 0.977934161 | 3 | 114 | 0.009340434 |
| 0.978576863 | 0.978576863 | 4 | 243 | 0.019909873 |

| | | | | |
|---|---|---|---|---|
| 0.978577058 | 0.978577058 | 26 | 261 | 0.021384678 |
| 0.97968731 | 0.97968731 | 5 | 48 | 0.003932814 |
| 0.980842221 | 0.980842221 | 40 | 156 | 0.012781647 |
| 0.980854317 | 0.980854317 | 12 | 99 | 0.00811143 |
| 0.981954827 | 0.981954827 | 6 | 19 | 0.001556739 |
| 0.982167283 | 0.982167283 | 5 | 54 | 0.004424416 |
| 0.98289154 | 0.98289154 | 6 | 82 | 0.006718558 |
| 0.98443932 | 0.98443932 | 6 | 29 | 0.002376075 |
| 0.985422065 | 0.985422065 | 1 | 107 | 0.008766899 |
| 0.985508293 | 0.985508293 | 3 | 18 | 0.001474805 |
| 0.985931911 | 0.985931911 | 5 | 43 | 0.003523146 |
| 0.985931911 | 0.985931911 | 5 | 43 | 0.003523146 |
| 0.986261947 | 0.986261947 | 9 | 53 | 0.004342483 |
| 0.986261947 | 0.986261947 | 9 | 53 | 0.004342483 |
| 0.986439318 | 0.986439318 | 59 | 322 | 0.02638263 |
| 0.988707581 | 0.988707581 | 12 | 56 | 0.004588283 |
| 0.990885284 | 0.990885284 | 6 | 107 | 0.008766899 |
| 0.991774877 | 0.991774877 | 14 | 90 | 0.007374027 |
| 0.99181241 | 0.99181241 | 2 | 150 | 0.012290045 |
| 0.991861082 | 0.991861082 | 3 | 318 | 0.026054896 |
| 0.992193851 | 0.992193851 | 5 | 44 | 0.00360508 |
| 0.993778393 | 0.993778393 | 5 | 22 | 0.00180254 |
| 0.993778393 | 0.993778393 | 5 | 22 | 0.00180254 |
| 0.994277953 | 0.994277953 | 4 | 191 | 0.015649324 |
| 0.994503585 | 0.994503585 | 30 | 259 | 0.021220811 |
| 0.99456172 | 0.99456172 | 34 | 551 | 0.045145432 |
| 0.994642388 | 0.994642388 | 15 | 152 | 0.012453912 |
| 0.995247027 | 0.995247027 | 13 | 139 | 0.011388775 |
| 0.995616154 | 0.995616154 | 140 | 872 | 0.071446129 |
| 0.995681612 | 0.995681612 | 13 | 152 | 0.012453912 |
| 0.995758597 | 0.995758597 | 5 | 284 | 0.023269152 |
| 0.995932977 | 0.995932977 | 5 | 66 | 0.00540762 |
| 0.995981481 | 0.995981481 | 9 | 158 | 0.012945514 |
| 0.996683943 | 0.996683943 | 34 | 234 | 0.01917247 |
| 0.997560257 | 0.997560257 | 2 | 177 | 0.014502253 |
| 0.997986323 | 0.997986323 | 132 | 811 | 0.066448177 |
| 0.998129004 | 0.998129004 | 10 | 162 | 0.013273249 |
| 0.998383655 | 0.998383655 | 145 | 983 | 0.080540762 |
| 0.998797075 | 0.998797075 | 18 | 98 | 0.008029496 |
| 0.999073957 | 0.999073957 | 39 | 327 | 0.026792298 |
| 0.999164372 | 0.999164372 | 3 | 326 | 0.026710365 |
| 0.999182458 | 0.999182458 | 67 | 410 | 0.03359279 |
| 0.999199512 | 0.999199512 | 35 | 297 | 0.024334289 |
| 0.999227916 | 0.999227916 | 9 | 110 | 0.0090127 |
| 0.999425131 | 0.999425131 | 44 | 699 | 0.05727161 |
| 0.999456228 | 0.999456228 | 56 | 942 | 0.077181483 |
| 0.999480522 | 0.999480522 | 9 | 131 | 0.010733306 |
| 0.999728247 | 0.999728247 | 35 | 524 | 0.042933224 |
| 0.999904909 | 0.999904909 | 20 | 329 | 0.026956166 |
| 0.999938046 | 0.999938046 | 17 | 130 | 0.010651372 |

| | | | | |
|---|---|---|---|---|
| 0.999957637 | 0.999957637 | 17 | 139 | 0.011388775 |
| 0.99996001 | 0.99996001 | 25 | 255 | 0.020893077 |
| 0.99996825 | 0.99996825 | 22 | 685 | 0.056124539 |
| 0.999970423 | 0.999970423 | 39 | 187 | 0.01532159 |
| 0.999983418 | 0.999983418 | 91 | 1586 | 0.129946743 |
| 0.999983965 | 0.999983965 | 6 | 437 | 0.035804998 |
| 0.999985621 | 0.999985621 | 64 | 511 | 0.041868087 |
| 0.999987803 | 0.999987803 | 61 | 890 | 0.072920934 |
| 0.999996951 | 0.999996951 | 27 | 432 | 0.03539533 |
| 0.999997995 | 0.999997995 | 61 | 959 | 0.078574355 |
| 0.999999785 | 0.999999785 | 333 | 2230 | 0.182712003 |
| 0.999999999 | 0.999999999 | 81 | 2218 | 0.1817288 |

| Species identifier | Species name |
|---|---|
| 9606 | Homo sapiens |
| 9606 | Homo sapiens |
| 9606 | Homo sapiens |
| 9606 | Homo sapiens |
| 9606 | Homo sapiens |
| 9606 | Homo sapiens |
| 9606 | Homo sapiens |
| 9606 | Homo sapiens |
| 9606 | Homo sapiens |
| 9606 | Homo sapiens |
| 9606 | Homo sapiens |
| 9606 | Homo sapiens |
| 9606 | Homo sapiens |
| 9606 | Homo sapiens |
| 9606 | Homo sapiens |
| 9606 | Homo sapiens |
| 9606 | Homo sapiens |
| 9606 | Homo sapiens |
| 9606 | Homo sapiens |
| 9606 | Homo sapiens |
| 9606 | Homo sapiens |
| 9606 | Homo sapiens |
| 9606 | Homo sapiens |
| 9606 | Homo sapiens |
| 9606 | Homo sapiens |
| 9606 | Homo sapiens |
| 9606 | Homo sapiens |
| 9606 | Homo sapiens |
| 9606 | Homo sapiens |
| 9606 | Homo sapiens |
| 9606 | Homo sapiens |
| 9606 | Homo sapiens |
| 9606 | Homo sapiens |
| 9606 | Homo sapiens |
| 9606 | Homo sapiens |
| 9606 | Homo sapiens |
| 9606 | Homo sapiens |
| 9606 | Homo sapiens |
| 9606 | Homo sapiens |
| 9606 | Homo sapiens |
| 9606 | Homo sapiens |
| 9606 | Homo sapiens |
| 9606 | Homo sapiens |
| 9606 | Homo sapiens |
| 9606 | Homo sapiens |
| 9606 | Homo sapiens |
| 9606 | Homo sapiens |
| 9606 | Homo sapiens |
| 9606 | Homo sapiens |

| | |
|---|---|
| 9606 | Homo sapiens |
| 9606 | Homo sapiens |
| 9606 | Homo sapiens |
| 9606 | Homo sapiens |
| 9606 | Homo sapiens |
| 9606 | Homo sapiens |
| 9606 | Homo sapiens |
| 9606 | Homo sapiens |
| 9606 | Homo sapiens |
| 9606 | Homo sapiens |
| 9606 | Homo sapiens |
| 9606 | Homo sapiens |
| 9606 | Homo sapiens |
| 9606 | Homo sapiens |
| 9606 | Homo sapiens |
| 9606 | Homo sapiens |
| 9606 | Homo sapiens |
| 9606 | Homo sapiens |
| 9606 | Homo sapiens |
| 9606 | Homo sapiens |
| 9606 | Homo sapiens |
| 9606 | Homo sapiens |
| 9606 | Homo sapiens |
| 9606 | Homo sapiens |
| 9606 | Homo sapiens |
| 9606 | Homo sapiens |
| 9606 | Homo sapiens |
| 9606 | Homo sapiens |
| 9606 | Homo sapiens |
| 9606 | Homo sapiens |
| 9606 | Homo sapiens |
| 9606 | Homo sapiens |
| 9606 | Homo sapiens |
| 9606 | Homo sapiens |
| 9606 | Homo sapiens |
| 9606 | Homo sapiens |
| 9606 | Homo sapiens |
| 9606 | Homo sapiens |
| 9606 | Homo sapiens |
| 9606 | Homo sapiens |
| 9606 | Homo sapiens |
| 9606 | Homo sapiens |
| 9606 | Homo sapiens |
| 9606 | Homo sapiens |
| 9606 | Homo sapiens |
| 9606 | Homo sapiens |
| 9606 | Homo sapiens |
| 9606 | Homo sapiens |
| 9606 | Homo sapiens |
| 9606 | Homo sapiens |

| | |
|---|---|
| 9606 | Homo sapiens |
| 9606 | Homo sapiens |
| 9606 | Homo sapiens |
| 9606 | Homo sapiens |
| 9606 | Homo sapiens |
| 9606 | Homo sapiens |
| 9606 | Homo sapiens |
| 9606 | Homo sapiens |
| 9606 | Homo sapiens |
| 9606 | Homo sapiens |
| 9606 | Homo sapiens |
| 9606 | Homo sapiens |
| 9606 | Homo sapiens |
| 9606 | Homo sapiens |
| 9606 | Homo sapiens |
| 9606 | Homo sapiens |
| 9606 | Homo sapiens |
| 9606 | Homo sapiens |
| 9606 | Homo sapiens |
| 9606 | Homo sapiens |
| 9606 | Homo sapiens |
| 9606 | Homo sapiens |
| 9606 | Homo sapiens |
| 9606 | Homo sapiens |
| 9606 | Homo sapiens |
| 9606 | Homo sapiens |
| 9606 | Homo sapiens |
| 9606 | Homo sapiens |
| 9606 | Homo sapiens |
| 9606 | Homo sapiens |
| 9606 | Homo sapiens |
| 9606 | Homo sapiens |
| 9606 | Homo sapiens |
| 9606 | Homo sapiens |
| 9606 | Homo sapiens |
| 9606 | Homo sapiens |
| 9606 | Homo sapiens |
| 9606 | Homo sapiens |
| 9606 | Homo sapiens |
| 9606 | Homo sapiens |
| 9606 | Homo sapiens |
| 9606 | Homo sapiens |
| 9606 | Homo sapiens |
| 9606 | Homo sapiens |
| 9606 | Homo sapiens |
| 9606 | Homo sapiens |
| 9606 | Homo sapiens |
| 9606 | Homo sapiens |
| 9606 | Homo sapiens |
| 9606 | Homo sapiens |

| | |
|---|---|
| 9606 | Homo sapiens |
| 9606 | Homo sapiens |
| 9606 | Homo sapiens |
| 9606 | Homo sapiens |
| 9606 | Homo sapiens |
| 9606 | Homo sapiens |
| 9606 | Homo sapiens |
| 9606 | Homo sapiens |
| 9606 | Homo sapiens |
| 9606 | Homo sapiens |
| 9606 | Homo sapiens |
| 9606 | Homo sapiens |
| 9606 | Homo sapiens |
| 9606 | Homo sapiens |
| 9606 | Homo sapiens |
| 9606 | Homo sapiens |
| 9606 | Homo sapiens |
| 9606 | Homo sapiens |
| 9606 | Homo sapiens |
| 9606 | Homo sapiens |
| 9606 | Homo sapiens |
| 9606 | Homo sapiens |
| 9606 | Homo sapiens |
| 9606 | Homo sapiens |
| 9606 | Homo sapiens |
| 9606 | Homo sapiens |
| 9606 | Homo sapiens |
| 9606 | Homo sapiens |
| 9606 | Homo sapiens |
| 9606 | Homo sapiens |
| 9606 | Homo sapiens |
| 9606 | Homo sapiens |
| 9606 | Homo sapiens |
| 9606 | Homo sapiens |
| 9606 | Homo sapiens |
| 9606 | Homo sapiens |
| 9606 | Homo sapiens |
| 9606 | Homo sapiens |
| 9606 | Homo sapiens |
| 9606 | Homo sapiens |
| 9606 | Homo sapiens |
| 9606 | Homo sapiens |
| 9606 | Homo sapiens |
| 9606 | Homo sapiens |
| 9606 | Homo sapiens |
| 9606 | Homo sapiens |
| 9606 | Homo sapiens |
| 9606 | Homo sapiens |
| 9606 | Homo sapiens |
| 9606 | Homo sapiens |

| | |
|---|---|
| 9606 | Homo sapiens |
| 9606 | Homo sapiens |
| 9606 | Homo sapiens |
| 9606 | Homo sapiens |
| 9606 | Homo sapiens |
| 9606 | Homo sapiens |
| 9606 | Homo sapiens |
| 9606 | Homo sapiens |
| 9606 | Homo sapiens |
| 9606 | Homo sapiens |
| 9606 | Homo sapiens |
| 9606 | Homo sapiens |
| 9606 | Homo sapiens |
| 9606 | Homo sapiens |
| 9606 | Homo sapiens |
| 9606 | Homo sapiens |
| 9606 | Homo sapiens |
| 9606 | Homo sapiens |
| 9606 | Homo sapiens |
| 9606 | Homo sapiens |
| 9606 | Homo sapiens |
| 9606 | Homo sapiens |
| 9606 | Homo sapiens |
| 9606 | Homo sapiens |
| 9606 | Homo sapiens |
| 9606 | Homo sapiens |
| 9606 | Homo sapiens |
| 9606 | Homo sapiens |
| 9606 | Homo sapiens |
| 9606 | Homo sapiens |
| 9606 | Homo sapiens |
| 9606 | Homo sapiens |
| 9606 | Homo sapiens |
| 9606 | Homo sapiens |
| 9606 | Homo sapiens |
| 9606 | Homo sapiens |
| 9606 | Homo sapiens |
| 9606 | Homo sapiens |
| 9606 | Homo sapiens |
| 9606 | Homo sapiens |
| 9606 | Homo sapiens |
| 9606 | Homo sapiens |
| 9606 | Homo sapiens |
| 9606 | Homo sapiens |
| 9606 | Homo sapiens |
| 9606 | Homo sapiens |
| 9606 | Homo sapiens |
| 9606 | Homo sapiens |
| 9606 | Homo sapiens |
| 9606 | Homo sapiens |

| | |
|---|---|
| 9606 | Homo sapiens |
| 9606 | Homo sapiens |
| 9606 | Homo sapiens |
| 9606 | Homo sapiens |
| 9606 | Homo sapiens |
| 9606 | Homo sapiens |
| 9606 | Homo sapiens |
| 9606 | Homo sapiens |
| 9606 | Homo sapiens |
| 9606 | Homo sapiens |
| 9606 | Homo sapiens |
| 9606 | Homo sapiens |
| 9606 | Homo sapiens |
| 9606 | Homo sapiens |
| 9606 | Homo sapiens |
| 9606 | Homo sapiens |
| 9606 | Homo sapiens |
| 9606 | Homo sapiens |
| 9606 | Homo sapiens |
| 9606 | Homo sapiens |
| 9606 | Homo sapiens |
| 9606 | Homo sapiens |
| 9606 | Homo sapiens |
| 9606 | Homo sapiens |
| 9606 | Homo sapiens |
| 9606 | Homo sapiens |
| 9606 | Homo sapiens |
| 9606 | Homo sapiens |
| 9606 | Homo sapiens |
| 9606 | Homo sapiens |
| 9606 | Homo sapiens |
| 9606 | Homo sapiens |
| 9606 | Homo sapiens |
| 9606 | Homo sapiens |
| 9606 | Homo sapiens |
| 9606 | Homo sapiens |
| 9606 | Homo sapiens |
| 9606 | Homo sapiens |
| 9606 | Homo sapiens |
| 9606 | Homo sapiens |
| 9606 | Homo sapiens |
| 9606 | Homo sapiens |
| 9606 | Homo sapiens |
| 9606 | Homo sapiens |
| 9606 | Homo sapiens |
| 9606 | Homo sapiens |
| 9606 | Homo sapiens |
| 9606 | Homo sapiens |
| 9606 | Homo sapiens |
| 9606 | Homo sapiens |

| | |
|---|---|
| 9606 | Homo sapiens |
| 9606 | Homo sapiens |
| 9606 | Homo sapiens |
| 9606 | Homo sapiens |
| 9606 | Homo sapiens |
| 9606 | Homo sapiens |
| 9606 | Homo sapiens |
| 9606 | Homo sapiens |
| 9606 | Homo sapiens |
| 9606 | Homo sapiens |
| 9606 | Homo sapiens |
| 9606 | Homo sapiens |
| 9606 | Homo sapiens |
| 9606 | Homo sapiens |
| 9606 | Homo sapiens |
| 9606 | Homo sapiens |
| 9606 | Homo sapiens |
| 9606 | Homo sapiens |
| 9606 | Homo sapiens |
| 9606 | Homo sapiens |
| 9606 | Homo sapiens |
| 9606 | Homo sapiens |
| 9606 | Homo sapiens |
| 9606 | Homo sapiens |
| 9606 | Homo sapiens |
| 9606 | Homo sapiens |
| 9606 | Homo sapiens |
| 9606 | Homo sapiens |
| 9606 | Homo sapiens |
| 9606 | Homo sapiens |
| 9606 | Homo sapiens |
| 9606 | Homo sapiens |
| 9606 | Homo sapiens |
| 9606 | Homo sapiens |
| 9606 | Homo sapiens |
| 9606 | Homo sapiens |
| 9606 | Homo sapiens |
| 9606 | Homo sapiens |
| 9606 | Homo sapiens |
| 9606 | Homo sapiens |
| 9606 | Homo sapiens |
| 9606 | Homo sapiens |
| 9606 | Homo sapiens |
| 9606 | Homo sapiens |
| 9606 | Homo sapiens |
| 9606 | Homo sapiens |
| 9606 | Homo sapiens |
| 9606 | Homo sapiens |
| 9606 | Homo sapiens |
| 9606 | Homo sapiens |

| | |
|---|---|
| 9606 | Homo sapiens |
| 9606 | Homo sapiens |
| 9606 | Homo sapiens |
| 9606 | Homo sapiens |
| 9606 | Homo sapiens |
| 9606 | Homo sapiens |
| 9606 | Homo sapiens |
| 9606 | Homo sapiens |
| 9606 | Homo sapiens |
| 9606 | Homo sapiens |
| 9606 | Homo sapiens |
| 9606 | Homo sapiens |
| 9606 | Homo sapiens |
| 9606 | Homo sapiens |
| 9606 | Homo sapiens |
| 9606 | Homo sapiens |
| 9606 | Homo sapiens |
| 9606 | Homo sapiens |
| 9606 | Homo sapiens |
| 9606 | Homo sapiens |
| 9606 | Homo sapiens |
| 9606 | Homo sapiens |
| 9606 | Homo sapiens |
| 9606 | Homo sapiens |
| 9606 | Homo sapiens |
| 9606 | Homo sapiens |
| 9606 | Homo sapiens |
| 9606 | Homo sapiens |
| 9606 | Homo sapiens |
| 9606 | Homo sapiens |
| 9606 | Homo sapiens |
| 9606 | Homo sapiens |
| 9606 | Homo sapiens |
| 9606 | Homo sapiens |
| 9606 | Homo sapiens |
| 9606 | Homo sapiens |
| 9606 | Homo sapiens |
| 9606 | Homo sapiens |
| 9606 | Homo sapiens |
| 9606 | Homo sapiens |
| 9606 | Homo sapiens |
| 9606 | Homo sapiens |
| 9606 | Homo sapiens |
| 9606 | Homo sapiens |
| 9606 | Homo sapiens |
| 9606 | Homo sapiens |
| 9606 | Homo sapiens |
| 9606 | Homo sapiens |
| 9606 | Homo sapiens |
| 9606 | Homo sapiens |

| | |
|---|---|
| 9606 | Homo sapiens |
| 9606 | Homo sapiens |
| 9606 | Homo sapiens |
| 9606 | Homo sapiens |
| 9606 | Homo sapiens |
| 9606 | Homo sapiens |
| 9606 | Homo sapiens |
| 9606 | Homo sapiens |
| 9606 | Homo sapiens |
| 9606 | Homo sapiens |
| 9606 | Homo sapiens |
| 9606 | Homo sapiens |
| 9606 | Homo sapiens |
| 9606 | Homo sapiens |
| 9606 | Homo sapiens |
| 9606 | Homo sapiens |
| 9606 | Homo sapiens |
| 9606 | Homo sapiens |
| 9606 | Homo sapiens |
| 9606 | Homo sapiens |
| 9606 | Homo sapiens |
| 9606 | Homo sapiens |
| 9606 | Homo sapiens |
| 9606 | Homo sapiens |
| 9606 | Homo sapiens |
| 9606 | Homo sapiens |
| 9606 | Homo sapiens |
| 9606 | Homo sapiens |
| 9606 | Homo sapiens |
| 9606 | Homo sapiens |
| 9606 | Homo sapiens |
| 9606 | Homo sapiens |
| 9606 | Homo sapiens |
| 9606 | Homo sapiens |
| 9606 | Homo sapiens |
| 9606 | Homo sapiens |
| 9606 | Homo sapiens |
| 9606 | Homo sapiens |
| 9606 | Homo sapiens |
| 9606 | Homo sapiens |
| 9606 | Homo sapiens |
| 9606 | Homo sapiens |
| 9606 | Homo sapiens |
| 9606 | Homo sapiens |
| 9606 | Homo sapiens |
| 9606 | Homo sapiens |
| 9606 | Homo sapiens |
| 9606 | Homo sapiens |
| 9606 | Homo sapiens |
| 9606 | Homo sapiens |

| | |
|---|---|
| 9606 | Homo sapiens |
| 9606 | Homo sapiens |
| 9606 | Homo sapiens |
| 9606 | Homo sapiens |
| 9606 | Homo sapiens |
| 9606 | Homo sapiens |
| 9606 | Homo sapiens |
| 9606 | Homo sapiens |
| 9606 | Homo sapiens |
| 9606 | Homo sapiens |
| 9606 | Homo sapiens |
| 9606 | Homo sapiens |
| 9606 | Homo sapiens |
| 9606 | Homo sapiens |
| 9606 | Homo sapiens |
| 9606 | Homo sapiens |
| 9606 | Homo sapiens |
| 9606 | Homo sapiens |
| 9606 | Homo sapiens |
| 9606 | Homo sapiens |
| 9606 | Homo sapiens |
| 9606 | Homo sapiens |
| 9606 | Homo sapiens |
| 9606 | Homo sapiens |
| 9606 | Homo sapiens |
| 9606 | Homo sapiens |
| 9606 | Homo sapiens |
| 9606 | Homo sapiens |
| 9606 | Homo sapiens |
| 9606 | Homo sapiens |
| 9606 | Homo sapiens |
| 9606 | Homo sapiens |
| 9606 | Homo sapiens |
| 9606 | Homo sapiens |
| 9606 | Homo sapiens |
| 9606 | Homo sapiens |
| 9606 | Homo sapiens |
| 9606 | Homo sapiens |
| 9606 | Homo sapiens |
| 9606 | Homo sapiens |
| 9606 | Homo sapiens |
| 9606 | Homo sapiens |
| 9606 | Homo sapiens |
| 9606 | Homo sapiens |
| 9606 | Homo sapiens |
| 9606 | Homo sapiens |
| 9606 | Homo sapiens |
| 9606 | Homo sapiens |
| 9606 | Homo sapiens |
| 9606 | Homo sapiens |

| | |
|---|---|
| 9606 | Homo sapiens |
| 9606 | Homo sapiens |
| 9606 | Homo sapiens |
| 9606 | Homo sapiens |
| 9606 | Homo sapiens |
| 9606 | Homo sapiens |
| 9606 | Homo sapiens |
| 9606 | Homo sapiens |
| 9606 | Homo sapiens |
| 9606 | Homo sapiens |
| 9606 | Homo sapiens |
| 9606 | Homo sapiens |
| 9606 | Homo sapiens |
| 9606 | Homo sapiens |
| 9606 | Homo sapiens |
| 9606 | Homo sapiens |
| 9606 | Homo sapiens |
| 9606 | Homo sapiens |
| 9606 | Homo sapiens |
| 9606 | Homo sapiens |
| 9606 | Homo sapiens |
| 9606 | Homo sapiens |
| 9606 | Homo sapiens |
| 9606 | Homo sapiens |
| 9606 | Homo sapiens |
| 9606 | Homo sapiens |
| 9606 | Homo sapiens |
| 9606 | Homo sapiens |
| 9606 | Homo sapiens |
| 9606 | Homo sapiens |
| 9606 | Homo sapiens |
| 9606 | Homo sapiens |
| 9606 | Homo sapiens |
| 9606 | Homo sapiens |
| 9606 | Homo sapiens |
| 9606 | Homo sapiens |
| 9606 | Homo sapiens |
| 9606 | Homo sapiens |
| 9606 | Homo sapiens |
| 9606 | Homo sapiens |
| 9606 | Homo sapiens |
| 9606 | Homo sapiens |
| 9606 | Homo sapiens |
| 9606 | Homo sapiens |
| 9606 | Homo sapiens |
| 9606 | Homo sapiens |
| 9606 | Homo sapiens |
| 9606 | Homo sapiens |
| 9606 | Homo sapiens |
| 9606 | Homo sapiens |

| | |
|---|---|
| 9606 | Homo sapiens |
| 9606 | Homo sapiens |
| 9606 | Homo sapiens |
| 9606 | Homo sapiens |
| 9606 | Homo sapiens |
| 9606 | Homo sapiens |
| 9606 | Homo sapiens |
| 9606 | Homo sapiens |
| 9606 | Homo sapiens |
| 9606 | Homo sapiens |
| 9606 | Homo sapiens |
| 9606 | Homo sapiens |
| 9606 | Homo sapiens |
| 9606 | Homo sapiens |
| 9606 | Homo sapiens |
| 9606 | Homo sapiens |
| 9606 | Homo sapiens |
| 9606 | Homo sapiens |
| 9606 | Homo sapiens |
| 9606 | Homo sapiens |
| 9606 | Homo sapiens |
| 9606 | Homo sapiens |
| 9606 | Homo sapiens |
| 9606 | Homo sapiens |
| 9606 | Homo sapiens |
| 9606 | Homo sapiens |
| 9606 | Homo sapiens |
| 9606 | Homo sapiens |
| 9606 | Homo sapiens |
| 9606 | Homo sapiens |
| 9606 | Homo sapiens |
| 9606 | Homo sapiens |
| 9606 | Homo sapiens |
| 9606 | Homo sapiens |
| 9606 | Homo sapiens |
| 9606 | Homo sapiens |
| 9606 | Homo sapiens |
| 9606 | Homo sapiens |
| 9606 | Homo sapiens |
| 9606 | Homo sapiens |
| 9606 | Homo sapiens |
| 9606 | Homo sapiens |
| 9606 | Homo sapiens |
| 9606 | Homo sapiens |
| 9606 | Homo sapiens |
| 9606 | Homo sapiens |
| 9606 | Homo sapiens |
| 9606 | Homo sapiens |
| 9606 | Homo sapiens |
| 9606 | Homo sapiens |

| | |
|---|---|
| 9606 | Homo sapiens |
| 9606 | Homo sapiens |
| 9606 | Homo sapiens |
| 9606 | Homo sapiens |
| 9606 | Homo sapiens |
| 9606 | Homo sapiens |
| 9606 | Homo sapiens |
| 9606 | Homo sapiens |
| 9606 | Homo sapiens |
| 9606 | Homo sapiens |
| 9606 | Homo sapiens |
| 9606 | Homo sapiens |
| 9606 | Homo sapiens |
| 9606 | Homo sapiens |
| 9606 | Homo sapiens |
| 9606 | Homo sapiens |
| 9606 | Homo sapiens |
| 9606 | Homo sapiens |
| 9606 | Homo sapiens |
| 9606 | Homo sapiens |
| 9606 | Homo sapiens |
| 9606 | Homo sapiens |
| 9606 | Homo sapiens |
| 9606 | Homo sapiens |
| 9606 | Homo sapiens |
| 9606 | Homo sapiens |
| 9606 | Homo sapiens |
| 9606 | Homo sapiens |
| 9606 | Homo sapiens |
| 9606 | Homo sapiens |
| 9606 | Homo sapiens |
| 9606 | Homo sapiens |
| 9606 | Homo sapiens |
| 9606 | Homo sapiens |
| 9606 | Homo sapiens |
| 9606 | Homo sapiens |
| 9606 | Homo sapiens |
| 9606 | Homo sapiens |
| 9606 | Homo sapiens |
| 9606 | Homo sapiens |
| 9606 | Homo sapiens |
| 9606 | Homo sapiens |
| 9606 | Homo sapiens |
| 9606 | Homo sapiens |
| 9606 | Homo sapiens |
| 9606 | Homo sapiens |
| 9606 | Homo sapiens |
| 9606 | Homo sapiens |
| 9606 | Homo sapiens |
| 9606 | Homo sapiens |

| | |
|---|---|
| 9606 | Homo sapiens |
| 9606 | Homo sapiens |
| 9606 | Homo sapiens |
| 9606 | Homo sapiens |
| 9606 | Homo sapiens |
| 9606 | Homo sapiens |
| 9606 | Homo sapiens |
| 9606 | Homo sapiens |
| 9606 | Homo sapiens |
| 9606 | Homo sapiens |
| 9606 | Homo sapiens |
| 9606 | Homo sapiens |
| 9606 | Homo sapiens |
| 9606 | Homo sapiens |
| 9606 | Homo sapiens |
| 9606 | Homo sapiens |
| 9606 | Homo sapiens |
| 9606 | Homo sapiens |
| 9606 | Homo sapiens |
| 9606 | Homo sapiens |
| 9606 | Homo sapiens |
| 9606 | Homo sapiens |
| 9606 | Homo sapiens |
| 9606 | Homo sapiens |
| 9606 | Homo sapiens |
| 9606 | Homo sapiens |
| 9606 | Homo sapiens |
| 9606 | Homo sapiens |
| 9606 | Homo sapiens |
| 9606 | Homo sapiens |
| 9606 | Homo sapiens |
| 9606 | Homo sapiens |
| 9606 | Homo sapiens |
| 9606 | Homo sapiens |
| 9606 | Homo sapiens |
| 9606 | Homo sapiens |
| 9606 | Homo sapiens |
| 9606 | Homo sapiens |
| 9606 | Homo sapiens |
| 9606 | Homo sapiens |
| 9606 | Homo sapiens |
| 9606 | Homo sapiens |
| 9606 | Homo sapiens |
| 9606 | Homo sapiens |
| 9606 | Homo sapiens |
| 9606 | Homo sapiens |
| 9606 | Homo sapiens |
| 9606 | Homo sapiens |
| 9606 | Homo sapiens |
| 9606 | Homo sapiens |

| | |
|---|---|
| 9606 | Homo sapiens |
| 9606 | Homo sapiens |
| 9606 | Homo sapiens |
| 9606 | Homo sapiens |
| 9606 | Homo sapiens |
| 9606 | Homo sapiens |
| 9606 | Homo sapiens |
| 9606 | Homo sapiens |
| 9606 | Homo sapiens |
| 9606 | Homo sapiens |
| 9606 | Homo sapiens |
| 9606 | Homo sapiens |
| 9606 | Homo sapiens |
| 9606 | Homo sapiens |
| 9606 | Homo sapiens |
| 9606 | Homo sapiens |
| 9606 | Homo sapiens |
| 9606 | Homo sapiens |
| 9606 | Homo sapiens |
| 9606 | Homo sapiens |
| 9606 | Homo sapiens |
| 9606 | Homo sapiens |
| 9606 | Homo sapiens |
| 9606 | Homo sapiens |
| 9606 | Homo sapiens |
| 9606 | Homo sapiens |
| 9606 | Homo sapiens |
| 9606 | Homo sapiens |
| 9606 | Homo sapiens |
| 9606 | Homo sapiens |
| 9606 | Homo sapiens |
| 9606 | Homo sapiens |
| 9606 | Homo sapiens |
| 9606 | Homo sapiens |
| 9606 | Homo sapiens |
| 9606 | Homo sapiens |
| 9606 | Homo sapiens |
| 9606 | Homo sapiens |
| 9606 | Homo sapiens |
| 9606 | Homo sapiens |
| 9606 | Homo sapiens |
| 9606 | Homo sapiens |
| 9606 | Homo sapiens |
| 9606 | Homo sapiens |
| 9606 | Homo sapiens |
| 9606 | Homo sapiens |
| 9606 | Homo sapiens |
| 9606 | Homo sapiens |
| 9606 | Homo sapiens |
| 9606 | Homo sapiens |

| | |
|---|---|
| 9606 | Homo sapiens |
| 9606 | Homo sapiens |
| 9606 | Homo sapiens |
| 9606 | Homo sapiens |
| 9606 | Homo sapiens |
| 9606 | Homo sapiens |
| 9606 | Homo sapiens |
| 9606 | Homo sapiens |
| 9606 | Homo sapiens |
| 9606 | Homo sapiens |
| 9606 | Homo sapiens |
| 9606 | Homo sapiens |
| 9606 | Homo sapiens |
| 9606 | Homo sapiens |
| 9606 | Homo sapiens |
| 9606 | Homo sapiens |
| 9606 | Homo sapiens |
| 9606 | Homo sapiens |
| 9606 | Homo sapiens |
| 9606 | Homo sapiens |
| 9606 | Homo sapiens |
| 9606 | Homo sapiens |
| 9606 | Homo sapiens |
| 9606 | Homo sapiens |
| 9606 | Homo sapiens |
| 9606 | Homo sapiens |
| 9606 | Homo sapiens |
| 9606 | Homo sapiens |
| 9606 | Homo sapiens |
| 9606 | Homo sapiens |
| 9606 | Homo sapiens |
| 9606 | Homo sapiens |
| 9606 | Homo sapiens |
| 9606 | Homo sapiens |
| 9606 | Homo sapiens |
| 9606 | Homo sapiens |
| 9606 | Homo sapiens |
| 9606 | Homo sapiens |
| 9606 | Homo sapiens |
| 9606 | Homo sapiens |
| 9606 | Homo sapiens |
| 9606 | Homo sapiens |
| 9606 | Homo sapiens |
| 9606 | Homo sapiens |
| 9606 | Homo sapiens |
| 9606 | Homo sapiens |
| 9606 | Homo sapiens |
| 9606 | Homo sapiens |
| 9606 | Homo sapiens |
| 9606 | Homo sapiens |

| | |
|---|---|
| 9606 | Homo sapiens |
| 9606 | Homo sapiens |
| 9606 | Homo sapiens |
| 9606 | Homo sapiens |
| 9606 | Homo sapiens |
| 9606 | Homo sapiens |
| 9606 | Homo sapiens |
| 9606 | Homo sapiens |
| 9606 | Homo sapiens |
| 9606 | Homo sapiens |
| 9606 | Homo sapiens |
| 9606 | Homo sapiens |

**Submitted entities found**

Rtn3;Ptprd;Lrrn2;Dlg4;Nrxn3;Lrrc4b;Dlgap3
Lrrn2;Dlg4;Nrxn3;Dlgap3
Dlg4;Grik5
Dlg4;Grik5
Rtn3;Ptprd;Dlg4
Ptprd;Lrrc4b
Cdk6
Tcf4
Cdk6
Dlg4;Grik5
Tsc2;Pten
Pten
Ptpn11;Ptpn5
Dlg4
Ptpn11;Ptk2b
Arhgdia
Rtn3;Ptprd;Lrrn2;Dlg4;Grik5;Nrxn3;Lrrc4b;Tspan7;Kcnh7;Slc17a7;Cplx1;Dlgap3
Dlg4
Dnm1

Nptxr;Pten

Dctn1;Dynll1
Dnm1

Dctn1;Dynll1
Sympk

Cdk6
Cdk6
Cdk6

Nptxr;Smarca4

Dctn1;Dynll1
Dlg4

Dlg4
Arhgdia
Arhgdia

Dctn1;Dynll1

Slc17a7
Pten
Sympk

Tmem132a;Sparcl1
Ptpn11
Ptpn11

Dctn1;Dynll1
Tsc2

Dctn1;Dnm1;Dynll1
Kcnh7

Dctn1;Dynll1
Dctn1;Dynll1

Ppargc1a

Arhgef10l;Arhgdia
Dlg4
Ttll7

Slc17a7;Cplx1

Dctn1;Dynll1

Cdk6;Ptpn11

Dctn1;Dynll1

Ptpn11;Pten;Pip5k1b

Cbx3

Pikfyve;Pten;Pip5k1b
Dynll1

Sympk

Dgkz

Ptpn11

Ogdh

Dnm1

Dctn1;Dynll1

Tsc2

Pten

Ptpn11
Dctn1;Dynll1

Ptpn11

Sympk

Ptpn11
Ptpn11

Ptpn11

Nptxr;Ptpn11;Tsc2;Pten;Pip5k1b

Tmem132a;Sparcl1
Mprip

Pten;Pip5k1b

Ptpn11;Ptpn5

Golga5
Ogdh
Nfya
Dlg4

Tsc2;Dynll1
Ptpn11
Nfya

Nfya
Ptpn11;Tsc2;Pten
Dlg4

Dlg4

Ogdh

Cbx3

Ppargc1a

Dynll1
Ptk2b
Dnm1;Epn1
Dctn1;Dynll1
Ash1l

Dctn1;Dynll1

Dlg4

Tsc2
Ptpn11

Dnm1

Ptpn11

Ptpn11;Pip5k1b
Ptpn11
Ptpn11

Pip5k1b

Pikfyve
Mprip

Ppargc1a

Pikfyve
Arhgef10l

Pten

Nptxr;Ptpn11;Tsc2;Pten;Pip5k1b

Nptxr;Pten
Cdk6
Cdk6

Dynll1
Dnm1

Ptpn11

Nfya;Ppargc1a;Med16

Nfya

Elmo2
Kcnh7
Tcf4

Golga5;Dctn1;Dynll1

Ptpn11

Tcf4

Nfya;Ppargc1a;Med16

Tcf4

Ptpn11
Sympk

Ptpn11
Cdk6

Slc17a7;Cplx1

Eef1a2

Epn1

Tcf4

Nptxr;Cdk6
Dynll1
Dynll1

Foxk2
Tcf4
Dnm1

Ptpn11

Eef1a2

Pikfyve

Dnm1

Tsc2

Ptpn11
Sympk

Ogdh
Tcf4
Nfya

Nfya

Smarca4;Tcf4

Cplx1
Cplx1

Ptpn11
Smarca4

Cdk6
Ptpn11
Dynll1
Cplx1

Dctn1;Dynll1

Nptxr;Cdk6;Ptpn11;Smarca4;Zfpm1

Ptpn11

Dynll1

Pten
Cplx1

Ptpn11

Ptpn11
Ptpn11

Dynll1
Dynll1
Ptpn11
Dctn1

Ptpn11

Dlg4;Tspan7
Ptpn11

Ptpn11

Ppargc1a
Ptpn11

Ptpn11
Ptpn11
Dynll1

Ptpn11
Dlg4;Tspan7

Ptpn11

Ptpn11
Ptpn11
Ptpn11

Ptpn11
Nfya

Tcf4
Cplx1
Ptpn11

Pten
Ptpn11
Tcf4
Ptpn11
Dctn1

Ints1
Dctn1;Dynll1

Dlg4;Ptpn11
Ppargc1a;Med16

Tsc2

Ppargc1a

Pikfyve;Pten;Pip5k1b
Ptpn11

Ptpn11
Arhgef10l

Arhgef10l;Arhgdia

Ptpn11

Slc17a7

Dctn1;Ppargc1a;Dynll1

Cbx3

Epn1

Dlg4
Dynll1

Ppargc1a
Tsc2;Dynll1

Ptpn11

Ptpn11
Dynll1
Ptpn11;Ptk2b

Smarca4
Cbx3
Ptpn11;Epn1

Dctn1;Dynll1
Dlg4;Grik5;Tspan7;Slc17a7;Cplx1

Cbx3

Tsc2

Ptpn11
Ptpn11

Dctn1;Dynll1

Dlg4;Grik5;Tspan7
Eif4g3

Trim9
Zfpm1

Elmo2;Ptk2b
Sympk

Dctn1;Nfya
Dctn1;Dynll1
Pten
Dctn1;Dynll1

Nfya;Ppargc1a
Arhgef10l
Pten
Ptpn11
Dlg4

Elmo2;Ptk2b

Ash1l;Smarca4
Ash1l;Smarca4
Cbx3

Ptpn11
Pten
Tsc2

Mprip;Ptpn11;Tsc2;Tcf4;Pten

Arhgef10l;Dlg4;Arhgdia;Dynll1

Dlg4;Ptpn11

Ptpn11
Tspan7

Elmo2
Ptpn11

Ptpn11
Eif4g3

Ptpn11

Dlg4

Ptpn11
Ptpn11

Ptpn11;Dnm1
Ogdh
Ptk2b
Nptxr;Cdk6

Ptpn11
Smarca4;Tcf4;Pip5k1b
Ppargc1a

Ptpn11;Dnm1

Tcf4

Pten
Dctn1;Dynll1

Golga5;Dctn1;Tsc2;Dnm1;Dynll1;Epn1

Ptpn11

Ptpn11;Dnm1
Dnm1

Dynll1

Elmo2
Dlg4
Tcf4
Ptpn11
Dlg4;Ptpn11
Hnrnpul1;Sympk
Tcf4
Ptpn11
Ptpn11

Tcf4

Hnrnpul1;Sympk
Ptpn11
Dlg4;Dnm1
Pten
Ptpn11
Nptxr;Cdk6;Dctn1;Tsc2;Dynll1
Foxk2;Pten
Nptxr;Cdk6;Dctn1;Dynll1

Ptpn11

Tcf4

Dgkz
Dynll1
Cbx3

Zfpm1

Golga5;Dctn1;Tsc2;Dnm1;Dynll1;Epn1
Dgkz;Ptpn11

Ptpn11;Tspan7
Ptpn11
Cdk6

Ptpn11
Ppargc1a

Dlg4;Ptpn11
Dlg4;Dynll1

Dlg4;Elmo2;Ptpn11;Ptk2b;Dnm1;Epn1
Ptpn11;Eif4g3

Hnrnpul1;Sympk

Trim9;Dctn1;Ptpn11;Pten;Dnm1;Dynll1
Trim9
Smarca4;Tcf4;Pip5k1b
Eef1a2

Elmo2
Elmo2
Cdk6;Dctn1;Dynll1
Dnm1

Arhgef10l;Arhgdia

Slc17a7
Tsc2;Pten
Dlg4;Ptpn11;Ptk2b;Smarca4;Ptpn5

Dynll1
Cbx3;Nfya;Sympk;Ppargc1a;Tsc2;Pten;Zfpm1;Nptxr;Cdk6;Ints1;Ptpn11;Smarca4;Tcf4;Med16
Dynll1
Ogdh

Nfya

Nptxr;Cdk6;Cbx3;Nfya;Ptpn11;Ppargc1a;Tsc2;Smarca4;Tcf4;Pten;Zfpm1;Med16

Cbx3;Nfya;Sympk;Ppargc1a;Tsc2;Pten;Zfpm1;Nptxr;Cdk6;Ints1;Ptpn11;Smarca4;Tcf4;Med16

Dgkz;Ptpn11;Tspan7;Zfpm1

Cdk6;Dctn1;Dynll1
Dlg4;Ptpn11;Dnm1

Dlg4;Ptpn11;Ptk2b;Smarca4;Eif4g3;Ptpn5
Nfya;Pikfyve;Ppargc1a;Pten;Pip5k1b;Med16

Eef1a2;Ttll7;Dctn1;Foxk2;Tmem132a;Ppargc1a;Sparcl1;Pten;Dynll1

Ppargc1a

Ppargc1a
Dgkz;Arhgef10l
Elmo2;Ptpn11;Dnm1;Dynll1
Hnrnpul1;Sympk
Trim9;Dctn1;Dlg4;Elmo2;Ptpn11;Ptk2b;Smarca4;Pten;Eif4g3;Dynll1;Ptpn5;Dnm1
Slc17a7
Dlg4;Ptpn11;Ppargc1a;Tcf4;Med16;Dnm1
Eef1a2;Ttll7;Dctn1;Nfya;Foxk2;Tmem132a;Ppargc1a;Sparcl1;Tcf4;Pten;Dynll1
Dgkz;Arhgef10l
Cdk6;Mprip;Ptpn11;Tsc2;Tcf4;Pten
Tsc2;Ptk2b;Pten;Arhgdia;Dnm1;Epn1;Dgkz;Arhgef10l;Nptxr;Dlg4;Elmo2;Ptpn11;Smarca4;Tcf4;Pip5k1b;Dynll1
Nfya;Pikfyve;Ppargc1a;Ogdh;Pten;Pip5k1b;Med16

**Mapped entities**

O95886;Q9Y4C0;O43300;P23468;Q9HDB5;P78352;Q9NT99;O95197

P23468;P78352;O95197

ENST00000371953;P60484;P49815;ENSG00000171862

ENST00000371953;P60484

O95886;O43300;Q9NS40;Q9HDB5;Q9P2U7;Q9NT99;O95197;Q9Y4C0;P23468;P78352;Q16478;O14810;P41732

P63167;Q14203

Q05193;P63167;Q14203

P52565;Q9HCE6
P78352

Q00534;Q06124

P63167;Q14203-2

P60484;Q06124;O14986

P60484;Q9Y2I7;O14986

Q06124

Q02218

Q05193

P60484

ENST00000371953;P60484;O95503;Q06124;O14986;P49815;ENSG00000171862

P60484;O14986

Q02218

P63167;P49815

P60484;Q06124;P49815
P78352

P78352

Q13185

ENSG00000109189;Q9UBK2

P63167

P63167;Q14203

P78352

Q06124

Q05193

Q06124
Q06124

Q9UBK2

Q9HCE6

ENST00000371953;P60484;O95503;Q06124;O14986;P49815;ENSG00000171862

ENST00000371953;P60484;O95503;ENSG00000171862
Q00534
Q00534

P63167
Q05193

P23511;Q9Y2X0;Q9UBK2

Q96JJ3

Q9NQB0

P63167;Q14203;Q8TBA6

P23511;Q9Y2X0;Q9UBK2

Q06124
Q92797

Q00534

Q05639

Q00534;O95503

Q01167

Q05193

Q9Y2I7

Q05193

P49815

Q06124
Q92797

Q02218
Q9NQB0

P23511

ENSG00000196628;Q9NQB0;P51532

Q06124

Q00534

P63167;Q14203

Q00534;O95503;Q06124;P51532;Q8IX07

Q06124

ENSG00000204843;Q14203

P78352;P41732

ENSG00000109189;Q9UBK2
Q06124

Q06124

P78352;P41732

Q06124

Q06124

Q06124
P23511

Q9NQB0

Q06124

ENST00000371953;P60484
Q06124
Q9NQB0
Q06124
ENSG00000204843;Q14203

Q8N201
P63167;Q14203

Q06124;P78352
Q9Y2X0;Q9UBK2

P49815

Q9UBK2

P60484;Q9Y2I7;O14986
Q06124

Q06124
Q9HCE6

P52565;Q9HCE6

Q06124

P63167;Q14203-2;ENSG00000109189;Q9UBK2

Q13185

P78352
P63167

ENSG00000109189;Q9UBK2
P63167;P49815

Q06124

Q06124

Q06124;Q14289

P51532
Q13185
Q06124;Q9Y6I3

P63167;Q14203-2
P78352;Q16478;Q9P2U7;O14810;P41732

Q13185

P49815

Q06124
Q06124

P63167;Q14203-2

P78352;Q16478;P41732
O43432

Q8IX07

Q96JJ3;Q14289
Q92797

ENSG00000204843;Q14203;P23511
P63167;Q14203-2
P60484
P63167;Q14203-2

P23511;Q9UBK2
Q9HCE6
P60484
Q06124
P78352

Q96JJ3;Q14289

Q9NR48;P51532
Q9NR48;P51532
Q13185

Q06124

P49815

P60484;Q9NQB0;Q06124;Q6WCQ1;P49815

P52565;P63167;Q9HCE6;P78352

Q06124;P78352

Q06124
P41732

Q96JJ3

Q06124
O43432

Q06124

P78352

Q06124
Q06124

Q02218
Q14289
Q00534;O95503

ENSG00000196628;Q9NQB0;O14986;P51532
Q9UBK2

Q05193;Q06124

Q9NQB0

P60484
P63167;Q14203

Q05193;P63167;Q14203;Q9Y6I3;P49815;Q8TBA6

Q06124

Q05193

Q96JJ3
P78352
Q9NQB0
Q06124
Q06124;P78352
Q92797;Q9BUJ2
Q9NQB0

Q9NQB0

Q92797;Q9BUJ2
Q06124
Q05193;P78352
P60484
Q06124
P63167;Q14203;Q00534;O95503;P49815
Q01167;P60484
P63167;Q14203;Q00534;O95503

Q06124

Q9NQB0

Q13574
P63167
Q13185

Q8IX07

Q05193;P63167;Q14203;Q9Y6I3;P49815;Q8TBA6
Q06124;Q13574

Q06124;P41732
Q06124
Q00534

Q06124
Q9UBK2

Q06124;P78352
P63167;P78352

Q96JJ3;Q05193;Q06124;Q9Y6I3;P78352;Q14289
O43432;Q06124

Q92797;Q9BUJ2

Q05193;P63167;Q14203;Q9C026;P60484;Q06124
Q9C026
ENSG00000196628;Q9NQB0;O14986;P51532
Q05639

Q96JJ3
Q96JJ3
P63167;Q14203-2;Q00534
Q05193

P52565;Q9HCE6

Q9P2U7
ENST00000371953;P60484;P49815;ENSG00000171862
ENSG00000110786;P54829;Q06124;P78352;ENSG00000179295;Q14289;P51532

P63167
O95503;Q06124;P51532;Q8IX07;ENSG00000171862;ENST00000371953;P60484;Q00534;Q92797;P23511;Q13185;Q[illegible]
P63167
Q02218

P23511

O95503;Q06124;P51532;Q8IX07;ENSG00000171862;ENST00000371953;P60484;Q00534;P23511;Q13185;Q9NQB0;Q[illegible]

O95503;Q06124;P51532;Q8IX07;ENSG00000171862;ENST00000371953;P60484;Q00534;Q92797;P23511;Q13185;Q[illegible]

Q06124;Q13574;Q8IX07;P41732

P63167;Q14203-2;Q00534
Q05193;Q06124;P78352

ENSG00000110786;O43432;P54829;Q06124;P78352;ENSG00000179295;Q14289;P51532
P60484;P23511;Q9Y2I7;O14986;Q9Y2X0;Q9UBK2

P63167;Q14203;Q01167;P60484;Q6ZT98;P68104;Q14515;Q24JP5;Q9UBK2

Q9UBK2

Q9UBK2
Q9HCE6;Q13574
Q96JJ3;Q05193;P63167;Q06124
Q92797;Q9BUJ2
Q05193;Q14203;Q9C026;ENSG00000110786;O43432;Q06124;P51532;Q96JJ3;P63167;P60484;P54829;P78352;ENSG
Q9P2U7
Q05193;Q06124;P78352;Q9Y2X0;Q9UBK2;P15884
Q14203;Q6ZT98;Q14515;Q24JP5;ENSG00000204843;P63167;Q01167;Q05639;P60484;P68104;P23511;Q9NQB0;Q9U
Q9HCE6;Q13574
P60484;Q00534;Q9NQB0;Q06124;Q6WCQ1;P49815
Q05193;P52565;O95503;Q9HCE6;Q06124;Q9Y6I3;P51532;ENSG00000171862;Q96JJ3;ENST00000371953;ENSG00000
Q02218;P60484;P23511;Q9Y2I7;O14986;Q9Y2X0;Q9UBK2

**Submitted entities hit interactor**

Rtn3

Rtn3

Pten;Tsc2

Pten

Frmpd4;Ptk2b;Dlg4;Rtn3;Dctn1;Dlg4;Speg;Dnm1

Tcf4

Tcf4

Tcf4

Ptpn11

Ptpn11

Dhx30

Tcf4

Ptpn11

Cplx1

Cplx1

Cplx1

Smarca4

Smarca4

Cplx1

Dnm1

Dnm1
Dnm1
Smarca4
Smarca4

Dnm1
Dnm1
Tcf4;Tcf4
Dynll1
Dctn1;Dnm1

Dnm1
Dnm1
Dynll1
Tcf4
Cplx1
Smarca4
Smarca4
Dnm1

Dctn1
Dnm1

Smarca4
Dnm1

Cdk6
Tcf4
Dnm1
Dnm1

Dnm1

Dnm1
Dynll1
Dnm1
Dlg4
Ppargc1a

Ptprd
Dynll1
Tcf4
Tcf4
Ptk2b;Cdk6

Dnm1
Dhx30;Fam120a
Dnm1
Ptpn11
Dnm1
Dynll1
Dnm1
Pten
Tcf4;Tcf4
Dnm1
Tcf4;Tcf4

Dnm1
Ptpn11
Cdk6
Dgkz
Dnm1
Pten;Rtn3

Dnm1

Dnm1
Dnm1
Dnm1
Dnm1
Dnm1
Ptk2b

Dnm1
Cplx1
Ptk2b;Pten
Smarca4
Ptpn11
Dctn1
Dgkz
Ptpn11
Dnm1
Dlg4;Dlg4;Dnm1
Dnm1
Dnm1
Dynll1;Dnm1
Dnm1
Dnm1
Dnm1
Dnm1
Dhx30
Dnm1
Eif4g3;Eif4g3

Dnm1

Dynll1
C2cd2l
Dnm1
Dnm1
Pten;Ptpn11
Dnm1
Zfpm1
Pten;Pten
Zfpm1
Dnm1

Ptpn11

Dnm1
Tcf4
Ptpn11
Ptpn11
Tcf4
Dnm1
Smarca4
Eif4g3
Zfpm1
Smarca4
C2cd2l

Dnm1

Arhgdia
Ptk2b

Dhx30
Pten;Pten;Dlg4
Ptk2b
Dctn1
Ptpn11;Dnm1
Dhx30
Ptpn11

Ptpn11;Dnm1
Pten
Rtn3
Dnm1
Dnm1
Dnm1
Dnm1
Dnm1

Smarca4

Dnm1
Dnm1
Tcf4

Dnm1

Dnm1
Dynll1;Dlg4

Dlg4

Dlg4
Dnm1
Dnm1
Ptk2b
Dnm1
Cdk6

Dgkz
Cbx3
Dnm1
Ppargc1a;Dynll1
Dnm1
Dlg4
Dynll1

Dgkz
Smarca4
Dnm1
Tcf4;Tcf4
Ptk2b;Ptpn11
Dnm1
Dnm1
Tcf4
Dctn1;Dnm1
Ptpn11;Dnm1
C2cd2l
Smarca4
Zfpm1
Dlg4;Dlg4
Zfpm1
Eif4g3;Eif4g3
Dnm1

Ptk2b
Dynll1
Dnm1
Ptk2b
Ptpn11
Dnm1
Dnm1
Dynll1
C2cd2l
Dynll1

Dnm1
Dnm1
Dnm1

Ptpn11
Ptpn11
Tcf4;Tcf4
Dnm1
Dnm1
Dlg4;Ptpn11;Dlg4
Zfpm1
Dnm1

Zfpm1
Ptpn11

Rtn3
Smarca4;Ptpn11
Rtn3
Nisch

Tcf4;Dnm1
Dnm1

Dnm1
Pten;Pten;Dlg4
Dnm1
Dnm1
Pten;Pten
Dgkz
Dgkz
Nfya
Smarca4
Dnm1
Eml2
C2cd2l
C2cd2l

Cdk6
Zfpm1;Cdk6
Ptpn11
Eif4g3;Dhx30;Tcf4

Dnm1
Ppargc1a
Ptpn11

Dnm1
Ptk2b
Ptpn11;Dnm1

Tcf4;Cdk6
Pten;Ptpn11
Tcf4;Dctn1;Dnm1
Ptk2b;Dnm1

Smarca4
Dynll1

Hnrnpul1;Eif4g3
Cbx3
Tcf4;Tcf4
Ppargc1a
Smarca4

Tcf4;Ptpn11
Gas7
Ptk2b
Eif4g3
Rtn3
Dlg4
Dnm1
Cbx3
Tcf4
Mprip;Arhgdia
Ptpn11
Hnrnpul1;Eif4g3
Eif4g3;Tcf4;Eif4g3

Cdk6
Tcf4;Tcf4

Ptpn11;Dnm1
Dnm1
Ptk2b;Dnm1

Tcf4;Tcf4
Ptpn11

Ptpn11
Dnm1
Rtn3

Tcf4
Zfpm1;Cdk6


C2cd2l
C2cd2l
Ptk2b
Foxk2

Dnm1
Dnm1
Tcf4
Pten;Ptpn11
Hnrnpul1;Eif4g3

Dnm1

Tcf4
Dlg4
Rtn3
Tcf4;Trim9;Ptpn11
Hnrnpul1;Dhx30;Fam120a
C2cd2l
Zfpm1
Ptprd
Dlg4
Zfpm1
Dnm1
Hnrnpul1;Eif4g3
Mprip;Arhgdia
Dnm1
Zfpm1

Dnm1
Mprip;Samd4b;Tsc2
Nfya
Dnm1
Tcf4
Tcf4;Smarca4;Tcf4
Ptk2b
Rtn3
Dlg4
Tcf4


Eif4g3

Map1b;Hnrnpul1;Nfya;Tcf4
Ptpn11;Dnm1
Ptpn11

Dgkz;Tcf4;Pten;Ptpn11
Ptk2b;Arhgdia
Dlg4
Dnm1
Tcf4;Dgkz;Cdk6;Nfya;Tcf4

Speg;Ptprd
Cdk6
Ptpn11;Dnm1
Trim9;Dnm1
Zfpm1;Tcf4;Tcf4
Tcf4;Tcf4
Ptk2b;Speg;Cdk6
Trim9;Trim9;Rtn3

Pten;Ptpn11

Smarca4

Dgkz
Dnm1
Cdk6
Tcf4;Ptpn11;Dnm1
Tcf4
Ptpn11;Dnm1
Ptk2b;Tcf4
Tcf4

Eif4g3
Eif4g3
Cdk6
Dnm1

Gas7
Dlg4
Cplx1;Tcf4;Dnm1

Dnm1
Frmpd4;Speg

Mprip;Samd4b;Tsc2

Zfpm1;Dnm1
Ppargc1a;Dynll1
Tcf4;Ptpn11;Dnm1
Fibp;Tsc2;Pten
Tcf4;Ptpn11;Dnm1

Tcf4

Frmpd4;Speg
Nfya
Samd4b;Tsc2
Dnm1
Pten;Ptpn11
Dnm1
Tcf4;Ptpn11;Dnm1
Dnm1

Dnm1
Ptpn11;Dnm1
Ptpn11;Dnm1
Tcf4;Ptpn11;Dnm1
Nfya
Tcf4
Ptprd
Smarca4
Tcf4
Map1b
Smarca4
Tcf4

Dnm1
Ptpn11
Zfpm1
Tcf4;Ptpn11
Cbx3;Ppargc1a;Dlg4;Pten;Tsc2;Ptpn11;Dlg4
Dnm1
Dnm1
Dnm1
Gas7
Ptk2b;Dnm1
Dnm1
Tcf4
Trim9;Dnm1
Nfya;Dnm1
Trim9;Dnm1
Smarca4;Cdk6;Dlg4

Ptpn11;Dnm1
Dnm1
Cdk6;Ppargc1a;Smarca4;Cdk6
Eif4g3;Tcf4;Eif4g3;Dnm1
Tcf4;Dlg4
Tcf4;Dlg4
Eif4g3;Tcf4;Eif4g3;Dnm1
Nisch
Dnm1
Ptpn11;Dnm1
Dnm1
Dgkz
Zfpm1;Tcf4;Tcf4
Tcf4;Trim9;Rtn3
Dnm1
Eif4g3
Dnm1
C2cd2l;Pten;Rtn3
Dlg4;Dnm1
Ptk2b;Arhgdia
Tcf4;Dlg4
Ptpn11;Dnm1
Ptpn11;Dnm1
Dnm1
Tcf4;Dynll1
Pten;Ptpn11
Zfpm1
Tcf4;Tcf4
Hnrnpul1;Dhx30;Smarca4;Fam120a
Mprip;Arhgdia
Dnm1
Dlg4;Dnm1
Dnm1
Dnm1
Dnm1
Dgkz
Dynll1;Dnm1
Cplx1;Dnm1
Dnm1

Smarca4
Smarca4
Dnm1
C2cd2l;Ppargc1a;Eml2;Dynll1;Dctn1;Dnm1
Dlg4
Tcf4
Tcf4;Dnm1
Dnm1
C2cd2l;Dctn1;Dnm1
Dnm1

Dgkz

Ptk2b;Dnm1
Ptpn11;Dnm1
Ptk2b;Dctn1;Dnm1
Dynll1
Ptk2b;Dnm1
Tcf4
Dgkz;Tcf4
Smarca4;Trim9;Trim9
Tcf4;Dynll1;Dlg4;Dnm1
Cbx3
Smarca4
Smarca4
Smarca4
Smarca4
Smarca4
Dgkz
Smarca4;Cdk6;Dlg4
Dnm1
Dnm1
Zfpm1
Ptpn11
Ptpn11;Ptpn11;Dnm1

Ptk2b;Tcf4;Trim9;Ptpn11
Foxk2;Zfpm1
Speg
Tcf4
Ptpn11;Dnm1
Tcf4;Dnm1
C2cd2l;Eml2;Dctn1;Dnm1
Frmpd4;Ptk2b;Dlg4;Dctn1;Dlg4;Speg;Dnm1
Cbx3;C2cd2l
Tcf4
Tcf4;Dgkz;Tcf4
Smarca4
Zfpm1;Cbx3
Cbx3;Map1b;Eef1a2;Dnm1
Dlg4
Ptpn11;Ptpn11;Dnm1
Ptk2b;Dnm1
C2cd2l;Samd4b;Tsc2;Ptpn11
Cplx1;Cdk6
Ptpn11;Dnm1
Dgkz
Tcf4
Frmpd4;Ptk2b;Dlg4;Dctn1;Dlg4;Speg;Dnm1
Tcf4;Pten
Tcf4

Speg
Dnm1
Ptk2b;Pten;Ptpn11;Dnm1
Hnrnpul1;Nfya;Eif4g3
Dnm1
Dnm1
Gas7
Tcf4;Tcf4;Ptprd
Pten;Pten;Ngrn
Tcf4;Tcf4;Ptprd
Cplx1;Dnm1
Smarca4;Dgkz;Tcf4;Pten;Ptpn11
Dlg4;Dnm1
Pten
Dlg4;Tcf4;Ptpn11;Dlg4;Dnm1
Ptk2b;Dlg4;Dctn1;Dlg4;Dnm1
Dnm1
Ptk2b;Pten;Ptpn11;Dnm1
Epn1;Ptpn11
Speg
Speg
Zfpm1;Cbx3;Smarca4
C2cd2l;Tcf4
Ptk2b;Dnm1

Mprip;Eif4g3;Samd4b;Dhx30;Tcf4;Tsc2;Dnm1
Tcf4
Zfpm1;Smarca4;Ptpn11;Dlg4;Dnm1
Dnm1
Smarca4;Tcf4;Dctn1;Dnm1
Map1b;Hnrnpul1;Nfya;Tcf4;Tcf4
Smarca4
Hnrnpul1;Tcf4
Tcf4;Trim9;Ptpn11
Cdk6
Hnrnpul1;Arhgdia;Dctn1
Ptpn11;Dnm1
Ptpn11;Dnm1
Ptk2b;Ptpn11;Dnm1
Tcf4;Dnm1
Ptk2b;Ptpn11;Dnm1
Speg
Tcf4;Pten;Tsc2;Dlg4
Cdk6;Tcf4
Dnm1

Nisch
Tcf4;Ptpn11;Dnm1
Dhx30;Tcf4;Pten

Speg;Tcf4;Ptprd
Tcf4;Ptpn11;Dnm1
Tcf4;Tcf4;Dnm1
Dnm1
Cdk6;Nfya
Ptpn11;Dnm1
C2cd2l;Tcf4
Dnm1
Tcf4;Ptpn11;Dnm1
Tcf4;Ptpn11;Dnm1
Ptk2b;Tcf4;Tcf4;Dnm1

Dnm1
Ptpn11;Ptpn11;Dnm1
Zfpm1;Tcf4;Cdk6;Dgkz;Nfya;Tcf4
Tcf4;Tcf4
Ptk2b;Speg;Cdk6;Dlg4
Ppargc1a;Ptpn11;Ptpn11

Zfpm1;Tcf4;Samd4b;Smarca4;Tcf4;Dynll1;Trim9;Trim9;Tsc2
Cbx3;Tcf4;Cbx3
C2cd2l
Ptk2b;Mprip;Arhgdia;Dynll1;Dlg4;Tcf4;Ptpn11;Dlg4;Dnm1
Nisch;Dhx30
Rtn3;Dnm1
Tcf4;Samd4b;Tcf4;Dynll1;Trim9;Trim9;Tsc2
Cdk6;Cdk6
Pten
Trim9;Dnm1
Zfpm1;Tcf4;Cdk6;Tcf4
C2cd2l;Cplx1;Dynll1;Tcf4;Trim9;Dctn1;Dlg4;Dnm1
Dnm1
Tcf4;Dgkz;Tcf4;Ptpn11
Ptk2b;Ptpn11;Dnm1
Rtn3;Dnm1
Hnrnpul1;Arhgdia;Dctn1

Mprip;Arhgdia;Dynll1;Ptpn11;Dnm1
Dhx30;Tcf4

Tcf4;Tcf4
Dnm1
Ptk2b;Pten;Dnm1
Ptk2b;Tcf4;Cdk6;Tcf4
Ptk2b;Arhgdia;Dnm1
Cdk6;Cdk6;Dnm1
Hnrnpul1;Nfya;Dhx30;Fam120a
Zfpm1;Tcf4;Tcf4

Dnm1
Fibp;Tsc2;Pten;Dnm1
C2cd2l;Cdk6;Rtn3
Hnrnpul1;Ppargc1a
Hnrnpul1;Nfya;Dhx30;Fam120a
Tcf4;Ptpn11;Dnm1
C2cd2l;Tcf4;Ptpn11;Dnm1
Pten;Dnm1
Tcf4;Ptpn11;Dnm1
Zfpm1;Tcf4;Cdk6;Dgkz;Nfya;Tcf4;Dynll1;Dlg4;Dnm1
Foxk2;Pten;Tcf4;Pten;Ngrn
Zfpm1;Tcf4;Cdk6;Dgkz;Nfya;Tcf4;Dnm1
Dlg4;Dnm1
Dhx30;Tcf4;Trim9
Tcf4;Ptpn11;Dnm1
Tcf4
Hnrnpul1;Cbx3;Smarca4
Tcf4;Cdk6;Nfya;Tcf4
Tcf4
Tcf4;Trim9;Rtn3
Map1b;Hnrnpul1;Nfya;Tcf4;Pten;Tcf4;Tsc2;Dlg4
Mprip;Samd4b;Tsc2;Eif4g3
Ptpn11;Dnm1
Tcf4;Dlg4
Zfpm1;Cbx3
C2cd2l;Tcf4;Tcf4;Dnm1
Foxk2;Zfpm1
Foxk2;Cbx3;Fibp;Smarca4;Dctn1;Dnm1
Ptk2b;Tcf4;Eif4g3;Dnm1
C2cd2l;Cplx1;Dynll1;Tcf4;Trim9;Dctn1;Dlg4;Dnm1
Dlg4;Dnm1
Dhx30;Tcf4;Trim9
Pten;Ptpn11;Speg;Dnm1
Tcf4;Ptpn11;Dnm1
Tcf4;Dgkz;Tcf4;Ptpn11
Tcf4;Dnm1
Tcf4;Dynll1
Tcf4;Ptpn11;Dnm1
Zfpm1;Tcf4;Dgkz;Tcf4
Epn1;Ptpn11
Ptk2b;Ptpn11;Dnm1
Smarca4;Tcf4;Dctn1
Ptpn11;Dnm1
Ptk2b;Mprip;Arhgdia;Tcf4;Dynll1;Dlg4;Tcf4;Pten;Ptpn11;Ptpn11;Dlg4;Dnm1
Dhx30;Tcf4;Pten
Dnm1
Hnrnpul1;Nfya;Eif4g3;Dhx30;Fam120a
Dhx30;Tcf4;Trim9
Hnrnpul1;Arhgdia;Dctn1
Ptpn11;Dnm1

Ptk2b;Epn1;Pten;Ptpn11;Dnm1
Dnm1
Zfpm1;Tcf4;Samd4b;Smarca4;Tcf4;Dynll1;Trim9;Trim9;Tsc2;Dnm1
Eif4g3;Tcf4;Eif4g3;Dnm1
Cplx1;Dynll1;Dlg4;Dnm1
Ptk2b;Arhgdia;Trim9;Trim9;Rtn3;Dnm1
Tcf4;Cdk6;Nfya;Tcf4
Tcf4;Dynll1
Tcf4
Ptk2b;Dnm1
Smarca4
Smarca4
Cbx3;Map1b;Eef1a2;Gas7;Ptpn11;Dnm1
Cbx3;Map1b;Eef1a2;Gas7;Ptpn11;Dnm1
Zfpm1;Tcf4;Tcf4;Cdk6;Dgkz;Tcf4;Ptpn11;Ptprd
C2cd2l;Tcf4;Tcf4;Dnm1
Tcf4;Cdk6;Nfya;Tcf4
Tcf4;Dynll1;Dlg4;Dnm1
C2cd2l;Tcf4
Epn1;Ptpn11
Ppargc1a;Tcf4;Ptpn11;Ptpn11
Dlgap3;Fibp;Pten;Dlg4
Dlgap3;Fibp;Pten;Dlg4
C2cd2l;Cdk6;Rtn3
Hnrnpul1;Map1b;Nfya;Tcf4;Pten;Speg;Tcf4;Pten;Tsc2;Dlg4;Ptprd
Hnrnpul1;Ptk2b;C2cd2l;Samd4b;Arhgdia;Tsc2;Ptpn11;Dctn1;Ptpn11;Dnm1
Hnrnpul1;Dhx30;Smarca4;Cdk6;Fam120a;Dlg4
Tcf4;Dynll1;Tcf4
Hnrnpul1;Cbx3;Ppargc1a;Pten;Tsc2;Ptpn11;Ptpn11;Med16;Dnm1;Map1b;Nfya;Tcf4;Dgkz;Tcf4;Dlg4;Zfpm1;Cdk6;Dlg4;[illegible]
Tcf4;Dynll1;Tcf4
Dnm1
Speg;Smarca4;Cdk6;Nfya
Ptk2b;Arhgdia;Trim9;Trim9;Rtn3;Dnm1
Cbx3;C2cd2l;Nfya
C2cd2l;Tcf4
Zfpm1;Hnrnpul1;Cbx3;Ppargc1a;Cdk6;Dlg4;Pten;Tsc2;Ptpn11;Ptpn11;Med16;Map1b;Nfya;Ptk2b;Tcf4;Pten;Speg;Smar[illegible]
Ptk2b;Fibp;Tsc2;Pten;Dnm1
Hnrnpul1;Cbx3;Ppargc1a;Pten;Tsc2;Ptpn11;Ptpn11;Med16;Dnm1;Map1b;Nfya;Tcf4;Dgkz;Tcf4;Dlg4;Zfpm1;Cdk6;Dlg4;[illegible]
Hnrnpul1;Map1b;Nfya;Tcf4;Pten;Tcf4;Tsc2;Dlg4
Foxk2;Ptk2b;Cbx3;Fibp;Smarca4;Pten;Ptpn11;Dctn1;Speg;Dlg4;Dnm1
Smarca4;Tcf4;Tcf4
Zfpm1;Tcf4;Smarca4;Tcf4;Cdk6;Dgkz;Tcf4;Ptpn11;Dlg4;Ptprd
Zfpm1;Ptk2b;C2cd2l;Arhgdia;Tcf4;Tcf4;Ptpn11;Eif4g3;Dnm1
Ptk2b;Speg;Smarca4;Cdk6;Nfya;Dlg4
Hnrnpul1;Samd4b;Tsc2;Pten;Ptpn11;Ptpn11;Dnm1;Ptk2b;C2cd2l;Arhgdia;Dhx30;Tcf4;Dctn1
Cbx3;C2cd2l;Ppargc1a;Nfya;Pten;Ptpn11;Rtn3
Ptk2b;Speg;Smarca4;Cdk6;Nfya;Dlg4
Hnrnpul1;Cbx3;Ppargc1a;Trim9;Pten;Dnm1;Map1b;Foxk2;Nfya;Tcf4;Pten;Smarca4;Tcf4;Cbx3;Dlg4;Ngrn;Ptprd
Nisch;Ptk2b;Cplx1;Arhgdia;Dhx30;Cdk6;Trim9;Trim9;Rtn3;Ptpn11;Dnm1
Map1b;Hnrnpul1;Nfya;Tcf4;Cbx3;Ppargc1a;Smarca4;Tcf4;Cbx3;Dlg4;Ptprd

Map1b;Hnrnpul1;Nfya;Tcf4;Cbx3;Ppargc1a;Smarca4;Tcf4;Cbx3;Dlg4;Ptprd
Ptk2b;Mprip;Fibp;Arhgdia;Tcf4;Tsc2;Pten;Ptpn11;Dnm1
Tcf4;Dhx30;Cdk6;Tcf4;Ptpn11;Dnm1
Hnrnpul1;Mprip;Nfya;Eif4g3;Samd4b;Dhx30;Fam120a;Tcf4;Tsc2;Trim9;Eif4g3;Dnm1
Hnrnpul1;Samd4b;Epn1;Cdk6;Tsc2;Pten;Ptpn11;Ptpn11;Dnm1;Ptk2b;Tcf4;C2cd2l;Dhx30;Arhgdia;Tcf4;Dctn1
C2cd2l;Cdk6;Tcf4;Rtn3;Dnm1
Zfpm1;Cdk6;Ppargc1a;Cdk6;Ptpn11;Dnm1;Ptk2b;C2cd2l;Tcf4;Arhgdia;Smarca4;Tcf4;Eif4g3
Hnrnpul1;Eif4g3;Cbx3;Ppargc1a;Pten;Trim9;Gas7;Dnm1;Foxk2;Map1b;Nfya;Tcf4;Pten;Smarca4;Tcf4;Tcf4;Cbx3;Rtn3;Ei
Ptk2b;Mprip;Fibp;C2cd2l;Arhgdia;Tcf4;Tsc2;Pten;Ptpn11;Dnm1
Zfpm1;Cplx1;Cdk6;Trim9;Trim9;Pten;Ptpn11;Dnm1;Dlgap3;Nisch;Ptk2b;Fibp;Smarca4;Arhgdia;Dhx30;Rtn3;Dlg4
Hnrnpul1;Eif4g3;Mprip;Cbx3;Cdk6;Samd4b;Trim9;Trim9;Pten;Tsc2;Ptpn11;Ptpn11;Dnm1;Map1b;Fibp;C2cd2l;Fam120
Cbx3;C2cd2l;Cplx1;Ppargc1a;Nfya;Tcf4;Pten;Ptpn11;Rtn3;Dnm1

**Interacts with**

O95197

O95197

P60484;Q92574

P60484

Q13224;P12931;Q12959;O95197;15996;Q9NRD5;P10636-8;Q14957;Q12879

O43929

Q9Y4Z2;Q13562

Q13761

P49247

P49247

P19525

Q13761

P49247

P63027

P63027

P63027

P46531

P46531

P63027

| | |
|---|---|
| | 15996 |
| | 15996 |
| P46531 | |
| P46531 | |
| | 15996 |
| | 15996 |
| Q9Y316 | |
| Q9NR28 | |
| P43034;15996 | |
| | 15996 |
| | 15996 |
| P63244 | |
| Q9Y4Z2 | |
| P63027 | |
| P46531 | |
| P46531 | |
| | 15996 |
| P30622 | |
| | 15996 |
| O75376 | |
| | 15996 |
| Q01196 | |
| Q8NFF5 | |
| | 15996 |
| | 15996 |
| | 15996 |
| | 15996 |
| P98170 | |
| | 15996 |
| Q05513 | |
| P37231 | |
| O14965 | |
| P98170 | |
| P38936 | |
| P38936 | |
| P12931;Q01196 | |

15996
P67809
15996
P08581
15996
Q96LC9
15996
P60484
Q9UI95
15996
Q13761;P38936

15996
P08581
Q01196
P06400
15996
P60484;Q9NTJ5

15996

15996
15996
15996
15996
15996
P06241

15996
P63027
P46934;P12931
O14746
Q15116
P10636-8
P06400
P49023
15996
Q13224;15996
15996
15996
O96015;15996
15996
15996
15996
15996
P19525
15996
P11940;P06730

15996

| | |
|---|---|
| P98170 | |
| P08034 | |
| | 15996 |
| | 15996 |
| Q03135 | |
| | 15996 |
| P49841 | |
| P60484;Q93009 | |
| P49841 | |
| | 15996 |
| P49023 | |
| | 15996 |
| Q96FJ0 | |
| O75496 | |
| P08581 | |
| O43929 | |
| | 15996 |
| P46531 | |
| P11940 | |
| P49841 | |
| Q06330 | |
| P49682 | |
| | 15996 |
| P60953 | |
| P12931 | |
| Q9Y6K5 | |
| P60484;Q93009;Q00987;P42685 | |
| P12931 | |
| Q96RK4 | |
| 15996;Q8WU20 | |
| P19525 | |
| P49247 | |
| 15996;Q8WU20 | |
| P60484 | |
| P05919 | |
| | 15996 |
| | 15996 |
| | 15996 |
| | 15996 |
| | 15996 |

Q06330

15996
15996
P38936

15996

15996
Q13501;Q9C0C7

Q13501

Q00987
15996
15996
P12931
15996
P24385

P06400
Q13185
15996
Q16656;P11474
15996
Q13501
Q96LC9

P28749
Q06330
15996
P35222
P08581;P12931
15996
15996
Q08050
P43034;15996
15996;Q8WU20
P08034
Q06330
P49841
Q13224;Q14957;Q12879
P49841
P11940;P06730
15996

| | |
|---|---|
| P12931 | |
| P98170 | |
| | 15996 |
| P12931 | |
| P49247 | |
| Q05193;15996 | |
| | 15996 |
| P98170 | |
| Q8N661 | |
| P98170 | |
| | 15996 |
| | 15996 |
| | 15996 |
| P16284 | |
| P16284 | |
| P35222 | |
| | 15996 |
| | 15996 |
| P08581;Q13224;Q12879 | |
| P49841 | |
| | 15996 |
| P49841 | |
| P10721 | |
| Q9NTJ5 | |
| P04040 | |
| Q8IWU4 | |
| P03496 | |
| Q92888;15996 | |
| | 15996 |
| | 15996 |
| P60484;Q93009;Q00987;P42685 | |
| | 15996 |
| | 15996 |
| P60484;Q93009;P42685 | |
| P28749;Q08999;P06400 | |
| P28749;Q08999;P06400 | |
| Q07955 | |
| O14746 | |
| | 15996 |
| Q13099 | |
| P08034 | |
| Q3SXY8 | |

P08238
P24385;P49841
Q13480
Q13541;P06730;P62753

15996
P11474
Q13480

15996
P42679
P49023;15996

P24385;Q13761
Q03135
Q9UIL1;P43034;15996
P06241;15996

O15360
Q9NR28;P98170

P06730;Q9UBU9
O43463
Q9UI95
P11474
Q06330

O75496;O43929
Q07666;P23246
P06241
P06730
P05919
Q13501;P41743
15996
Q14739
O43639
P61586
Q8WU20
P06730;Q9UBU9
Q13541;P11940;P06730

P42772
P61024

15996;Q8WU20
15996
P12931;15996

Q9UI95
P00533

| | |
|---|---|
| P00533 | |
| | 15996 |
| Q8IWU4 | |
| O75771 | |
| Q09028;P42771 | |
| Q15125 | |
| Q15125 | |
| P06241 | |
| Q92560 | |
| | 15996 |
| | 15996 |
| P38936 | |
| Q03135 | |
| P06730;Q9UBU9 | |
| | 15996 |
| O75771 | |
| Q13501 | |
| Q9NTJ5 | |
| P49023;Q96AC1;P50552 | |
| O00308;P67809 | |
| P08034 | |
| P49841 | |
| O14965 | |
| Q00987 | |
| P49841 | |
| | 15996 |
| P06730;Q9UBU9 | |
| P61586 | |
| | 15996 |
| P49841 | |
| | 15996 |
| P31946 | |
| P08047 | |
| | 15996 |
| Q13562 | |
| O14746;P35222 | |
| P12931 | |
| Q8IWU4 | |
| Q13501;P41743 | |
| Q08050 | |
| P11940 | |

P04637
P00533;15996
Q13480

Q03135;P38936;Q08999
P63000;P06241
Q00987
15996
P06400;P42771;P08047;P41134

Q99873;O14965
P08238
15996;Q8WU20
15996;P48730
P49841;P35222
P09874;Q9UPN9
P12931;Q01196;Q99873
Q9Y297;P05919

Q03135

O14746

P06400
15996
P08238
O43559;15996;Q8WU20
P38936
15996;Q13480
Q13761;P12931
P54646

P11940
P11940
P24385
15996

Q95070
Q00987
Q9BZE9;15996;P63027
15996
Q12959;Q9NRD5

P31946

15996;P49841
Q16656;P11474
O43559;15996;Q8WU20
Q00535;P62136
O43559;15996;Q8WU20

O15160;O95602

Q12959;Q9NRD5
Q07955
P63104
15996
Q03135
15996
O43559;15996;Q8WU20
15996;P16333

15996
15996;Q8WU20
15996;Q8WU20
O43559;15996;Q8WU20
P08047
P54646
O14965
O60341
Q13526
Q13043;Q13188
O14746
Q08117

15996
Q9UQC2
P49841
O75496;O43929
P37231;P60484;P08581;Q13224;O00141;Q12879;Q14739
15996
15996
15996
O95070
P06241;15996
15996
Q9Y4Z2;Q13562
15996;P48730
15996;P08047
15996;P48730
P41743;Q01196;P46531

15996;Q8WU20
15996
P37231;P30281;O75376
Q13541;P11940;15996;P06730
P38936;Q00987
P38936;Q00987
Q13541;P11940;15996;P06730
P03496
15996
P00533;15996
Q15811;15996
P06400
P35548;Q15672;P49841
Q8IWU4;Q8IYI6;Q9NV70
15996
P11940
15996
P60484;Q9NTJ5;Q8N661
Q05513;15996
P63000;P06241
P38936;Q00987
P32239;P00533;15996
P16284;15996
15996
P63244;Q13077
Q03135
P49841
P38936;P61024
O00308;P67809;Q06330
P60953;P61586
15996
Q13501;P41743;15996
15996
15996
15996
P28749;Q08999
Q14677;15996
15996;P63027
15996

O60341
Q06330
15996
Q16656;Q3SXY8;Q96RK4;Q13099;15996;P11474
P05067
O15160;O95602
Q9UIL1;15996
15996
Q3SXY8;Q96RK4;15996
P16333

P06400

P12931;15996
15996;Q8WU20
P12931;15996;P10636-8
Q96LC9;Q9NR28;P98170
P06241;15996
Q13526
P38936;P06400
O75376;Q9Y297
Q13501;P54646;Q9C0C7;15996
O43463
P46531
P46531
P46531
P46531
P46531
P06400
P41743;Q01196;P46531
P16333
15996
P49841
P49247
Q9UQC2;P10721;15996

P49023;Q96AC1;P06241;P50552
O75386;P49841
Q99873
O15160;O95602
P00533;15996;Q13480
P54646;15996
Q3SXY8;Q96RK4;Q13099;15996
Q13224;P12931;Q12959;15996;Q9NRD5;P10636-8;Q14957;Q12879
Q15125;Q14739
O75771
P38936;P06400;P61024
P46531
Q13185;P49841
Q5S007
Q00987
Q9UQC2;P62993
P12931;15996
P32927;P26951;P63104
P08238;P63027
15996;Q8WU20
P06400
Q13526
Q13224;P12931;Q12959;15996;Q9NRD5;P10636-8;Q14957;Q12879
P46934;Q13526;O60573
O76003

Q99873
15996
P49023;Q03135;P06241;15996
Q07955;P06730;Q9UBU9
15996
15996
O95070
P38936;Q08050;O14965
P60484;Q5VVQ6;Q5T2D3
P38936;Q08050;O14965
15996;P63027
Q03135;P38936;Q08999;P04040
Q13501;15996
P60484
Q13224;O43559;15996;Q8WU20
Q13224;P12931;15996;P10636-8;Q14957;Q12879
15996
P49023;Q03135;P06241;15996
P19320;P16284
Q99873
Q99873
Q09028;Q13185;P45973
O00560
P12931;15996

Q13541;P31946;15996;P06730;P62753
P09874
P00533;P46531;15996;Q8WU20;P49841;Q00987
15996
P30622;Q13526;O60341;15996
Q14526;P04637
P46531
Q9NPI1;Q13526
P49023;Q96AC1;P50552
P08238
P52597;P52907;P60953
P00533;15996
P00533;15996
Q9UQC2;P06241;P36888;15996
15996;Q8NFF5
Q15116;P06241;15996
Q9NRD5
Q93009;Q9UER7;O00141;Q00987
P38936;Q01196
15996

P03496
O43559;15996;Q8WU20
P46934;Q13526;Q9Y6K5;O60573

P38936;Q99873;O14965
O43559;15996;Q8WU20
15996;P35222

15996

P42772;P08047
P00533;15996
P49682;Q9GZQ4

15996

O43559;15996;Q8WU20
O43559;15996;Q8WU20
P42679;15996;Q9Y316

15996

Q9UQC2;P62993
Q09028;P38936;P42772;P06400;P42771;P08047;P41134
P35222
P24385;Q05513;P12931;Q99873
P00533;P10721;P03372

O14746;Q9Y297;P49841;P35222;P98170;P63104
Q13263;Q9UER7
Q8N661
O96015;Q13224;P12931;O43559;15996;Q8WU20;P61586
P19525;P03496
15996;P05919
Q9Y297;P35222;P98170;P63104
P30281
P60484;P42685
15996;P48730
P24385;P38936;P49841;P61024
Q14677;Q9UIL1;Q9BZE9;P08034;P43034;15996;P48730;P05067;P63027
15996;P16333
O75496;P38936;P06400;P61024
P08581;P12931;15996;Q13480
15996;P05919
P52597;P52907;P60953

O96015;15996;Q8WU20;P61586
Q96C10;Q13526

P35222

15996

P46934;P12931;15996
P24385;Q13761;P38936;P12931
P15311;P12931;15996;P16333
P30281;15996
P52597;O75400;Q9BUJ2;P67809;Q07955
Q08117;P49841;P35222

15996
Q00535;15996;P62136
Q8IWU4;Q3KNW5;Q01196
P37231;O75928-2;P03372
P52597;O75400;Q9BUJ2;P67809;Q07955
O43559;15996;Q8WU20
P00533;O00560;15996
P60484;P16333
O43559;15996;Q8WU20
Q13501;Q09028;P38936;Q9C0C7;15996;P42772;P49841;P06400;P42771;P08047;P41134
Q92560;P60484;Q5VVQ6;Q5T2D3;Q96FJ0
Q09028;P38936;15996;P42772;P49841;P06400;P42771;P08047;P41134
15996;P05067
Q5SY16;P48730;P62753
O43559;15996;Q8WU20
O75771
Q12873;O75928-2;P45973
P09874;Q9UPN9;P42772;P08047
O75771
Q8IWU4;Q8IYI6;Q9NV70
P04637;Q93009;Q9UER7;O00141;Q00987
P31946;P11940;P63104
P32239;P00533;15996
P38936;Q00987
O43463;Q13185;P49841
O43639;O00560;15996
O75386;P49841
Q00535;P14316;15996;P47756;P45973
P12931;P11940;O43639;15996
Q14677;Q9UIL1;Q9BZE9;P08034;P43034;15996;P48730;P05067;P63027
Q05513;15996
Q5SY16;P48730;P62753
Q03135;P16284;15996;Q9NRD5;Q9BZW8
O43559;15996;Q8WU20
O75496;P38936;P28749;Q08999;P06400;P61024
15996;Q8NFF5
P63244;Q13077
O43559;15996;Q8WU20
P61296;P35548;P38936;Q15672;P49841;P06400
P43626;P19320
Q9UQC2;P06241;P36888;15996
P30622;Q13526;O60341
Q9UQC2;15996
P49023;O96015;P00533;P10721;Q13224;P06241;Q8WU20;P61586;Q13480;Q9UQC2;P42679;P46934;P08581;Q0313[illegible]
P46934;Q13526;Q9Y6K5;O60573
15996
P52597;O75400;Q9BUJ2;P67809;Q07955;P06730;Q9UBU9
Q5SY16;P48730;P62753
P52597;P52907;P60953
15996;P49247

P60484;P43626;Q15116;P06241;P19320;15996;P16333

15996

O14746;Q9Y297;Q08117;15996;P49841;P35222;P98170;P63104
Q13541;P11940;15996;P06730
Q14677;15996;P05067;P63027
P63000;Q9Y297;P06241;15996;P05919
P09874;Q9UPN9;P42772;P08047
P63244;Q13077
O75771
P12931;15996
O75376
O75376
P49023;Q5S007;Q07666;P23246;15996
P49023;Q5S007;Q07666;P23246;15996
P24385;O75496;P38936;P28749;Q08999;Q08050;O14965;P49841;P06400;P61024
Q15811;Q05193;O43639;O00560;15996
P09874;Q9UPN9;P42772;P08047
Q13501;P63244;P41743;Q13077;15996
P49682;Q9GZQ4
P19320;P16284
P00533;P10721;P38936;P03372
Q00535;Q08379;Q06830;P05067
Q00535;Q08379;Q06830;P05067
Q8IWU4;Q3KNW5;Q01196
Q9NPI1;P60484;Q13526;P04637;P38936;Q93009;Q9UER7;O00141;Q99873;O14965;Q00987;Q92574
P52597;Q9UQC2;P52907;P06241;P60953;P32927;P36888;15996;P62993;P26951;P63104
P41743;O00308;Q01196;P67809;P46531;Q06330
Q96LC9;Q9NR28;P35222;P98170
O95402;Q13526;Q09028;Q13761;P35548;P10721;Q13224;P38936;Q13185;Q08999;P09874;P04040;O14965;Q00987
Q96LC9;Q9NR28;P35222;P98170

15996

P24385;O60341;Q99873;P08047
P63000;Q9Y297;P06241;15996;P05919
Q15125;P08047;Q14739
P49682;Q9GZQ4
O95402;Q13526;Q09028;Q13761;P35548;P10721;Q13224;P38936;Q13185;Q08999;P09874;P04040;O14965;Q00987
Q00535;P12931;15996;P62136
O95402;O43463;O95602;Q13526;Q09028;Q13761;P35548;P10721;Q13224;P38936;Q13185;Q08999;P09874;P04040
Q9NPI1;Q13526;P04637;Q93009;Q9UER7;O00141;Q00987
Q00535;Q05513;P14316;Q03135;P06241;P16284;15996;Q9NRD5;P47756;Q9BZW8;P45973
O15360;O75771;Q9UI95
P24385;O14746;O75496;P38936;P28749;Q08999;Q08050;O14965;P49841;Q00987;P06400;P61024
Q15811;Q05193;P00533;P06241;Q8WU20;P49841;P15311;P12931;P11940;O43639;O00560;15996;P16333
P24385;Q05513;O60341;P12931;Q99873;P08047
P52597;Q13526;P06241;P60953;P32927;P62993;P26951;P63104;Q9UQC2;P46934;P52907;Q9Y6K5;O60573;P36888;[illegible]
P60484;Q15125;Q03135;Q9NTJ5;Q8N661;P11474;P08047;Q14739
P24385;Q05513;O60341;P12931;Q99873;P08047
Q14526;P04637;Q5VVQ6;Q12873;O75928-2;Q5T2D3;P09874;Q13263;Q9UER7;O14965;Q00987;P48730;P03372;P45[illegible]
P63000;P19525;P08238;P08581;Q9Y297;P06241;P12931;15996;P03496;P05919;P63027
P37231;Q14526;P04637;Q12873;O75928-2;P09874;Q13263;Q9UER7;O14965;Q00987;P03372;P45973

P37231;Q14526;P04637;Q12873;O75928-2;P09874;Q13263;Q9UER7;O14965;Q00987;P03372;P45973
P32239;Q00535;P00533;P12931;P60953;Q92888;15996;P62136;P61586
Q9UQC2;Q96C10;Q13526;P08238;15996;P35222
P52597;P31946;O75400;Q9BUJ2;Q07955;P06730;Q5SY16;P48730;Q9UBU9;P62753;P63104;P11940;P67809;15996
P52597;Q96C10;Q13526;P08238;Q15116;P06241;P60953;P32927;P62993;P26951;P63104;Q9UQC2;P46934;P60484;
Q8IWU4;O76003;Q3KNW5;Q01196;15996
Q15811;Q05193;P00533;P38936;P06241;Q9Y4Z2;Q13562;Q8WU20;P49841;P15311;P37231;P30281;O75376;P12931
Q13541;Q12873;P09874;Q13263;O14965;Q00987;P03372;P45973;Q92560;P60484;Q8IYI6;Q9NV70;Q96FJ0;Q14526;
P49682;P32239;Q9GZQ4;Q00535;P00533;P12931;P60953;Q92888;15996;P62136;P61586
P08238;Q08379;P00533;Q9Y297;P06241;P46531;Q8WU20;P49841;P49247;Q00987;P03496;P05067;P63027;P63000;
Q13501;Q13541;Q13224;Q9Y297;Q07666;Q08117;P06241;P09874;Q13188;Q92888;Q00987;Q9UQC2;P60484;O147
P60484;Q15125;Q03135;Q9NTJ5;Q8N661;15996;Q8NFF5;P11474;P49247;P08047;Q14739;P63027

**Found reaction identifiers**

R-HSA-6794338;R-HSA-6794336;R-HSA-6794349;R-HSA-6794348;R-HSA-6797803;R-HSA-6794346;R-HSA-6794345;R-H
R-HSA-6794338;R-HSA-6794336;R-HSA-6797568;R-HSA-6794349;R-HSA-6794348;R-HSA-6794346;R-HSA-6794345;R-H
R-HSA-451283;R-HSA-451311
R-HSA-451283;R-HSA-451311
R-HSA-8849878;R-HSA-8849891;R-HSA-8849906;R-HSA-8855648;R-HSA-8849882
R-HSA-6798258;R-HSA-6797810;R-HSA-6800380;R-HSA-6797803;R-HSA-388824
R-HSA-9630795
R-HSA-5334052
R-HSA-9630795
R-HSA-451283;R-HSA-451311
R-HSA-198609;R-HSA-2321904;R-HSA-2318752;R-HSA-5632993;R-HSA-380979;R-HSA-165179;R-HSA-5632732;R-HSA-
R-HSA-8944650;R-HSA-8944522;R-HSA-2318752;R-HSA-8944684;R-HSA-8945709;R-HSA-8944706;R-HSA-8944483;R-H
R-HSA-9008894
R-HSA-376134
R-HSA-391152;R-HSA-391150
R-HSA-193668;R-HSA-193646
R-HSA-888589;R-HSA-9609746;R-HSA-9609744;R-HSA-9611030;R-HSA-6794338;R-HSA-6794336;R-HSA-421007;R-HSA
R-HSA-5682794
R-HSA-196017
R-HSA-68610
R-HSA-8944104;R-HSA-8943817;R-HSA-6807077;R-HSA-8944397;R-HSA-8944078;R-HSA-8944047;R-HSA-8944099;R-H
R-HSA-210913;R-HSA-210886;R-HSA-210921;R-HSA-210920
R-HSA-380316;R-HSA-380508
R-HSA-196017;R-HSA-190519;R-HSA-190829
R-HSA-8951676
R-HSA-5660013
R-HSA-5660015
R-HSA-8853405;R-HSA-8853419
R-HSA-77589;R-HSA-77591;R-HSA-77590;R-HSA-77593;R-HSA-77592
R-HSA-168896
R-HSA-8878193;R-HSA-8878220
R-HSA-9630795
R-HSA-9630795
R-HSA-9630795
R-HSA-5660015;R-HSA-5660013
R-HSA-5228578
R-HSA-8938217;R-HSA-8937989
R-HSA-5250892
R-HSA-5250606
R-HSA-5618080
R-HSA-9617326
R-HSA-2220978
R-HSA-2220978
R-HSA-432164;R-HSA-438037;R-HSA-432162;R-HSA-432172;R-HSA-9614185
R-HSA-9012761;R-HSA-193696;R-HSA-193668;R-HSA-193646
R-HSA-9012761;R-HSA-193696
R-HSA-194796
R-HSA-380311;R-HSA-380455;R-HSA-380283
R-HSA-187746

R-HSA-428052
R-HSA-2317387
R-HSA-77589;R-HSA-77591;R-HSA-77590;R-HSA-77593;R-HSA-77592
R-HSA-109817
R-HSA-446221
R-HSA-2666278
R-HSA-2666278
R-HSA-8952289
R-HSA-8865994
R-HSA-9645137;R-HSA-9645126;R-HSA-9645134
R-HSA-72722
R-HSA-9032798
R-HSA-6785636
R-HSA-9627106;R-HSA-114307
R-HSA-8849350;R-HSA-8849353;R-HSA-8850041
R-HSA-165162;R-HSA-165182
R-HSA-428535
R-HSA-8875591;R-HSA-8875568
R-HSA-5626982
R-HSA-210836
R-HSA-5250962
R-HSA-2730752
R-HSA-2730752
R-HSA-2213248;R-HSA-2130641;R-HSA-8951498;R-HSA-2130725;R-HSA-2130619
R-HSA-1296127
R-HSA-5672329
R-HSA-1250498;R-HSA-1306972
R-HSA-380272;R-HSA-380303
R-HSA-380294;R-HSA-380272;R-HSA-380303
R-HSA-5663252
R-HSA-392870
R-HSA-1592233;R-HSA-1592244
R-HSA-549241
R-HSA-196929
R-HSA-194913;R-HSA-194854;R-HSA-195146
R-HSA-442732;R-HSA-442760
R-HSA-8865774
R-HSA-2485180
R-HSA-210426;R-HSA-210430;R-HSA-210444
R-HSA-428917;R-HSA-428750
R-HSA-9627350
R-HSA-5623508;R-HSA-5623521
R-HSA-437192
R-HSA-9017441
R-HSA-380311;R-HSA-380455;R-HSA-380294;R-HSA-380272;R-HSA-380303;R-HSA-380283
R-HSA-8853429;R-HSA-8853444
R-HSA-9627107;R-HSA-9627104;R-HSA-9627350;R-HSA-9627369
R-HSA-6803411
R-HSA-6803411
R-HSA-8935785;R-HSA-8937728;R-HSA-8937744;R-HSA-8937682;R-HSA-8938867;R-HSA-8938853;R-HSA-8937767;R-H

R-HSA-167415;R-HSA-167429
R-HSA-9017797
R-HSA-5626228;R-HSA-5626227;R-HSA-5626220;R-HSA-5626223;R-HSA-5626681;R-HSA-5617816;R-HSA-5626699;R-H
R-HSA-6800298
R-HSA-400092
R-HSA-139908;R-HSA-139909
R-HSA-177938
R-HSA-2316434;R-HSA-199456;R-HSA-1676082
R-HSA-110322
R-HSA-1168636
R-HSA-8878117;R-HSA-8878130;R-HSA-8952128;R-HSA-8878143;R-HSA-8878186
R-HSA-74986
R-HSA-392195
R-HSA-8875371
R-HSA-549241
R-HSA-113503
R-HSA-169904
R-HSA-1675866;R-HSA-199456;R-HSA-1675773;R-HSA-1676124;R-HSA-1675921;R-HSA-1676145;R-HSA-1676082;R-HS
R-HSA-139918
R-HSA-9036970
R-HSA-72180;R-HSA-77592
R-HSA-9026891
R-HSA-9619803
R-HSA-9033292
R-HSA-5686318
R-HSA-392152
R-HSA-432295
R-HSA-426240
R-HSA-171026
R-HSA-5250962;R-HSA-5250892;R-HSA-5228578;R-HSA-5250606;R-HSA-194796
R-HSA-9612219;R-HSA-1973956
R-HSA-164616
R-HSA-388828;R-HSA-389759;R-HSA-389758
R-HSA-9619542;R-HSA-9619515
R-HSA-113503
R-HSA-446322
R-HSA-947535
R-HSA-9032798;R-HSA-9032582
R-HSA-6788611;R-HSA-71037
R-HSA-6787533
R-HSA-177479;R-HSA-177501
R-HSA-416588
R-HSA-72663;R-HSA-72669
R-HSA-5654647
R-HSA-171026;R-HSA-5686071
R-HSA-168896
R-HSA-5654426
R-HSA-429955;R-HSA-429992
R-HSA-3000319;R-HSA-2574845;R-HSA-3000310;R-HSA-2574840
R-HSA-389954

R-HSA-380979;R-HSA-380927
R-HSA-114261;R-HSA-114252
R-HSA-190681;R-HSA-190687
R-HSA-5654600
R-HSA-399938
R-HSA-204549
R-HSA-416588
R-HSA-4839734
R-HSA-6807126;R-HSA-6807106;R-HSA-6807105;R-HSA-6807118
R-HSA-4839734
R-HSA-5654402
R-HSA-388829
R-HSA-6809006;R-HSA-6809003
R-HSA-446322
R-HSA-1369114;R-HSA-1369115
R-HSA-3928651
R-HSA-5696547
R-HSA-69299
R-HSA-8875374
R-HSA-68610
R-HSA-1474146
R-HSA-157632;R-HSA-157926;R-HSA-157353
R-HSA-156808
R-HSA-399951
R-HSA-2197588
R-HSA-374248
R-HSA-8849157;R-HSA-72180;R-HSA-72185
R-HSA-70600;R-HSA-70589
R-HSA-109858
R-HSA-877308
R-HSA-8964634
R-HSA-8876948;R-HSA-112333
R-HSA-109857
R-HSA-8985157
R-HSA-8943817;R-HSA-6807206;R-HSA-6807077;R-HSA-8944650;R-HSA-8944522;R-HSA-2318752;R-HSA-198599;R-H[illegible]
R-HSA-9011241
R-HSA-5624125
R-HSA-170979;R-HSA-170964
R-HSA-168896
R-HSA-177784;R-HSA-71306
R-HSA-8952289
R-HSA-6802935;R-HSA-6802934;R-HSA-6802933;R-HSA-6802932;R-HSA-6802927
R-HSA-170979;R-HSA-169904;R-HSA-170964
R-HSA-199456;R-HSA-1675773;R-HSA-1676145;R-HSA-1676082;R-HSA-1675810;R-HSA-1676149;R-HSA-1676134
R-HSA-3149440;R-HSA-3149432;R-HSA-3149433
R-HSA-177945
R-HSA-9029158
R-HSA-825631
R-HSA-1250383
R-HSA-77081

R-HSA-9013631
R-HSA-9008894
R-HSA-418579
R-HSA-111524
R-HSA-6803411
R-HSA-8847635;R-HSA-8847544
R-HSA-71401;R-HSA-71775
R-HSA-1791150;R-HSA-1791107;R-HSA-1791121
R-HSA-5671993
R-HSA-180687
R-HSA-1632857;R-HSA-1632843;R-HSA-5205673;R-HSA-5672017;R-HSA-5678315;R-HSA-5678313
R-HSA-997309
R-HSA-1791107
R-HSA-205008
R-HSA-1791150;R-HSA-1791107;R-HSA-1791121
R-HSA-2317387;R-HSA-2399981;R-HSA-2399982;R-HSA-2400009
R-HSA-442732;R-HSA-442760
R-HSA-416530
R-HSA-445367;R-HSA-9612085;R-HSA-9618750
R-HSA-381727;R-HSA-381706
R-HSA-8952058
R-HSA-6792572
R-HSA-4647594
R-HSA-427383
R-HSA-114552;R-HSA-114558
R-HSA-1592249;R-HSA-1592250;R-HSA-1592252;R-HSA-2466369;R-HSA-1605535;R-HSA-2466370;R-HSA-1592242;R-[illegible]
R-HSA-8851877;R-HSA-8851827;R-HSA-8851899
R-HSA-5205673
R-HSA-139908;R-HSA-139909;R-HSA-139918
R-HSA-508451;R-HSA-508513
R-HSA-8868071;R-HSA-8866279;R-HSA-8868230;R-HSA-8866275;R-HSA-8867756;R-HSA-8868236;R-HSA-8868651;R-[illegible]
R-HSA-2484822;R-HSA-375302;R-HSA-380316;R-HSA-380508;R-HSA-1638803;R-HSA-2468287;R-HSA-1638821
R-HSA-5638157;R-HSA-4827383
R-HSA-8964525
R-HSA-2197588
R-HSA-111524
R-HSA-3134904;R-HSA-3134901;R-HSA-3134883;R-HSA-3134914
R-HSA-8876948;R-HSA-112333;R-HSA-8875371
R-HSA-428917;R-HSA-418579;R-HSA-392195;R-HSA-428750
R-HSA-1250463
R-HSA-4088141;R-HSA-4088024
R-HSA-8849350;R-HSA-6811414;R-HSA-8849353;R-HSA-8850041
R-HSA-171026;R-HSA-187746;R-HSA-170979;R-HSA-5686071;R-HSA-169904;R-HSA-170964
R-HSA-190693;R-HSA-190681;R-HSA-190687
R-HSA-9013693
R-HSA-4791278
R-HSA-9609746;R-HSA-9610195;R-HSA-9609744;R-HSA-9611030;R-HSA-9610327;R-HSA-9610270;R-HSA-9610879;R-[illegible]
R-HSA-4791278
R-HSA-429955;R-HSA-429992
R-HSA-432195

R-HSA-8854302
R-HSA-418863;R-HSA-418859
R-HSA-9627078
R-HSA-156930
R-HSA-8874083
R-HSA-5660015;R-HSA-5660013
R-HSA-3928654;R-HSA-3928642;R-HSA-3928628
R-HSA-354173;R-HSA-939265
R-HSA-114261;R-HSA-9627078;R-HSA-114252
R-HSA-8874435
R-HSA-9627078
R-HSA-166271
R-HSA-749453
R-HSA-74885
R-HSA-1474146
R-HSA-2316434;R-HSA-1676082
R-HSA-210290;R-HSA-210304;R-HSA-210294;R-HSA-210277;R-HSA-210283;R-HSA-210285
R-HSA-210294;R-HSA-210277;R-HSA-432143;R-HSA-435244
R-HSA-202969
R-HSA-389954
R-HSA-5419268
R-HSA-9006507;R-HSA-8986940;R-HSA-9020513
R-HSA-4839746
R-HSA-5419264
R-HSA-3772434;R-HSA-3772436
R-HSA-4839746
R-HSA-1433508
R-HSA-1676168;R-HSA-1676174
R-HSA-6802935;R-HSA-6802934;R-HSA-6802933;R-HSA-6802932;R-HSA-6802927
R-HSA-1676124
R-HSA-9622738;R-HSA-9623168;R-HSA-9623280;R-HSA-9623281;R-HSA-9623192;R-HSA-9623240
R-HSA-437136
R-HSA-192841
R-HSA-1675866;R-HSA-1676020;R-HSA-1675910
R-HSA-114544;R-HSA-114548;R-HSA-419166;R-HSA-751029
R-HSA-180687
R-HSA-1855205
R-HSA-74885;R-HSA-2485180
R-HSA-8943817;R-HSA-6807206;R-HSA-6807077;R-HSA-8944650;R-HSA-8944522;R-HSA-2318752;R-HSA-198599;R-H[illegible]
R-HSA-70241;R-HSA-372819
R-HSA-6782434
R-HSA-8943817;R-HSA-6807206;R-HSA-6807077;R-HSA-8944650;R-HSA-8944522;R-HSA-2318752;R-HSA-8851011;R-[illegible]
R-HSA-182594;R-HSA-8942607;R-HSA-9018017;R-HSA-69227;R-HSA-8941895;R-HSA-8941915;R-HSA-1363328;R-HSA-[illegible]
R-HSA-182594;R-HSA-8942607;R-HSA-9018017;R-HSA-69227;R-HSA-8941895;R-HSA-8941915;R-HSA-1363328;R-HSA-[illegible]
R-HSA-75080
R-HSA-164616
R-HSA-432195
R-HSA-5624949;R-HSA-5625424;R-HSA-5625426;R-HSA-5625421;R-HSA-5617820;R-HSA-5624952;R-HSA-5625416
R-HSA-196017;R-HSA-190519;R-HSA-190693;R-HSA-190681;R-HSA-190687;R-HSA-190829
R-HSA-5624953

R-HSA-5336420
R-HSA-75824;R-HSA-75820;R-HSA-75823
R-HSA-8851908;R-HSA-8851919
R-HSA-165726;R-HSA-165708
R-HSA-1112690;R-HSA-1112708;R-HSA-1112703
R-HSA-399995
R-HSA-560473;R-HSA-1655848;R-HSA-1655850;R-HSA-1592238;R-HSA-1655826;R-HSA-560517
R-HSA-9027281
R-HSA-1791107
R-HSA-416530
R-HSA-1963563
R-HSA-8848596;R-HSA-8848611;R-HSA-8848606;R-HSA-8848618
R-HSA-1296127
R-HSA-4411367;R-HSA-8951442;R-HSA-8951443;R-HSA-8951428;R-HSA-4411357
R-HSA-163418
R-HSA-8849350;R-HSA-6811414;R-HSA-6811429;R-HSA-8847635;R-HSA-6814096;R-HSA-8849353;R-HSA-8850041;R-H
R-HSA-389348;R-HSA-389350;R-HSA-389354
R-HSA-109857;R-HSA-109858
R-HSA-6785126
R-HSA-9627106;R-HSA-9627107;R-HSA-114307;R-HSA-9627104;R-HSA-114261;R-HSA-9627350;R-HSA-9627078;R-HSA
R-HSA-381799
R-HSA-158484;R-HSA-111439
R-HSA-427528
R-HSA-110322
R-HSA-560473;R-HSA-1655848;R-HSA-1655850;R-HSA-1592238;R-HSA-1655826;R-HSA-560517
R-HSA-212356;R-HSA-350058
R-HSA-448962;R-HSA-448963
R-HSA-69299;R-HSA-68610
R-HSA-8849463;R-HSA-8848993;R-HSA-8848975;R-HSA-8848939
R-HSA-8855237
R-HSA-72619
R-HSA-3149440;R-HSA-3149432;R-HSA-3149433;R-HSA-3149454
R-HSA-193684;R-HSA-193694
R-HSA-392152
R-HSA-9018594
R-HSA-8985812
R-HSA-202692
R-HSA-8855508;R-HSA-8855564
R-HSA-77594;R-HSA-158447
R-HSA-72631;R-HSA-156808;R-HSA-72622
R-HSA-2671742;R-HSA-2671747
R-HSA-182594;R-HSA-3857348
R-HSA-187545
R-HSA-888589;R-HSA-380901;R-HSA-380869;R-HSA-380905;R-HSA-374922;R-HSA-210444;R-HSA-372529;R-HSA-3748
R-HSA-8853310;R-HSA-8853307
R-HSA-156930;R-HSA-156913;R-HSA-156909
R-HSA-8875591;R-HSA-8874083;R-HSA-8875568
R-HSA-8866279;R-HSA-8866275;R-HSA-8866283
R-HSA-110322
R-HSA-1248677

R-HSA-1248677
R-HSA-180687
R-HSA-437136
R-HSA-381799
R-HSA-5685318
R-HSA-182594;R-HSA-3229089;R-HSA-3223200;R-HSA-3209114;R-HSA-3240957;R-HSA-3229102
R-HSA-141409;R-HSA-141431;R-HSA-141422;R-HSA-141439
R-HSA-141409;R-HSA-141431;R-HSA-141422;R-HSA-141439
R-HSA-6807052
R-HSA-195690
R-HSA-8981534
R-HSA-5690790;R-HSA-5690785;R-HSA-5689649;R-HSA-5689630
R-HSA-4411351;R-HSA-4411367;R-HSA-4411357;R-HSA-4411372
R-HSA-555065
R-HSA-8876615;R-HSA-8876454;R-HSA-8876837;R-HSA-8877475;R-HSA-8876193;R-HSA-8877760;R-HSA-8877998;R-[illegible]
R-HSA-8865256
R-HSA-163418
R-HSA-77587;R-HSA-159050
R-HSA-1112690;R-HSA-1112708;R-HSA-1112703
R-HSA-1168636
R-HSA-8931974
R-HSA-5685318
R-HSA-5205673
R-HSA-1675921;R-HSA-1676114;R-HSA-1676005;R-HSA-1676133
R-HSA-430341;R-HSA-446322;R-HSA-446364
R-HSA-9021520;R-HSA-9017797
R-HSA-196017;R-HSA-190519;R-HSA-190693;R-HSA-190681;R-HSA-190687;R-HSA-190829
R-HSA-5610730
R-HSA-8853496
R-HSA-198609;R-HSA-198599
R-HSA-5610732
R-HSA-210977;R-HSA-204873
R-HSA-77587;R-HSA-158484;R-HSA-159050;R-HSA-77594;R-HSA-158447;R-HSA-111439
R-HSA-202692
R-HSA-71401;R-HSA-71775
R-HSA-5334052;R-HSA-4791278;R-HSA-4839746;R-HSA-4839734
R-HSA-9622579;R-HSA-9622604;R-HSA-9622572
R-HSA-3244614
R-HSA-450394
R-HSA-1655833;R-HSA-2426148;R-HSA-1655832;R-HSA-2426151;R-HSA-2426150;R-HSA-1655837;R-HSA-2426147;R-[illegible]
R-HSA-5419264;R-HSA-5419268
R-HSA-211346
R-HSA-3299569;R-HSA-3322422;R-HSA-4411367;R-HSA-3451153;R-HSA-3364026;R-HSA-3322427;R-HSA-3322424;R-[illegible]
R-HSA-9603419
R-HSA-437136
R-HSA-193684;R-HSA-193694
R-HSA-4088134;R-HSA-4088439;R-HSA-4088309;R-HSA-4088162;R-HSA-4088130;R-HSA-4088306;R-HSA-4088307
R-HSA-372529;R-HSA-372505
R-HSA-380901;R-HSA-380905
R-HSA-450551

R-HSA-6804188;R-HSA-6804762
R-HSA-2179387;R-HSA-2179407
R-HSA-177923;R-HSA-177920;R-HSA-177926;R-HSA-177924;R-HSA-177944;R-HSA-177935
R-HSA-8865605
R-HSA-6803411;R-HSA-9624985;R-HSA-9620813
R-HSA-200952;R-HSA-200908
R-HSA-6804741;R-HSA-6804955
R-HSA-389954
R-HSA-182594;R-HSA-3209109;R-HSA-3209096;R-HSA-3209114;R-HSA-3209165
R-HSA-109699
R-HSA-141409;R-HSA-141431;R-HSA-141422;R-HSA-141439
R-HSA-374899;R-HSA-374922
R-HSA-3215152;R-HSA-6791235
R-HSA-874087
R-HSA-5655351;R-HSA-5655347
R-HSA-6809006;R-HSA-203977;R-HSA-6809003;R-HSA-5694441;R-HSA-5694409
R-HSA-201685;R-HSA-5323526
R-HSA-2187330;R-HSA-870538;R-HSA-2187325;R-HSA-870449
R-HSA-8937097;R-HSA-8937037;R-HSA-8937728;R-HSA-8937989;R-HSA-8938887;R-HSA-8937050;R-HSA-8937744;R-H
R-HSA-180606;R-HSA-180591
R-HSA-1549564
R-HSA-202144;R-HSA-203712
R-HSA-2467809;R-HSA-2467811
R-HSA-164616
R-HSA-1855205
R-HSA-380869;R-HSA-380574
R-HSA-2172666
R-HSA-5621573
R-HSA-874087
R-HSA-9028730;R-HSA-9031981;R-HSA-9032067;R-HSA-9032402
R-HSA-8852362
R-HSA-9027281;R-HSA-9029158
R-HSA-8952408;R-HSA-8952382;R-HSA-8952399;R-HSA-8937807;R-HSA-8937792;R-HSA-8865454;R-HSA-8952419;R-H
R-HSA-9613513;R-HSA-9613565
R-HSA-5654709;R-HSA-5654643;R-HSA-5654641
R-HSA-5654669;R-HSA-5654714;R-HSA-5654667
R-HSA-927830
R-HSA-927830
R-HSA-9623355
R-HSA-389954
R-HSA-5666160;R-HSA-5666129;R-HSA-5666169
R-HSA-2484822;R-HSA-1638803;R-HSA-2468287;R-HSA-1638821
R-HSA-5654692;R-HSA-5654596;R-HSA-5654594
R-HSA-1791143;R-HSA-1791092
R-HSA-5228523;R-HSA-3000434
R-HSA-1449574;R-HSA-2255342;R-HSA-2255343
R-HSA-5654679;R-HSA-1549564;R-HSA-5654730
R-HSA-983422
R-HSA-416320;R-HSA-421007;R-HSA-416639;R-HSA-416985
R-HSA-5654622;R-HSA-5654620;R-HSA-5654697

R-HSA-450406;R-HSA-482788
R-HSA-5654734;R-HSA-5654684;R-HSA-1549564
R-HSA-399938;R-HSA-399951;R-HSA-416588
R-HSA-1592249;R-HSA-1592250;R-HSA-1592252;R-HSA-2466369;R-HSA-1605535;R-HSA-2466370;R-HSA-1592242;R-H
R-HSA-5654631;R-HSA-8941628;R-HSA-5654623;R-HSA-5654413;R-HSA-5654633;R-HSA-5654409
R-HSA-180038;R-HSA-180047
R-HSA-5654422;R-HSA-5654663;R-HSA-5654658;R-HSA-8941613;R-HSA-5654655;R-HSA-5654651
R-HSA-2400009
R-HSA-2467809;R-HSA-2467811
R-HSA-73865
R-HSA-8987132;R-HSA-8987070;R-HSA-8987236;R-HSA-8987014
R-HSA-416320;R-HSA-421007;R-HSA-416639;R-HSA-416985
R-HSA-75096
R-HSA-450620
R-HSA-6811414
R-HSA-202144
R-HSA-70241;R-HSA-372819
R-HSA-5654571;R-HSA-5654587;R-HSA-5654584;R-HSA-5654392;R-HSA-8941623;R-HSA-5654569
R-HSA-373716;R-HSA-418850;R-HSA-418856
R-HSA-5654677;R-HSA-1549564;R-HSA-5654729
R-HSA-5654672;R-HSA-5654673;R-HSA-1549564
R-HSA-109699;R-HSA-109817
R-HSA-5655262;R-HSA-8853310;R-HSA-5655277;R-HSA-8853307
R-HSA-5655262;R-HSA-5655277
R-HSA-5654608;R-HSA-5654607;R-HSA-5654399;R-HSA-8941618;R-HSA-5654618;R-HSA-5654603
R-HSA-1655833;R-HSA-2426148;R-HSA-1655832;R-HSA-2426151;R-HSA-2426150;R-HSA-1655837;R-HSA-2426147;R-H
R-HSA-9613513;R-HSA-9613565
R-HSA-4655404
R-HSA-5625870;R-HSA-5625849
R-HSA-9626816
R-HSA-2028697;R-HSA-2028692
R-HSA-164616
R-HSA-4649028;R-HSA-5334050;R-HSA-8944349;R-HSA-8944352;R-HSA-4641231;R-HSA-5229348
R-HSA-888589;R-HSA-917744
R-HSA-109699;R-HSA-109817
R-HSA-2730848
R-HSA-5250947
R-HSA-69299;R-HSA-68610
R-HSA-9615570;R-HSA-9615571;R-HSA-9006507;R-HSA-9022764;R-HSA-8986940;R-HSA-9017441;R-HSA-9020513;R-H
R-HSA-109699;R-HSA-5686318;R-HSA-109817
R-HSA-8944362;R-HSA-4411383;R-HSA-4411365;R-HSA-3965444
R-HSA-109699;R-HSA-5686318;R-HSA-109817
R-HSA-1791143;R-HSA-1791092
R-HSA-392054;R-HSA-391867
R-HSA-6787533;R-HSA-446221
R-HSA-211346;R-HSA-210913;R-HSA-210886;R-HSA-210836;R-HSA-210921;R-HSA-210920
R-HSA-203977;R-HSA-5694441;R-HSA-5694409
R-HSA-6814885;R-HSA-6814549;R-HSA-6814559;R-HSA-6814555;R-HSA-6814554;R-HSA-6807496
R-HSA-6809006;R-HSA-203977;R-HSA-6809003;R-HSA-5694441;R-HSA-5694409
R-HSA-1912412;R-HSA-9021357;R-HSA-9604703

R-HSA-5655339;R-HSA-5655241
R-HSA-109857;R-HSA-5672965;R-HSA-109858
R-HSA-1183058;R-HSA-560491;R-HSA-560493;R-HSA-381283;R-HSA-1183003;R-HSA-381262;R-HSA-560517;R-HSA-38
R-HSA-72722;R-HSA-72631;R-HSA-72663;R-HSA-72619;R-HSA-156808;R-HSA-72669;R-HSA-72622
R-HSA-6804741;R-HSA-6803411;R-HSA-6804955
R-HSA-6804741;R-HSA-6803411;R-HSA-6804955
R-HSA-72722;R-HSA-72631;R-HSA-72663;R-HSA-72619;R-HSA-156808;R-HSA-72669;R-HSA-72622
R-HSA-192841
R-HSA-8876615;R-HSA-8876454;R-HSA-8876837;R-HSA-8877475;R-HSA-8876193;R-HSA-8877760;R-HSA-8877998;R-H
R-HSA-445069;R-HSA-445064
R-HSA-3928632;R-HSA-4093336;R-HSA-3928592;R-HSA-3928644;R-HSA-3928612
R-HSA-8985460
R-HSA-8939929;R-HSA-9008475;R-HSA-8939900;R-HSA-9008476;R-HSA-9007759;R-HSA-8940032
R-HSA-437136;R-HSA-265177
R-HSA-2424476;R-HSA-2424477
R-HSA-9014610
R-HSA-5696197
R-HSA-1675866;R-HSA-199456;R-HSA-8874435;R-HSA-1675773;R-HSA-1676124;R-HSA-1675921;R-HSA-1676145;R-HS
R-HSA-442291;R-HSA-453200;R-HSA-5684801;R-HSA-442273;R-HSA-442314;R-HSA-437192;R-HSA-5684169
R-HSA-200952;R-HSA-200908
R-HSA-6804741;R-HSA-6803411;R-HSA-6804955
R-HSA-870269;R-HSA-2179387;R-HSA-2179407
R-HSA-392870;R-HSA-210294;R-HSA-210277;R-HSA-432143;R-HSA-435244;R-HSA-392152
R-HSA-205039
R-HSA-5357776;R-HSA-5357780
R-HSA-202144;R-HSA-203712
R-HSA-3371435
R-HSA-187828;R-HSA-187545
R-HSA-9013631;R-HSA-9021520;R-HSA-9017797
R-HSA-8964634;R-HSA-202692
R-HSA-6782434
R-HSA-9012761;R-HSA-193696;R-HSA-205008;R-HSA-193684;R-HSA-193668;R-HSA-193694;R-HSA-193646;R-HSA-205
R-HSA-1225957;R-HSA-1225951
R-HSA-1225957;R-HSA-1225951
R-HSA-109699;R-HSA-109817;R-HSA-109807
R-HSA-8964525;R-HSA-1363314;R-HSA-1363306
R-HSA-434362;R-HSA-421835
R-HSA-416530;R-HSA-265166;R-HSA-400092;R-HSA-381727;R-HSA-399995;R-HSA-381706
R-HSA-2424476;R-HSA-2424477
R-HSA-428052
R-HSA-5625870;R-HSA-5625849
R-HSA-9013693
R-HSA-6787533;R-HSA-446221
R-HSA-5623508;R-HSA-5625424;R-HSA-1605535;R-HSA-5625426;R-HSA-5617820;R-HSA-5617816;R-HSA-1368140;R-H
R-HSA-8871494;R-HSA-9010091;R-HSA-9010096;R-HSA-5692495;R-HSA-9010034
R-HSA-73865;R-HSA-74986
R-HSA-6811429;R-HSA-6814096
R-HSA-74885;R-HSA-2485180
R-HSA-5623508;R-HSA-5623521;R-HSA-5624125;R-HSA-5624953
R-HSA-430190

R-HSA-2172666
R-HSA-8867047;R-HSA-183072;R-HSA-8867044;R-HSA-8867041;R-HSA-182986;R-HSA-182990
R-HSA-9036970;R-HSA-9603419
R-HSA-5655269;R-HSA-5655326
R-HSA-445367;R-HSA-9619542;R-HSA-9612085;R-HSA-9619515;R-HSA-9619803;R-HSA-442732;R-HSA-9618750;R-HSA
R-HSA-9627106;R-HSA-9627107;R-HSA-114307;R-HSA-9627104;R-HSA-114261;R-HSA-9627350;R-HSA-9627078;R-HSA
R-HSA-389348;R-HSA-389350;R-HSA-389354
R-HSA-936462;R-HSA-936380
R-HSA-4647613;R-HSA-4647594
R-HSA-5663252;R-HSA-1368087;R-HSA-549475;R-HSA-5663258;R-HSA-400267;R-HSA-400342;R-HSA-1368069;R-HSA-
R-HSA-1632857;R-HSA-1632843;R-HSA-9626038;R-HSA-5205673;R-HSA-5672017;R-HSA-9613513;R-HSA-5678315;R-H
R-HSA-427528
R-HSA-2220978;R-HSA-2730752;R-HSA-2220988;R-HSA-2666278
R-HSA-2220988
R-HSA-2220988
R-HSA-2220988
R-HSA-2220988
R-HSA-8985460
R-HSA-1912412;R-HSA-9021357;R-HSA-9604703
R-HSA-9606151
R-HSA-109699;R-HSA-109817;R-HSA-109807
R-HSA-3371435
R-HSA-5660015;R-HSA-5660013
R-HSA-1433428;R-HSA-1433488;R-HSA-1562641;R-HSA-205321;R-HSA-1433471;R-HSA-205231;R-HSA-205234;R-HSA-
R-HSA-2467809;R-HSA-2467811
R-HSA-430341;R-HSA-446322;R-HSA-391152;R-HSA-8981534;R-HSA-446364;R-HSA-391150
R-HSA-5610726;R-HSA-5610732;R-HSA-5610730
R-HSA-5205867;R-HSA-5205799;R-HSA-3215448;R-HSA-5205861
R-HSA-73865;R-HSA-74986
R-HSA-8867047;R-HSA-177923;R-HSA-183072;R-HSA-177920;R-HSA-8867044;R-HSA-177926;R-HSA-8867041;R-HSA-1
R-HSA-9626038;R-HSA-9613513;R-HSA-9613565
R-HSA-5624949;R-HSA-5626228;R-HSA-5623508;R-HSA-5625424;R-HSA-5626227;R-HSA-5625426;R-HSA-5624125;R-H
R-HSA-888589;R-HSA-380869;R-HSA-9609746;R-HSA-9610195;R-HSA-9609744;R-HSA-9619542;R-HSA-9611030;R-HSA
R-HSA-194674;R-HSA-195690;R-HSA-6807052
R-HSA-5685318
R-HSA-188386;R-HSA-187828;R-HSA-187545
R-HSA-1912396;R-HSA-1912388;R-HSA-1912391;R-HSA-1912385
R-HSA-427383;R-HSA-5250947
R-HSA-8857565
R-HSA-2399981;R-HSA-2399982
R-HSA-8983425;R-HSA-8983394
R-HSA-186834;R-HSA-186819;R-HSA-186778
R-HSA-909738;R-HSA-912757;R-HSA-914036;R-HSA-914022;R-HSA-450074;R-HSA-879937
R-HSA-5250962;R-HSA-5336420;R-HSA-5250892;R-HSA-5228578;R-HSA-5250606;R-HSA-194796
R-HSA-5655351;R-HSA-5655269;R-HSA-5655347;R-HSA-5655262;R-HSA-5655326;R-HSA-8853310;R-HSA-5655277;R-H
R-HSA-2484822;R-HSA-375302;R-HSA-2467809;R-HSA-2467811;R-HSA-380316;R-HSA-380508;R-HSA-1638803;R-HSA-
R-HSA-8877692
R-HSA-9609746;R-HSA-9610195;R-HSA-9609744;R-HSA-9619542;R-HSA-9611030;R-HSA-9610327;R-HSA-416320;R-HS
R-HSA-936462;R-HSA-1678843;R-HSA-1678842;R-HSA-1169399
R-HSA-5690886

R-HSA-983147;R-HSA-983157;R-HSA-983156;R-HSA-983140
R-HSA-8936616;R-HSA-8937097;R-HSA-8937037;R-HSA-8936621;R-HSA-8936481;R-HSA-8936995;R-HSA-8937016;R-H
R-HSA-111524
R-HSA-5218830;R-HSA-5218829;R-HSA-202144;R-HSA-5218811;R-HSA-5218824;R-HSA-5218839;R-HSA-5218838;R-HS
R-HSA-77587;R-HSA-158484;R-HSA-159050;R-HSA-75096;R-HSA-77594;R-HSA-158447;R-HSA-111439
R-HSA-354173;R-HSA-939265
R-HSA-354173;R-HSA-939265
R-HSA-1791150;R-HSA-1791107;R-HSA-1791121;R-HSA-1791143;R-HSA-1791092
R-HSA-8853444;R-HSA-380294;R-HSA-4088439;R-HSA-380303;R-HSA-4088309;R-HSA-4088306;R-HSA-4088307;R-HSA
R-HSA-6781953;R-HSA-8873946
R-HSA-8853444;R-HSA-380294;R-HSA-4088439;R-HSA-380303;R-HSA-4088309;R-HSA-4088306;R-HSA-4088307;R-HSA
R-HSA-825631;R-HSA-416530;R-HSA-265166;R-HSA-400092;R-HSA-381727;R-HSA-399995;R-HSA-381706
R-HSA-9622738;R-HSA-9622579;R-HSA-9623168;R-HSA-9623280;R-HSA-9623281;R-HSA-6803411;R-HSA-9623192;R-H
R-HSA-205008;R-HSA-205039
R-HSA-199456
R-HSA-9026891;R-HSA-9028730;R-HSA-9031981;R-HSA-9033292;R-HSA-9032798;R-HSA-9032067;R-HSA-9032402;R-H
R-HSA-9609746;R-HSA-9610195;R-HSA-9609744;R-HSA-9619542;R-HSA-9611030;R-HSA-9610327;R-HSA-9619803;R-H
R-HSA-354173;R-HSA-939265
R-HSA-5218830;R-HSA-5218829;R-HSA-202144;R-HSA-5218811;R-HSA-5218824;R-HSA-5218839;R-HSA-5218838;R-HS
R-HSA-265428;R-HSA-210304;R-HSA-198941
R-HSA-8865605;R-HSA-5205867;R-HSA-5205799;R-HSA-3215448;R-HSA-5205861;R-HSA-5638157;R-HSA-4827383
R-HSA-8865605;R-HSA-5205867;R-HSA-5205799;R-HSA-3215448;R-HSA-5205861;R-HSA-5638157;R-HSA-4827383
R-HSA-9007465;R-HSA-9007464;R-HSA-9007640;R-HSA-9007467;R-HSA-8953452;R-HSA-9007646;R-HSA-9007283;R-H
R-HSA-373738
R-HSA-186834;R-HSA-186819;R-HSA-186778
R-HSA-5689950
R-HSA-380979;R-HSA-165162;R-HSA-165195;R-HSA-380927;R-HSA-165182;R-HSA-165726;R-HSA-5672824;R-HSA-165
R-HSA-4551604;R-HSA-4551768
R-HSA-6802927;R-HSA-5655262;R-HSA-5655326;R-HSA-2399981;R-HSA-2399982;R-HSA-4839746;R-HSA-1225951;R-H
R-HSA-2730840;R-HSA-2424477
R-HSA-9626816;R-HSA-194913;R-HSA-194854;R-HSA-5666160;R-HSA-5666129;R-HSA-195146;R-HSA-5625870;R-HSA-
R-HSA-4090281;R-HSA-2997706
R-HSA-1912396;R-HSA-157632;R-HSA-157926;R-HSA-1912388;R-HSA-1912391;R-HSA-157353;R-HSA-1912385
R-HSA-8877692;R-HSA-3222093
R-HSA-430341;R-HSA-446322;R-HSA-446364
R-HSA-874087
R-HSA-8950634;R-HSA-8950637;R-HSA-8950415
R-HSA-5637808;R-HSA-5637806;R-HSA-1247999
R-HSA-5637808;R-HSA-5637806;R-HSA-1247999
R-HSA-9606784;R-HSA-9645137;R-HSA-109699;R-HSA-9606624;R-HSA-6786789;R-HSA-5672965;R-HSA-9645126;R-HS
R-HSA-947535;R-HSA-196929
R-HSA-389941;R-HSA-389348;R-HSA-389350;R-HSA-388829;R-HSA-388828;R-HSA-389759;R-HSA-389758;R-HSA-3893
R-HSA-421007;R-HSA-416639;R-HSA-416985
R-HSA-6804741;R-HSA-6795460;R-HSA-198599;R-HSA-6793666;R-HSA-3222072;R-HSA-3215295;R-HSA-6795667;R-HS
R-HSA-9616214;R-HSA-9624668
R-HSA-2197697;R-HSA-2029467;R-HSA-2029451;R-HSA-2029445
R-HSA-997309;R-HSA-909726
R-HSA-192841
R-HSA-5654631;R-HSA-5654647;R-HSA-5654709;R-HSA-8941628;R-HSA-5654643;R-HSA-5654641;R-HSA-5654623;R-H
R-HSA-936462;R-HSA-1678843;R-HSA-1678842;R-HSA-1169399;R-HSA-8985157

R-HSA-3215152;R-HSA-6791235;R-HSA-6803411
R-HSA-5654422;R-HSA-5654663;R-HSA-5654658;R-HSA-8941613;R-HSA-5654655;R-HSA-5654669;R-HSA-5654714;R-H
R-HSA-3134904;R-HSA-3244614;R-HSA-3134901;R-HSA-3134883;R-HSA-3134914
R-HSA-5672965
R-HSA-2187303;R-HSA-2187309
R-HSA-1248677;R-HSA-5637808;R-HSA-1225957;R-HSA-5637806;R-HSA-1247999;R-HSA-1225951
R-HSA-981832;R-HSA-374248
R-HSA-392152
R-HSA-5654402;R-HSA-5654608;R-HSA-5654622;R-HSA-5654607;R-HSA-5654399;R-HSA-5654620;R-HSA-8941618;R-H
R-HSA-5654692;R-HSA-5654596;R-HSA-5654594;R-HSA-5654571;R-HSA-5654587;R-HSA-5654584;R-HSA-5654392;R-H
R-HSA-1250498;R-HSA-6785636;R-HSA-1963563;R-HSA-1250463;R-HSA-1306972
R-HSA-2201293;R-HSA-166271
R-HSA-71401;R-HSA-71775
R-HSA-508451;R-HSA-508513;R-HSA-8983425;R-HSA-8983394
R-HSA-182594;R-HSA-3229089;R-HSA-3223200;R-HSA-3209109;R-HSA-3209096;R-HSA-3209114;R-HSA-4647613;R-HS
R-HSA-202969
R-HSA-9632182;R-HSA-9021596;R-HSA-9623355;R-HSA-437192
R-HSA-8937369;R-HSA-8874797;R-HSA-8864395;R-HSA-8864698
R-HSA-877308
R-HSA-3299569;R-HSA-3322422;R-HSA-201669;R-HSA-3769393;R-HSA-3364026;R-HSA-3322427;R-HSA-3322424;R-HS
R-HSA-3903017;R-HSA-4085992;R-HSA-3782535
R-HSA-8874435
R-HSA-187746;R-HSA-170979;R-HSA-5686071;R-HSA-9026891;R-HSA-9036970;R-HSA-9031981;R-HSA-177479;R-HSA-
R-HSA-168896;R-HSA-192841
R-HSA-3149440;R-HSA-3149432;R-HSA-3149433;R-HSA-180687;R-HSA-3149454
R-HSA-3322434;R-HSA-3361751;R-HSA-5368580;R-HSA-3772430;R-HSA-3769393;R-HSA-3364042;R-HSA-3322431;R-H
R-HSA-5692764
R-HSA-6807206;R-HSA-8850997;R-HSA-8851011;R-HSA-8850961;R-HSA-8850945;R-HSA-8850992;R-HSA-8847968;R-H
R-HSA-6809006;R-HSA-6787533;R-HSA-203977;R-HSA-446221;R-HSA-6809003;R-HSA-5694441;R-HSA-5694409
R-HSA-75824;R-HSA-187828;R-HSA-187545;R-HSA-75820;R-HSA-75823
R-HSA-8868230;R-HSA-5229132;R-HSA-8868236;R-HSA-8877451;R-HSA-8847635;R-HSA-8876191;R-HSA-8854302;R-H
R-HSA-9606151;R-HSA-1168636
R-HSA-188386;R-HSA-69299;R-HSA-187828;R-HSA-187545;R-HSA-113503
R-HSA-8875591;R-HSA-8851877;R-HSA-8851908;R-HSA-8874083;R-HSA-8851827;R-HSA-8875568;R-HSA-8851919;R-H
R-HSA-3149440;R-HSA-3149432;R-HSA-3149433;R-HSA-180687;R-HSA-3149454
R-HSA-8950634;R-HSA-8950637;R-HSA-8950415
R-HSA-2201293;R-HSA-166271
R-HSA-171026;R-HSA-187746;R-HSA-170979;R-HSA-5686071;R-HSA-169904;R-HSA-177479;R-HSA-170964;R-HSA-202
R-HSA-936462;R-HSA-936380;R-HSA-168935
R-HSA-6798747;R-HSA-6800426
R-HSA-202969
R-HSA-2197697;R-HSA-2029467;R-HSA-2029451;R-HSA-2029445
R-HSA-9612593;R-HSA-9612219;R-HSA-1973956;R-HSA-1250383
R-HSA-8878117;R-HSA-4411367;R-HSA-8952399;R-HSA-8937807;R-HSA-8937792;R-HSA-8952128;R-HSA-8865454;R-H
R-HSA-373716;R-HSA-374662;R-HSA-418850;R-HSA-418863;R-HSA-418859;R-HSA-418856
R-HSA-109857;R-HSA-5672965;R-HSA-109858;R-HSA-5692764
R-HSA-72160;R-HSA-72130;R-HSA-72231;R-HSA-156661;R-HSA-72107;R-HSA-72139;R-HSA-72124;R-HSA-72143;R-HSA
R-HSA-195251;R-HSA-4649028;R-HSA-5334050;R-HSA-8944349;R-HSA-8944352;R-HSA-4641231;R-HSA-195304;R-HSA
R-HSA-166271
R-HSA-166271

R-HSA-8944362;R-HSA-4411383;R-HSA-4411365;R-HSA-3965444
R-HSA-180038;R-HSA-167415;R-HSA-167429;R-HSA-180047
R-HSA-549241;R-HSA-437136;R-HSA-433089
R-HSA-4717461;R-HSA-4090408;R-HSA-4719423;R-HSA-4719447
R-HSA-72160;R-HSA-72130;R-HSA-72231;R-HSA-156661;R-HSA-75080;R-HSA-72107;R-HSA-72139;R-HSA-72124;R-HS
R-HSA-5654422;R-HSA-8941613;R-HSA-5654655;R-HSA-5654684;R-HSA-1549564;R-HSA-5654714;R-HSA-5654426;R-H
R-HSA-555065;R-HSA-376134;R-HSA-445069;R-HSA-445064;R-HSA-373738
R-HSA-199456;R-HSA-430190
R-HSA-5654679;R-HSA-5654647;R-HSA-5654709;R-HSA-5654643;R-HSA-5654641;R-HSA-5654623;R-HSA-1549564;R-H
R-HSA-182594;R-HSA-1632857;R-HSA-3209109;R-HSA-5205673;R-HSA-5672017;R-HSA-3209114;R-HSA-4647594;R-HS
R-HSA-5690790;R-HSA-5696547;R-HSA-6781953;R-HSA-5690785;R-HSA-5689649;R-HSA-5689950;R-HSA-5689630;R-H
R-HSA-182594;R-HSA-3209109;R-HSA-3209114;R-HSA-4647594;R-HSA-3240957;R-HSA-3229089;R-HSA-3223200;R-HS
R-HSA-5229132;R-HSA-432707
R-HSA-6791223;R-HSA-6791221;R-HSA-6791219
R-HSA-5654672;R-HSA-5654673;R-HSA-1549564;R-HSA-5654587;R-HSA-5654584;R-HSA-5654392;R-HSA-5654692;R-H
R-HSA-5685318
R-HSA-8956365;R-HSA-4615873;R-HSA-4615933
R-HSA-2187330;R-HSA-2187303;R-HSA-870538;R-HSA-2187325;R-HSA-870449;R-HSA-2187309
R-HSA-5685318
R-HSA-381799;R-HSA-437136;R-HSA-265177
R-HSA-6804741;R-HSA-6795460;R-HSA-198599;R-HSA-6793666;R-HSA-3222072;R-HSA-3215295;R-HSA-6795667;R-HS
R-HSA-450406;R-HSA-450551;R-HSA-482788;R-HSA-450620;R-HSA-450394
R-HSA-870269;R-HSA-426240;R-HSA-2179387;R-HSA-379048;R-HSA-2179407
R-HSA-6804741;R-HSA-141409;R-HSA-6803411;R-HSA-141431;R-HSA-141422;R-HSA-6804955;R-HSA-141439
R-HSA-427383;R-HSA-5250947;R-HSA-427528
R-HSA-3928633;R-HSA-3928639;R-HSA-4093329
R-HSA-5610726;R-HSA-5610732;R-HSA-5610730
R-HSA-994106;R-HSA-1013833;R-HSA-1008248;R-HSA-994038;R-HSA-996755;R-HSA-879459
R-HSA-9011241;R-HSA-428535;R-HSA-9014610;R-HSA-8985812
R-HSA-8868230;R-HSA-5229132;R-HSA-8868236;R-HSA-8877451;R-HSA-8847635;R-HSA-8876191;R-HSA-8854302;R-H
R-HSA-426240;R-HSA-392195;R-HSA-442273;R-HSA-428750;R-HSA-442314;R-HSA-437192;R-HSA-939265;R-HSA-428
R-HSA-6791223;R-HSA-6791221;R-HSA-6791219
R-HSA-210290;R-HSA-202722;R-HSA-210977;R-HSA-210304;R-HSA-8858435;R-HSA-210294;R-HSA-204549;R-HSA-210
R-HSA-5654677;R-HSA-5654608;R-HSA-5654622;R-HSA-5654399;R-HSA-5654620;R-HSA-1549564;R-HSA-5654618;R-H
R-HSA-182594;R-HSA-188386;R-HSA-8964525;R-HSA-1363314;R-HSA-8942607;R-HSA-9018017;R-HSA-69227;R-HSA-8
R-HSA-947535;R-HSA-196929;R-HSA-1474146
R-HSA-5634228;R-HSA-5626962;R-HSA-5634221
R-HSA-5654608;R-HSA-5654622;R-HSA-5654623;R-HSA-5654620;R-HSA-5654618;R-HSA-5654596;R-HSA-5654594;R-H
R-HSA-8939929;R-HSA-9007816;R-HSA-9008456;R-HSA-9008475;R-HSA-8939900;R-HSA-9008476;R-HSA-9007759;R-H
R-HSA-199558;R-HSA-198941
R-HSA-9606784;R-HSA-9645137;R-HSA-109699;R-HSA-9606624;R-HSA-6786789;R-HSA-5672965;R-HSA-9645126;R-HS
R-HSA-9626816;R-HSA-5666160;R-HSA-5666129;R-HSA-5625870;R-HSA-5625849;R-HSA-5672329;R-HSA-5666169;R-H
R-HSA-2730840;R-HSA-2730848;R-HSA-2424477
R-HSA-177923;R-HSA-177920;R-HSA-6785636;R-HSA-177926;R-HSA-177924;R-HSA-5654608;R-HSA-1562641;R-HSA-5
R-HSA-936462;R-HSA-877308;R-HSA-1678843;R-HSA-997309;R-HSA-1678842;R-HSA-909726;R-HSA-1169399;R-HSA-8
R-HSA-5621573
R-HSA-72160;R-HSA-8849157;R-HSA-72130;R-HSA-72231;R-HSA-72103;R-HSA-159050;R-HSA-75080;R-HSA-72107;R-H
R-HSA-6791223;R-HSA-6791221;R-HSA-6791219
R-HSA-8950634;R-HSA-8950637;R-HSA-8950415
R-HSA-70241;R-HSA-372819;R-HSA-177784;R-HSA-71306

R-HSA-8951498;R-HSA-199558;R-HSA-2130725;R-HSA-939265;R-HSA-9606151;R-HSA-388829;R-HSA-388828;R-HSA-2
R-HSA-983422;R-HSA-983147;R-HSA-983157;R-HSA-983156;R-HSA-983140
R-HSA-4649028;R-HSA-3299569;R-HSA-3322422;R-HSA-201669;R-HSA-3769393;R-HSA-4641231;R-HSA-3364026;R-HS
R-HSA-156930;R-HSA-5419264;R-HSA-72722;R-HSA-156913;R-HSA-72631;R-HSA-72663;R-HSA-5419268;R-HSA-7261
R-HSA-5229132;R-HSA-376369;R-HSA-350769;R-HSA-432707;R-HSA-434362;R-HSA-421835
R-HSA-200952;R-HSA-180606;R-HSA-180591;R-HSA-180687;R-HSA-200908
R-HSA-2187330;R-HSA-2187303;R-HSA-870538;R-HSA-2187325;R-HSA-870449;R-HSA-2187309
R-HSA-5357776;R-HSA-5634228;R-HSA-5626982;R-HSA-5357780;R-HSA-5626962;R-HSA-5634221
R-HSA-5685318
R-HSA-379044;R-HSA-8964287;R-HSA-8964278
R-HSA-5617431;R-HSA-5617862;R-HSA-5617859;R-HSA-5617452;R-HSA-5617896
R-HSA-5617431;R-HSA-5617862;R-HSA-5617859;R-HSA-5617452;R-HSA-5617896
R-HSA-8849463;R-HSA-8848596;R-HSA-8848611;R-HSA-8848993;R-HSA-8848975;R-HSA-8848606;R-HSA-8857565;R-H
R-HSA-8849463;R-HSA-8848596;R-HSA-8848611;R-HSA-8848993;R-HSA-8848975;R-HSA-8848606;R-HSA-8857565;R-H
R-HSA-380294;R-HSA-375302;R-HSA-8942607;R-HSA-380303;R-HSA-1638803;R-HSA-8852362;R-HSA-380311;R-HSA-1
R-HSA-3928651;R-HSA-3928633;R-HSA-3928632;R-HSA-3928639;R-HSA-3928654;R-HSA-4093336;R-HSA-3928642;R-H
R-HSA-2187330;R-HSA-2187303;R-HSA-870538;R-HSA-2187325;R-HSA-870449;R-HSA-2187309
R-HSA-5357776;R-HSA-5634228;R-HSA-193696;R-HSA-5357780;R-HSA-193668;R-HSA-5626962;R-HSA-193646;R-HSA-
R-HSA-981832;R-HSA-374248
R-HSA-265428;R-HSA-210304;R-HSA-198941
R-HSA-8937369;R-HSA-8874797;R-HSA-8864395;R-HSA-8864698;R-HSA-8865256
R-HSA-8868567;R-HSA-8870710;R-HSA-8863013;R-HSA-8868260;R-HSA-8863009
R-HSA-8868567;R-HSA-8870710;R-HSA-8863013;R-HSA-8868260;R-HSA-8863009
R-HSA-549241;R-HSA-428052;R-HSA-437136;R-HSA-433089
R-HSA-3215152;R-HSA-2318752;R-HSA-198599;R-HSA-5632732;R-HSA-3222072;R-HSA-3215295;R-HSA-198609;R-HSA
R-HSA-8865605;R-HSA-9645137;R-HSA-109699;R-HSA-6786789;R-HSA-8950415;R-HSA-8983425;R-HSA-879937;R-HSA
R-HSA-1912412;R-HSA-157632;R-HSA-157926;R-HSA-9021357;R-HSA-157353;R-HSA-9017797;R-HSA-9604703;R-HSA-
R-HSA-114307;R-HSA-9627350;R-HSA-139908;R-HSA-139909;R-HSA-139918;R-HSA-114252;R-HSA-9627106;R-HSA-96
R-HSA-3215152;R-HSA-8937097;R-HSA-2187303;R-HSA-9007759;R-HSA-3222072;R-HSA-9623192;R-HSA-8878220;R-H
R-HSA-114307;R-HSA-9627350;R-HSA-139908;R-HSA-139909;R-HSA-139918;R-HSA-114252;R-HSA-9627106;R-HSA-96
R-HSA-6788611;R-HSA-70600;R-HSA-6792572;R-HSA-71037;R-HSA-70589
R-HSA-9011985;R-HSA-9011984;R-HSA-9023840;R-HSA-9009378;R-HSA-9023861
R-HSA-3149440;R-HSA-3149432;R-HSA-3149433;R-HSA-200952;R-HSA-180606;R-HSA-180591;R-HSA-180687;R-HSA-3
R-HSA-1655833;R-HSA-2426148;R-HSA-1655832;R-HSA-2426151;R-HSA-2426150;R-HSA-1655837;R-HSA-2426147;R-H
R-HSA-981832;R-HSA-374248
R-HSA-3215152;R-HSA-8937097;R-HSA-2187303;R-HSA-9007759;R-HSA-3222072;R-HSA-9623192;R-HSA-8878220;R-H
R-HSA-8964329;R-HSA-180038;R-HSA-74885;R-HSA-420883;R-HSA-167415;R-HSA-167429;R-HSA-2485180;R-HSA-380
R-HSA-3215152;R-HSA-8937097;R-HSA-2187303;R-HSA-9007759;R-HSA-3222072;R-HSA-9623192;R-HSA-8878220;R-H
R-HSA-6804741;R-HSA-6795460;R-HSA-198599;R-HSA-6793666;R-HSA-3222072;R-HSA-3215295;R-HSA-6795667;R-HS
R-HSA-210304;R-HSA-8858435;R-HSA-426240;R-HSA-1013833;R-HSA-204549;R-HSA-392195;R-HSA-428750;R-HSA-43
R-HSA-5685318;R-HSA-6785126;R-HSA-110322
R-HSA-380294;R-HSA-375302;R-HSA-8942607;R-HSA-164616;R-HSA-380303;R-HSA-1638803;R-HSA-8852362;R-HSA-3
R-HSA-8855237;R-HSA-3928633;R-HSA-376134;R-HSA-3928632;R-HSA-374662;R-HSA-3928639;R-HSA-399938;R-HSA-
R-HSA-9632182;R-HSA-9021596;R-HSA-9011985;R-HSA-9623355;R-HSA-9011984;R-HSA-9023840;R-HSA-9009378;R-H
R-HSA-936462;R-HSA-8865605;R-HSA-9645137;R-HSA-109699;R-HSA-6786789;R-HSA-8950415;R-HSA-8983425;R-HSA
R-HSA-1655833;R-HSA-2426148;R-HSA-1655832;R-HSA-2426151;R-HSA-2426150;R-HSA-1655837;R-HSA-2426147;R-H
R-HSA-9632182;R-HSA-9021596;R-HSA-9011985;R-HSA-9623355;R-HSA-9011984;R-HSA-9023840;R-HSA-9009378;R-H
R-HSA-8956365;R-HSA-4615873;R-HSA-5689950;R-HSA-5689630;R-HSA-6809006;R-HSA-203977;R-HSA-446221;R-HSA
R-HSA-5250962;R-HSA-168896;R-HSA-3149432;R-HSA-3149433;R-HSA-192841;R-HSA-112333;R-HSA-8875371;R-HSA-
R-HSA-5228523;R-HSA-8956365;R-HSA-2997706;R-HSA-4615873;R-HSA-4655404;R-HSA-4551604;R-HSA-3903017;R-H

R-HSA-5228523;R-HSA-8956365;R-HSA-2997706;R-HSA-4615873;R-HSA-4655404;R-HSA-4551604;R-HSA-3903017;R-H
R-HSA-180038;R-HSA-74885;R-HSA-426240;R-HSA-749453;R-HSA-202692;R-HSA-167429;R-HSA-180047;R-HSA-89646
R-HSA-936462;R-HSA-2197697;R-HSA-2029467;R-HSA-168935;R-HSA-2730848;R-HSA-874087;R-HSA-3134883;R-HSA-
R-HSA-8849157;R-HSA-72130;R-HSA-429955;R-HSA-6782434;R-HSA-159050;R-HSA-75080;R-HSA-72139;R-HSA-72143
R-HSA-936462;R-HSA-109699;R-HSA-6786789;R-HSA-2029467;R-HSA-199558;R-HSA-2130725;R-HSA-8950415;R-HSA-
R-HSA-5690886;R-HSA-549241;R-HSA-428052;R-HSA-432195;R-HSA-437136;R-HSA-433089
R-HSA-3928633;R-HSA-3928632;R-HSA-374662;R-HSA-3928639;R-HSA-1183003;R-HSA-8855564;R-HSA-445069;R-HSA
R-HSA-156930;R-HSA-4615873;R-HSA-5689950;R-HSA-1791121;R-HSA-156808;R-HSA-446221;R-HSA-8931974;R-HSA-
R-HSA-180038;R-HSA-74885;R-HSA-981832;R-HSA-426240;R-HSA-749453;R-HSA-202692;R-HSA-167429;R-HSA-18004
R-HSA-6802927;R-HSA-5655262;R-HSA-4839746;R-HSA-2666278;R-HSA-6802935;R-HSA-6802934;R-HSA-6802933;R-H
R-HSA-4649028;R-HSA-6785636;R-HSA-3322422;R-HSA-2187303;R-HSA-5654608;R-HSA-1562641;R-HSA-5626962;R-H
R-HSA-1655833;R-HSA-2426148;R-HSA-1655832;R-HSA-2426151;R-HSA-2426150;R-HSA-1655837;R-HSA-2426147;R-H

-ISA-6794344;R-HSA-8849878;R-HSA-6794357;R-HSA-6794356;R-HSA-6798258;R-HSA-6797810;R-HSA-679755[illegible]
-ISA-6794344;R-HSA-6797974;R-HSA-6794357;R-HSA-6794356;R-HSA-6797554;R-HSA-6794354;R-HSA-679435[illegible]

A-6794349;R-HSA-6794348;R-HSA-6797803;R-HSA-9610270;R-HSA-6794346;R-HSA-442760;R-HSA-6794345;R-

SA-1676114;R-HSA-1676149;R-HSA-1676020;R-HSA-1676168;R-HSA-1676174;R-HSA-1675810;R-HSA-1676005;

SA-8851011;R-HSA-8944397;R-HSA-8850945;R-HSA-8944078;R-HSA-8948800;R-HSA-8944706;R-HSA-8943811;

HSA-1605428;R-HSA-1592245;R-HSA-1592246;R-HSA-1592247;R-HSA-1592232;R-HSA-2466391;R-HSA-159223

HSA-8866283;R-HSA-8867754;R-HSA-8868072;R-HSA-8868648;R-HSA-8868661;R-HSA-8868660;R-HSA-886865

HSA-9610653;R-HSA-9609738;R-HSA-9609739;R-HSA-9611368;R-HSA-9609737;R-HSA-9609742;R-HSA-960974

;A-8851011;R-HSA-8944397;R-HSA-8850945;R-HSA-8944078;R-HSA-8948800;R-HSA-8944706;R-HSA-8943811;

-ISA-8944397;R-HSA-8850945;R-HSA-8944078;R-HSA-8948800;R-HSA-8944706;R-HSA-8943811;R-HSA-894856

HSA-8877612;R-HSA-8877451;R-HSA-8876616;R-HSA-8875318;R-HSA-8877813;R-HSA-8876191;R-HSA-887731[illegible]

HSA-1655839;R-HSA-2426146;R-HSA-2426157;R-HSA-2426156;R-HSA-1655824;R-HSA-1655827;R-HSA-242615[illegible]

HSA-8937682;R-HSA-8936979;R-HSA-8934742;R-HSA-8936616;R-HSA-8935785;R-HSA-8938217;R-HSA-893662[illegible]

HSA-1605428;R-HSA-1592244;R-HSA-1592245;R-HSA-1592246;R-HSA-1592247;R-HSA-1592232;R-HSA-159223[illegible]

HSA-1655839;R-HSA-2426146;R-HSA-2426157;R-HSA-2426156;R-HSA-1655824;R-HSA-1655827;R-HSA-242615[illegible]

HSA-8877612;R-HSA-8877451;R-HSA-8876616;R-HSA-8875318;R-HSA-8877813;R-HSA-8876191;R-HSA-887731

SA-1676082;R-HSA-1676114;R-HSA-1676149;R-HSA-1676020;R-HSA-1676168;R-HSA-1676174;R-HSA-1675810

HSA-5625421;R-HSA-2466367;R-HSA-8940309;R-HSA-5625416;R-HSA-5626699;R-HSA-5624949;R-HSA-562622

.77924;R-HSA-182986;R-HSA-182990;R-HSA-177935;R-HSA-177938;R-HSA-177942;R-HSA-177944;R-HSA-1779

HSA-5617820;R-HSA-5624953;R-HSA-5626681;R-HSA-5617816;R-HSA-5624952;R-HSA-5638009;R-HSA-562352

A-9610327;R-HSA-416320;R-HSA-421007;R-HSA-9619803;R-HSA-9610270;R-HSA-442760;R-HSA-374922;R-HSA

SA-421007;R-HSA-9619803;R-HSA-9610270;R-HSA-442760;R-HSA-9610653;R-HSA-438037;R-HSA-451283;R-HS

HSA-8937050;R-HSA-8935740;R-HSA-8936979;R-HSA-8935731;R-HSA-8935732;R-HSA-8936628;R-HSA-89347

A-8852362;R-HSA-3000319;R-HSA-380311;R-HSA-8853405;R-HSA-2574845;R-HSA-3000310;R-HSA-4088162;R-

A-8852362;R-HSA-3000319;R-HSA-380311;R-HSA-8853405;R-HSA-2574845;R-HSA-3000310;R-HSA-4088162;R-

HSA-9610270;R-HSA-442760;R-HSA-9610653;R-HSA-438037;R-HSA-9609738;R-HSA-9609739;R-HSA-9609737;R

HSA-2666278;R-HSA-6802935;R-HSA-6802934;R-HSA-6802933;R-HSA-6802932;R-HSA-2220978;R-HSA-222098

SA-6795473;R-HSA-3700992;R-HSA-6792863;R-HSA-6793661;R-HSA-6804955;R-HSA-6795290;R-HSA-3215310

SA-3322431;R-HSA-4411351;R-HSA-5368580;R-HSA-201685;R-HSA-5323526;R-HSA-3364014;R-HSA-5665608;R

-9033292;R-HSA-202692;R-HSA-9603419;R-HSA-9032067;R-HSA-9032582;R-HSA-171026;R-HSA-169904;R-HSA

HSA-8876190;R-HSA-8876188;R-HSA-8871194;R-HSA-8871193;R-HSA-5694409;R-HSA-8876454;R-HSA-887683

HSA-8878220;R-HSA-8952419;R-HSA-8951428;R-HSA-8878186;R-HSA-8952408;R-HSA-8952058;R-HSA-895167

SA-3240957;R-HSA-3229089;R-HSA-3223200;R-HSA-1632843;R-HSA-5696197;R-HSA-5618080;R-HSA-3209096;

SA-5696197;R-HSA-5618080;R-HSA-3209096;R-HSA-3371435;R-HSA-4647613;R-HSA-3209165;R-HSA-3857348;

SA-6795473;R-HSA-3700992;R-HSA-6792863;R-HSA-6793661;R-HSA-6804188;R-HSA-6804955;R-HSA-6795290;

HSA-8876190;R-HSA-8876188;R-HSA-8871194;R-HSA-8871193;R-HSA-5694409;R-HSA-8876454;R-HSA-887683

17;R-HSA-418579;R-HSA-442291;R-HSA-453200;R-HSA-5684801;R-HSA-354173;R-HSA-114552;R-HSA-568416

941895;R-HSA-69299;R-HSA-8941915;R-HSA-1363328;R-HSA-187828;R-HSA-1363276;R-HSA-8942803;R-HSA-

HSA-5654734;R-HSA-5654607;R-HSA-5654730;R-HSA-5654603;R-HSA-5654600;R-HSA-5654729;R-HSA-565464

654622;R-HSA-5654623;R-HSA-9032067;R-HSA-5654620;R-HSA-205321;R-HSA-5654618;R-HSA-177935;R-HSA

HSA-72139;R-HSA-158447;R-HSA-72143;R-HSA-111439;R-HSA-77587;R-HSA-72180;R-HSA-158484;R-HSA-1566

2130619;R-HSA-198941;R-HSA-2213248;R-HSA-389348;R-HSA-199456;R-HSA-389350;R-HSA-983147;R-HSA-43

SA-3322427;R-HSA-3322424;R-HSA-3322431;R-HSA-4411351;R-HSA-8944349;R-HSA-5368580;R-HSA-201685;

363328;R-HSA-380316;R-HSA-8853405;R-HSA-2468287;R-HSA-187545;R-HSA-380455;R-HSA-8964525;R-HSA-

-205039;R-HSA-9012761;R-HSA-5626982;R-HSA-205008;R-HSA-193684;R-HSA-5634221;R-HSA-193694

A-2321904;R-HSA-6793661;R-HSA-6804741;R-HSA-6795460;R-HSA-6791235;R-HSA-6793666;R-HSA-6803411;
A-8987014;R-HSA-9606784;R-HSA-5672965;R-HSA-1112708;R-HSA-9645126;R-HSA-8854736;R-HSA-9605259;
-1912396;R-HSA-9013693;R-HSA-2197588;R-HSA-9013631;R-HSA-1912388;R-HSA-9021520;R-HSA-1912391;R-

HSA-9624985;R-HSA-8864395;R-HSA-8951428;R-HSA-2187309;R-HSA-8938887;R-HSA-8939929;R-HSA-893736

HSA-1655839;R-HSA-2426146;R-HSA-2426157;R-HSA-2426156;R-HSA-1655824;R-HSA-1655827;R-HSA-242615

HSA-9624985;R-HSA-8864395;R-HSA-8951428;R-HSA-2187309;R-HSA-8938887;R-HSA-8939929;R-HSA-893736

HSA-9624985;R-HSA-8864395;R-HSA-8951428;R-HSA-2187309;R-HSA-427528;R-HSA-8938887;R-HSA-8939929;
SA-6795473;R-HSA-3700992;R-HSA-6792863;R-HSA-6793661;R-HSA-8877692;R-HSA-6804188;R-HSA-6804955;
2143;R-HSA-204873;R-HSA-442314;R-HSA-202702;R-HSA-437192;R-HSA-939265;R-HSA-418579;R-HSA-45320

80311;R-HSA-1363328;R-HSA-380316;R-HSA-8853405;R-HSA-2468287;R-HSA-187545;R-HSA-380455;R-HSA-8
-399951;R-HSA-416588;R-HSA-8855564;R-HSA-445069;R-HSA-445064;R-HSA-3928628;R-HSA-373716;R-HSA-8

A-8985157;R-HSA-879937;R-HSA-1369114;R-HSA-1369115;R-HSA-8987014;R-HSA-9606784;R-HSA-5672965;R-
HSA-1655839;R-HSA-2426146;R-HSA-2426157;R-HSA-1675921;R-HSA-2426156;R-HSA-1655824;R-HSA-165582

A-6809003;R-HSA-8931974;R-HSA-3903017;R-HSA-4717461;R-HSA-4090408;R-HSA-4085992;R-HSA-4090281;
-5228578;R-HSA-180591;R-HSA-180687;R-HSA-194796;R-HSA-200908;R-HSA-3149440;R-HSA-8876948;R-HSA-
HSA-4717461;R-HSA-4090408;R-HSA-4085992;R-HSA-4090281;R-HSA-4719423;R-HSA-4615933;R-HSA-455176

HSA-4717461;R-HSA-4090408;R-HSA-4085992;R-HSA-4090281;R-HSA-4719423;R-HSA-4615933;R-HSA-455176
34;R-HSA-420883;R-HSA-419166;R-HSA-2485180;R-HSA-2179387;R-HSA-8964311;R-HSA-8964329;R-HSA-379
-2201293;R-HSA-6800426;R-HSA-3134914;R-HSA-5621573;R-HSA-2730840;R-HSA-936380;R-HSA-2029451;R-H
;R-HSA-111439;R-HSA-6791223;R-HSA-77587;R-HSA-6791221;R-HSA-6791219;R-HSA-77589;R-HSA-77591;R-H
-8983425;R-HSA-3134883;R-HSA-1369114;R-HSA-1369115;R-HSA-8987014;R-HSA-939265;R-HSA-5621573;R-H

A-560517;R-HSA-445064;R-HSA-3928628;R-HSA-211346;R-HSA-5617862;R-HSA-5617859;R-HSA-210836;R-HSA
-3903017;R-HSA-72722;R-HSA-4090408;R-HSA-437136;R-HSA-4090281;R-HSA-1791107;R-HSA-5692495;R-HSA
7;R-HSA-8964634;R-HSA-420883;R-HSA-419166;R-HSA-2485180;R-HSA-2179387;R-HSA-8964311;R-HSA-8964
HSA-6802932;R-HSA-2220978;R-HSA-8876948;R-HSA-5250892;R-HSA-2220988;R-HSA-5655241;R-HSA-886301
HSA-167429;R-HSA-3769393;R-HSA-5654622;R-HSA-3322427;R-HSA-5654623;R-HSA-3322424;R-HSA-5654620;
HSA-1655839;R-HSA-2426146;R-HSA-2426157;R-HSA-1675921;R-HSA-2426156;R-HSA-1655824;R-HSA-165582

4;R-HSA-6794354;R-HSA-6794353;R-HSA-6794352;R-HSA-6800380;R-HSA-388824;R-HSA-8849882;R-HSA-8849

HSA-374922;R-HSA-6794344;R-HSA-9610653;R-HSA-210444;R-HSA-6794357;R-HSA-438037;R-HSA-6794356;R-

;R-HSA-8948569;R-HSA-198609;R-HSA-8944026;R-HSA-8944538;R-HSA-8850961;R-HSA-8944599;R-HSA-89441

4;R-HSA-1592236;R-HSA-1592238;R-HSA-1368140;R-HSA-2466367;R-HSA-2466392;R-HSA-8940309;R-HSA-159

;R-HSA-8948569;R-HSA-198609;R-HSA-8944026;R-HSA-8944538;R-HSA-8850961;R-HSA-8944599;R-HSA-89441

9;R-HSA-8944026;R-HSA-8944538;R-HSA-8850961;R-HSA-8944599;R-HSA-8944104;R-HSA-8944684;R-HSA-680

3;R-HSA-1655826;R-HSA-2426153;R-HSA-2426152;R-HSA-2426155;R-HSA-2426164;R-HSA-2426161;R-HSA-242

1;R-HSA-8936481;R-HSA-8936995;R-HSA-8938853;R-HSA-8937767;R-HSA-8865320;R-HSA-8937016;R-HSA-893

3;R-HSA-2466391;R-HSA-1592234;R-HSA-1592236;R-HSA-1592238;R-HSA-1368140;R-HSA-2466367;R-HSA-246

3;R-HSA-1655826;R-HSA-2426153;R-HSA-2426152;R-HSA-2426155;R-HSA-2426164;R-HSA-2426161;R-HSA-242

3;R-HSA-1592249;R-HSA-1592250;R-HSA-1592252;R-HSA-2466369;R-HSA-5626227;R-HSA-2466370;R-HSA-562

-9610653;R-HSA-210444;R-HSA-438037;R-HSA-451283;R-HSA-9609738;R-HSA-9609739;R-HSA-380574;R-HSA-

A-9609738;R-HSA-9609739;R-HSA-9609737;R-HSA-9609742;R-HSA-9609743;R-HSA-9609740;R-HSA-416985;R-

HSA-2574840;R-HSA-380455;R-HSA-4088024;R-HSA-8853419;R-HSA-8853429;R-HSA-4088141;R-HSA-380272;

HSA-2574840;R-HSA-380455;R-HSA-4088024;R-HSA-8853419;R-HSA-8853429;R-HSA-4088141;R-HSA-380272;

-HSA-9609742;R-HSA-9609743;R-HSA-9609740;R-HSA-432164;R-HSA-432162;R-HSA-9612085;R-HSA-9619515

;R-HSA-5655241;R-HSA-5655351;R-HSA-5655347;R-HSA-5637808;R-HSA-4791278;R-HSA-2317387;R-HSA-273

:-HSA-4411357;R-HSA-3769383;R-HSA-3361751;R-HSA-4411367;R-HSA-3451153;R-HSA-3772434;R-HSA-37724

7;R-HSA-8877475;R-HSA-8876193;R-HSA-8877998;R-HSA-8867756;R-HSA-8877612;R-HSA-8868651;R-HSA-886

6;R-HSA-8878130;R-HSA-8878193;R-HSA-8952382;R-HSA-8878143;R-HSA-8951442;R-HSA-8951443;R-HSA-895

;R-HSA-3371435;R-HSA-4647613;R-HSA-5678315;R-HSA-3209165;R-HSA-3857348;R-HSA-5678313;R-HSA-3229

7;R-HSA-8877475;R-HSA-8876193;R-HSA-8877998;R-HSA-8867756;R-HSA-8877612;R-HSA-8868651;R-HSA-886

187545;R-HSA-8942836;R-HSA-1226094;R-HSA-1226095;R-HSA-1363306;R-HSA-113503;R-HSA-1363274

7;R-HSA-5654643;R-HSA-8941613;R-HSA-5654641;R-HSA-5654655;R-HSA-5654399;R-HSA-5654651;R-HSA-565

-9032582;R-HSA-171026;R-HSA-177938;R-HSA-5654596;R-HSA-5654594;R-HSA-177942;R-HSA-5654607;R-HS

0190;R-HSA-983140;R-HSA-1168636;R-HSA-389354;R-HSA-389941;R-HSA-2130641;R-HSA-983422;R-HSA-3541

-HSA-5323526;R-HSA-3364014;R-HSA-5665608;R-HSA-4411357;R-HSA-3769383;R-HSA-8944362;R-HSA-33617

9018017;R-HSA-8853419;R-HSA-75820;R-HSA-75823;R-HSA-75824;R-HSA-8853429;R-HSA-69299;R-HSA-18782

-HSA-6795667;R-HSA-5632993;R-HSA-380979;R-HSA-6795473;R-HSA-3700992;R-HSA-6792863;R-HSA-165179
-HSA-9607304;R-HSA-450074;R-HSA-9645134;R-HSA-8950634;R-HSA-1112690;R-HSA-508451;R-HSA-8950637

9;R-HSA-9623168;R-HSA-77592;R-HSA-8951442;R-HSA-8951443;R-HSA-2187325;R-HSA-8936616;R-HSA-90065

8;R-HSA-1655826;R-HSA-2426153;R-HSA-2426152;R-HSA-2426155;R-HSA-2426164;R-HSA-2426161;R-HSA-242

9;R-HSA-9623168;R-HSA-8951442;R-HSA-8951443;R-HSA-2187325;R-HSA-8936616;R-HSA-9006507;R-HSA-679

;R-HSA-8937369;R-HSA-9623168;R-HSA-77592;R-HSA-8951442;R-HSA-8951443;R-HSA-2187325;R-HSA-893661

0;R-HSA-5684801;R-HSA-996755;R-HSA-392152;R-HSA-5684169;R-HSA-432295;R-HSA-202722;R-HSA-210977;

964525;R-HSA-9018017;R-HSA-8853419;R-HSA-75820;R-HSA-75823;R-HSA-75824;R-HSA-8853429;R-HSA-692
855508;R-HSA-9014610;R-HSA-8985812;R-HSA-3928612;R-HSA-9011241;R-HSA-418850;R-HSA-4093336;R-HS

HSA-909726;R-HSA-1112708;R-HSA-9645126;R-HSA-8854736;R-HSA-9605259;R-HSA-9607304;R-HSA-450074;
7;R-HSA-1655826;R-HSA-2426158;R-HSA-2426153;R-HSA-560517;R-HSA-2426152;R-HSA-2426155;R-HSA-2426

-HSA-4551768;R-HSA-8873946;R-HSA-3000434;R-HSA-4719447;R-HSA-5694409;R-HSA-5228523;R-HSA-67819

044;R-HSA-380073;R-HSA-379048;R-HSA-2179407;R-HSA-870269;R-HSA-114544;R-HSA-114548;R-HSA-167415
SA-3134904;R-HSA-3244614;R-HSA-2424476;R-HSA-2424477;R-HSA-3134901;R-HSA-2029445;R-HSA-6798747
ISA-158484;R-HSA-77590;R-HSA-77593;R-HSA-927830;R-HSA-77081;R-HSA-75096;R-HSA-77592;R-HSA-77594;
SA-9606784;R-HSA-909726;R-HSA-1112708;R-HSA-9606151;R-HSA-2029451;R-HSA-3134904;R-HSA-9605259;

-3928612;R-HSA-9011241;R-HSA-418850;R-HSA-4093336;R-HSA-418863;R-HSA-3928592;R-HSA-418859;R-HSA
-4551768;R-HSA-4719447;R-HSA-5694409;R-HSA-9010091;R-HSA-5228523;R-HSA-72619;R-HSA-6787533;R-HS
329;R-HSA-379044;R-HSA-380073;R-HSA-374248;R-HSA-379048;R-HSA-2179407;R-HSA-870269;R-HSA-114544
3;R-HSA-8868260;R-HSA-8863009;R-HSA-4791278;R-HSA-2730752;R-HSA-2400009;R-HSA-3149440;R-HSA-887
;R-HSA-205321;R-HSA-2187309;R-HSA-5654618;R-HSA-3322431;R-HSA-171026;R-HSA-5654596;R-HSA-565459
7;R-HSA-1655826;R-HSA-2426158;R-HSA-71306;R-HSA-2426153;R-HSA-560517;R-HSA-2426152;R-HSA-242615

-HSA-6797810;R-HSA-6797554;R-HSA-6794354;R-HSA-6794353;R-HSA-6794352;R-HSA-9609738;R-HSA-96097[illegible]

.04;R-HSA-199456;R-HSA-8944684;R-HSA-6807106;R-HSA-8945709;R-HSA-6807105;R-HSA-8847968;R-HSA-89

.04;R-HSA-199456;R-HSA-8944684;R-HSA-6807106;R-HSA-8945709;R-HSA-6807105;R-HSA-8847968;R-HSA-89

)7106;R-HSA-8945709;R-HSA-6807105;R-HSA-8847968;R-HSA-8944047;R-HSA-8948832;R-HSA-6807118;R-HSA

:6163;R-HSA-1655823;R-HSA-2426162;R-HSA-2065966;R-HSA-1655848;R-HSA-1655850;R-HSA-1655852;R-HSA

:6163;R-HSA-1655823;R-HSA-2426162;R-HSA-2065966;R-HSA-1655848;R-HSA-1655850;R-HSA-1655852;R-HSA

:4125;R-HSA-1592242;R-HSA-1605428;R-HSA-1592244;R-HSA-5624953;R-HSA-5626681;R-HSA-5624952;R-HSA

-9609737;R-HSA-372505;R-HSA-9609742;R-HSA-9609743;R-HSA-9609740;R-HSA-416985;R-HSA-380901;R-HSA

-HSA-432164;R-HSA-432162;R-HSA-9612085;R-HSA-9619515;R-HSA-451311;R-HSA-442732;R-HSA-432172;R-H [illegible];R-HSA-442732;R-HSA-432172;R-HSA-9618750;R-HSA-9610879;R-HSA-445367;R-HSA-9611368;R-HSA-961418[illegible]

[illegible]0752;R-HSA-1247999;R-HSA-2400009;R-HSA-5655269;R-HSA-5334052;R-HSA-1248677;R-HSA-1225957;R-HSA

436;R-HSA-4411372;R-HSA-5626938;R-HSA-3322434;R-HSA-201712;R-HSA-3772430;R-HSA-3364042;R-HSA-345

57754;R-HSA-8868648;R-HSA-8875318;R-HSA-8868661;R-HSA-8868660;R-HSA-196017;R-HSA-376369;R-HSA-88

[illegible]7754;R-HSA-8868648;R-HSA-8875318;R-HSA-8868661;R-HSA-8868660;R-HSA-196017;R-HSA-376369;R-HSA-8[illegible]

[illegible]4392;R-HSA-5654631;R-HSA-8941628;R-HSA-8941618;R-HSA-5654633;R-HSA-8941623;R-HSA-5654422;R-HSA

[illegible]A-177944;R-HSA-177945;R-HSA-5654603;R-HSA-5654600;R-HSA-5654647;R-HSA-183072;R-HSA-8851877;R-HS

'51;R-HSA-4411367;R-HSA-3451153;R-HSA-4411365;R-HSA-8944352;R-HSA-195304;R-HSA-5229348;R-HSA-37

!8;R-HSA-1638821;R-HSA-182594;R-HSA-8853444;R-HSA-2467809;R-HSA-2467811;R-HSA-4088439;R-HSA-408

);R-HSA-380927;R-HSA-8877692;R-HSA-6804188;R-HSA-6804955;R-HSA-3222093;R-HSA-6795290;R-HSA-6804
';R-HSA-508513;R-HSA-909738;R-HSA-914022;R-HSA-8983394;R-HSA-8987236;R-HSA-1112703;R-HSA-960662

i07;R-HSA-6791235;R-HSA-8936621;R-HSA-8865454;R-HSA-6814549;R-HSA-380979;R-HSA-3700992;R-HSA-68

!6163;R-HSA-1655823;R-HSA-2426162;R-HSA-195690;R-HSA-6807052;R-HSA-194674;R-HSA-2065966;R-HSA-1

)1235;R-HSA-8936621;R-HSA-8865454;R-HSA-380979;R-HSA-3700992;R-HSA-165179;R-HSA-9007283;R-HSA-8

.6;R-HSA-9006507;R-HSA-6791235;R-HSA-8936621;R-HSA-8865454;R-HSA-6814549;R-HSA-380979;R-HSA-370

R-HSA-392870;R-HSA-210277;R-HSA-442273;R-HSA-210283;R-HSA-435244;R-HSA-879459;R-HSA-210285;R-HS

)9;R-HSA-6803411;R-HSA-187828;R-HSA-1638821;R-HSA-6804955;R-HSA-182594;R-HSA-8853444;R-HSA-2467
\-418863;R-HSA-3928592;R-HSA-418859;R-HSA-418856;R-HSA-4093329;R-HSA-373738;R-HSA-3928651;R-HSA

R-HSA-9645134;R-HSA-8950634;R-HSA-1678843;R-HSA-1112690;R-HSA-1678842;R-HSA-508451;R-HSA-89506
i164;R-HSA-1676168;R-HSA-2426161;R-HSA-2426163;R-HSA-1655823;R-HSA-2426162;R-HSA-1676174;R-HSA-

)53;R-HSA-2997706;R-HSA-5689649;R-HSA-8952289;R-HSA-8865774;R-HSA-6787533;R-HSA-4655404;R-HSA-5

R-HSA-450394;R-HSA-450406;R-HSA-72160;R-HSA-482788;R-HSA-72231;R-HSA-72103;R-HSA-72107;R-HSA-15
R-HSA-3134901;R-HSA-9607304;R-HSA-2130619;R-HSA-2029445;R-HSA-450074;R-HSA-198941;R-HSA-221324

A-418856;R-HSA-4093329;R-HSA-555065;R-HSA-5682794;R-HSA-5617896;R-HSA-391867;R-HSA-1183058;R-HS
SA-72622;R-HSA-1791150;R-HSA-72631;R-HSA-9010096;R-HSA-4615933;R-HSA-1791143;R-HSA-3782535;R-HS

70710;R-HSA-5655269;R-HSA-5336420;R-HSA-1248677;R-HSA-5660015;R-HSA-8853310;R-HSA-5660013;R-HSA
4;R-HSA-9021596;R-HSA-420883;R-HSA-5654607;R-HSA-2187325;R-HSA-5654603;R-HSA-3769383;R-HSA-565
5;R-HSA-381706;R-HSA-2426164;R-HSA-1676168;R-HSA-825631;R-HSA-2426161;R-HSA-416530;R-HSA-24261

39;R-HSA-380574;R-HSA-6800380;R-HSA-9609737;R-HSA-372505;R-HSA-9609742;R-HSA-9609743;R-HSA-9609

44047;R-HSA-8948832;R-HSA-6807118;R-HSA-1676082;R-HSA-8944099;R-HSA-8944483;R-HSA-8943780;R-HSA

44047;R-HSA-8948832;R-HSA-6807118;R-HSA-1676082;R-HSA-8944099;R-HSA-8944483;R-HSA-8943780;R-HS

-8944099;R-HSA-8944483;R-HSA-8943780;R-HSA-8944420;R-HSA-8847977;R-HSA-8948775;R-HSA-8944632;R

-1592245;R-HSA-1592246;R-HSA-1592247;R-HSA-5638009;R-HSA-1592232;R-HSA-1592233;R-HSA-2466391;R-

-432164;R-HSA-432162;R-HSA-9612085;R-HSA-9619515;R-HSA-451311;R-HSA-442732;R-HSA-432172;R-HSA-3

SA-9618750;R-HSA-9610879;R-HSA-445367;R-HSA-416639;R-HSA-9611368;R-HSA-9614185;R-HSA-9617326

368659;R-HSA-350769;R-HSA-8868658;R-HSA-190519;R-HSA-8869438;R-HSA-434362;R-HSA-5694441;R-HSA-8

868659;R-HSA-350769;R-HSA-8868658;R-HSA-190519;R-HSA-8869438;R-HSA-434362;R-HSA-5694441;R-HSA-8

-5654679;R-HSA-5654677;R-HSA-5654672;R-HSA-5654673;R-HSA-5654684;R-HSA-5654426;R-HSA-5654663;R-

A-9036970;R-HSA-5654643;R-HSA-5654641;R-HSA-9612219;R-HSA-5654655;R-HSA-5654399;R-HSA-5654651;

72434;R-HSA-3772436;R-HSA-4411372;R-HSA-5626938;R-HSA-3322434;R-HSA-195251;R-HSA-5334050;R-HSA-

8309;R-HSA-4088306;R-HSA-4088307;R-HSA-8941895;R-HSA-3000319;R-HSA-8941915;R-HSA-1363276;R-HSA-

14559;R-HSA-165179;R-HSA-9007283;R-HSA-870449;R-HSA-6814555;R-HSA-8936628;R-HSA-6814554;R-HSA-8

655848;R-HSA-1655850;R-HSA-1655852;R-HSA-1655843;R-HSA-1655844;R-HSA-1655847;R-HSA-1655846

70449;R-HSA-8936628;R-HSA-8864698;R-HSA-9622738;R-HSA-9615570;R-HSA-9007816;R-HSA-9615571;R-HSA

0992;R-HSA-6814559;R-HSA-165179;R-HSA-9007283;R-HSA-870449;R-HSA-6814555;R-HSA-8936628;R-HSA-68

A-210290;R-HSA-428917;R-HSA-994106;R-HSA-210294;R-HSA-442291;R-HSA-1008248;R-HSA-354173;R-HSA-9

809;R-HSA-2467811;R-HSA-4088439;R-HSA-4088309;R-HSA-4088306;R-HSA-4088307;R-HSA-8941895;R-HSA-3
-555065;R-HSA-428535;R-HSA-392054;R-HSA-3928654;R-HSA-3928642;R-HSA-391867;R-HSA-3928644

37;R-HSA-1169399;R-HSA-508513;R-HSA-909738;R-HSA-914022;R-HSA-8983394;R-HSA-8987236;R-HSA-11127
1675910;R-HSA-199456;R-HSA-1676082;R-HSA-1655848;R-HSA-2065966;R-HSA-1655850;R-HSA-1655852;R-HS

690790;R-HSA-4551604;R-HSA-5696547;R-HSA-4719423;R-HSA-5690785;R-HSA-4615933;R-HSA-3782535;R-HS

8447;R-HSA-429992;R-HSA-450551;R-HSA-72180;R-HSA-156661;R-HSA-72185;R-HSA-450620;R-HSA-72124;R-I
3;R-HSA-199456;R-HSA-508451;R-HSA-909738;R-HSA-389941;R-HSA-936380;R-HSA-997309;R-HSA-8987070;R-

A-5617431;R-HSA-8855237;R-HSA-376134;R-HSA-9616214;R-HSA-448962;R-HSA-399938;R-HSA-448963;R-HSA
A-5694441;R-HSA-8871494;R-HSA-8956365;R-HSA-389954;R-HSA-5689630;R-HSA-6809006;R-HSA-203977;R-I

-5655277;R-HSA-8853307;R-HSA-3149454;R-HSA-5250962;R-HSA-168896;R-HSA-3149432;R-HSA-5655326;R-I
4600;R-HSA-5654647;R-HSA-9036970;R-HSA-6807106;R-HSA-5654643;R-HSA-6807105;R-HSA-5654641;R-HSA-
63;R-HSA-1655823;R-HSA-2426162;R-HSA-1676174;R-HSA-381727;R-HSA-6792572;R-HSA-1675910;R-HSA-199

740;R-HSA-432164;R-HSA-432162;R-HSA-9612085;R-HSA-9619515;R-HSA-432172;R-HSA-9618750;R-HSA-445

\-8944420;R-HSA-8847977;R-HSA-8948775;R-HSA-8944632;R-HSA-6807126;R-HSA-2316434;R-HSA-8944569;R

\-8944420;R-HSA-8847977;R-HSA-8948775;R-HSA-8944632;R-HSA-6807126;R-HSA-2316434;R-HSA-8944569;R

-HSA-6807126;R-HSA-8944569;R-HSA-8850997;R-HSA-8850992;R-HSA-8943728;R-HSA-8944497;R-HSA-68071

-HSA-1592234;R-HSA-1592236;R-HSA-5623521;R-HSA-1592238;R-HSA-5626220;R-HSA-5626223;R-HSA-24663[illegible]

380905;R-HSA-9618750;R-HSA-9610879;R-HSA-445367;R-HSA-372529;R-HSA-374899;R-HSA-210426;R-HSA-41[illegible]

847544;R-HSA-8875320;R-HSA-8876615;R-HSA-8849350;R-HSA-6811429;R-HSA-432707;R-HSA-8877760;R-HS

847544;R-HSA-8875320;R-HSA-8876615;R-HSA-8849350;R-HSA-6811429;R-HSA-432707;R-HSA-8877760;R-HS

-HSA-5654658;R-HSA-5654402;R-HSA-5654669;R-HSA-5654413;R-HSA-5654667;R-HSA-5654409;R-HSA-56547

R-HSA-5654392;R-HSA-5654631;R-HSA-1963563;R-HSA-8851899;R-HSA-5654633;R-HSA-5218830;R-HSA-1433

-380508;R-HSA-2574845;R-HSA-8942803;R-HSA-3000310;R-HSA-2172666;R-HSA-4088162;R-HSA-113503;R-HS

8864698;R-HSA-9622738;R-HSA-9615570;R-HSA-9007816;R-HSA-9615571;R-HSA-9623240;R-HSA-8937682;R-H

A-9623240;R-HSA-8937682;R-HSA-8953556;R-HSA-8938217;R-HSA-9623280;R-HSA-6804741;R-HSA-9623281;R

314554;R-HSA-8864698;R-HSA-9622738;R-HSA-9615570;R-HSA-9007816;R-HSA-9615571;R-HSA-9623240;R-HS

3000319;R-HSA-8941915;R-HSA-1363276;R-HSA-380508;R-HSA-2574845;R-HSA-8942803;R-HSA-3000310;R-HS

703;R-HSA-877308;R-HSA-997309;R-HSA-9606624;R-HSA-8987132;R-HSA-9008894;R-HSA-8987070;R-HSA-912

SA-1592238;R-HSA-1655843;R-HSA-1675810;R-HSA-1655844;R-HSA-1655847;R-HSA-1655846;R-HSA-1675866;

-HSA-2424476;R-HSA-2424477;R-HSA-8865605;R-HSA-9645137;R-HSA-8951498;R-HSA-2730848;R-HSA-89851[illegible]

A-210886;R-HSA-399951;R-HSA-416588;R-HSA-381262;R-HSA-9624668;R-HSA-560472;R-HSA-560473;R-HSA-3[illegible]
HSA-6809003;R-HSA-4717461;R-HSA-5419264;R-HSA-4085992;R-HSA-72663;R-HSA-5419268;R-HSA-72669;R-H

HSA-2399981;R-HSA-3149433;R-HSA-2399982;R-HSA-192841;R-HSA-112333;R-HSA-180687;R-HSA-1225951;R-
-6807118;R-HSA-5654655;R-HSA-3772434;R-HSA-8943780;R-HSA-5654651;R-HSA-3772436;R-HSA-6807126;R-
[illegible]456;R-HSA-202144;R-HSA-111524;R-HSA-1676082;R-HSA-1655848;R-HSA-2065966;R-HSA-1655850;R-HSA-16[illegible]

367;R-HSA-372529;R-HSA-9617326;R-HSA-380869;R-HSA-9610195;R-HSA-9619542;R-HSA-9610327;R-HSA-416

A-6809006;R-HSA-203977;R-HSA-6809003;R-HSA-2255342;R-HSA-421835;R-HSA-2255343;R-HSA-8849353;R-H

A-6809006;R-HSA-203977;R-HSA-6809003;R-HSA-2255342;R-HSA-421835;R-HSA-2255343;R-HSA-8849353;R-H

)9;R-HSA-1549564;R-HSA-5654714;R-HSA-5654587;R-HSA-5654584;R-HSA-5654692;R-HSA-5654571;R-HSA-56

428;R-HSA-5218829;R-HSA-8851908;R-HSA-177479;R-HSA-5218824;R-HSA-8851919;R-HSA-8865994;R-HSA-13

A-1363274;R-HSA-2574840;R-HSA-188386;R-HSA-4088024;R-HSA-1363314;R-HSA-69227;R-HSA-2484822;R-HS

SA-8953556;R-HSA-8938217;R-HSA-9623280;R-HSA-6804741;R-HSA-9623281;R-HSA-2187330;R-HSA-9018594

-HSA-2187330;R-HSA-9018594;R-HSA-350058;R-HSA-8865256;R-HSA-8952058;R-HSA-3222093;R-HSA-679529

A-8937682;R-HSA-8953556;R-HSA-8938217;R-HSA-9623280;R-HSA-6804741;R-HSA-9623281;R-HSA-2187330;

A-2172666;R-HSA-4088162;R-HSA-113503;R-HSA-1363274;R-HSA-2574840;R-HSA-188386;R-HSA-6804741;R-H

R-HSA-8874435;R-HSA-1676124;R-HSA-1676114;R-HSA-560473;R-HSA-163418;R-HSA-1675773;R-HSA-195690

57;R-HSA-879937;R-HSA-6800426;R-HSA-5672965;R-HSA-9645126;R-HSA-8854736;R-HSA-388829;R-HSA-3888

73716;R-HSA-8855508;R-HSA-9014610;R-HSA-8985812;R-HSA-210913;R-HSA-560491;R-HSA-560493;R-HSA-38
SA-265177;R-HSA-8873946;R-HSA-3000434;R-HSA-381799;R-HSA-6781953;R-HSA-2997706;R-HSA-5689649;R-

HSA-200908;R-HSA-8868567;R-HSA-9630795;R-HSA-5655351;R-HSA-5655347;R-HSA-5637808;R-HSA-2317387
HSA-5654631;R-HSA-5626982;R-HSA-380979;R-HSA-1963563;R-HSA-3772430;R-HSA-8964287;R-HSA-3965444
55852;R-HSA-1592238;R-HSA-1655843;R-HSA-1675810;R-HSA-70589;R-HSA-1655844;R-HSA-1655847;R-HSA-1

[illegible]320;R-HSA-9619803;R-HSA-8849878;R-HSA-6798258;R-HSA-451283;R-HSA-388824;R-HSA-8849882;R-HSA-41[illegible]

HSA-8876616;R-HSA-190681;R-HSA-190687;R-HSA-8868071;R-HSA-8866279;R-HSA-8866275;R-HSA-190693;R-

ISA-8876616;R-HSA-190681;R-HSA-190687;R-HSA-8868071;R-HSA-8866279;R-HSA-8866275;R-HSA-190693;R-

306972;R-HSA-1433415;R-HSA-5218839;R-HSA-5218838;R-HSA-1433423;R-HSA-9032402;R-HSA-5218836;R-HS

;A-4088141;R-HSA-380272;R-HSA-4088134;R-HSA-8942836;R-HSA-1226094;R-HSA-4088130;R-HSA-1226095;R

-;R-HSA-350058;R-HSA-8865256;R-HSA-8952058;R-HSA-3222093;R-HSA-6795290;R-HSA-6804762;R-HSA-2123[illegible]

)0;R-HSA-6804762;R-HSA-212352;R-HSA-2318752;R-HSA-212356;R-HSA-870538;R-HSA-8937728;R-HSA-89400[illegible]

R-HSA-9018594;R-HSA-350058;R-HSA-8865256;R-HSA-427383;R-HSA-8952058;R-HSA-3222093;R-HSA-679529[illegible]

HSA-141409;R-HSA-4088024;R-HSA-1363314;R-HSA-69227;R-HSA-141422;R-HSA-2484822;R-HSA-4088141;R-H

);R-HSA-1676145;R-HSA-6807052;R-HSA-1676149;R-HSA-1676020;R-HSA-194674;R-HSA-1676005;R-HSA-1676[illegible]

328;R-HSA-9645134;R-HSA-2197697;R-HSA-389348;R-HSA-8950634;R-HSA-1678843;R-HSA-1112690;R-HSA-167

31283;R-HSA-210921;R-HSA-210920;R-HSA-381290;R-HSA-373738;R-HSA-3928651;R-HSA-428535;R-HSA-39205
-HSA-8952289;R-HSA-8865774;R-HSA-4655404;R-HSA-1791092;R-HSA-156909;R-HSA-5690790;R-HSA-4551604

7;R-HSA-1247999;R-HSA-8875371;R-HSA-5228578;R-HSA-180591;R-HSA-194796;R-HSA-5334052;R-HSA-122595
4;R-HSA-6807134;R-HSA-870449;R-HSA-8964278;R-HSA-5654633;R-HSA-1912412;R-HSA-8943817;R-HSA-52188
1655846;R-HSA-947535;R-HSA-196929;R-HSA-1675866;R-HSA-203712;R-HSA-8874435;R-HSA-1676124;R-HSA-1

6985;R-HSA-380901;R-HSA-8849891;R-HSA-8855648;R-HSA-6797568;R-HSA-451311;R-HSA-442732;R-HSA-380

HSA-8866283;R-HSA-8868072;R-HSA-190829;R-HSA-1449574;R-HSA-6811414;R-HSA-8877813;R-HSA-6814096

HSA-8866283;R-HSA-8868072;R-HSA-190829;R-HSA-1449574;R-HSA-6811414;R-HSA-8877813;R-HSA-6814096

A-1250383;R-HSA-177501;R-HSA-8867047;R-HSA-187746;R-HSA-5686071;R-HSA-8867044;R-HSA-8867041;R-H

52;R-HSA-2318752;R-HSA-212356;R-HSA-870538;R-HSA-8937728;R-HSA-8940032;R-HSA-5632732;R-HSA-8937

32;R-HSA-5632732;R-HSA-8937989;R-HSA-3215295;R-HSA-9007640;R-HSA-9007646;R-HSA-8937744;R-HSA-89

);R-HSA-6804762;R-HSA-212352;R-HSA-2318752;R-HSA-212356;R-HSA-870538;R-HSA-8937728;R-HSA-73865;I

SA-380272;R-HSA-141431;R-HSA-4088134;R-HSA-8942836;R-HSA-1226094;R-HSA-4088130;R-HSA-1226095;R-

78842;R-HSA-389350;R-HSA-983147;R-HSA-168935;R-HSA-8950637;R-HSA-1169399;R-HSA-508513;R-HSA-430

54;R-HSA-3928654;R-HSA-560510;R-HSA-381309;R-HSA-5617452;R-HSA-3928642;R-HSA-560498;R-HSA-39286

57;R-HSA-5637806;R-HSA-200952;R-HSA-180606;R-HSA-5655339;R-HSA-5250606;R-HSA-4839734
330;R-HSA-5218829;R-HSA-6807077;R-HSA-5218824;R-HSA-195146;R-HSA-8943811;R-HSA-8865994;R-HSA-19
70600;R-HSA-265166;R-HSA-1676114;R-HSA-560473;R-HSA-372819;R-HSA-163418;R-HSA-400092;R-HSA-7177

905;R-HSA-1296127;R-HSA-9610879;R-HSA-6797974;R-HSA-8849906;R-HSA-374899;R-HSA-210426;R-HSA-41

ISA-1433471;R-HSA-5218851;R-HSA-5654587;R-HSA-5218850;R-HSA-5654584;R-HSA-5686318;R-HSA-143345[illegible]

[illegible]989;R-HSA-3215295;R-HSA-9007640;R-HSA-9007646;R-HSA-8937744;R-HSA-8936979;R-HSA-4411357;R-HSA-[illegible]

[illegible]36979;R-HSA-4411357;R-HSA-212380;R-HSA-9622579;R-HSA-8878117;R-HSA-6795460;R-HSA-4411367;R-HSA

[illegible]R-HSA-8940032;R-HSA-5632732;R-HSA-8937989;R-HSA-3215295;R-HSA-9007640;R-HSA-9007646;R-HSA-8937[illegible]

)190;R-HSA-874087;R-HSA-983140;R-HSA-914022;R-HSA-8983394;R-HSA-2201293;R-HSA-3134914;R-HSA-898[illegible]

)12396;R-HSA-9626816;R-HSA-5672965;R-HSA-8848596;R-HSA-8944349;R-HSA-2671742;R-HSA-5218839;R-HS[illegible]
'5;R-HSA-70241;R-HSA-1675773;R-HSA-6788611;R-HSA-71401;R-HSA-195690;R-HSA-1676145;R-HSA-6807052;

4;R-HSA-5654571;R-HSA-5654569;R-HSA-109699;R-HSA-1433488;R-HSA-1250463;R-HS

212380;R-HSA-9622579;R-HSA-8878117;R-HSA-6795460;R-HSA-4411367;R-HSA-902276

-9022764;R-HSA-6793666;R-HSA-8936481;R-HSA-8874797;R-HSA-8936995;R-HSA-8937

744;R-HSA-8936979;R-HSA-4411357;R-HSA-212380;R-HSA-9622579;R-HSA-8878117;R-

7236;R-HSA-1168636;R-HSA-389354;R-HSA-1112703;R-HSA-877308;R-HSA-2130641;R

A-9027281;R-HSA-1912388;R-HSA-8848606;R-HSA-5218838;R-HSA-1912391;R-HSA-562
